# Bayesian inference of toothed whale lifespans

**DOI:** 10.1101/2023.02.22.529527

**Authors:** Samuel Ellis, Darren P Croft, Mia Lybkær Kronborg Nielsen, Daniel W Franks, Michael N Weiss

## Abstract

Accurate measures of lifespan and age-specific mortality are important both for understanding life history evolution and informing conservation and population management strategies. The most accurate data to estimate lifespan are from longitudinal studies, but for many species – especially those such as toothed whales that are wide-ranging and inhabit difficult-to-access environments - these longitudinal data are not available. However, other forms of age-structured data – such as from mass-strandings - are available for many toothed species, and using these data to infer patterns of age-specific mortality and lifespan remains an important outstanding challenge. Here we develop and test a Bayesian mortality model to derive parameters of a mortality function from age-structured data while accounting for potential error introduced by mistakes in age estimation, sampling biases and population growth. We then searched the literature to assemble a database of 269 published age-structured toothed whale datasets. We applied our mortality model to derive lifespan estimates for 32 species of female and 33 species of male toothed whale. We also use our model to characterise sex differences in lifespan in toothed whales. Our mortality model allows us to curate the most complete and accurate collection of toothed whale lifespans to date.

## Introduction

Quantifying age-specific mortality risk – the hazard of an individual dying given their age – is fundamental to the study of life history evolution (Shefferson et al. 2017) as well as vital to informing conservation and population management strategies (Wade 2018). Data from longitudinal studies of individually identified individuals are the benchmark for calculating age-specific mortality risk.

Tracking individuals from birth to death can give an unbiased measure of age-specific mortality risk over the time period the observations were conducted (Caughley 1966; Hamlin et al. 2000; Nuñez et al. 2008; Gaillard et al. 2017). Although the number of species for who these data are available is increasing (e.g. (Salguero-Gómez et al. 2016; Lemaître et al. 2020)) they are still rare and hard to collect, especially for long-lived and difficult-to-study species in the wild.

Toothed whales (Odontocetes) are difficult to follow longitudinally through their life due both to their wide-ranging habits in marine habitats and, in at least some species, their extreme longevity.

Addressing this challenge is crucial, as characterising age-specific mortality of toothed whale species is important to inform conservation efforts. Toothed whale populations are subject to a variety of stressors including fisheries bycatch (Lewison et al. 2004), whaling (Clapham et al. 2007), pollution (Reijnders et al. 2018), anthropogenic noise (National Research Council 2005) and a changing climate (van Weelden et al. 2021; Kebke et al. 2022). Lifespan and age-specific mortality are important variables when modelling how populations will react to stressors. As well as conservation implications, toothed whales represent a scientifically important case study of the links between sociality and life history - particularly as they show a diversity of social systems and life histories (Connor et al. 1998; Whitehead and Mann 2000; Gowans et al. 2007; Weiss et al. 2021) including being the only mammalian taxa where menopause has evolved multiple times (Ellis et al. 2018a, b, 2024). It is therefore important to develop robust measures of lifespan that can be applied to toothed whales.

Although long-term longitudinal mortality data are rare in toothed whales, age-structured data are relatively common. The ages of individual toothed whales can be estimated by counting the growth rings in a cross-section of a tooth (Perrin and Myrick 1980; Read et al. 2018). This characteristic, along with a tradition of collecting detailed anatomical data from deceased whales (for example from mass-strandings, or fisheries by-catch), has resulted in the widespread publication of age distributions of samples of toothed whales. However, reporting the ages of calves and younger whales is less common and less systematic than the reporting of adults, for this reason throughout this study we focus on lifespan from age-at-maturity. Anatomical and age distribution data from deceased whales have been an important - and in many species the only - source of information on demography, life history and behaviour in toothed whales. However, because data are cross-sectional they have the potential to be biased as a measure of lifespan. In particular, three sources of error in cross-sectional mortality data have been identified: population growth or shrinking, sample biases and age-estimation error (Caughley 1966; Hamlin et al. 2000; Nuñez et al. 2008; Gaillard et al. 2017). Population change (growth or reduction) will affect the realised age distribution of a sample independently of patterns of mortality. For example, when a population is increasing younger individuals will be over-represented in a given sample. Conversely, if a population is shrinking then older individuals will be over-represented. Sample biases may be introduced into the reported age distribution of whales both at the point of mortality-for example if younger individuals are more likely to be caught as fisheries bycatch – and during the sample – for example, if larger, older, whales are preferentially selected for anatomical sampling. Similarly, and as with any data collection methodology, errors in age estimation may be introduced during the process of counting growth rings in the teeth. Any method using age-structured data to derive age-specific mortality requires a mechanism to propagate error through to the final mortality parameter estimates.

In this study, we develop, test and apply a method that uses age-structured data to derive estimates of age-specific mortality for toothed whale species. Specifically, we develop a Bayesian mortality model to estimate the parameters of a mortality function from age-structured data, while capturing variation in these values introduced by population growth, sampling biases and age-estimation errors. We then use large-scale simulation to test our mortality model. And lastly, we collate a database of age-structured toothed whale data from the literature and apply our methods to get estimates of age-specific mortality for as many toothed whale species as possible. We also use our models to compare male and female lifespans across our sample. Throughout this study, we focus on lifespan as a tractable and biologically-relevant property of cumulative age-specific mortality.

## Methods

### Mortality model

The number of whales of age *i* in a sample will depend on four key variables (Nuñez et al. 2008; Gaillard et al. 2017): (1) the probability of a whale surviving to age *i*; (2) the population growth rate (positive or negative); (3) biases in sampling and (4) error in age estimation. The aim of our model is to get a distribution of potential parameter values for a mortality model (1) given the unknown magnitude and influence of the other effects (2-4). Our model, therefore, does not aim to get a single ‘best estimate’ for the lifespan of each species, but rather to generate a distribution of potential lifespans given the observed data and the other processes that might be influencing the observed data. Subsequently, the model is not designed to estimate the effects of (2-4) and in fact when fitted the posterior of parameters relating to (2-4) do not usually differ from their priors. The presence of these parameters in the model, however, means that the derived distribution of mortality parameters (1) are those most consistent with the data given the unknown effects 2-4.

In this section, we first describe the case of the model applied to a single dataset. We then move on to describe how this single dataset model can be generalised to combine data from multiple datasets, sexes and populations of the same species into a single model.

We consider the count, *c*, of whales of a given age *AGE* in a sample to be drawn from a multinomial distribution, with probability θ over *N* individuals. Where θ_i_ is a function of: survival to age *AGE*_*i*_, *L*_*i*_; the effect of population growth, *R*_i_; and sampling biases at that age *S*_*i*_ (equation 1). Throughout this section *AGE* refers to age adjusted to age at maturity, so that age 0 is species- and sex-specific age at maturity.

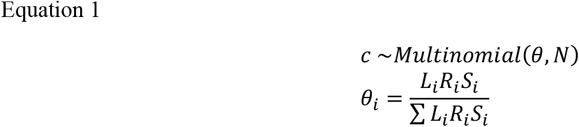

#### 1. Survival

We model the underlying probability of an individual surviving to age *i* as the survival curve (equation 2) derived from the Gompertz mortality model (Gompertz 1825).

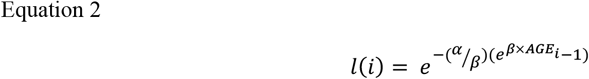

Where *α is* the baseline mortality and *β* is the increasing probability of mortality with age (ageing). As we are only modelling survival from age at maturity we do not use the Makeham/bathtub form of the Gompertz equation. We chose the Gompertz mortality model because it has few parameters, is widely used in studies of mammal senescence (Gaillard et al. 2017) and was recently found to be the most parsimonious mortality model in several killer whale populations (Nielsen et al. 2021a).

The probability that a sampled individual is of age *i, L*_*i*_ is drawn from the probability density function of the survival equation. Hence:

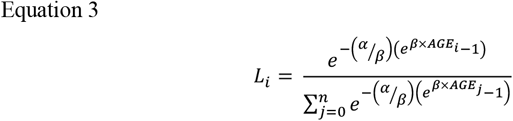

Where *n* is the maximum possible age, here we use *n* = 150 for all species.

As both Gompertz parameters are bounded between 0 and 1 we set up priors on *α* and *β* drawn from Beta distributions:

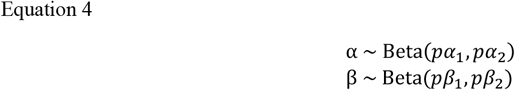

For this study, *pα*_*1*_ *= 6*.*79, pα*_*2*_ *= 166*.*31, pβ*_*1*_ *= 9*.*91, pβ*_*2*_ *= 37*.*93*. These priors are centred on the mean of published Gompertz *alpha* and *beta* values for Artiodactyla, with a variance double that of the published data (Gaillard et al. 2004; Toïgo et al. 2007; Foote 2008).

#### 2. Population growth rate

We model the increased probability of individuals of age *i* being observed due to population growth *R*_*i*_ as:

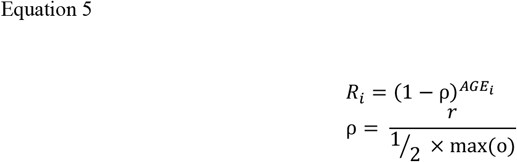

Where *r* is the population growth parameter, with *r* = 0 representing a stable population, *r* > 0 a growing population and *r* < 0 a shrinking population. For simplicity, we assume that the population growth rate is constant, or averages to a constant value, over the whole length of the study: *r*, therefore, represents annual population growth in any year of the study. *rho* is the growth parameter scaled to the maximum observed lifespan of the species *max(o)* so that the value of *r* can be directly compared between species and given a sensible and consistent prior. For example, r =0.1 means that the population will increase by 10% over half the maximum species lifespan. *Max(o)* is calculated from the raw data.

We use priors that allow us to leverage any data that may exist on the given population:

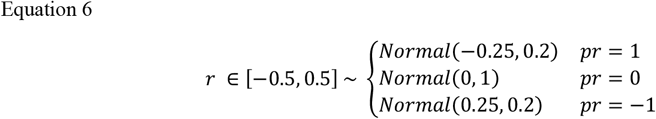

Where *pr = 1* indicates that there is strong evidence the population is growing, *pr = -1* that there is strong evidence that the population is shrinking and *pr = 0* that the population growth status of the population is unknown. Priors are moderately informative and limited between the population halving and doubling over half the maximum species lifetime.

#### 3. Sampling bias

We model sampling biases as an increased or decreased probability that whales within a given window *W* are under or over sampled. The sampling bias age window *W* is a pre-determined input into the model. Within the range a sampling bias effect *s* is applied, thus:

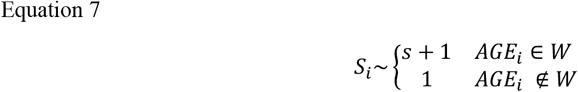

Where *s > 1* indicates that ages within range *W* are more likely to be sampled, *s* < 0 that they are less likely to be sampled, and *s* = 0 means no differences within and without the range.

The prior used for *s* in a given sample depends on whether the ages within the range are known to be over (*ps = 1*), under (*ps = -1*) sampled, or if the direction is unknown (*ps = 0*).

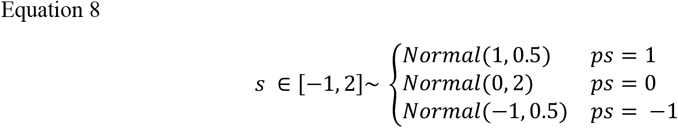

#### 4. Age-estimation error

We consider the true age of each whale *j* in the sample, *τ*_*j*_, to be drawn from a Gaussian distribution around the observed age *o*_*j*_ with standard deviation *ε*_*j*_. Where *B* is species-sex specific age at maturity at adding *B* therefore corrects between the real observed age of the sample and the model age (where age at maturity is age 0).

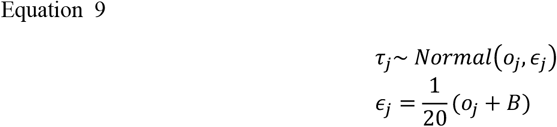

We use a chose a Gaussian distribution because it allows observed ages to be both over- and under-estimates of the true and allows assumptions around different ageing error with age pattern to be explored.

We assume that the error is additive such that *ε* is 5% of the observed age, and therefore that there is a greater error around older samples than younger samples. 5% was chosen to reflect a reasonable error rate in counting tooth rings, but changing this variable within reasonable bounds does not qualitatively affect the model outcomes. The outcome of this is that for a sample with an observed age of 10 the model assumes that there is a 90% probability that the true age is between 9 and 11, and for a whale of observed age 50 there is a 90% chance the sample is between 45 and 55.

We use estimated true ages, rounded to the nearest integer (notation: ⌊*x*⌉), of all the whales in the sample (N) to calculate the counts *c* of whales of a given age *i* in each sample used for the rest of the model.

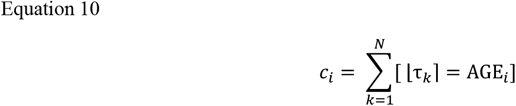

We combine these parts into a single model per species. Further, we design the model such that we run one model per species, combining multiple datasets and both sexes (where available). We estimate mortality parameters (*α & β*) for each sex with data from multiple datasets. Where multiple datasets are derived from the same population (most commonly, one for each sex) we estimate a single population growth parameter for these datasets. We calculate a separate sampling bias metric for each dataset. The complete model is shown below (notation key in table 1):

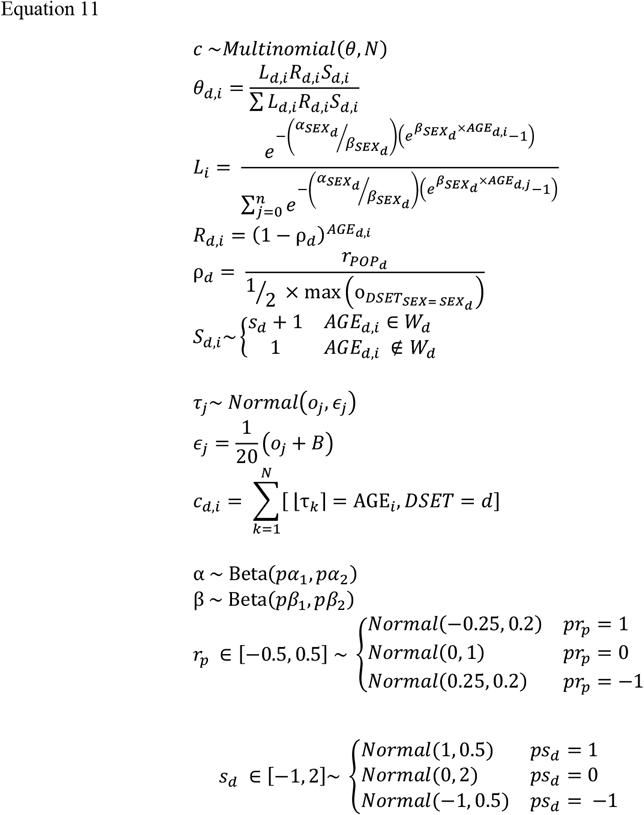

**Table 1.**
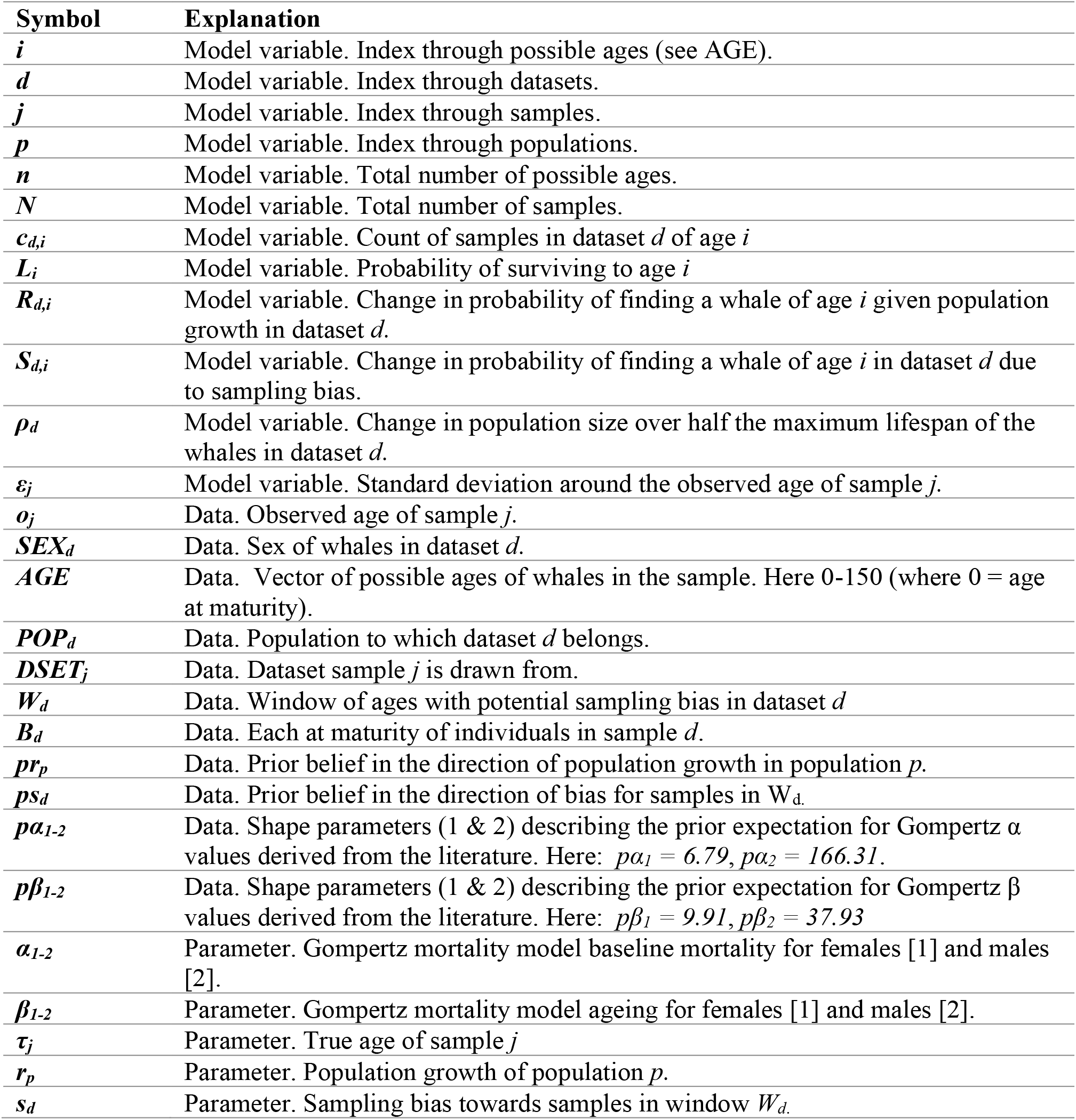
Explanation of symbols used in equation 1.

The model was implemented in STAN and R using functionality from the rstan, cmdstanr and rethinking packages (McElreath 2020; Stan Development Team 2020, 2021; R Development Core Team 2021). The model has been built into an R package - marinesurvival (github.com/samellisq/marinesurvival) – which includes all R and STAN code necessary to implement the models.

### Testing the model

We performed two analyses to test the ability of the model to estimate toothed whale age-specific mortality. First, we test the ability of the model to recover the known input parameters. To do this we generate 100 simulated datasets of each combination of seven sample sizes (n = 10, 20, 35, 50, 75, 100, 200) and seven population growth rates (r =-0.25, -0.1, -0.05, 0, 0.05, 0.1, 0.25), giving a total of 4900 simulated datasets. Rather than selecting mortality parameters α and β at random, we chose the parameters to represent a plausible range of Odontocete adult lifespans. For each five-year age window between 0 and 50, we generated 500 paired α and β parameters resulting in lifespans within that window by selecting at random from a systematic analysis of the αxβ parameter space. For each simulation, we selected α and β parameters from these 7500 pairs. The result of this is an equal probability of the population in any simulation having an (adult) lifespan in any 5-year age window between 0 and 50. All other model parameters are chosen at random for each simulation. We apply the mortality model to each simulation and generate a posterior distribution of ordinary maximum lifespans (see calculating lifespans section) predicted by the model. Models that did not converge were not included in any further analysis. As our metric of model accuracy, we use the minimum credible interval width from the posterior distribution of maximum lifespans required to capture the true maximum lifespan. An accuracy of 0 would indicate the median of the posterior matches the true maximum lifespan, while an accuracy of 0.52 would mean that the true lifespan is captured by the 52% credible interval of the posterior. We focus on the accuracy of the model in relation to sample size, population growth and whale lifespan as those most likely to systematically bias our measures of lifespan with real data. This approach is functionally similar to a simulation based calibration approach (Modrák et al. 2023).

Secondly, we compare the lifespan estimates from our mortality model to well-characterised lifespan estimates derived from longitudinal data in a well-studied toothed whale population. Southern-resident killer whales inhabiting the north-east Pacific ocean are regularly observed in inland waters in the Salish Sea (Bigg et al. 1990). The population has been studied intensively for 50 years, and all births (of whales surviving their first year) and deaths in the population have been collated by the Center for Whale Research since 1976 (details, including age estimate for whales born before 1976, in: (Nielsen et al. 2021a)). A previous study has used these detailed demographic records in conjunction with established Bayesian mark-recapture mortality modelling to estimate age-specific mortality in this population (Nielsen et al. 2021a). Mark-recapture mortality models derive parameters for a mortality function given known dates of birth and death, incorporating sightings data to improve parameter estimation and estimate unknown years of birth (further details in: (Colchero et al. 2012; Nielsen et al. 2021a)). We rescale the published age-specific survival estimates to apply only to adults by dividing all post-maturity survival estimates by the estimated survival to maturity. From these adult survival curves, we can derive a credible interval of ordinary maximum lifespans (hereafter: longitudinal lifespan). We tested our model by comparing the longitudinal lifespans to estimates derived from our model applied to data from the same population. We use a single year of southern-resident killer whale demographic data – the equivalent of a situation where the whole population of whales in a single year were sampled, sexed and aged – as the datasets input into the model. We use the year 1992 when the southern-residents population size peaked (1976-2015), the samples consist of 45 adult females and 16 adult males present in the population in that year.

### Literature search

We performed a systematic and opportunistic search of the literature in May-September 2021 to identify and extract all published age-structured toothed whale data. Age-structured data are data where the ages of individuals of known age and sex can be collected or inferred from published results. For each of the 75 species of toothed whale (IWC 2021) we searched the Web of Science database (webofscience.com) with the species name plus separately each of “life history”, “lifespan” and “age structure”. We repeated this sequence with both the currently accepted species name, any previously used species names, and all widely used common names for the species. In addition, limiting searches to publications in English can bias datasets in systematic reviews and meta-analyses (Konno et al. 2020). For each species, we, therefore, repeated the database query replacing the English search terms with the same terms translated into major languages commonly spoken around the species distribution. However, there is still likely to be some bias, especially against languages not written in the Latin alphabet. We also performed an opportunistic search by consulting the relevant chapters in the Handbook of Marine Mammals v4-7 (Ridgway and Harrison 1989, 1994, 1999) and the Encyclopaedia of Marine Mammals 3^rd^ Edition (Würsig et al. 2018) and identifying potential data sources referenced in these chapters, as well as searching backwards and forwards through the citation network of identified papers.

We extracted age-structured datasets from these publications to create a database of odontocete age-structured data. Where possible each dataset in the database represents data from a single population collected at a single time, however in some systems and publications this is not possible and a single dataset represents data from multiple populations or is collected over a longer timescale. Each dataset represents only whales of a single known sex, and whales of unknown sex are not included in the datasets. In each publication, age data are extracted from the table or figure giving the largest and most complete sample. Where a single publication contains information from multiple populations we extracted data from each publication as a separate dataset, if possible. Where the same dataset is presented in multiple publications we use the original publication unless later publications present a larger sample. Data were extracted from figures using WebPlotDigitizer (Rohatgi 2020). Data were extracted and included in the database if they contained more than three adult samples of a given sex. For each species-sex, we define the age at maturity based on expert consensus (Ridgway and Harrison 1989, 1994, 1999; Würsig et al. 2018). The complete database contains 269 datasets from 118 publications with data from 44 species (table 2; figure 1; supplementary 1). The database has been built into an R package: marinelifehistdata (github.com/samellisq/marinelifehistdata).

**Table 2.**
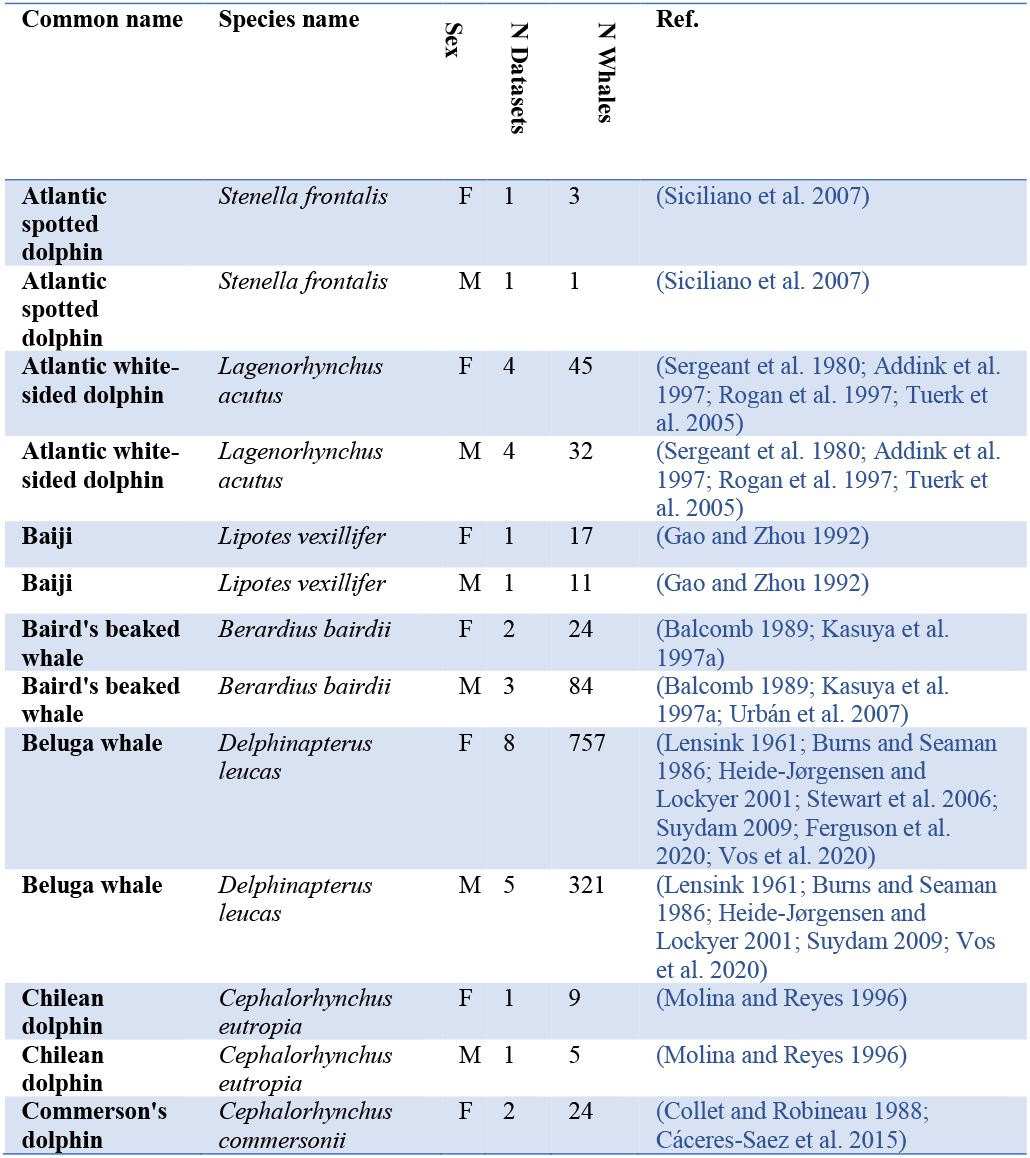
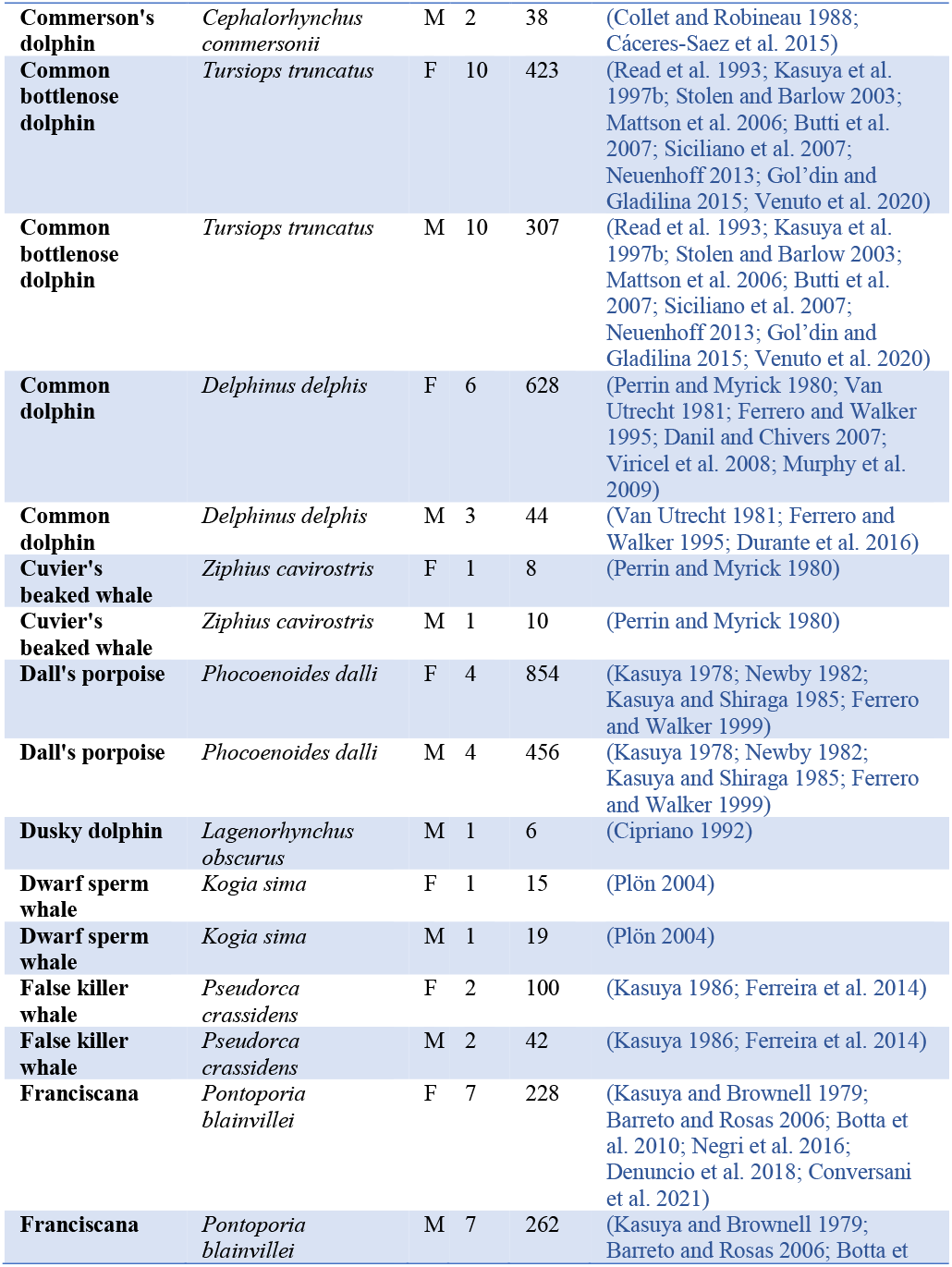
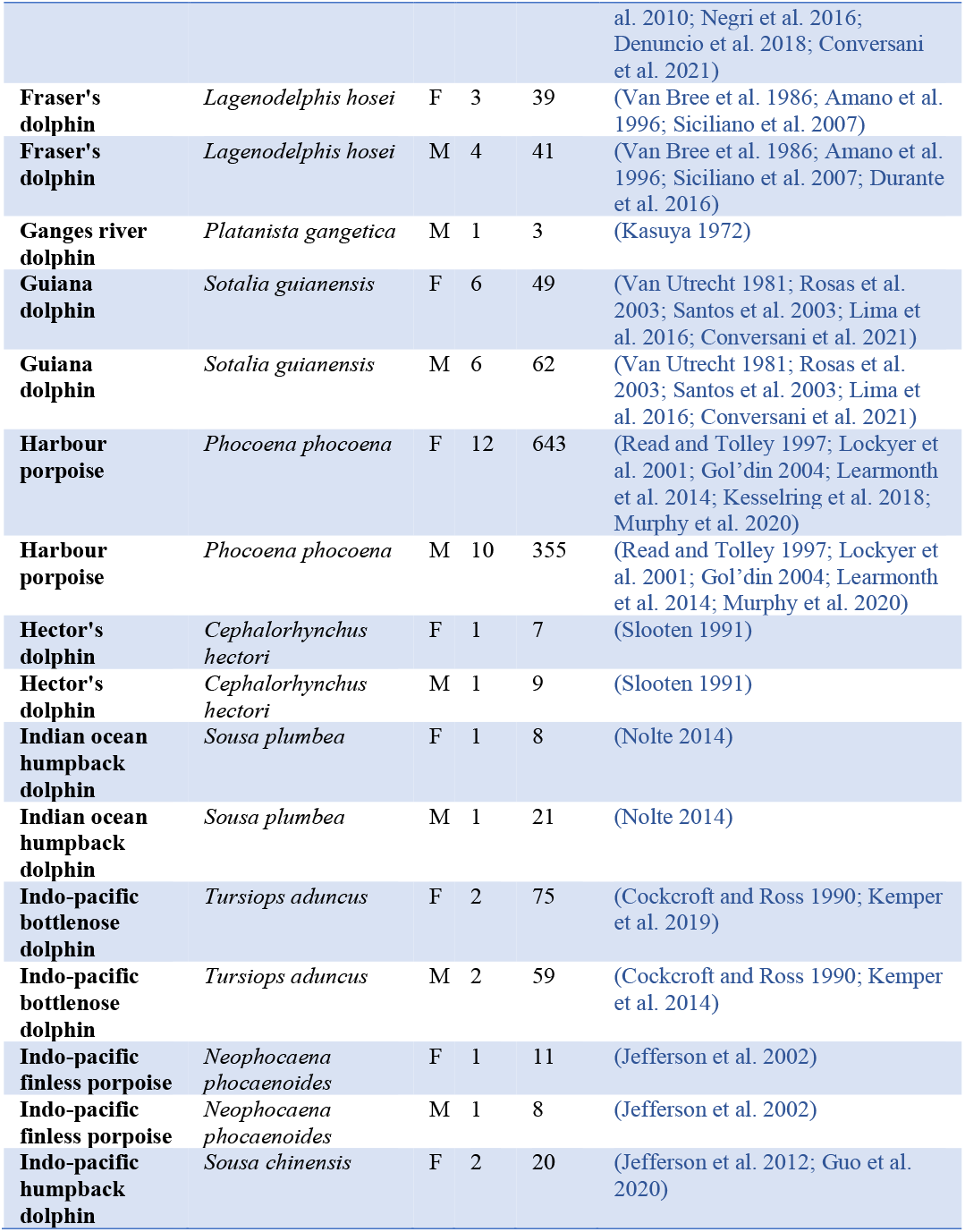
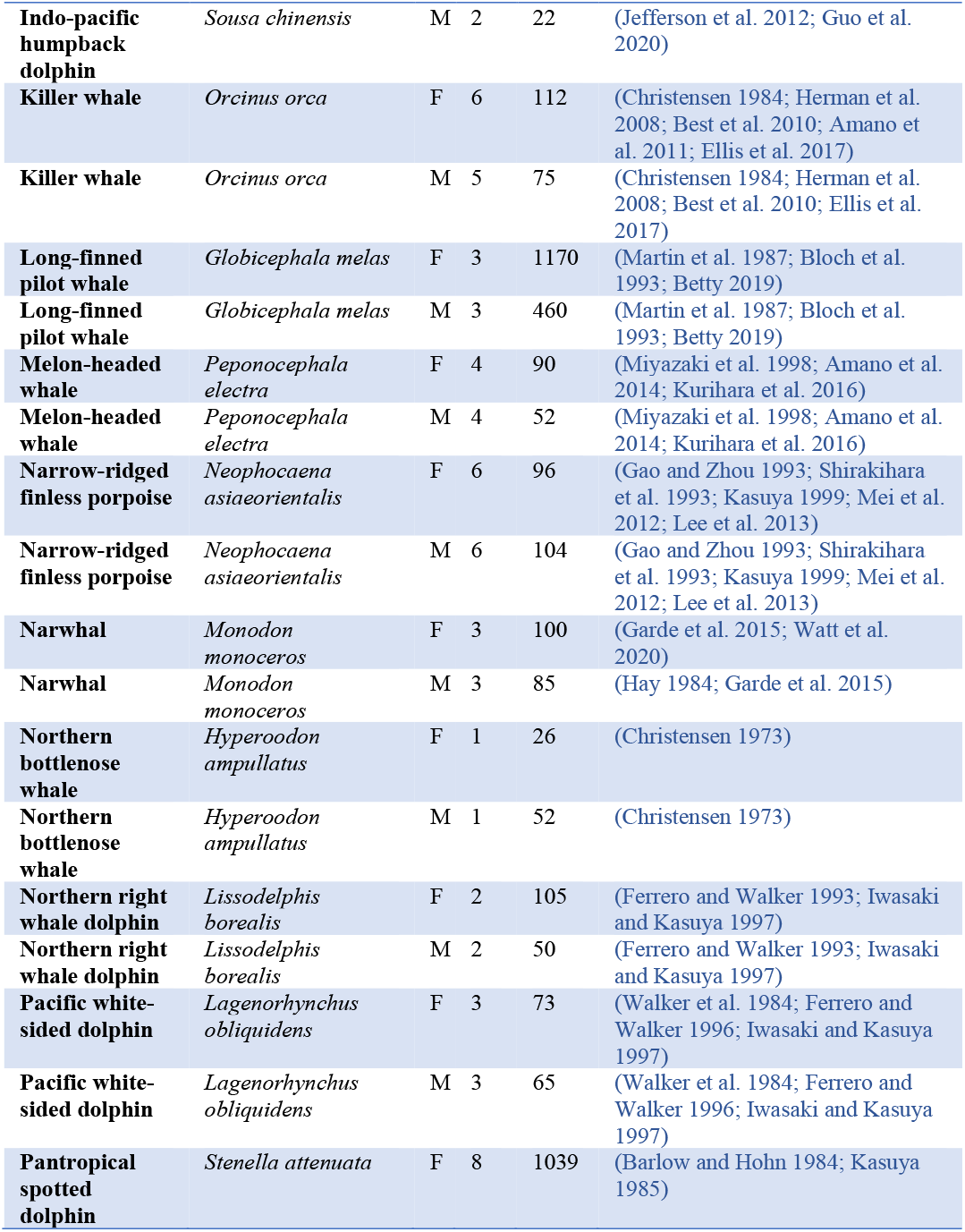
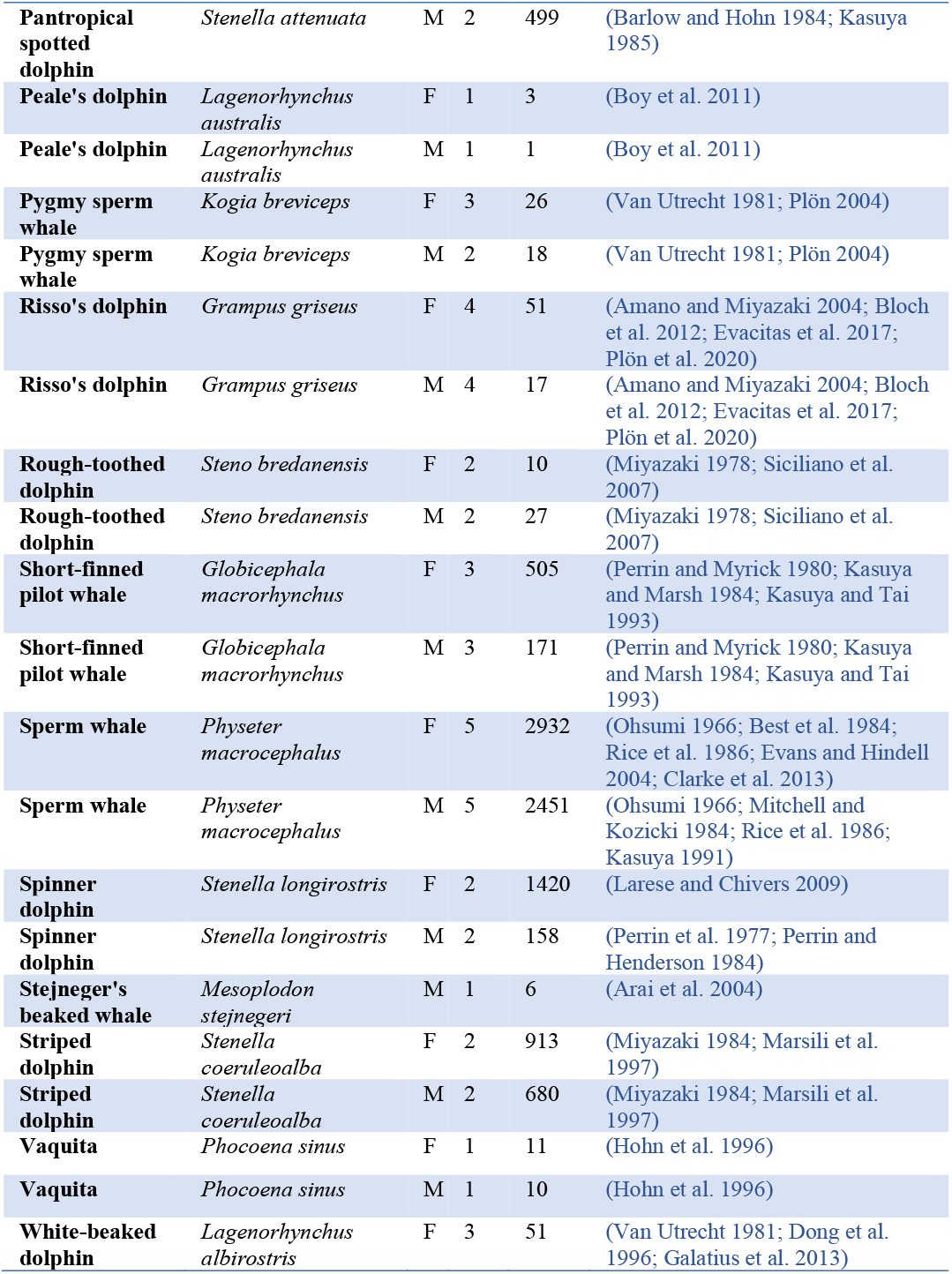
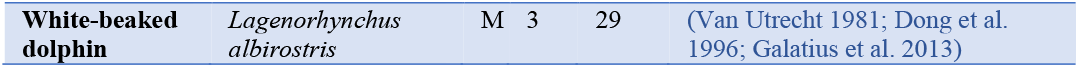
Details of the age-structured samples identified in this study. Note while all datasets and species where data were identified are included here we only analyse species-sex in further analysis if it exceeded sample size and sampling intensity conditions (see methods: calculating lifespan).

| Common name | Species name | Sex | N Datasets | N Whales | Ref. |
| --- | --- | --- | --- | --- | --- |
| Atlantic spotted dolphin | *Stenella frontalis* | F | 1 | 3 | (Siciliano et al. 2007) |
| Atlantic spotted dolphin | *Stenella frontalis* | M | 1 | 1 | (Siciliano et al. 2007) |
| Atlantic white-sided dolphin | *Lagenorhynchus acutus* | F | 4 | 45 | (Sergeant et al. 1980; Addink et al. 1997; Rogan et al. 1997; Tuerk et al. 2005) |
| Atlantic white-sided dolphin | *Lagenorhynchus acutus* | M | 4 | 32 | (Sergeant et al. 1980; Addink et al. 1997; Rogan et al. 1997; Tuerk et al. 2005) |
| Baiji | *Lipotes vexillifer* | F | 1 | 17 | (Gao and Zhou 1992) |
| Baiji | *Lipotes vexillifer* | M | 1 | 11 | (Gao and Zhou 1992) |
| Baird's beaked whale | *Berardius bairdii* | F | 2 | 24 | (Balcomb 1989; Kasuya et al. 1997a) |
| Baird's beaked whale | *Berardius bairdii* | M | 3 | 84 | (Balcomb 1989; Kasuya et al. 1997a; Urbán et al. 2007) |
| Beluga whale | *Delphinapterus leucas* | F | 8 | 757 | (Lensink 1961; Burns and Seaman 1986; Heide-Jørgensen and Lockyer 2001; Stewart et al. 2006; Suydam 2009; Ferguson et al. 2020; Vos et al. 2020) |
| Beluga whale | *Delphinapterus leucas* | M | 5 | 321 | (Lensink 1961; Burns and Seaman 1986; Heide-Jørgensen and Lockyer 2001; Suydam 2009; Vos et al. 2020) |
| Chilean dolphin | *Cephalorhynchus eutropia* | F | 1 | 9 | (Molina and Reyes 1996) |
| Chilean dolphin | *Cephalorhynchus eutropia* | M | 1 | 5 | (Molina and Reyes 1996) |
| Commerson's dolphin | *Cephalorhynchus commersonii* | F | 2 | 24 | (Collet and Robineau 1988; Cáceres-Saez et al. 2015) |
| Commerson's dolphin | *Cephalorhynchus commersonii* | M | 2 | 38 | (Collet and Robineau 1988; Cáceres-Saez et al. 2015) |
| Common bottlenose dolphin | *Tursiops truncatus* | F | 10 | 423 | (Read et al. 1993; Kasuya et al. 1997b; Stolen and Barlow 2003; Mattson et al. 2006; Butti et al. 2007; Siciliano et al. 2007; Neuenhoff 2013; Gol'din and Gladilina 2015; Venuto et al. 2020) |
| Common bottlenose dolphin | *Tursiops truncatus* | M | 10 | 307 | (Read et al. 1993; Kasuya et al. 1997b; Stolen and Barlow 2003; Mattson et al. 2006; Butti et al. 2007; Siciliano et al. 2007; Neuenhoff 2013; Gol'din and Gladilina 2015; Venuto et al. 2020) |
| Common dolphin | *Delphinus delphis* | F | 6 | 628 | (Perrin and Myrick 1980; Van Utrecht 1981; Ferrero and Walker 1995; Danil and Chivers 2007; Viricel et al. 2008; Murphy et al. 2009) |
| Common dolphin | *Delphinus delphis* | M | 3 | 44 | (Van Utrecht 1981; Ferrero and Walker 1995; Durante et al. 2016) |
| Cuvier's beaked whale | *Ziphius cavirostris* | F | 1 | 8 | (Perrin and Myrick 1980) |
| Cuvier's beaked whale | *Ziphius cavirostris* | M | 1 | 10 | (Perrin and Myrick 1980) |
| Dall's porpoise | *Phocoenoides dalli* | F | 4 | 854 | (Kasuya 1978; Newby 1982; Kasuya and Shiraga 1985; Ferrero and Walker 1999) |
| Dall's porpoise | *Phocoenoides dalli* | M | 4 | 456 | (Kasuya 1978; Newby 1982; Kasuya and Shiraga 1985; Ferrero and Walker 1999) |
| Dusky dolphin | *Lagenorhynchus obscurus* | M | 1 | 6 | (Cipriano 1992) |
| Dwarf sperm whale | *Kogia sima* | F | 1 | 15 | (Plön 2004) |
| Dwarf sperm whale | *Kogia sima* | M | 1 | 19 | (Plön 2004) |
| False killer whale | *Pseudorca crassidens* | F | 2 | 100 | (Kasuya 1986; Ferreira et al. 2014) |
| False killer whale | *Pseudorca crassidens* | M | 2 | 42 | (Kasuya 1986; Ferreira et al. 2014) |
| Franciscana | *Pontoporia blainvillei* | F | 7 | 228 | (Kasuya and Brownell 1979; Barreto and Rosas 2006; Botta et al. 2010; Negri et al. 2016; Denuncio et al. 2018; Conversani et al. 2021) |
| Franciscana | *Pontoporia blainvillei* | M | 7 | 262 | (Kasuya and Brownell 1979; Barreto and Rosas 2006; Botta et al. 2010; Negri et al. 2016; Denuncio et al. 2018; Conversani et al. 2021) |
| **Fraser's dolphin** | *Lagenodelphis hosei* | F | 3 | 39 | (Van Bree et al. 1986; Amano et al. 1996; Siciliano et al. 2007) |
| **Fraser's dolphin** | *Lagenodelphis hosei* | M | 4 | 41 | (Van Bree et al. 1986; Amano et al. 1996; Siciliano et al. 2007; Durante et al. 2016) |
| **Ganges river dolphin** | *Platanista gangetica* | M | 1 | 3 | (Kasuya 1972) |
| **Guiana dolphin** | *Sotalia guianensis* | F | 6 | 49 | (Van Utrecht 1981; Rosas et al. 2003; Santos et al. 2003; Lima et al. 2016; Conversani et al. 2021) |
| **Guiana dolphin** | *Sotalia guianensis* | M | 6 | 62 | (Van Utrecht 1981; Rosas et al. 2003; Santos et al. 2003; Lima et al. 2016; Conversani et al. 2021) |
| **Harbour porpoise** | *Phocoena phocoena* | F | 12 | 643 | (Read and Tolley 1997; Lockyer et al. 2001; Gol'din 2004; Learmonth et al. 2014; Kesselring et al. 2018; Murphy et al. 2020) |
| **Harbour porpoise** | *Phocoena phocoena* | M | 10 | 355 | (Read and Tolley 1997; Lockyer et al. 2001; Gol'din 2004; Learmonth et al. 2014; Murphy et al. 2020) |
| **Hector's dolphin** | *Cephalorhynchus hectori* | F | 1 | 7 | (Slooten 1991) |
| **Hector's dolphin** | *Cephalorhynchus hectori* | M | 1 | 9 | (Slooten 1991) |
| **Indian ocean humpback dolphin** | *Sousa plumbea* | F | 1 | 8 | (Nolte 2014) |
| **Indian ocean humpback dolphin** | *Sousa plumbea* | M | 1 | 21 | (Nolte 2014) |
| **Indo-pacific bottlenose dolphin** | *Tursiops aduncus* | F | 2 | 75 | (Cockcroft and Ross 1990; Kemper et al. 2019) |
| **Indo-pacific bottlenose dolphin** | *Tursiops aduncus* | M | 2 | 59 | (Cockcroft and Ross 1990; Kemper et al. 2014) |
| **Indo-pacific finless porpoise** | *Neophocaena phocaenoides* | F | 1 | 11 | (Jefferson et al. 2002) |
| **Indo-pacific finless porpoise** | *Neophocaena phocaenoides* | M | 1 | 8 | (Jefferson et al. 2002) |
| **Indo-pacific humpback dolphin** | *Sousa chinensis* | F | 2 | 20 | (Jefferson et al. 2012; Guo et al. 2020) |
| **Indo-pacific humpback dolphin** | *Sousa chinensis* | M | 2 | 22 | (Jefferson et al. 2012; Guo et al. 2020) |
| **Killer whale** | *Orcinus orca* | F | 6 | 112 | (Christensen 1984; Herman et al. 2008; Best et al. 2010; Amano et al. 2011; Ellis et al. 2017) |
| **Killer whale** | *Orcinus orca* | M | 5 | 75 | (Christensen 1984; Herman et al. 2008; Best et al. 2010; Ellis et al. 2017) |
| **Long-finned pilot whale** | *Globicephala melas* | F | 3 | 1170 | (Martin et al. 1987; Bloch et al. 1993; Betty 2019) |
| **Long-finned pilot whale** | *Globicephala melas* | M | 3 | 460 | (Martin et al. 1987; Bloch et al. 1993; Betty 2019) |
| **Melon-headed whale** | *Peponocephala electra* | F | 4 | 90 | (Miyazaki et al. 1998; Amano et al. 2014; Kurihara et al. 2016) |
| **Melon-headed whale** | *Peponocephala electra* | M | 4 | 52 | (Miyazaki et al. 1998; Amano et al. 2014; Kurihara et al. 2016) |
| **Narrow-ridged finless porpoise** | *Neophocaena asiaeorientalis* | F | 6 | 96 | (Gao and Zhou 1993; Shirakihara et al. 1993; Kasuya 1999; Mei et al. 2012; Lee et al. 2013) |
| **Narrow-ridged finless porpoise** | *Neophocaena asiaeorientalis* | M | 6 | 104 | (Gao and Zhou 1993; Shirakihara et al. 1993; Kasuya 1999; Mei et al. 2012; Lee et al. 2013) |
| **Narwhal** | *Monodon monoceros* | F | 3 | 100 | (Garde et al. 2015; Watt et al. 2020) |
| **Narwhal** | *Monodon monoceros* | M | 3 | 85 | (Hay 1984; Garde et al. 2015) |
| **Northern bottlenose whale** | *Hyperoodon ampullatus* | F | 1 | 26 | (Christensen 1973) |
| **Northern bottlenose whale** | *Hyperoodon ampullatus* | M | 1 | 52 | (Christensen 1973) |
| **Northern right whale dolphin** | *Lissodelphis borealis* | F | 2 | 105 | (Ferrero and Walker 1993; Iwasaki and Kasuya 1997) |
| **Northern right whale dolphin** | *Lissodelphis borealis* | M | 2 | 50 | (Ferrero and Walker 1993; Iwasaki and Kasuya 1997) |
| **Pacific white-sided dolphin** | *Lagenorhynchus obliquidens* | F | 3 | 73 | (Walker et al. 1984; Ferrero and Walker 1996; Iwasaki and Kasuya 1997) |
| **Pacific white-sided dolphin** | *Lagenorhynchus obliquidens* | M | 3 | 65 | (Walker et al. 1984; Ferrero and Walker 1996; Iwasaki and Kasuya 1997) |
| **Pantropical spotted dolphin** | *Stenella attenuata* | F | 8 | 1039 | (Barlow and Hohn 1984; Kasuya 1985) |
| **Pantropical spotted dolphin** | *Stenella attenuata* | M | 2 | 499 | (Barlow and Hohn 1984; Kasuya 1985) |
| **Peale's dolphin** | *Lagenorhynchus australis* | F | 1 | 3 | (Boy et al. 2011) |
| **Peale's dolphin** | *Lagenorhynchus australis* | M | 1 | 1 | (Boy et al. 2011) |
| **Pygmy sperm whale** | *Kogia breviceps* | F | 3 | 26 | (Van Utrecht 1981; Plön 2004) |
| **Pygmy sperm whale** | *Kogia breviceps* | M | 2 | 18 | (Van Utrecht 1981; Plön 2004) |
| **Risso's dolphin** | *Grampus griseus* | F | 4 | 51 | (Amano and Miyazaki 2004; Bloch et al. 2012; Evacitas et al. 2017; Plön et al. 2020) |
| **Risso's dolphin** | *Grampus griseus* | M | 4 | 17 | (Amano and Miyazaki 2004; Bloch et al. 2012; Evacitas et al. 2017; Plön et al. 2020) |
| **Rough-toothed dolphin** | *Steno bredanensis* | F | 2 | 10 | (Miyazaki 1978; Siciliano et al. 2007) |
| **Rough-toothed dolphin** | *Steno bredanensis* | M | 2 | 27 | (Miyazaki 1978; Siciliano et al. 2007) |
| **Short-finned pilot whale** | *Globicephala macrorhynchus* | F | 3 | 505 | (Perrin and Myrick 1980; Kasuya and Marsh 1984; Kasuya and Tai 1993) |
| **Short-finned pilot whale** | *Globicephala macrorhynchus* | M | 3 | 171 | (Perrin and Myrick 1980; Kasuya and Marsh 1984; Kasuya and Tai 1993) |
| **Sperm whale** | *Physeter macrocephalus* | F | 5 | 2932 | (Ohsumi 1966; Best et al. 1984; Rice et al. 1986; Evans and Hindell 2004; Clarke et al. 2013) |
| **Sperm whale** | *Physeter macrocephalus* | M | 5 | 2451 | (Ohsumi 1966; Mitchell and Kozicki 1984; Rice et al. 1986; Kasuya 1991) |
| **Spinner dolphin** | *Stenella longirostris* | F | 2 | 1420 | (Larese and Chivers 2009) |
| **Spinner dolphin** | *Stenella longirostris* | M | 2 | 158 | (Perrin et al. 1977; Perrin and Henderson 1984) |
| **Stejneger's beaked whale** | *Mesoplodon stejnegeri* | M | 1 | 6 | (Arai et al. 2004) |
| **Striped dolphin** | *Stenella coeruleoalba* | F | 2 | 913 | (Miyazaki 1984; Marsili et al. 1997) |
| **Striped dolphin** | *Stenella coeruleoalba* | M | 2 | 680 | (Miyazaki 1984; Marsili et al. 1997) |
| **Vaquita** | *Phocoena sinus* | F | 1 | 11 | (Hohn et al. 1996) |
| **Vaquita** | *Phocoena sinus* | M | 1 | 10 | (Hohn et al. 1996) |
| **White-beaked dolphin** | *Lagenorhynchus albirostris* | F | 3 | 51 | (Van Utrecht 1981; Dong et al. 1996; Galatius et al. 2013) |
| **White-beaked dolphin** | *Lagenorhynchus albirostris* | M | 3 | 29 | (Van Utrecht 1981; Dong et al. 1996; Galatius et al. 2013) |

**Figure 1.**
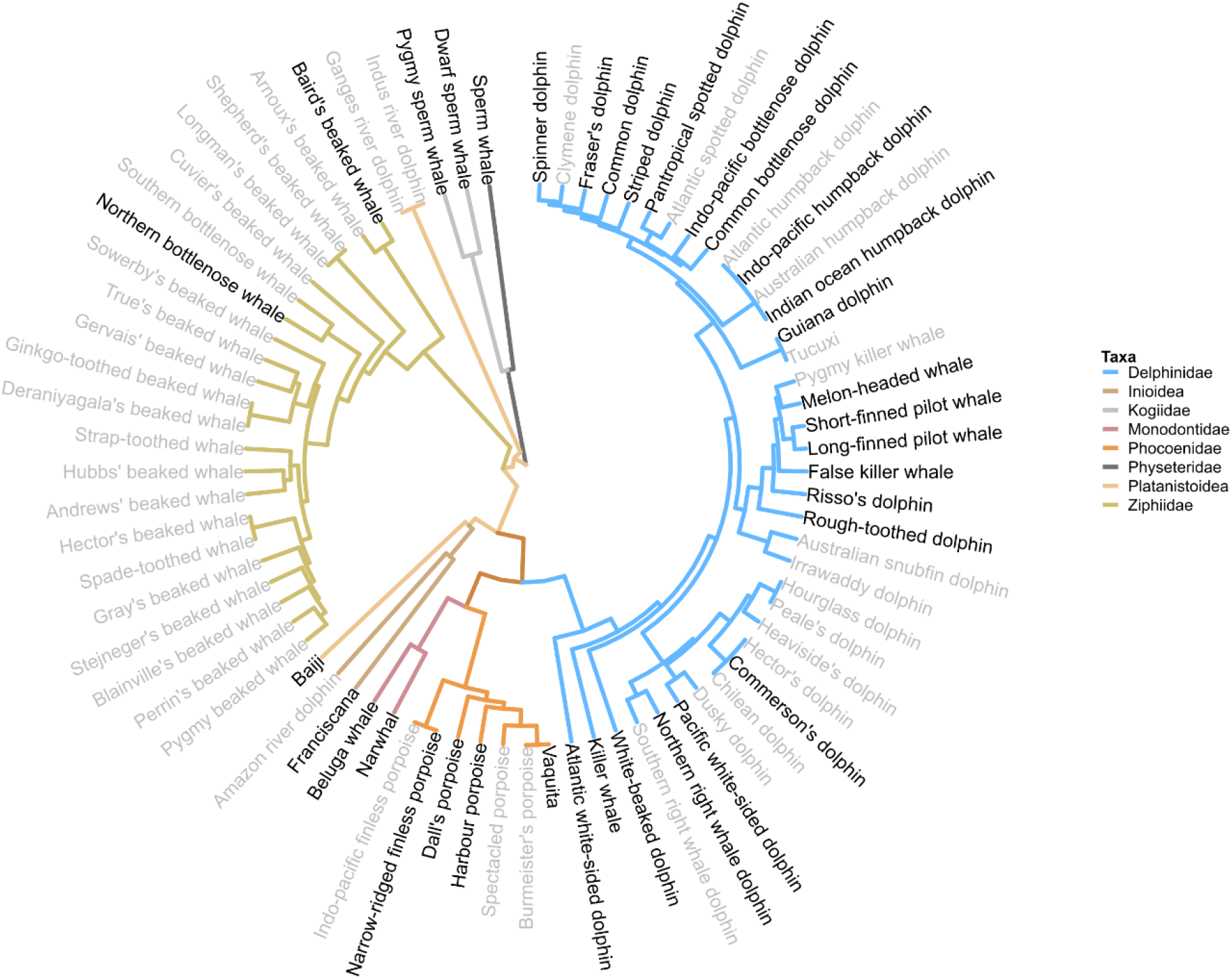
Toothed whale phylogeny (adapted from (McGowen et al. 2020)). Data are available to apply our model and estimate age-specific mortality for at least one sex in 35 species (black text; grey text = no data). Branch colours correspond to taxonomic family.

We also parametrise the mortality model with dataset-specific information on population growth, sampling biases and age estimation error. We base our population change prior on necessarily coarse assumptions on the source of the data. Where datasets are derived from bycatch (usually resulting from largescale bycatch in commercial fisheries, such as the pacific gillnet fishery; N = 72) or whaling/drive fisheries (N = 50) we assume that the population is decreasing (*pr* = -1). Where datasets are from Harvest (‘Aboriginal Subsistence Whaling’ as defined by the International Whaling Commission; N = 20) we assume that there is no population change based on the monitoring and management the populations are subject to. For all other data sources, we assume that the population could be increasing or decreasing (*pr* = 0). Datasets are assumed to have no systematic sampling biases unless noted by the authors of the original study (or subsequent publications analysing the same dataset). Where biased sampling is present we define the bias window *W* using the information provided by the authors. Unless the ages in a dataset were derived from long-term observations (n = 8) or aspartic acid racemisation (n = 6) they were assumed to have additive age estimation error as defined by the model (above).

### Calculating lifespan

We filtered the database before analysis. For each dataset, we calculate the number of samples, and the sampling rate (total samples/maximum observed age observed across all species-sex datasets). We only analyse species-sexes where at least one dataset has >10 samples and a sampling rate of >0.5.

We applied the mortality model to each species as described above (supplementary 1). We obtain estimates of lifespan for each species-sex using Gompertz α and β parameters. Specifically, we calculate the age at which X proportion of species-sex life years have been lived via:

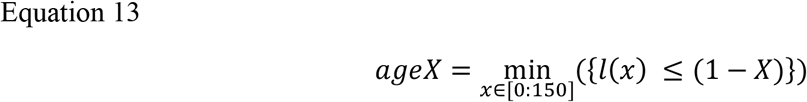

Here we use X=0.9 as ‘ordinary maximum lifespan’ (or age Z) as our measure of species-sex lifespan (Ronget and Gaillard 2020).

We calculate ordinary maximum lifespan from each of the 40 000 posterior draws of the fitted Bayesian model, resulting in a distribution of age Z’s for each species-sex.

We also performed a series of additional analyses to explore the robustness and implications of our models. Specifically we: (1) explore the robustness of our model to systematic mis-ageing of older individuals (Barratclough et al. 2023; supplementary 2); (2) establish if changing ageing methodologies through time have led to different observed age-distributions (Read et al. 2018; supplementary 3); (3) investigate the sensitivity of our model to our population change assumptions (supplementary 4); (4) reformulate and test our model as an age-at-death model (supplementary 5); (5) evaluate evidence for phylogenetic signal in toothed whale lifespan evolution (supplementary 6); and (6) explore the impact of alternative age estimation error structures on inference of lifespan (supplementary 7).

## Results

### Testing the model

Our first set of analyses explore the consistency and accuracy of the model across a realistic range of population growth rates and whale adult lifespans (figure 2). There is a trend for increased accuracy with larger sample sizes: with larger sample sizes the posterior median is closer to the true lifespan (figure 2). However, the increase in accuracy is relatively modest and quite variable, for example in stable populations (r = 0) mean minimal credible interval required to capture the true lifespan is 60± 35% (±std. dev.) for a sample size of 10 and 27± 37% for a sample size of 200 (figure 2; see also (Vaupel 2003)). There are no clear trends for increased accuracy for either shorter lifespans or more stable populations (figure 2). The model accuracy is maintained across these scenarios because credible intervals are wider under small sample sizes and for long-lived whales: in short, the model becomes less certain under more unsure information (Supplementary 8). Under all scenarios, the largest mean minimum credible interval needed to capture the true value is 77% (n=10, true age 40-49) suggesting that as long as measures of error are carried through subsequent analyses the model is likely to capture the true lifespan.

**Figure 2.**
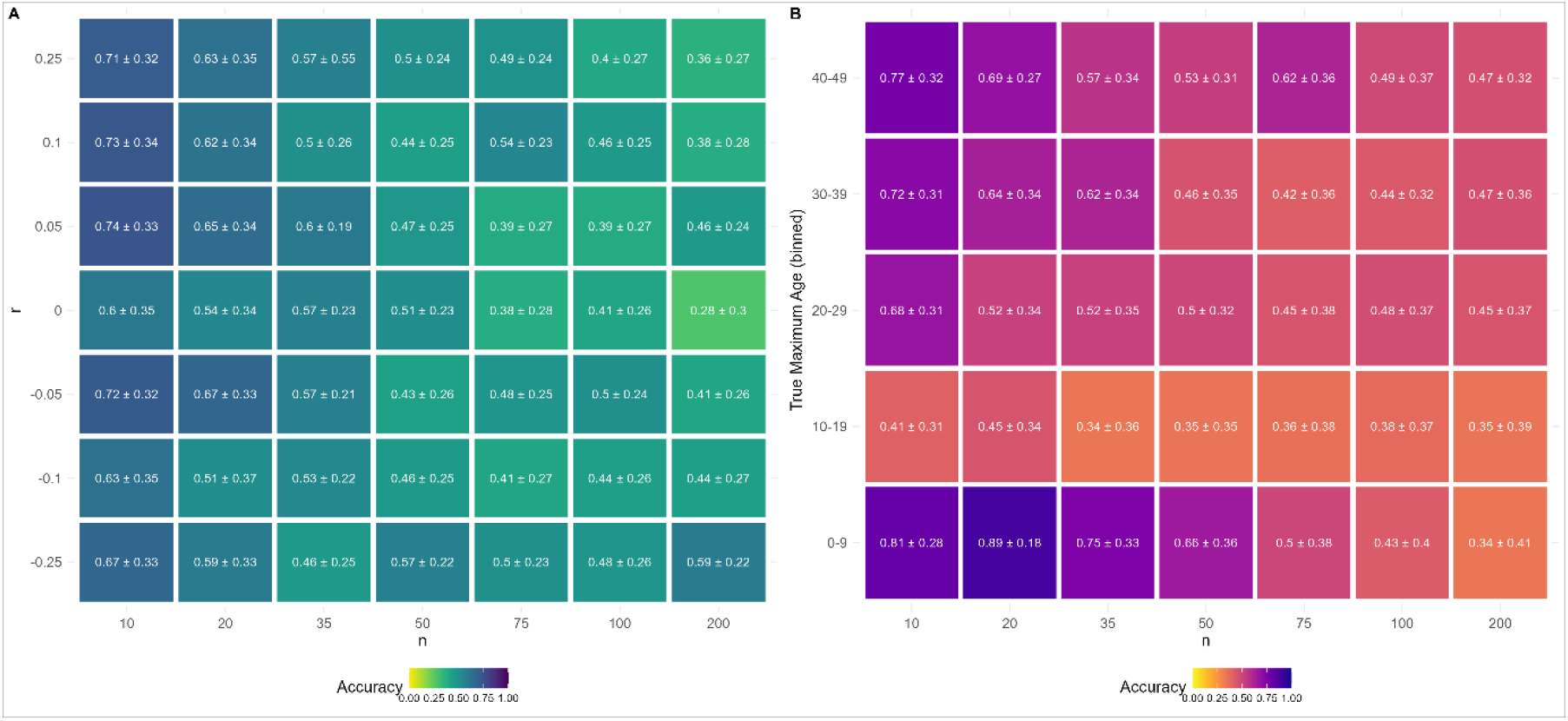
Results from simulations testing the accuracy of the mortality model to detect the true maximum lifespan of a population under a range of A) sample sizes (n) x population growth rates (r) and B) sample sizes x true maximum lifespans. Each cell shows the mean ± std dev. of accuracy over models in each scenario. Cell colour shows the mean accuracy, with lighter colours and smaller values indicating greater accuracy. Accuracy is measured as the minimum credible interval width from the posterior distribution of maximum lifespans required to capture the true maximum lifespan. An accuracy of 0 would indicate the median of the posterior matches the true maximum lifespan, while an accuracy of 0.5 would mean that the true lifespan is captured by the 50% credible interval of the posterior. 100 models were run for each population growth x sample size scenario, resulting in a total of 4900 fitted models. B) is based on the same 4900 fitted models, true maximum lifespans and has a uniform probability of being selected but the number of models applied in each cell will vary. Note that neither sample size nor population growth rate are a regular sequence.

In our second analysis, we compare the mortality estimates from our model to those derived from longitudinal data when applied to the southern-resident killer whales. The longitudinally-derived adult lifespan of female and male southern-resident killer whales is, respectively, 69 (62-79) years [mean (95 % cred. int.)] and 44 (40 -51) years (derived from: (Nielsen et al. 2021a). Our model applied to the southern-resident killer whale population in 1992 estimates lifespans of: 76 (65-88) for females and 44 (38-53) for males. For both males and females there is considerable credible interval overlap between estimates derived from the two models.

### Toothed Whale lifespans

Applying our mortality model to the real Odontocete age-structured data allowed us to generate adult lifespan estimates for 32 female and 33 male toothed whale species (table 3; table 4; figure 1; figure 3). The estimates for each species-sex were based on a median of 75 whales (1^st^ quartile = 39; 3^rd^ quartile = 355). The phylogenetic spread of samples is broad with all of the major Odontocete clades represented (figure 1).

**Table 3.** Estimated female ordinary maximum adult lifespan for 32 species of toothed whale. Where ordinary maximum lifespan is the age at which 90% of adult life years have been lived. All metrics describe the distribution of the posterior derived from the Bayesian mortality model (see methods). Each metric (apart from standard deviation) is the adult lifespan estimated by the model plus the age at maturity to give the expected maximum lifespan for females who reach maturity (Age at Maturity). lCI, and uCI represent the lower and upper bounds respectively of the 50% (l/uCI50) and 95% (l/uCI95) credible intervals.

| Common name | Age at Maturity | Median | ICI50 | uCI50 | ICI95 | uCI95 | Post. Mean | Post. sd |
| --- | --- | --- | --- | --- | --- | --- | --- | --- |
| Atlantic white-sided dolphin | 9 | 21 | 20 | 21 | 19 | 23 | 20.88 | 0.97 |
| Baiji | 6 | 22.5 | 21 | 24 | 20 | 27 | 22.71 | 1.85 |
| Baird's beaked whale | 13 | 52 | 50 | 55 | 46 | 62 | 52.76 | 4.09 |
| Beluga whale | 10 | 54 | 53 | 55 | 52 | 56 | 53.85 | 1.15 |
| Commerson's dolphin | 5 | 17 | 17 | 18 | 15 | 20 | 17.48 | 1.32 |
| Common bottlenose dolphin | 9 | 40 | 39 | 40 | 38 | 41 | 39.76 | 0.82 |
| Common dolphin | 7 | 25 | 25 | 25 | 24 | 26 | 25.04 | 0.42 |
| Dall's porpoise | 5 | 13 | 13 | 13 | 12 | 13 | 12.88 | 0.32 |
| Dwarf sperm whale | 4 | 20 | 19 | 22 | 17 | 25 | 20.53 | 1.92 |
| False killer whale | 10 | 67 | 66 | 69 | 62 | 74 | 67.51 | 2.84 |
| Franciscana | 2 | 13 | 13 | 13 | 12 | 14 | 13.14 | 0.55 |
| Fraser's dolphin | 6 | 20 | 20 | 21 | 18 | 23 | 20.38 | 1.22 |
| Guiana dolphin | 6 | 33 | 32 | 35 | 30 | 38 | 33.45 | 2.09 |
| Harbour porpoise | 3 | 17 | 17 | 18 | 17 | 18 | 17.3 | 0.5 |
| Indo-pacific bottlenose dolphin | 12 | 37 | 36 | 38 | 34 | 41 | 36.92 | 1.85 |
| Indo-pacific humpback dolphin | 9 | 38 | 36 | 40 | 34 | 45 | 38.55 | 3 |
| Killer whale | 13 | 60 | 58 | 62 | 54 | 66 | 59.81 | 3.31 |
| Long-finned pilot whale | 8 | 45 | 45 | 46 | 44 | 46 | 45.17 | 0.68 |
| Melon-headed whale | 7 | 41 | 40 | 42 | 38 | 45 | 41.35 | 1.83 |
| Narrow-ridged finless porpoise | 5 | 23 | 23 | 24 | 21 | 25 | 23.24 | 1.11 |
| Narwhal | 8 | 77 | 74 | 80 | 69 | 86 | 76.81 | 4.4 |
| Northern bottlenose whale | 10 | 27 | 26 | 28 | 24 | 31 | 26.92 | 1.69 |
| Northern right whale dolphin | 9 | 28 | 28 | 29 | 26 | 31 | 28.5 | 1.15 |
| Pacific white-sided dolphin | 9 | 40 | 39 | 42 | 37 | 45 | 40.5 | 1.98 |
| Pantropical spotted dolphin | 10 | 35 | 35 | 35 | 34 | 36 | 35.09 | 0.47 |
| Pygmy sperm whale | 5 | 22 | 21 | 24 | 20 | 26 | 22.58 | 1.73 |
| Risso's dolphin | 9 | 34 | 32 | 35 | 31 | 38 | 33.74 | 1.82 |
| Short-finned pilot whale | 9 | 56 | 55 | 57 | 54 | 58 | 55.81 | 1.14 |
| Sperm whale | 9 | 49 | 48 | 49 | 48 | 50 | 48.7 | 0.61 |
| Spinner dolphin | 8 | 23 | 23 | 23 | 23 | 24 | 23.07 | 0.26 |
| Striped dolphin | 9 | 31 | 31 | 31 | 30 | 32 | 31.11 | 0.51 |
| Vaquita | 5 | 23 | 22 | 24 | 20 | 28 | 23.16 | 2.24 |
| White-beaked dolphin | 8 | 36 | 34 | 37 | 32 | 40 | 35.74 | 2.03 |

**Table 4.** Estimated male ordinary maximum adult lifespan for 33 species of toothed whale. Where ordinary maximum lifespan is the age at which 90% of adult life years have been lived. All metrics describe the distribution of the posterior derived from the Bayesian mortality model (see methods). Each metric (apart from standard deviation) is the adult lifespan estimated by the model plus the age at maturity to give the expected maximum lifespan for males who reach maturity (Age at Maturity). lCI, and uCI represent the lower and upper bounds respectively of the 50% (l/uCI50) and 95% (l/uCI95) credible intervals.

| Common name | Age at Maturity | Median | ICI50 | uCI50 | ICI95 | uCI95 | Post. Mean | Post. sd |
| --- | --- | --- | --- | --- | --- | --- | --- | --- |
| Atlantic white-sided dolphin | 9 | 22 | 21 | 23 | 20 | 25 | 22.21 | 1.2 |
| Baiji | 6 | 20 | 19 | 22 | 18 | 25 | 20.48 | 1.93 |
| Baird's beaked whale | 13 | 87 | 84 | 90 | 78 | 97 | 87.31 | 4.91 |
| Beluga whale | 10 | 49 | 48 | 50 | 46 | 52 | 48.72 | 1.67 |
| Commerson's dolphin | 5 | 22 | 21 | 23 | 19 | 25 | 21.78 | 1.39 |
| Common bottlenose dolphin | 9 | 37 | 36 | 38 | 35 | 39 | 36.89 | 0.99 |
| Common dolphin | 7 | 25 | 24 | 26 | 23 | 28 | 25.32 | 1.46 |
| Dall's porpoise | 5 | 14 | 14 | 14 | 13 | 14 | 13.96 | 0.26 |
| Dwarf sperm whale | 4 | 20 | 19 | 21 | 17 | 24 | 20.12 | 1.75 |
| False killer whale | 10 | 55 | 53 | 58 | 49 | 63 | 55.54 | 3.52 |
| Franciscana | 2 | 13 | 12 | 13 | 12 | 13 | 12.55 | 0.55 |
| Fraser's dolphin | 6 | 19 | 18 | 19 | 17 | 21 | 18.62 | 1.06 |
| Guiana dolphin | 6 | 28 | 27 | 29 | 25 | 31 | 27.64 | 1.61 |
| Harbour porpoise | 3 | 16 | 16 | 17 | 15 | 17 | 16.28 | 0.51 |
| Indian ocean humpback dolphin | 10 | 26 | 25 | 27 | 23 | 30 | 25.99 | 1.65 |
| Indo-pacific bottlenose dolphin | 12 | 36 | 35 | 37 | 33 | 39 | 35.83 | 1.73 |
| Indo-pacific humpback dolphin | 9 | 29 | 27 | 30 | 25 | 33 | 28.75 | 1.96 |
| Killer whale | 13 | 38 | 37 | 39 | 35 | 42 | 38.13 | 1.73 |
| Long-finned pilot whale | 8 | 35 | 35 | 36 | 34 | 37 | 35.45 | 0.84 |
| Melon-headed whale | 7 | 35 | 34 | 37 | 32 | 40 | 35.57 | 1.97 |
| Narrow-ridged finless porpoise | 5 | 23 | 22 | 23 | 21 | 25 | 22.64 | 0.99 |
| Narwhal | 8 | 66 | 63 | 69 | 58 | 75 | 66.27 | 4.28 |
| Northern bottlenose whale | 10 | 36 | 35 | 37 | 32 | 40 | 35.89 | 1.93 |
| Northern right whale dolphin | 9 | 24 | 23 | 25 | 22 | 26 | 23.93 | 1.2 |
| Pacific white-sided dolphin | 9 | 38 | 37 | 39 | 34 | 42 | 37.88 | 1.92 |
| Pantropical spotted dolphin | 10 | 32 | 31 | 32 | 31 | 33 | 31.76 | 0.58 |
| Rough-toothed dolphin | 10 | 30 | 29 | 32 | 27 | 35 | 30.63 | 1.91 |
| Short-finned pilot whale | 9 | 37 | 36 | 38 | 34 | 39 | 36.64 | 1.31 |
| Sperm whale | 9 | 53 | 52 | 53 | 51 | 54 | 52.5 | 0.63 |
| Spinner dolphin | 8 | 19 | 19 | 20 | 18 | 20 | 19.3 | 0.56 |
| Striped dolphin | 9 | 30 | 30 | 31 | 29 | 31 | 30.12 | 0.68 |
| Vaquita | 5 | 21 | 20 | 23 | 18 | 27 | 21.63 | 2.32 |
| White-beaked dolphin | 8 | 29 | 28 | 31 | 26 | 34 | 29.52 | 2.08 |

**Figure 3.**
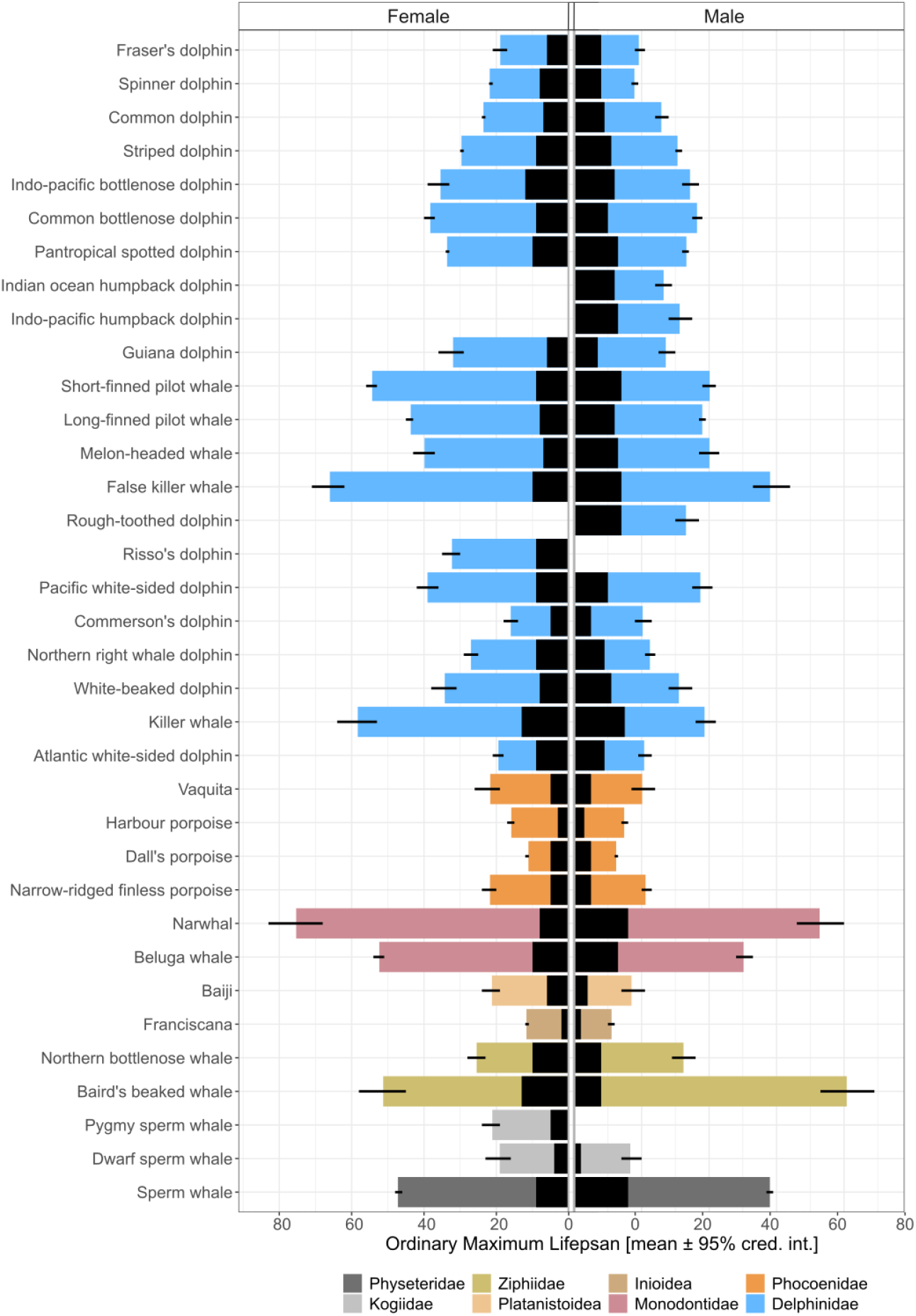
Adult lifespans of female (left panel) and male (right panel) toothed whales estimated by our mortality model. Bars show a mean estimate of the ordinary maximum lifespan - the age by which 90% of adult life years have been lived - derived from our mortality model. Error bars show the 95% credible interval of lifespan estimates. Black areas show the juvenile years not estimated by our model. Ordinary maximum lifespan, therefore, represents lifespan given survival to maturity. Species order and bar colours reflect phylogeny (figure 1). For some species data were available for only one sex - resulting in an empty bar.

Male Baird’s beaked whales have the longest lifespan in our sample, with an ordinary maximum lifespan of 84-90 years (50% credible interval [50CI]). The longest-lived females in our sample are Narwhals with an ordinary maximum lifespan of 74-80 years (50CI). The shortest-lived species in our sample are female Dall’s Porpoises and Franciscana of both sexes all of which have a median ordinary maximum lifespan of 13 years. Our additional analyses demonstrated that the ordinary maximum lifespans derived from the model are largely robust to differences in potential systematic errors in ageing of older whales (supplementary 2), changing ageing methodologies through time (supplementary 3), our population changes assumptions (supplementary 4) and the structure of our model (supplementary 7).

There is good evidence in 10 species that females have a longer adult lifespan than males (proportion of posterior where female – male lifespan is greater than 0 > 0.97; figure 4). However, in 4 species – Baird’s beaked whales, Commerson’s dolphin, northern bottlenose whales and sperm whales - the opposite is true and there is clear evidence that males live longer as adults than females (proportion of posterior female – male lifespan is less than 0 >0.975; figure 4). In the remaining species, there is no clear evidence that the sexes have different lifespans. Combining all species female toothed whales live a median of 10% longer than males (female – male lifespan / female lifespan), but overall there is no clear evidence of a sex difference in lifespan (proportion of posterior female – male lifespan > 0 = 0.66).

**Figure 4.**
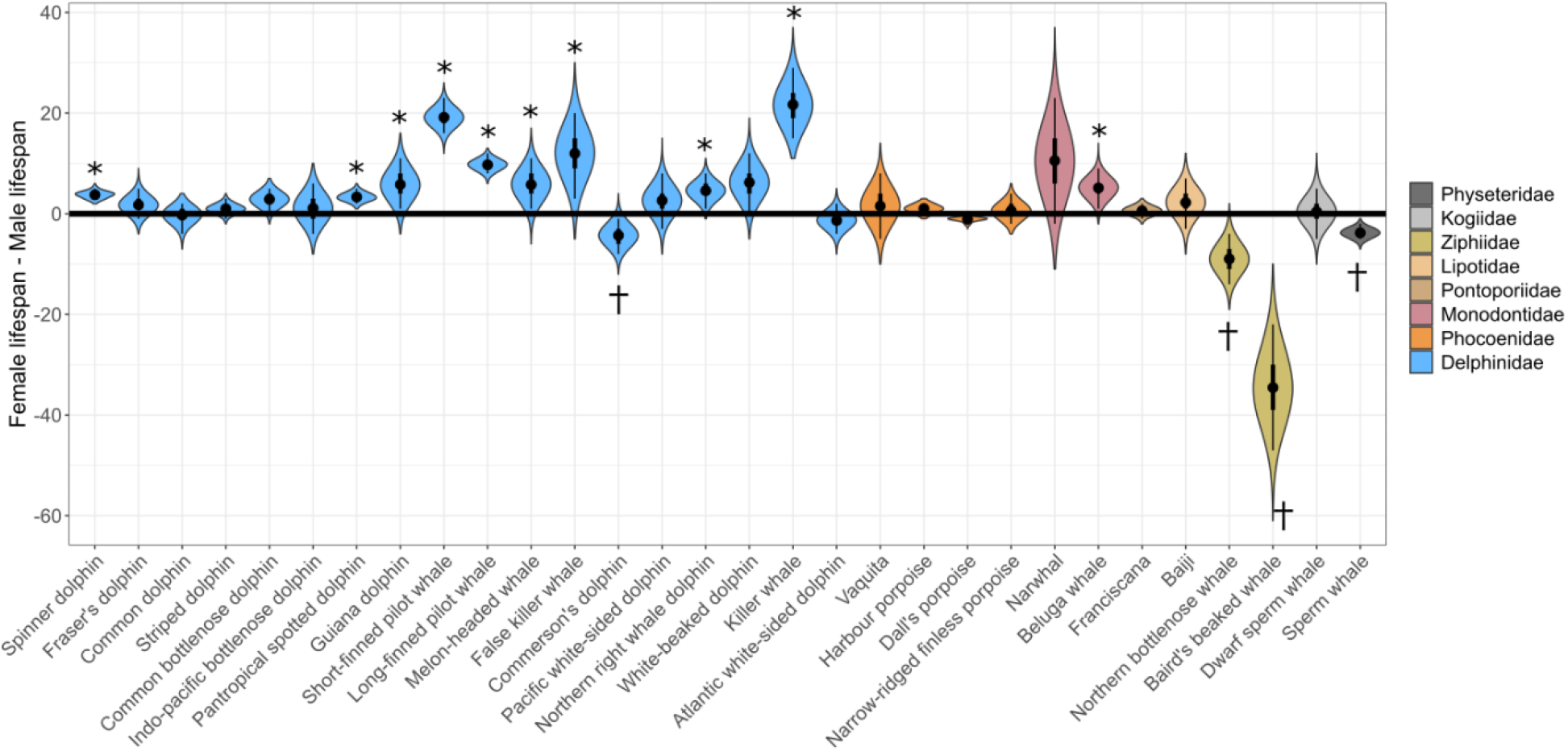
Comparing female and male adult lifespan in toothed whales. Violins show the posterior distribution of female–male lifespan derived from our mortality models. Both sexes are assessed in the same model so the posterior lifespan of each sex at each draw can be directly compared. * indicate species where there is good evidence that females live longer than males (proportion of posterior greater than 0 >0.975). † indicate species where there is good evidence that males live longer than females (proportion of posterior less than 0 >0.975). Species order and violin colour reflect phylogeny (figure 1).

## Discussion

Here we have developed and applied a method to estimate lifespan parameters from age-structured cross-sectional samples. Simulations demonstrate that the models can capture true lifespan given population growth, sampling error and age-estimation error under a realistic range of sample sizes and species maximum lifespans. Applying the model to data collected from the literature provides adult lifespan estimates for 32 female and 33 male species of toothed whale.

We find that there is almost an order of magnitude difference between the ordinary maximum lifespan of the shortest-lived and longest-lived toothed whales. This variation is particularly remarkable given that many other aspects of toothed whale life history do not differ between species. For example, all toothed whales: give birth to a single offspring per breeding event; provide several years of maternal year to their dependent offspring; and exhibit a polygynous mating system with no evidence of paternal care (Whitehead and Mann 2000; Chivers 2018). At the broadest level, toothed whale species also share a similar ecological niche, all are aquatic carnivores and most live and often hunt in social groupings. Despite these similarities here, we have shown that lifespan can vary considerably between species. This variation highlights the importance of toothed whales as a model of life history evolution. Future work using toothed whales to understand how ecological and social factors predict lifespan evolution has the potential to provide important fundamental insights.

We found that in 10 species of toothed whale there is good evidence that females live longer as adults than males - and overall female toothed whales live 10% longer than males. Longer-lived females are the general pattern in mammals: for example, a recent analysis of 101 mammal species – including four toothed whales - found that female mammals live a mean of 18.6% longer than males (Lemaître et al. 2020). The evolutionary drivers of sex differences in adult lifespan in mammals remain unresolved but both genetic conflict and life history trade-offs have been implicated (Lemaître et al. 2020). However, in toothed whales, we found that there are exceptions to this pattern. Perhaps most notably, in Baird’s beaked whales’ males live considerably longer than females: the median predicted ordinary maximum lifespan of females is 52 years compared to 87 for males. In Baird’s beaked whales, this sex difference in lifespan leads to heavily male-biased adult populations but the evolutionary reasons for the extreme lifespan differences in this species remain unknown (Kasuya et al. 1997a).

On the other hand, the comparative rarity of sex differences in toothed whale adult lifespan we find here is also evidence in favour of the argument that sex differences in mammals have been overstated. For example, a recent broad scale analysis of mammals found that, contrary to established wisdom, only a 45% of mammals demonstrate male-biased sex-size dimorphism (Tombak et al. 2024).

Similarly, while Lemaître et al (2020) found that on average female mammals live-longer than males, the pattern was in fact very variable between species. Reflecting this, we only found evidence of sex differences in lifespan in 53% of toothed whale species in our sample. Overall, there is considerable variability in the presence, direction and size of sex differences in adult lifespan in this taxa making toothed whales an important taxonomic group in which to test theories investigating the presence and absence of sex differences in lifespans.

This result of sex differences analysis highlights a strength of our modelling approach as rather than simply comparing the final realised maximum lifespans we can compare female and male lifespans at each posterior model draw. More generally, by taking a Bayesian approach we can embrace the uncertainty inherent in age-structured data and still draw useful inferences from uncertain data. The important implication of this is that this analysis aims to capture a distribution of plausible values of lifespan for toothed whale species-rather than to define a single ‘best-guess’ point estimate. Any future studies using these lifespan estimates should take this into account by carrying error through their analysis and not simply relying on a point-estimate of lifespan (e.g. supplementary 6; Ellis et al. 2024). It is also important to note that although throughout this study we have focussed on lifespan as our metric of species survival, our methods could also be used to derive distributions of other measures of age-specific mortality (e.g. measures of mortality shape (Healy et al. 2019) which might be more relevant for answering some questions.

Our approach builds upon and generalises concepts and methods that have been used to estimate toothed whale lifespans from age-structured data. Conceptually, our Bayesian approach – which uses priors from other species – has similarities to earlier methods based on scaling known ‘model’ lifetables from other species to toothed whales (Barlow and Boveng 1991; Caswell et al. 1998).

Unlike previous Bayesian approaches to using age-structured data to estimate toothed whale life history, we aim to capture the distribution of mortality parameters consistent with the data and the unknown effects of population growth and sampling bias, rather than explicitly estimating populating growth and sampling bias parameters (Moore and Read 2008; Saavedra 2018; Rouby et al. 2021). As a consequence our approach can estimate usefully narrow distributions of life history parameters based on relatively smaller sample sizes than previous methods, but – unlike those methods – is not suited in its current form to calculating growth rates or the demographic consequences of human-induced mortality in a given population.

The methods presented here derive the same or similar estimates of ordinary maximum lifespan estimates as lifespan estimates derived from other methods applied to the same datasets (table 5). However, these ‘single dataset’ lifespan measures often differ from the species-level measure derived by applying the model to multiple datasets (table 5). This highlights the value of our approach-by combining datasets from multiple populations we can both increase our sample size and also derive a more general species lifespan which is less likely to be driven by characteristics of a single population. Lifespan is only one of the traits that contribute to a species’ life history, it is nevertheless important to quantify and understand in its own right.

**Table 5.** Comparison of adult ordinary maximum lifespan estimates derived from the method developed and applied here (species estimate, dataset estimate) to other methods (published estimate). ‘Published estimates’ are sex-specific estimates of age-specific mortality that have been converted into adult ordinary maximum lifespan for comparison with our model. ‘Species estimate’ is the ordinary maximum lifespan as reported in table 3 & 4 using all of the data available for a particular species-sex. ‘Dataset estimate’ is the ordinary maximum lifespan derived by applying our mortality model just to the datasets used to derive the Published estimate. Both species and dataset estimate are reported as a 95% credible interval. Published methods are a Siler model applied to age-at-death data, ‘mark-recapture’ from a longitudinal study (see text) and a ‘scaled mammal’ estimate derived by rescaling a known mammal life history to fit the toothed whale data.

| Species | Sex | Species estimate<br>(95% cred. int.) | Dataset estimate<br>(95% cred. int.) | Published estimate | Published method used<br>(reference) |
| --- | --- | --- | --- | --- | --- |
| Common Bottlenose Dolphin | F | 38-41 | 33-42 | 30 | Age-at-death Siler model<br>(Stolen and Barlow 2003) |
| Common Bottlenose Dolphin | M | 35-39 | 29-37 | 26 | Age-at-death Siler model<br>(Stolen and Barlow 2003) |
| Killer Whale | F | 54-66 | 65-88 | 62-79<br>(95% cred. int.) | Mark-recapture<br>(Nielsen et al. 2021b) |
| Killer Whale | M | 35-42 | 38-53 | 40-51<br>(95% cred. int.) | Mark-recapture<br>(Nielsen et al. 2021b) |
| Pantropical Spotted Dolphin | F | 34-36 | 32-34 | 31 | Scaled mammal<br>(Barlow and Hohn 1984) |
| Pantropical Spotted Dolphin | M | 31-33 | 34-36 | 32 | Scaled mammal<br>(Barlow and Hohn 1984) |

Lifespan can be viewed as an emergent property of other process such as senescence (Jones et al. 2014; Shefferson et al. 2017). To understand senescence therefore requires an understanding of species lifespans and shapes of mortality (Ronget and Gaillard 2020), both of which can be derived from the outputs of our model. Lifespan is also hypothesised to have an important role in social evolution (e.g. (Silk and Hodgson 2021; Aubier and Kokko 2022)). For example, in a number of species (Péron et al. 2019) including toothed whales (Foster et al. 2012; Nattrass et al. 2019) older individuals have been shown to have a positive influence on the fitness of their younger relatives and group mates. In killer whales, one of the mechanisms driving this positive influence is the value of long-term ecological knowledge in allowing older females to lead their group to resources (Brent et al. 2015). The first step to understanding how the distribution of ecological knowledge differs across a social group, and how this therefore allows social species to thrive in spatially and temporally complex ecologies, is to understand how long individuals live. Quantifying lifespan is important, therefore, to understanding social as well as life history evolution.

It will be decades before longitudinal studies can provide the data necessary to provide estimates of life history metrics by currently established methods for most toothed whale species. For many species, utilising age-structured data are the only way we will be able to begin to understand their life history for the foreseeable future. In this study, we utilised modern statistical methodology, and the power of modern computing, to derive age-specific mortality estimates for 44% of extant toothed whale species, representing an important step in understanding the evolution and diversity of Odontocete life history.

## Data availability statement

The data underlying this article are avalaible in the *marinelifehistdata* R package at github.com/samellisq/marinelifehistdata.

## Acknowledgements

This project was funded as part of a Leverhulme Trust Early Career Research Fellowship awarded to SE. DPC, DWF & MNW acknowledge funding from a NERC Standard Grant (no. NE S010327/1) and MLKN acknowledges funding from a NERC PhD studentship. The authors would also like to thank members of the Centre for Research in Animal Behaviour at the University of Exeter for comments throughout the development of this project.

## Supporting information

S1 Modelling pathway applied to toothed whale data.

S2 Exploring the effect of systematic uncertainty in older whales

S3 Investigating the effect of changing whale ageing techniques

S4 Sensitivity analysis for population change assumptions

S5 Age-at-death model reformulation S6 Phylogenetic signal analysis

S7 Age estimation error model structure comparison S8 Simulation Credible Interval Widths

## References

Addink M, Hartmann MG, Couperus B (1997) A note on life-history parameters of the Atlantic white-sided dolphin (Lagenorhynchus acutus) from animals bycaught in the Northeastern Atlantic. Reports of the International Whaling Commission 47:637–639

Amano M, Miyazaki N (2004) Composition of a school of Risso’s dolphins, Grampus griseus. Marine Mammal Science 20:152–160

Amano M, Miyazaki N, Yanagisawa F (1996) Life hisotry of Fraser’s dolphin, Lagenodelphis hosei, based on a school captured off the pacific coast of Japan. Marine Mammal Science 12:199– 214

Amano M, Yamada TK, Brownell RL, Uni Y (2011) Age determination and reproductive traits of killer whales entrapped in ice off Aidomari, Hokkaido, Japan. Journal of Mammalogy 92:275–282. 10.1644/10-MAMM-A-276.1

Amano M, Yamada TK, Kuramochi T, et al (2014) Life history and group composition of melon-headed whales based on mass strandings in Japan. Marine Mammal Science 30:480–493. 10.1111/mms.12050

Arai K, K. Yamada T, Takano Y (2004) Age estimation of male Stejneger’s beaked whales (Mesoplodon stejnegeri) based on counting of growth layers in tooth cementum. Mammal Study 29:125–136. 10.3106/mammalstudy.29.125

Aubier TG, Kokko H (2022) Volatile social environments can favour investments in quality over quantity of social relationships. Proceedings of the Royal Society B: Biological Sciences 289:20220281. 10.1098/rspb.2022.0281

Balcomb KC (1989) Baird’s Beaked Whale Berardius bairdii Stejneger, 1883: Arnoux’s Beaked Whale Berardius arnuxii Duvernoy, 1851. In: Ridgway SH, Harrison R (eds) Handbook of Marine Mammals volume 4: River Dolphins and the Larger Toothed Whales. Academic Press, London, pp 261–288

Barlow J, Boveng P (1991) Modeling Age-Specific Mortality for Marine Mammal Populations. Marine Mammal Science 7:50–65. 10.1111/j.1748-7692.1991.tb00550.x

Barlow J, Hohn A (1984) Interpreting spotted dolphin age distributions. NOAA Techincal Memoradum NMFS

Barratclough A, McFee WE, Stolen M, et al (2023) How to estimate age of old bottlenose dolphins (Tursiops truncatus); by tooth or pectoral flipper? Frontiers in Marine Science 10:

Barreto AS, Rosas FCW (2006) Comparative growth analysis of two populations of Pontoporia blainvillei on the Brazilian coast. Marine Mammal Science 22:644–653. 10.1111/j.1748-7692.2006.00040.x

Best PB, Canham PAS, Macleod N (1984) Patterns of reproduction in sperm whales, Physeter macrocephalus. Report of the International Whaling Commission 51–80

Best PB, Meÿer MA, Lockyer C (2010) Killer whales in South African waters -a review of their biology. African Journal of Marine Science 32:171–186. 10.2989/1814232x.2010.501544

Betty EL (2019) Life history of the long-finned pilot whale (Globicephala melas edwardii); insights from strandings on the New Zealand coast. Auckland University of Technology

Bigg MA, Olesiuk PF, Ellis GM, et al (1990) Social organization and genealogy of resident killer whales (Orcinus orca) in the coastal waters of British Columbia and Washington State. Report of the International Whaling Commission, Special 383–405

Bloch D, Desportes G, Harvey P, et al (2012) Life history of Risso’s dolphin (Grampus griseus) (G. Cuvier, 1812) in the Faroe Islands. Aquatic Mammals 38:250–266. 10.1578/AM.38.3.2012.250

Bloch D, Lockyer C, Zachariassen M (1993) Age and growth parameters of the long-finned pilot whale off the Faroe Islands. Report of the International Whaling Commission 163–207

Botta S, Secchi ER, Muelbert MMC, et al (2010) Age and growth of franciscana dolphins, Pontoporia blainvillei (Cetacea: Pontoporiidae) incidentally caught off southern Brazil and northern Argentina. Journal of the Marine Biological Association of the United Kingdom 90:1493– 1500. 10.1017/S0025315410001141

Boy CC, Dellabianca N, Goodall RNP, Schiavini ACM (2011) Age and growth in Peale’s dolphin (Lagenorhynchus australis) in subantarctic waters off southern South America. Mammalian Biology 76:634–639. 10.1016/j.mambio.2011.03.001

Brent LJN, Franks DW, Foster EA, et al (2015) Ecological knowledge, leadership, and the evolution of menopause in killer whales. Current Biology 25:746–750. 10.1016/j.cub.2015.01.037

Burns JJ, Seaman GA (1986) Investigations of belukha whales in coastal waters of western and northern Alaska. II. Biology and Ecology

Butti C, Corain L, Cozzi B, et al (2007) Age estimation in the Mediterranean bottlenose dolphin Tursiops truncatus (Montagu 1821) by bone density of the thoracic limb. Journal of Anatomy 211:639–646. 10.1111/j.1469-7580.2007.00805.x

Cáceres-Saez I, Dellabianca NA, Pimper LE, et al (2015) Sexual dimorphism and morphometric relationships in pelvic bones of Commerson’s dolphins (Cephalorhynchus c. commersonii>) from Tierra del Fuego, Argentina. Marine Mammal Science 31:734–747. 10.1111/mms.12172

Caswell H, Brault S, Read AJ, Smith TD (1998) Harbor Porpoise and Fisheries: An Uncertainty Analysis of Incidental Mortality. Ecological Applications 8:1226–1238. 10.1890/1051-0761(1998)008[1226:HPAFAU]2.0.CO;2

Caughley G (1966) Mortality Patterns in Mammals. Ecology 47:906–918. 10.2307/1935638

Chivers SJ (2018) Cetecean life history. In: Würsig B, Thewissen JGM, Kovacs KM (eds) Encyclopedia of Marine Mammals, 3rd edn. Academic Press, London, pp 186–187

Christensen I (1984) Growth and reproduction of killer whales, Orcinus orca, in Norwegian coastal waters. Report of the International Whaling Commission 253–258

Christensen I (1973) Age determination, age distribution and growth of bottlenose whales, Hyperoodon ampullatus (Foster) in the Labrador sea. Norwegian Journal of Zoology 21:331– 340

Cipriano FW (1992) Behavior and occurrence patterns, feeding ecology, and life history of dusky dolphins (Lagenorhynchus obscurus) off Kaikoura, New Zealand. University of Arizona

Clapham PJ, Childerhouse S, Gales NJ, et al (2007) The whaling issue: Conservation, confusion, and casuistry. Marine Policy 31:314–319. 10.1016/j.marpol.2006.09.004

Clarke R, Paliza O, Van Waerebeek K (2013) Sperm whales of the Southeast Pacific. Part VII. Reproduction and growth in the female. Latin American Journal of Aquatic Mammals 9:8–39. 10.5597/lajam00172

Cockcroft VG, Ross GJB (1990) Age Growth and Reproduction of Bottlenose Dolphins Tursiops truncatus from the East Coast of Southern Africa. Fishery Bulletin 88:289–302

Colchero F, Jones OR, Rebke M (2012) BaSTA: An R package for Bayesian estimation of age-specific survival from incomplete mark-recapture/recovery data with covariates. Methods in Ecology and Evolution 3:466–470. 10.1111/j.2041-210X.2012.00186.x

Collet A, Robineau D (1988) Data on the genital tract and reproduction of Commerson’s dolphin, Cephalorynchus commersonii (Lacepede, 1804), from the Kergulen Islands. Report of the International Whaling Commission (Special Issue 9) 119–141

Connor RC, Mann J, Tyack PL, Whitehead H (1998) Social evolution in toothed whales. Trends in Ecology and Evolution 13:228–232. 10.1016/S0169-5347(98)01326-3

Conversani VRM, Silva DF, Barbosa RA, et al (2021) Age and growth of franciscana, Pontoporia blainvillei, and Guiana, Sotalia guianensis, dolphins from southeastern Brazil. Marine Mammal Science 37:702–716. 10.1111/mms.12763

Danil K, Chivers SJ (2007) Growth and reproduction of female short-beaked common dolphins, Delphinus delphis, in the eastern tropical Pacific. Canadian Journal of Zoology 85:108–121. 10.1139/Z06-188

Denuncio P, Negri MF, Bastida R, Rodríguez D (2018) Age and growth of Franciscana dolphins from northern Argentina. Journal of the Marine Biological Association of the United Kingdom 98:1197–1203. 10.1017/S0025315417000765

Dong JH, Lien J, Nelson D, Curren K (1996) A contribution to the biology of the white-beaked dolphin, Lagenorhynchus albirostris, in waters off Newfoundland. The Canadian Field-Naturalist 110:278–287

Durante CA, Santos-Neto EB, Azevedo A, et al (2016) POPs in the South Latin America: Bioaccumulation of DDT, PCB, HCB, HCH and Mirex in blubber of common dolphin (Delphinus delphis) and Fraser’s dolphin (Lagenodelphis hosei) from Argentina. Science of the Total Environment 572:352–360. 10.1016/j.scitotenv.2016.07.176

Ellis S, Franks DW, Nattrass S, et al (2018a) Postreproductive lifespans are rare in mammals. Ecology and Evolution 8:2482–2494. 10.1002/ece3.3856

Ellis S, Franks DW, Nattrass S, et al (2018b) Analyses of ovarian activity reveal repeated evolution of post-reproductive lifespans in toothed whales. Scientific Reports 8:1–10. 10.1038/s41598-018-31047-8

Ellis S, Franks DW, Nattrass S, et al (2017) Mortality risk and social network position in resident killer whales: sex differences and the importance of resource abundance. Proceedings of the Royal Society B 284:20171313. http://dx.doi.org/10.1098/rspb.2017.1313

Ellis S, Franks DW, Nielsen MLK, et al (2024) The evolution of menopause in toothed whales. Nature 627:579–585. 10.1038/s41586-024-07159-9

Evacitas FC, Kao WY, Worthy GAJ, Chou LS (2017) Annual variability in dentin δ15N and δ13C reveal sex differences in weaning age and feeding habits in Risso’s dolphins (Grampus griseus). Marine Mammal Science 33:748–770. 10.1111/mms.12396

Evans K, Hindell MA (2004) The age structure and growth of female sperm whales (Physeter macrocephalus) in southern Australian waters. Journal of Zoology 263:237–250. 10.1017/S0952836904005096

Ferguson SH, Willing C, Kelley TC, et al (2020) Reproductive parameters for female beluga whales (Delphinapterus leucas) of baffin bay and hudson bay, Canada. Arctic 73:405–420. 10.14430/arctic71435

Ferreira IM, Kasuya T, Marsh H, Best PB (2014) False killer whales (Pseudorca crassidens) from Japan and South Africa: Differences in growth and reproduction. Marine Mammal Science 30:64–84. 10.1111/mms.12021

Ferrero RC, Walker A (1999) Age, Growth, and Reproductive Patterns of Dall’s Porpoise (Phocoenoides dalli) in the Central North Pacific Ocean. Marine Mammal Science 15:273– 313. 10.1111/j.1748-7692.1999.tb00803.x

Ferrero RC, Walker WA (1995) Growth and reproduction of the common dolphin, Delphinus delphis Linnaeus, in the offshore waters of the North Pacific Ocean. Fishery Bulletin 93:483–494

Ferrero RC, Walker WA (1993) Growth and reproduction of the northern right whale dolphin Lissodelphis borealis in the offshore waters. Canadian Journal of Zoology 71:2335–2344

Ferrero RC, Walker WA (1996) Age, growth, and reproductive patterns of the Pacific white-sided dolphin (Lagenorhynchus obliquidens) taken in high seas drift nets in the central North Pacific Ocean. Canadian Journal of Zoology 74:1673–1687. 10.1139/z96-185

Foote AD (2008) Mortality rate acceleration and post-reproductive lifespan in matrilineal whale species. Biology Letters 4:189–191. 10.1098/rsbl.2008.0006

Foster EA, Franks DW, Mazzi S, et al (2012) Adaptative prolonged postreproductive life span in killer whales. Science 337:1313

Gaillard J-M, Garratt M, Lemaitre JF (2017) Senescence in mammalian life history traits. In: Shefferson RP, Jones OR, Salguero-Gómez R (eds) The evolution of senescence in the tree of life. Cambridge University Press, Cambridge, pp 126–155

Gaillard J-M, Viallefont A, Loison A, Festa–Bianchet M (2004) Assessing senescence patterns in populations of large mammals. Animal Biodiversity and Conservation 27:47–58

Galatius A, Jansen OE, Kinze CC (2013) Parameters of growth and reproduction of white-beaked dolphins (Lagenorhynchus albirostris) from the North Sea. Marine Mammal Science 29:348– 355. 10.1111/j.1748-7692.2012.00568.x

Gao A, Zhou K (1993) Growth and reproduction of three populations of finless porpoise (Neophocaena phocaenoides) in Chinese waters. Aquatic Mammals 19:3–12

Gao A, Zhou K (1992) Sexual dimorphism in the baiji, Lipotes vexillifer. Canadian Journal of Zoology 70:1484–1493. 10.1139/z92-205

Garde E, Hansen SH, Ditlevsen S, et al (2015) Life history parameters of narwhals (Monodon monoceros) from Greenland. Journal of Mammalogy 96:866–879. 10.1093/jmammal/gyv110

Gol’din P (2004) Growth and Body Size of the Harbour Porpoise, Phocoena phocoena (Cetacea, Phocoenidae), in the Sea of Azov and the Black Sea. Vestnik zoologii 38:59–73

Gol’din P, Gladilina E (2015) Small dolphins in a small sea: age, growth and life-history aspects of the black sea common bottlenose dolphin Tursiops truncatus. Aquatic Biology 23:159–166. 10.3354/ab00617

Gompertz B (1825) On the nature of the function expressive of the law of human mortality, and on a new mode of determinng the value of life contingencies. Philosophical Transactions of the Royal Society 115:513–583

Gowans S, Würsig B, Karczmarski L (2007) The social structure and strategies of delphinids: predictions based on an ecological framework. Advances in Marine Biology 53:195–294. 10.1016/S0065-2881(07)53003-8

Guo L, Lin W, Zeng C, et al (2020) Investigating the age composition of Indo-Pacific humpback dolphins in the Pearl River Estuary based on their pigmentation pattern. Marine Biology 167:1–12. 10.1007/s00227-020-3650-x

Hamlin KL, Pac DF, Sime CA, et al (2000) Evaluating the Accuracy of Ages Obtained by Two Methods for Montana Ungulates. The Journal of Wildlife Management 64:441–449. 10.2307/3803242

Hay KA (1984) The life history of the Narwal (Mondon monoceros L.) in the Eastern Canadian Arctic. PhD Thesis, McGill University

Healy K, Ezard THG, Jones OR, et al (2019) Animal life history is shaped by the pace of life and the distribution of age-specific mortality and reproduction. Nature Ecology & Evolution. 10.1038/s41559-019-0938-7

Heide-Jørgensen MP, Lockyer C (2001) Age and sex distributions in the catches of belugas, Delphinapterus leucas, in West Greenland and in western Russia. Mammalian Biology 66:215–227

Herman DP, Matkin CO, Ylitalo M. G, et al (2008) Assessing age distributions of killer whale Orcinus orca populations from the composition of endogenous fatty acids in their outer blubber layers. Marine Ecology Progress Series 372:289–302. 10.3354/meps07709

Hohn AA, Read AJ, Fernandez S, et al (1996) Life history of the vaquita, Phocoena sinus (Phocoenidae, Cetacea). Journal of Zoolgy 239:235–251

Iwasaki T, Kasuya T (1997) Life-history and catch bias of Pacific white-sided (Lagenorhynchus obliquidens) and Northern right whale dolphins(Lissodelphis borealis) incidentally taken by the Japanese high seas squid driftnet fisher. Report of the International Whaling Commission 47:683–692

IWC (2021) Taxonomy: classification of Cetacea. In: International Whaling Commission/About Whales

Jefferson TA, Hung SK, Robertson KM, Archer FI (2012) Life history of the Indo-Pacific humpback dolphin in the Pearl River Estuary, southern China. Marine Mammal Science 28:84–104. 10.1111/j.1748-7692.2010.00462.x

Jefferson TA, Robertson KM, Wang JY (2002) Growth and reproduction of the finless porpoise in southern China. Raffles Bulletin of Zoology 105–113

Jones OR, Scheuerlein A, Salguero-Gómez R, et al (2014) Diversity of ageing across the tree of life. Nature 505:169–173. 10.1038/nature12789

Kasuya T (1978) The life history of Dall’s porpoise with special reference to the stock off the Pacific coast of Japan. Scientific Reports of the Whales Research Institute, Tokyo 1–63

Kasuya T (1986) False killer whales. In: Tamura T, Ohsumi S, Arai S (eds) Report of the investigation in search of solution for dolphin fishery conflict in the Iki island. Japan Fisheries Agency, pp 178–187

Kasuya T (1972) Some informations on the growth of the Ganges Dolphin with a comment on the Indus Dolphin. Scientific Reports of the Whales Research Institute 24:87–108

Kasuya T (1999) Finless Porpoise Neophocaena phocaenoides (G. Cuvier, 1829). In: Ridgway SH, Harrison R (eds) Handbook of Marine Mammals Volume 6: The Second Book of Dolphins and the Porpoises. Academic Press, London, pp 411–442

Kasuya T (1985) Effect of exploitation on reproductive parameters of the spotted and striped dolphins off the Pacific coast of Japan. Sci Rep Whales Res Inst, Tokyo 107–138

Kasuya T (1991) Density Dependent Growth in North Pacific Sperm Whales. Marine Mammal Science 7:230–257. 10.1111/j.1748-7692.1991.tb00100.x

Kasuya T, Brownell Jr RL, Balcomb KC (1997a) Life history of Baird’s beaked whales. Report of the International Whaling Commission 47:969–979

Kasuya T, Brownell RLJ (1979) Age determination, reproduction, and growth of the Franciscana Dolphin Pontoporia blainvillei. Scientific Reports of the Whales Research Institute Tokyo 31:45–67

Kasuya T, Izumisawa Y, Komyo Y, et al (1997b) Life history parameters Bottlenose dolphins off Japan. IBI Reports 7:71–107

Kasuya T, Marsh H (1984) Life history and reproductive biology of the short-finned pilot whale, Globicephala macrorhynchus, off the Pacific coast of Japan. Report of the International Whaling Commission (Special Issue 6) 259–310

Kasuya T, Shiraga S (1985) Growth of Dall’s Porpoise in the western North Pacific and suggested geographical growth differentiation. Scientific Reports of the Whales Research Institute, Tokyo 36:139–152

Kasuya T, Tai S (1993) Life history of short-finned pilot whale stocks off Japan and a description of the fishery. Report of the International Whaling Commission SI14:35

Kebke A, Samarra F, Derous D (2022) Climate change and cetacean health: impacts and future directions. Philosophical Transactions of the Royal Society B: Biological Sciences 377:20210249. 10.1098/rstb.2021.0249

Kemper C, Talamonti M, Bossley M, Steiner A (2019) Sexual maturity and estimated fecundity in female Indo-Pacific bottlenose dolphins (Tursiops aduncus) from South Australia: Combining field observations and postmortem results. Marine Mammal Science 35:40–57. 10.1111/mms.12509

Kemper CM, Trentin E, Tomo I (2014) Sexual maturity in male Indo-Pacific bottlenose dolphins (Tursiops aduncus): Evidence for regressed/pathological adults. Journal of Mammalogy 95:357–368. 10.1644/13-MAMM-A-007.1

Kesselring T, Viquerat S, Brehm R, Siebert U (2018) Coming of age: - Do female harbour porpoises (Phocoena phocoena) from the North Sea and Baltic Sea have sufficient time to reproduce in a human influenced environment? PLoS ONE 12:e0186951. 10.1371/journal.pone.0199633

Konno K, Akasaka M, Koshida C, et al (2020) Ignoring non-English-language studies may bias ecological meta-analyses. Ecology and Evolution 10:6373–6384. 10.1002/ece3.6368

Kurihara N, Amano M, Yamada TK (2016) Decrease in tooth count in melon-headed whales. Journal of Zoology 300:8–17. 10.1111/jzo.12363

Larese JP, Chivers SJ (2009) Growth and reproduction of female eastern and whitebelly spinner dolphins incidentally killed in the eastern tropical Pacific tuna purse-seine fishery. Canadian Journal of Zoology 87:537–552. 10.1139/Z09-038

Learmonth JA, Murphy S, Luque PL, et al (2014) Life history of harbor porpoises (Phocoena phocoena) in Scottish (UK) waters. Marine Mammal Science 30:1427–1455. 10.1111/mms.12130

Lee YR, An YR, Park KJ, et al (2013) Age and reproduction of the finless porpoises, Neophocaena asiaeorientalis, in the Yellow Sea, Korea. Animal Cells and Systems 17:366–373. 10.1080/19768354.2013.851116

Lemaître JF, Ronget V, Tidière M, et al (2020) Sex differences in adult lifespan and aging rates of mortality across wild mammals. Proceedings of the National Academy of Sciences of the United States of America 117:8546–8553. 10.1073/pnas.1911999117

Lensink CJ (1961) Status report: beluga studies. Division of Biological Research, Alaska Department of Fish and Game, Juneau, Alaska

Lewison RL, Crowder LB, Read AJ, Freeman SA (2004) Understanding impacts of fisheries bycatch on marine megafauna. Trends in Ecology & Evolution 19:598–604. 10.1016/j.tree.2004.09.004

Lima JY, Carvalho APM, Azevedo CT, et al (2016) Variation of age and total length in Sotalia guianensis (Van Bénéden, 1864) (Cetacea, Delphinidae), on the coast of Espírito Santo state, Brazil. Brazilian Journal of Biology 77:437–443. 10.1590/1519-6984.13215

Lockyer C, Heide-Jørgensen MP, Jensen J, et al (2001) Age, length and reproductive parameters of harbour porpoises Phocoena phocoena (L.) from West greenland. ICES Journal of Marine Science 58:154–162. 10.1006/jmsc.2000.0998

Marsili L, Casini C, Marini L, et al (1997) Age, growth and organochlorines (HCB, DDTs and PCBs) in Mediterranean striped dolphins Stenella coeruleoalba stranded in 1988-1994 on the coasts of Italy. Marine Ecology Progress Series 151:273–282. 10.3354/meps151273

Martin AR, Reynolds P, Richardson MG (1987) Aspects of the biology of Pilot whales (Globicephala melaena) in recent mass strandings on the British coast. Journal of Zoology 211:11–23. 10.1111/j.1469-7998.1987.tb07449.x

Mattson MC, Mullin KD, Ingram GW, Hoggard W (2006) Age structure and growth of the bottlenose dolphin (Tursiops truncatus) from strandings in the Mississippi sound region of the north-central Gulf of Mexico from 1986 to 2003. Marine Mammal Science 22:654–666. 10.1111/j.1748-7692.2006.00057.x

McElreath R (2020) rethinking: Statistical Rethinking book package

McGowen MR, Tsagkogeorga G, Álvarez-Carretero S, et al (2020) Phylogenomic Resolution of the Cetacean Tree of Life Using Target Sequence Capture. Systematic Biology 69:479–501. 10.1093/sysbio/syz068

Mei Z, Huang SL, Hao Y, et al (2012) Accelerating population decline of Yangtze finless porpoise (Neophocaena asiaeorientalis asiaeorientalis). Biological Conservation 153:192–200. 10.1016/j.biocon.2012.04.029

Mitchell E, Kozicki M (1984) Reproductive condition of male sperm whales, Physeter macrocephalus, taken off Nova Scotia. Reports of the International Whaling Commission 243–252

Miyazaki N (1978) Preliminary note on age determination and growth of the rough-toothed dolphin, Steno bredanensis, off the Pacific coast of Japan. Report of the International Whaling Commission SI3:171–180

Miyazaki N (1984) Further analyses of reproduction in the striped dolphin, Stenella coeruleoalba, off the Pacific Coast of Japan. Reports of the International Whaling Commission SI6:343–353

Miyazaki N, Fujise Y, Iwata K (1998) Biological analysis of a mass stranding of Melon-headed whales (Peponocephala electra) at Aoshima, Japan. Bulletin of the National Science Museum Tokyo, Japan Series A 24:31–60

Modrák M, Moon AH, Kim S, et al (2023) Simulation-Based Calibration Checking for Bayesian Computation: The Choice of Test Quantities Shapes Sensitivity. Bayesian Anal 1: 10.1214/23-BA1404

Molina DM, Reyes JC (1996) Determinación de edad en el delfín chileno Cephalorhynchus eutropia (Cetacea: Delphinidae). Revista Chilena de Historia Natural 69:183–191

Moore JE, Read AJ (2008) A Bayesian Uncertainty Analysis of Cetacean Demography and Bycatch Mortality Using Age-at-Death Data. Ecological Applications 18:1914–1931. 10.1890/07-0862.1

Murphy S, Petitguyot MAC, Jepson PD, et al (2020) Spatio-Temporal Variability of Harbor Porpoise Life History Parameters in the North-East Atlantic. Frontiers in Marine Science 7:. 10.3389/fmars.2020.502352

Murphy S, Winship A, Dabin W, et al (2009) Importance of biological parameters in assessing the status of Delphinus delphis. Marine Ecology Progress Series 388:273–291. 10.3354/meps08129

National Research Council (ed) (2005) Marine mammal populations and ocean noise: determining when noise causes biologically significant effects. National Academies Press, Washington, D.C

Nattrass S, Croft DP, Ellis S, et al (2019) Postreproductive killer whale grandmothers improve the survival of their grandoffspring. Proceedings of the National Academy of Sciences. 10.1073/pnas.1903844116

Negri MF, Panebianco MV, Denuncio P, et al (2016) Biological parameters of franciscana dolphins, Pontoporia blainvillei, by-caught in artisanal fisheries off southern Buenos Aires, Argentina. Journal of the Marine Biological Association of the United Kingdom 96:821–829. 10.1017/S0025315414000393

Neuenhoff RD (2013) Age, growth, and population dynamics of common Bottlenose dolphin (Tursiops truncatus) along coastal Texas. Master of Science Thesis, Texas A&M University

Newby TC (1982) Life history of Dall’s porpoise (Phocoenoides dalli, True 1885) incidentally taken by the Japanese high seas salmon mothership fishery in the northwestern North Pacific and western Bering sea, 1978 and 1980. Univeristy of Washington

Nielsen MLK, Ellis S, Towers JR, et al (2021a) A long postreproductive life span is a shared trait among genetically distinct killer whale populations. Ecology and Evolution 11:9123–9136. 10.1002/ece3.7756

Nielsen MLK, Ellis S, Towers JR, et al (2021b) A long postreproductive life span is a shared trait among genetically distinct killer whale populations. Ecology and Evolution 11:9123–9136. 10.1002/ece3.7756

Nolte Z (2014) The natural history of the humpback dolphin, Sousa chinensis, in KwaZulu-Natal, South Africa: age, growth and reproduction. Rhodes University

Nuñez CMV, Coulson T, Festa-Bianchet M, Gaillard J-M (2008) Measuring senescence in wild animal populations: Towards a longitudinal approach. Functional Ecology 22:393–406. 10.1111/j.1365-2435.2008.01408.x

Ohsumi S (1966) Sexual segregation of the sperm whale in the North Pacific. Sci Rep Whales Res Inst 20:1–16

Péron G, Bonenfant C, Lemaitre JF, et al (2019) Does grandparental care select for a longer lifespan in non-human mammals? Biological Journal of the Linnean Society 128:360–372. 10.1093/biolinnean/blz078

Perrin WF, Henderson JR (1984) Growth and reproductive rates in two populations of spinner dolphins, Stenella longirostris, with different histories of exploitation. Report of the International Whaling Commission SI6:417–430

Perrin WF, Holts DB, Miller RB (1977) Growth and reproduction of the eastern spinner dolphin, a geographical form of Stenella longirostris in the eastern tropical Pacific. Fish Bull US 75:725–750

Perrin WF, Myrick AC (1980) Age determination of toothed whales and sirenians: report of the workshop

Plön S (2004) The status and natural history of pygmy (Kogia breviceps) and dwarf (K. sima) sperm whales off Southern Africa. PhD Thesis

Plön S, Heyns-Veale ER, Smale MJ, Froneman PW (2020) Life history parameters and diet of Risso’s dolphins, Grampus griseus, from southeastern South Africa. Marine Mammal Science 36:786–801. 10.1111/mms.12675

R Development Core Team (2021) R: A language and environment for statistical computing

Read AJ, Tolley KA (1997) Postnatal growth and allometry of harbour porpoises from the Bay of Fundy. Canadian Journal of Zoology 75:122–130. 10.1139/z97-016

Read AJ, Wells RS, Hohn AA, Scott MD (1993) Patterns of growth in wild bottlenose dolphins, Tursiops truncatus. Journal of Zoology 231:107–123

Read FL, Hohn AA, Lockyer CH (2018) A review of age estimation methods in marine mammals with special reference to monodontids. NAMMCO Scientific Publications 8:1–67. 10.7557/3.4474

Reijnders PJH, Borrell A, van Franeker JA, Aguilar A (2018) Pollution. In: Würsig B, Thewissen JGM, Kovacs KM (eds) Encyclopedia of Marine Mammals, 3rd edn. Academic Press, London, pp 746–753

Rice DW, Wolman AA, Mate BR, Harvey JT (1986) A mass stranding of sperm whales in Oregon: sex and age composition of the school. Marine Mammal Science 2:64–69. 10.1111/j.1748-7692.1986.tb00027.x

Ridgway SH, Harrison RJ (eds) (1989) Handbook of marine mammals. Vol. 4: River dolphins and the larger toothed whales. Academic Press, London

Ridgway SH, Harrison RJ (eds) (1994) Handbook of marine mammals. Vol. 5: The first book of dolphins. Academic Press, London

Ridgway SH, Harrison RJ (eds) (1999) Handbook of marine mammals. Vol. 6: The second book of dolphins and the porpoises. Academic Press, London

Rogan E, Baker JR, Jepson PD, et al (1997) A mass stranding of white-sided dolphins (Lagenorhynchus acutus) in Ireland: biological and pathological studies. Journal of Zoology 242:217–227

Rohatgi A (2020) WebPlotDigitizer

Ronget V, Gaillard J-M (2020) Assessing ageing patterns for comparative analyses of mortality curves: Going beyond the use of maximum longevity. Functional Ecology 34:65–75. 10.1111/1365-2435.13474

Rosas FCW, Barreto AS, Monteiro-Filho ELDA (2003) Age and growth of the estuarine dolphin (Sotalia guianensis) (Cetacea, Delphinidae) on the sParaná coast southern Brazil. Fishery Bulletin 101:377–383

Rouby E, Ridoux V, Authier M (2021) Flexible parametric modeling of survival from age at death data: A mixed linear regression framework. Population Ecology 63:108–122. 10.1002/1438-390X.12069

Saavedra C (2018) strandCet: R package for estimating natural and non-natural mortality-at-age of cetaceans from age-structured strandings. PeerJ 6:e5768. 10.7717/peerj.5768

Salguero-Gómez R, Jones OR, Archer CR, et al (2016) COMADRE: a global data base of animal demography. Journal of Animal Ecology 85:371–384. 10.1111/1365-2656.12482

Santos MC de. O, Rosso S, Ramos RMA (2003) Age estimation of marine tucuxi dolphins (Sotalia fluviatilis) in south-eastern Brazil. Journal of the Marine Biological Association of the United Kingdom 83:233–236. 10.1017/s0025315403007021h

Sergeant DE, St.Aubin DJ, Geraci JR (1980) Life history and Northwest Atlantic status of the Atlantic white-sided dolphin, Lagenorhynchus acutus. Cetology 37:1–12

Shefferson RP, Jones OR, Salguero-Gómez R (eds) (2017) The Evolution of Senescence in the Tree of Life. Cambridge University Press, Cambridge

Shirakihara M, Takemura A, Shirakihara K (1993) Age, growth, and reproduction of the finless porpoise, Neophocaena phocaenoides, in the coastal waters of western Kyushu, Japan. Marine Mammal Science 9:392–406

Siciliano S, Ramos RMA, Di Beneditto APM, et al (2007) Age and growth of some delphinids in south-eastern Brazil. Journal of the Marine Biological Association of the United Kingdom 87:293–303. 10.1017/S0025315407053398

Silk MJ, Hodgson DJ (2021) Differentiated Social Relationships and the Pace-of-Life-History. Trends in Ecology & Evolution 36:498–506. 10.1016/j.tree.2021.02.007

Slooten E (1991) Age, growth, and reproduction in Hector’s dolphins. Canadian Journal of Zoology 69:1689–1700. 10.1139/z91-234

Stan Development Team (2020) Rstan: the R interface to stan

Stan Development Team (2021) CmdStanR: the R interface to CmdStan

Stewart REA, Campana SE, Jones CM, Stewart BE (2006) Bomb radiocarbon dating calibrates beluga (Delphinapterus leucas) age estimates. Canadian Journal of Zoology 1852:1840–1852. 10.1139/Z06-182

Stolen MK, Barlow J (2003) A model life table for bottlenose dolphins (Tursiops truncatus) from the Indian River Lagoon system, Florida, U.S.A. Marine Mammal Science 19:630–649. 10.1111/j.1748-7692.2003.tb01121.x

Suydam RS (2009) Age, growth, reproduction, and movements of beluga whales (Delphinapterus leucas) from the eastern Chukchi Sea, Robert Scott Suydam. PhD Thesis, University of Washington

Toïgo C, Gaillard JM, Festa-Bianchet M, et al (2007) Sex- and age-specific survival of the highly dimorphic Alpine ibex: Evidence for a conservative life-history tactic. Journal of Animal Ecology 76:679–686. 10.1111/j.1365-2656.2007.01254.x

Tombak KJ, Hex SB, Rubenstein DI (2024) New estimates indicate that males are not larger than females in most mammal species. Nature Communications 15:1872

Tuerk KJS, Kucklick JR, McFee WE, et al (2005) Factors influencing persistent organic pollutant concentrations in the Atlantic white-sided dolphin (Lagenorhynchus acutus). Environmental Toxicology and Chemistry 24:1079–1087

Urbán JR, Cárdenas-Hinojosa G, Gómez-Gallardo U. A, et al (2007) Mass stranding of Baird’s beaked whales at San Jose Island, Gulf of California, Mexico. Latin American Journal of Aquatic Mammals 6:83–88. 10.5597/lajam00111

Van Bree PJH, Collet A, Desportes G, et al (1986) Le dauphin de Fraser, Lagenodephis hosei (Cetecea, Odontoceti), espèce nouvelle pour la faune d’Europe. Mammalia 50:57–86

Van Utrecht WL (1981) Comparison of accumulation patterns in leyered dentinal tissue od some odontocety and corresponding patterns in baleen plates and ear plugs of Balaenopteridae. Beaufortia 31:111–122

van Weelden C, Towers JR, Bosker T (2021) Impacts of climate change on cetacean distribution, habitat and migration. Climate Change Ecology 1:100009. 10.1016/j.ecochg.2021.100009

Vaupel JW (2003) Post-Darwinian Longevity. Population and Development Review 29:258–269

Venuto R, Botta S, Barreto AS, et al (2020) Age structure of strandings and growth of Lahille’s bottlenose dolphin (Tursiops truncatus gephyreus). Marine Mammal Science 36:813–827. 10.1111/mms.12683

Viricel A, Strand AE, Rosel PE, et al (2008) Insights on common dolphin (Delphinus delphis) social organization from genetic analysis of a mass-stranded pod. Behavioral Ecology and Sociobiology 63:173–185. 10.1007/s00265-008-0648-7

Vos DJ, Shelden KEW, Friday NA, Mahoney BA (2020) Age and growth analyses for the endangered belugas in Cook Inlet, Alaska. Marine Mammal Science 36:293–304. 10.1111/mms.12630

Wade PR (2018) Population dynamics. In: Würsug B, Thewissen JGM, Kovacs KM (eds) Encyclopedia of Marine Mammals, 3rd edn. Academic Press, London, pp 763–770

Walker WA, Goodrich KR, Leatherwood S, Stroud RK (1984) Population biology and ecology of the Pacific white-sided dolphin, Lagenorhynchus obliquidens, in the northeastern Pacific. Part II: Biology and geographical variation.

Watt CA, Stewart BE, Loseto L, et al (2020) Estimating narwhal (Monodon monoceros) age using tooth layers and aspartic acid racemization of eye lens nuclei. Marine Mammal Science 36:103–115. 10.1111/mms.12623

Weiss MN, Ellis S, Croft DP (2021) Diversity and consequences of social network structure in toothed whales. Frontiers in Marine Science 1:1

Whitehead H, Mann J (2000) Female reproductive strategies of ceteceans: life histories and calf care. In: Mann J, Connor RC, Tyack PL, Whitehead H (eds) Cetacean Societies: Field Studies of Whales and Dolphins. The University of Chicago Press, Chicago and London, pp 219–246

Würsig B, Thewissen JGM, Kovacs KM (eds) (2018) Encyclopedia of Marine Mammals, 3rd edn. Academic Press, London

