## Supplementary 8 for "Bayesian inference of toothed whale lifespans"

Credible interval widths from simulations testing the accuracy of the mortality model to detect the true maximum lifespan of a population under a range of A) sample sizes (n) x population growth rates (r) and B) sample sizes x true maximum lifespans. The colour and value in each cell represents the mean 95% credible interval width scaled to true maximum lifespan of simulations in the category. Balues show the mean ± std. dev. Relative credible interval width, and ligther colours reflect smaller values i.e. narrow relative 95% credible intervales. 100 models were run for each population growth x sample size scenario, resulting in a total of 4900 fitted models. B) is based on the same 4900 fitted models, true maximum lifespans and has a uniform probability of being selected but the number of models applied in each cell will vary. Note that neither sample size nor population growth rate are a regular sequence. All details of the simulations and their interpretation are given in the text and figure 2.

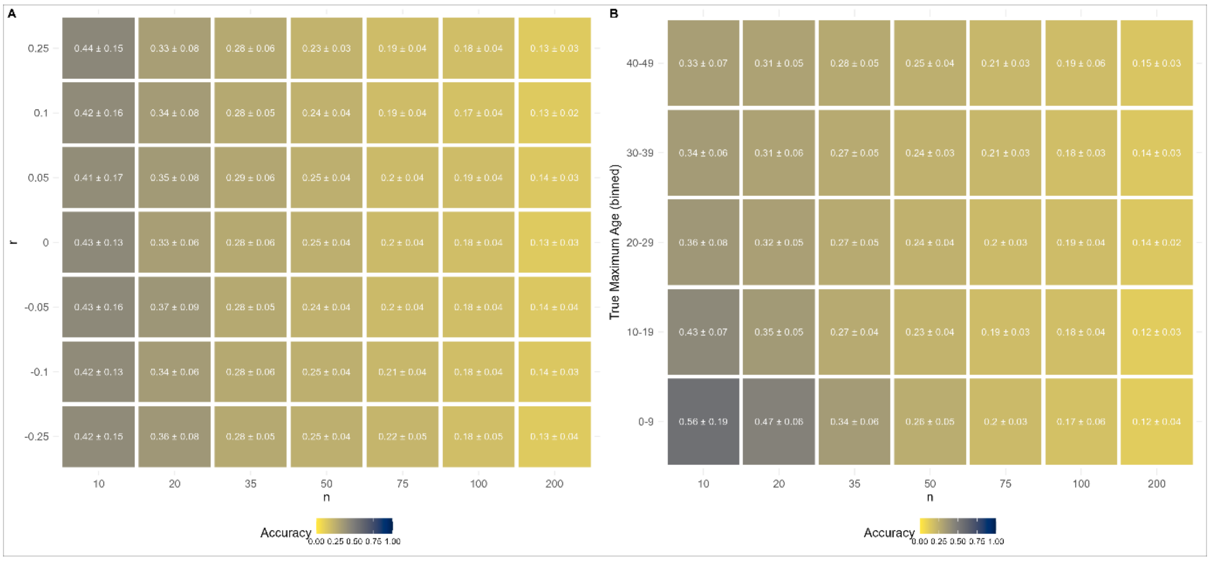
