## Supplementary 1 for "Bayesian inference of toothed whale lifespans"

The following pages illustrate the described modelling pathway applied to all toothed whale datasets in our database.

For each species there are a series of pages demonstrating the model applied to the valuable datasets:

(1) Dataset plots with derived model predictions applied. In all plots, x axis is age, y axis is count of whales of a given age. Grey bars show the dataset samples. Red line and ribbon shows the number of whales expected to be in the sample from the model. Red line shows the posterior mean and ribbon the 95% credible interval. Where multiple datasets are available for a given species they are shown as separate panels. As described in the text model mortality parameters are shared by all species-sex datasets but other parameters can vary between datasets and populations.
(2) As above but for males. Note that in some species data are only available for one sex and pages (1) or (2) will not be present.
(3) 100 posterior sample curves derived from the model when applied. Each curve shows the curve derived from a single posterior draw of the mortality parameters. Where both sexes are present the upper plot is female, and lower plot male.
(4) Mean and 95% credible interval of ordinary maximum lifespan for each species-sex serviced from the mortality parameters derived from the model.

All data are available in the R package marinelifehistdata, and the modelling pathway in the R package marinesurvival.

Note that this document shows the model applied to all datasets and species where data are identified. However, the final analysis filters down to a smaller proportion of these species with larger datasets and greater sampling intensity (see main text *calculating lifespan)*.

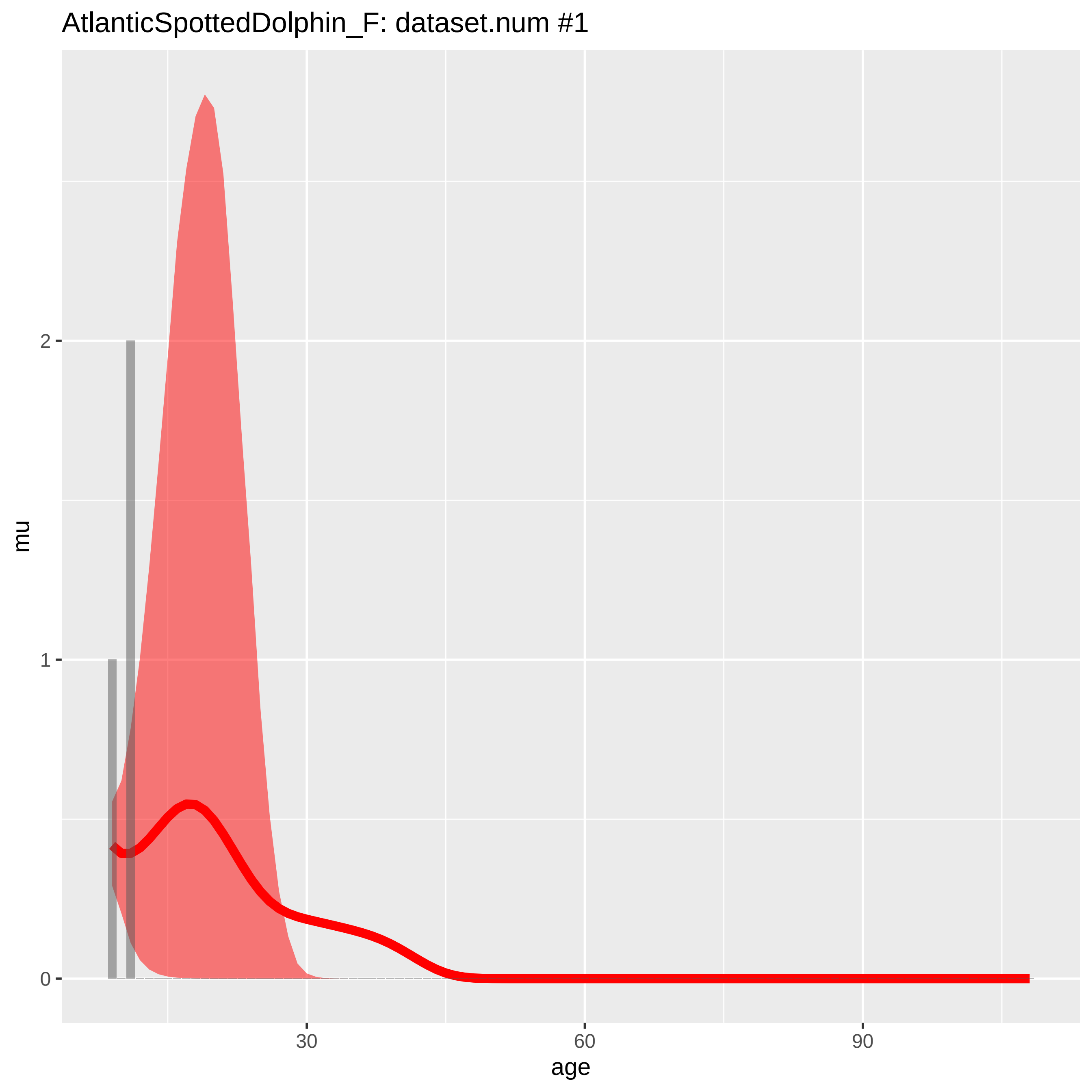
AtlanticSpottedDolphin_F: dataset.num #1
mu
0
1
2
age
30
60
90

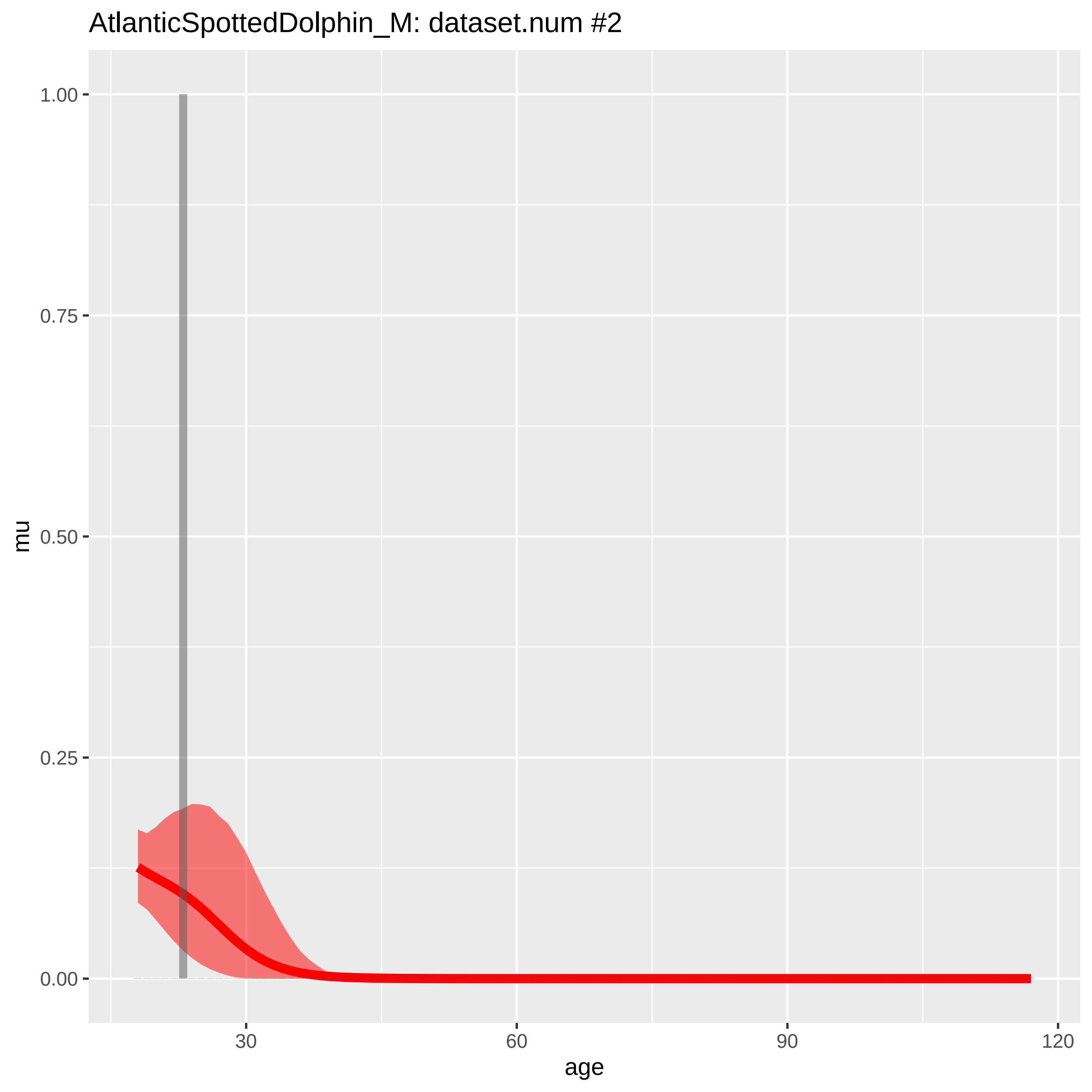
AtlanticSpottedDolphin_M: dataset.num #2
1.00
0.75
0.50
0.25
0.00
mu
30
60
90
120
age

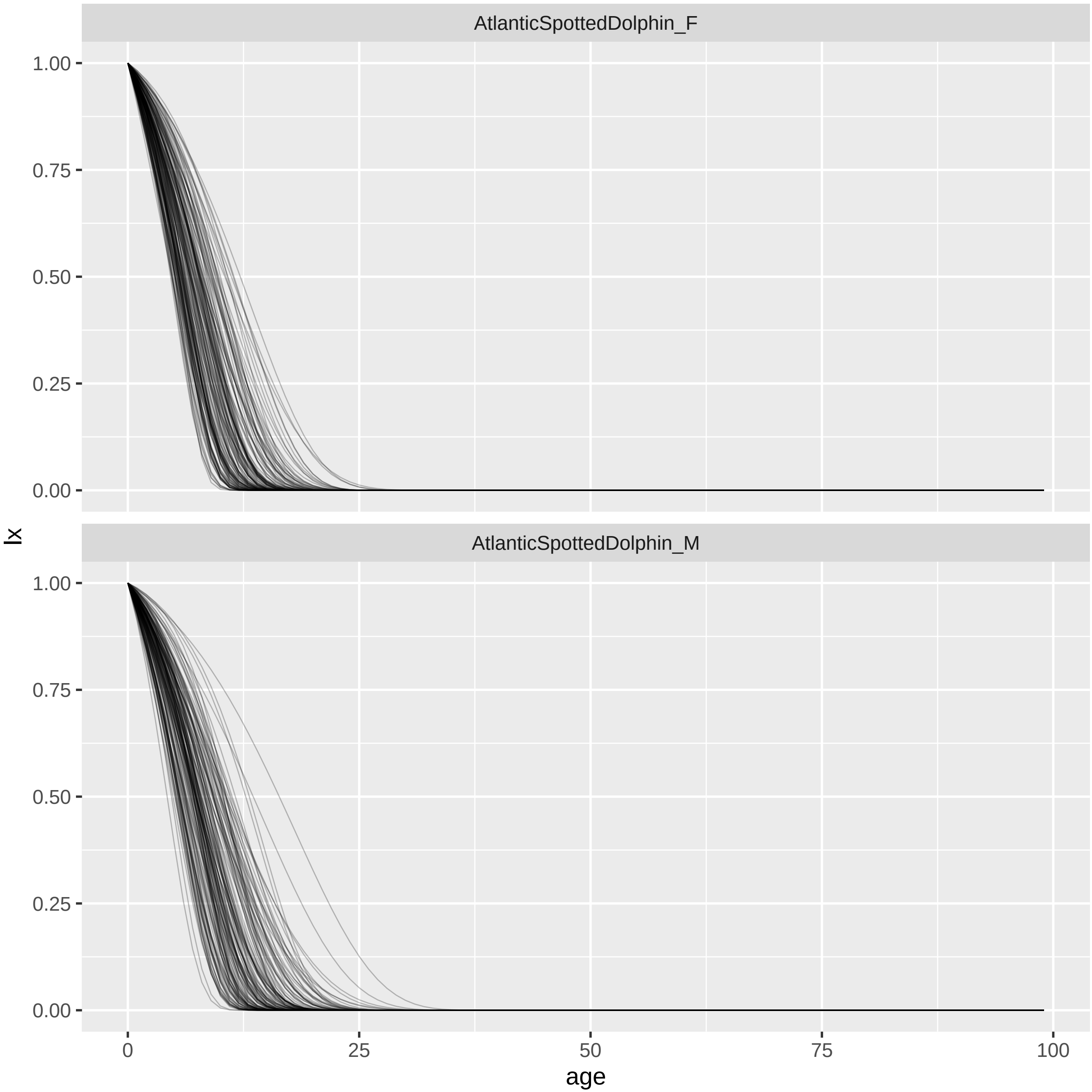
AtlanticSpottedDolphin_F
AtlanticSpottedDolphin_M
lx
1.00
0.75
0.50
0.25
0.00
0
25
50
75
100
age

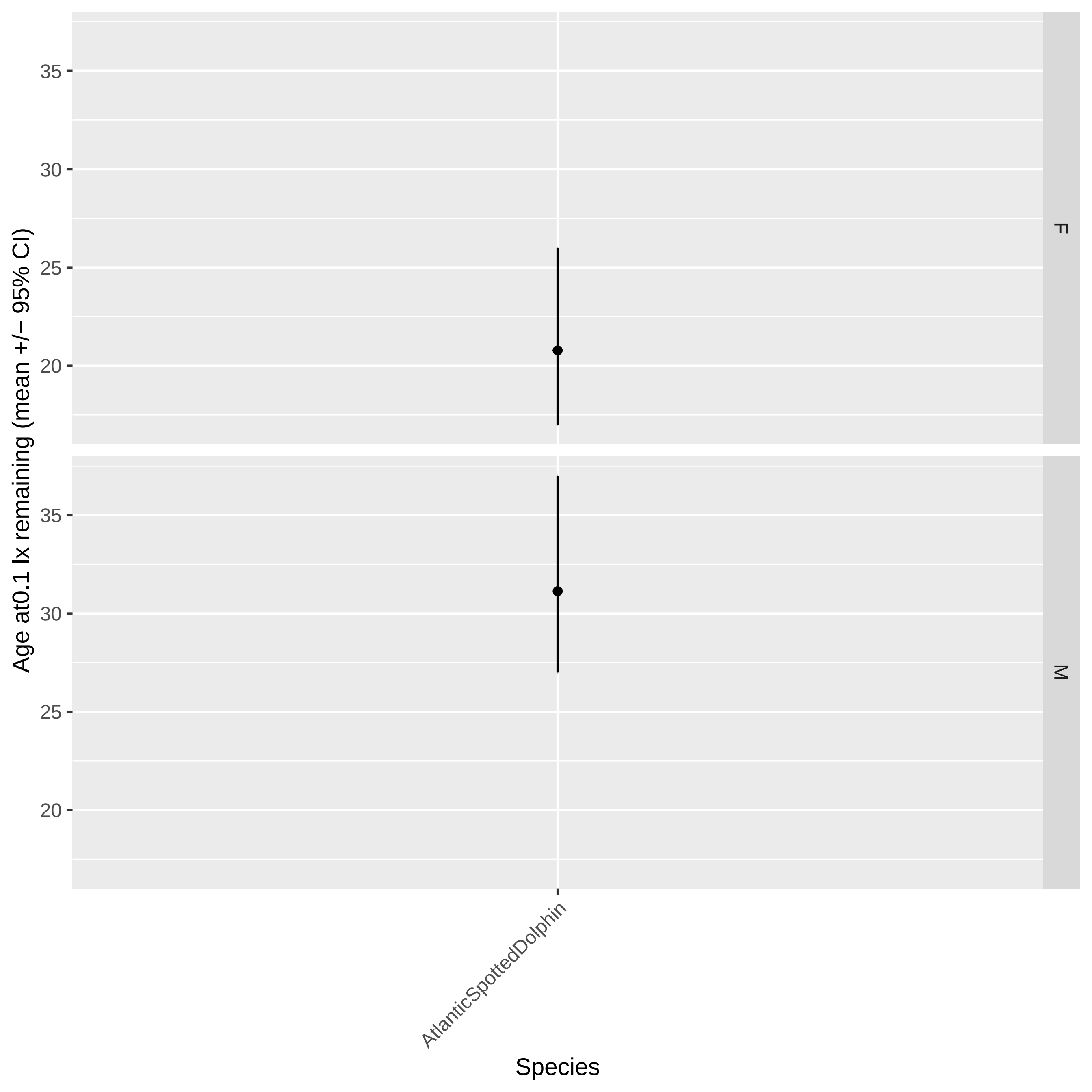
Age at0.1 lx remaining (mean +/− 95% CI)
35
30
25
20
F
M
AtlanticSpottedDolphin
Species

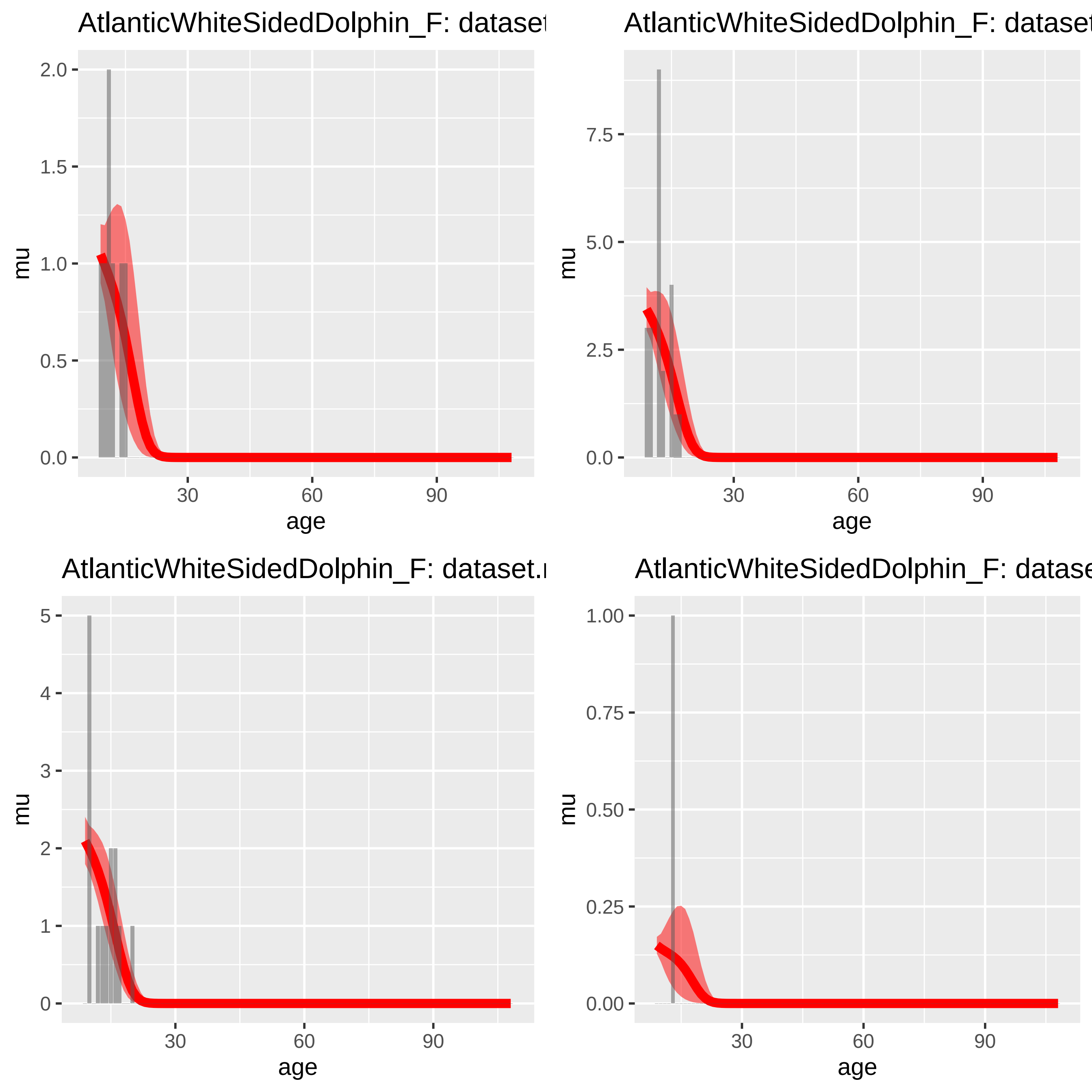

AtlanticWhiteSidedDolphin_F: dataset
AtlanticWhiteSidedDolphin_F: dataset
AtlanticWhiteSidedDolphin_F: dataset.
AtlanticWhiteSidedDolphin_F: datase
mu
age
0.0
0.5
1.0
1.5
2.0
2.5
5.0
7.5
0
1
2
3
4
5
0.00
0.25
0.50
0.75
1.00
30
60
90

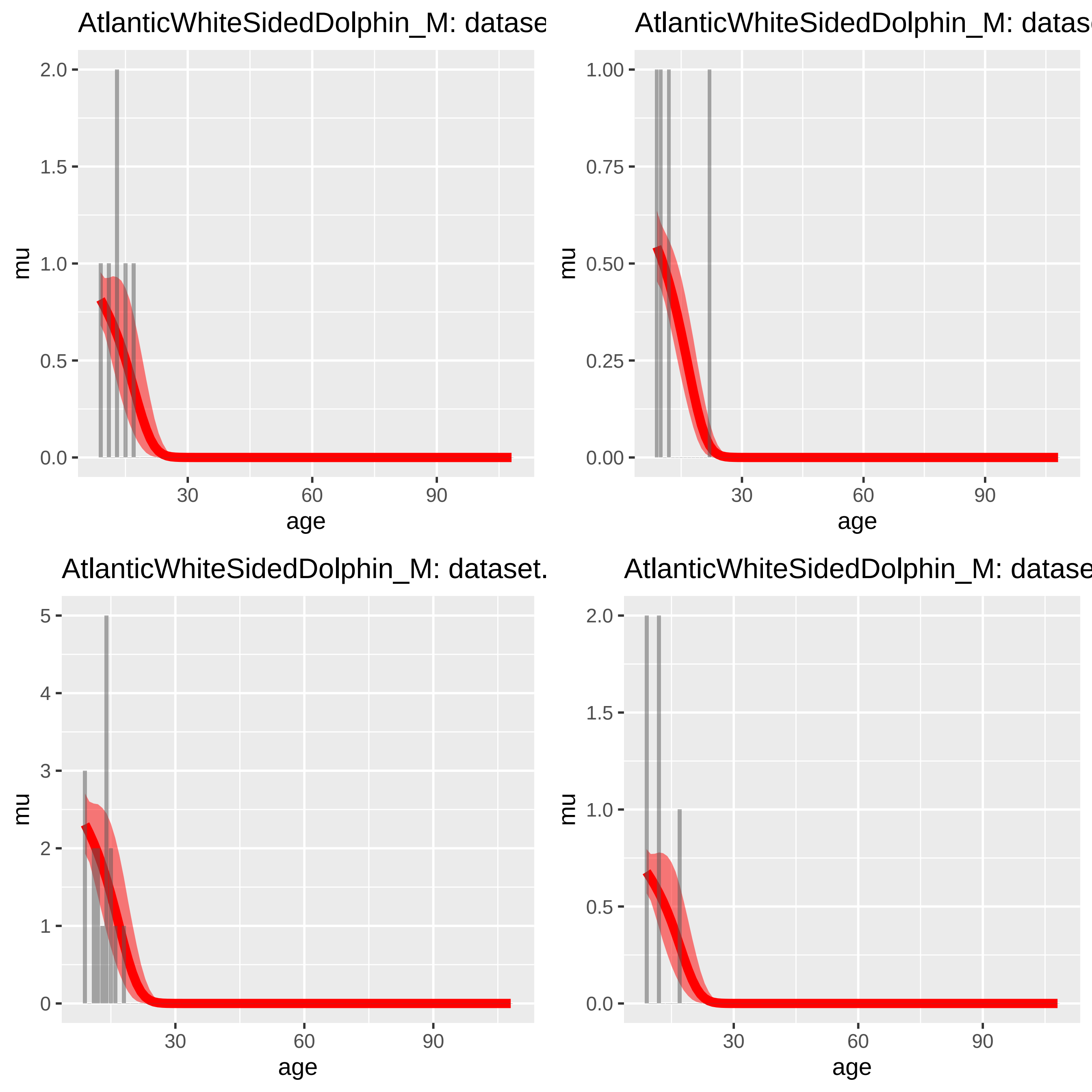
AtlanticWhiteSidedDolphin_M: datase
AtlanticWhiteSidedDolphin_M: datas
AtlanticWhiteSidedDolphin_M: dataset.
AtlanticWhiteSidedDolphin_M: datase
mu
age
0.0
0.5
1.0
1.5
2.0
0.00
0.25
0.50
0.75
1.00
0
1
2
3
4
5
30
60
90

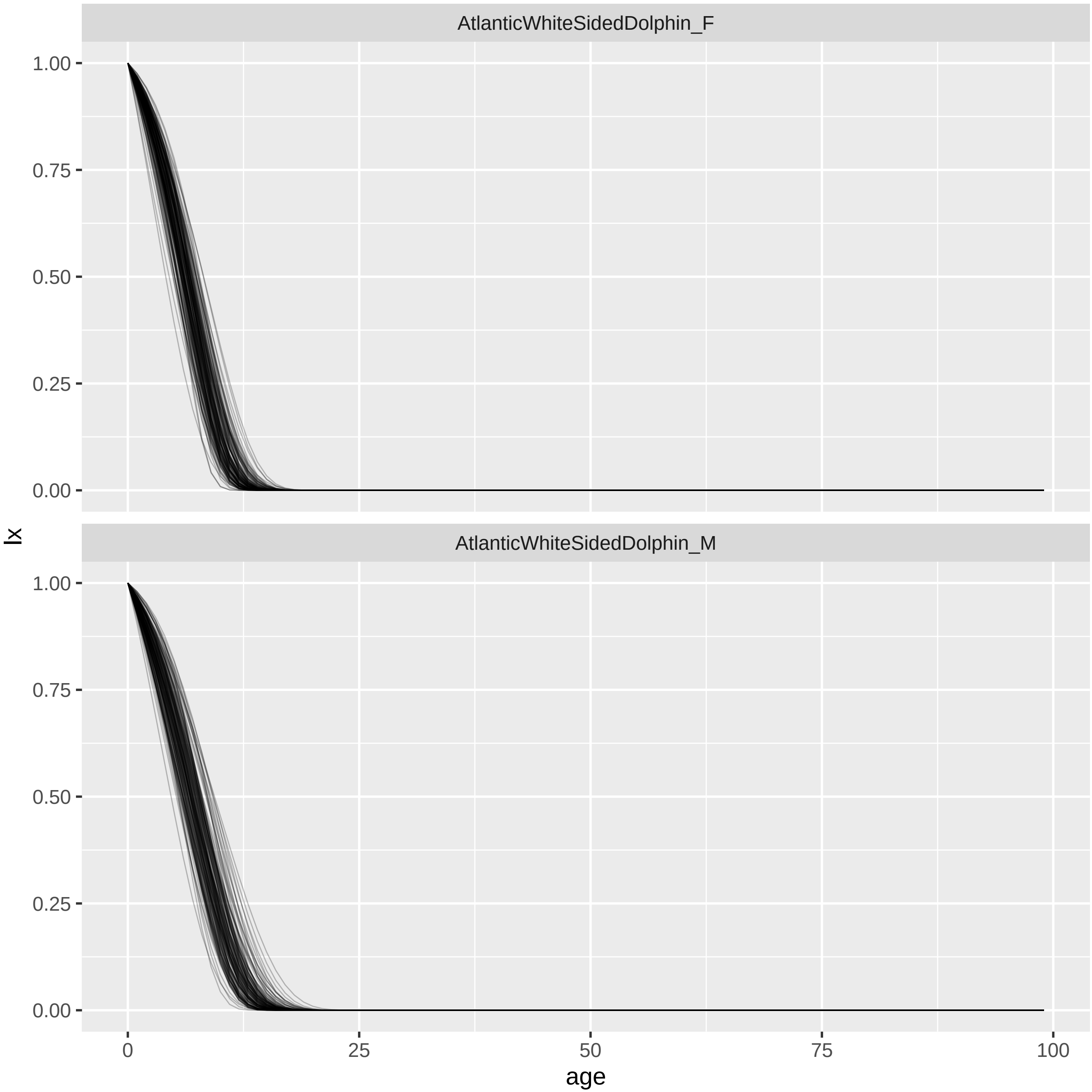
AtlanticWhiteSidedDolphin_F
AtlanticWhiteSidedDolphin_M
1.00
0.75
0.50
0.25
0.00
lx
0
25
50
75
100
age

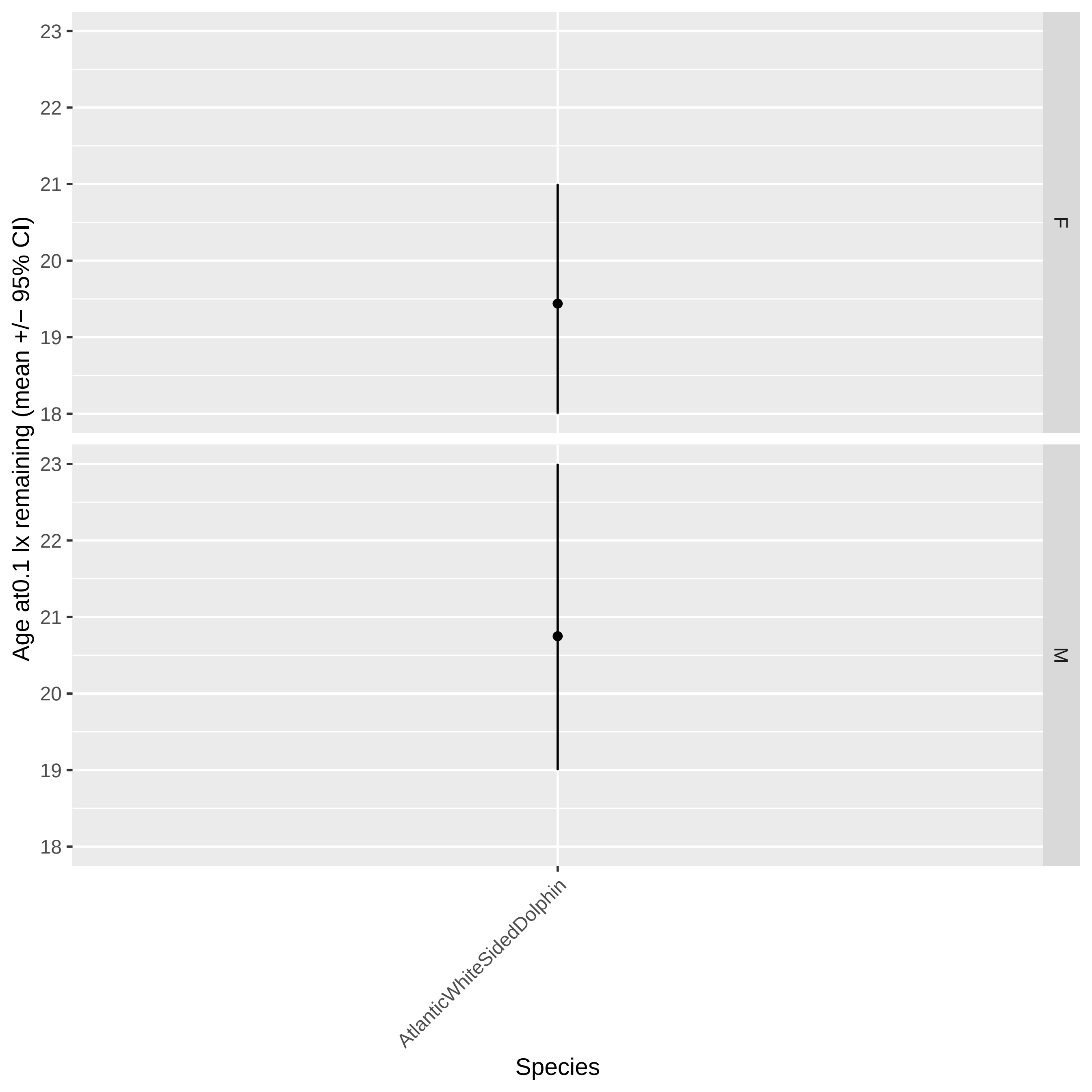

Age at0.1 lx remaining (mean +/− 95% CI)
23
22
21
20
19
18
F
M
AtlanticWhiteSidedDolphin
Species

Baiji_F: dataset.num #2

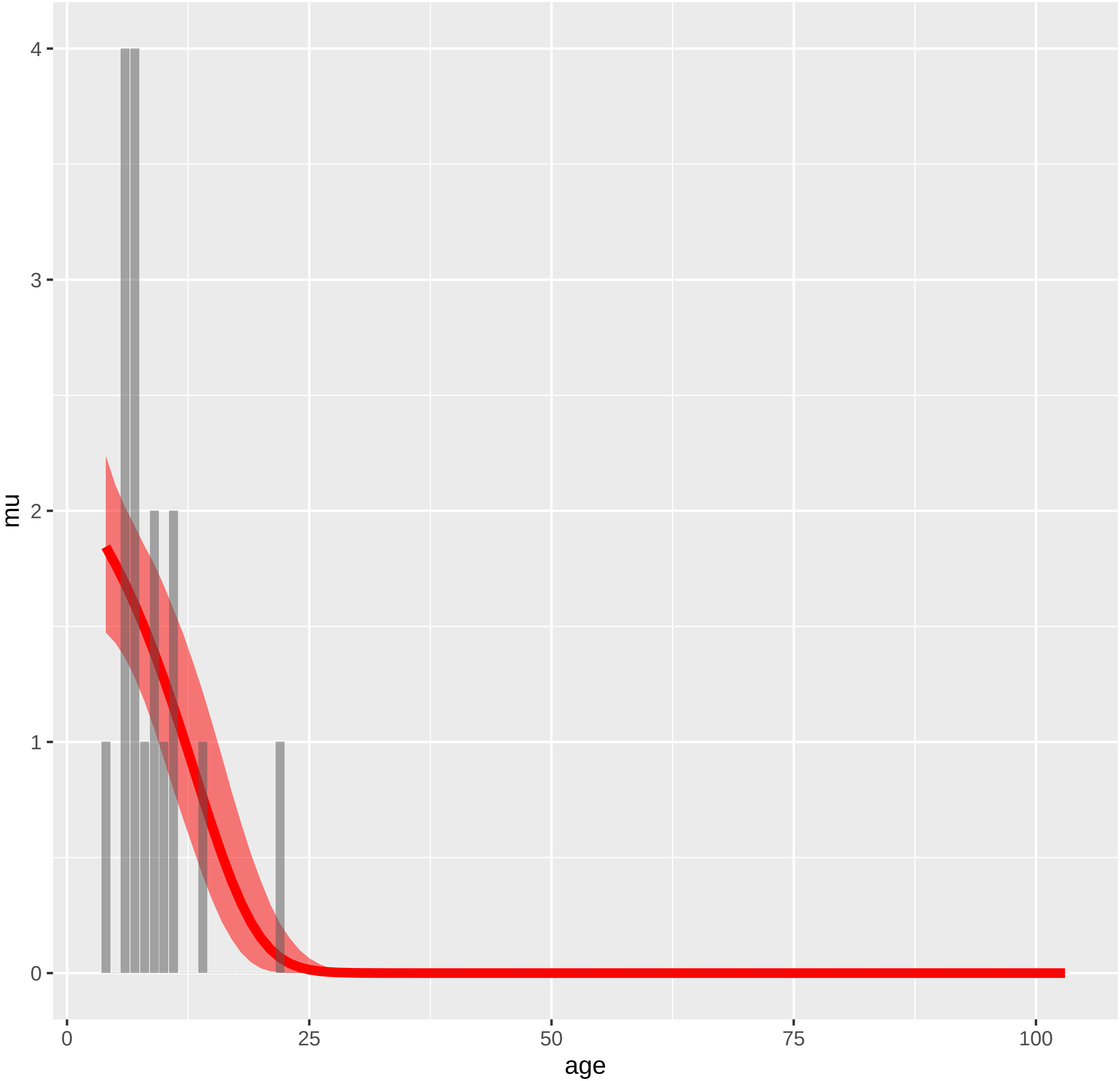

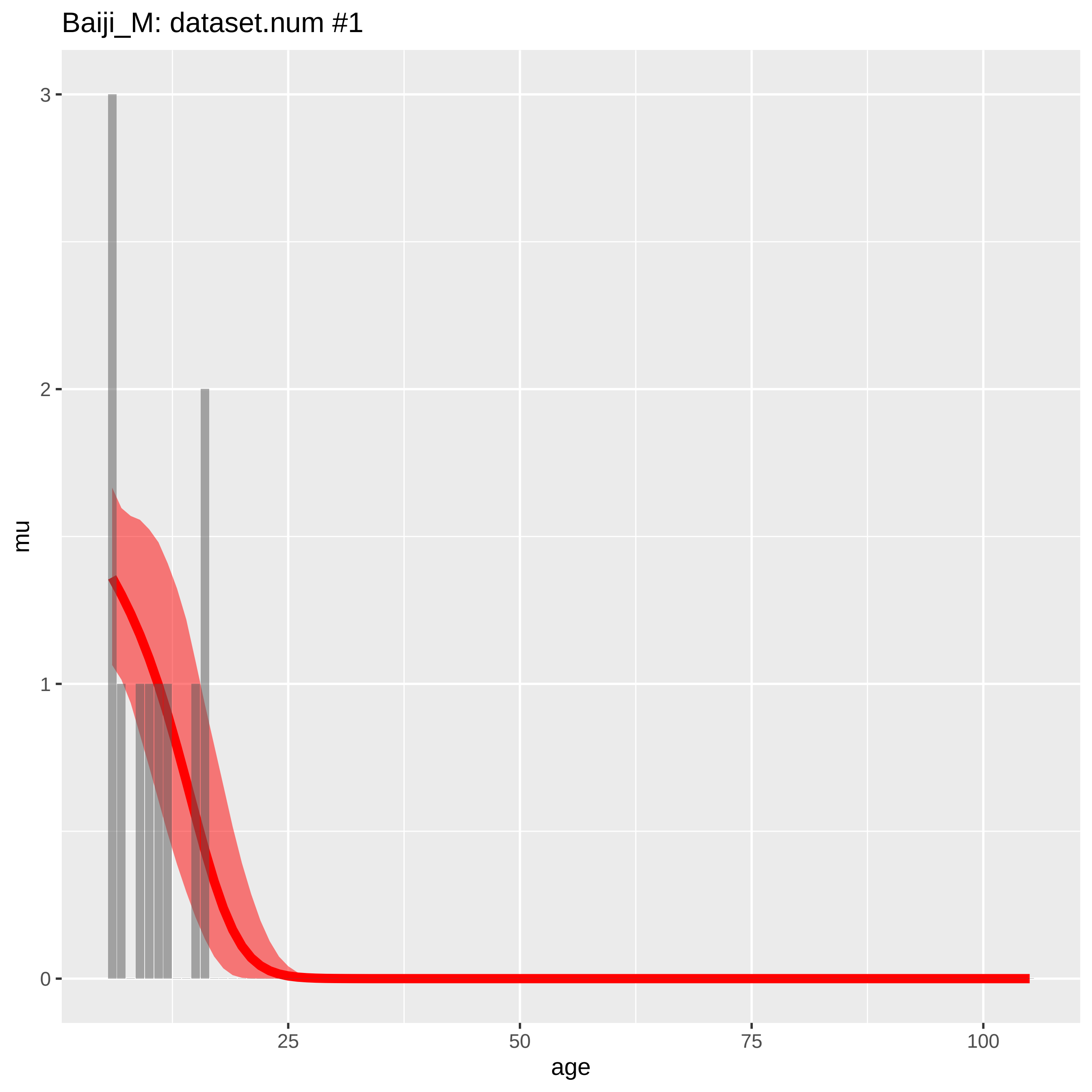

Baiji_M: dataset.num #1
mu
3
2
1
0
25
50
75
100
age

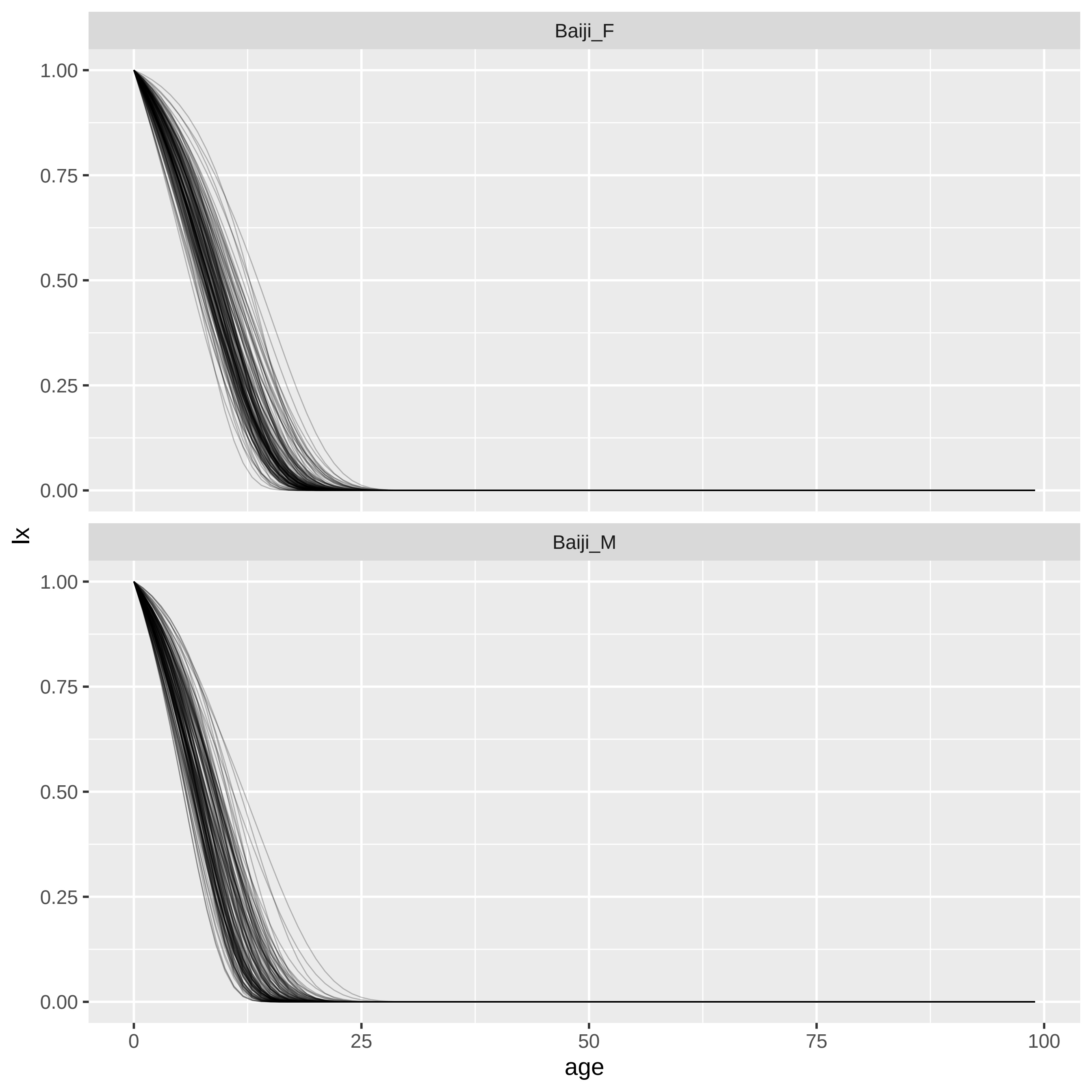
Baiji_F
Baiji_M
lx
1.00
0.75
0.50
0.25
0.00
0
25
50
75
100
age

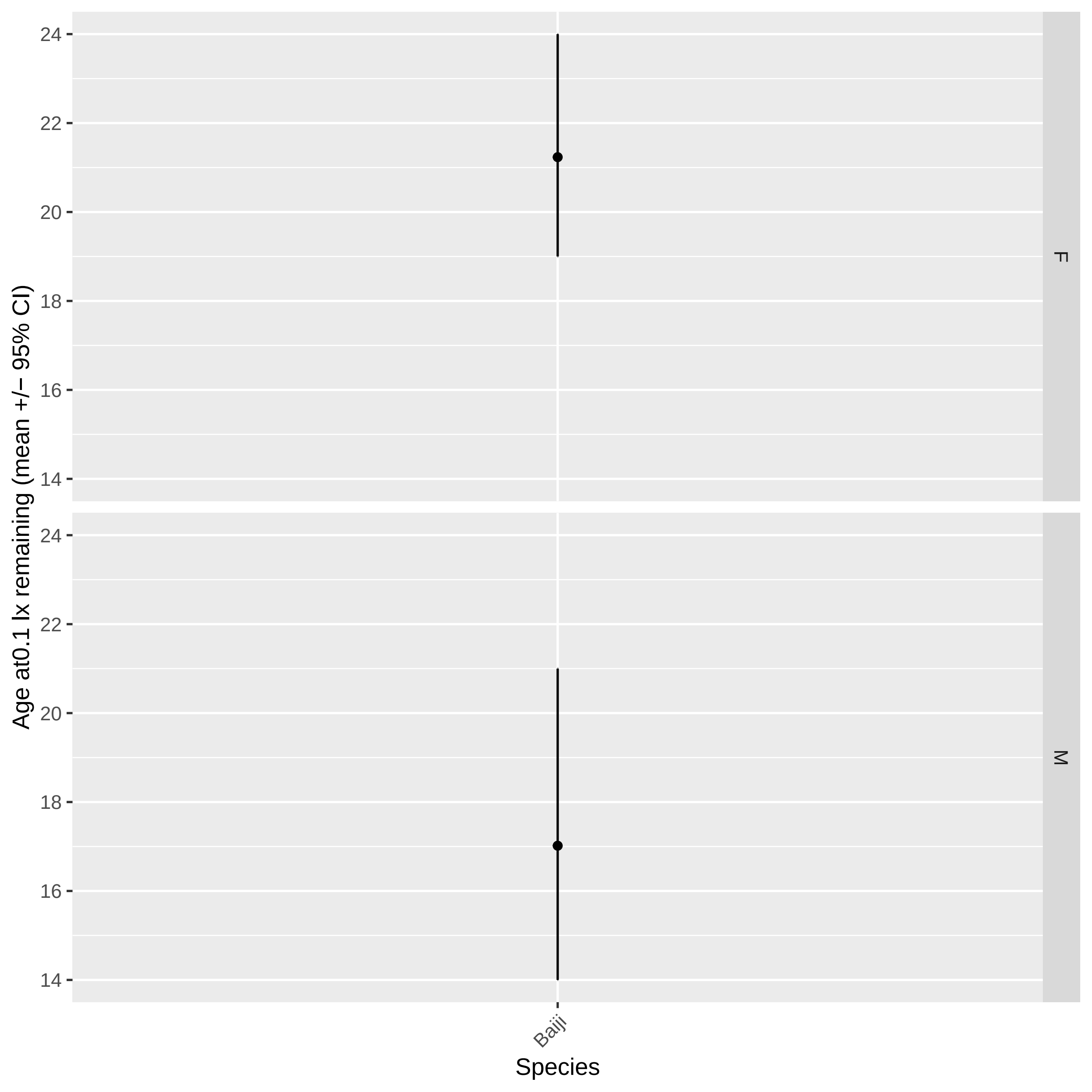
Age at0.1 lx remaining (mean +/− 95% CI)
24
22
20
18
16
14
F
M
Baiji
Species

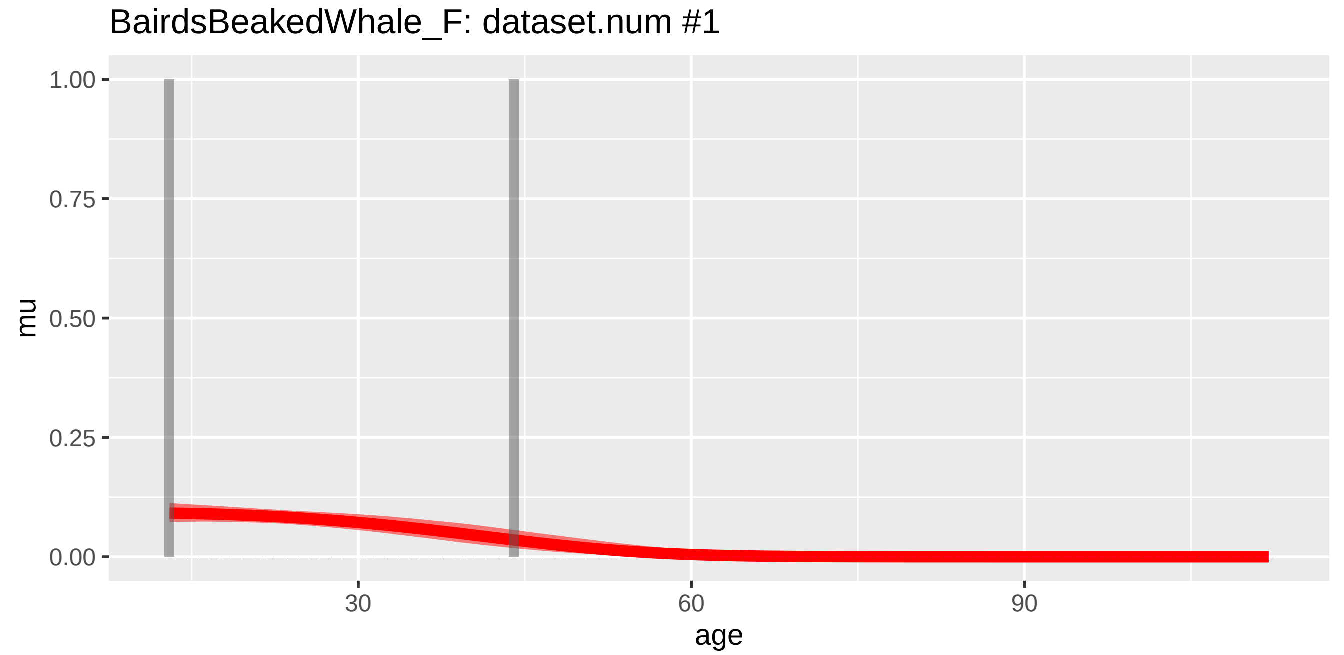
BairdsBeakedWhale_F: dataset.num #1
mu
1.00
0.75
0.50
0.25
0.00
30
60
90
age

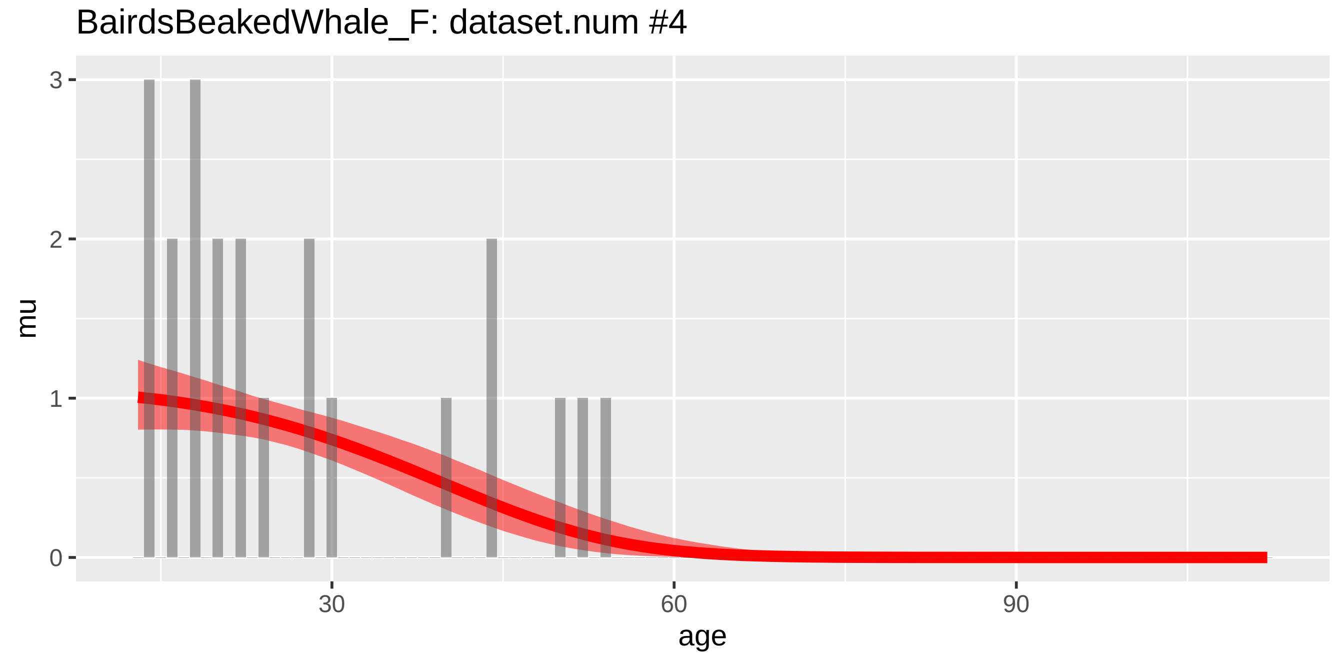
BairdsBeakedWhale_F: dataset.num #4
mu
3
2
1
0
30
60
90
age

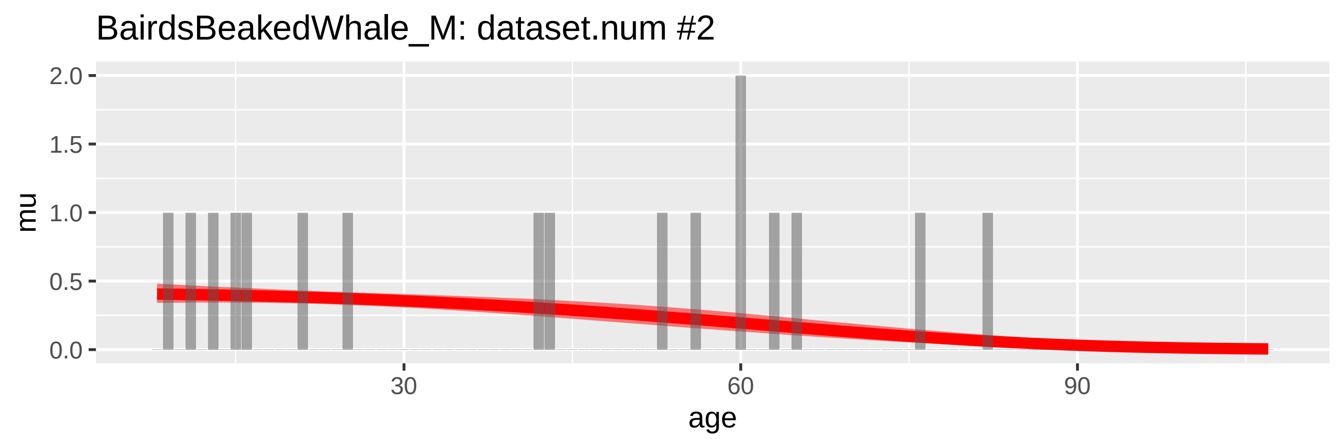
BairdsBeakedWhale_M: dataset.num #2
mu
age
0.0
0.5
1.0
1.5
2.0
30
60
90

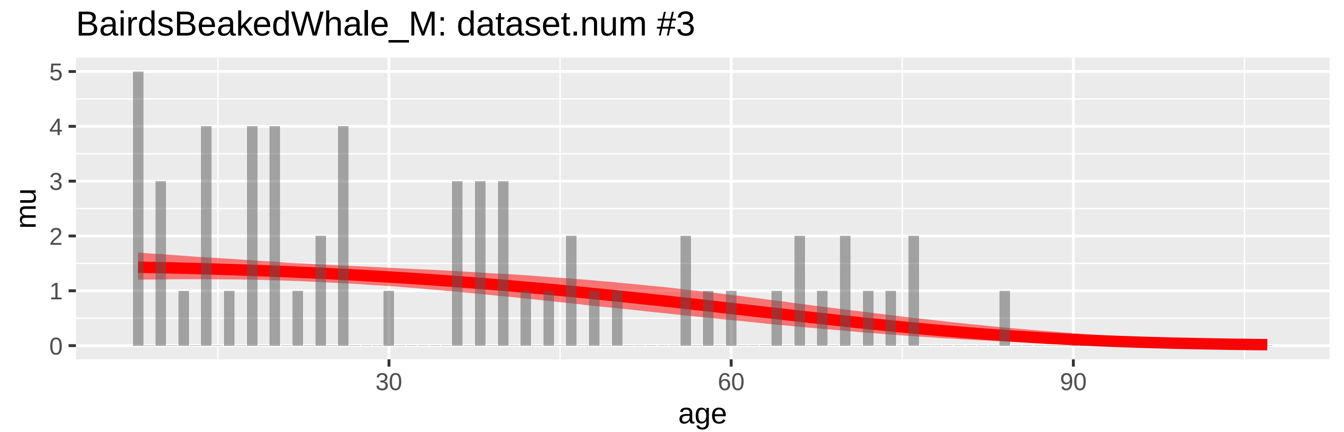
BairdsBeakedWhale_M: dataset.num #3
mu
age
0
1
2
3
4
5
30
60
90

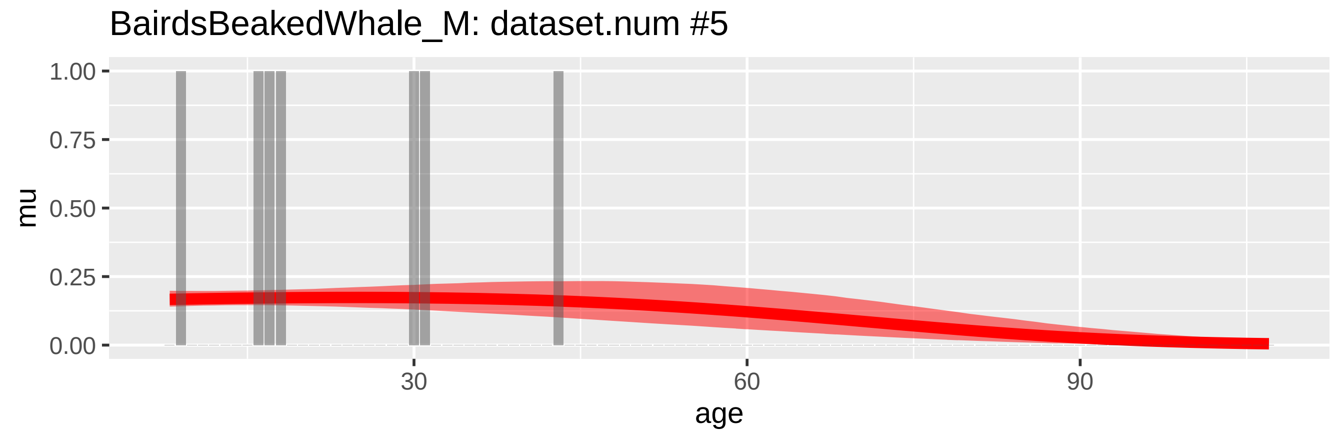
BairdsBeakedWhale_M: dataset.num #5
mu
age
0.00
0.25
0.50
0.75
1.00
30
60
90

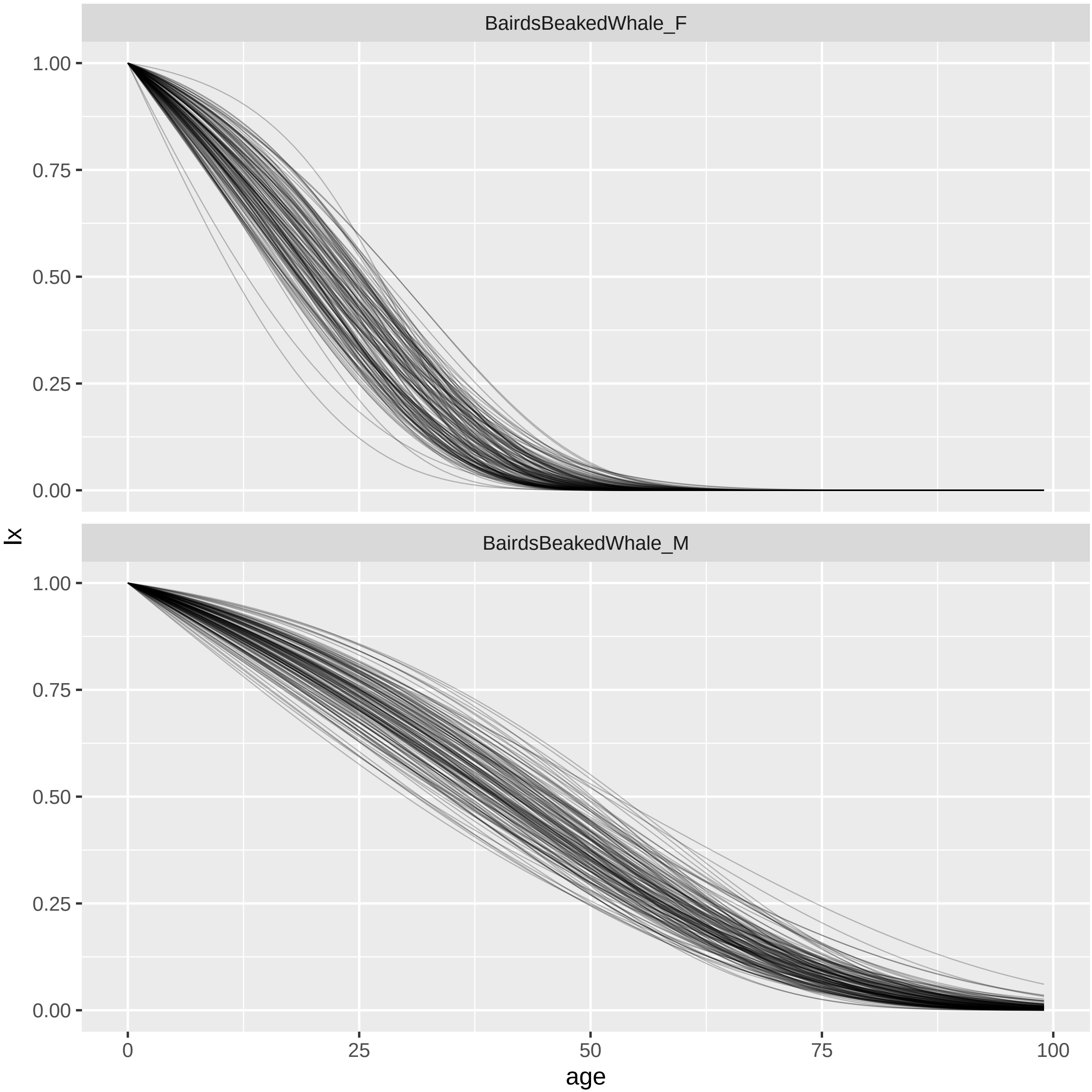

BairdsBeakedWhale_F
BairdsBeakedWhale_M
lx
1.00
0.75
0.50
0.25
0.00
0
25
50
75
100
age

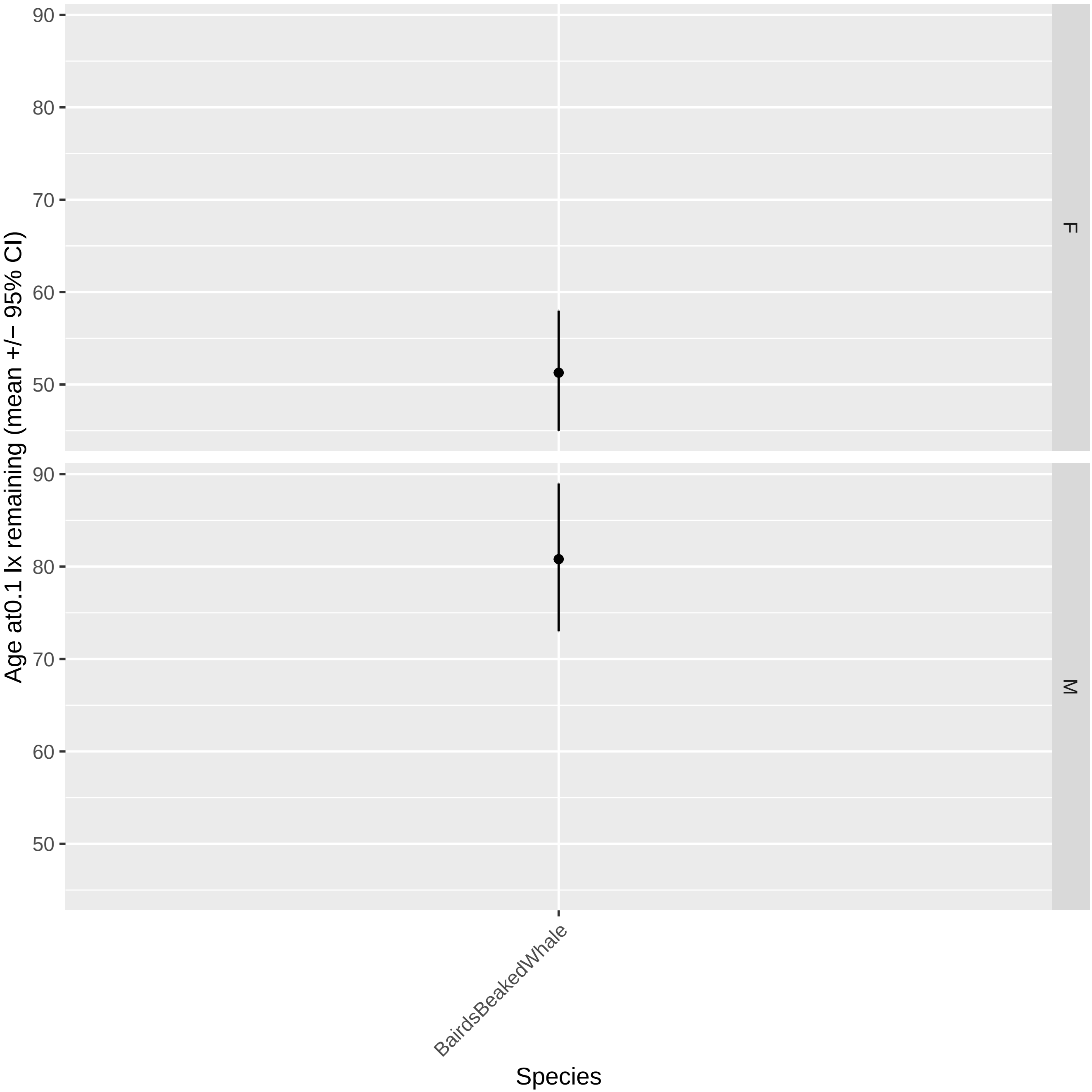

Age at0.1 lx remaining (mean +/− 95% CI)
90
80
70
60
50
F
M
BairdsBeakedWhale
Species

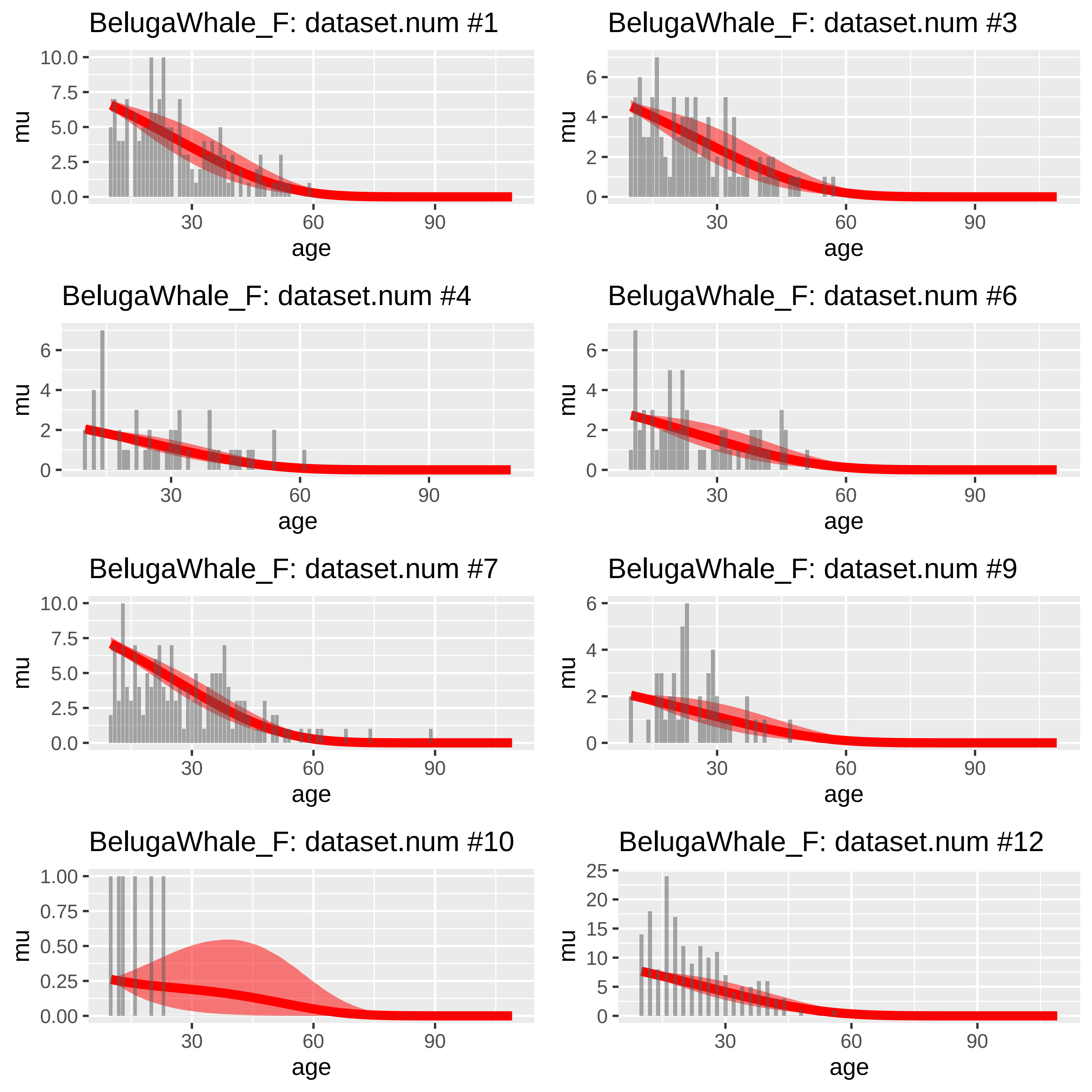
BelugaWhale_F: dataset.num #1
BelugaWhale_F: dataset.num #3
BelugaWhale_F: dataset.num #4
BelugaWhale_F: dataset.num #6
BelugaWhale_F: dataset.num #7
BelugaWhale_F: dataset.num #9
BelugaWhale_F: dataset.num #10
BelugaWhale_F: dataset.num #12
mu
age
30
60
90

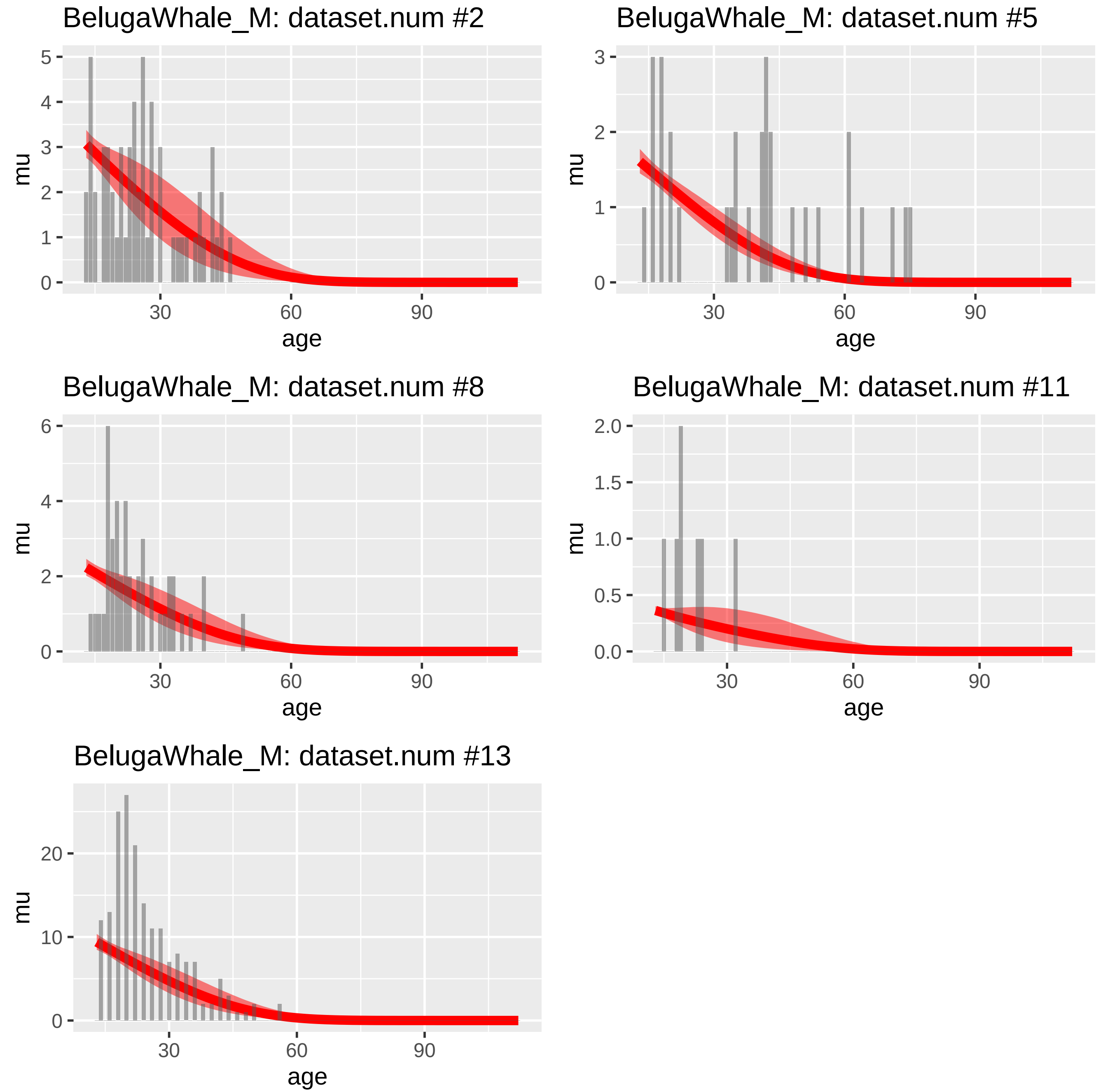
BelugaWhale_M: dataset.num #2
BelugaWhale_M: dataset.num #5
BelugaWhale_M: dataset.num #8
BelugaWhale_M: dataset.num #11
BelugaWhale_M: dataset.num #13
mu
age
0
1
2
3
4
5
6
0.0
0.5
1.0
1.5
2.0
10
20
30
60
90

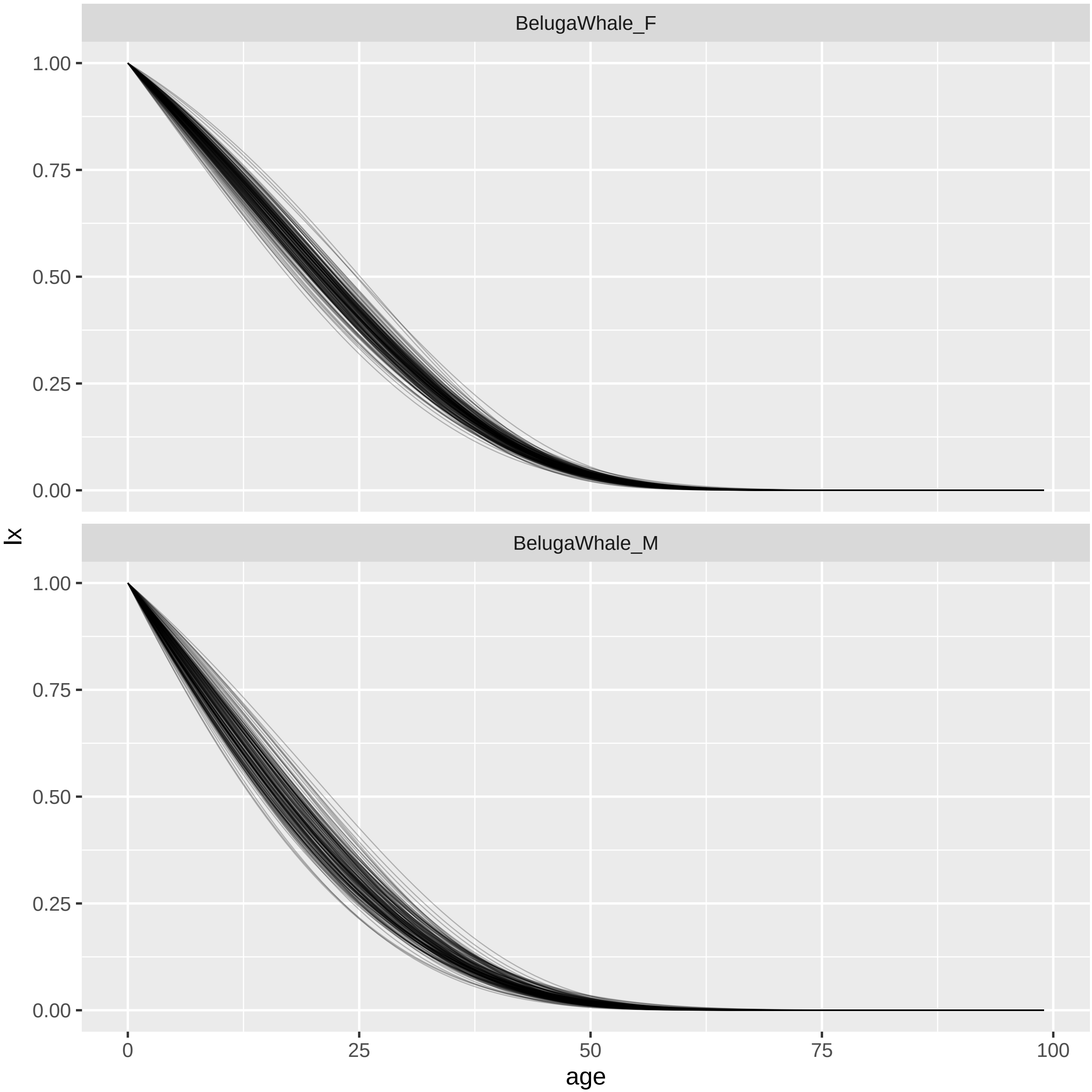

BelugaWhale_F
BelugaWhale_M
1.00
0.75
0.50
0.25
0.00
lx
0
25
50
75
100
age

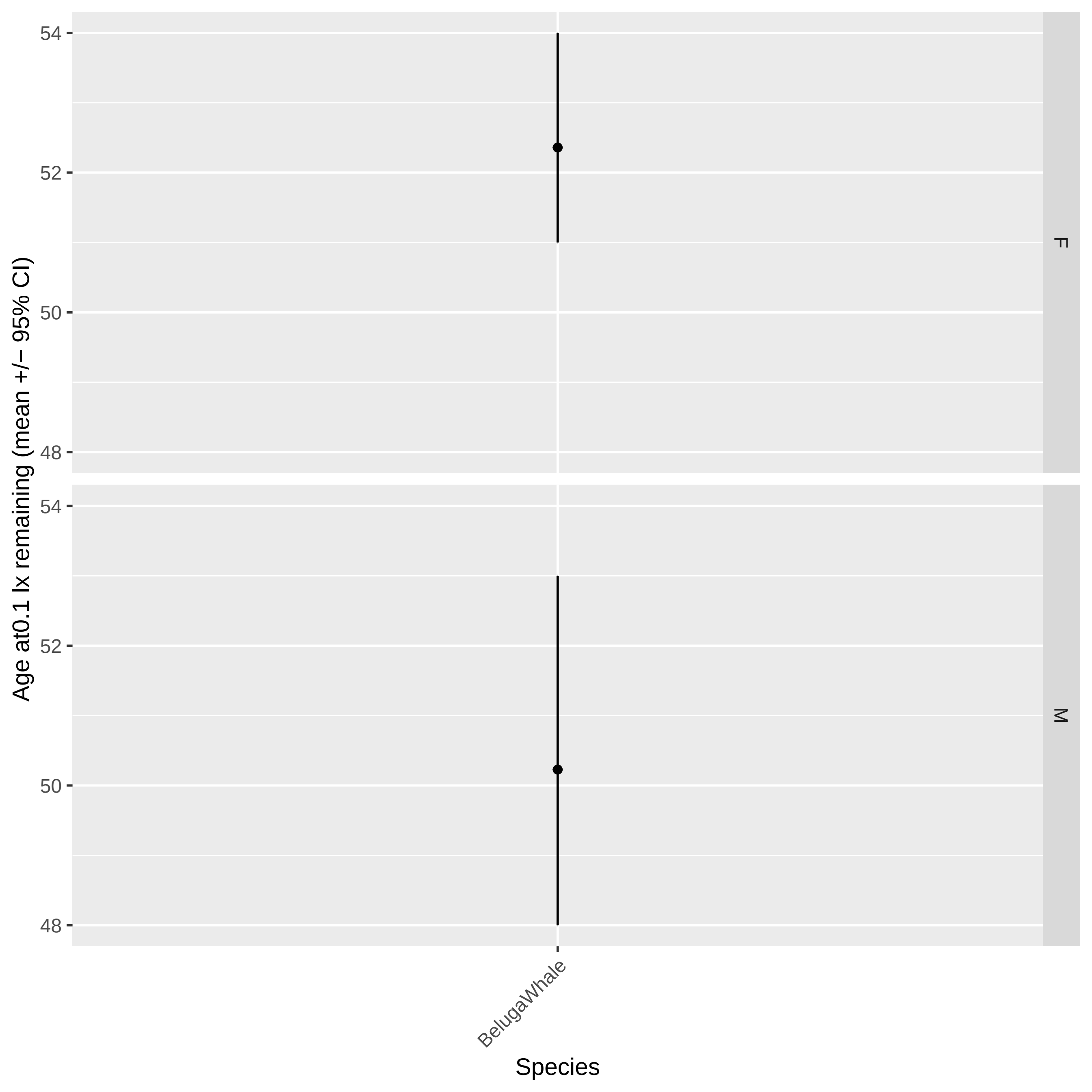

Age at0.1 lx remaining (mean +/− 95% CI)
54
52
50
48
F
M
BelugaWhale
Species

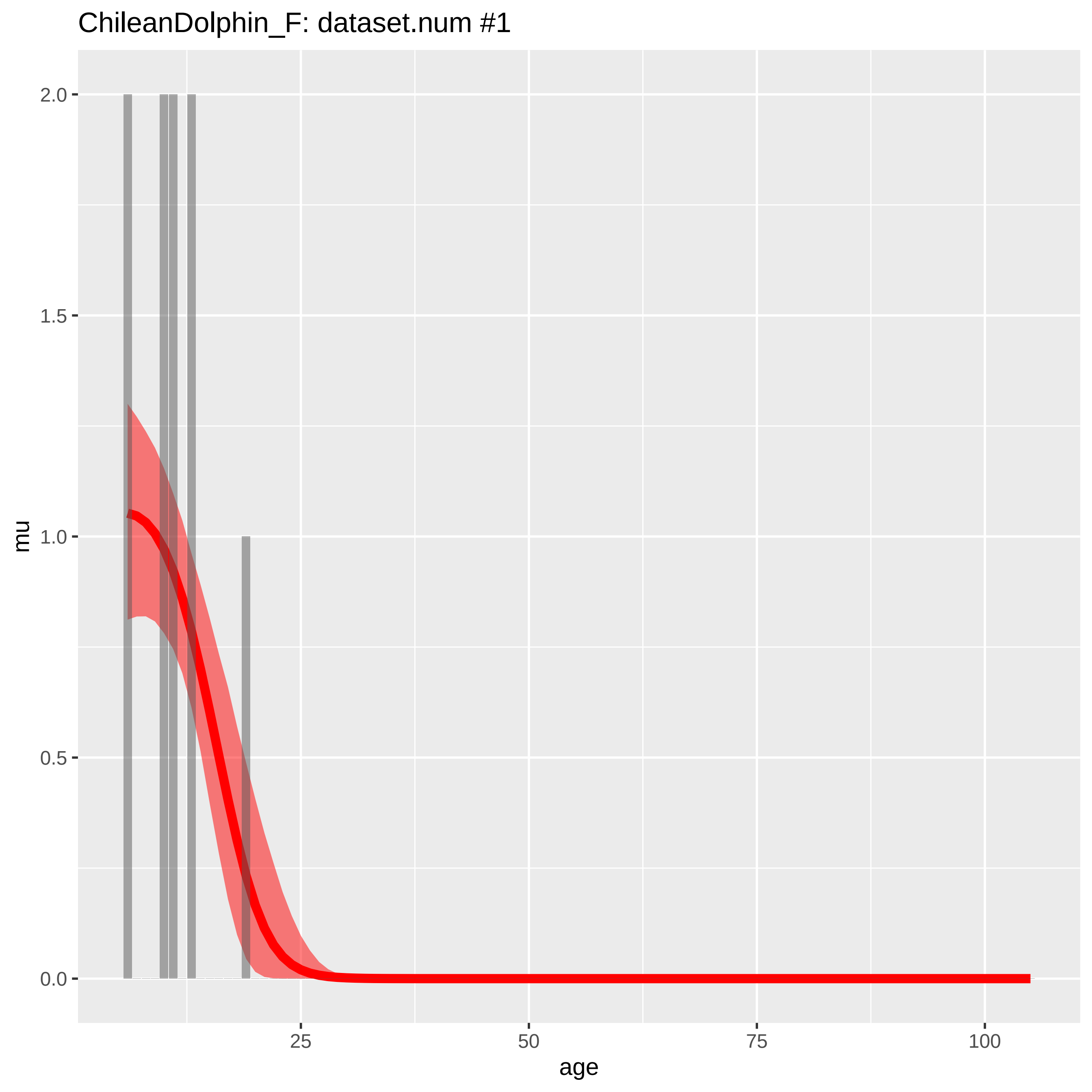

ChileanDolphin_F: dataset.num #1
mu
age
0.0
0.5
1.0
1.5
2.0
25
50
75
100

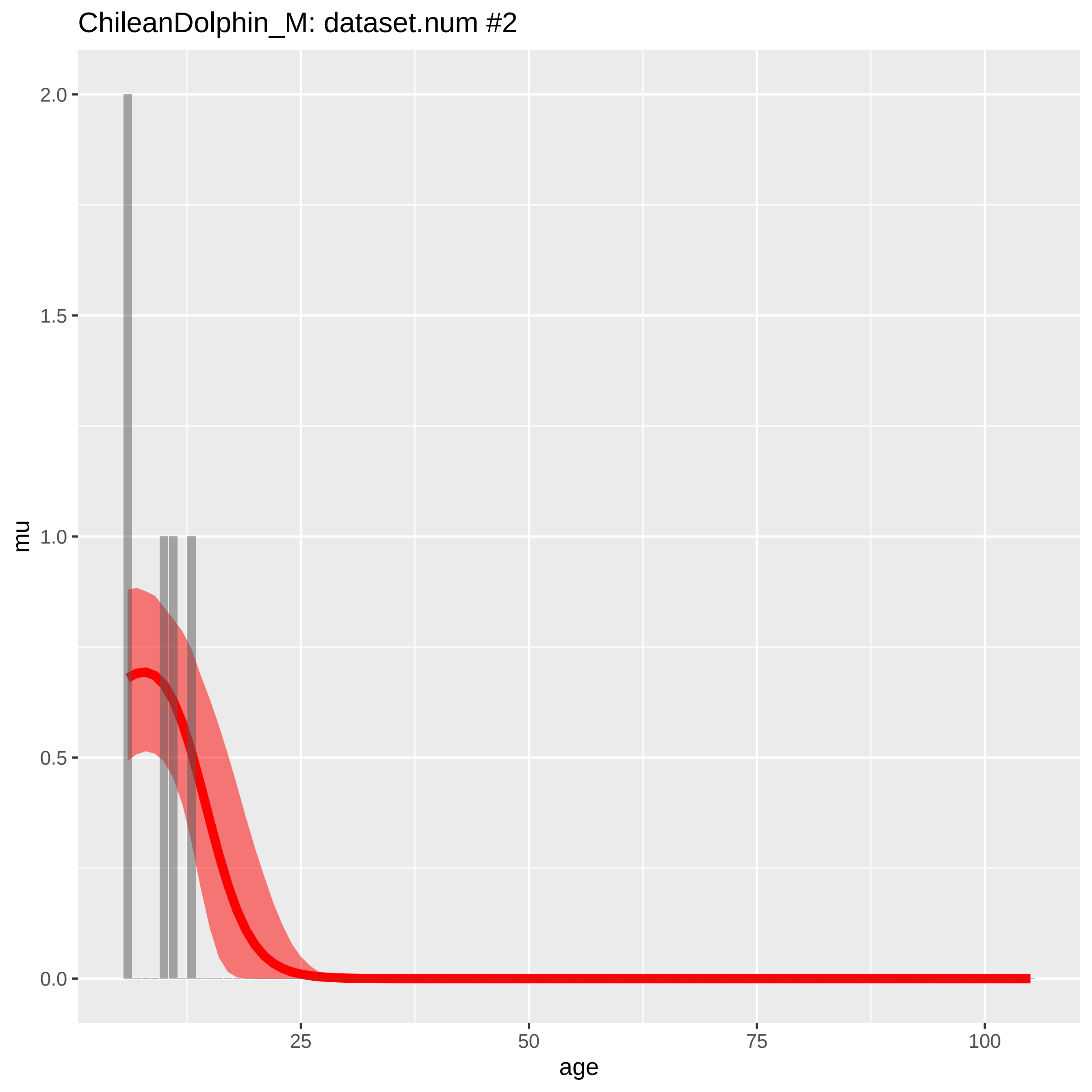
ChileanDolphin_M: dataset.num #2
mu
2.0
1.5
1.0
0.5
0.0
25
50
75
100
age

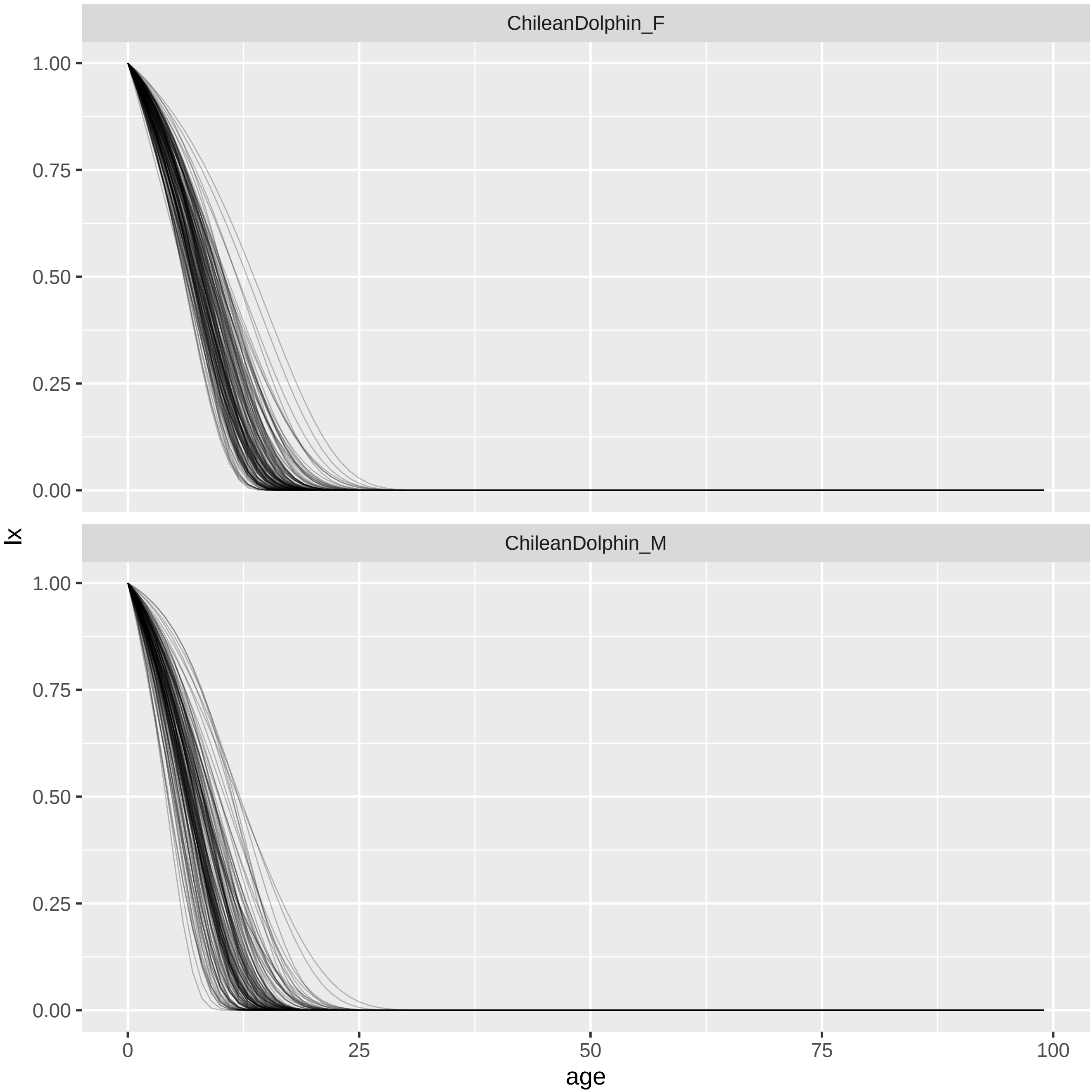
ChileanDolphin_F
ChileanDolphin_M
lx
age
1.00
0.75
0.50
0.25
0.00
0
25
50
75
100

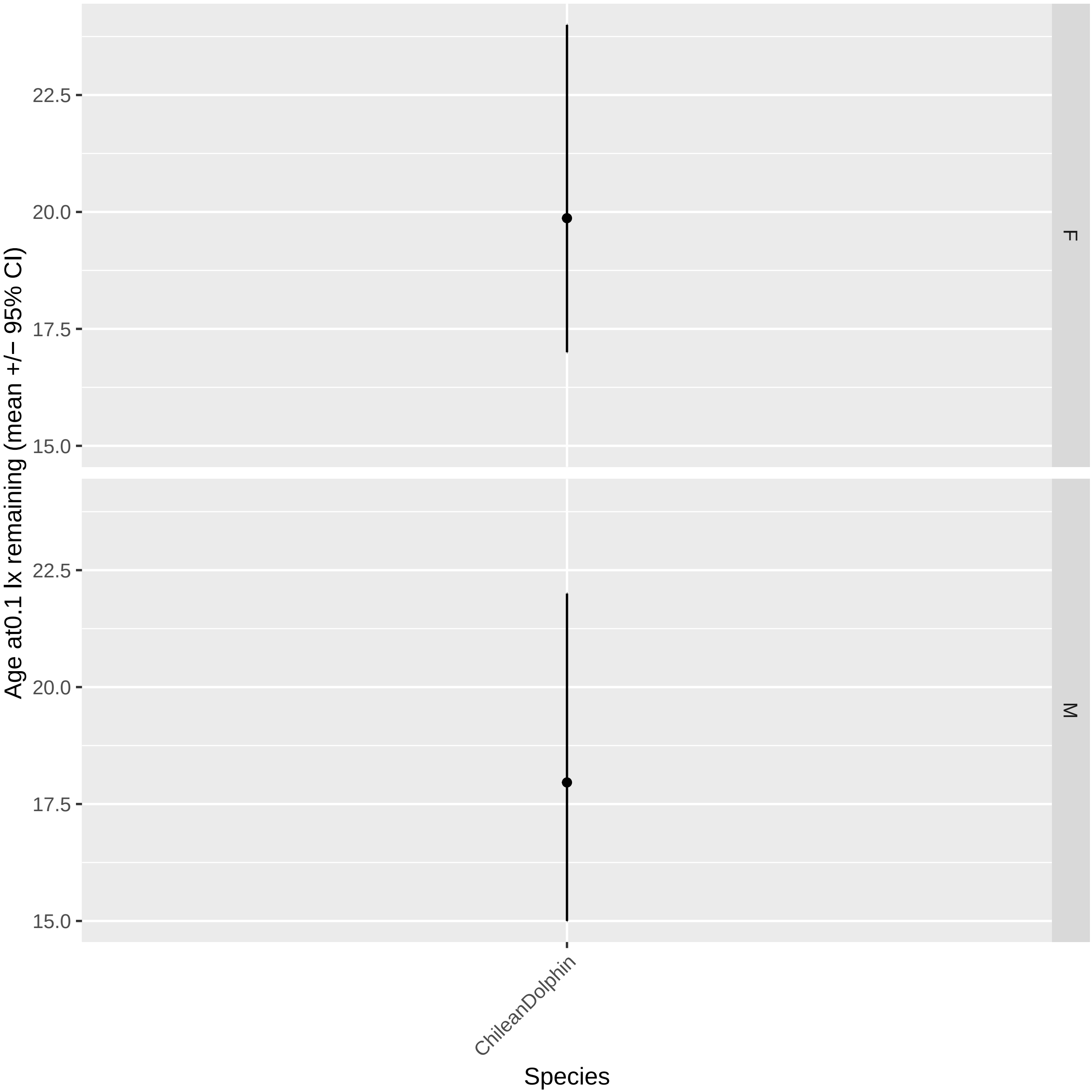

Age at0.1 lx remaining (mean +/− 95% CI)
22.5
20.0
17.5
15.0
F
22.5
20.0
17.5
15.0
M
ChileanDolphin
Species

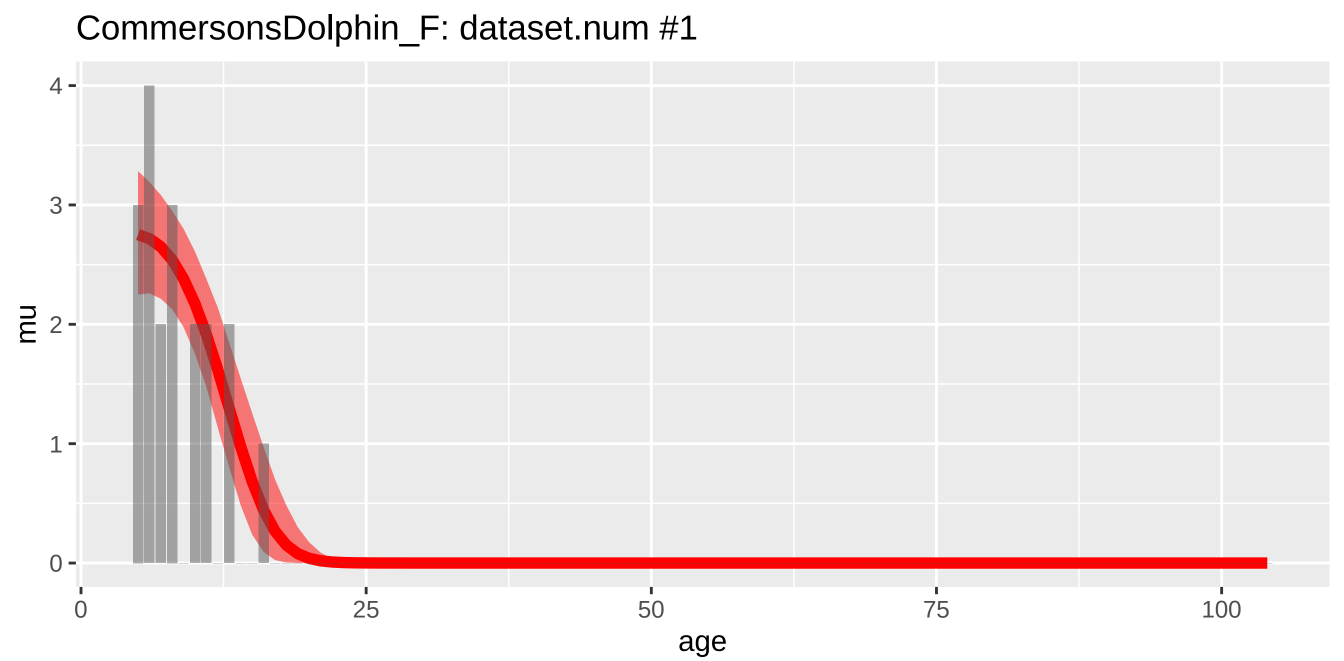

CommersonsDolphin_F: dataset.num #1
mu
4
3
2
1
0
0
25
50
75
100
age


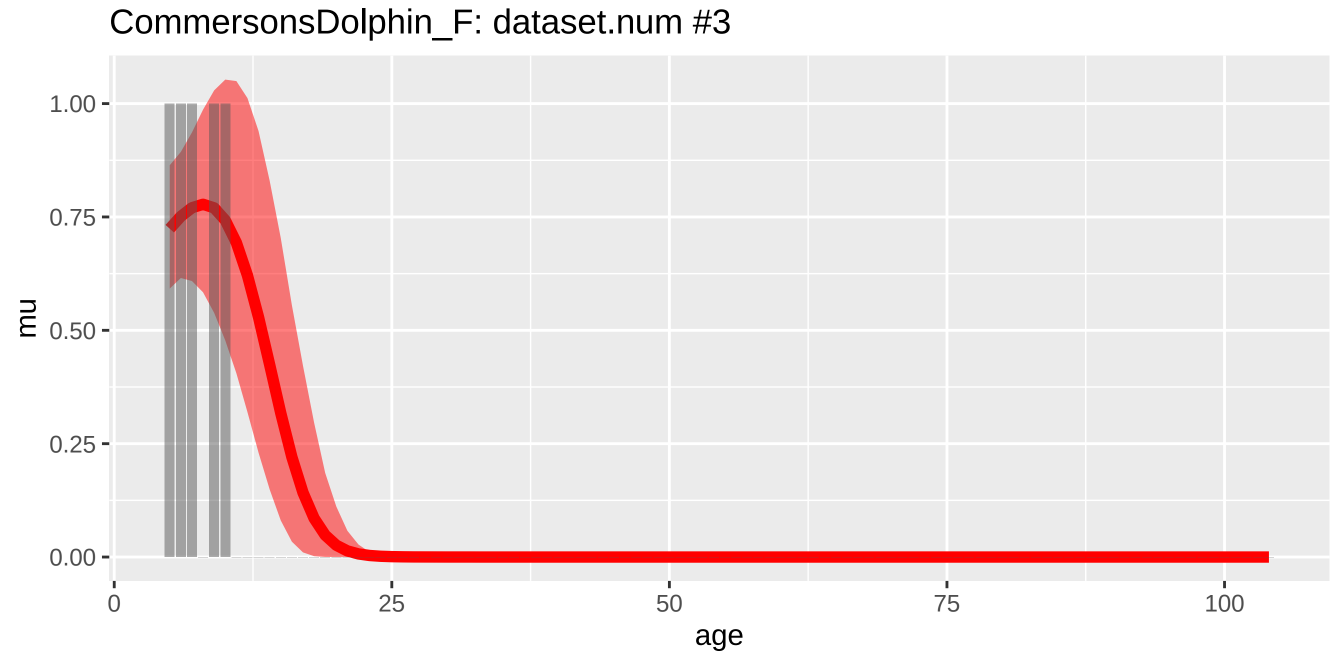

CommersonsDolphin_F: dataset.num #3
mu
1.00
0.75
0.50
0.25
0.00
0
25
50
75
100
age

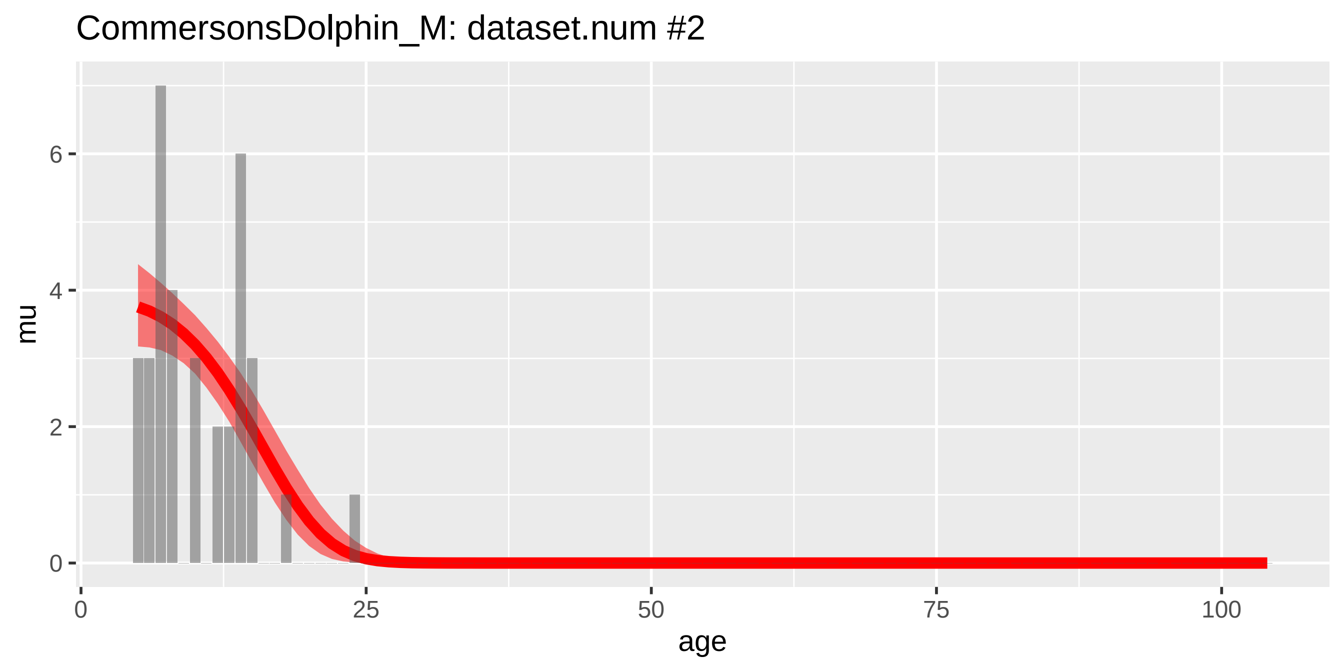
CommersonsDolphin_M: dataset.num #2
mu
age
0
2
4
6
0
25
50
75
100

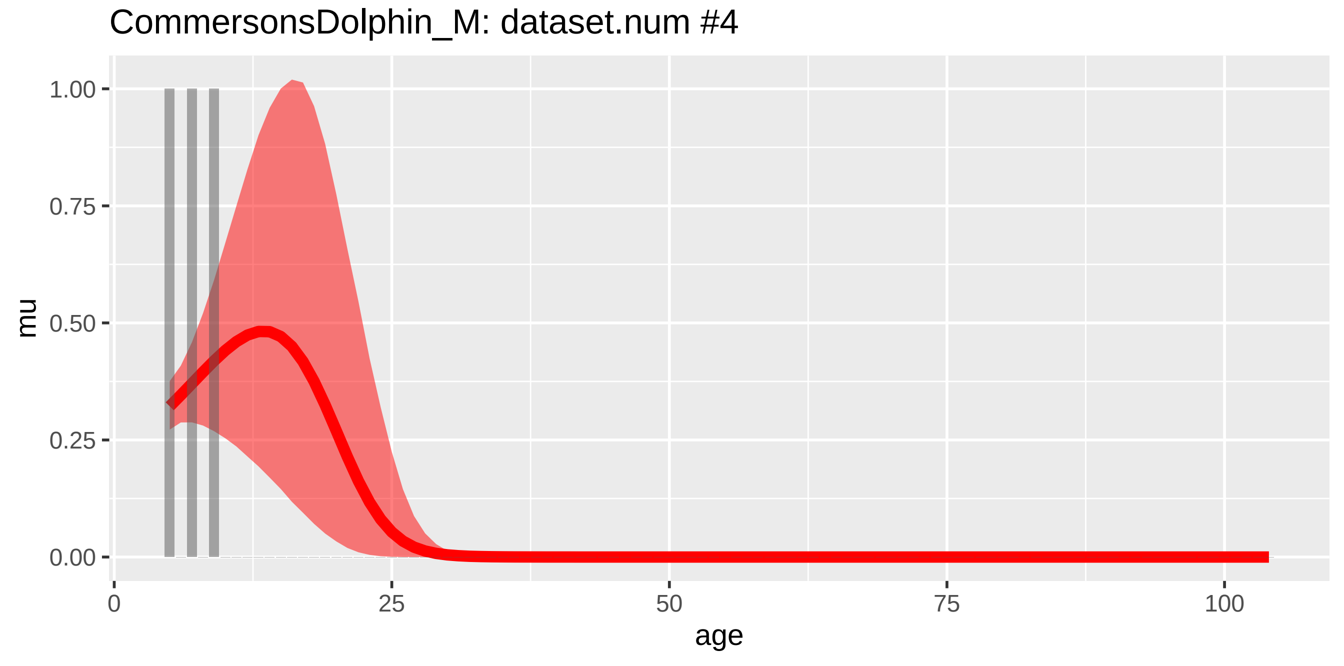
CommersonsDolphin_M: dataset.num #4
mu
age
0.00
0.25
0.50
0.75
1.00
0
25
50
75
100

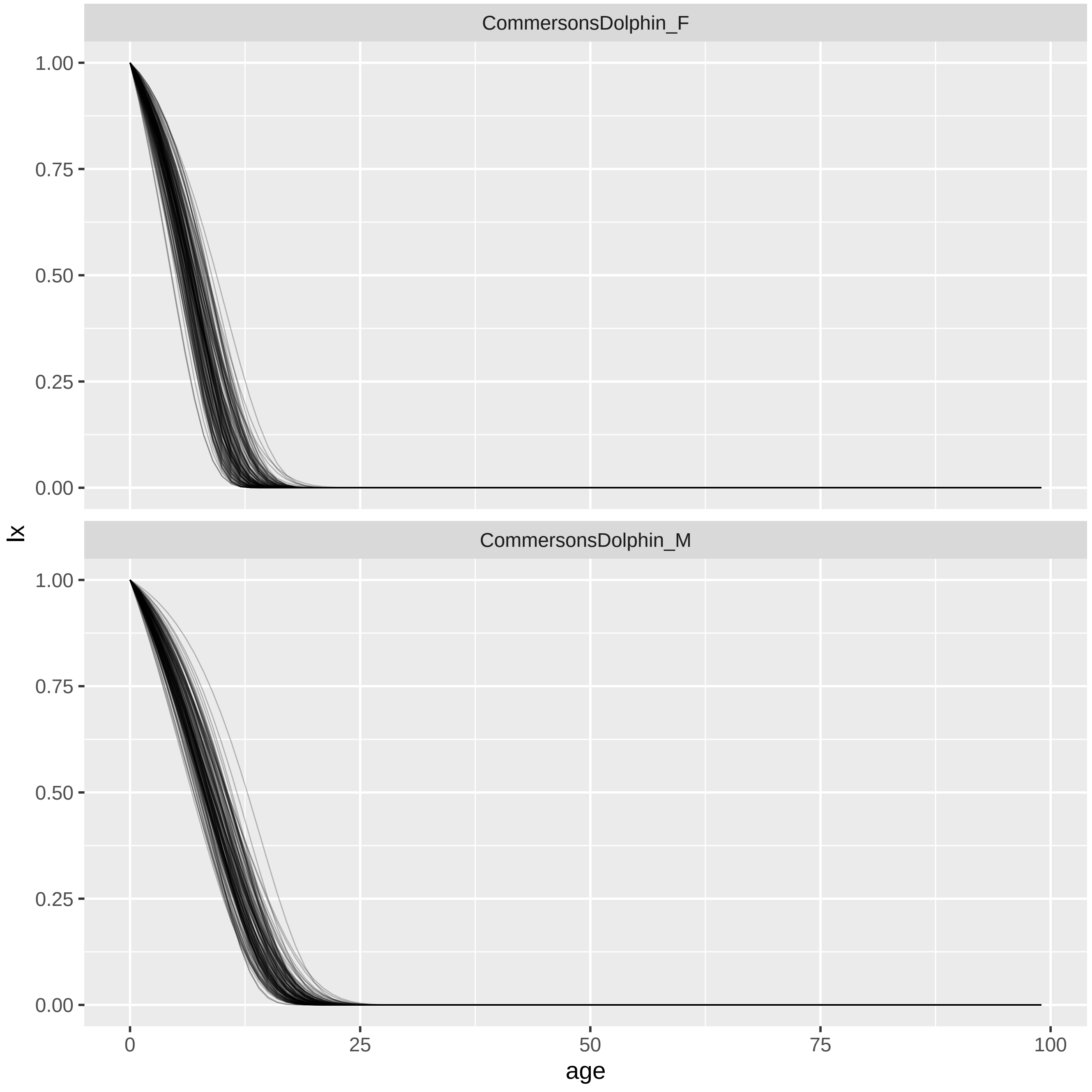
CommersonsDolphin_F
CommersonsDolphin_M
1.00
0.75
0.50
0.25
0.00
lx
0
25
50
75
100
age

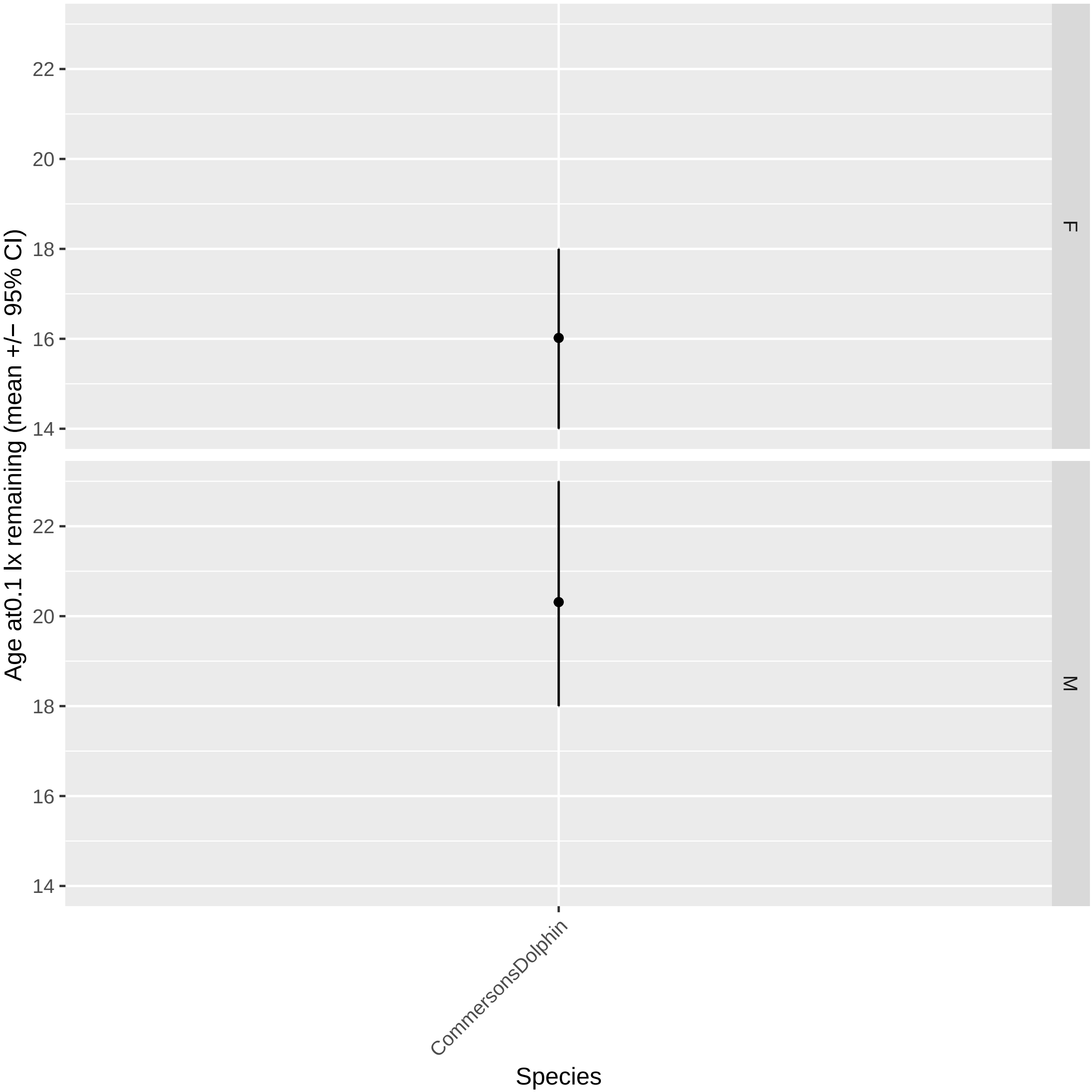

Age at0.1 lx remaining (mean +/− 95% CI)
22
20
18
16
14
F
M
CommersonsDolphin
Species

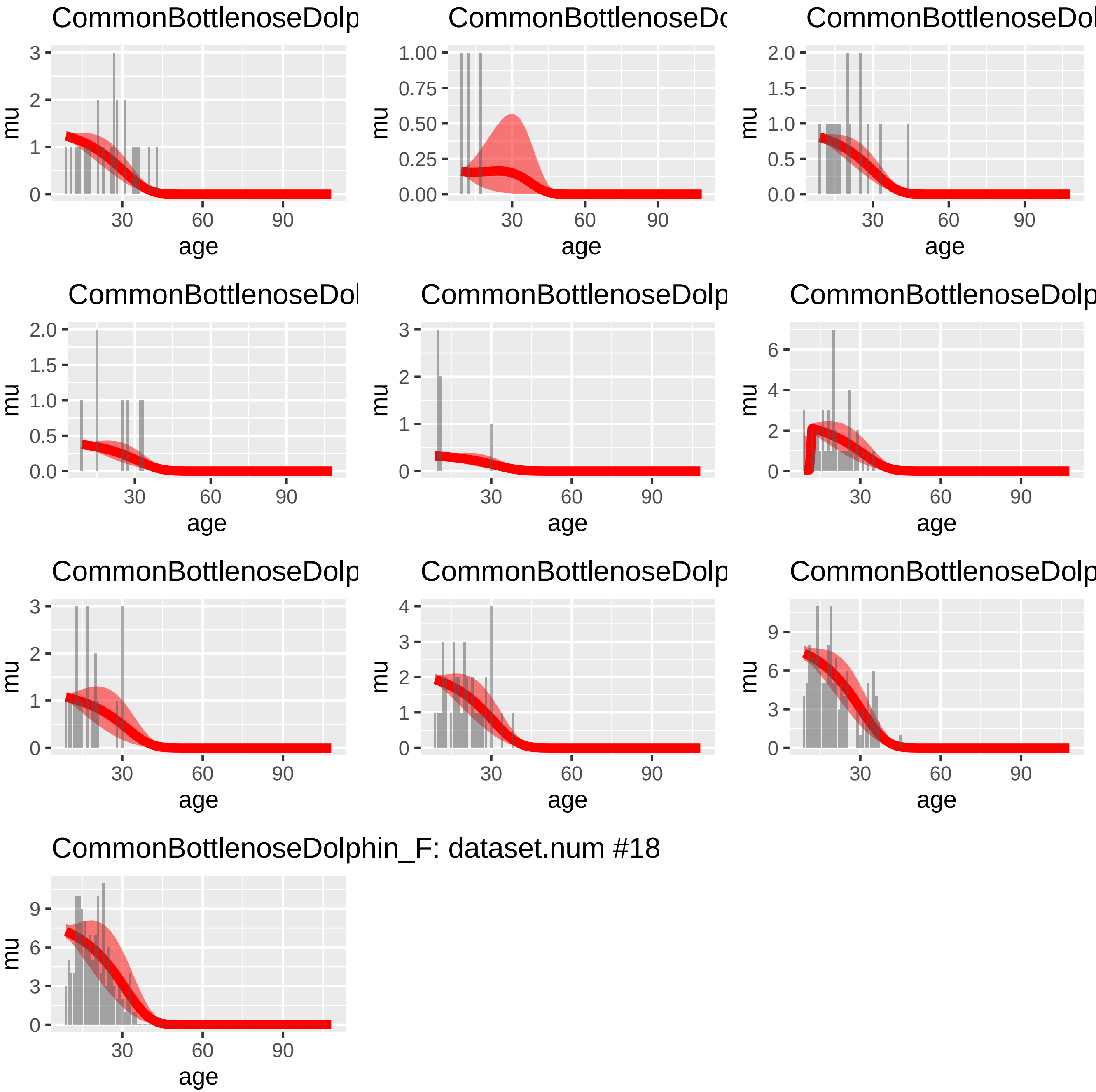
CommonBottlenoseDolp
CommonBottlenoseDo
CommonBottlenoseDol
CommonBottlenoseDol
CommonBottlenoseDolp
CommonBottlenoseDolp
CommonBottlenoseDolp
CommonBottlenoseDolp
CommonBottlenoseDolp
CommonBottlenoseDolphin_F: dataset.num #18
mu
age
30
60
90

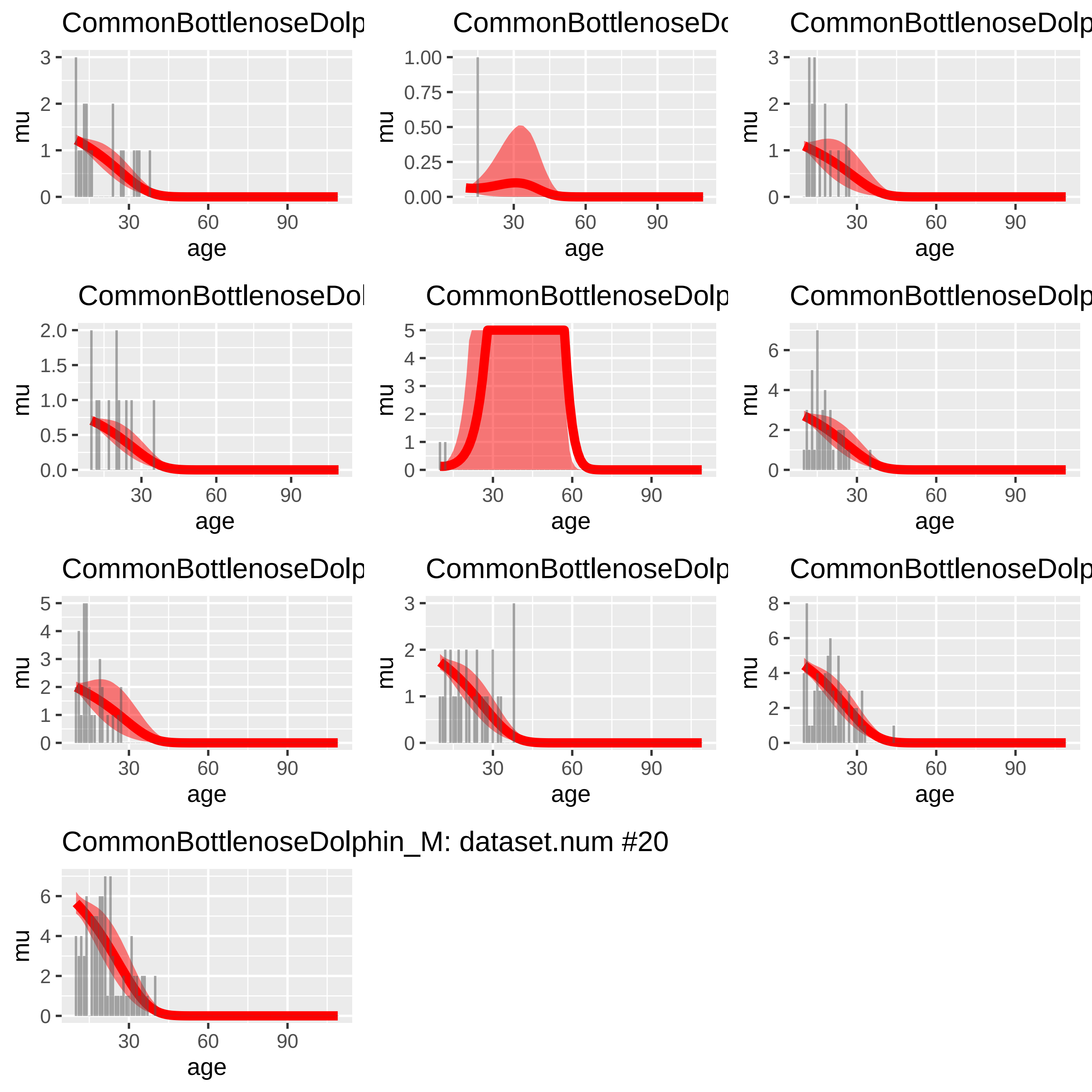
CommonBottlenoseDolp
CommonBottlenoseDc
CommonBottlenoseDolp
CommonBottlenoseDol
CommonBottlenoseDolp
CommonBottlenoseDolp
CommonBottlenoseDolp
CommonBottlenoseDolp
CommonBottlenoseDolp
CommonBottlenoseDolphin_M: dataset.num #20
mu
age

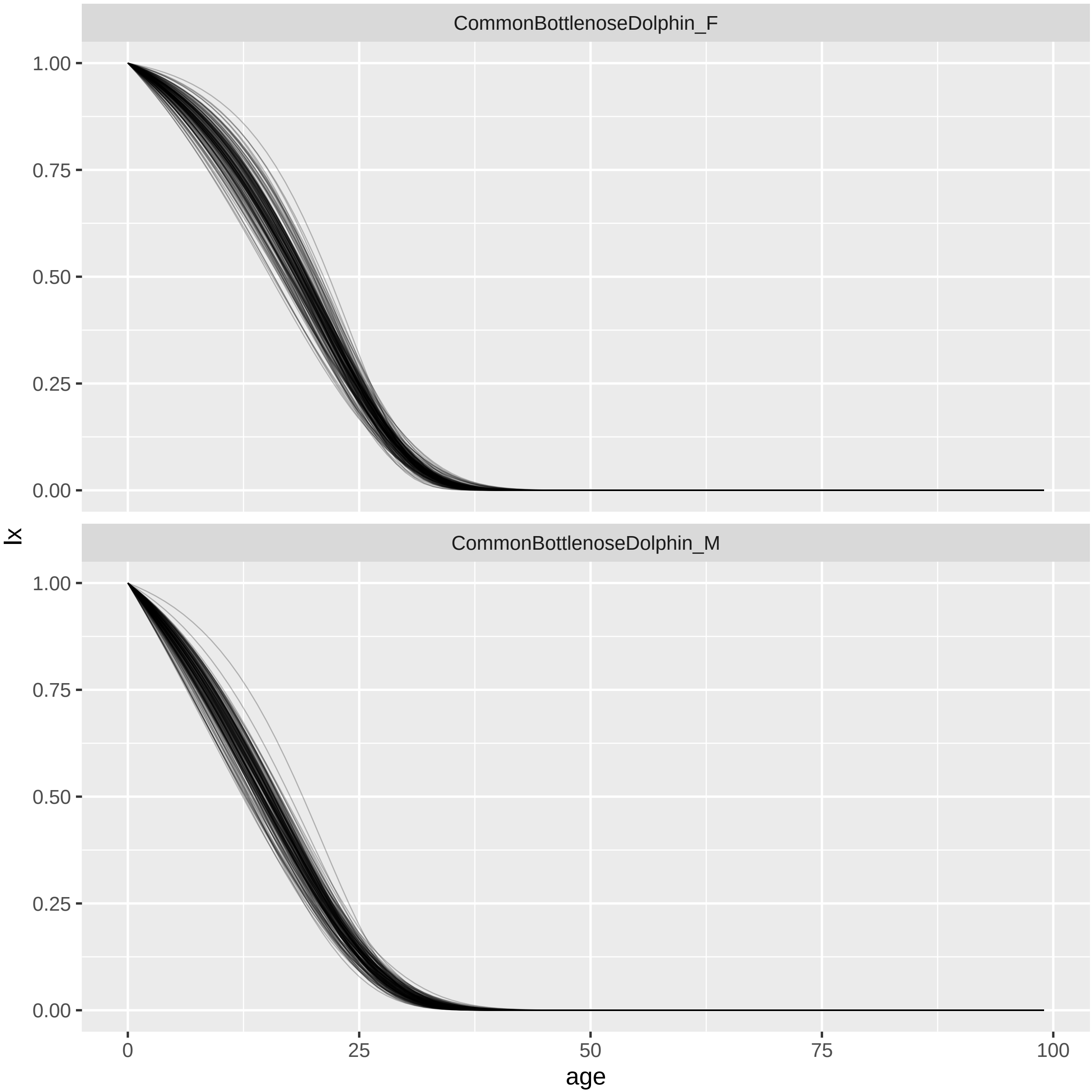
CommonBottlenoseDolphin_F
CommonBottlenoseDolphin_M
lx
age
1.00
0.75
0.50
0.25
0.00
0
25
50
75
100

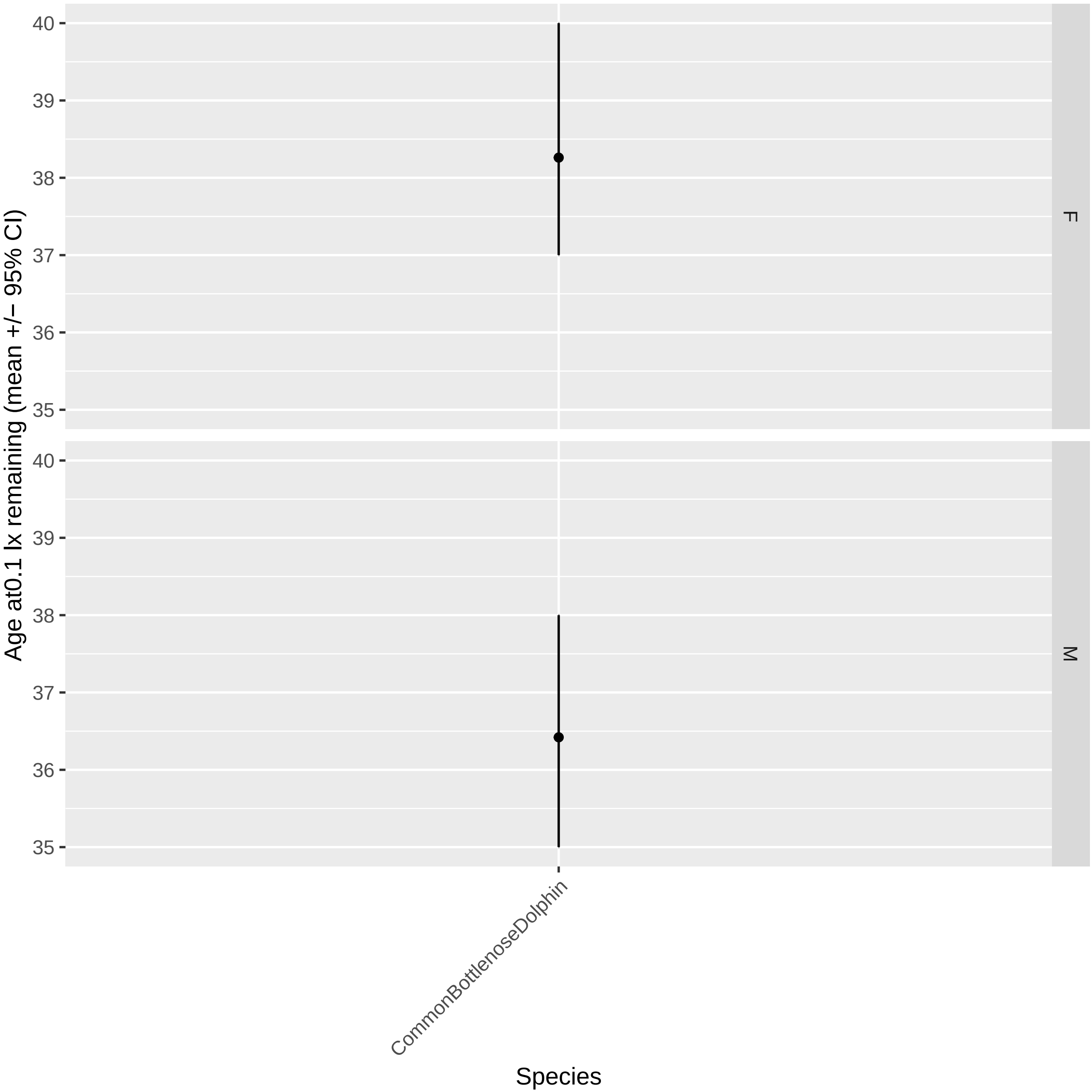

Age at0.1 lx remaining (mean +/− 95% CI)
40
39
38
37
36
35
F
M
CommonBottlenoseDolphin
Species

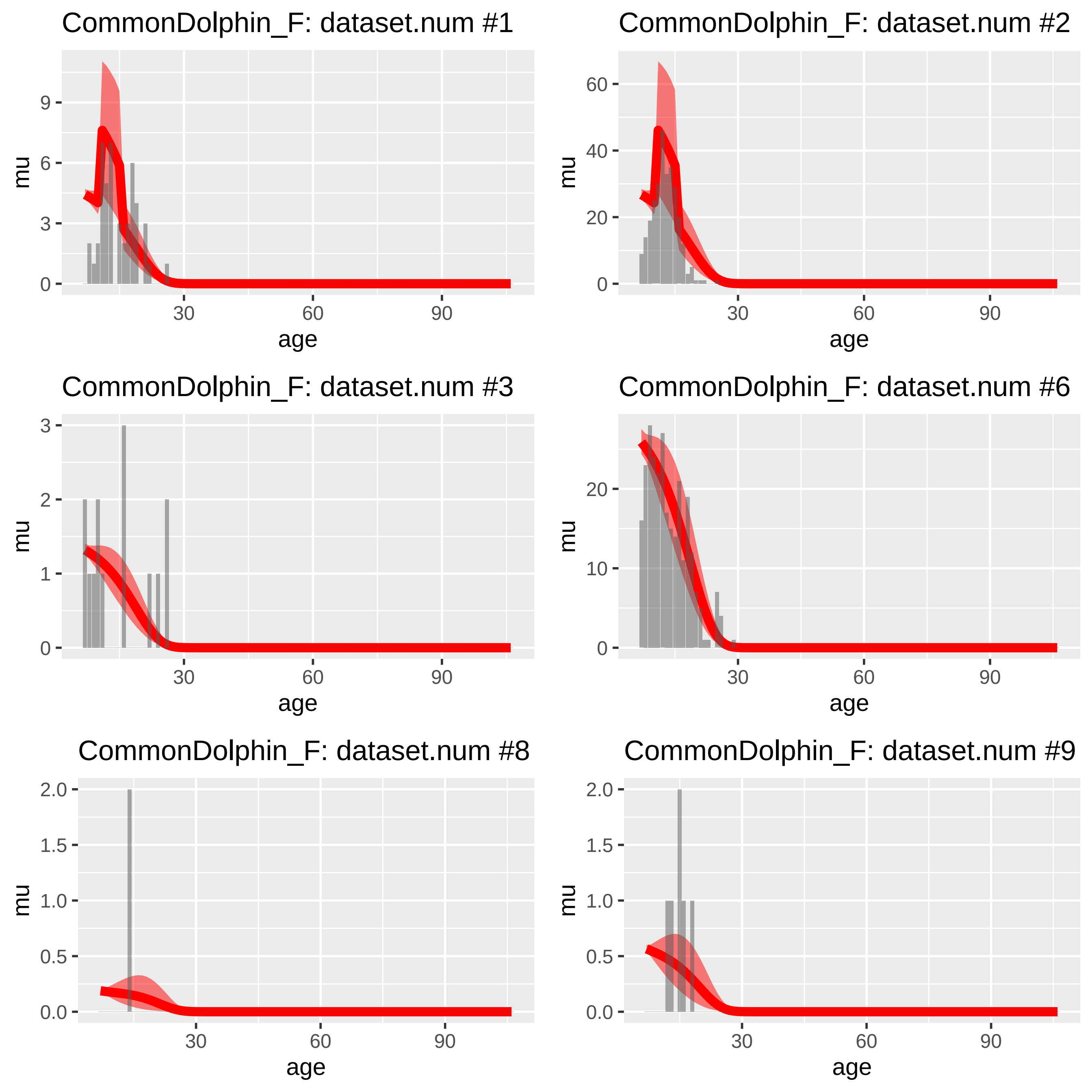
CommonDolphin_F: dataset.num #1
CommonDolphin_F: dataset.num #2
CommonDolphin_F: dataset.num #3
CommonDolphin_F: dataset.num #6
CommonDolphin_F: dataset.num #8
CommonDolphin_F: dataset.num #9
mu
age
0
3
6
9
20
40
60
1
2
10
0.0
0.5
1.0
1.5
2.0
30
60
90

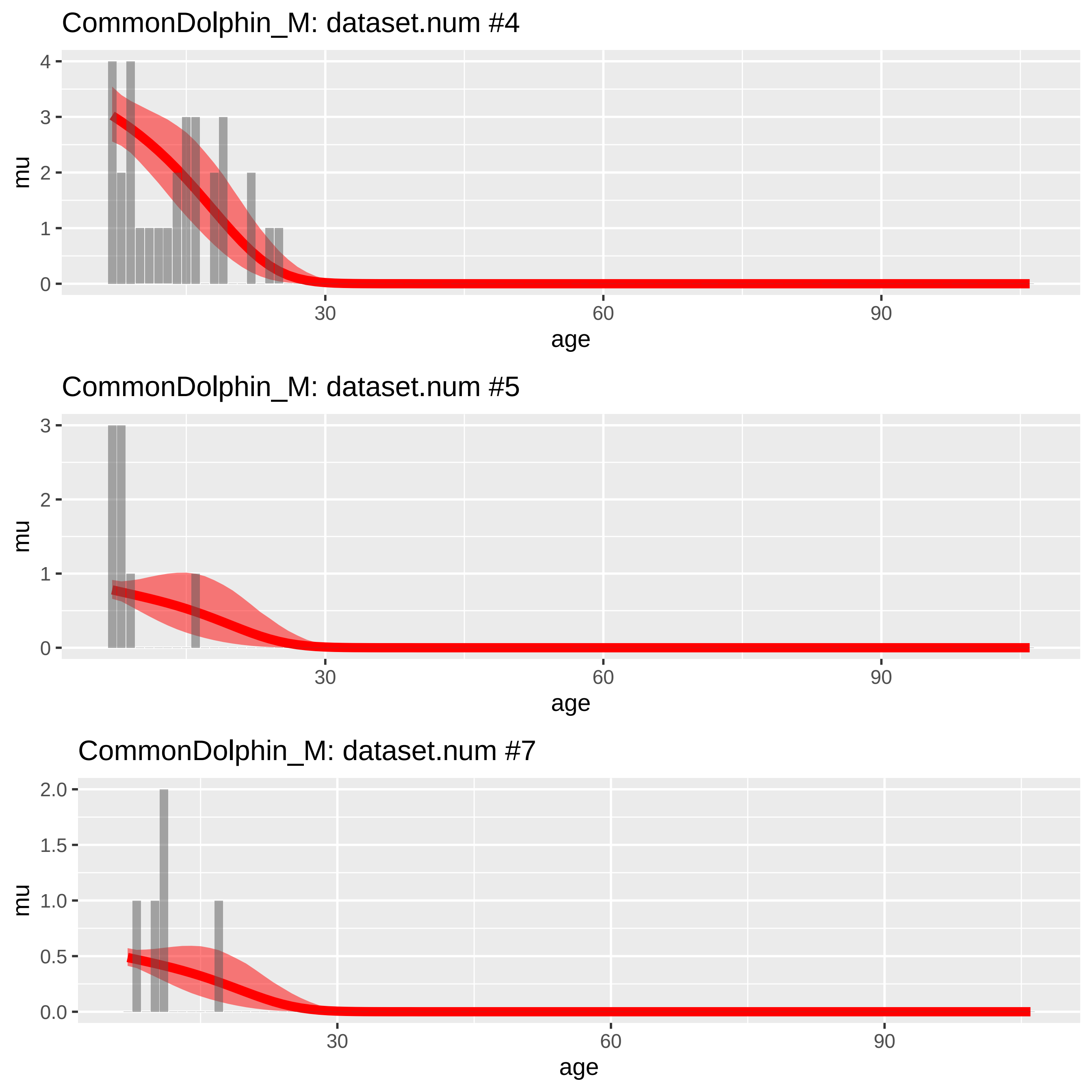
CommonDolphin_M: dataset.num #4
mu
0
1
2
3
4
30
60
90
age
CommonDolphin_M: dataset.num #5
mu
0
1
2
3
30
60
90
age
CommonDolphin_M: dataset.num #7
mu
0.0
0.5
1.0
1.5
2.0
30
60
90
age

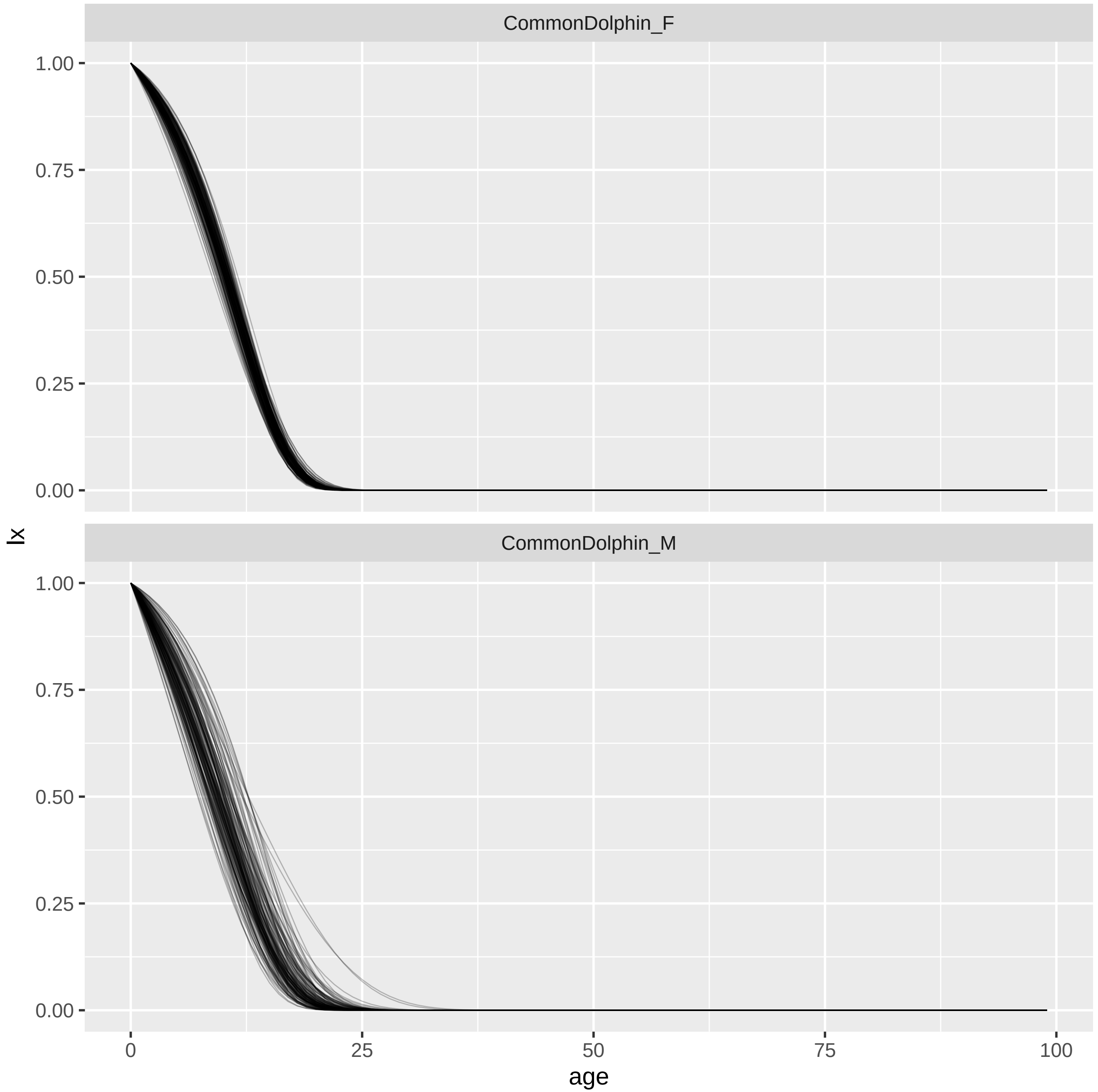

CommonDolphin_F
CommonDolphin_M
1.00
0.75
0.50
0.25
0.00
lx
0
25
50
75
100
age

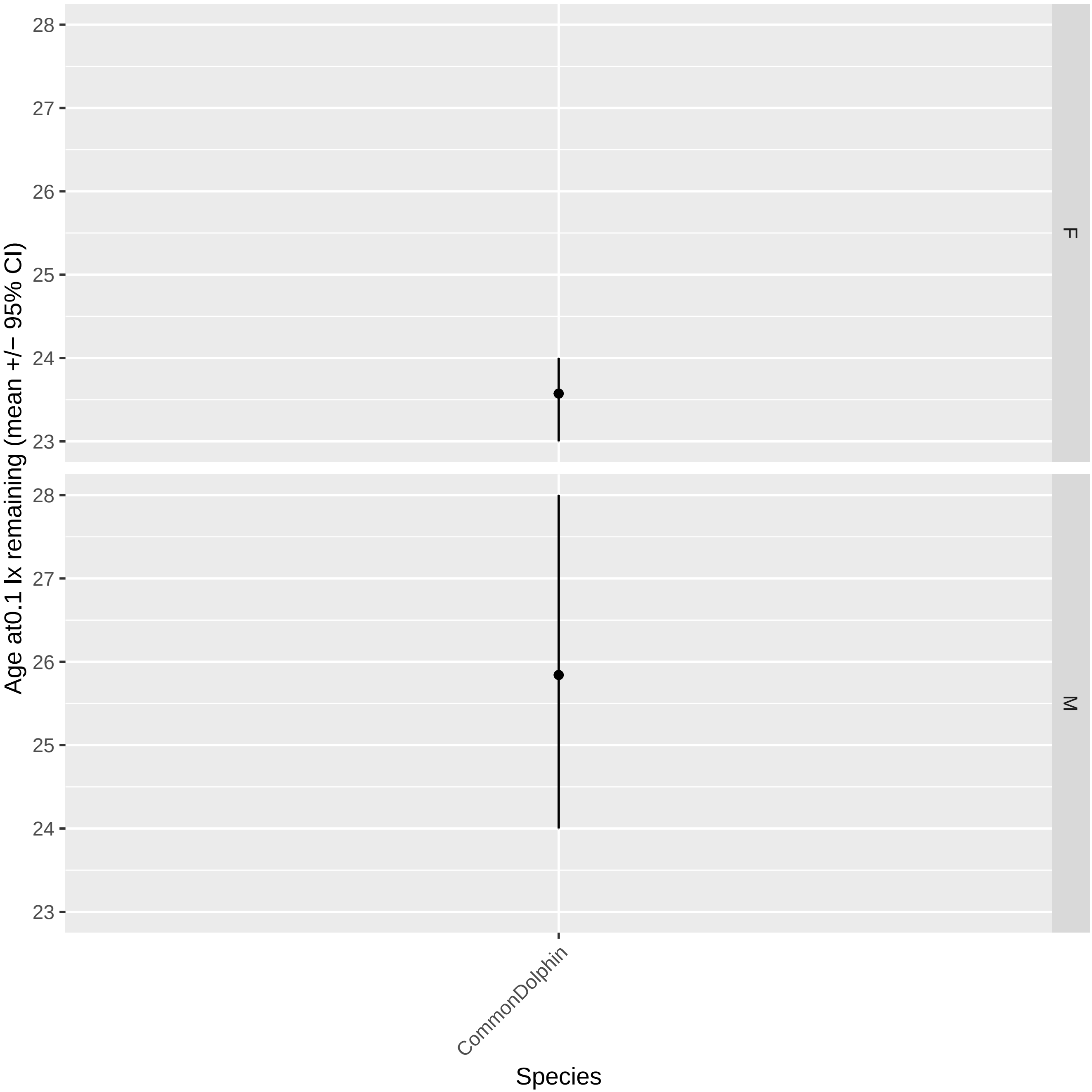

Age at0.1 lx remaining (mean +/− 95% CI)
28
27
26
25
24
23
F
M
CommonDolphin
Species

CuviersBeakedWhale_F: dataset.num #1

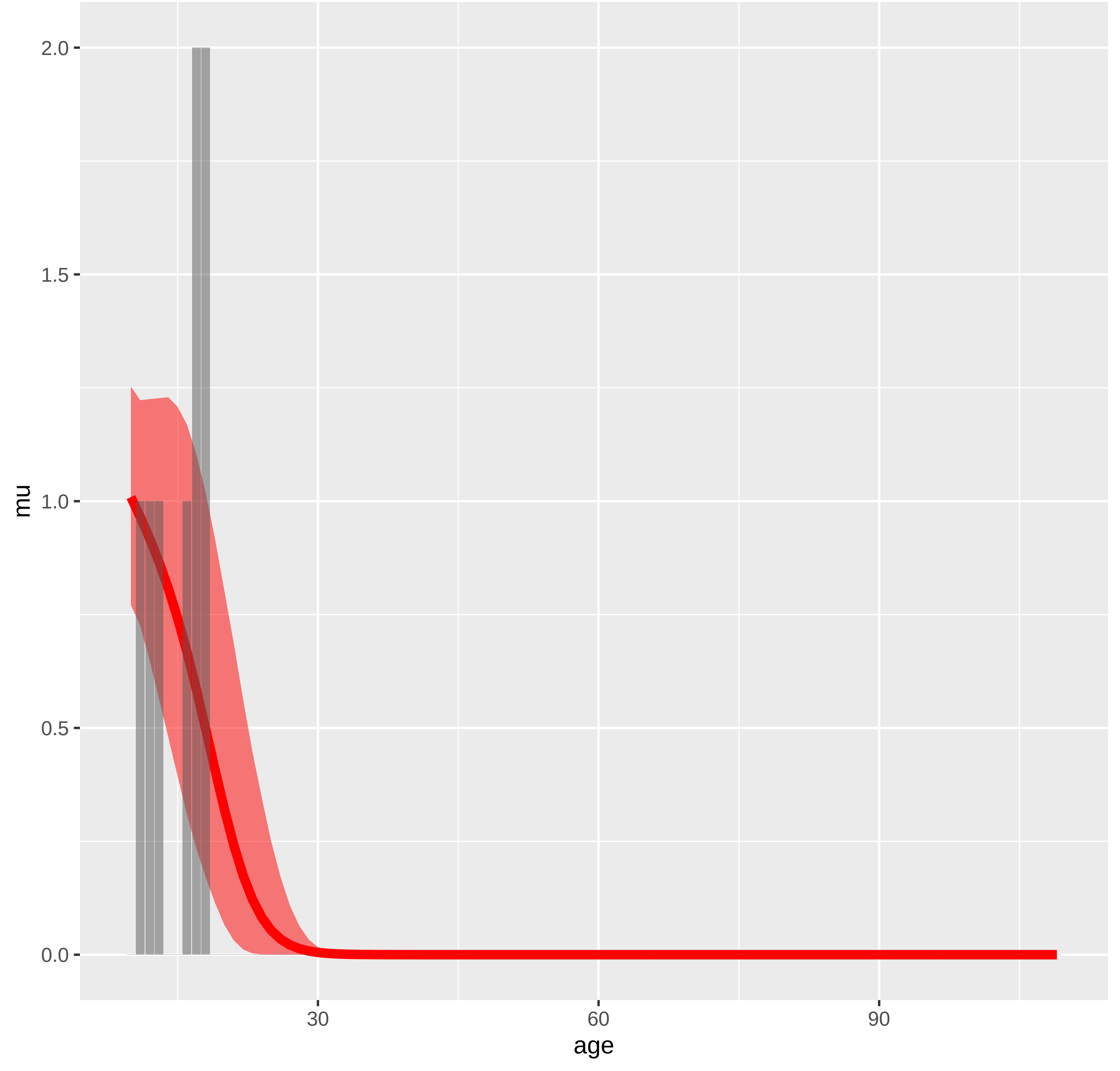

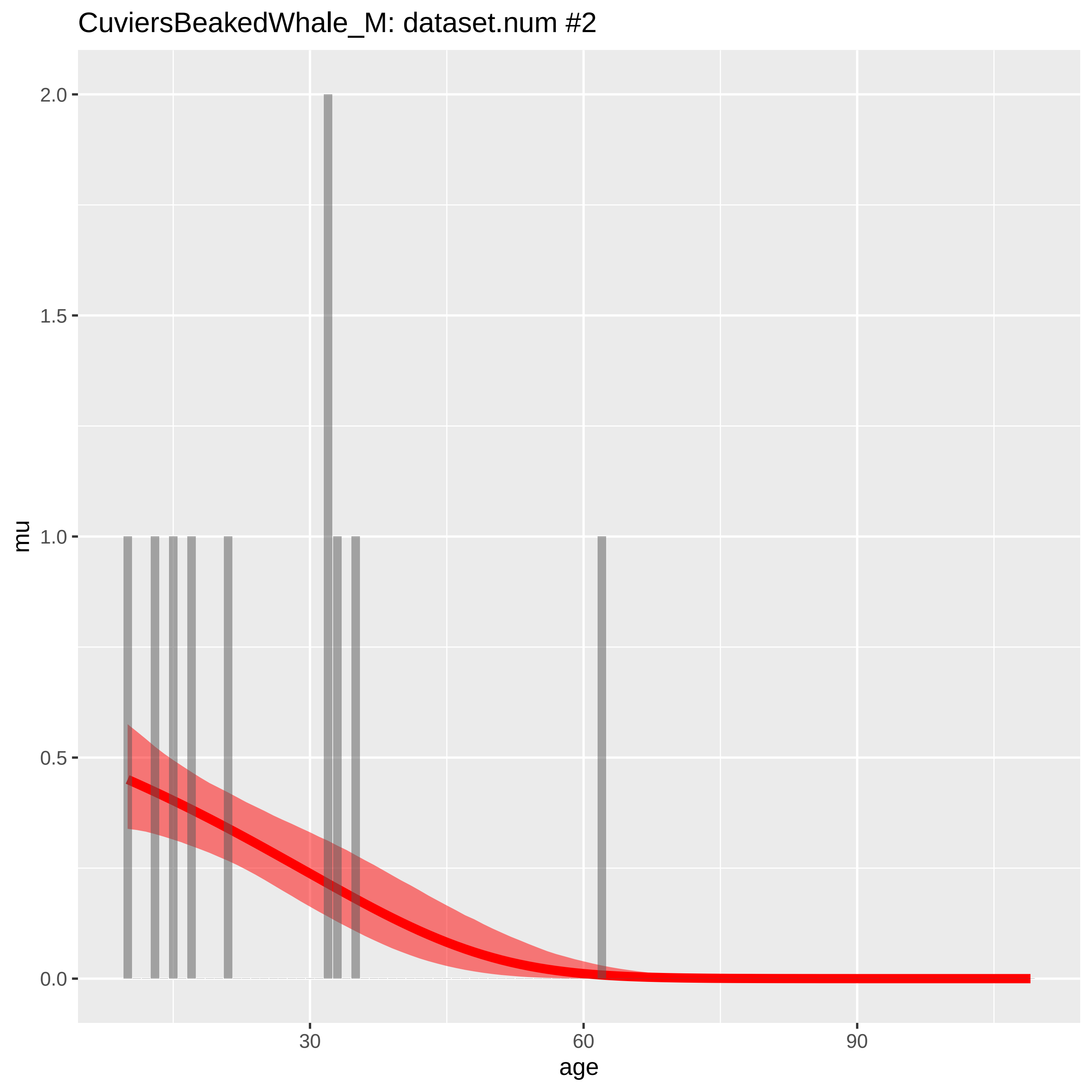
CuviersBeakedWhale_M: dataset.num #2
mu
2.0
1.5
1.0
0.5
0.0
30
60
90
age

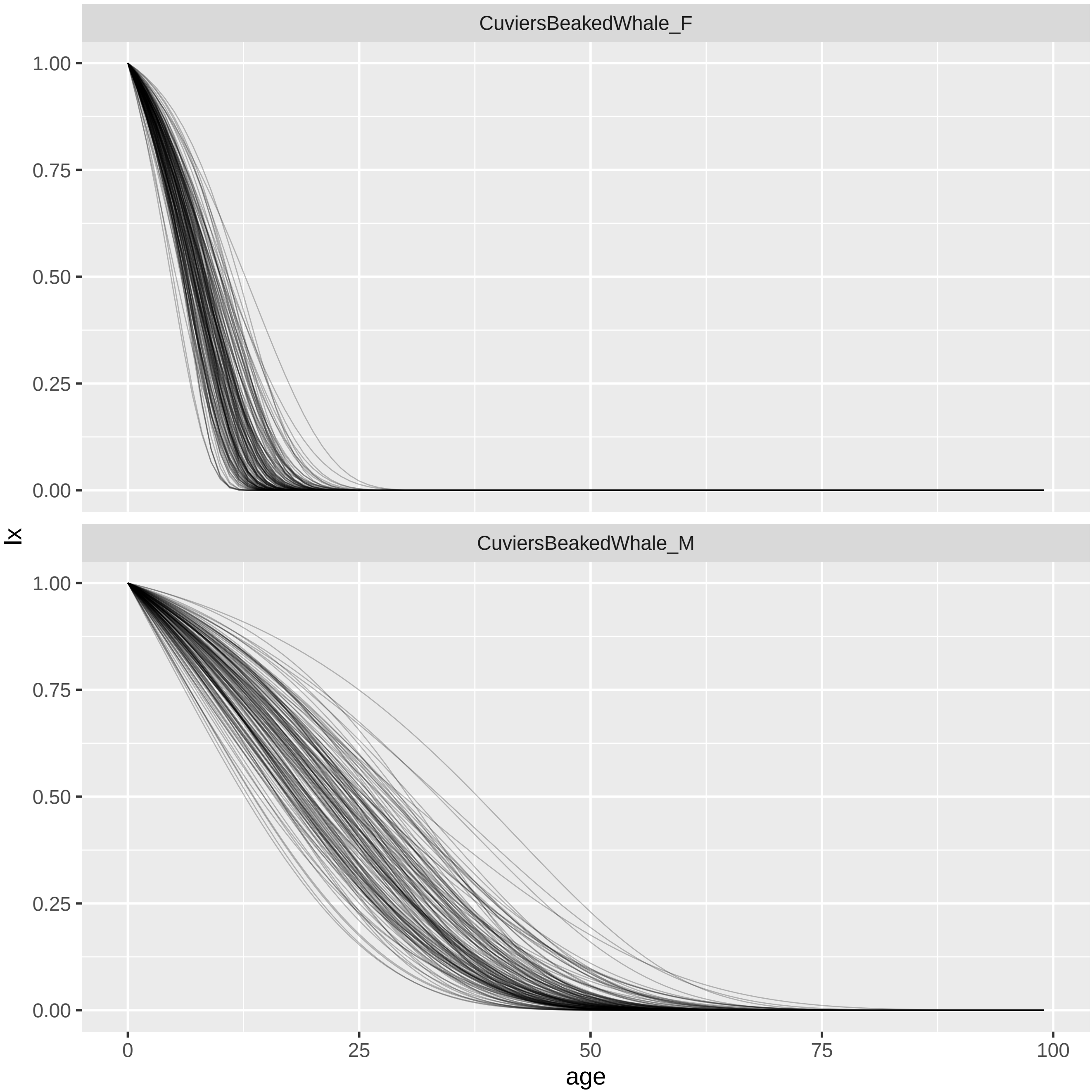
CuviersBeakedWhale_F
CuviersBeakedWhale_M
1.00
0.75
0.50
0.25
0.00
lx
0
25
50
75
100
age

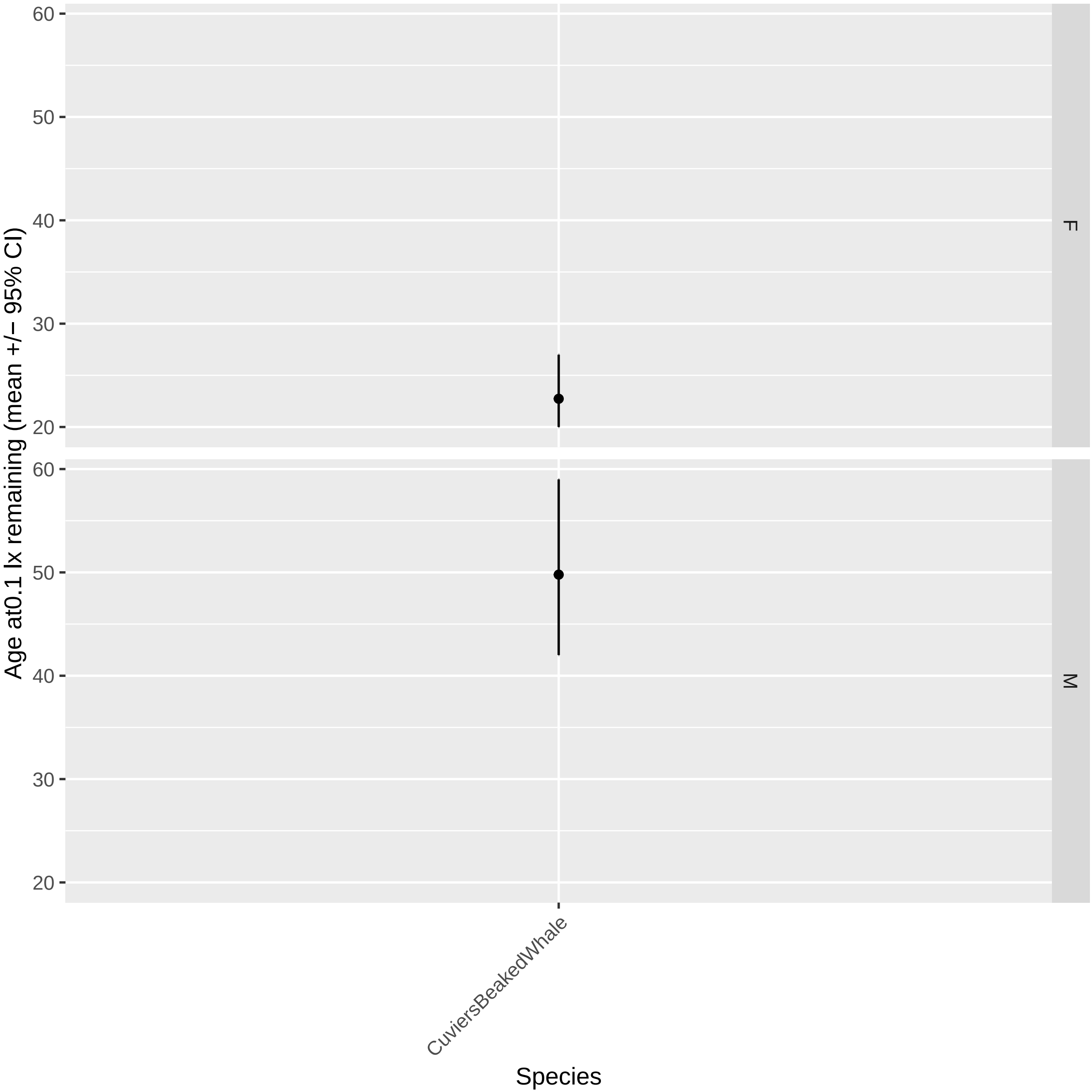

Age at0.1 lx remaining (mean +/− 95% CI)
60
50
40
30
20
F
M
CuviersBeakedWhale
Species

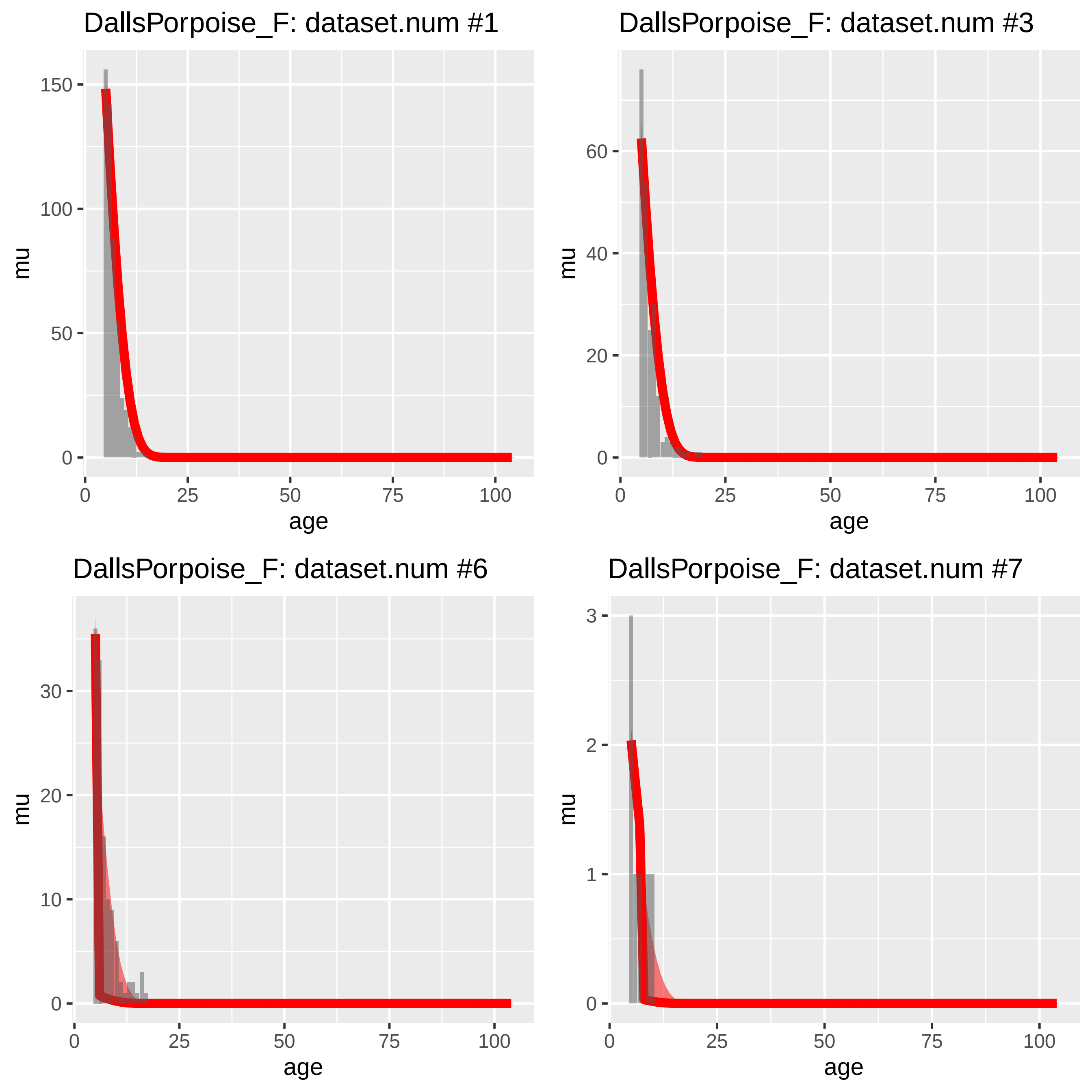
DallsPorpoise_F: dataset.num #1
DallsPorpoise_F: dataset.num #3
DallsPorpoise_F: dataset.num #6
DallsPorpoise_F: dataset.num #7
mu
age
0
25
50
75
100
150
100
50
60
40
20
30
10
3
2
1

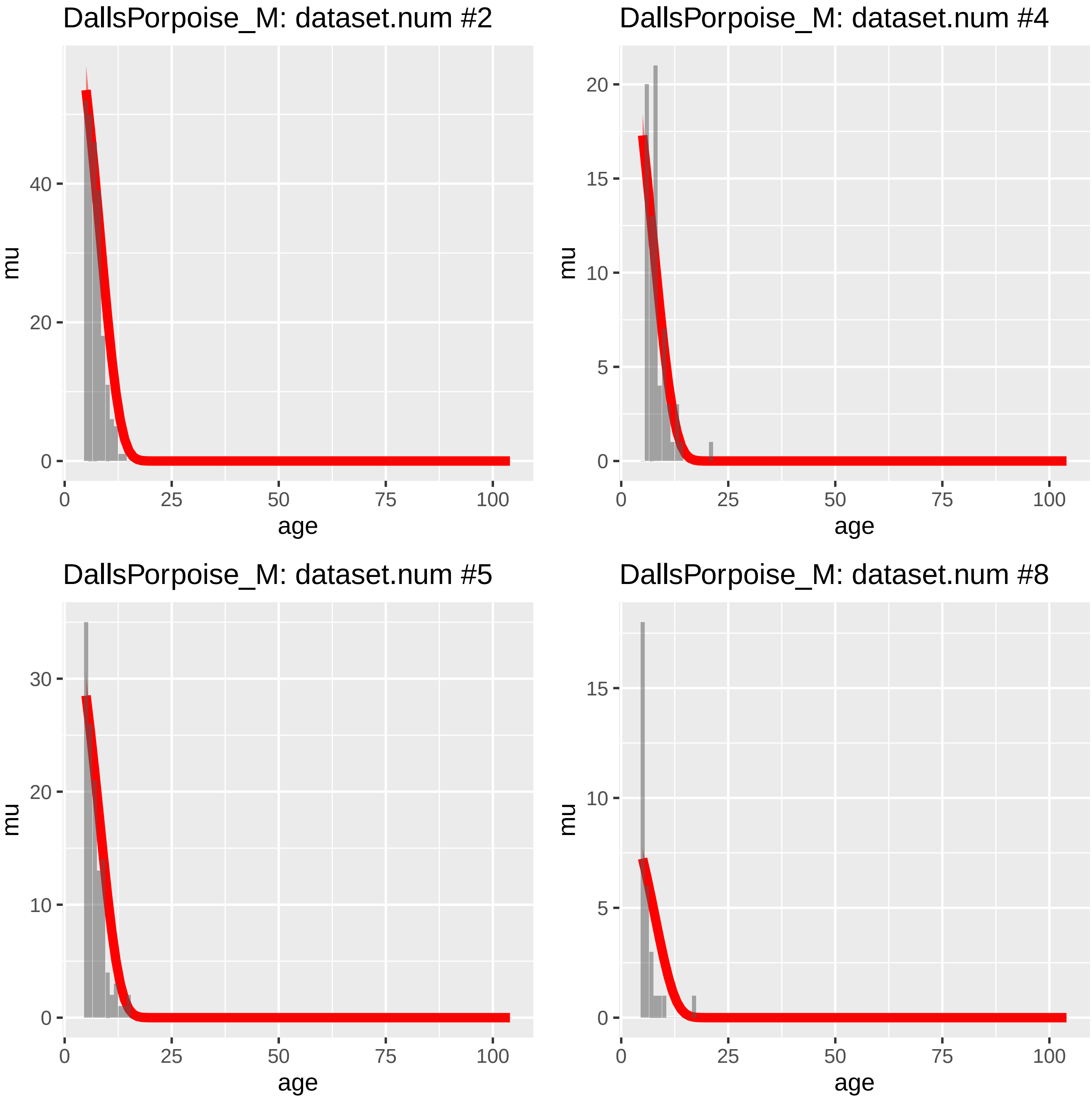
DallsPorpoise_M: dataset.num #2
mu
age
0
20
40
0
25
50
75
100
DallsPorpoise_M: dataset.num #4
mu
age
0
5
10
15
20
0
25
50
75
100
DallsPorpoise_M: dataset.num #5
mu
age
0
10
20
30
0
25
50
75
100
DallsPorpoise_M: dataset.num #8
mu
age
0
5
10
15
0
25
50
75
100

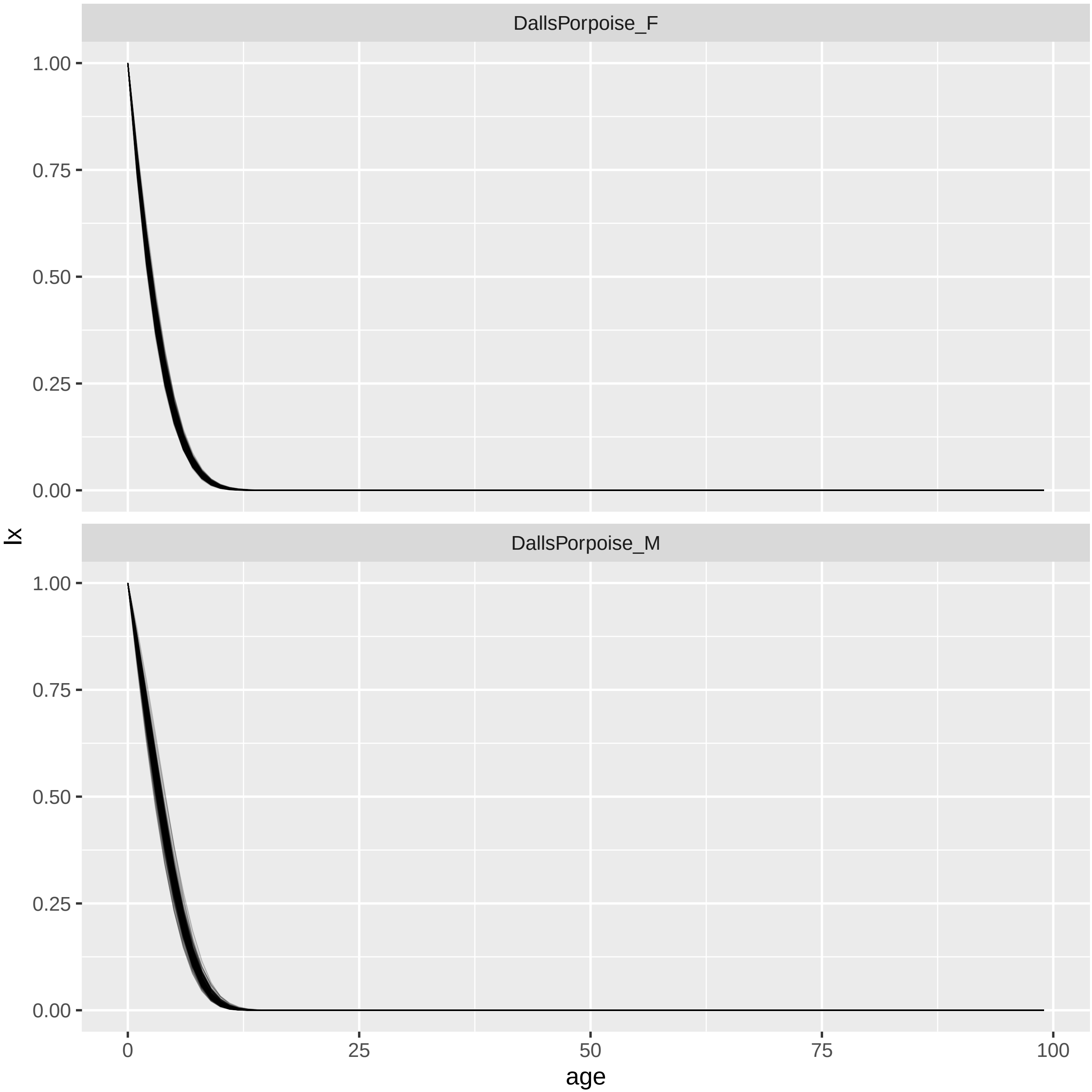
DallsPorpoise_F
DallsPorpoise_M
1.00
0.75
0.50
0.25
0.00
0
25
50
75
100
lx
age

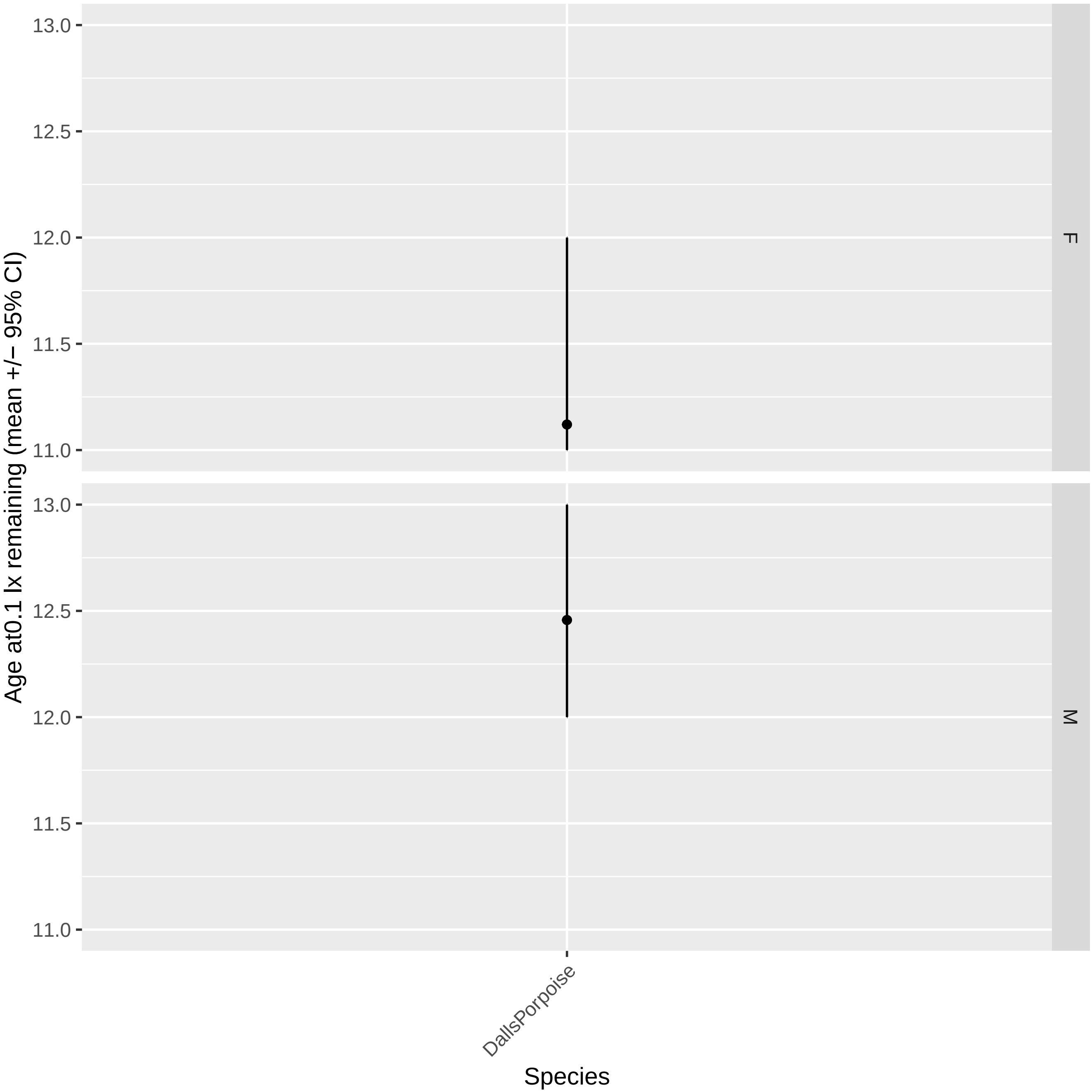

13.0
12.5
12.0
11.5
11.0
F
13.0
12.5
12.0
11.5
11.0
M
Age at0.1 lx remaining (mean +/− 95% CI)
DallsPorpoise
Species

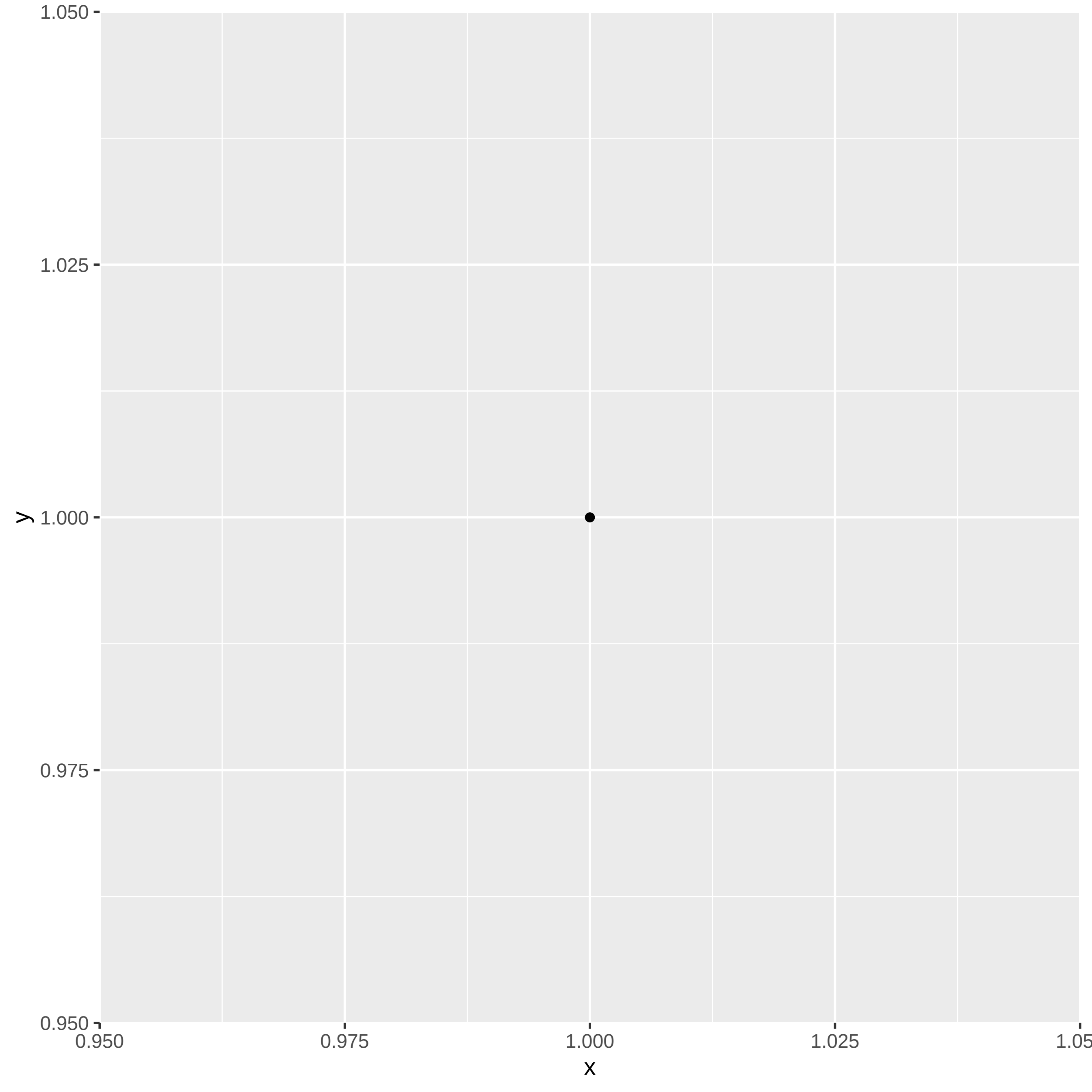
1.050
1.025
1.000
0.975
0.950
y
0.950
0.975
1.000
1.025
1.05
x

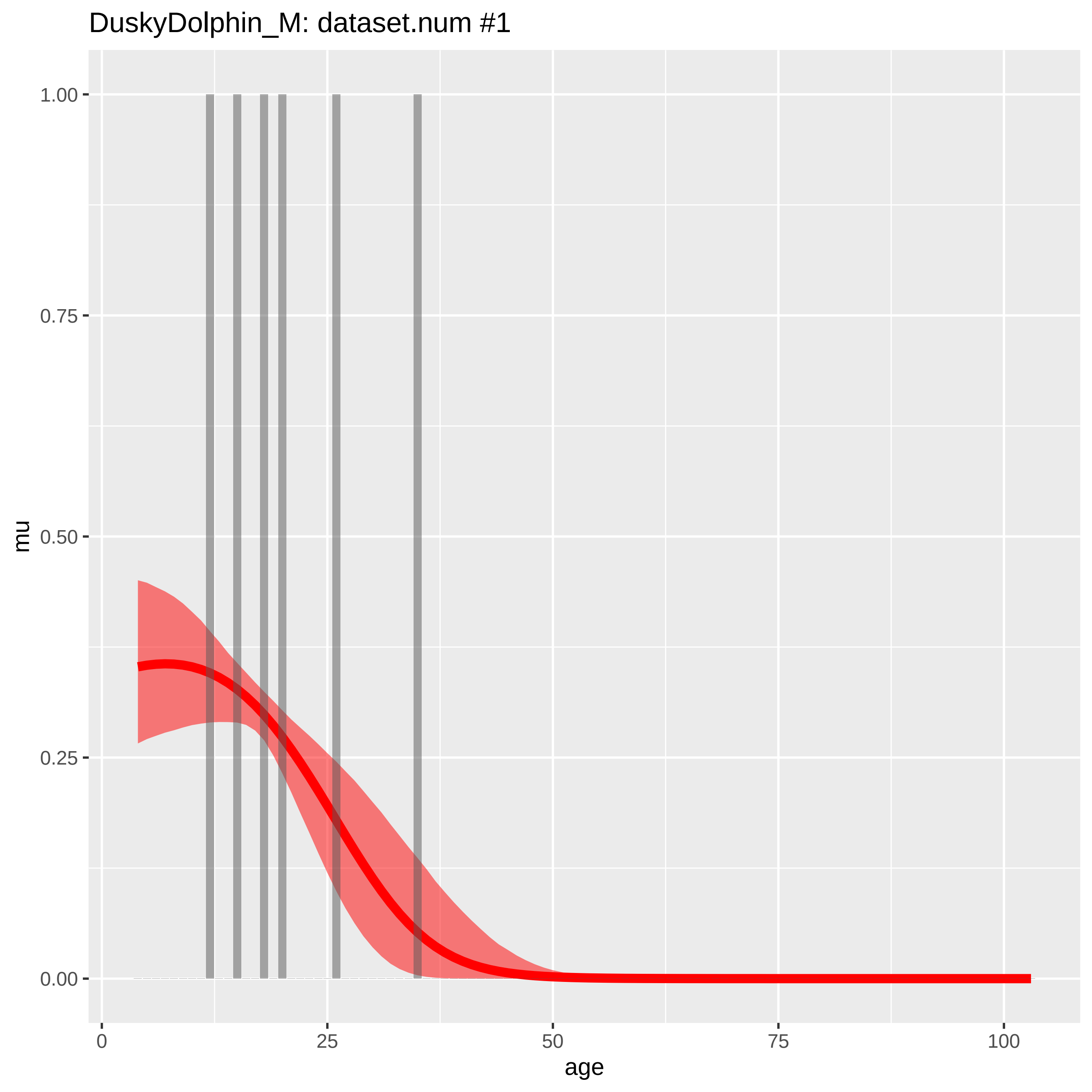
DuskyDolphin_M: dataset.num #1
1.00
0.75
0.50
0.25
0.00
mu
0
25
50
75
100
age

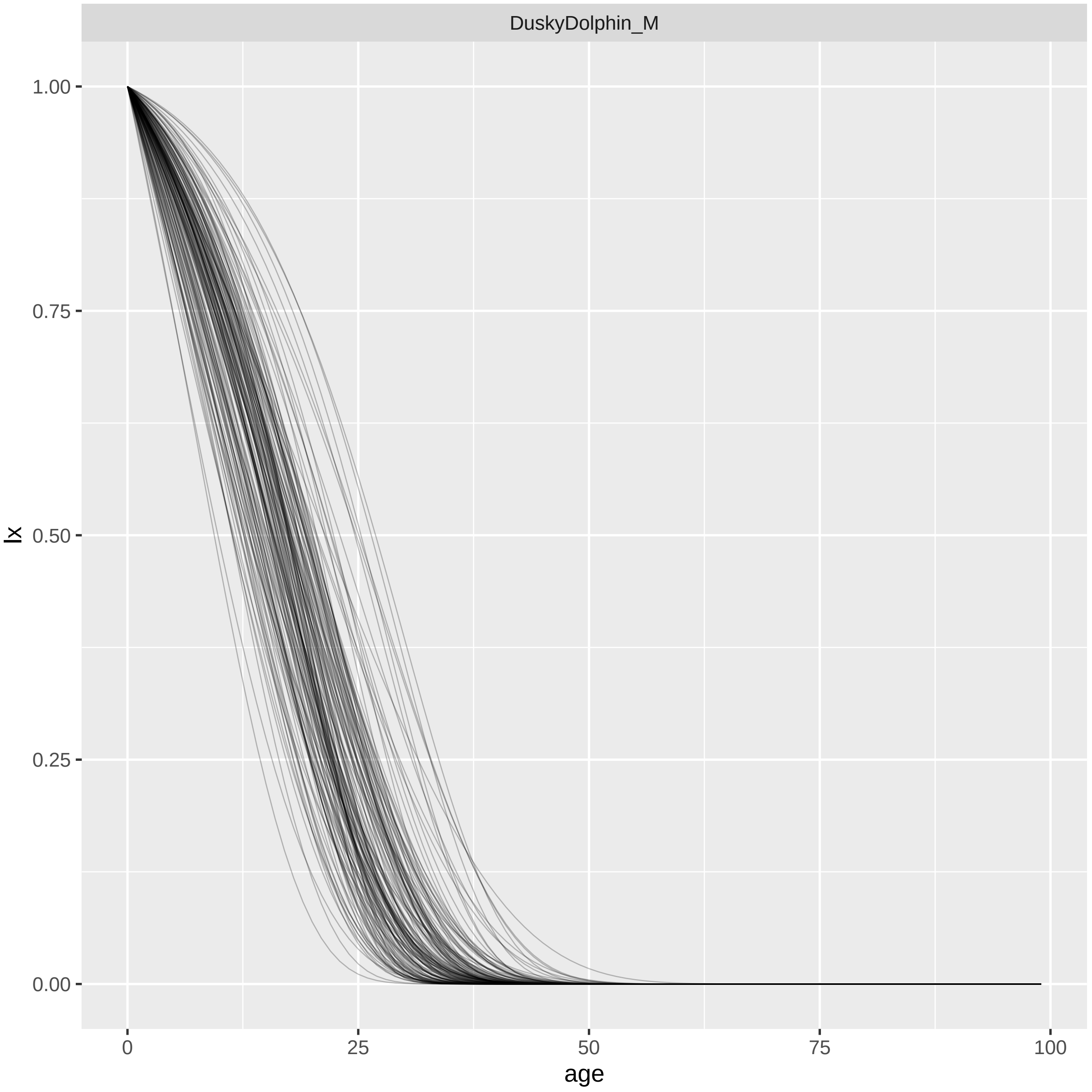
DuskyDolphin_M
1.00
0.75
0.50
0.25
0.00
lx
0
25
50
75
100
age

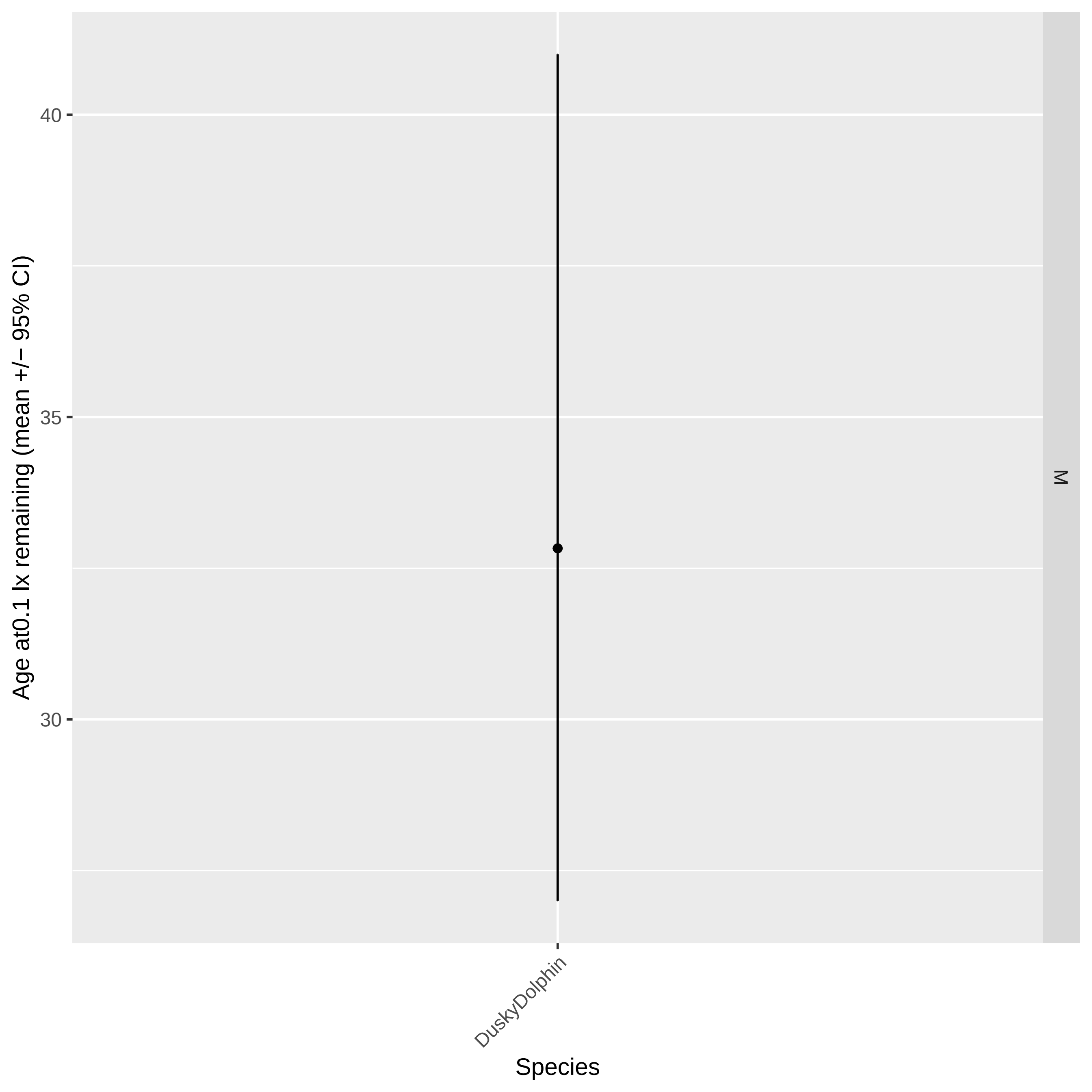

Age at0.1 lx remaining (mean +/− 95% CI)
40
35
30
M
DuskyDolphin
Species

DwarfSpermWhale_F: dataset.num #1

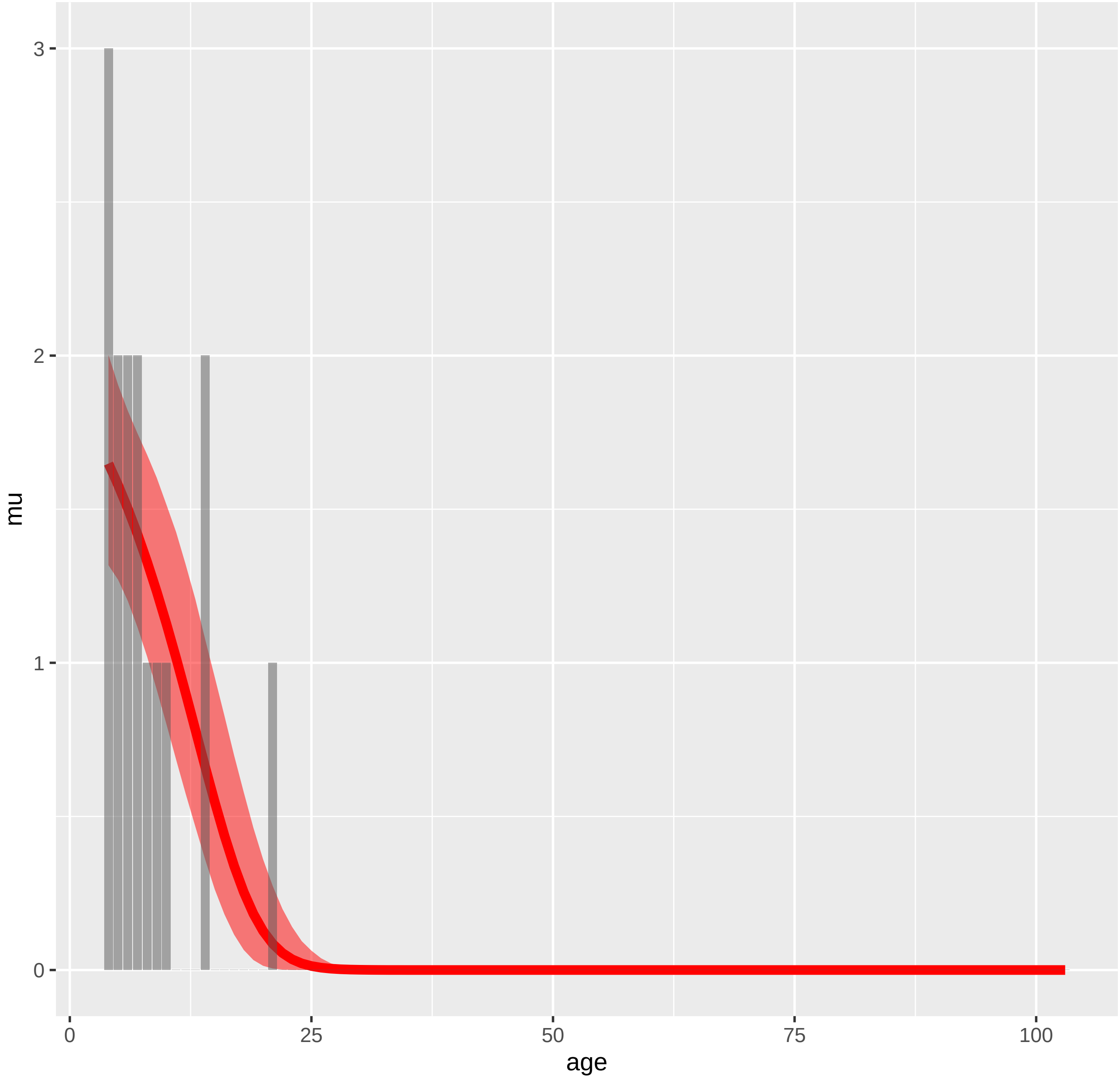

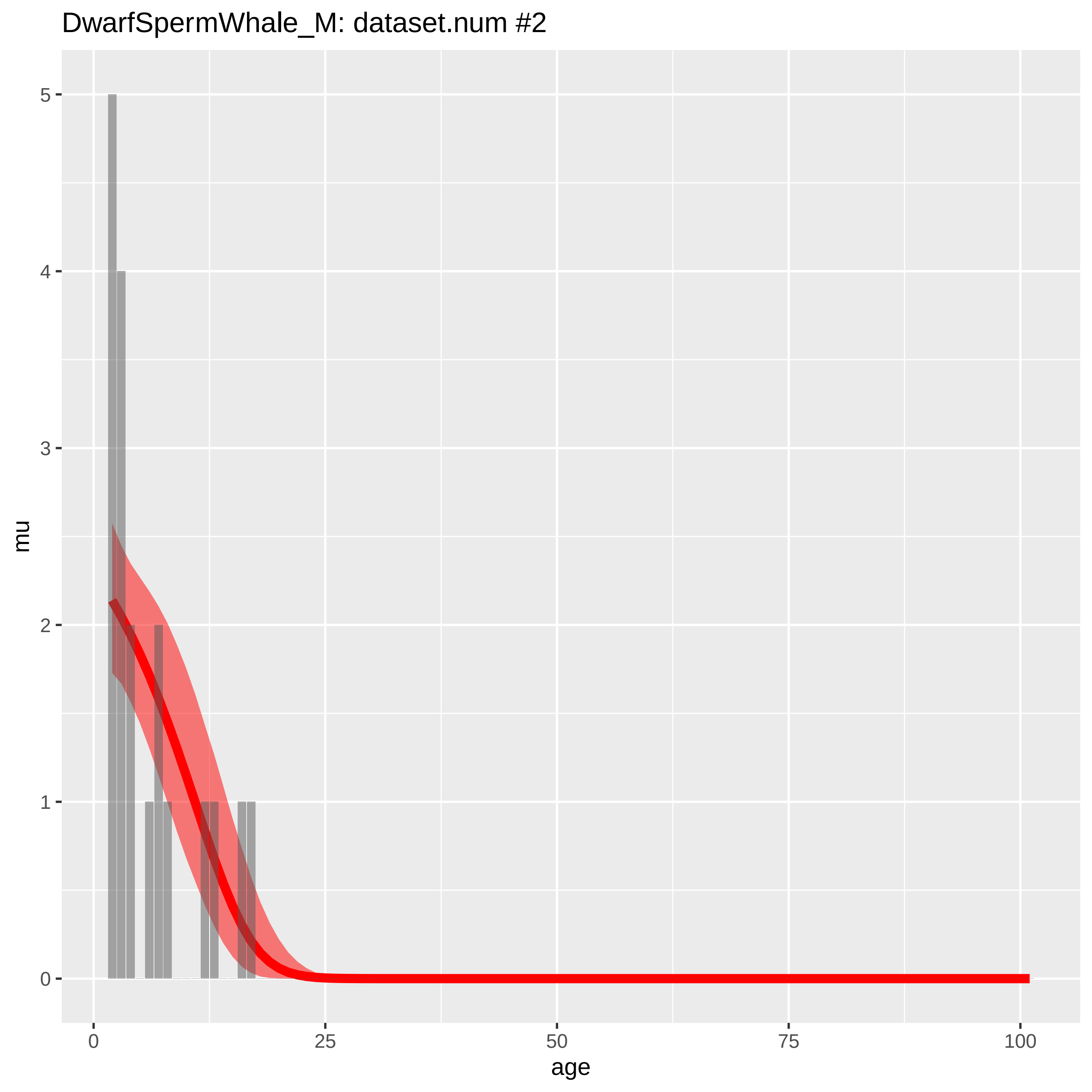

DwarfSpermWhale_M: dataset.num #2
mu
age
0
1
2
3
4
5
0
25
50
75
100

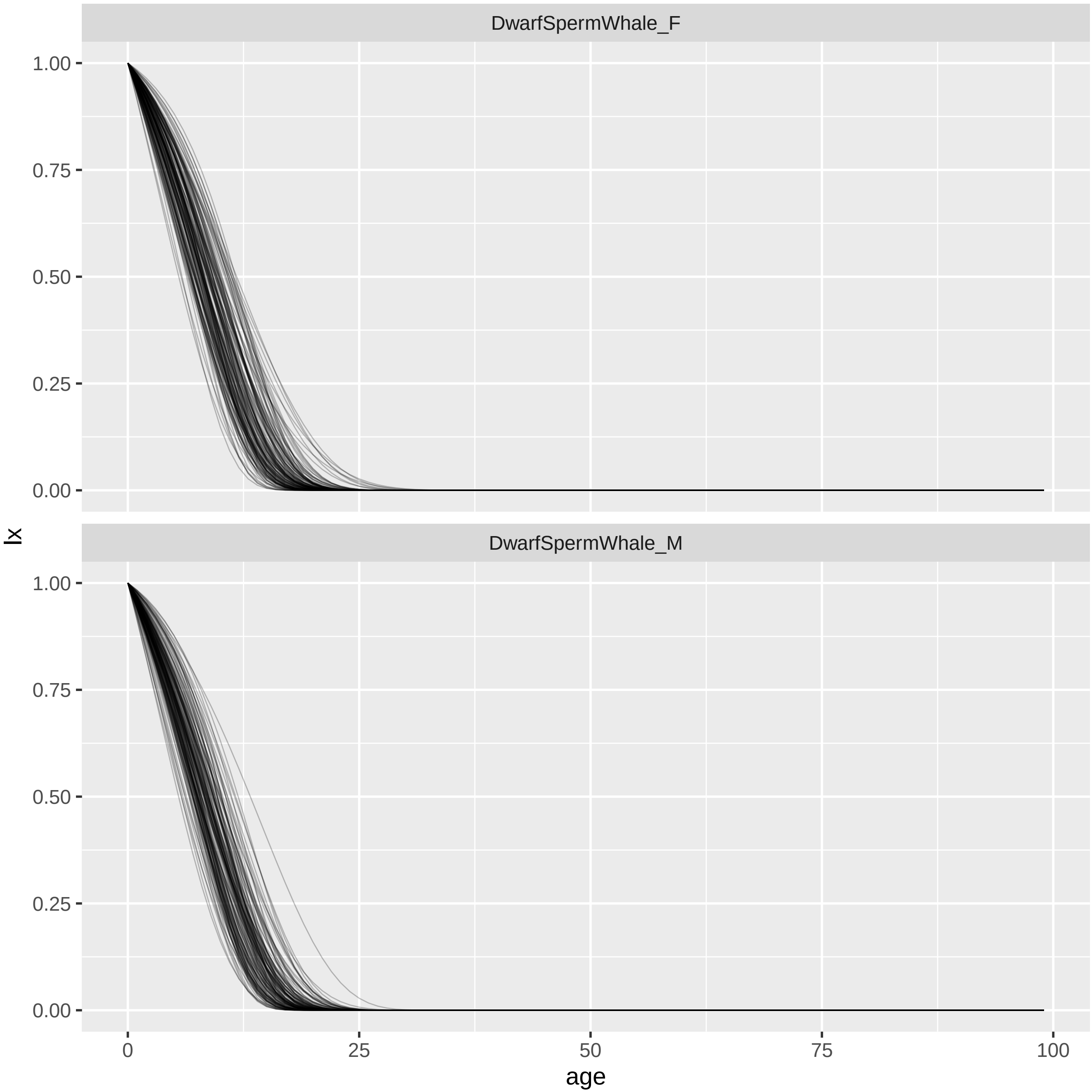

DwarfSpermWhale_F
DwarfSpermWhale_M
lx
1.00
0.75
0.50
0.25
0.00
0
25
50
75
100
age

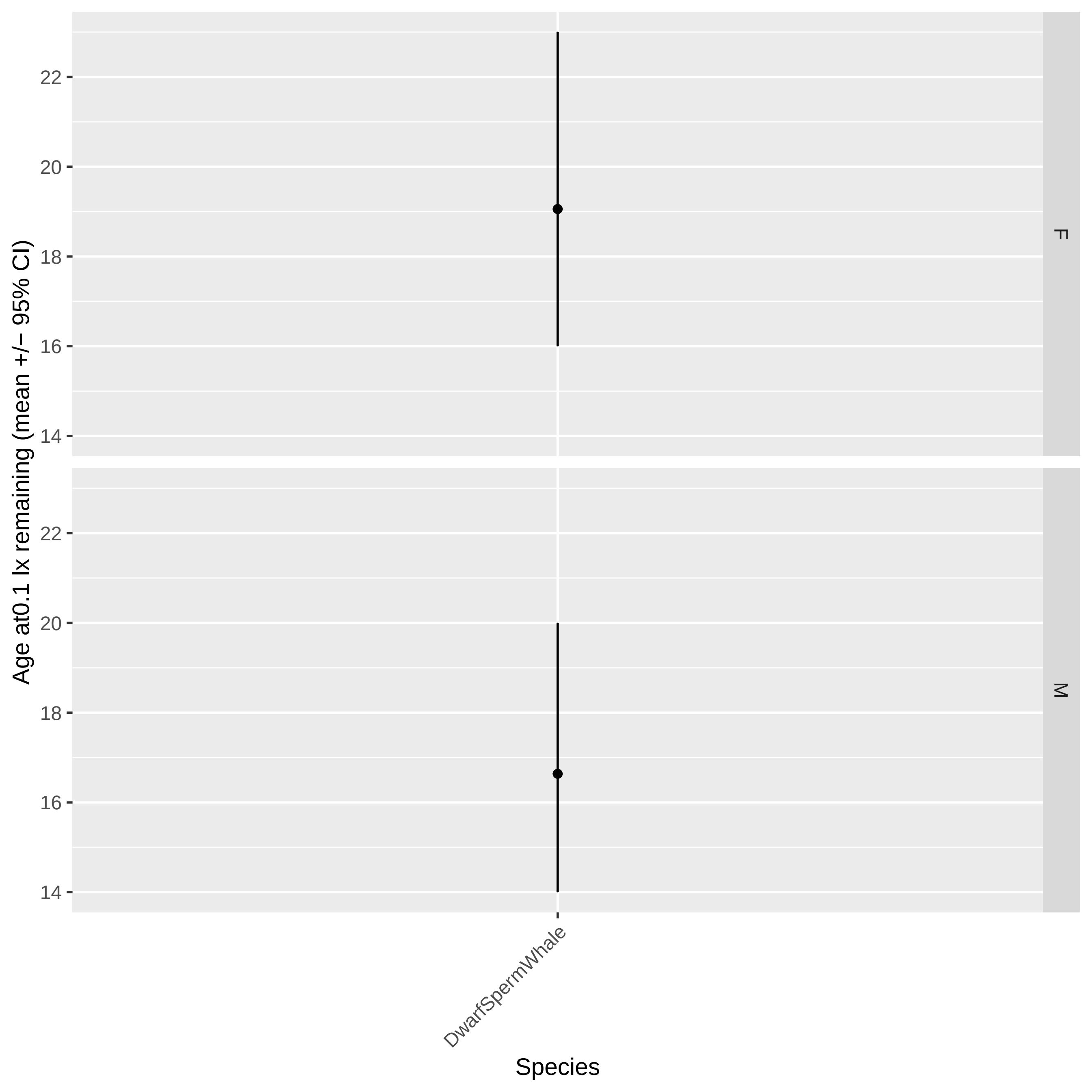

Age at0.1 lx remaining (mean +/− 95% CI)
22
20
18
16
14
F
22
20
18
16
14
M
DwarfSpermWhale
Species

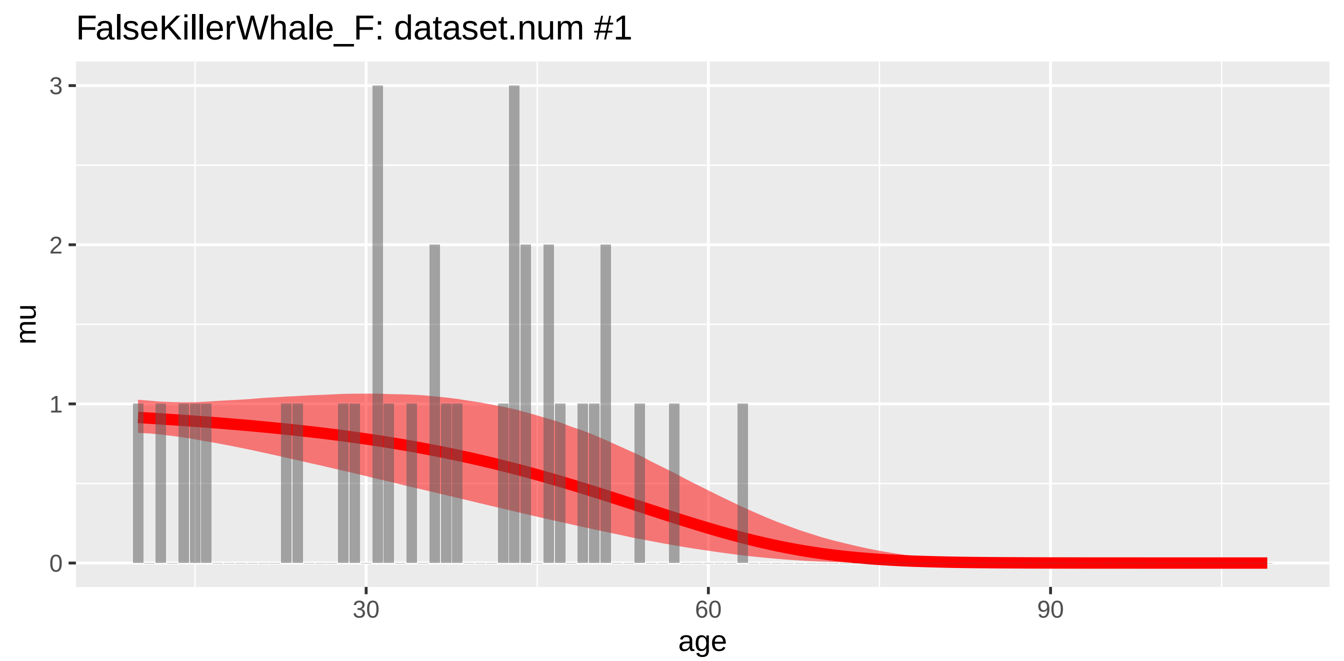

FalseKillerWhale_F: dataset.num #1
mu
3
2
1
0
30
60
90
age


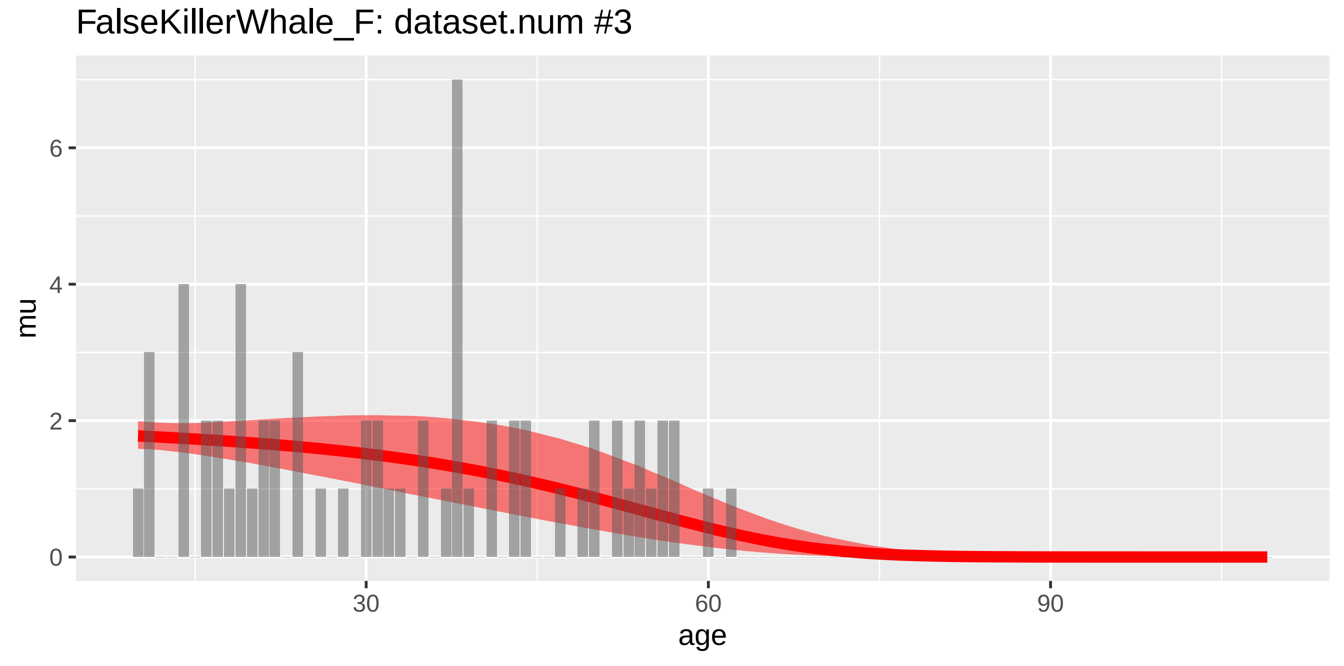

FalseKillerWhale_F: dataset.num #3
mu
6
4
2
0
30
60
90
age

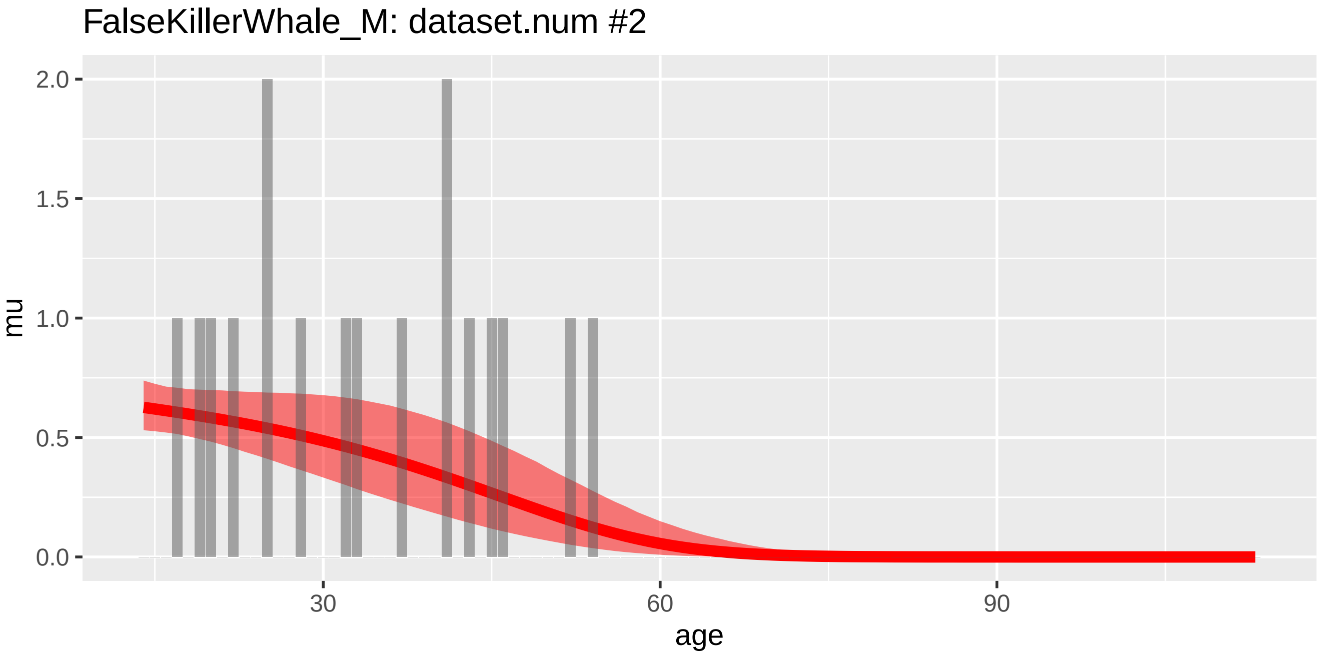
FalseKillerWhale_M: dataset.num #2
mu
2.0
1.5
1.0
0.5
0.0
30
60
90
age

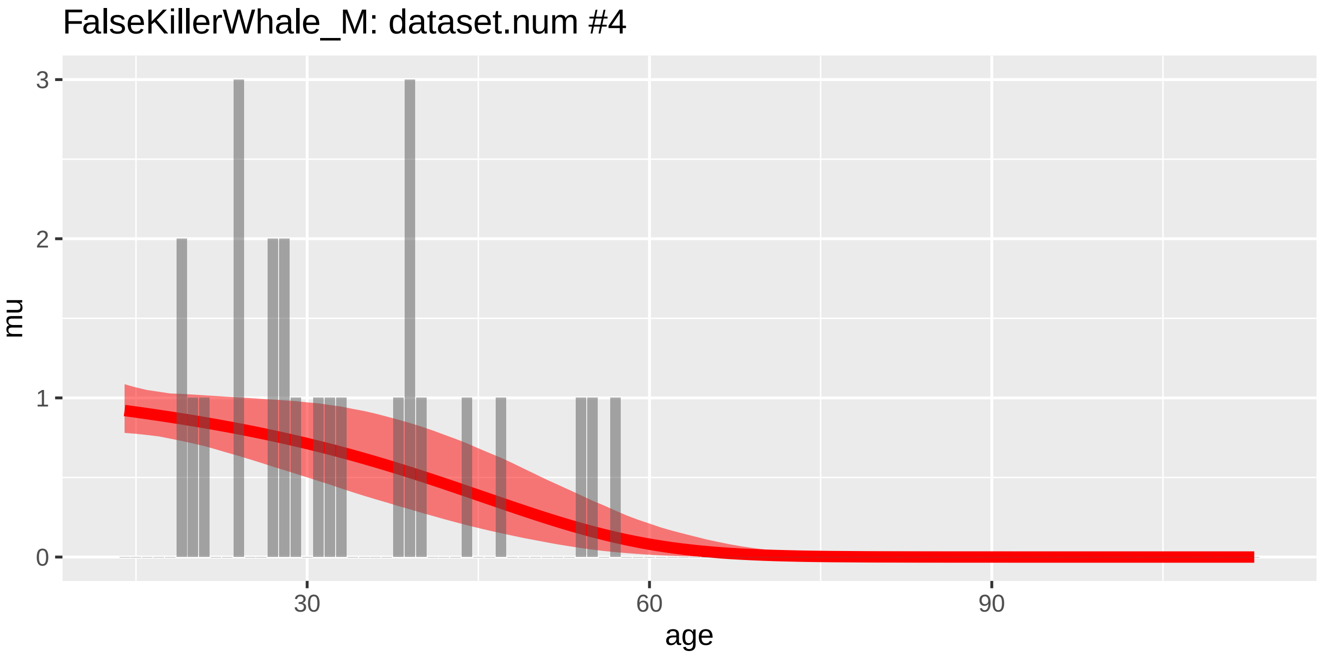
FalseKillerWhale_M: dataset.num #4
mu
3
2
1
0
30
60
90
age

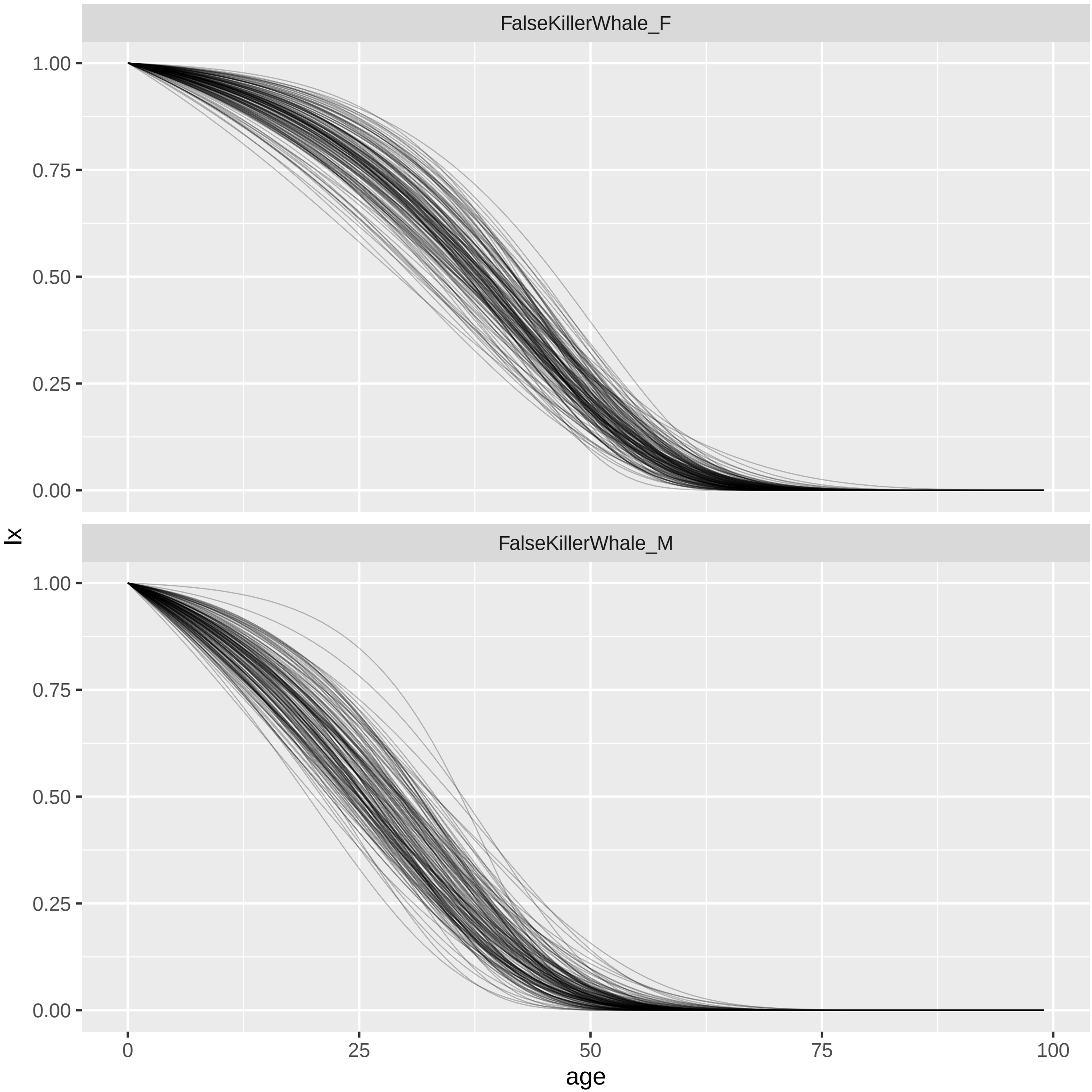
FalseKillerWhale_F
FalseKillerWhale_M
1.00
0.75
0.50
0.25
0.00
lx
0
25
50
75
100
age

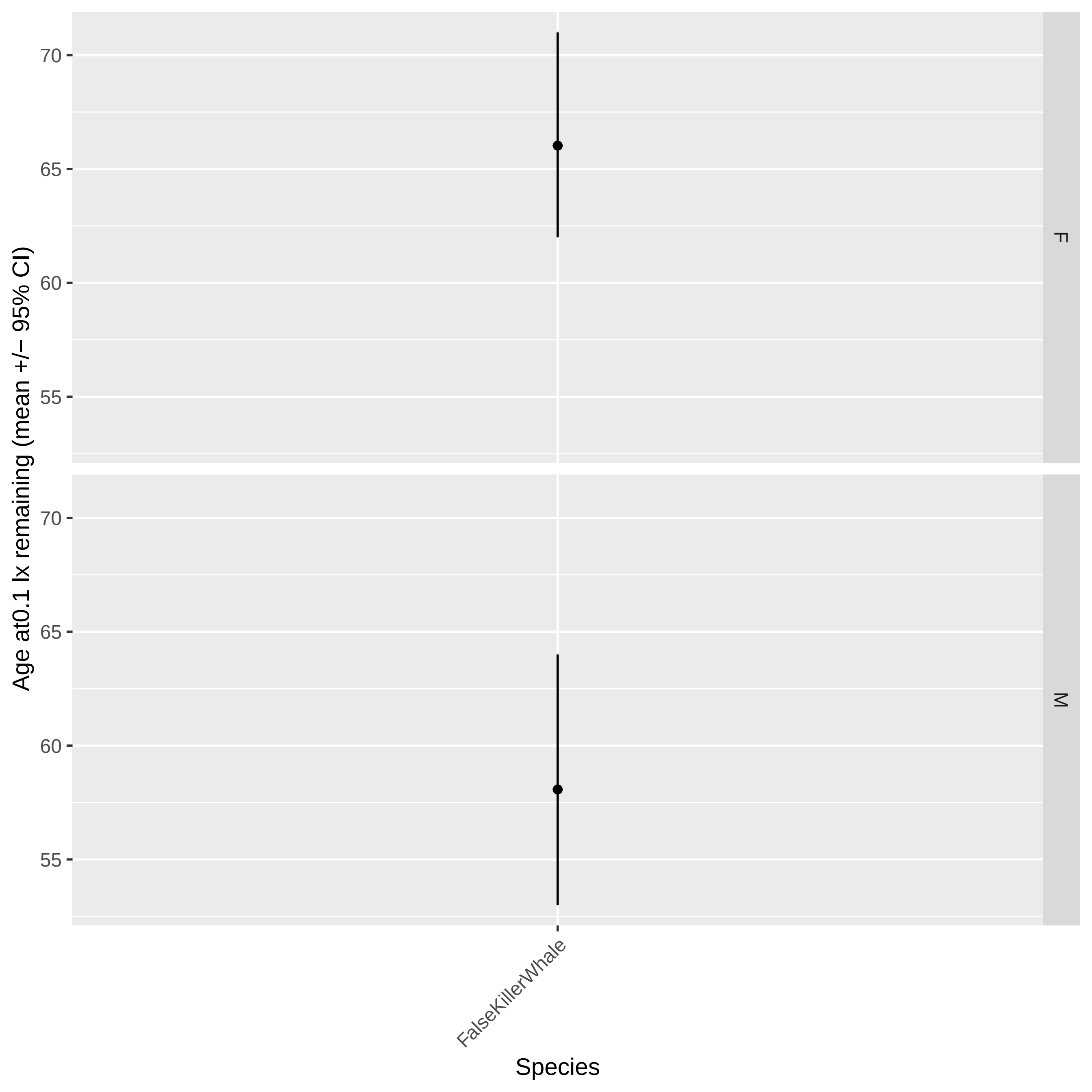

Age at0.1 lx remaining (mean +/− 95% CI)
70
65
60
55
F
M
FalseKillerWhale
Species

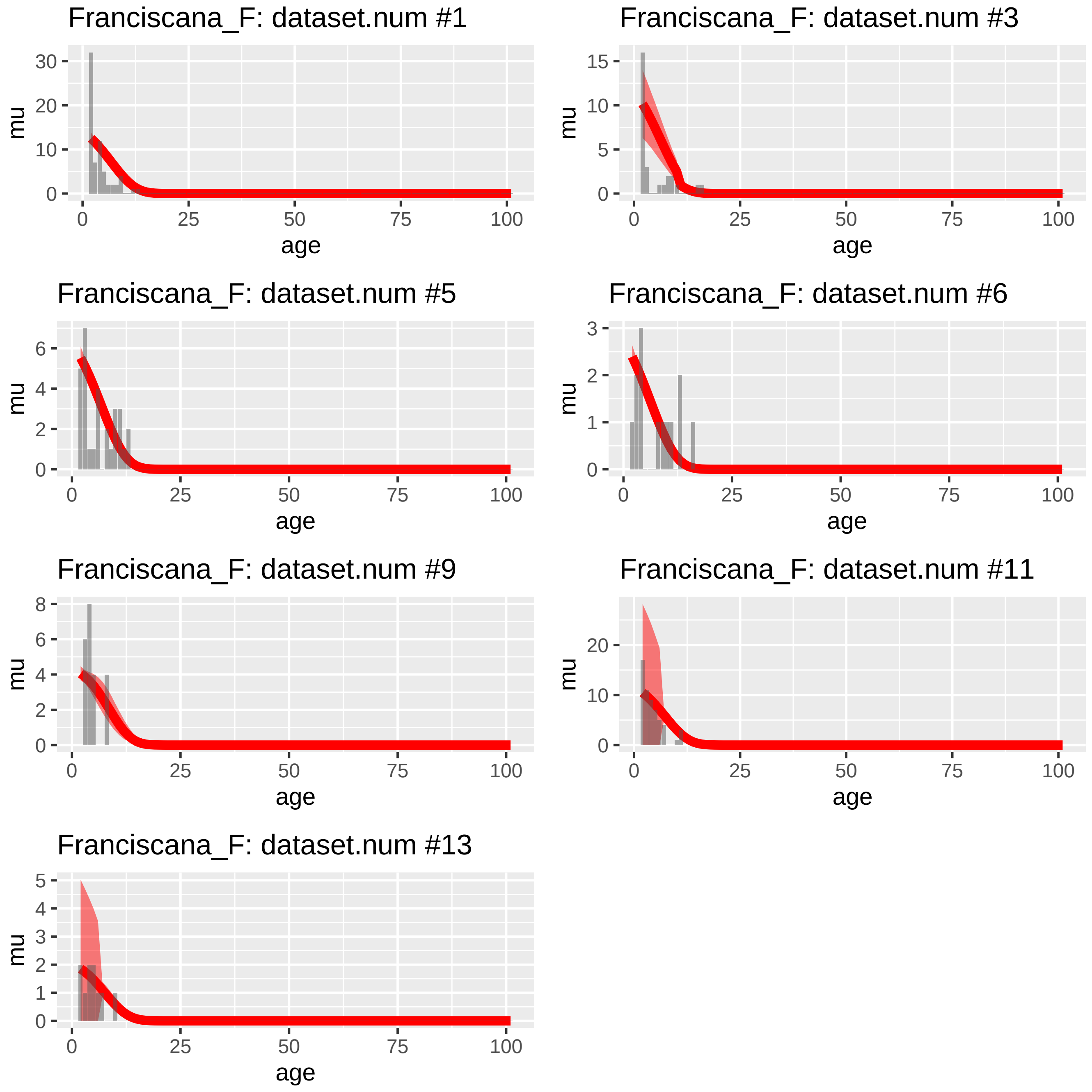
Franciscana_F: dataset.num #1
Franciscana_F: dataset.num #3
Franciscana_F: dataset.num #5
Franciscana_F: dataset.num #6
Franciscana_F: dataset.num #9
Franciscana_F: dataset.num #11
Franciscana_F: dataset.num #13
mu
age
0
25
50
75
100

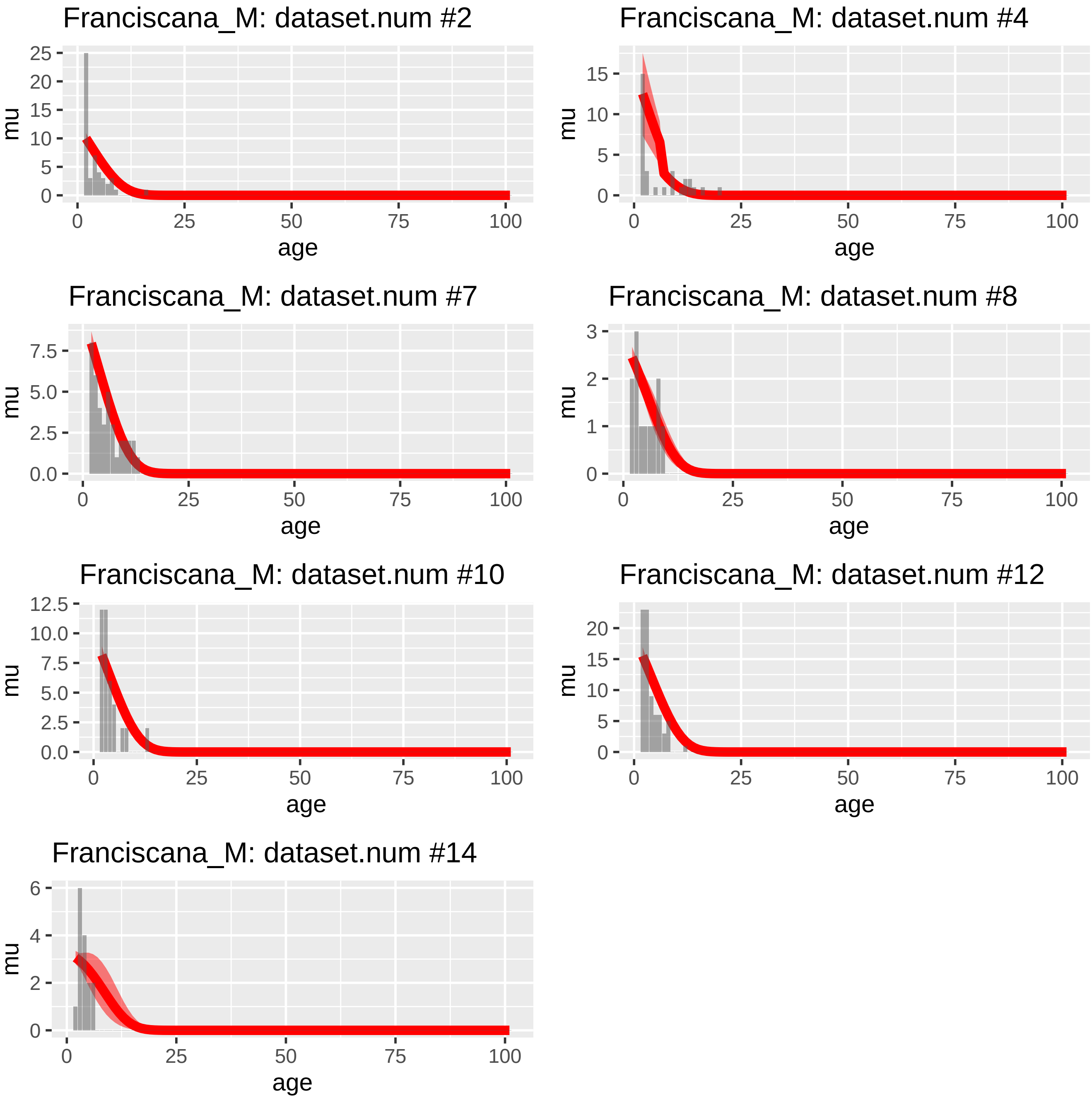
Franciscana_M: dataset.num #2
Franciscana_M: dataset.num #4
Franciscana_M: dataset.num #7
Franciscana_M: dataset.num #8
Franciscana_M: dataset.num #10
Franciscana_M: dataset.num #12
Franciscana_M: dataset.num #14
mu
age
0
25
50
75
100

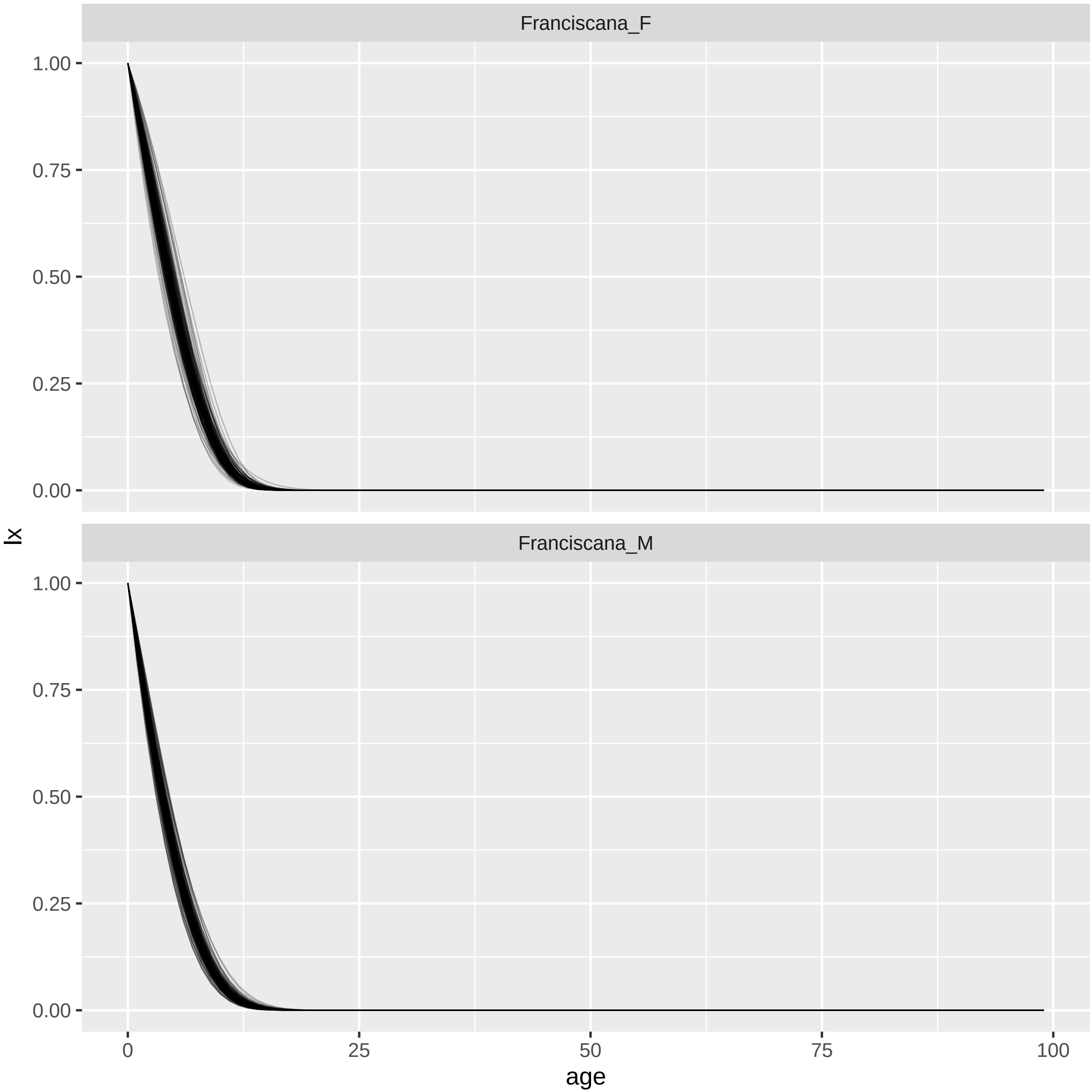

Franciscana_F
Franciscana_M
lx
1.00
0.75
0.50
0.25
0.00
0
25
50
75
100
age

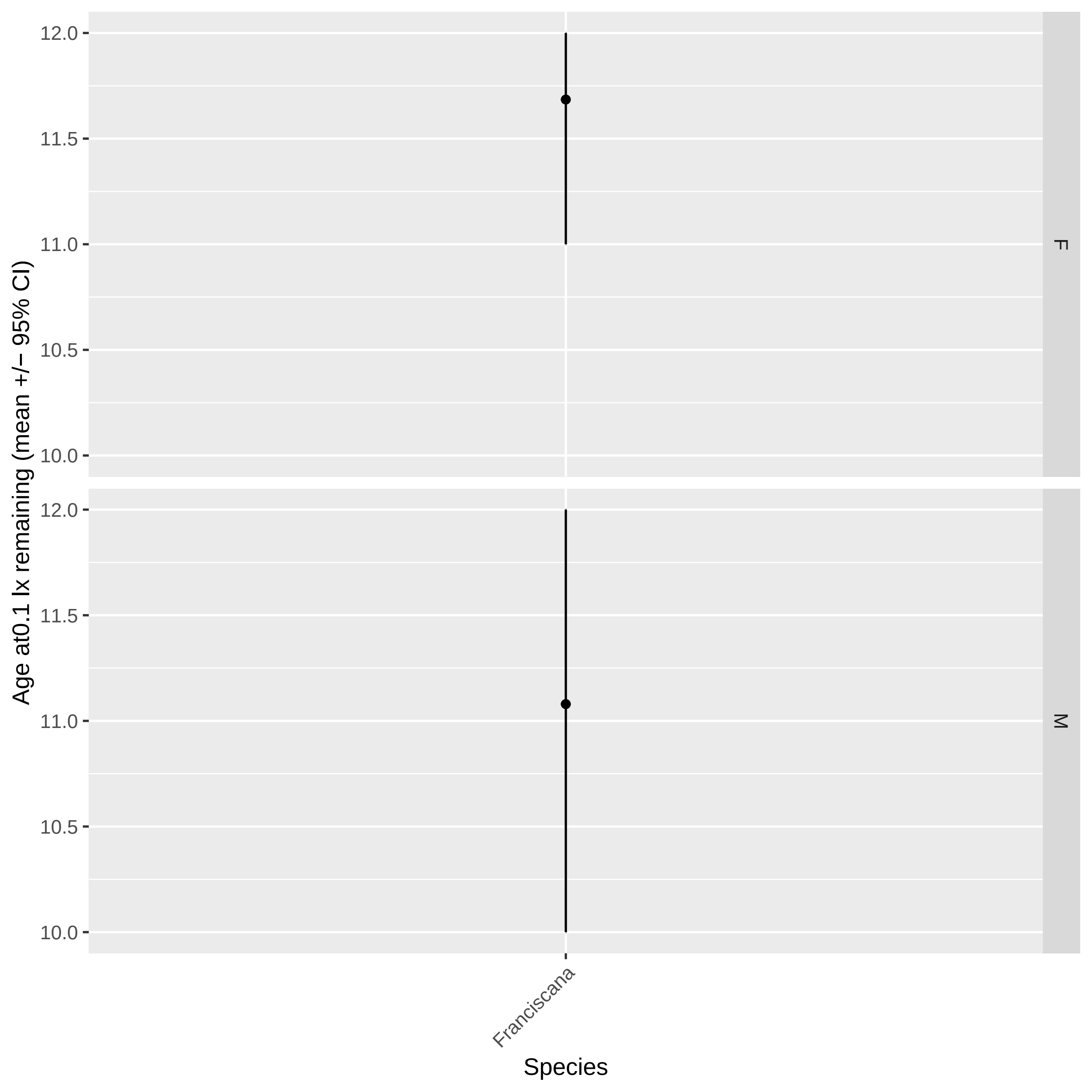

Age at0.1 lx remaining (mean +/− 95% CI)
12.0
11.5
11.0
10.5
10.0
F
M
Franciscana
Species

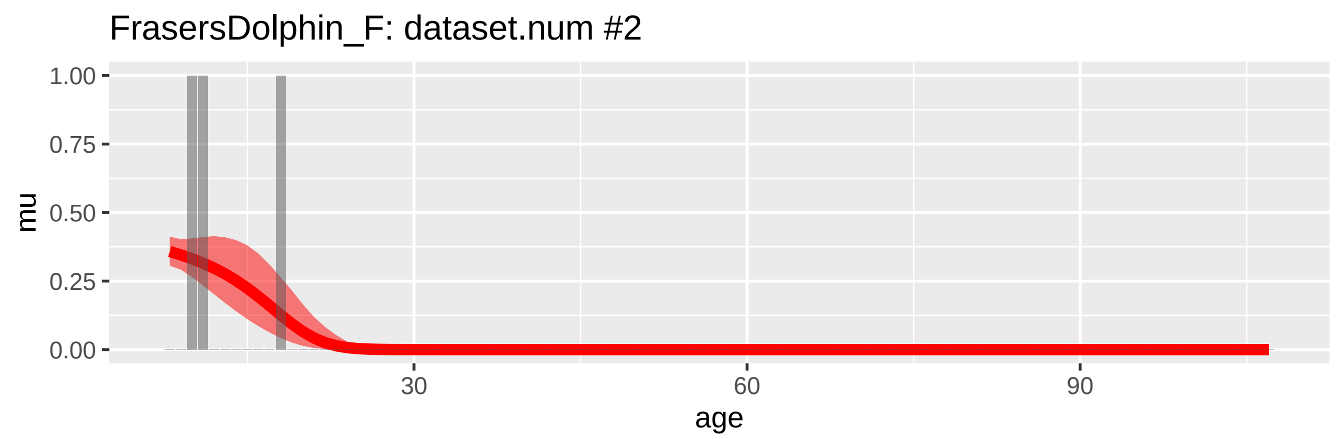

FrasersDolphin_F: dataset.num #2
mu
1.00
0.75
0.50
0.25
0.00
30
60
90
age


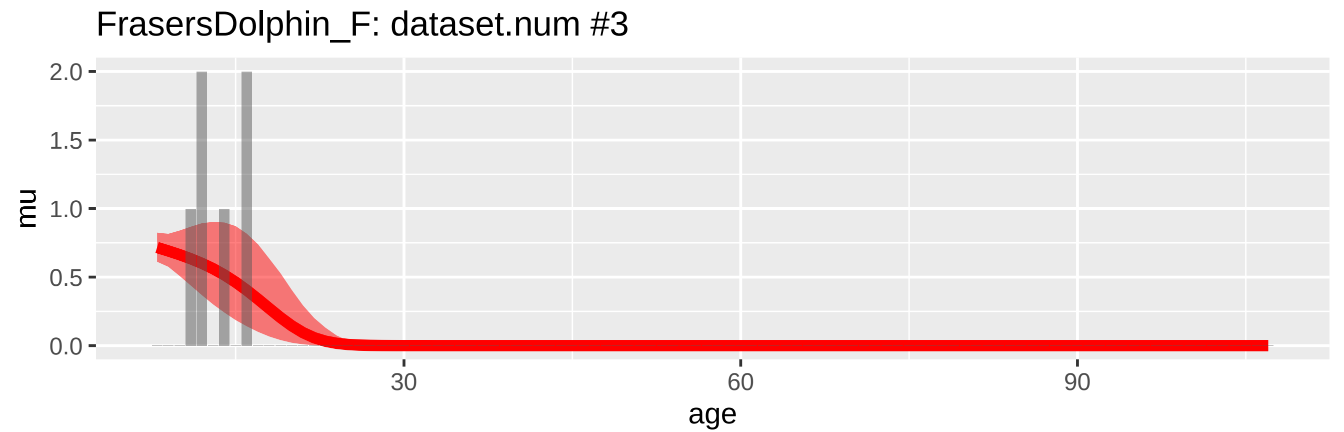

FrasersDolphin_F: dataset.num #3
mu
2.0
1.5
1.0
0.5
0.0
30
60
90
age


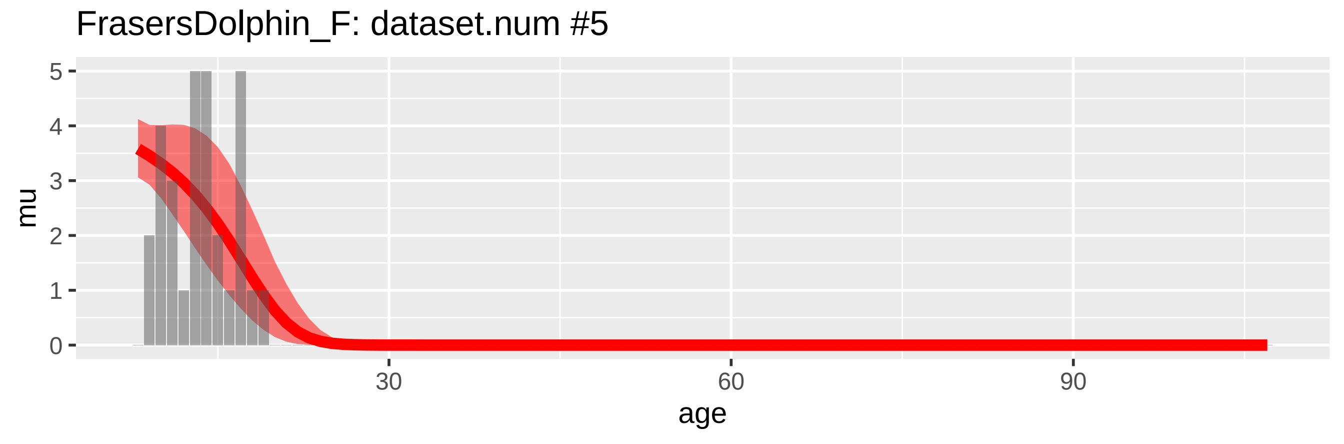

FrasersDolphin_F: dataset.num #5
mu
5
4
3
2
1
0
30
60
90
age

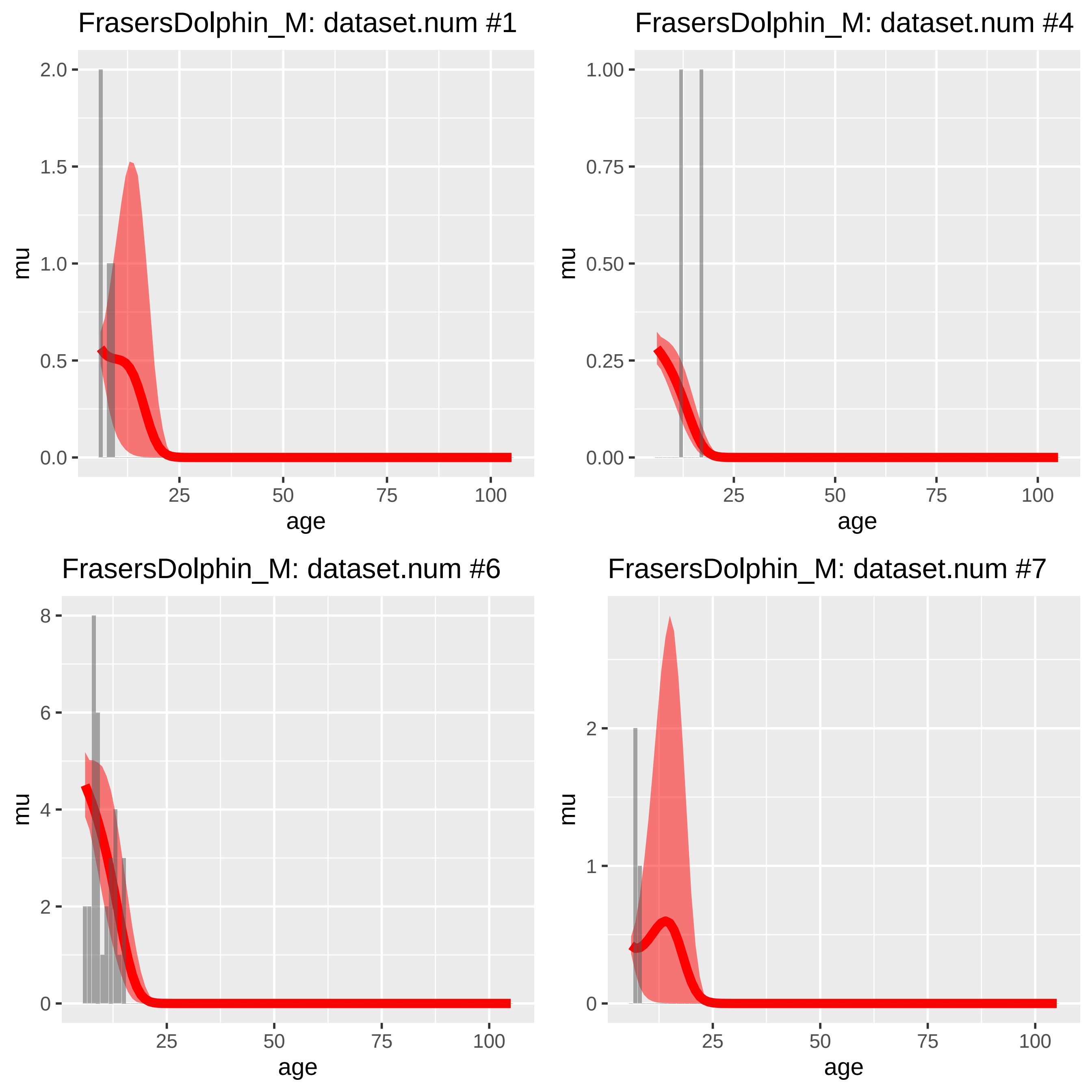
FrasersDolphin_M: dataset.num #1
FrasersDolphin_M: dataset.num #4
FrasersDolphin_M: dataset.num #6
FrasersDolphin_M: dataset.num #7
mu
age
0.0
0.5
1.0
1.5
2.0
0.00
0.25
0.50
0.75
1.00
0
2
4
6
8
1
25
50
75
100

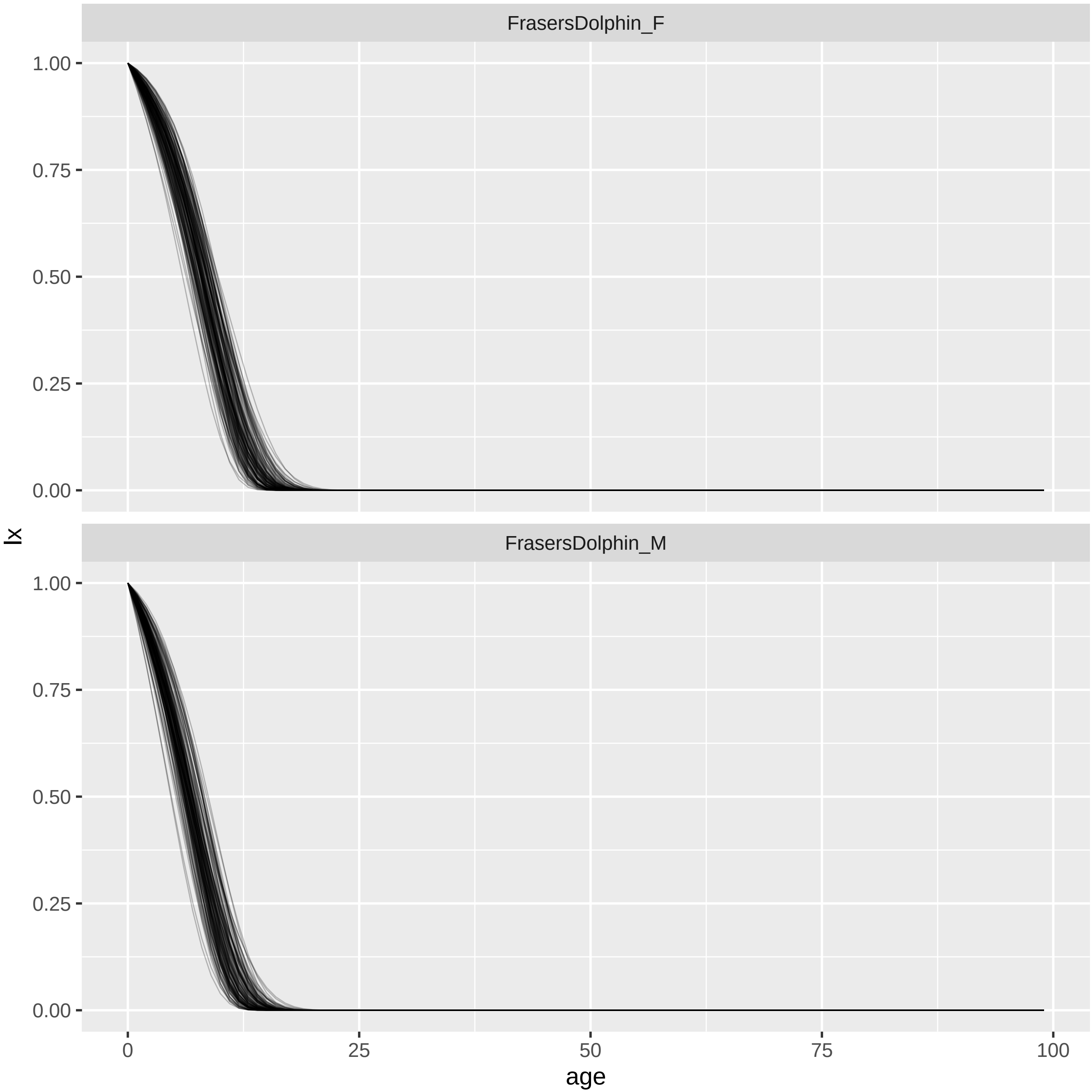

FrasersDolphin_F
FrasersDolphin_M
1.00
0.75
0.50
0.25
0.00
lx
0
25
50
75
100
age

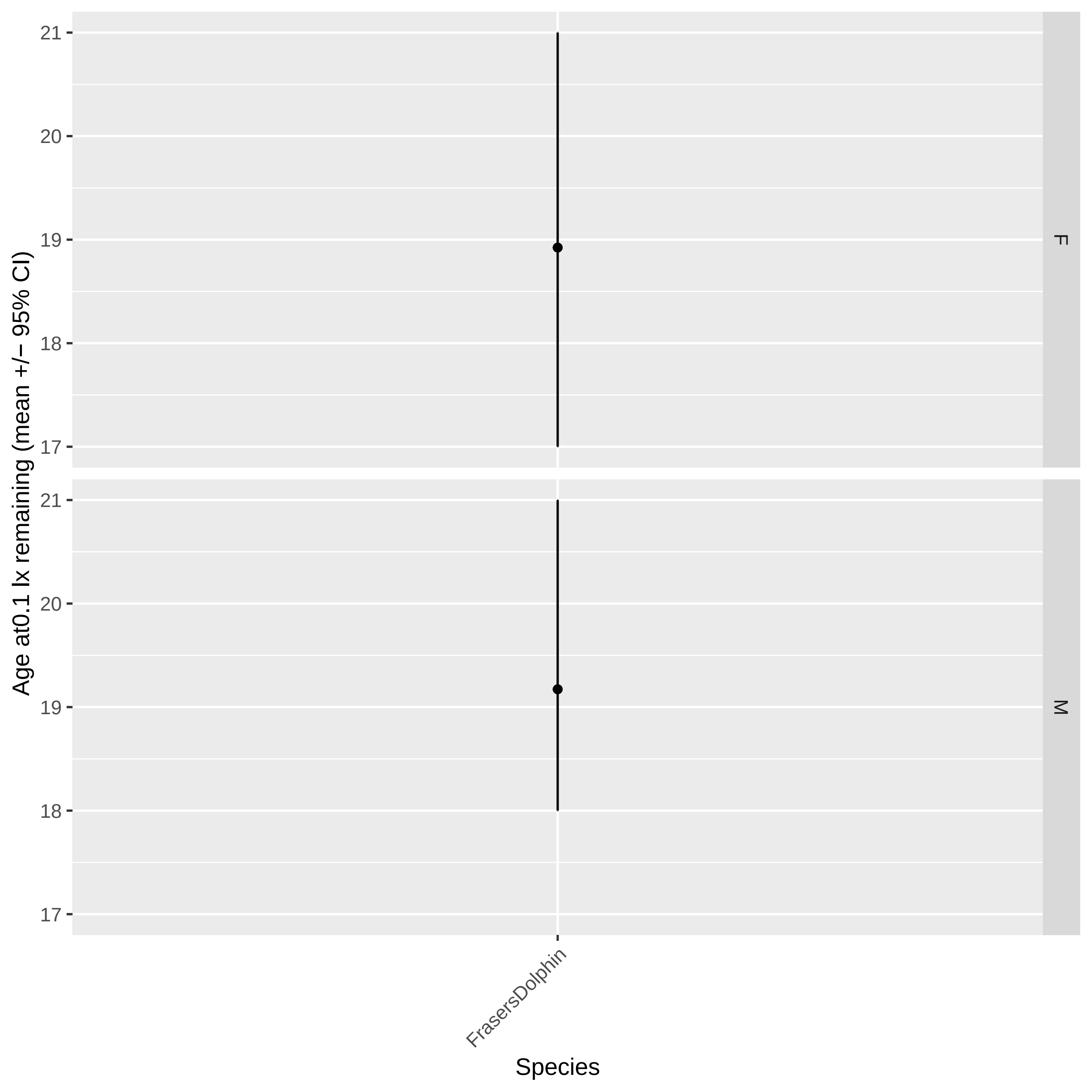
Age at0.1 lx remaining (mean +/− 95% CI)
21
20
19
18
17
F
M
FrasersDolphin
Species

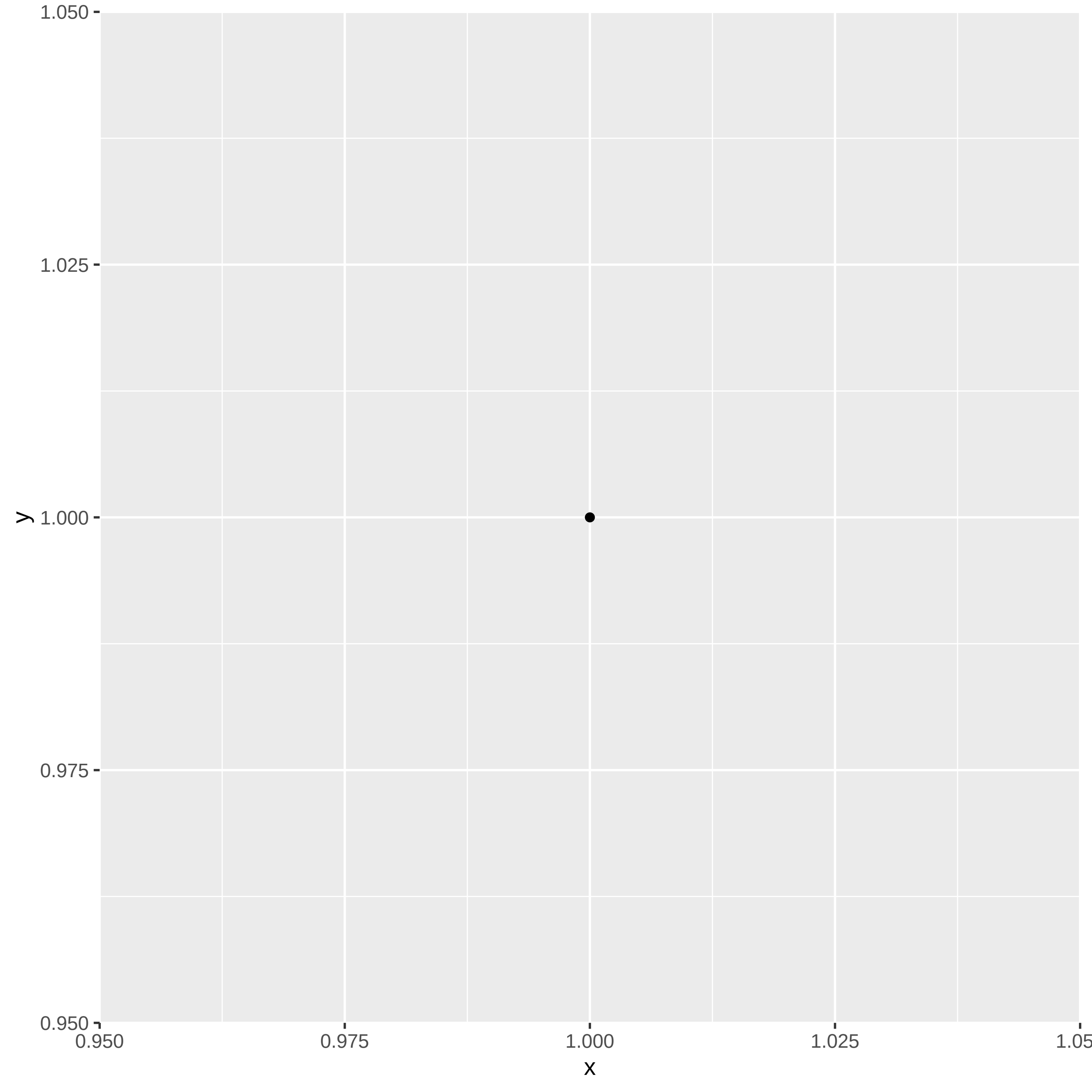
1.050
1.025
1.000
0.975
0.950
y
0.950
0.975
1.000
1.025
1.05
x

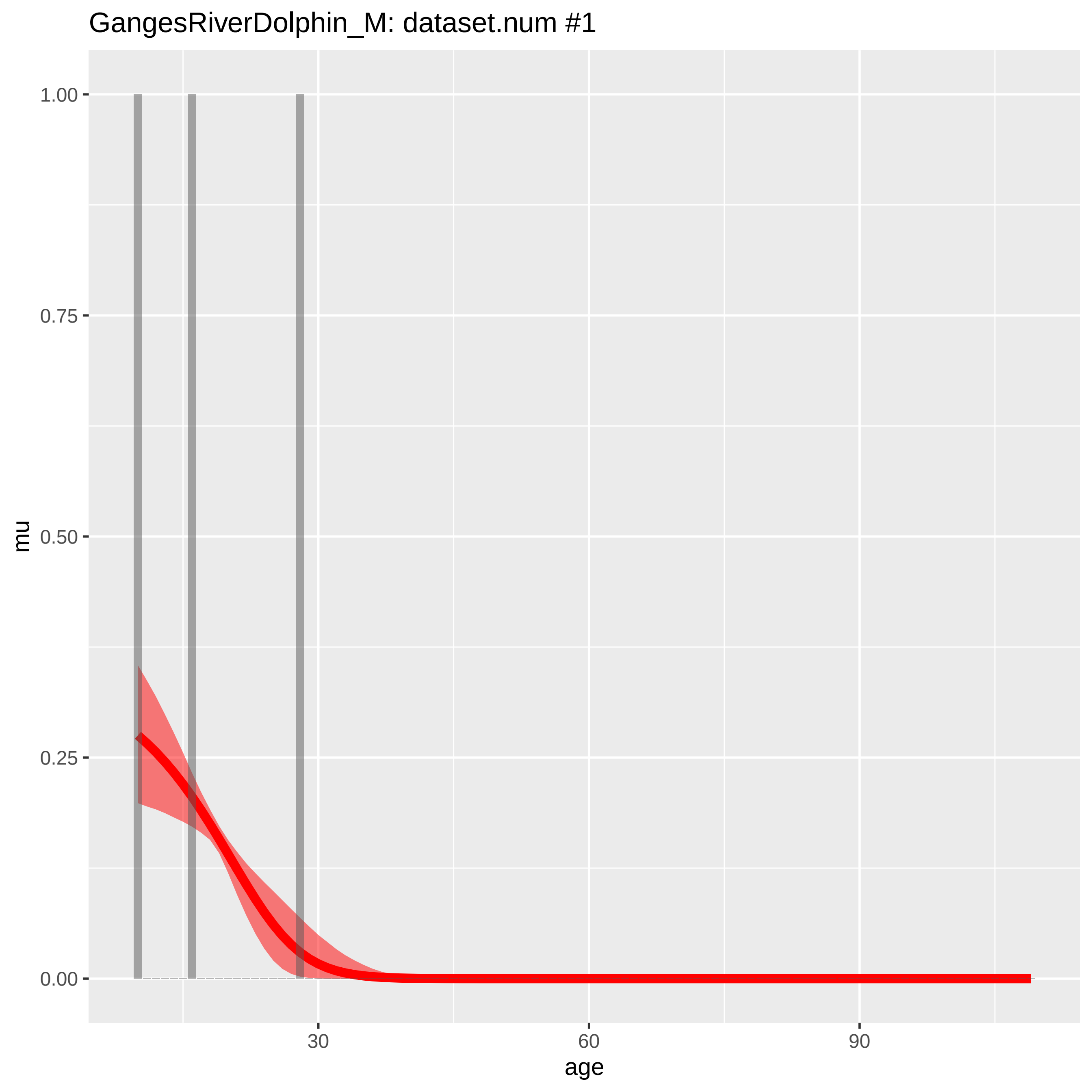

GangesRiverDolphin_M: dataset.num #1
1.00
0.75
0.50
0.25
0.00
mu
30
60
90
age

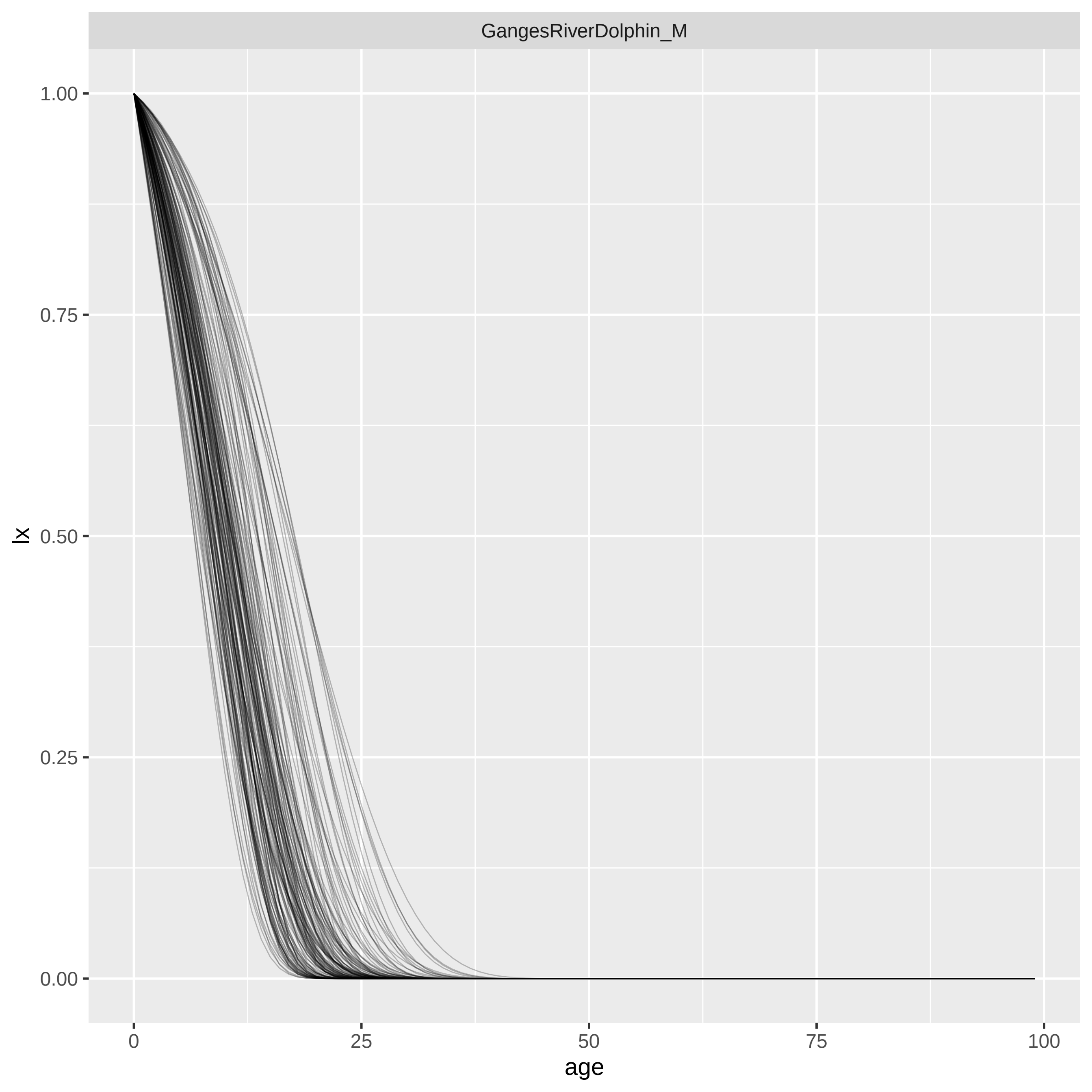
GangesRiverDolphin_M
lx
1.00
0.75
0.50
0.25
0.00
0
25
50
75
100
age

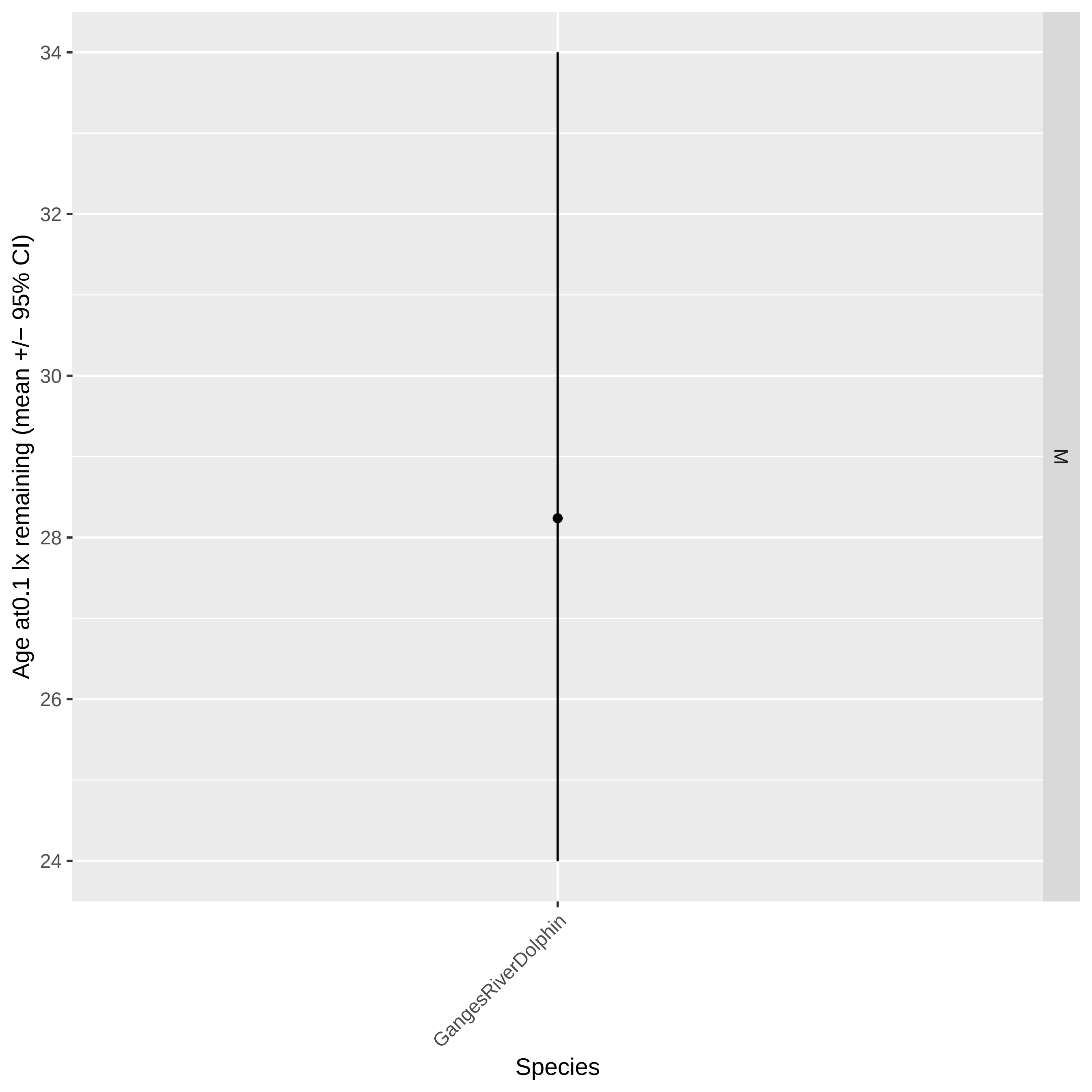

Age at0.1 lx remaining (mean +/− 95% CI)
34
32
30
28
26
24
M
GangesRiverDolphin
Species

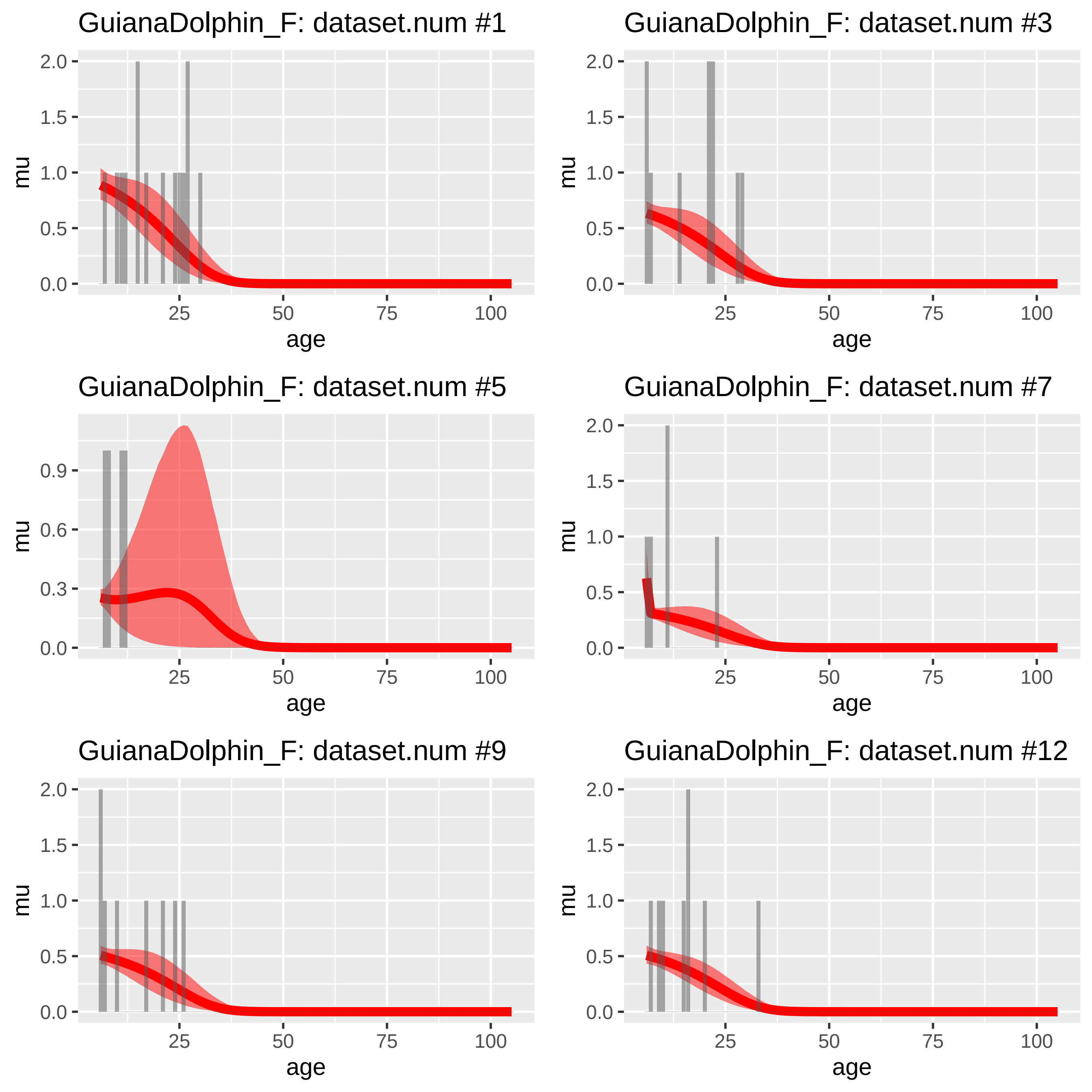
GuianaDolphin_F: dataset.num #1
GuianaDolphin_F: dataset.num #3
GuianaDolphin_F: dataset.num #5
GuianaDolphin_F: dataset.num #7
GuianaDolphin_F: dataset.num #9
GuianaDolphin_F: dataset.num #12
mu
age
0.0
0.5
1.0
1.5
2.0
0.3
0.6
0.9
25
50
75
100

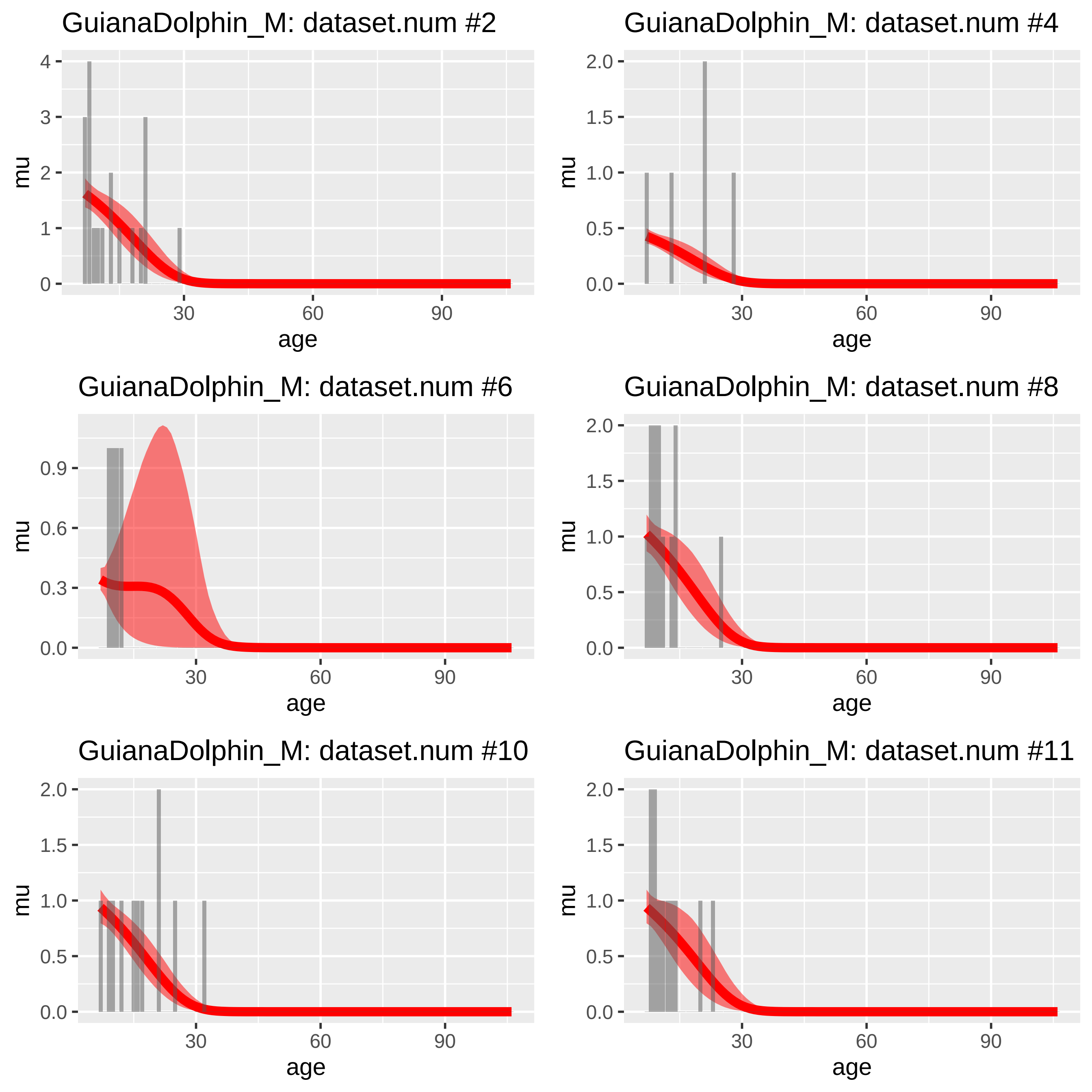
GuianaDolphin_M: dataset.num #2
GuianaDolphin_M: dataset.num #4
GuianaDolphin_M: dataset.num #6
GuianaDolphin_M: dataset.num #8
GuianaDolphin_M: dataset.num #10
GuianaDolphin_M: dataset.num #11
mu
age
0 1 2 3 4
0.0 0.5 1.0 1.5 2.0
0.0 0.3 0.6 0.9
30 60 90

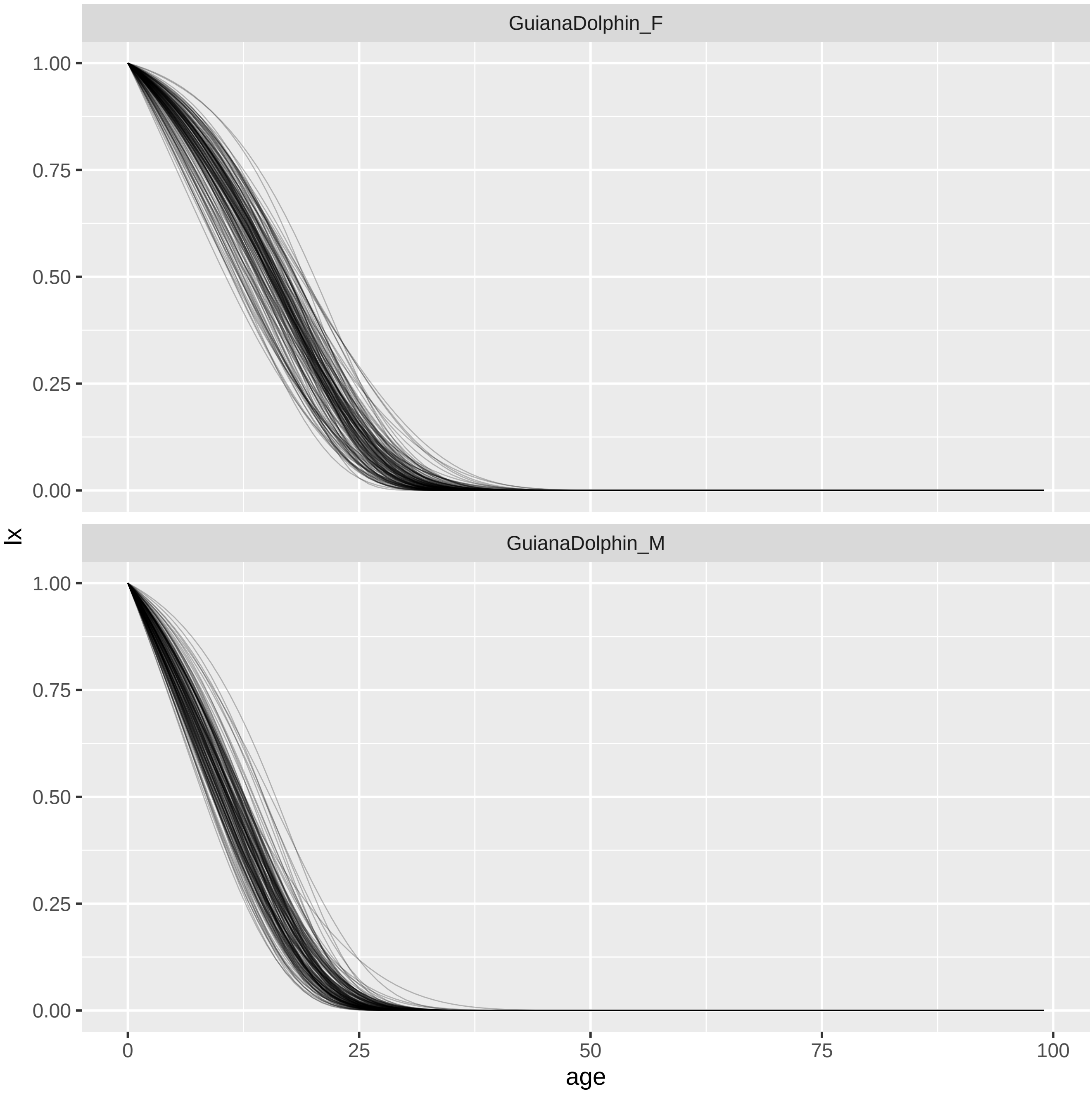

GuianaDolphin_F
GuianaDolphin_M
lx
1.00
0.75
0.50
0.25
0.00
0
25
50
75
100
age

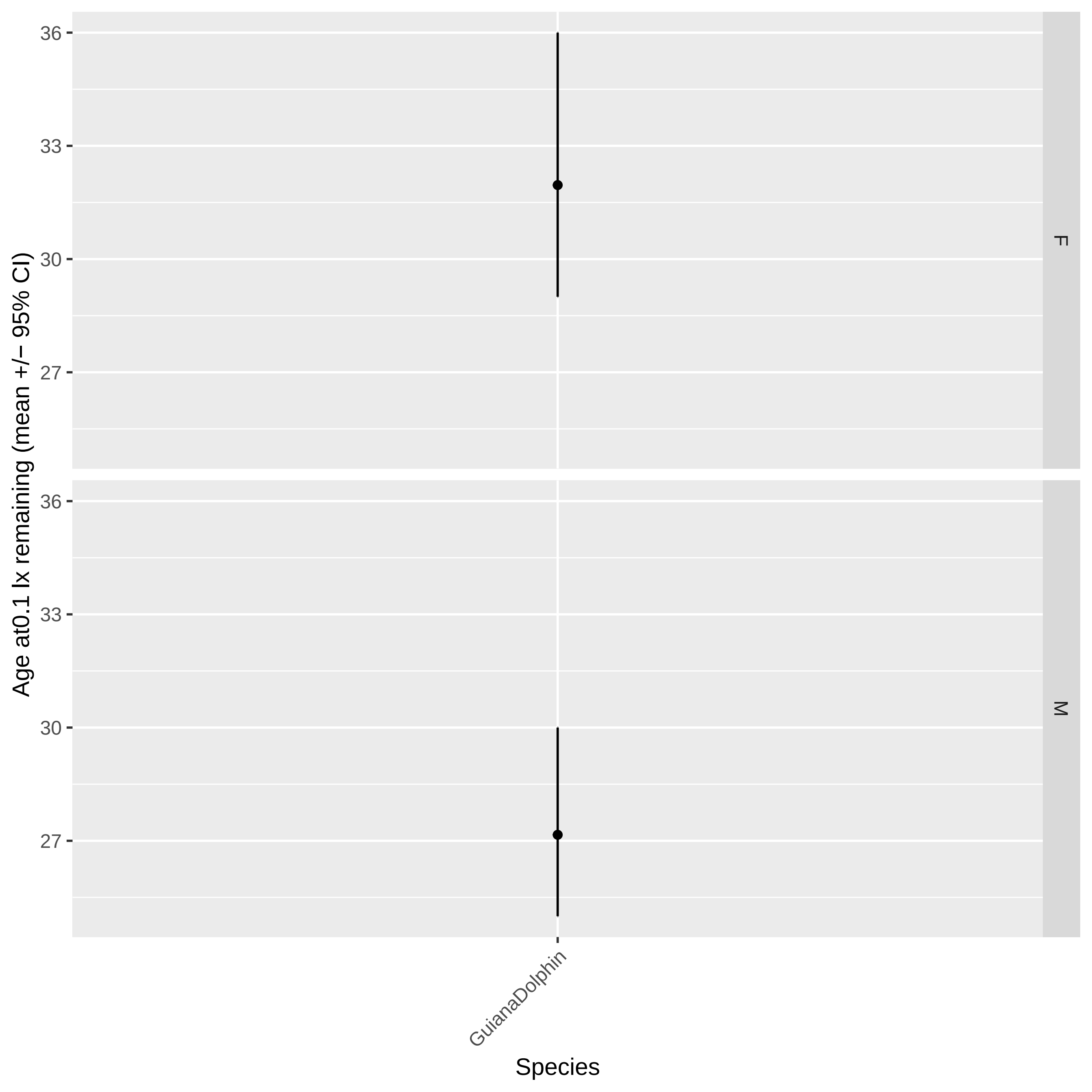

Age at0.1 lx remaining (mean +/− 95% CI)
36
33
30
27
F
M
GuianaDolphin
Species

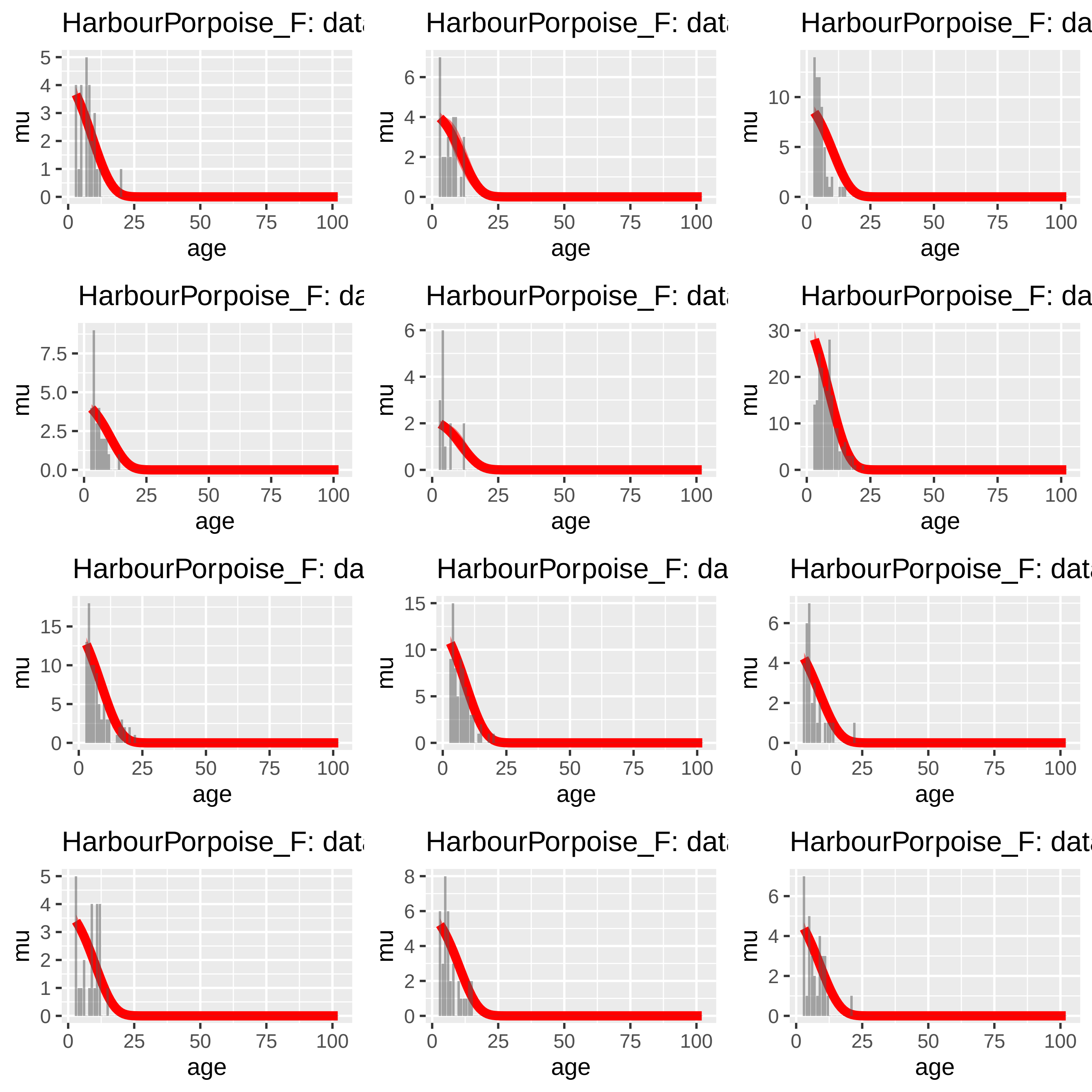

HarbourPorpoise_F: dat
HarbourPorpoise_F: dat
HarbourPorpoise_F: da
HarbourPorpoise_F: da
HarbourPorpoise_F: dat
HarbourPorpoise_F: da
HarbourPorpoise_F: da
HarbourPorpoise_F: da
HarbourPorpoise_F: dat
HarbourPorpoise_F: dat
HarbourPorpoise_F: dat
HarbourPorpoise_F: dat
mu
age
0
25
50
75
100

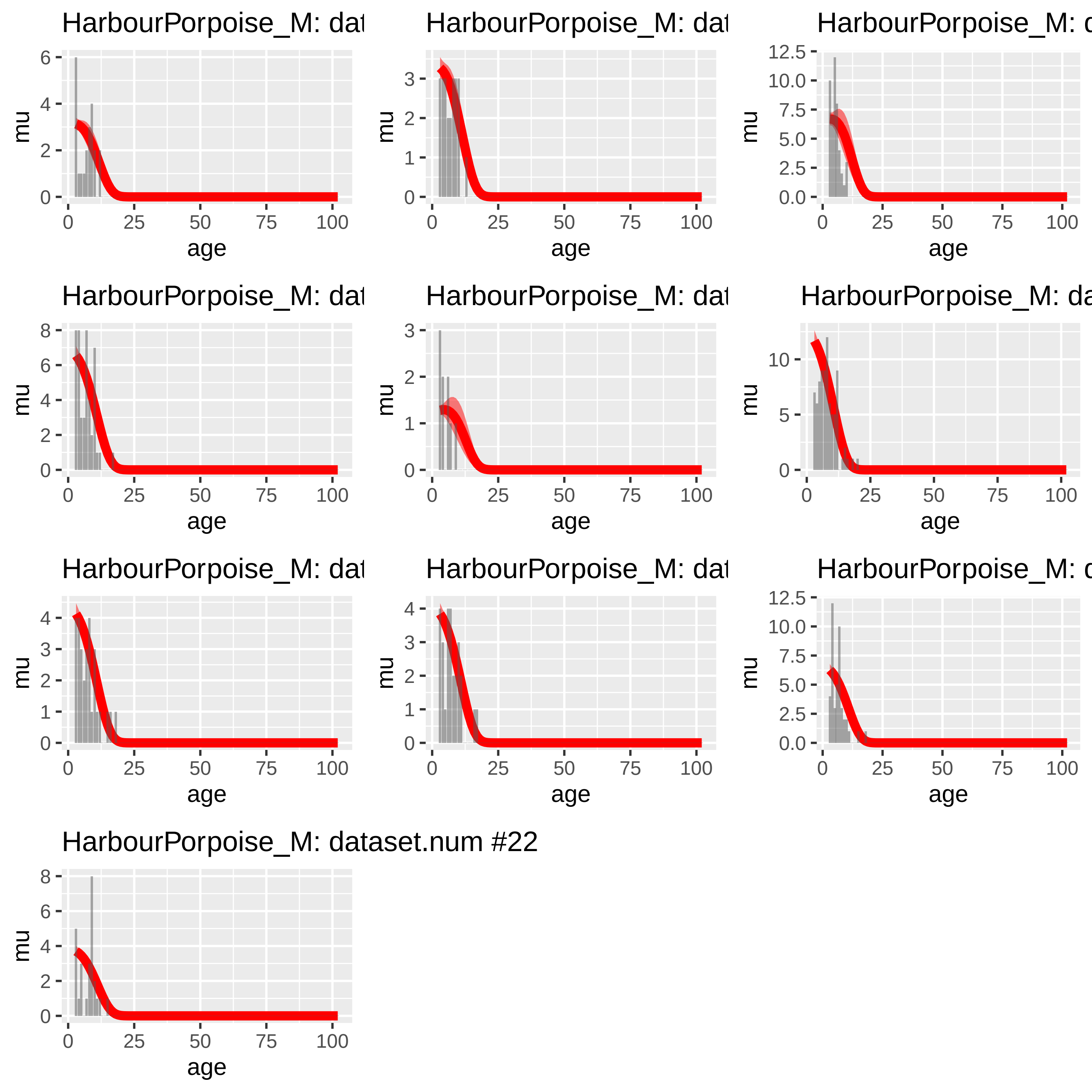
HarbourPorpoise_M: dat
HarbourPorpoise_M: dat
HarbourPorpoise_M: d
HarbourPorpoise_M: dat
HarbourPorpoise_M: dat
HarbourPorpoise_M: da
HarbourPorpoise_M: dat
HarbourPorpoise_M: dat
HarbourPorpoise_M: d
HarbourPorpoise_M: dataset.num #22
mu
age
0
25
50
75
100

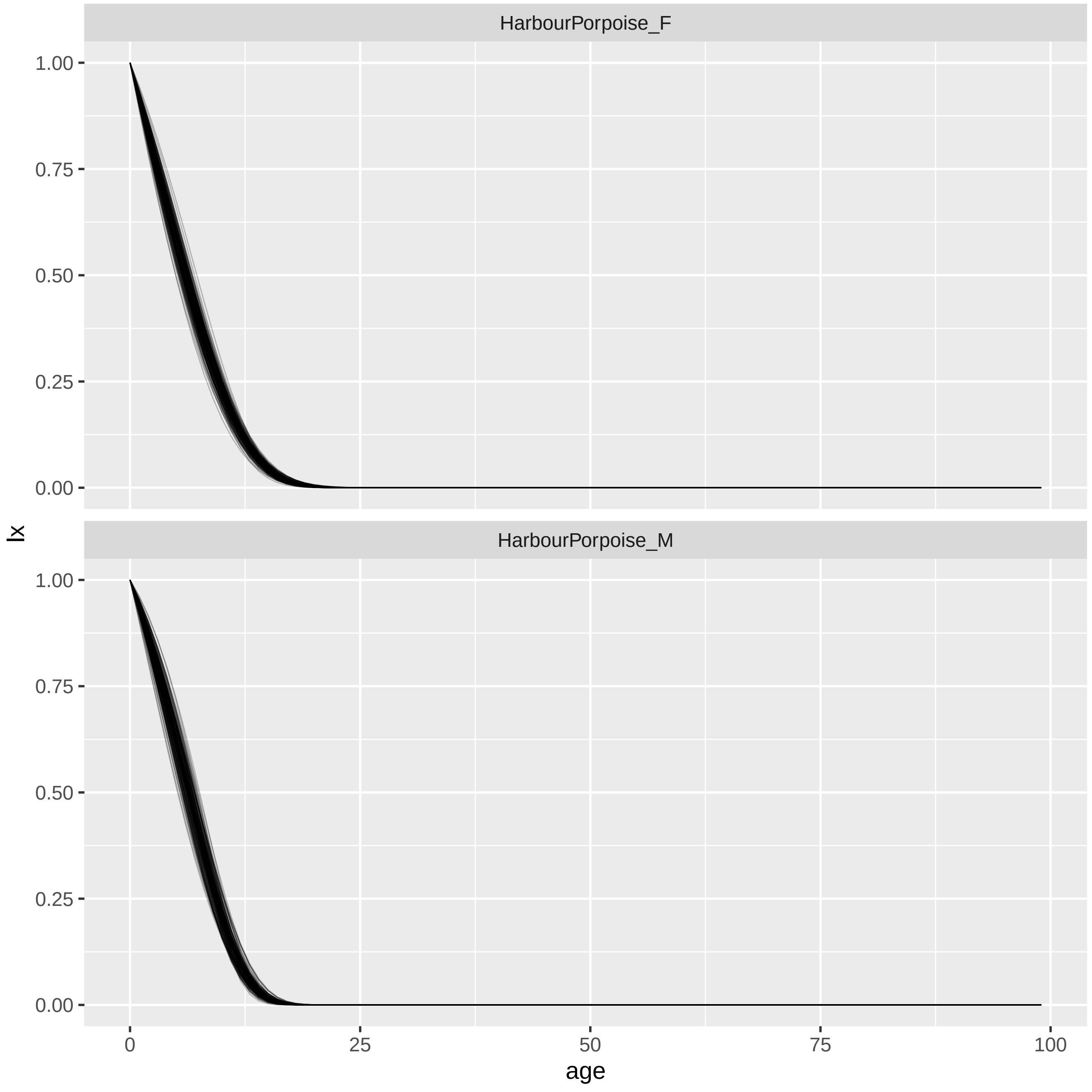

HarbourPorpoise_F
HarbourPorpoise_M
1.00
0.75
0.50
0.25
0.00
lx
0
25
50
75
100
age

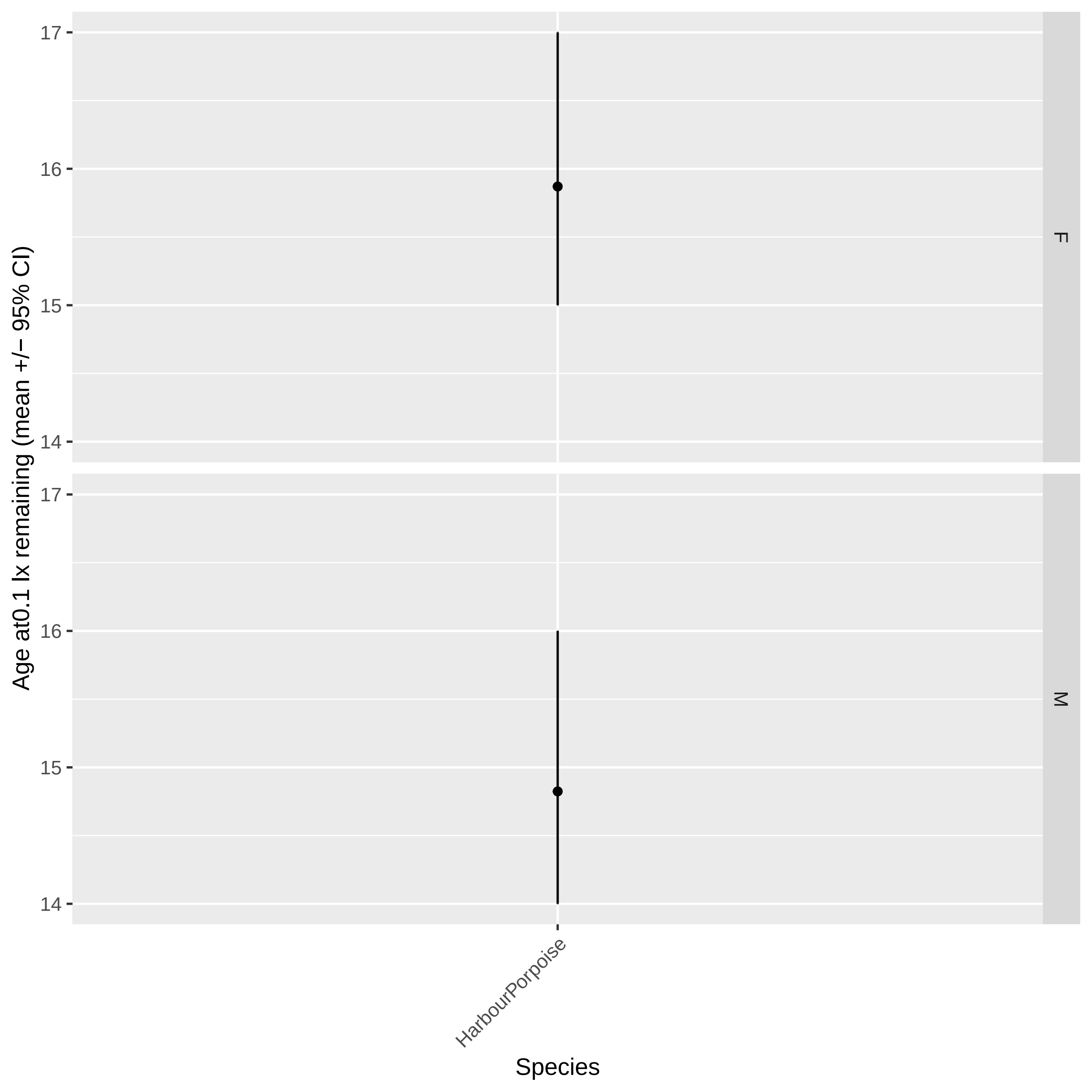
Age at0.1 lx remaining (mean +/− 95% CI)
17
16
15
14
F
M
HarbourPorpoise
Species

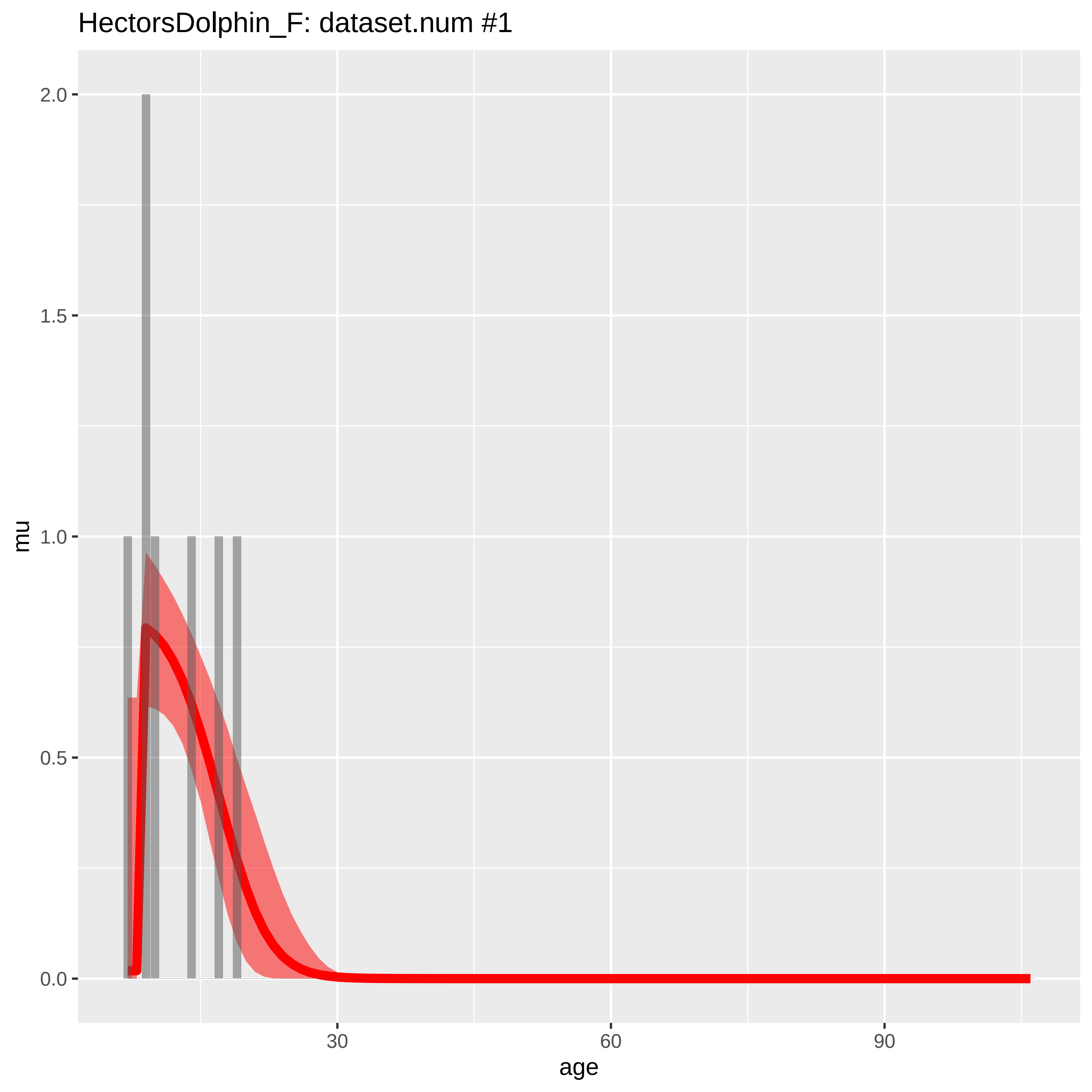
HectorsDolphin_F: dataset.num #1
mu
age
2.0
1.5
1.0
0.5
0.0
30
60
90

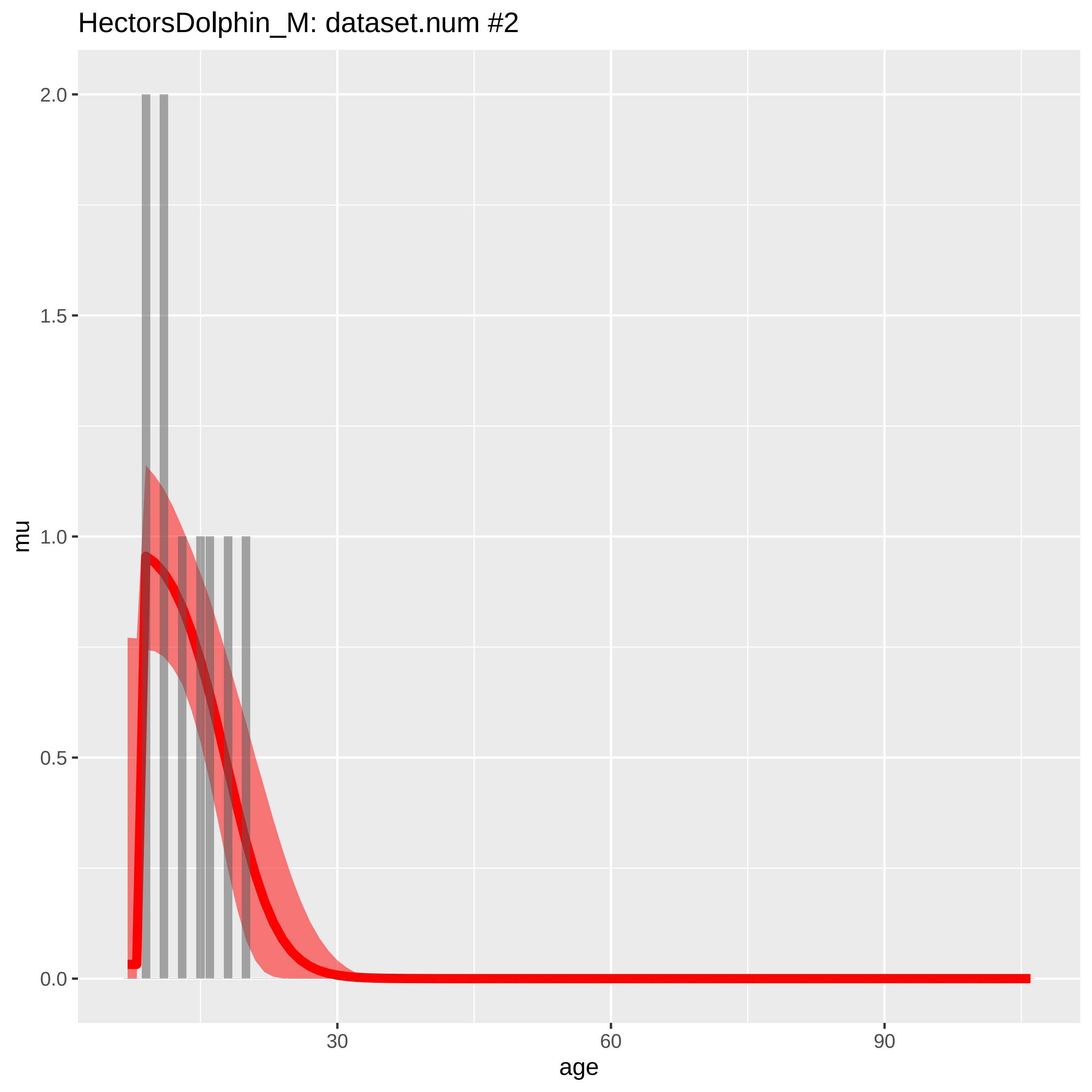
HectorsDolphin_M: dataset.num #2
2.0
1.5
1.0
0.5
0.0
mu
30
60
90
age

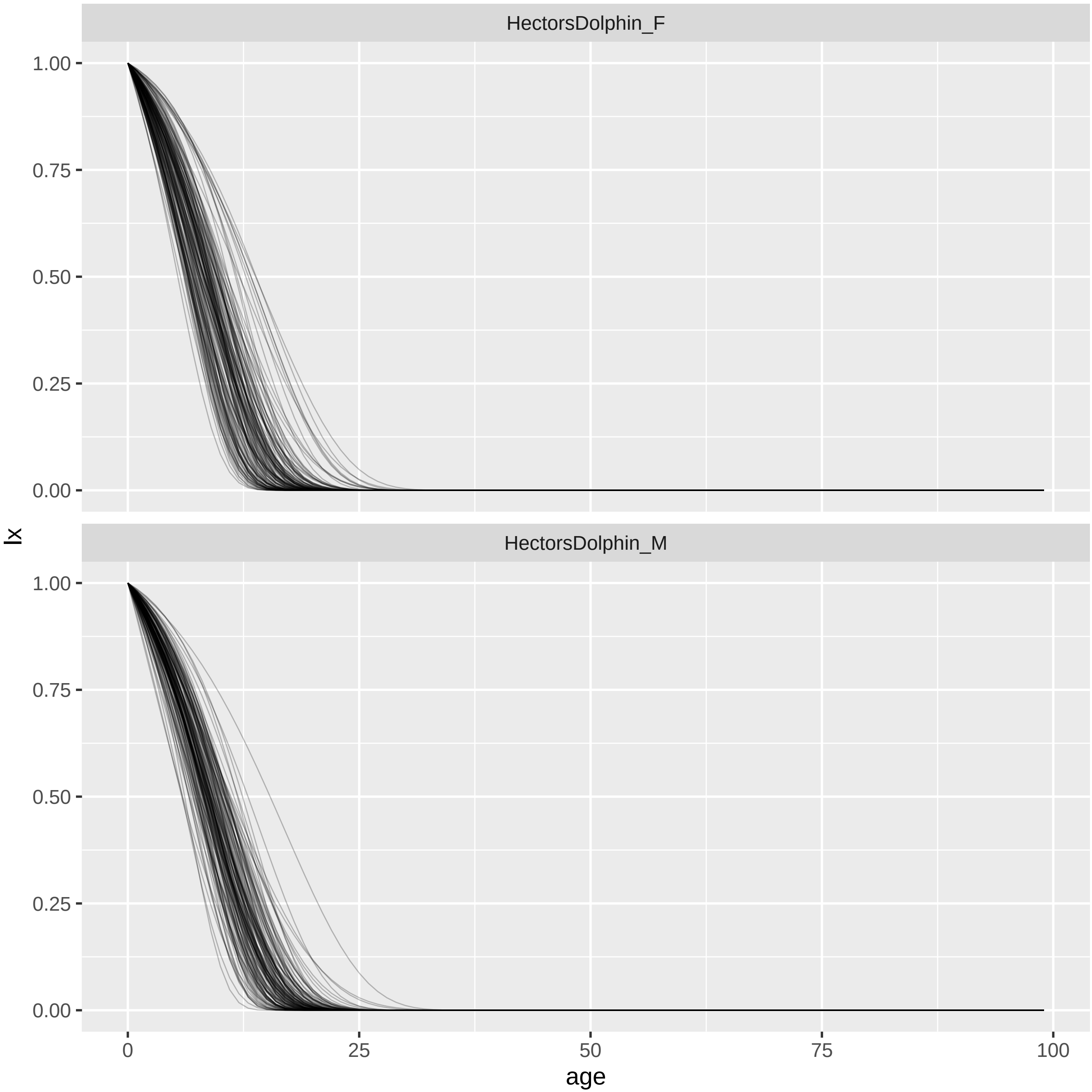
HectorsDolphin_F
HectorsDolphin_M
1.00
0.75
0.50
0.25
0.00
lx
0
25
50
75
100
age

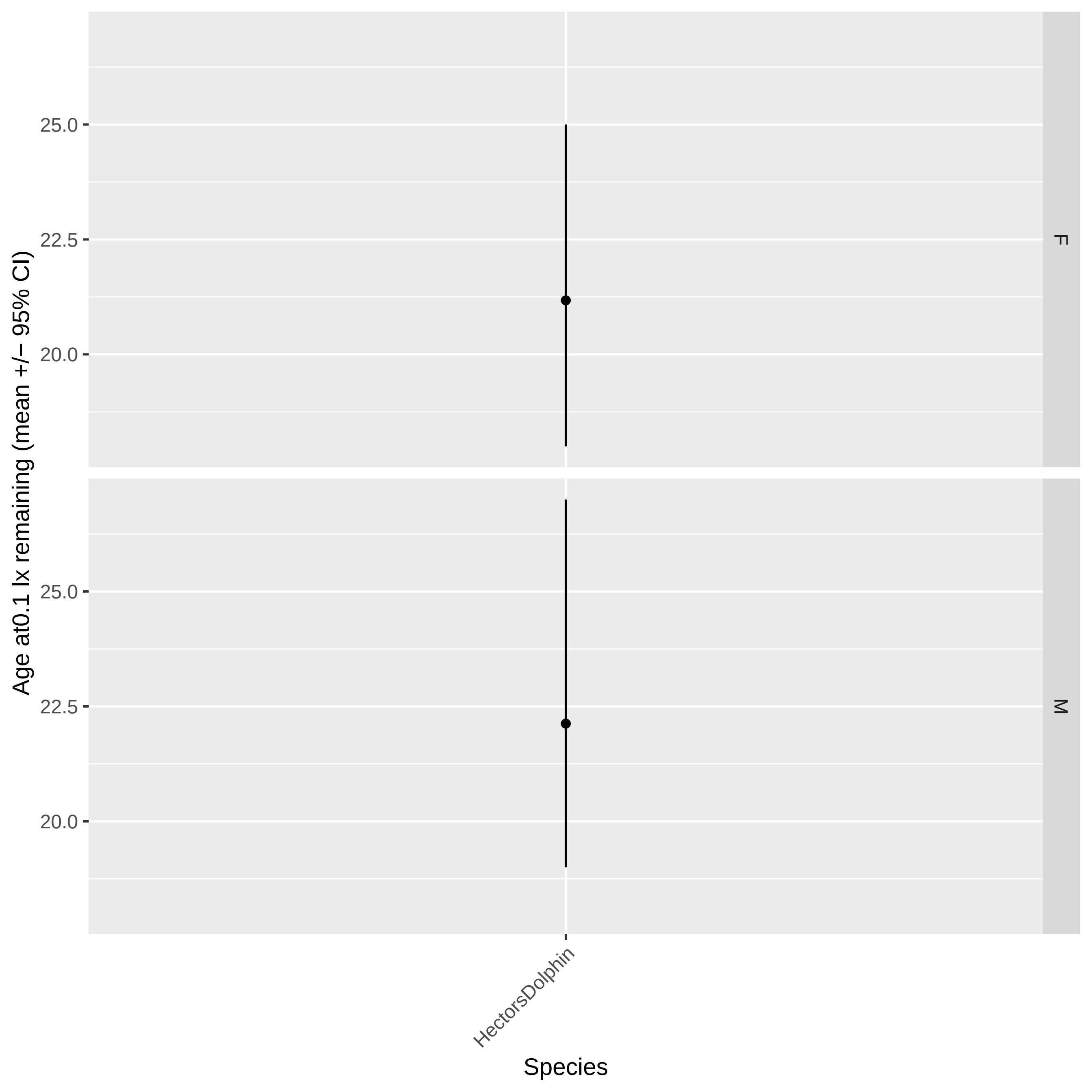

Age at0.1 lx remaining (mean +/− 95% CI)
25.0
22.5
20.0
F
M
HectorsDolphin
Species

IndianOceanHumpbackDolphin_F: dataset.num #1

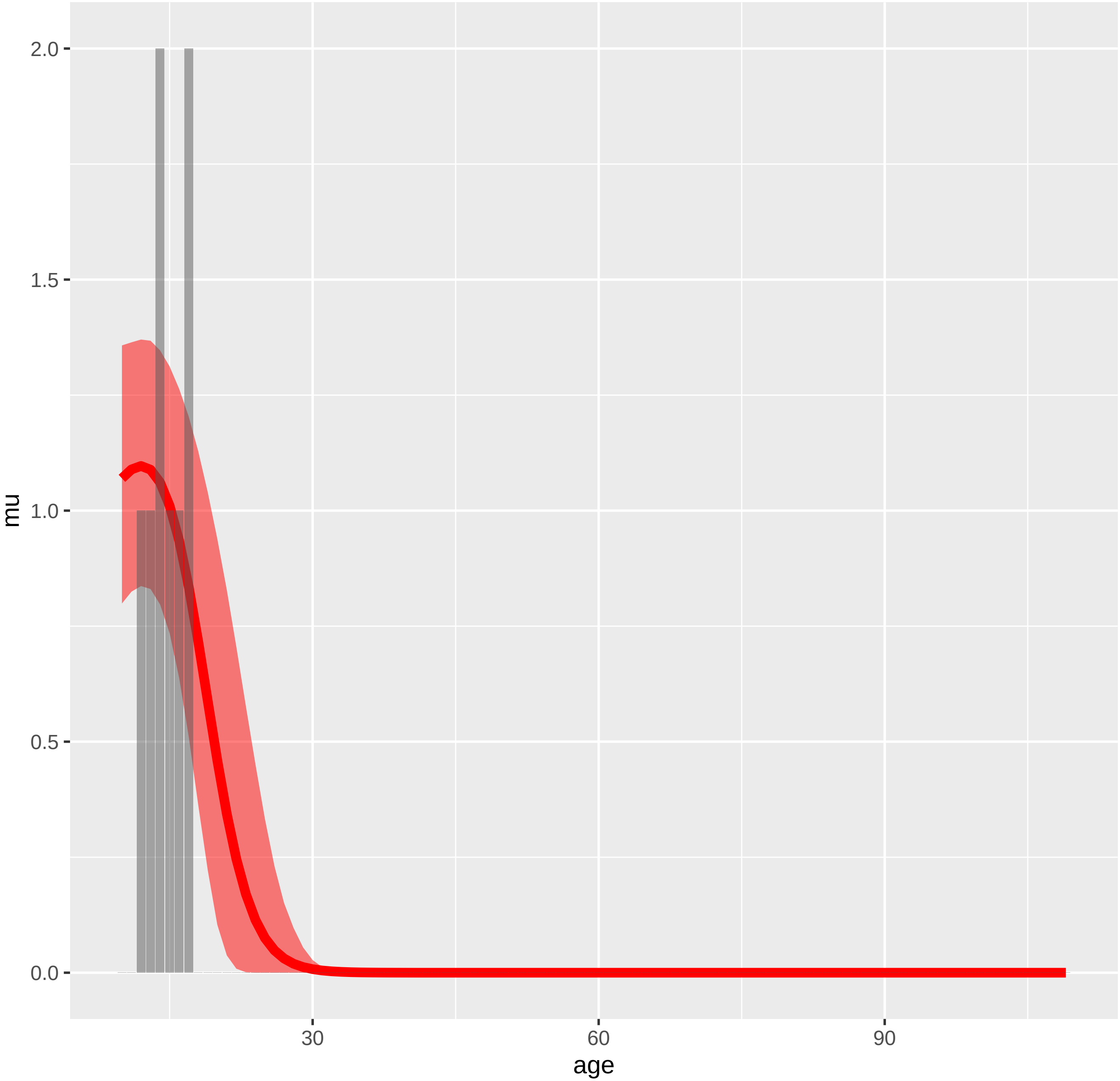

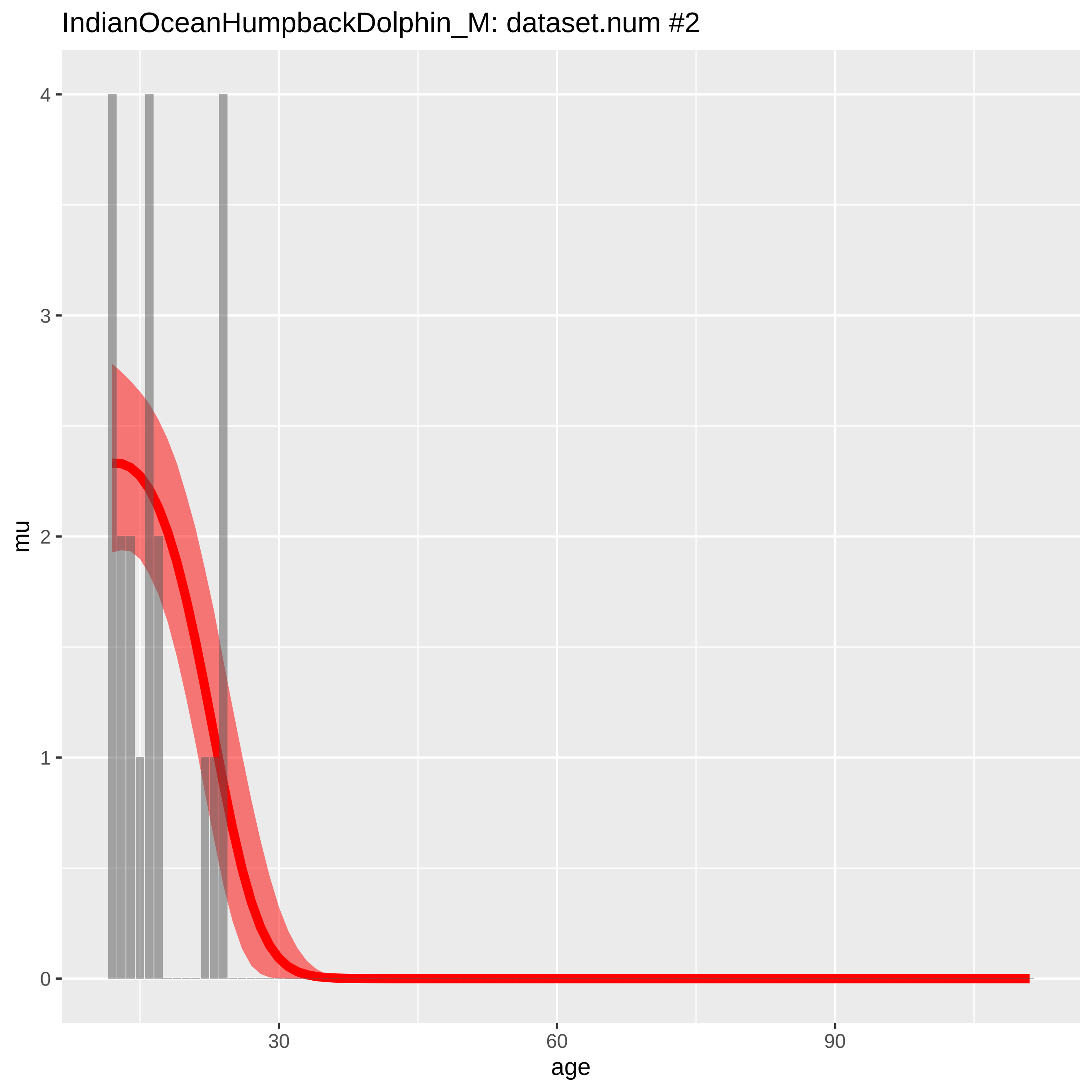
IndianOceanHumpbackDolphin_M: dataset.num #2
mu
age
0
1
2
3
4
30
60
90

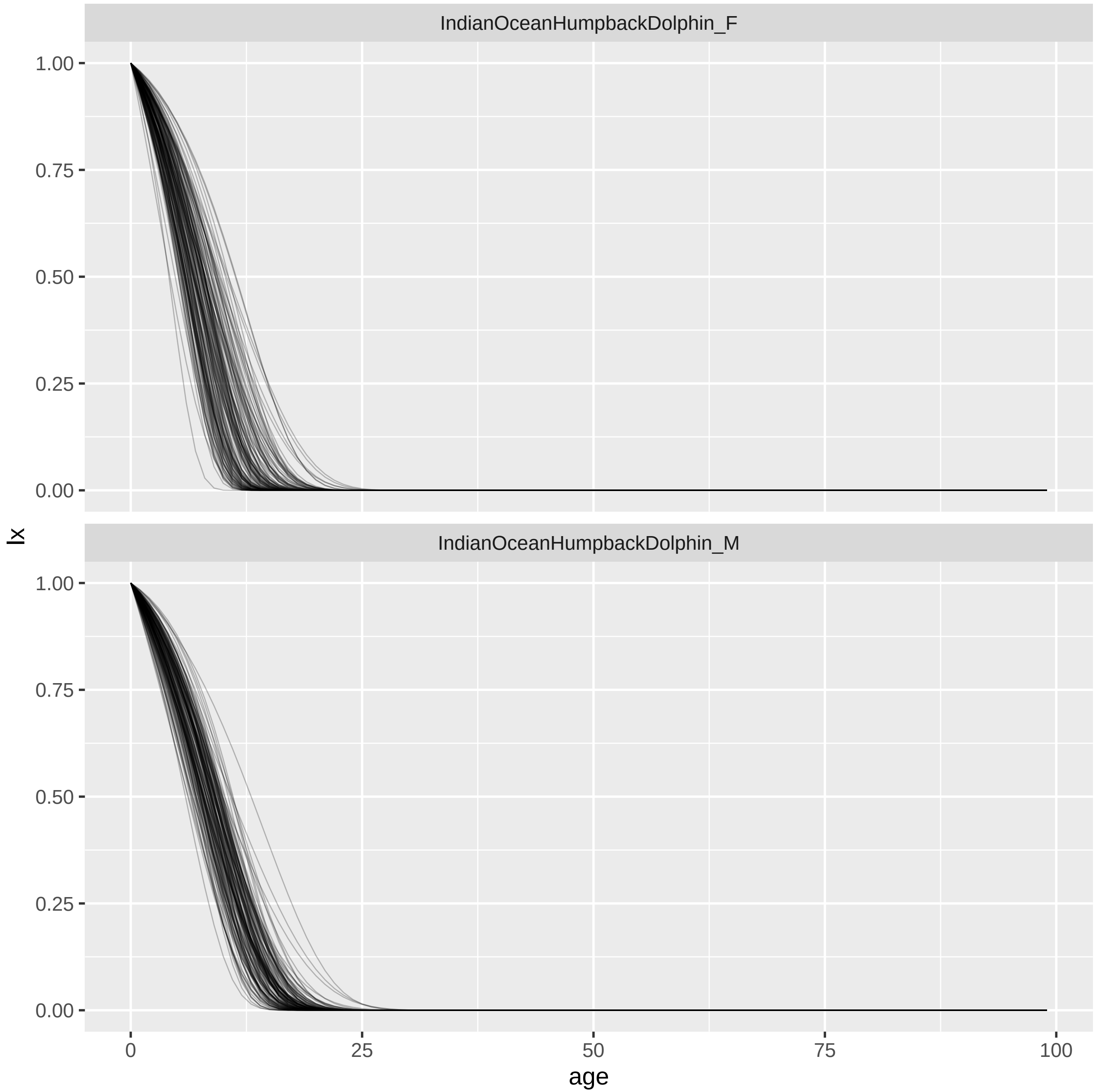
IndianOceanHumpbackDolphin_F
IndianOceanHumpbackDolphin_M
1.00
0.75
0.50
0.25
0.00
lx
0
25
50
75
100
age

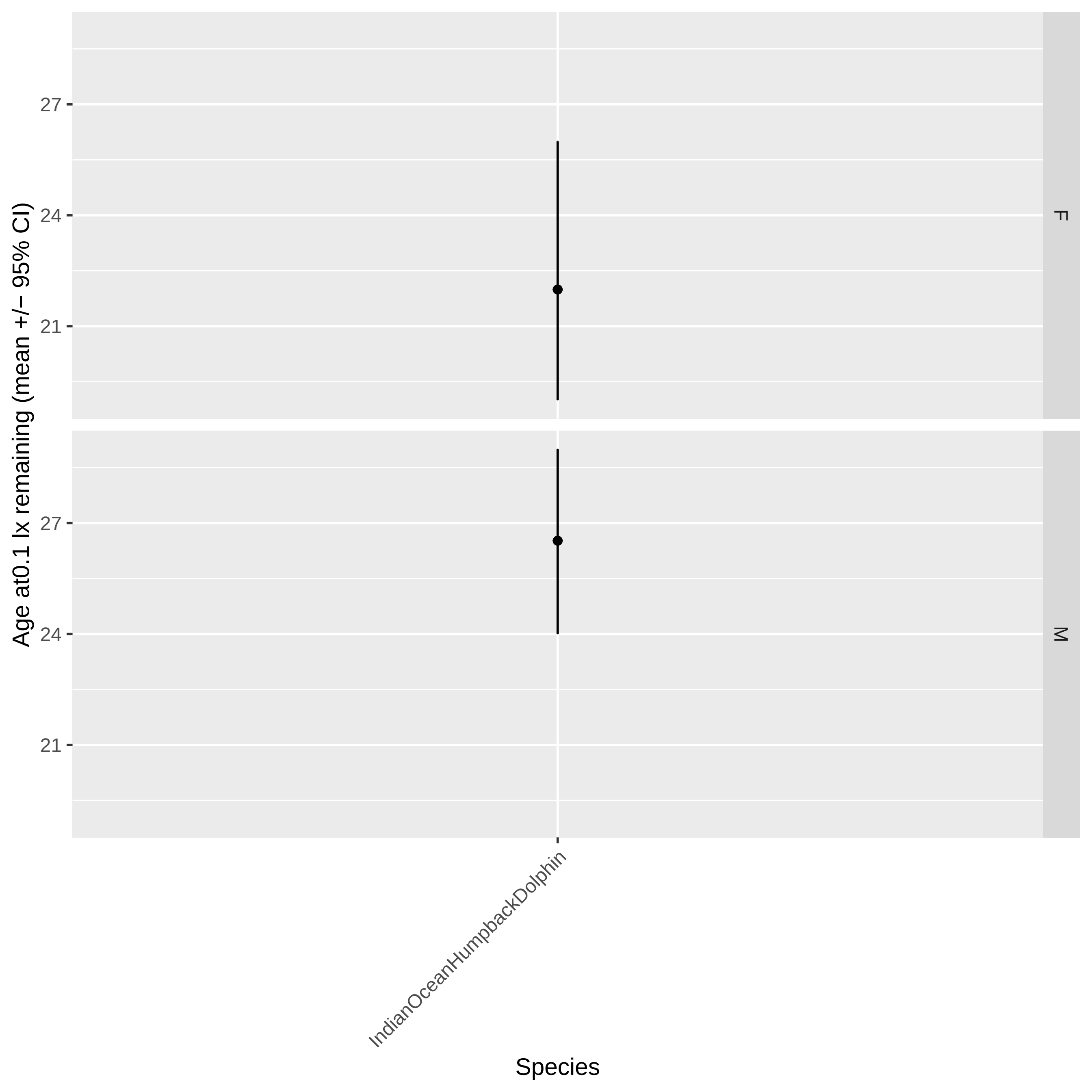

Age at0.1 lx remaining (mean +/− 95% CI)
27
24
21
F
M
IndianOceanHumpbackDolphin
Species

IndoPacificBottlenoseDolphin_F: dataset.num #2

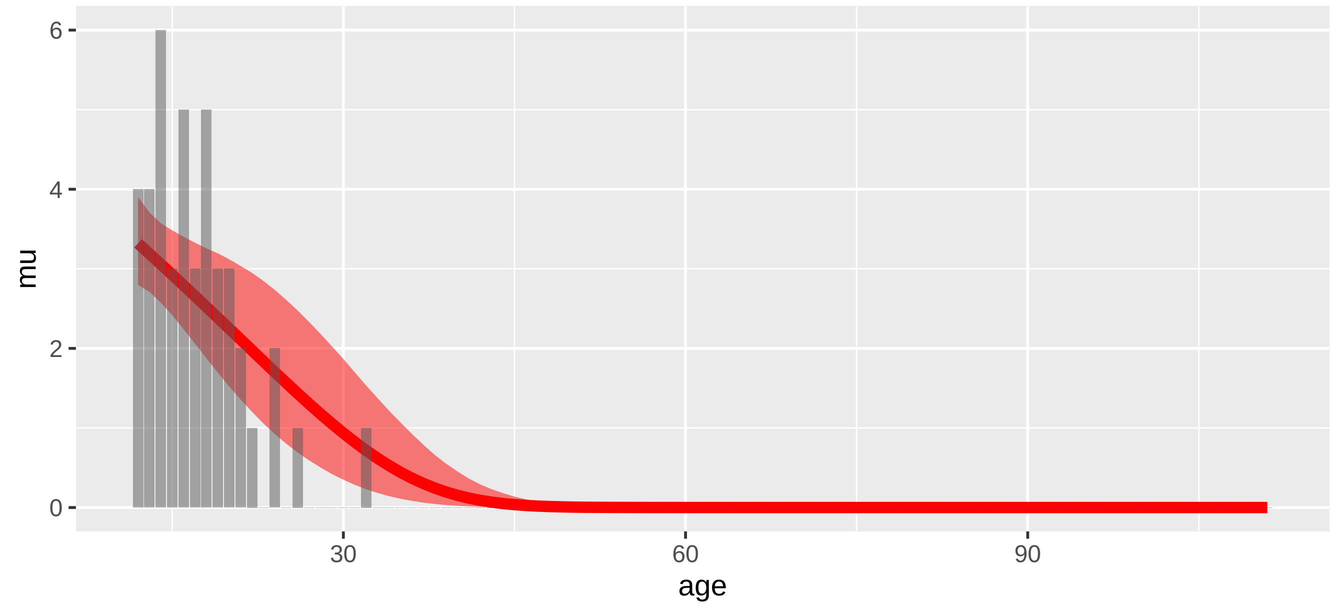


IndoPacificBottlenoseDolphin_F: dataset.num #3

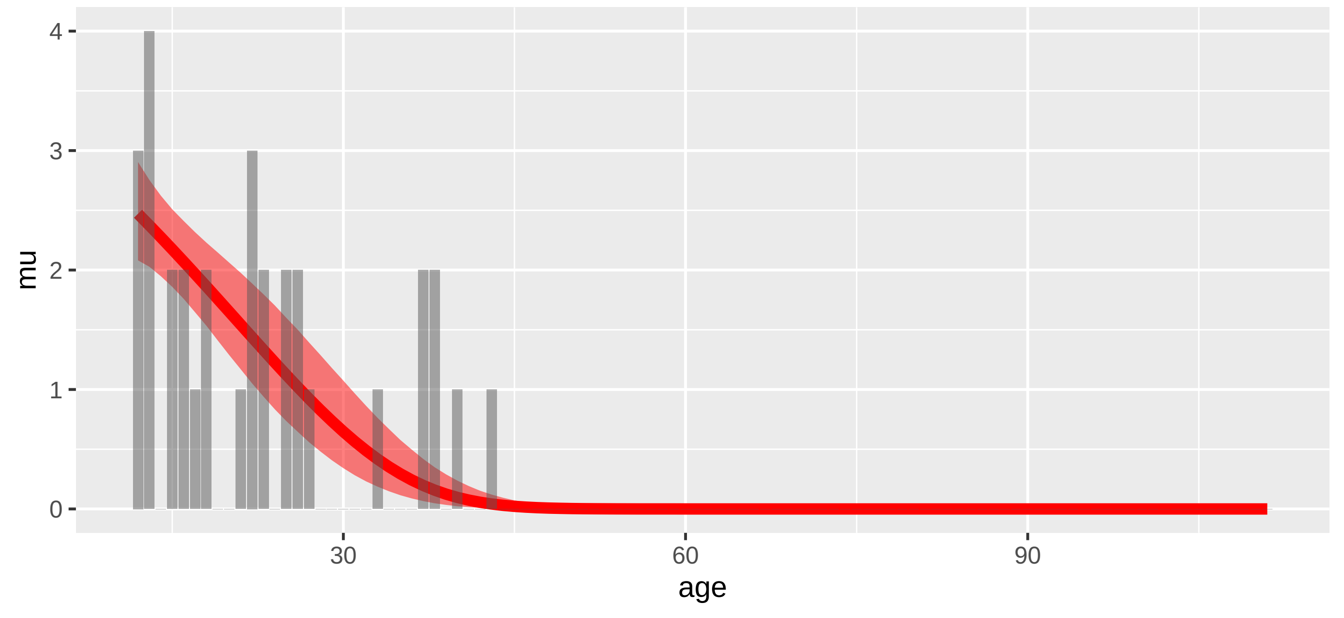

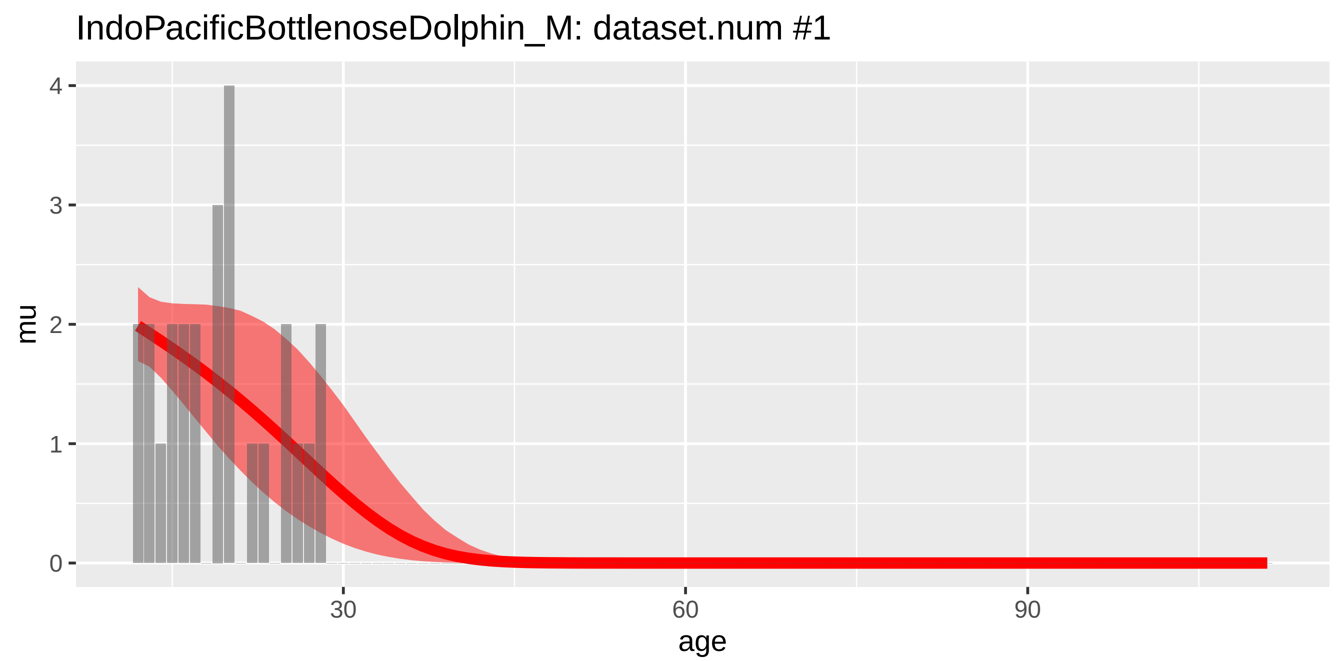
IndoPacificBottlenoseDolphin_M: dataset.num #1
mu
4
3
2
1
0
30
60
90
age

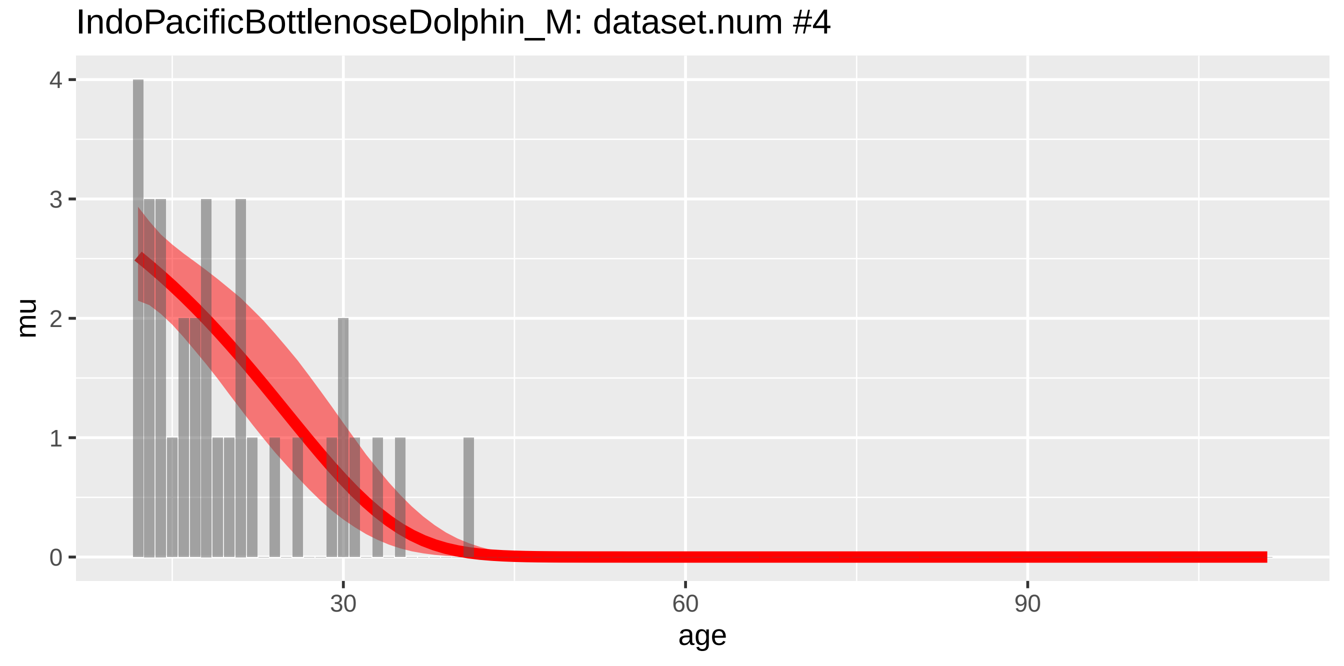
IndoPacificBottlenoseDolphin_M: dataset.num #4
mu
4
3
2
1
0
30
60
90
age

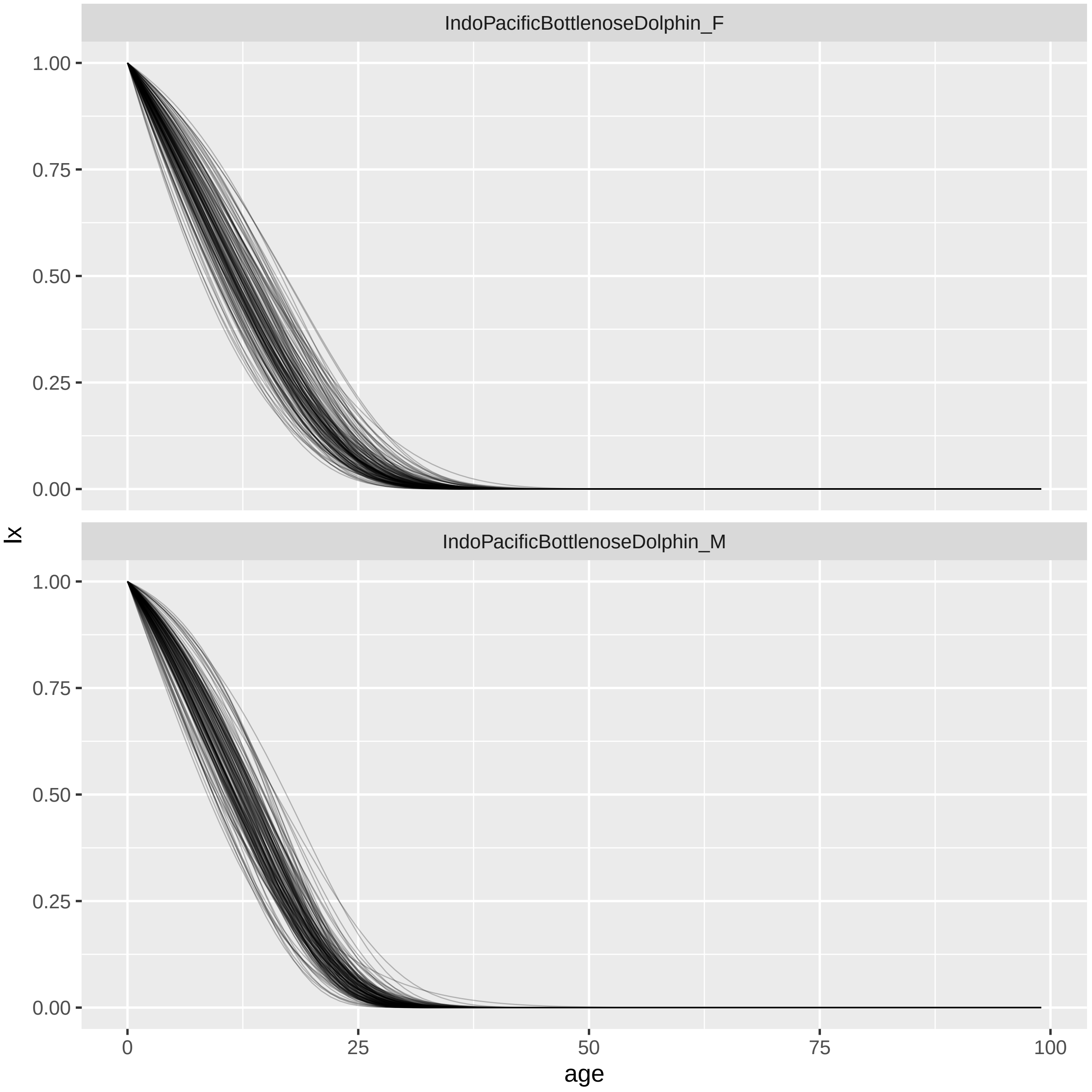

IndoPacificBottlenoseDolphin_F
IndoPacificBottlenoseDolphin_M
lx
1.00
0.75
0.50
0.25
0.00
0
25
50
75
100
age

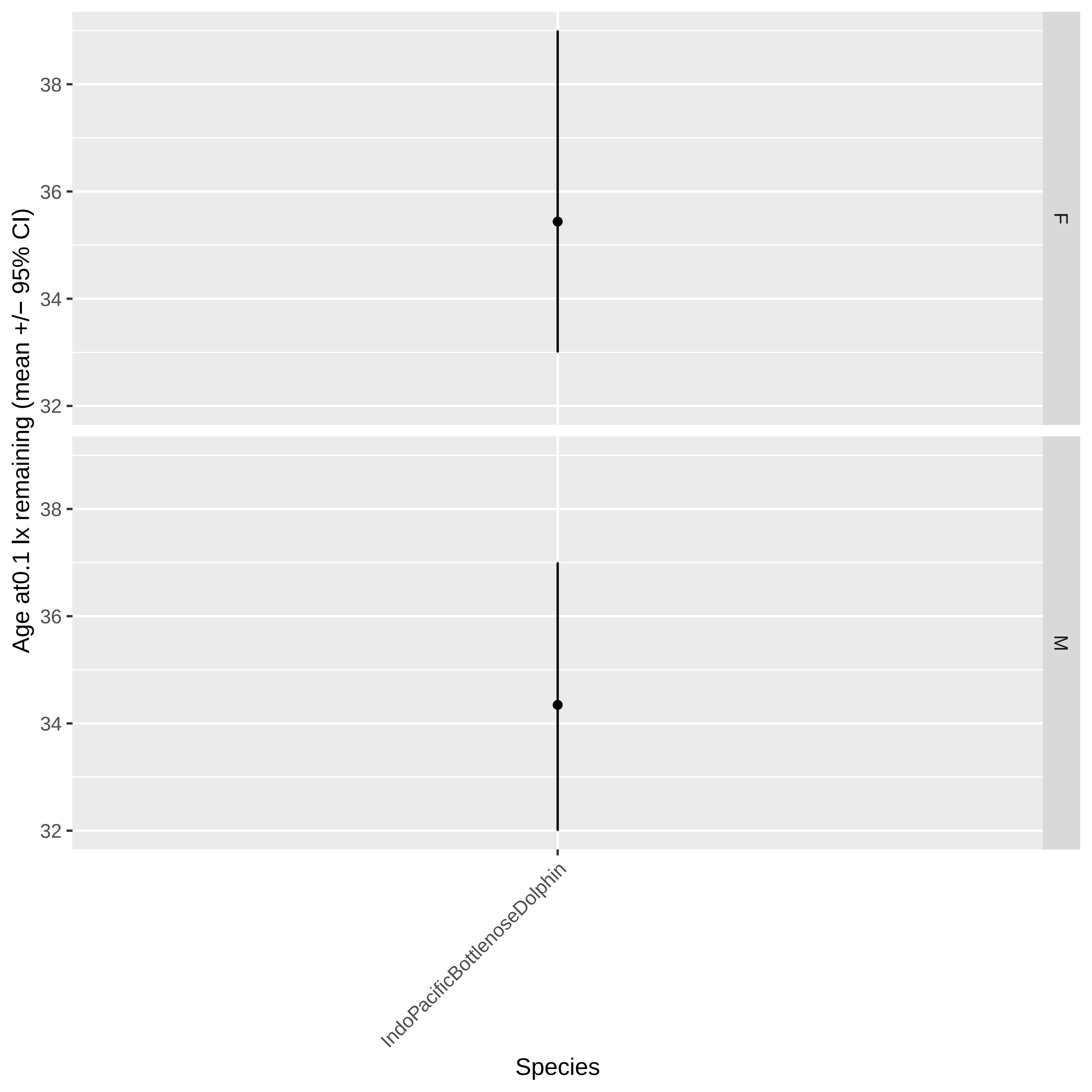

Age at0.1 lx remaining (mean +/− 95% CI)
38
36
34
32
F
M
IndoPacificBottlenoseDolphin
Species

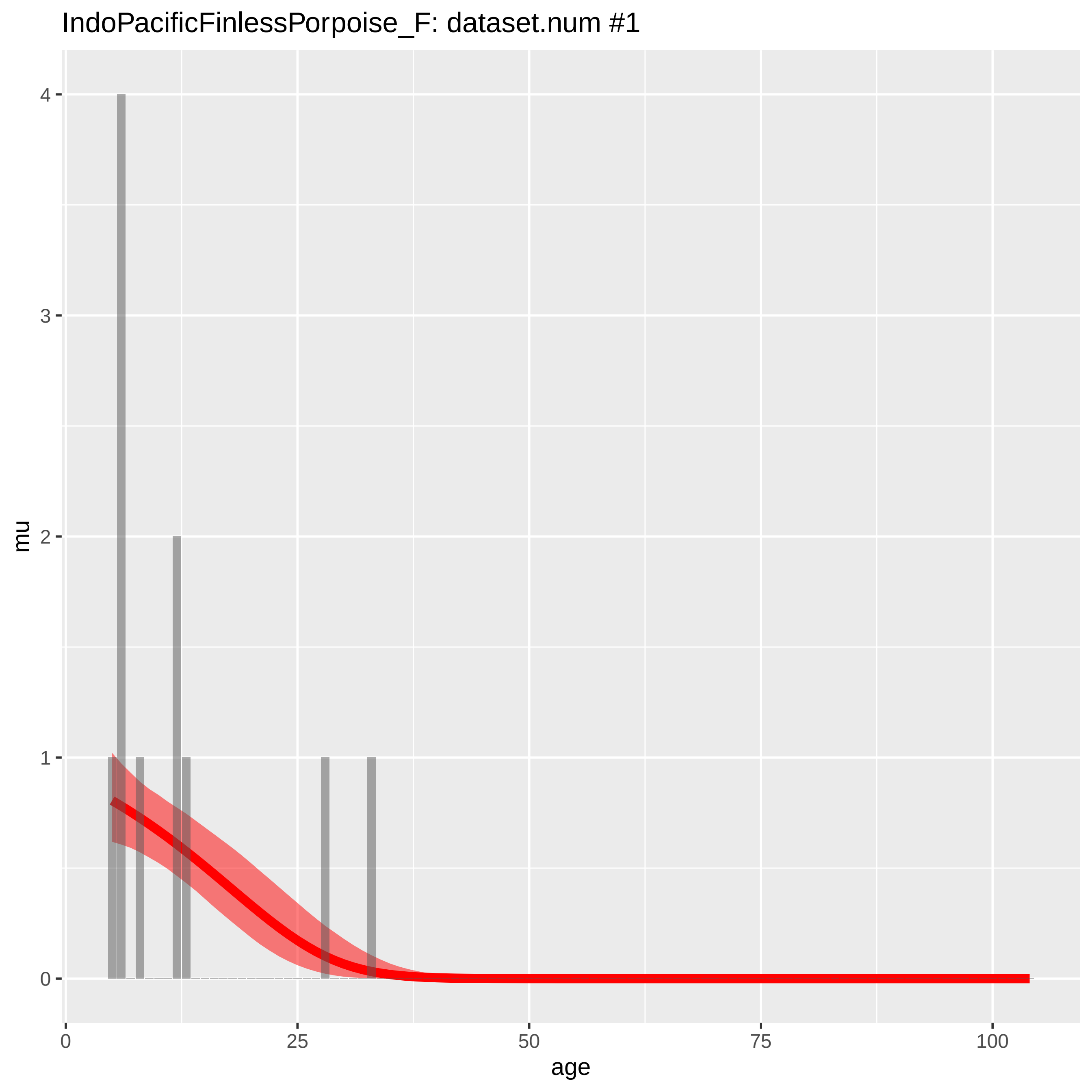

IndoPacificFinlessPorpoise_F: dataset.num #1
mu
4
3
2
1
0
0
25
50
75
100
age

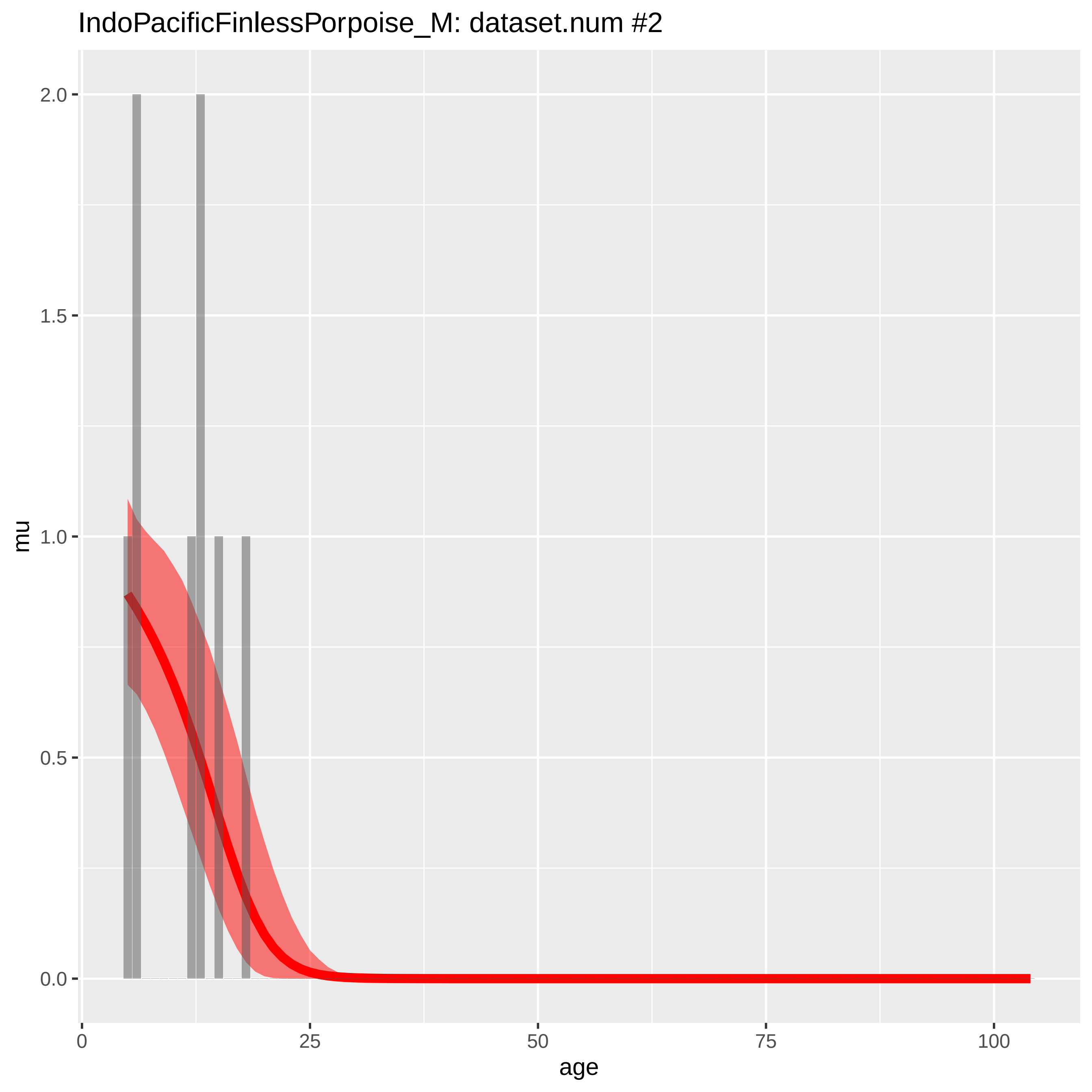

IndoPacificFinlessPorpoise_M: dataset.num #2
mu
2.0
1.5
1.0
0.5
0.0
0
25
50
75
100
age

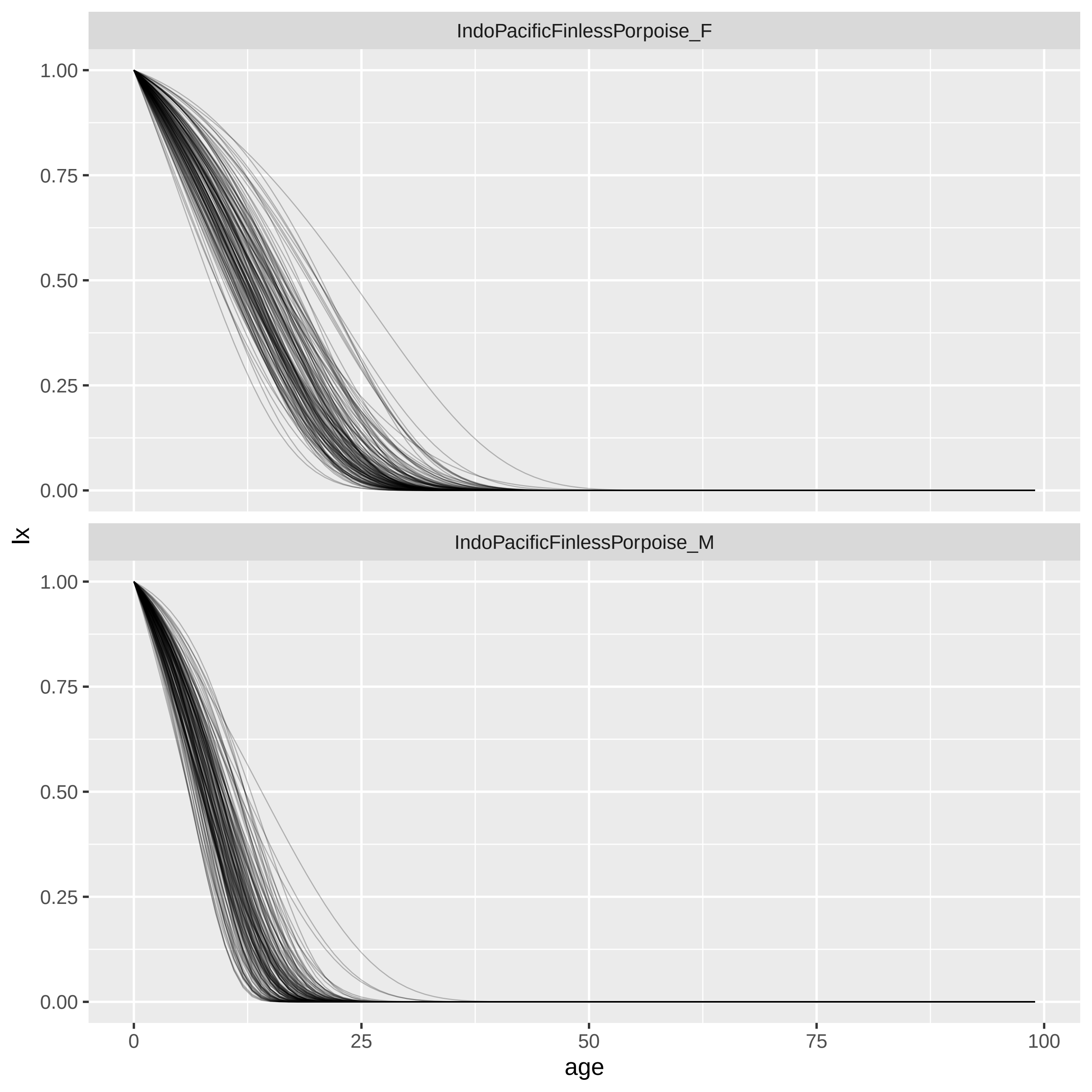
IndoPacificFinlessPorpoise_F
IndoPacificFinlessPorpoise_M
lx
1.00
0.75
0.50
0.25
0.00
age
0
25
50
75
100

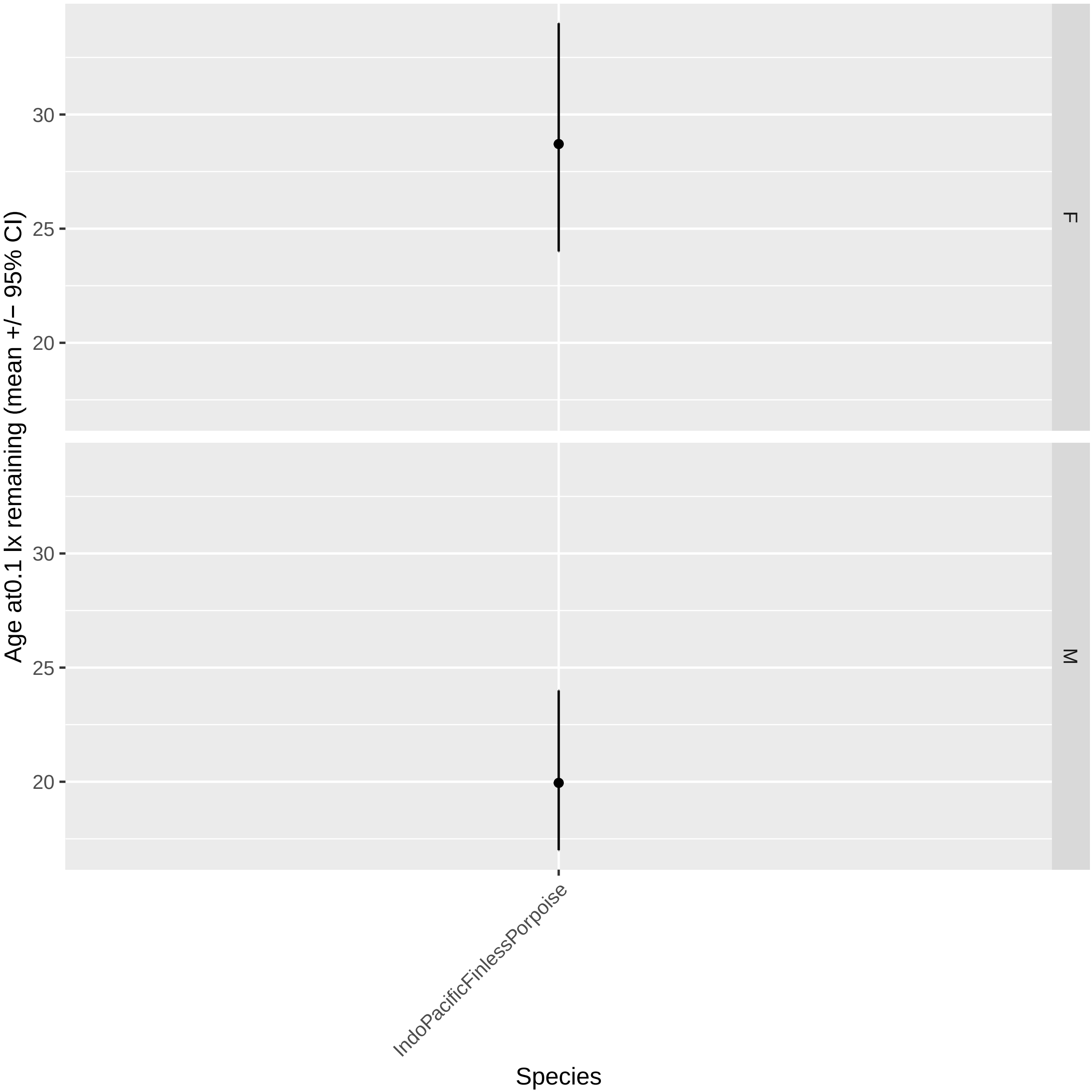

Age at0.1 lx remaining (mean +/− 95% CI)
F
M
30
25
20
IndoPacificFinlessPorpoise
Species

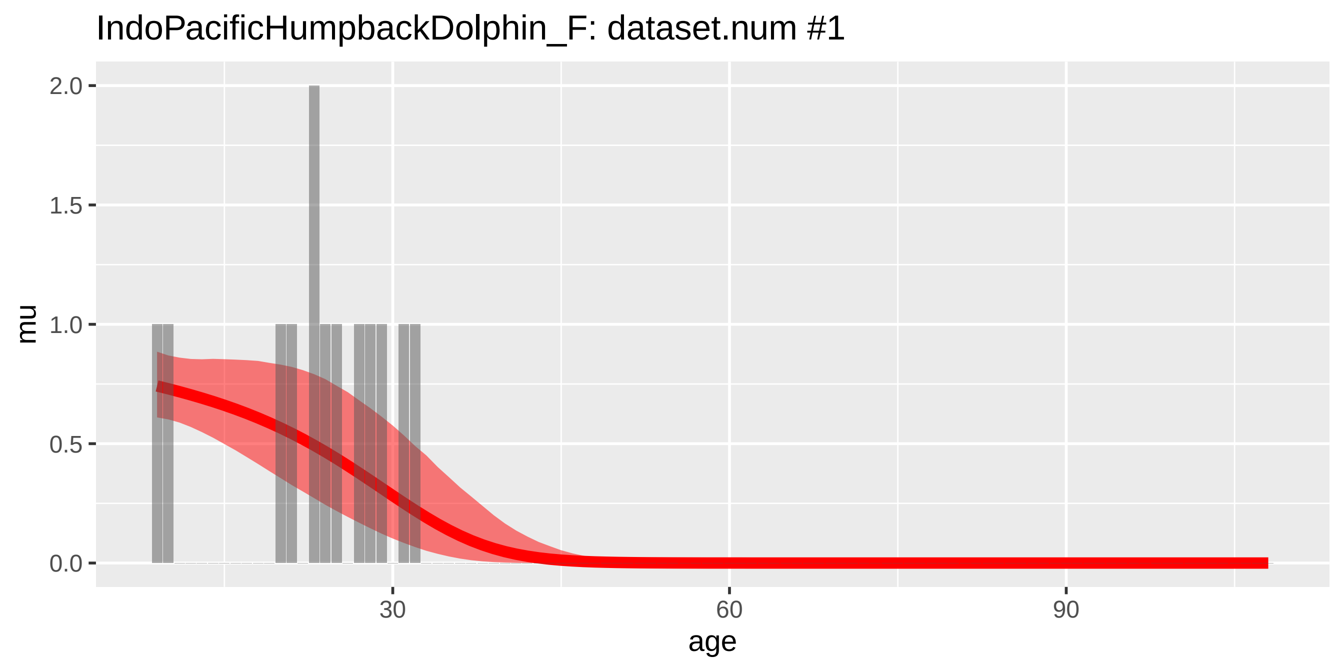
IndoPacificHumpbackDolphin_F: dataset.num #1
mu
age
2.0
1.5
1.0
0.5
0.0
30
60
90

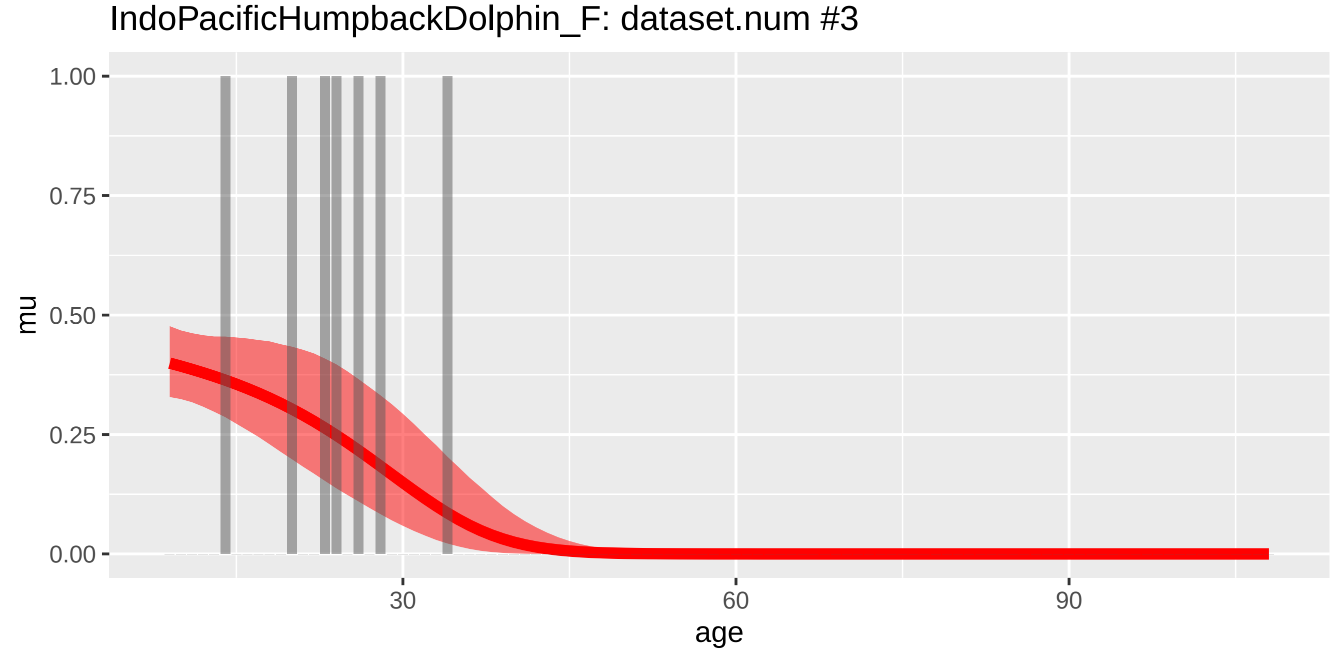
IndoPacificHumpbackDolphin_F: dataset.num #3
mu
age
1.00
0.75
0.50
0.25
0.00
30
60
90

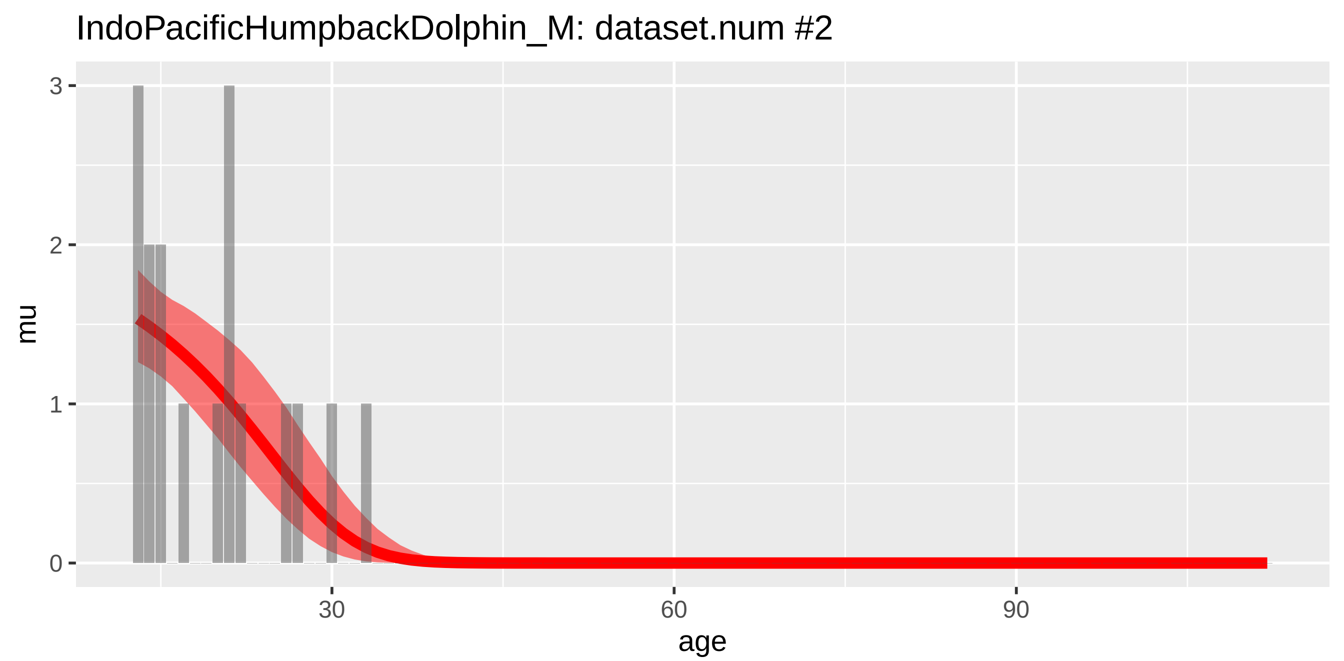
IndoPacificHumpbackDolphin_M: dataset.num #2
mu
3
2
1
0
30
60
90
age

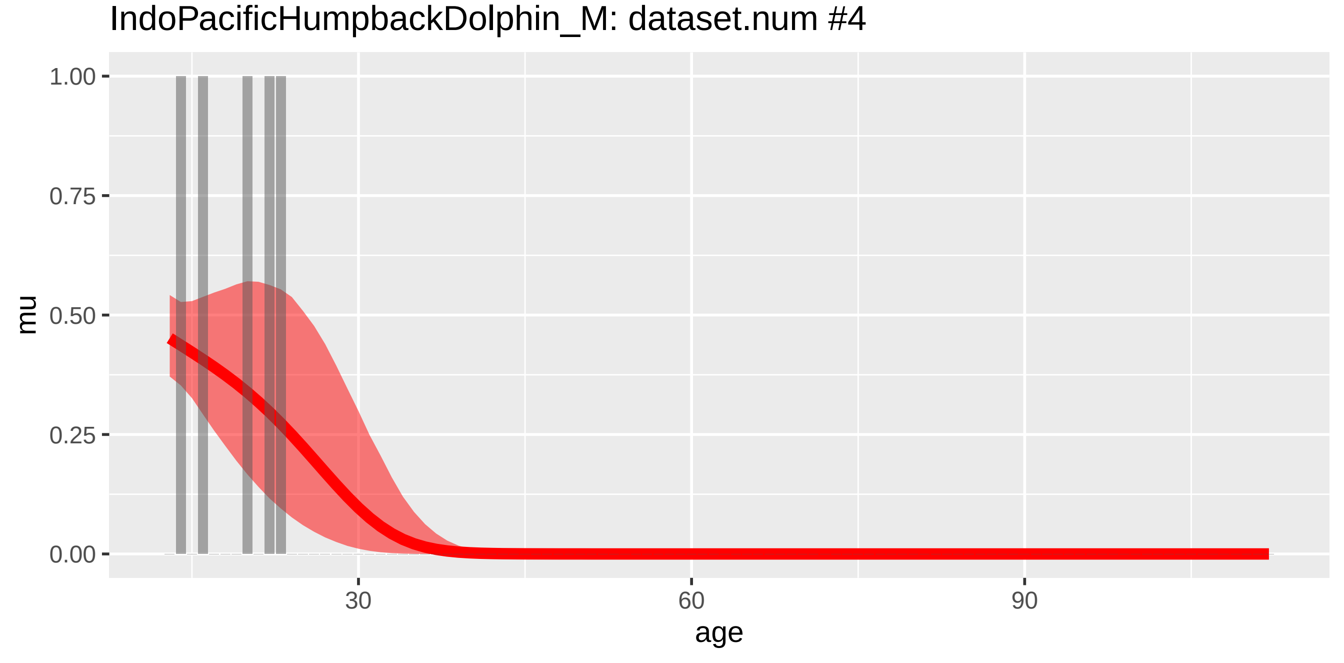
IndoPacificHumpbackDolphin_M: dataset.num #4
mu
1.00
0.75
0.50
0.25
0.00
30
60
90
age

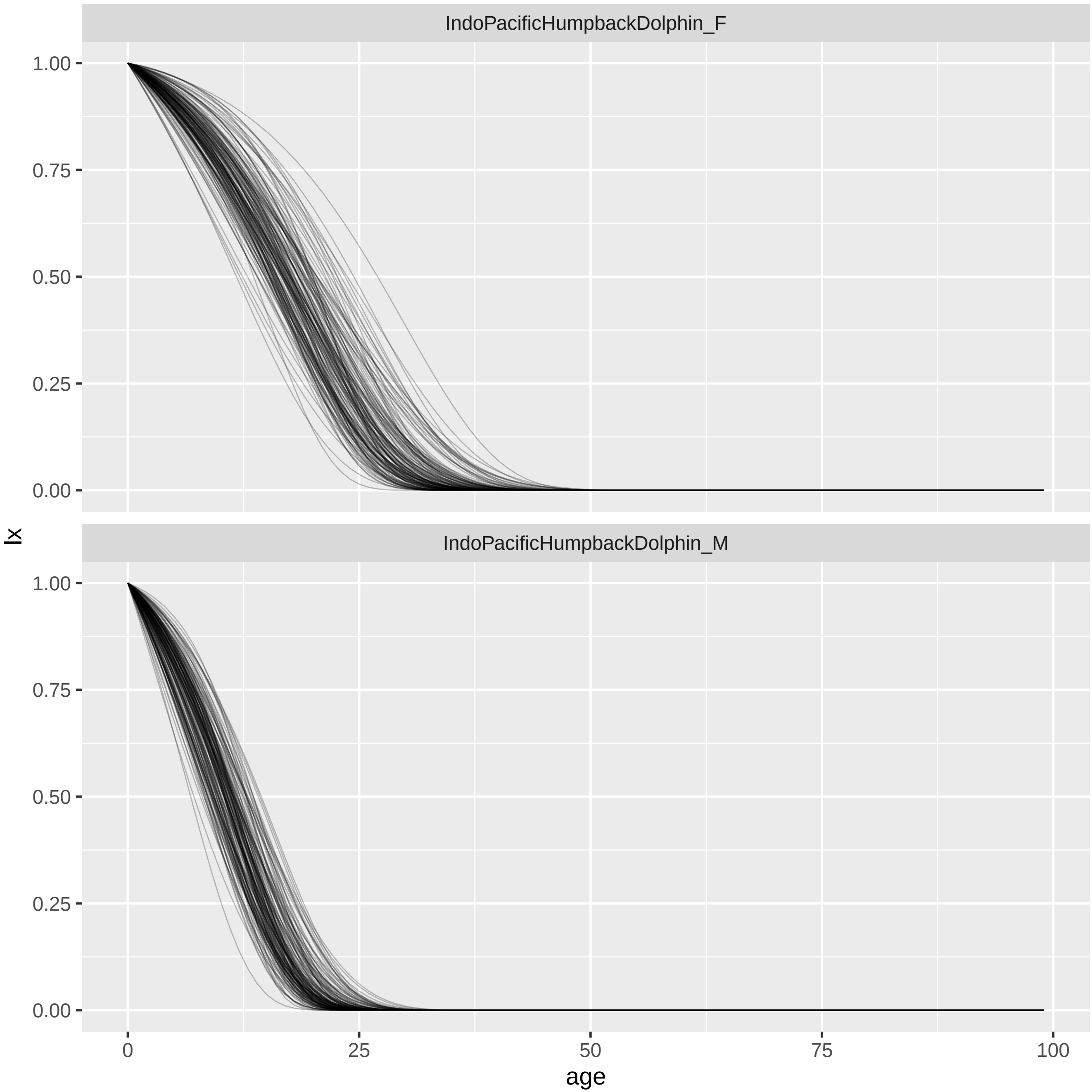
IndoPacificHumpbackDolphin_F
IndoPacificHumpbackDolphin_M
1.00
0.75
0.50
0.25
0.00
lx
0
25
50
75
100
age

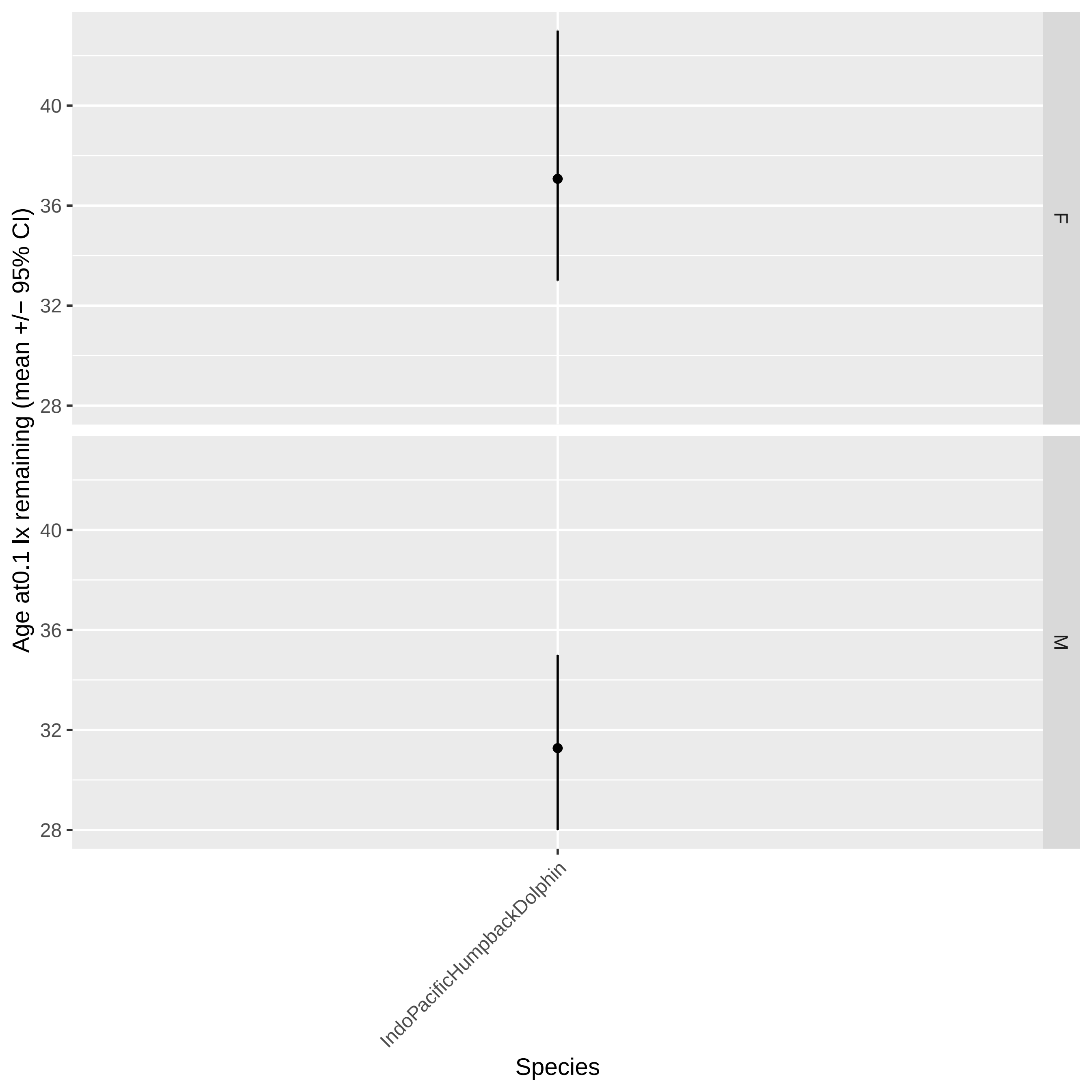

Age at0.1 lx remaining (mean +/− 95% CI)
40
36
32
28
F
40
36
32
28
M
IndoPacificHumpbackDolphin
Species

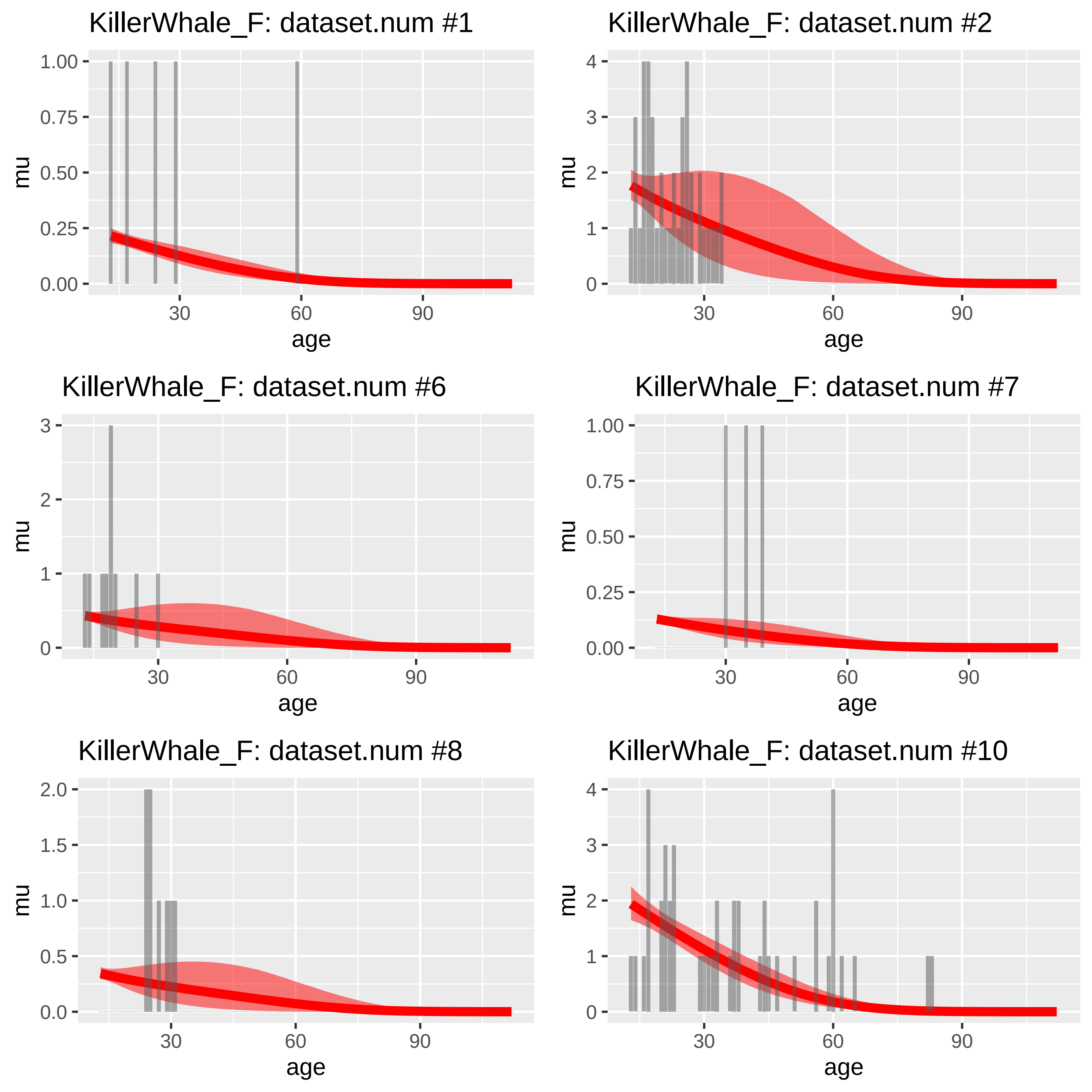
KillerWhale_F: dataset.num #1
KillerWhale_F: dataset.num #2
KillerWhale_F: dataset.num #6
KillerWhale_F: dataset.num #7
KillerWhale_F: dataset.num #8
KillerWhale_F: dataset.num #10
mu
age
0.00
0.25
0.50
0.75
1.00
0
1
2
3
4
0.0
0.5
1.0
1.5
2.0
30
60
90

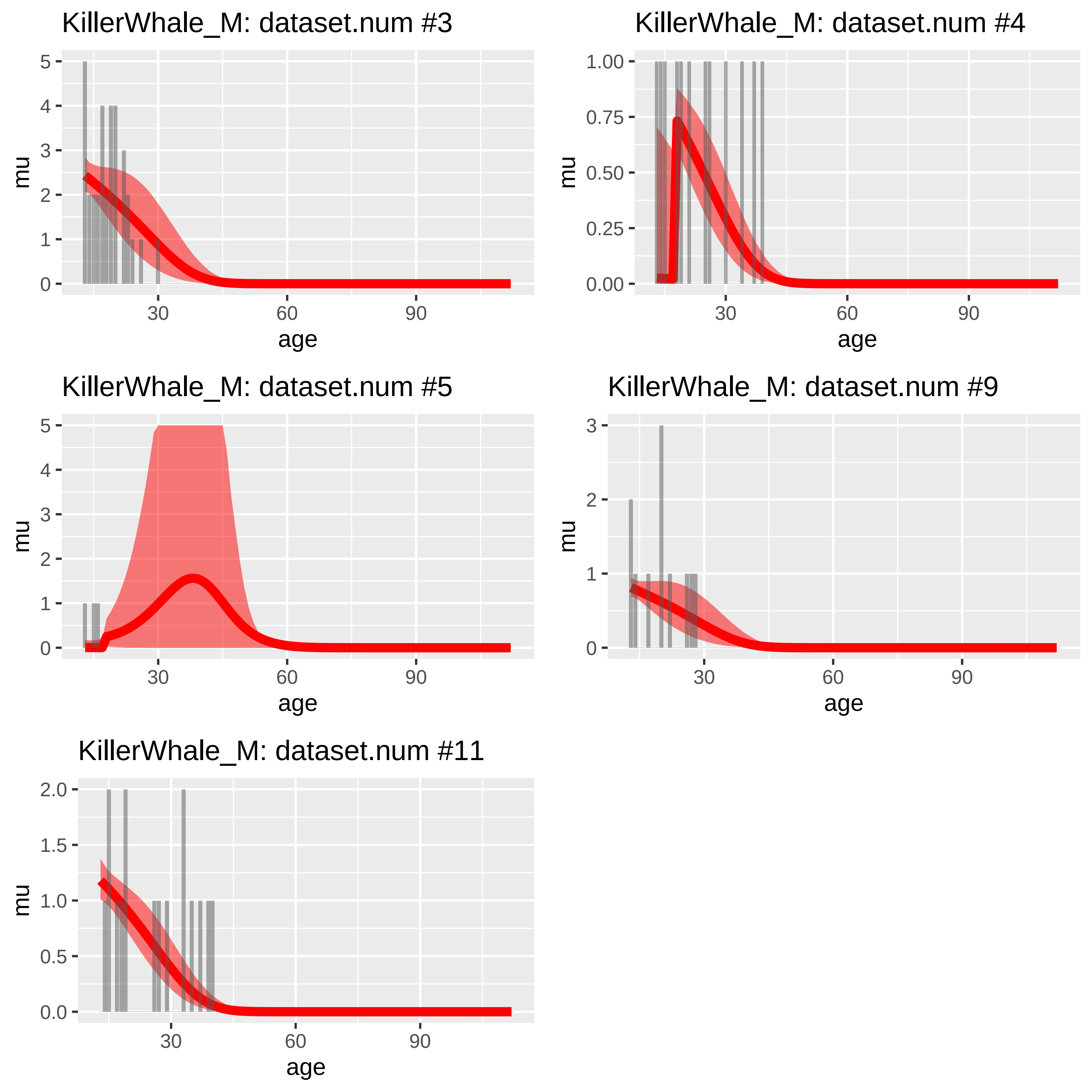

KillerWhale_M: dataset.num #3
mu
age
0
1
2
3
4
5
30
60
90
KillerWhale_M: dataset.num #4
mu
age
0.00
0.25
0.50
0.75
1.00
30
60
90
KillerWhale_M: dataset.num #5
mu
age
0
1
2
3
4
5
30
60
90
KillerWhale_M: dataset.num #9
mu
age
0
1
2
3
30
60
90
KillerWhale_M: dataset.num #11
mu
age
0.0
0.5
1.0
1.5
2.0
30
60
90

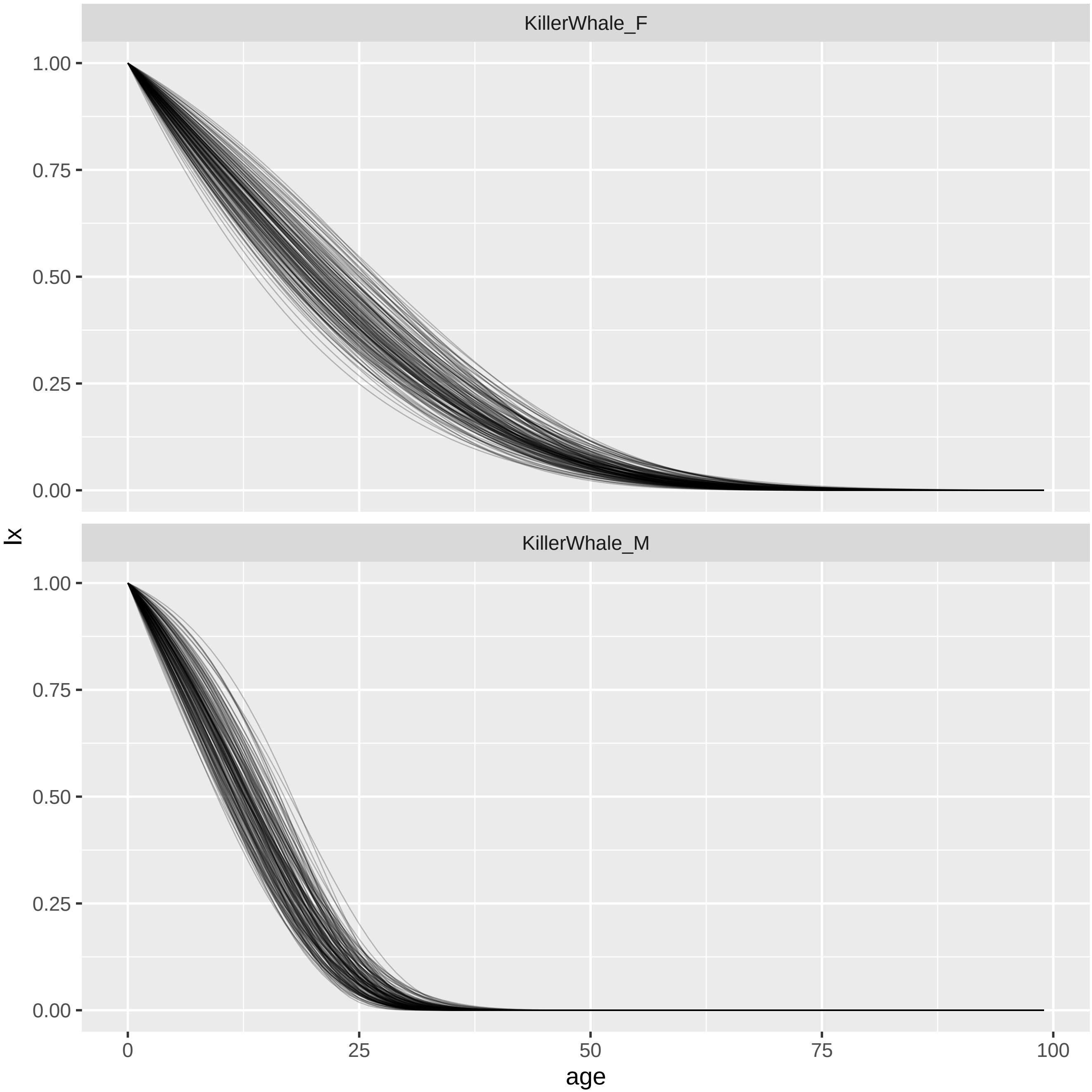

KillerWhale_F
KillerWhale_M
1.00
0.75
0.50
0.25
0.00
lx
0
25
50
75
100
age

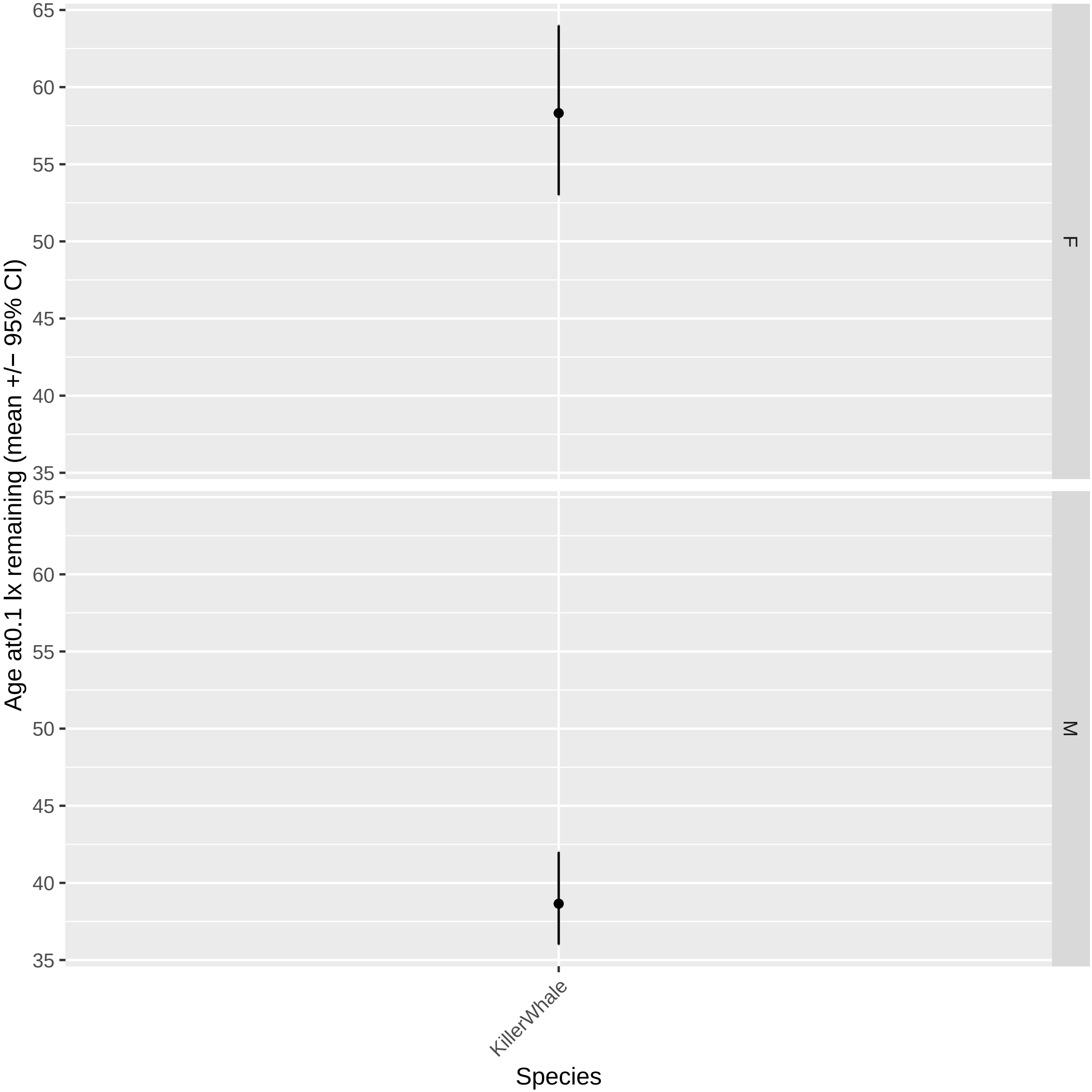

Age at0.1 lx remaining (mean +/− 95% CI)
65
60
55
50
45
40
35
F
M
KillerWhale
Species

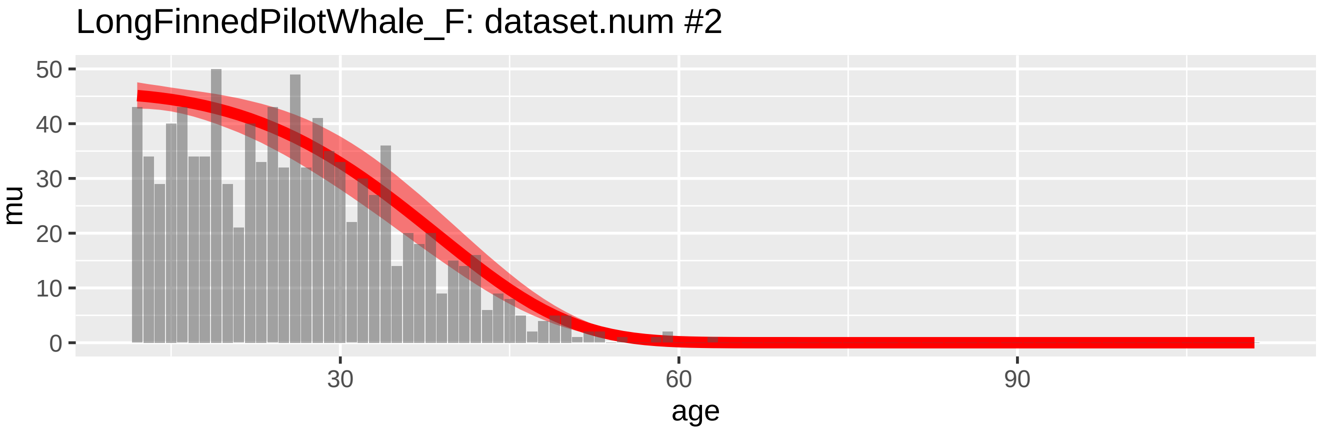
LongFinnedPilotWhale_F: dataset.num #2
mu
0
10
20
30
40
50
30
60
90
age

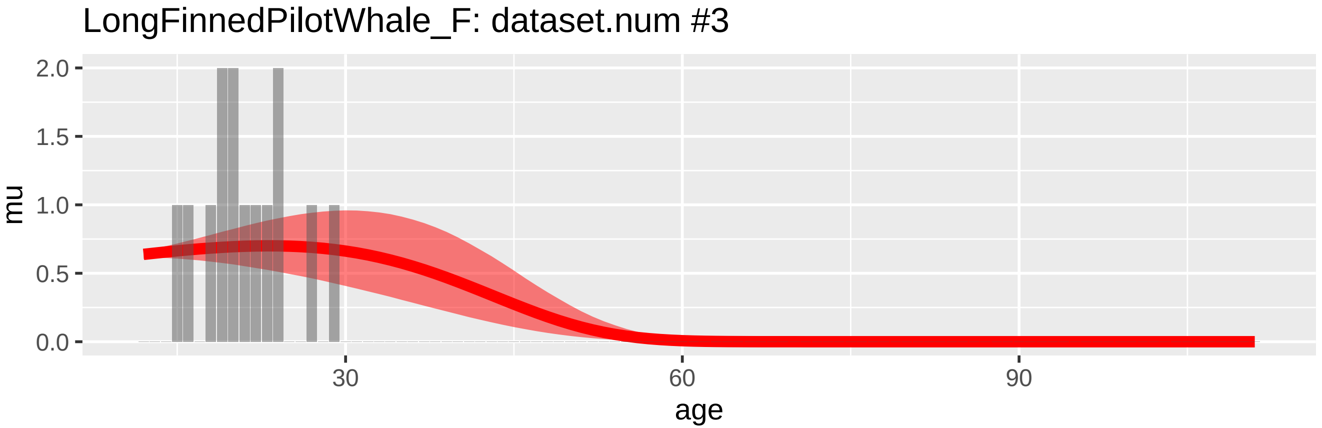
LongFinnedPilotWhale_F: dataset.num #3
mu
0.0
0.5
1.0
1.5
2.0
30
60
90
age

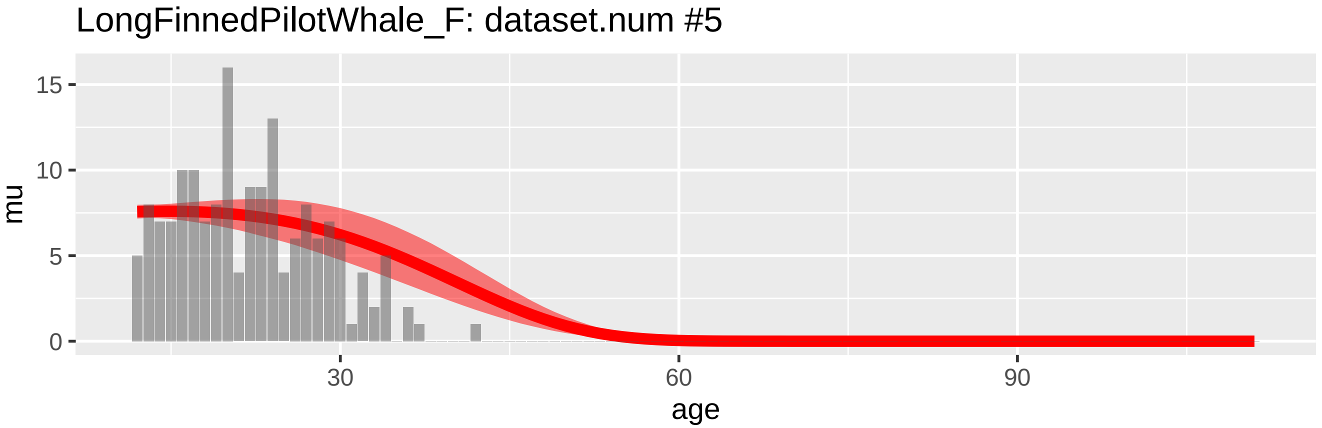
LongFinnedPilotWhale_F: dataset.num #5
mu
0
5
10
15
30
60
90
age

LongFinnedPilotWhale_M: dataset.num #1

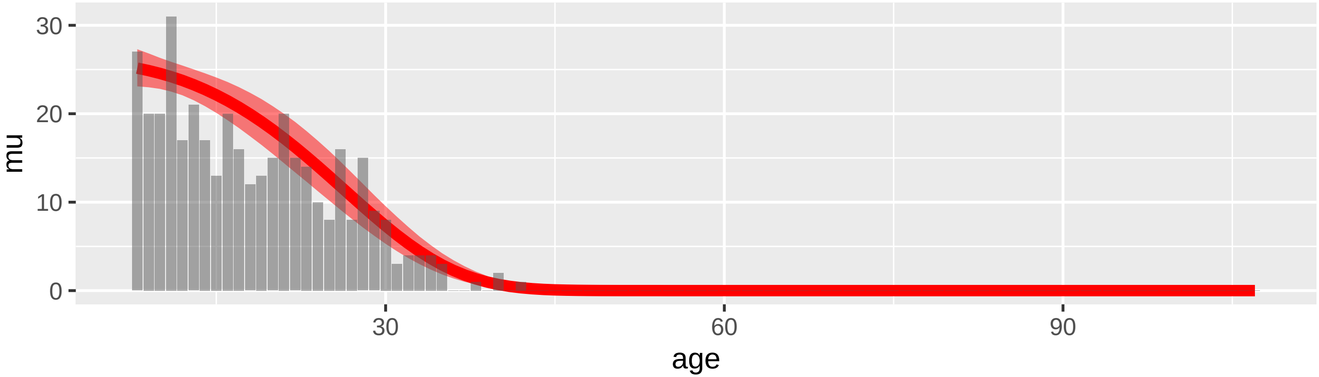


LongFinnedPilotWhale_M: dataset.num #4

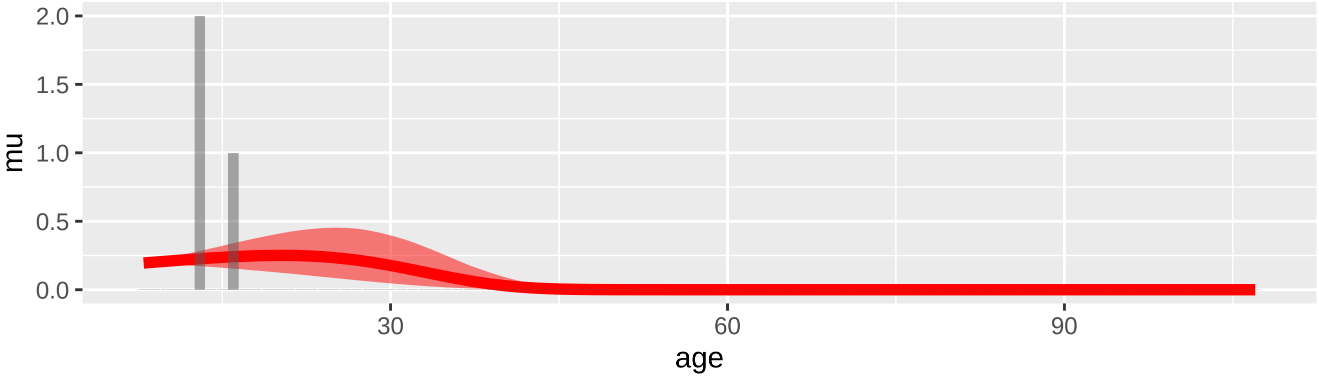


LongFinnedPilotWhale_M: dataset.num #6

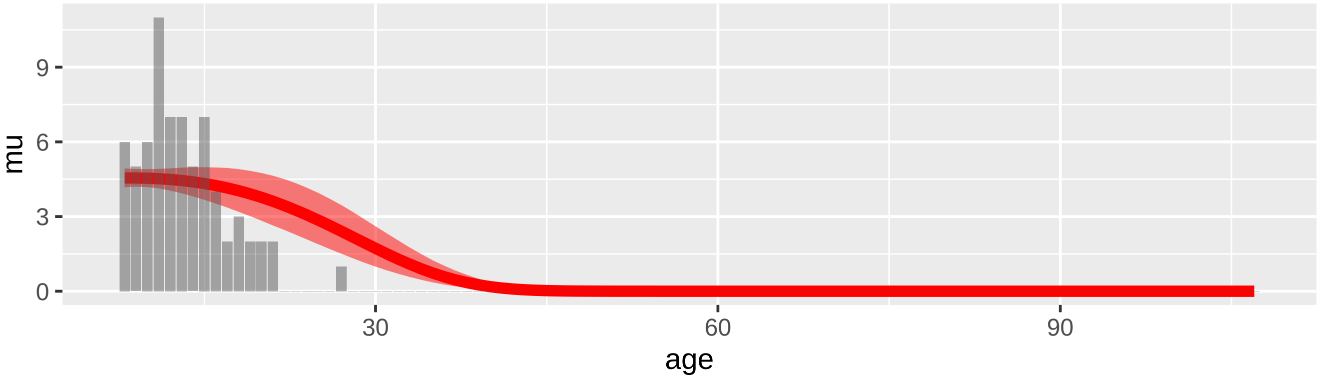

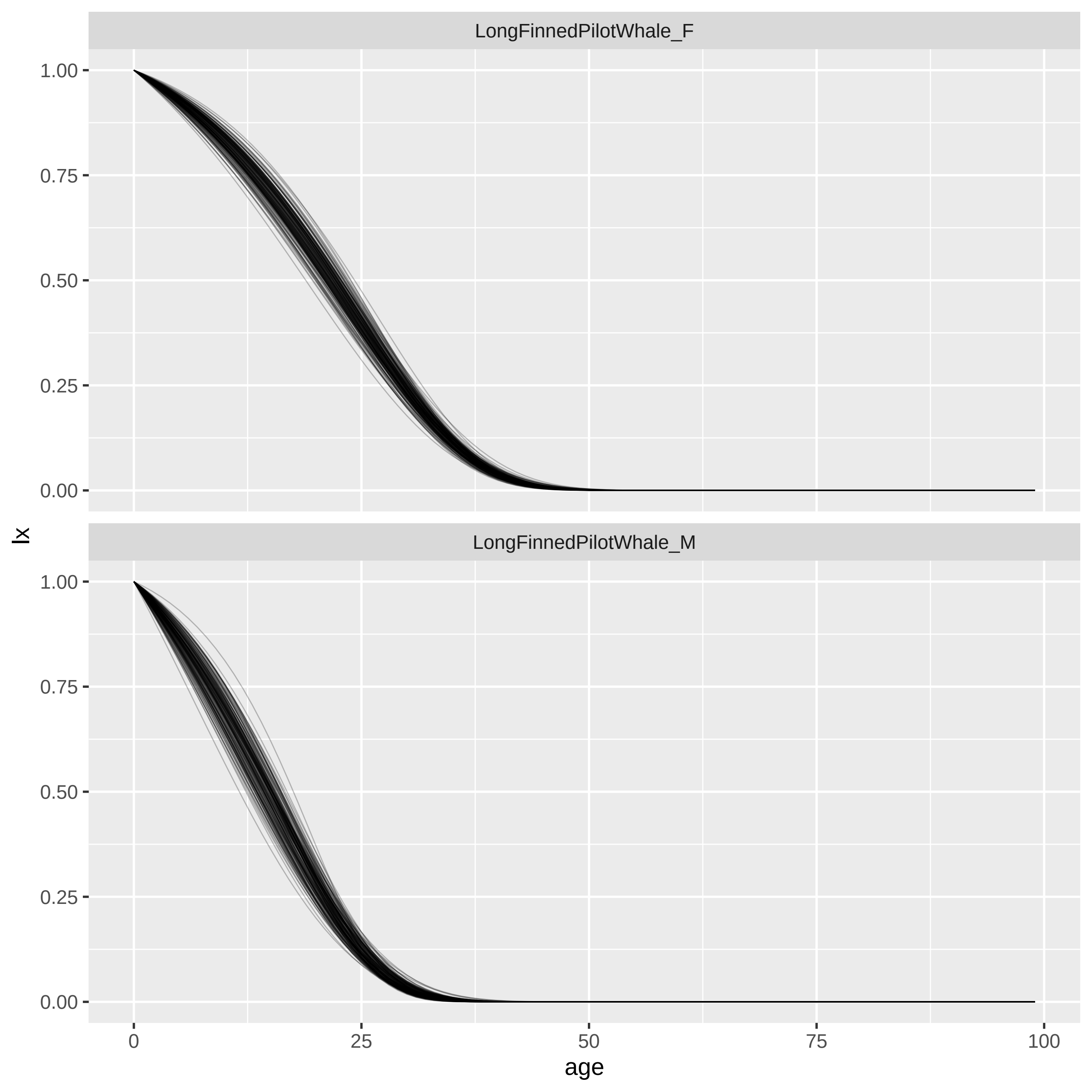

LongFinnedPilotWhale_F
LongFinnedPilotWhale_M
lx
age
1.00
0.75
0.50
0.25
0.00
0
25
50
75
100

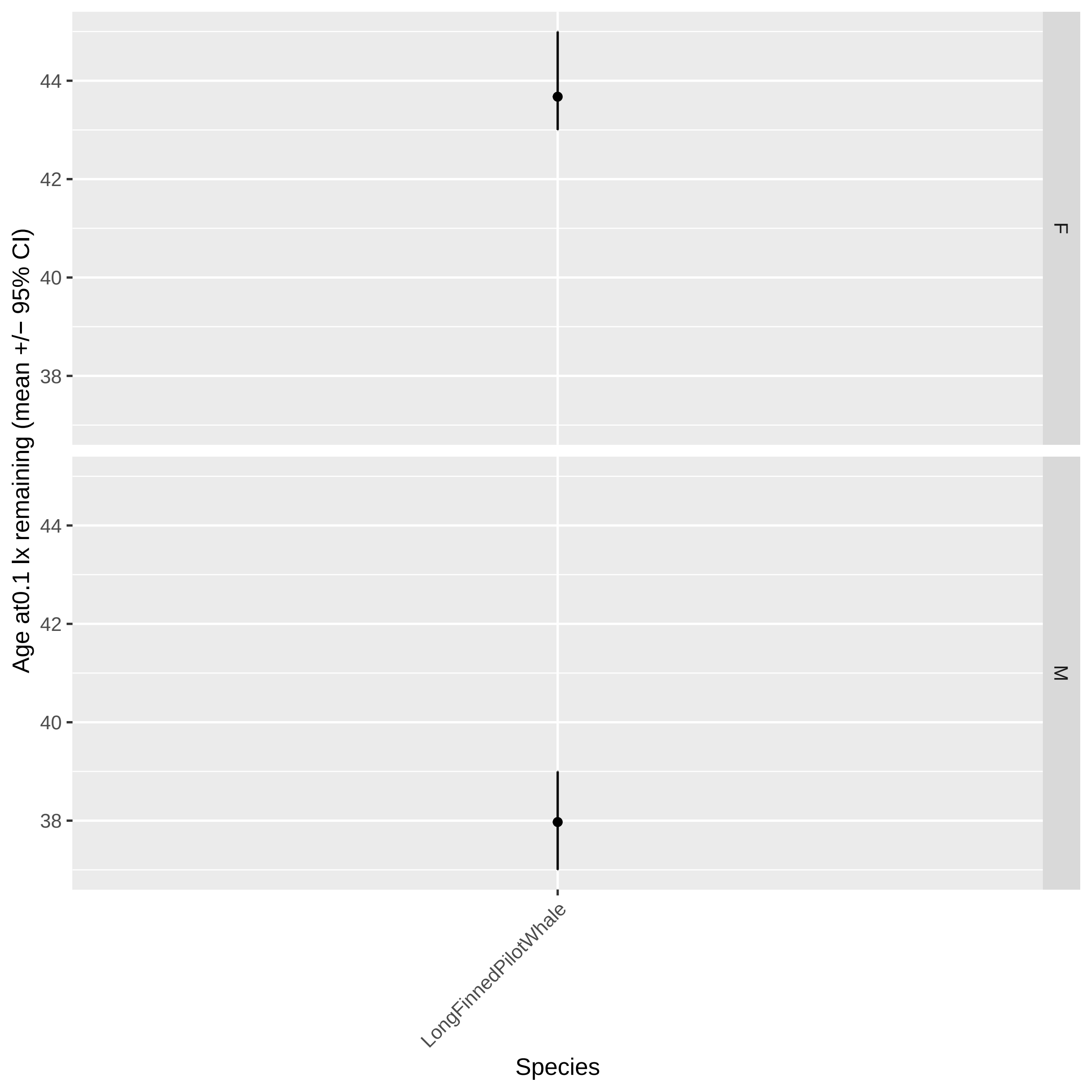

Age at0.1 lx remaining (mean +/− 95% CI)
44
42
40
38
F
M
LongFinnedPilotWhale
Species

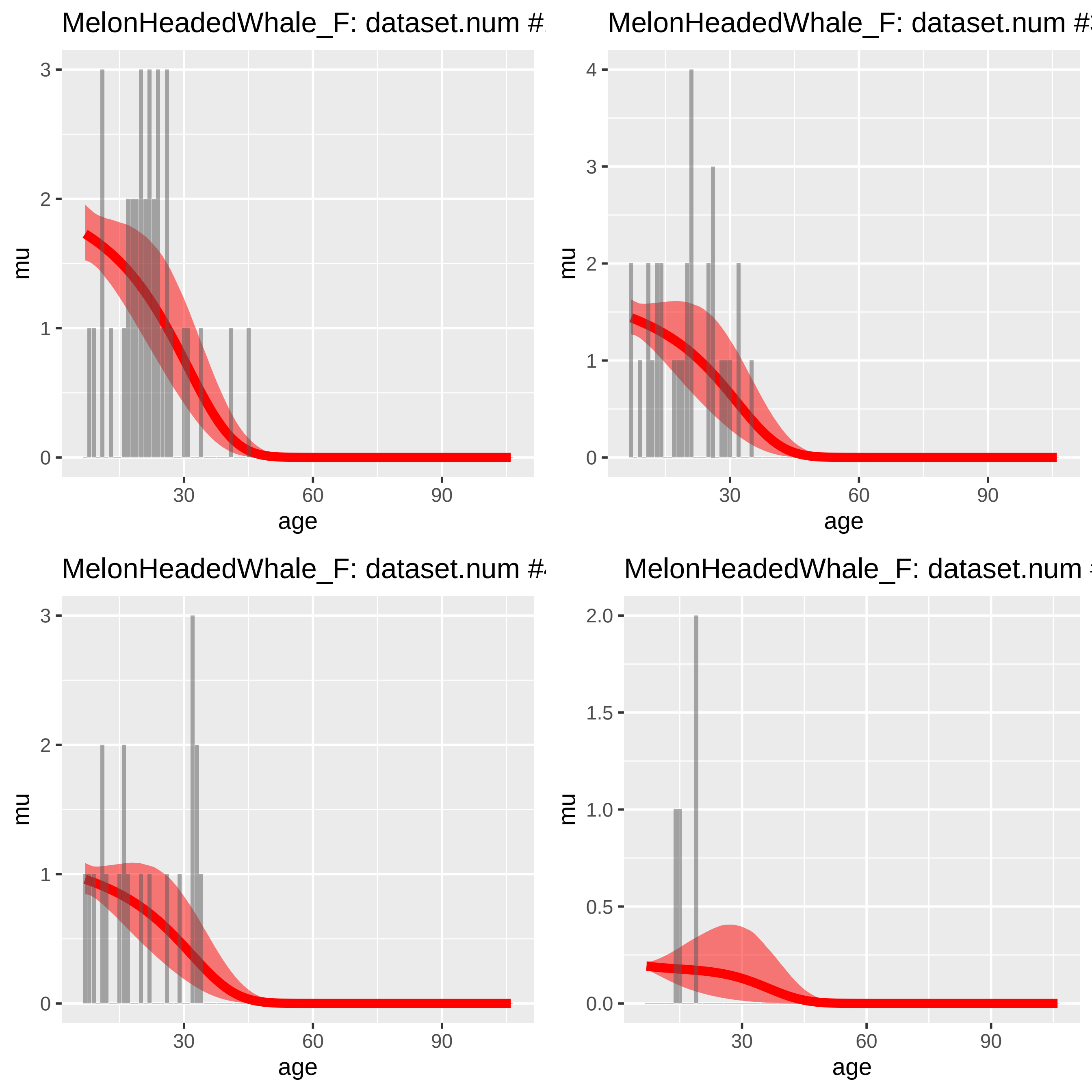

MelonHeadedWhale_F: dataset.num #
MelonHeadedWhale_F: dataset.num #
MelonHeadedWhale_F: dataset.num #
MelonHeadedWhale_F: dataset.num
mu
age

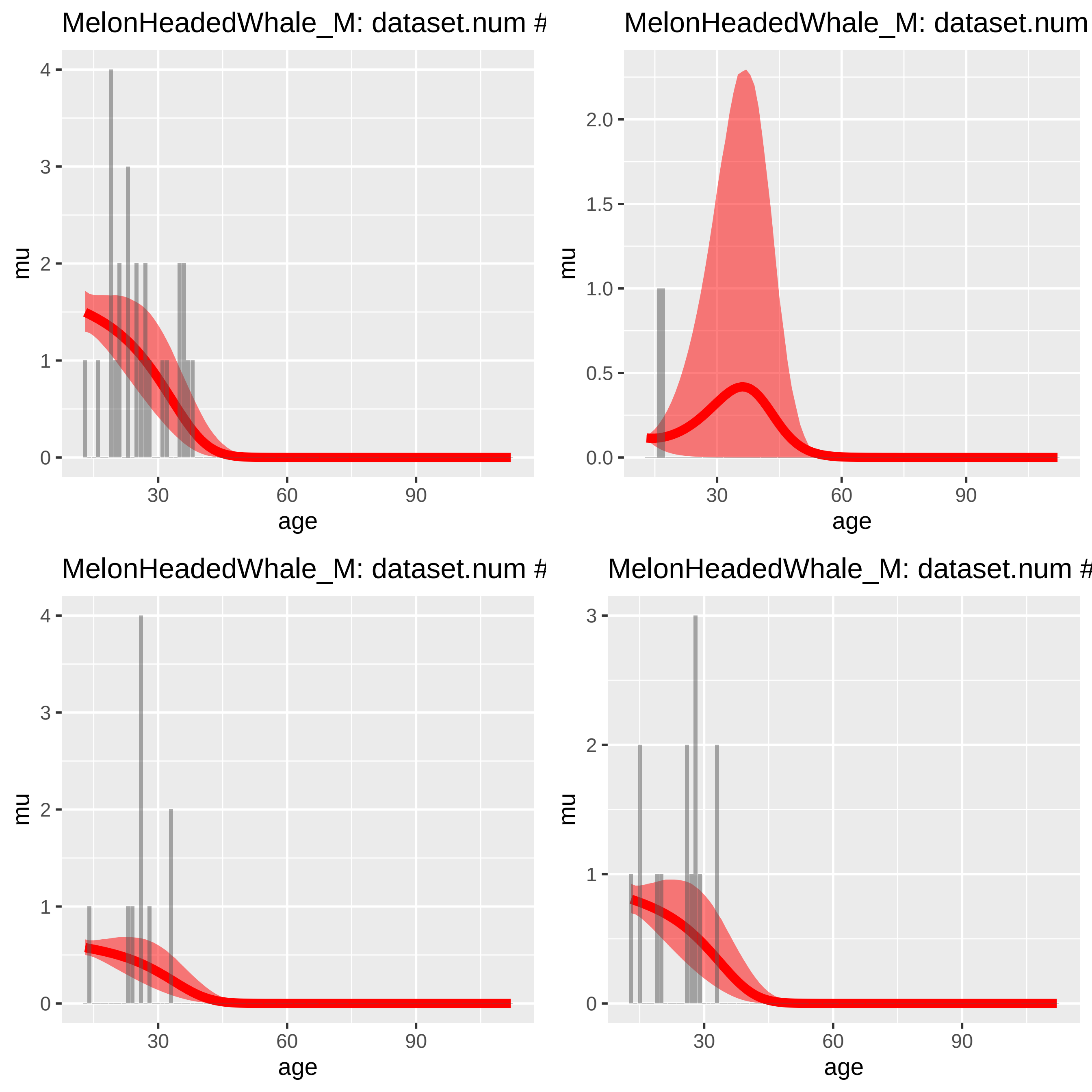

MelonHeadedWhale_M: dataset.num #
MelonHeadedWhale_M: dataset.num
MelonHeadedWhale_M: dataset.num #
MelonHeadedWhale_M: dataset.num #
mu
age
0
1
2
3
4
0.0
0.5
1.0
1.5
2.0
30
60
90

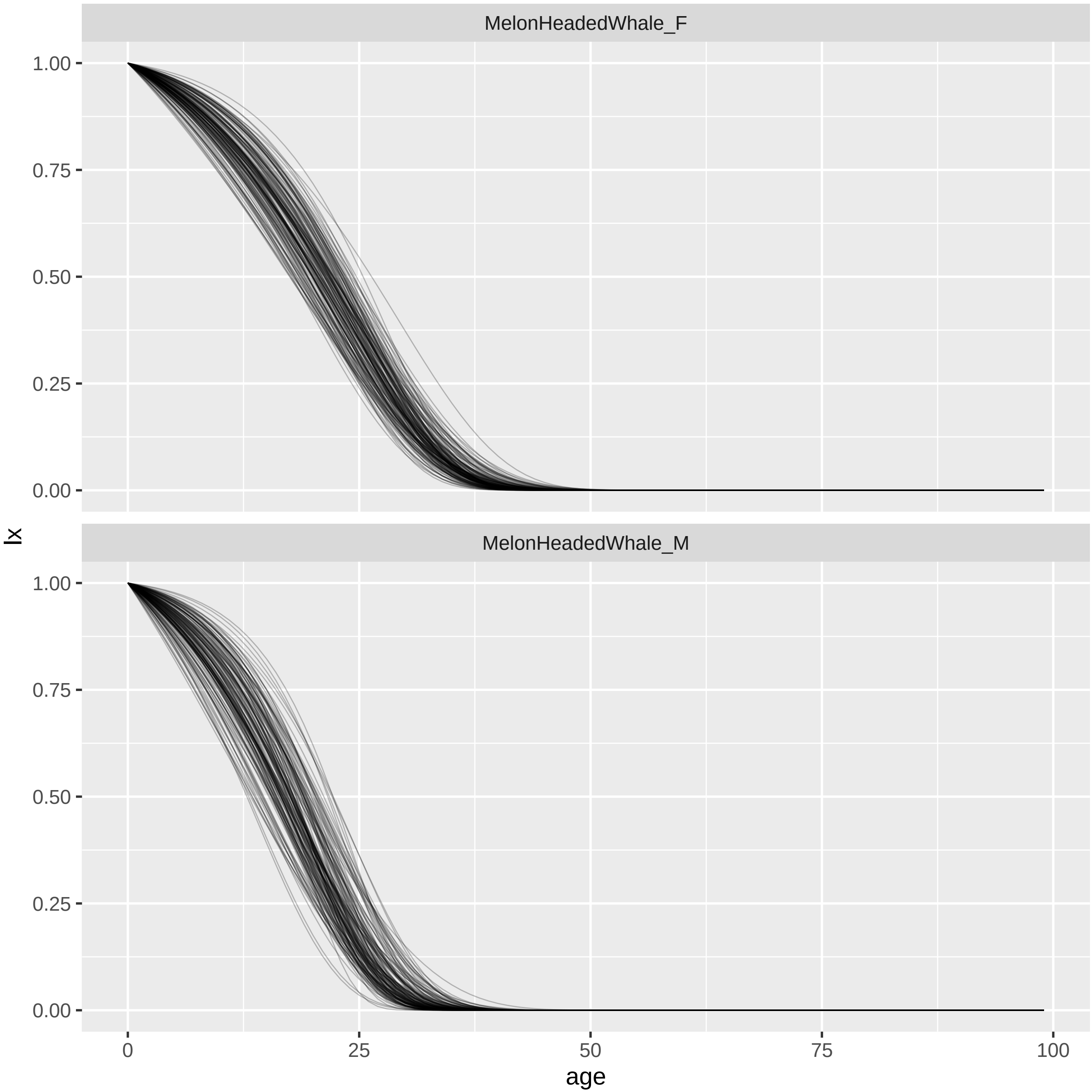
MelonHeadedWhale_F
MelonHeadedWhale_M
lx
1.00
0.75
0.50
0.25
0.00
age
0
25
50
75
100

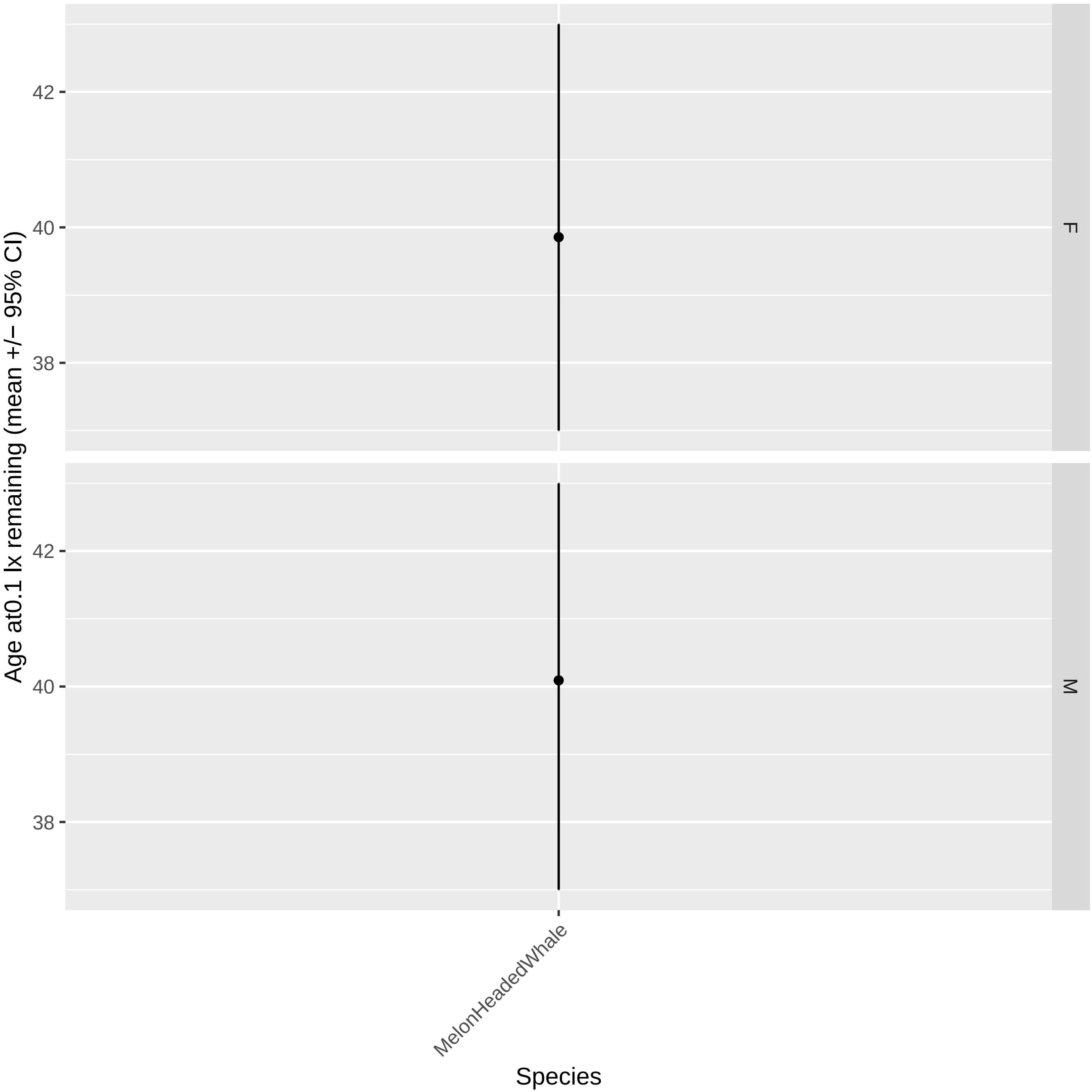

Age at0.1 lx remaining (mean +/− 95% CI)
42
40
38
F
42
40
38
M
MelonHeadedWhale
Species

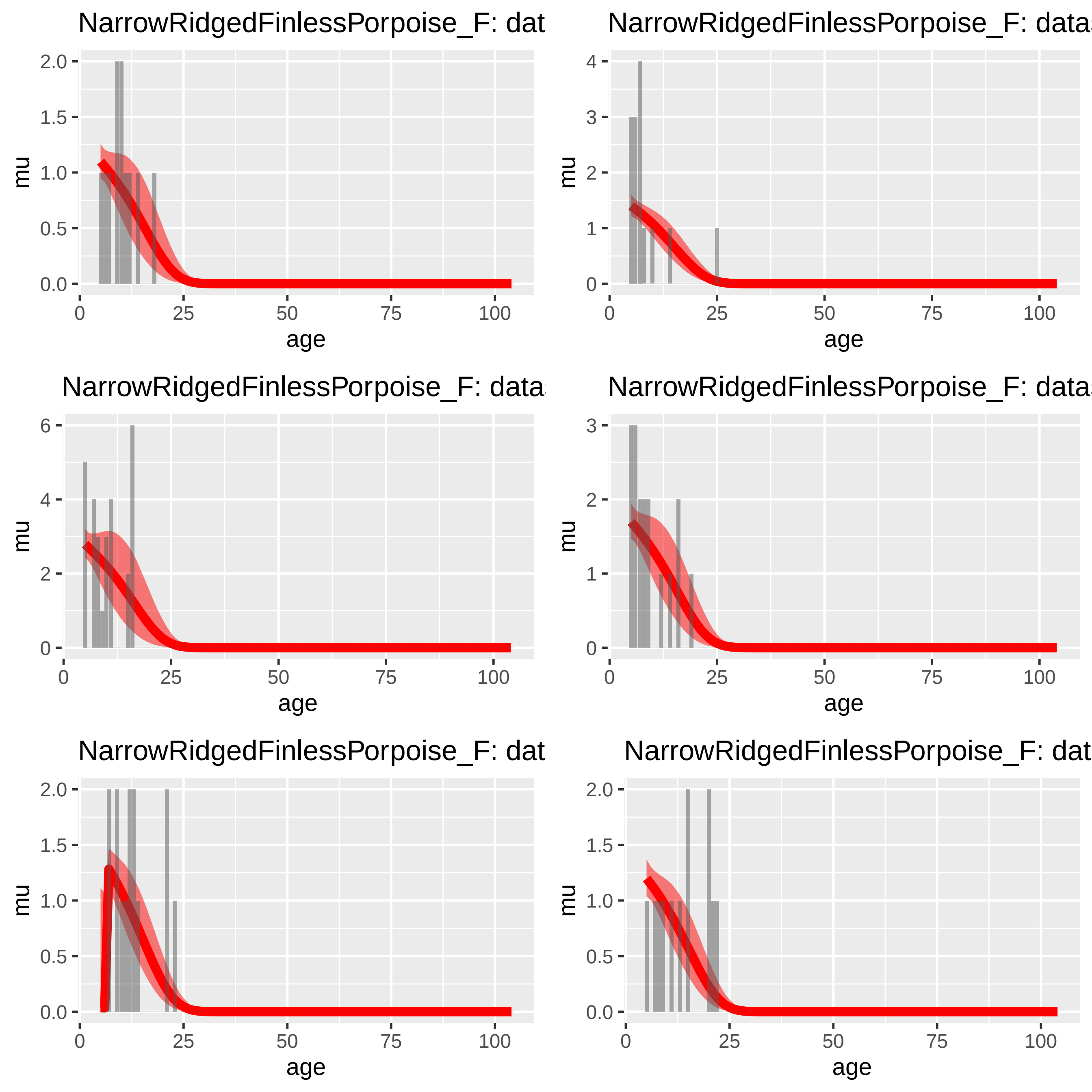
NarrowRidgedFinlessPorpoise_F: dat
NarrowRidgedFinlessPorpoise_F: data
NarrowRidgedFinlessPorpoise_F: data
NarrowRidgedFinlessPorpoise_F: data
NarrowRidgedFinlessPorpoise_F: dat
NarrowRidgedFinlessPorpoise_F: dat
mu
age
0
25
50
75
100

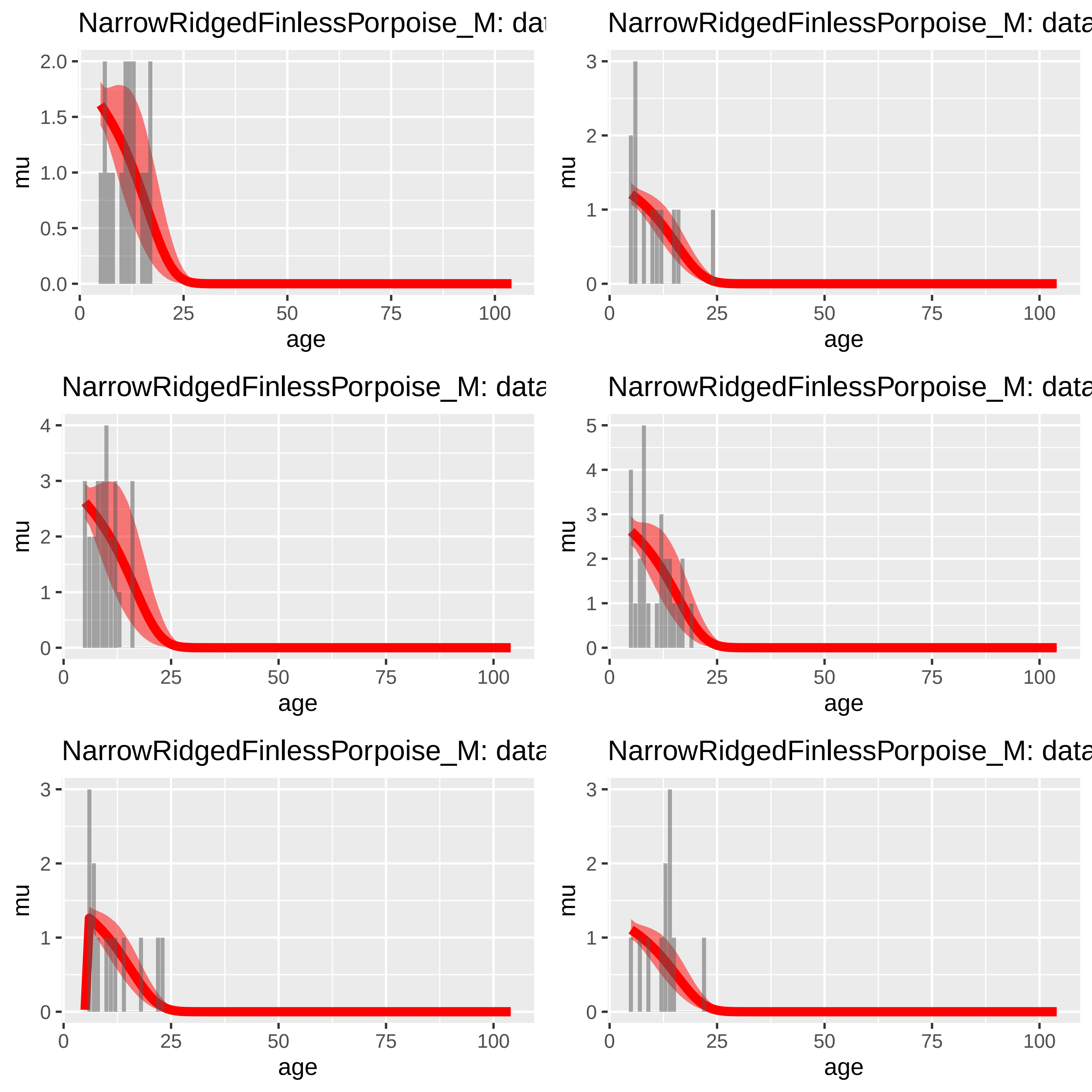
NarrowRidgedFinlessPorpoise_M: dat
NarrowRidgedFinlessPorpoise_M: data
NarrowRidgedFinlessPorpoise_M: data
NarrowRidgedFinlessPorpoise_M: data
NarrowRidgedFinlessPorpoise_M: data
NarrowRidgedFinlessPorpoise_M: data
mu
age
0
25
50
75
100
0.0
0.5
1.0
1.5
2.0
1
2
3
4
5

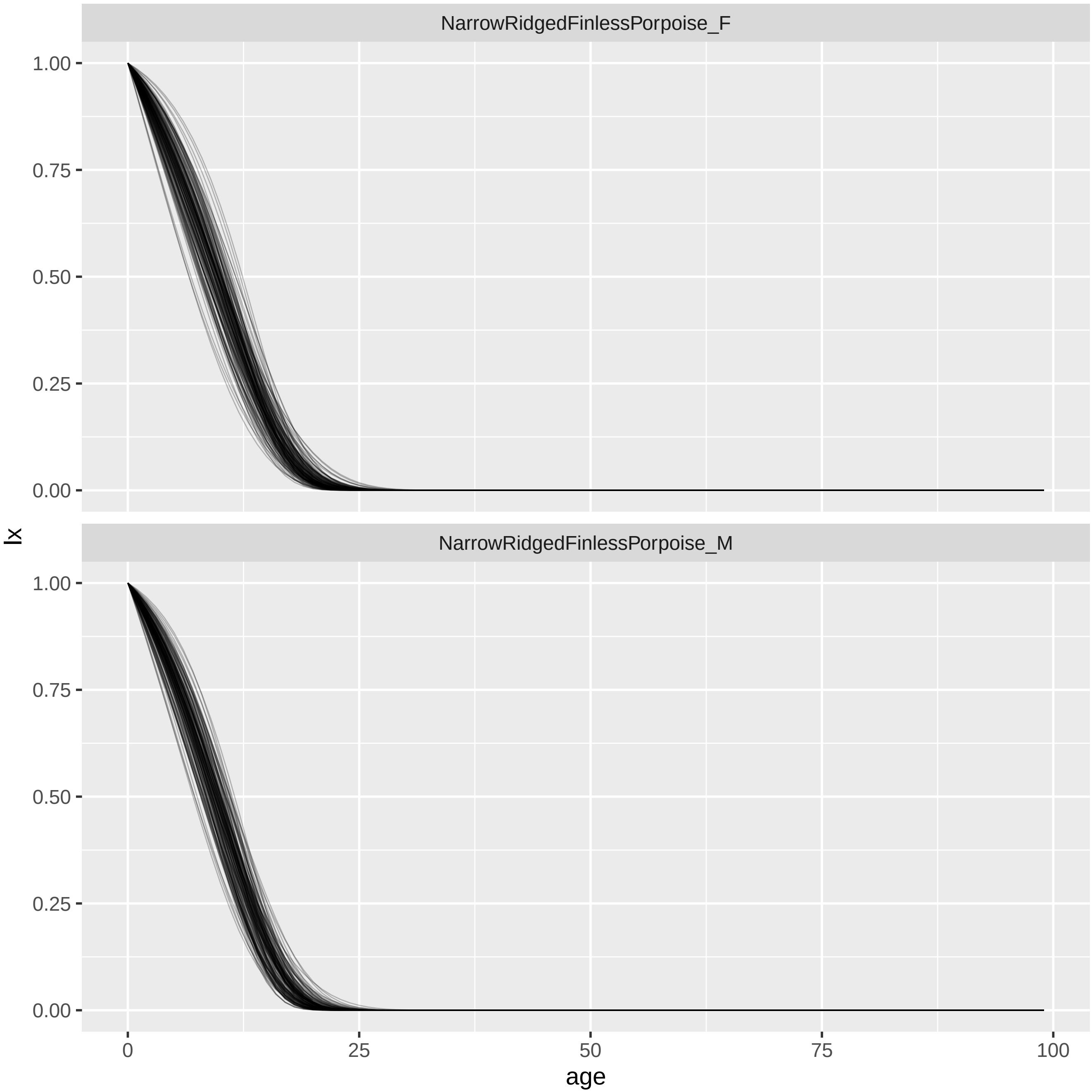

NarrowRidgedFinlessPorpoise_F
NarrowRidgedFinlessPorpoise_M
1.00
0.75
0.50
0.25
0.00
0
25
50
75
100
lx
age

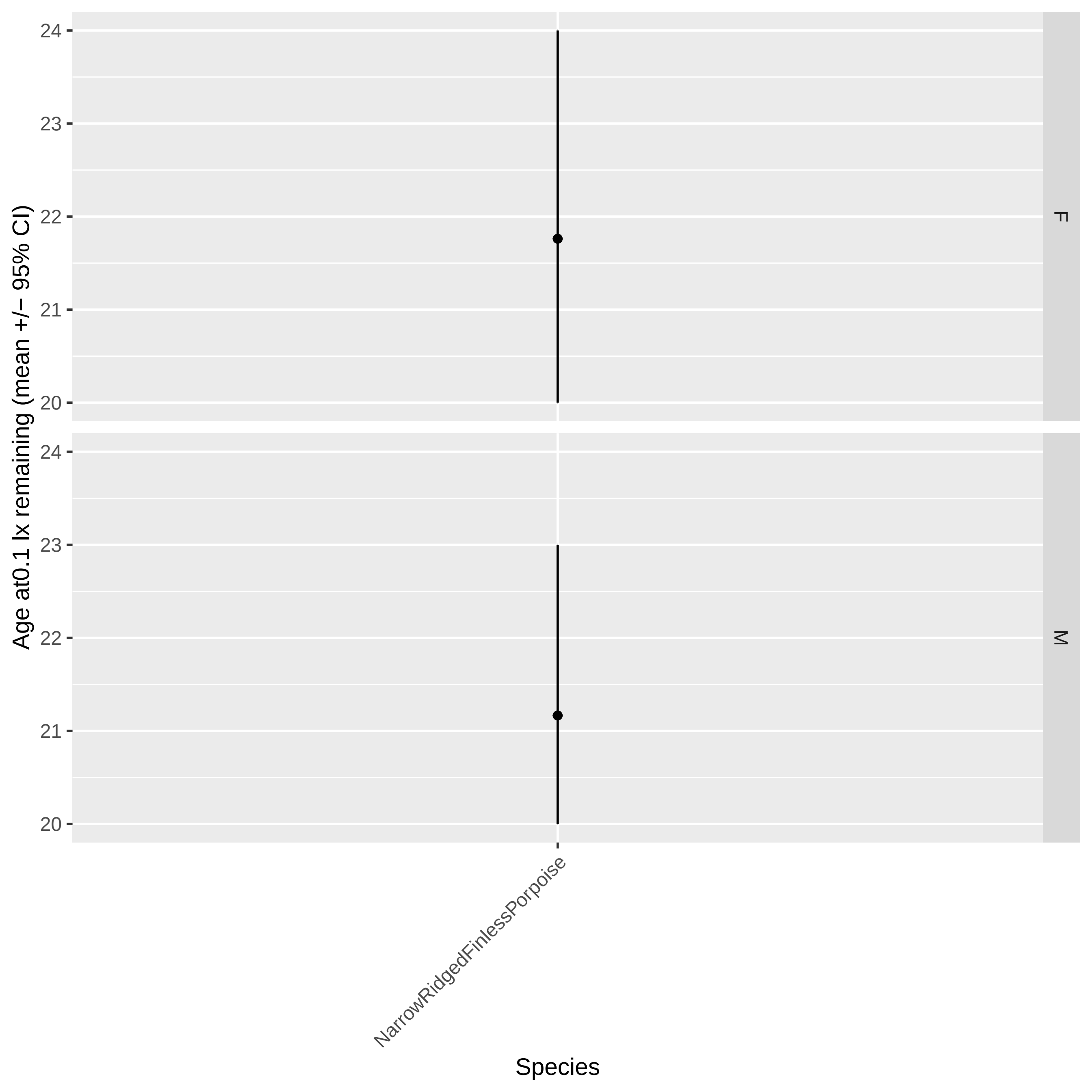
Age at0.1 lx remaining (mean +/− 95% CI)
24
23
22
21
20
F
M
NarrowRidgedFinlessPorpoise
Species

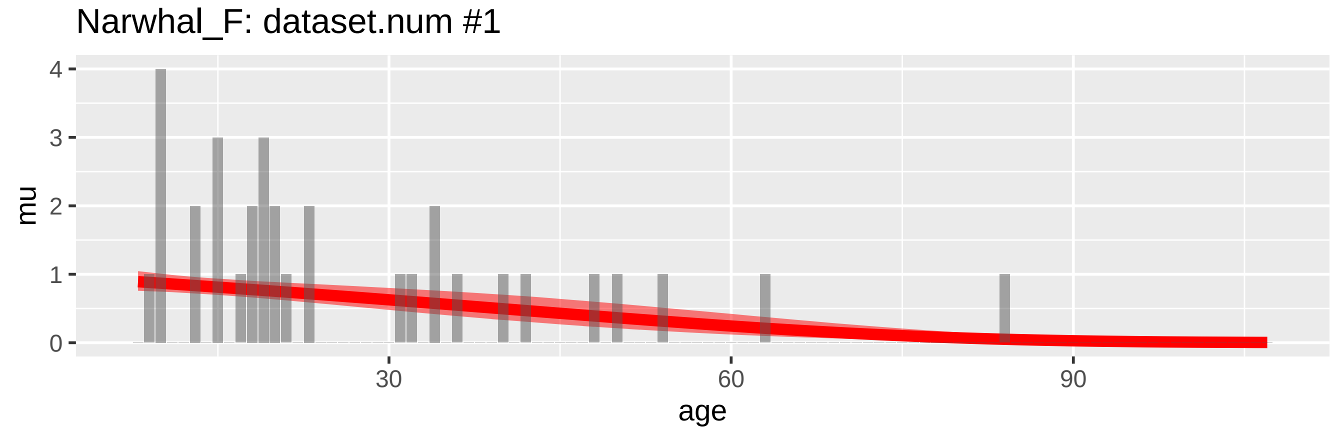
Narwhal_F: dataset.num #1
mu
0
1
2
3
4
30
60
90
age

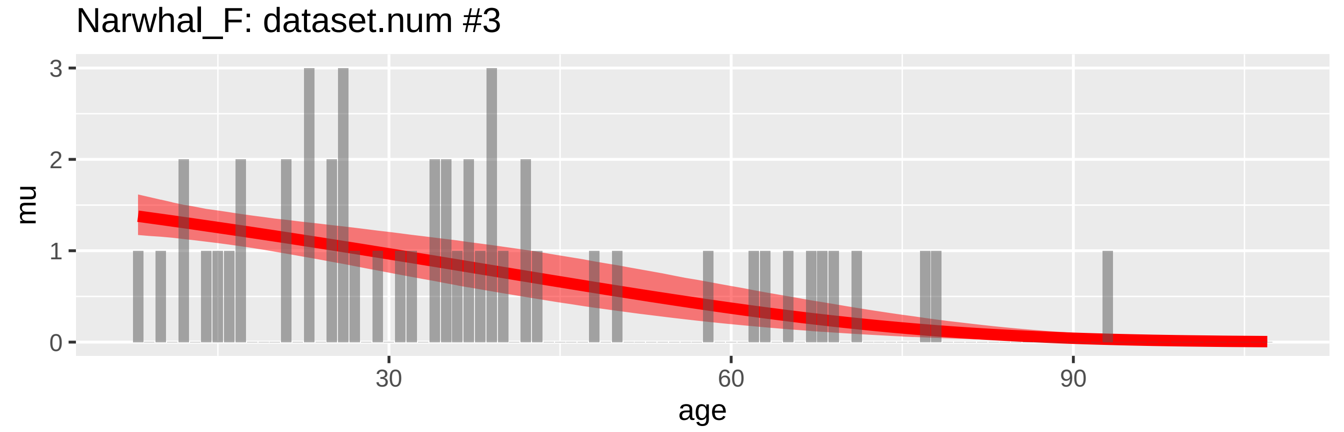
Narwhal_F: dataset.num #3
mu
0
1
2
3
30
60
90
age

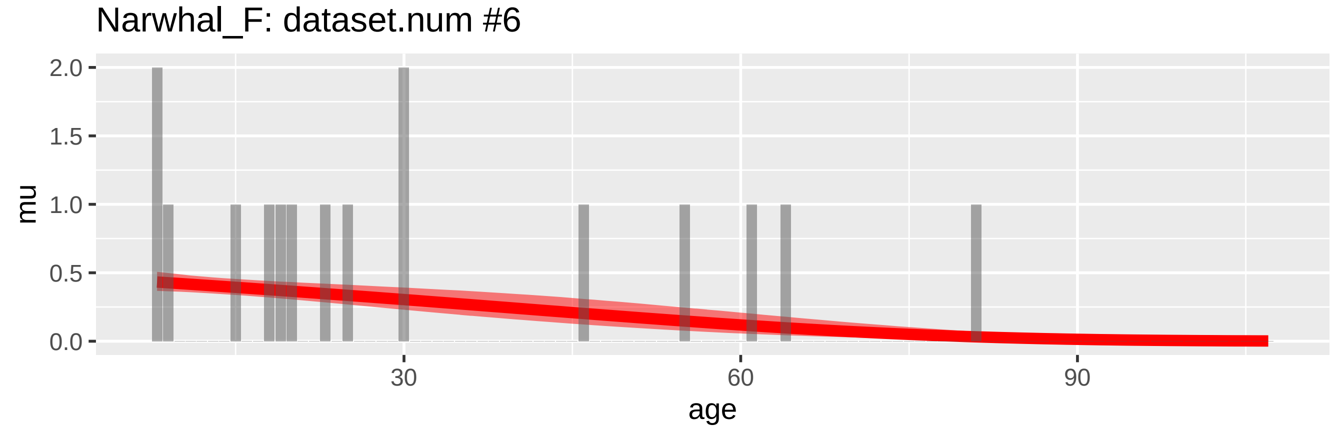
Narwhal_F: dataset.num #6
mu
0.0
0.5
1.0
1.5
2.0
30
60
90
age

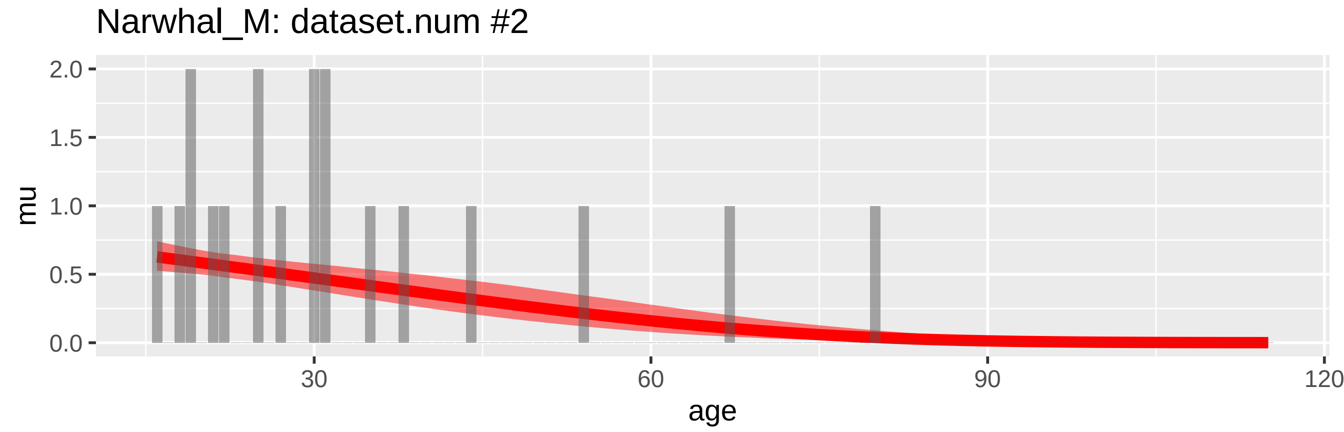
Narwhal_M: dataset.num #2
mu
2.0
1.5
1.0
0.5
0.0
30
60
90
120
age

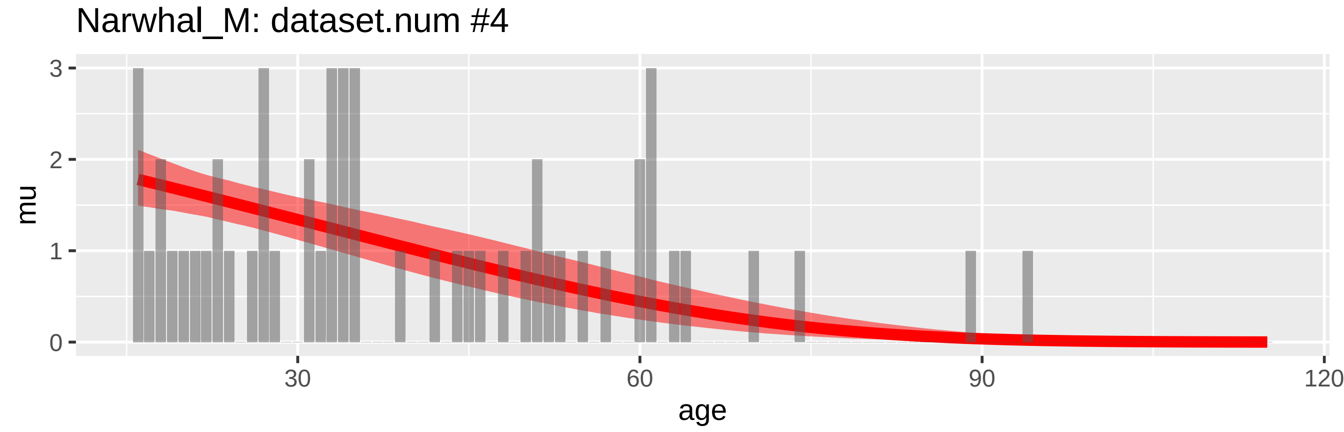
Narwhal_M: dataset.num #4
mu
3
2
1
0
30
60
90
120
age

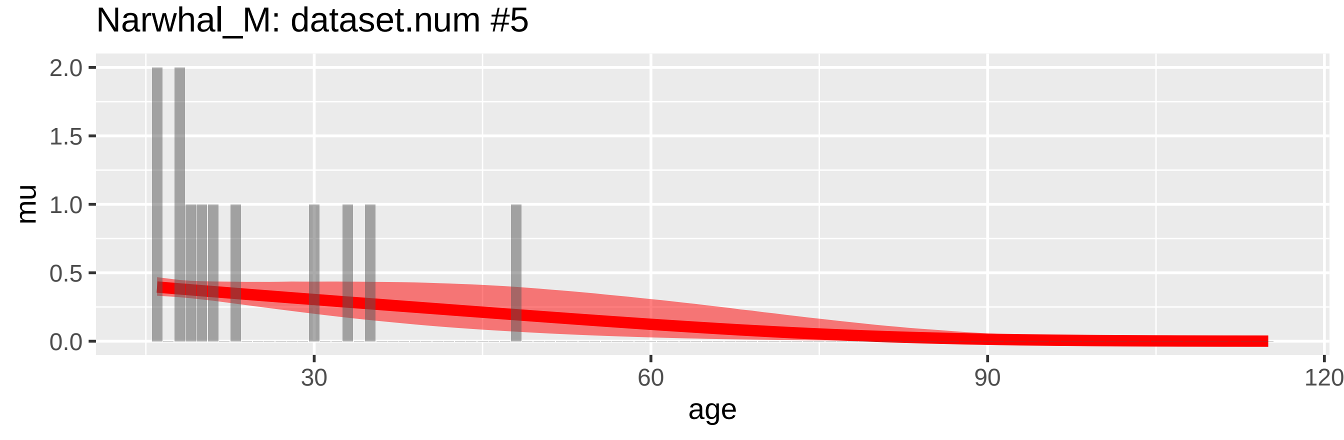
Narwhal_M: dataset.num #5
mu
2.0
1.5
1.0
0.5
0.0
30
60
90
120
age

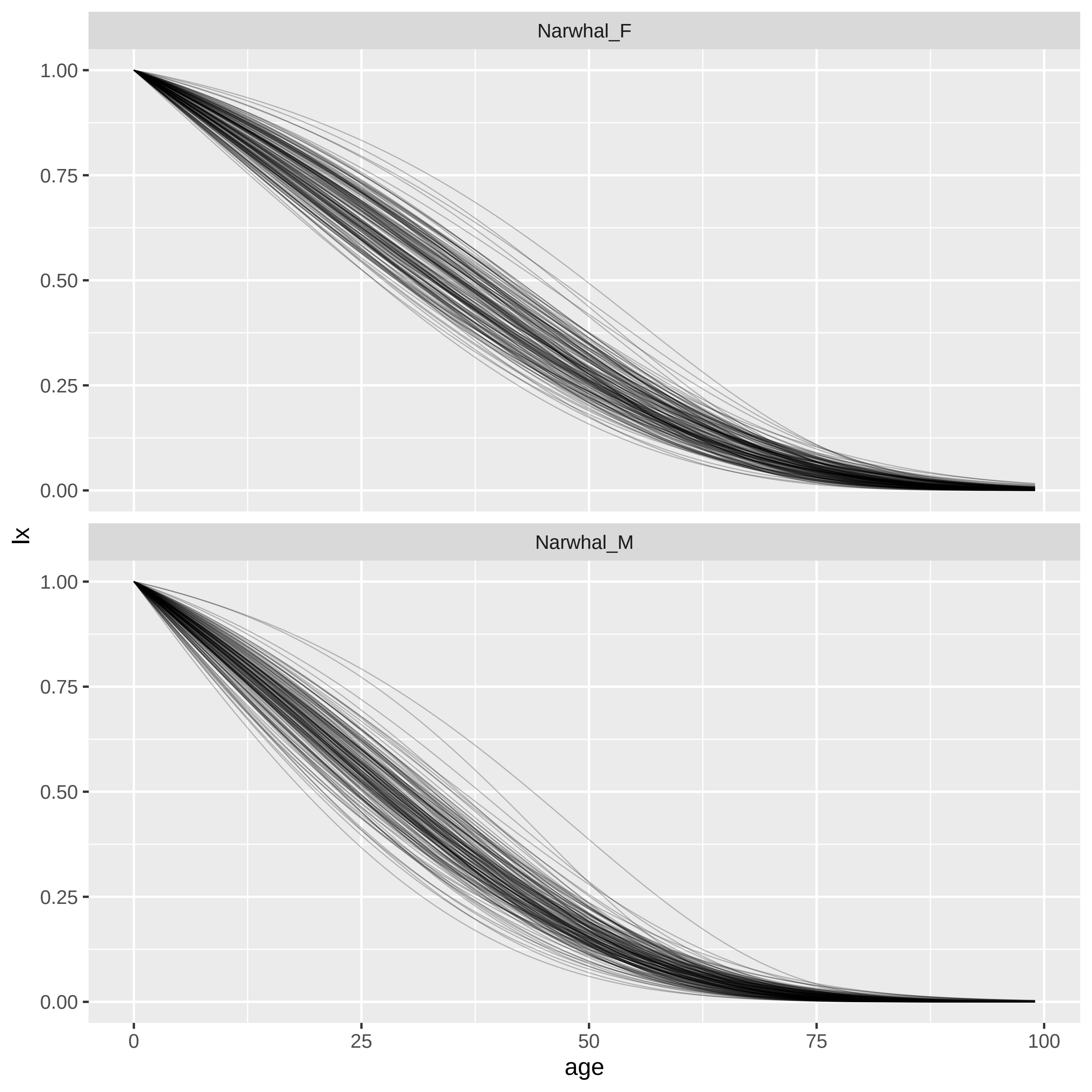
Narwhal_F
Narwhal_M
lx
age
1.00
0.75
0.50
0.25
0.00
0
25
50
75
100

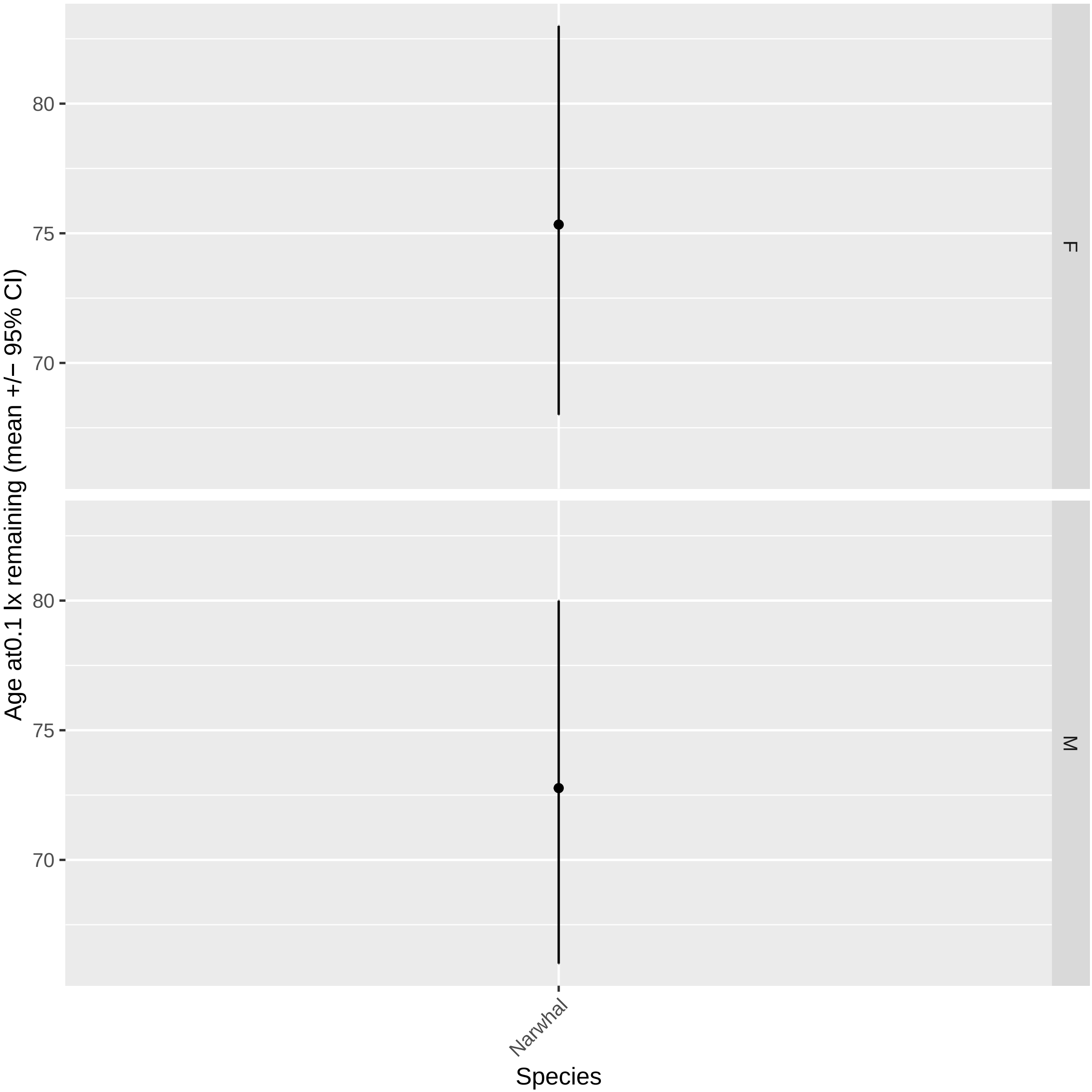
Age at0.1 lx remaining (mean +/− 95% CI)
80
75
70
F
80
75
70
M
Narwhal
Species

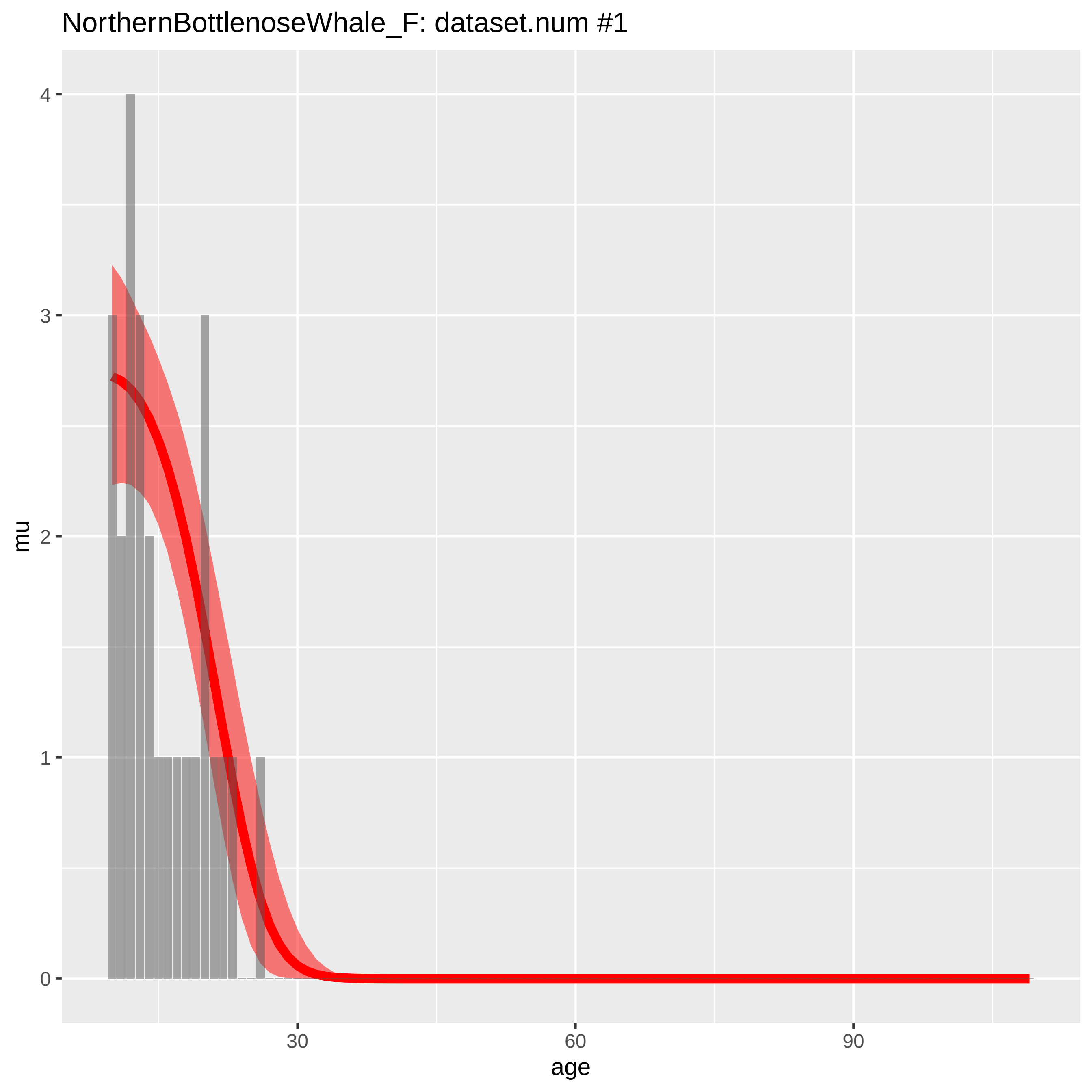

NorthernBottlenoseWhale_F: dataset.num #1
mu
4
3
2
1
0
30
60
90
age

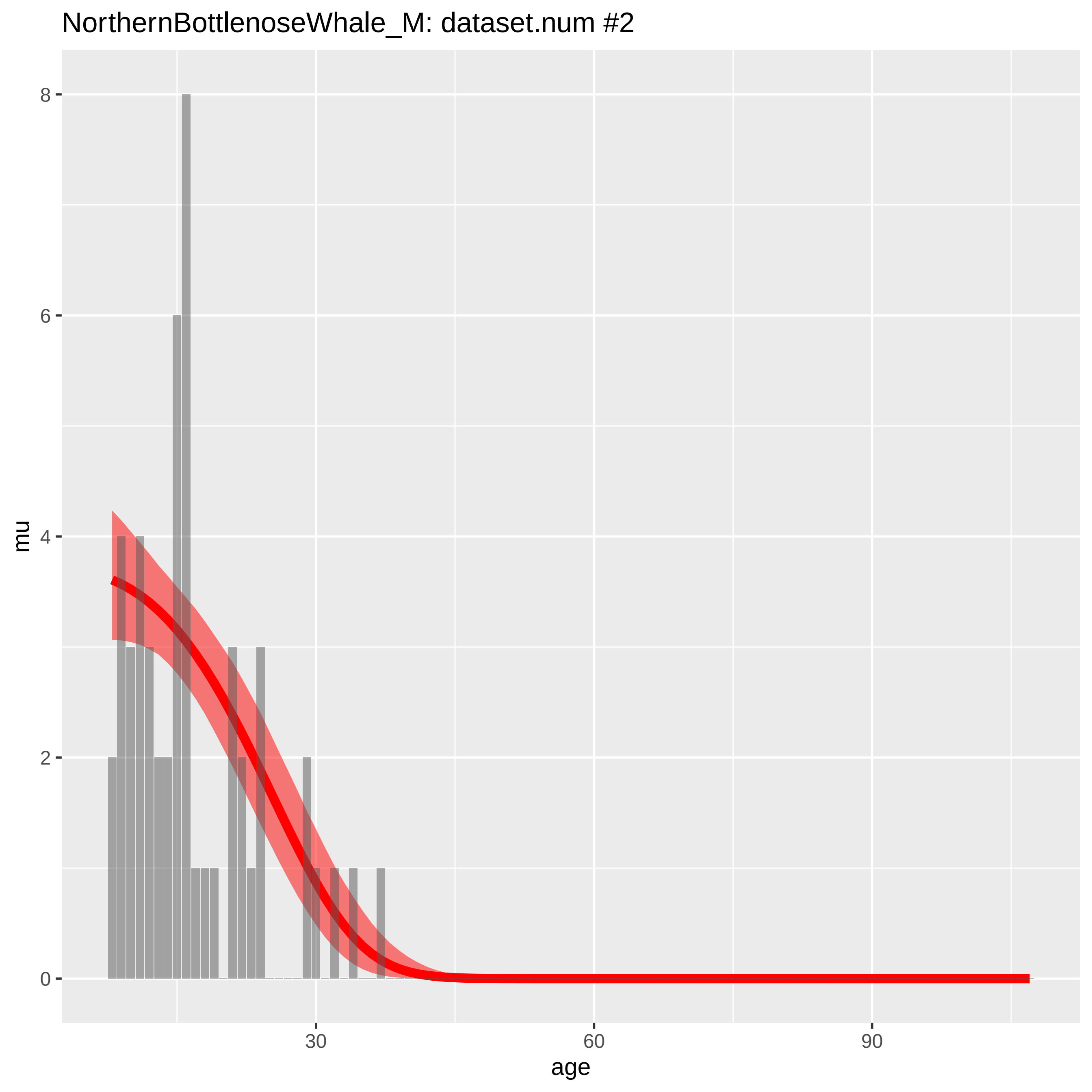

NorthernBottlenoseWhale_M: dataset.num #2
mu
0
2
4
6
8
30
60
90
age

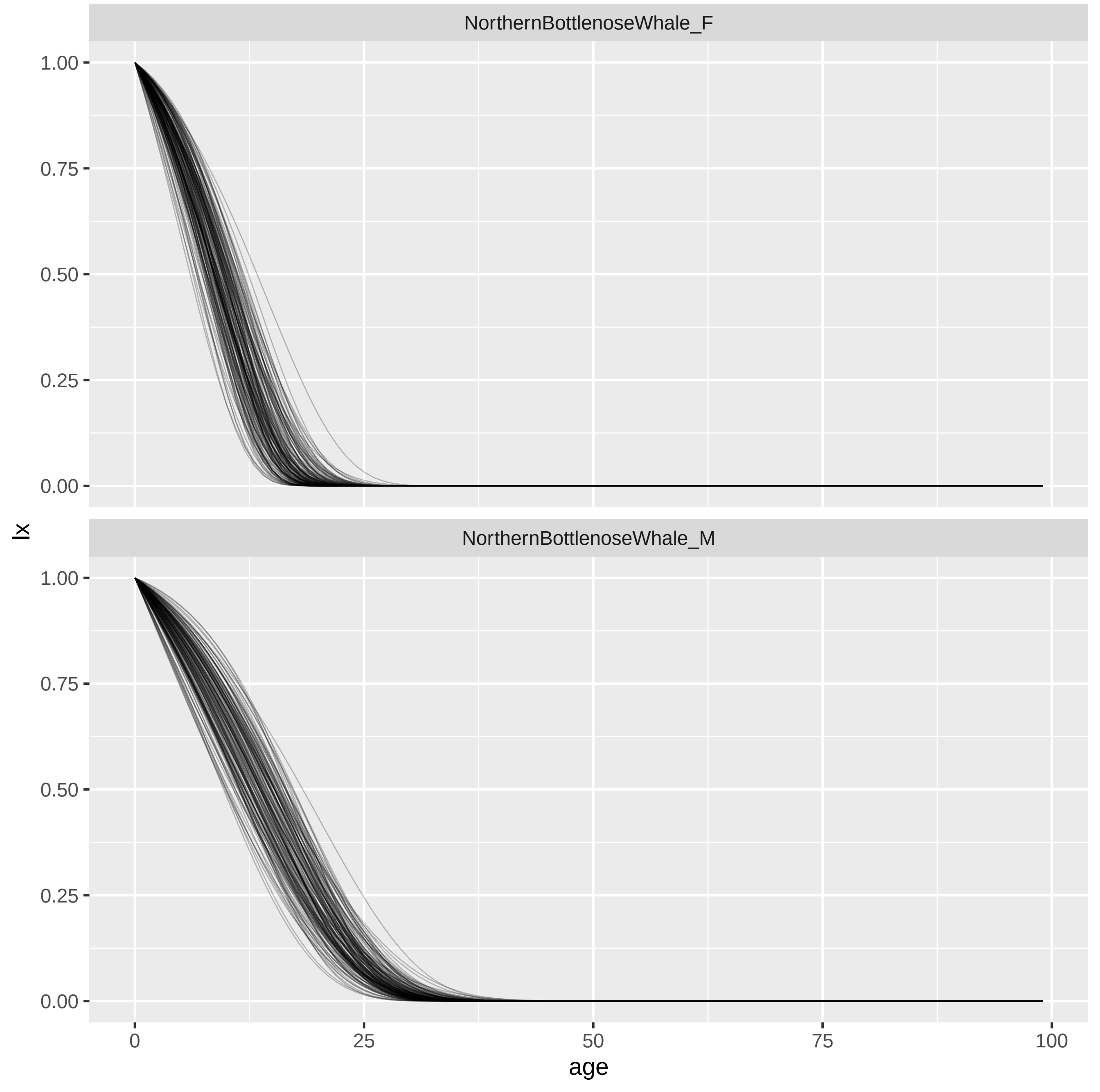
NorthernBottlenoseWhale_F
NorthernBottlenoseWhale_M
1.00
0.75
0.50
0.25
0.00
lx
0
25
50
75
100
age

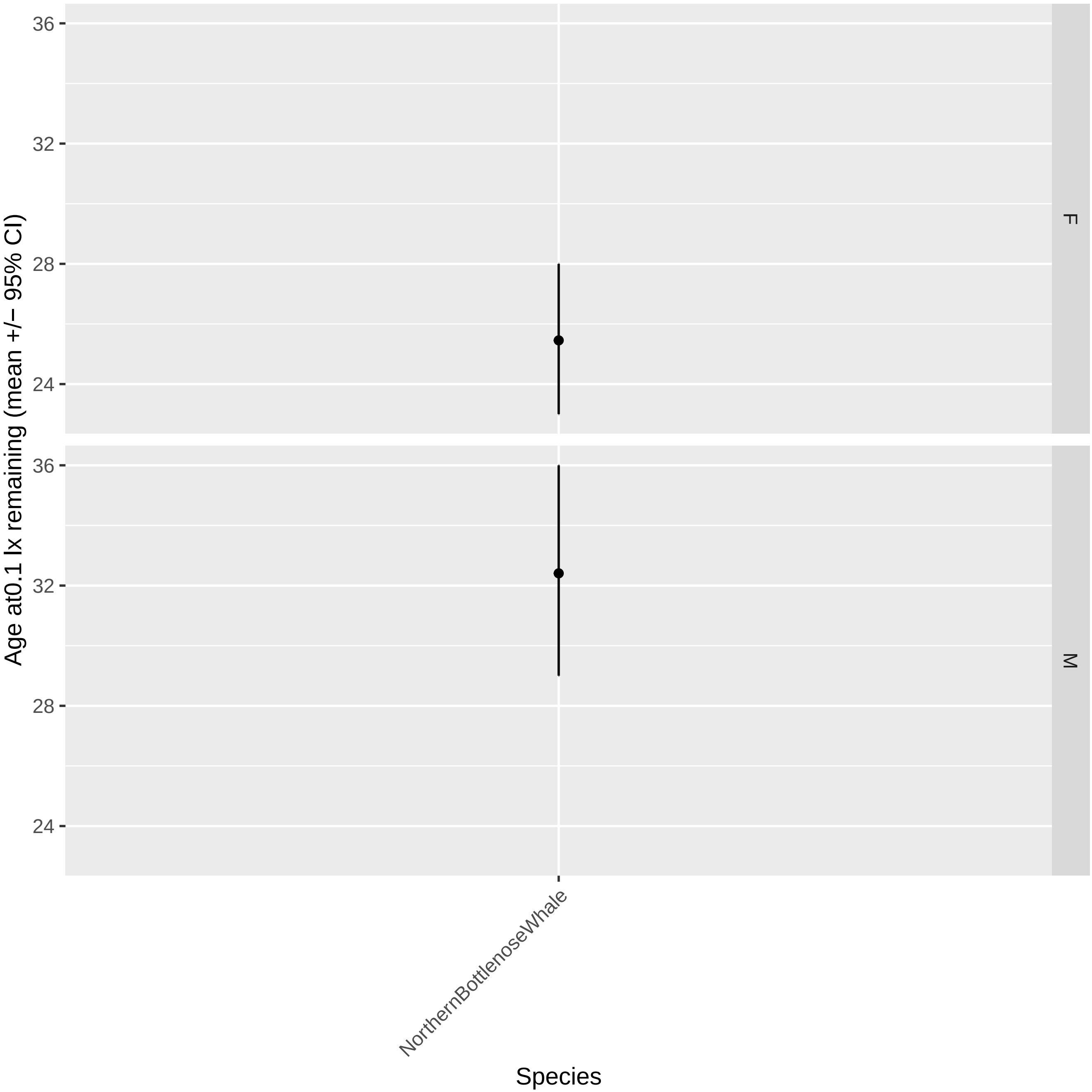
Age at0.1 lx remaining (mean +/− 95% CI)
36
32
28
24
F
M
NorthernBottlenoseWhale
Species

NorthernRightWhaleDolphin_F: dataset.num #1

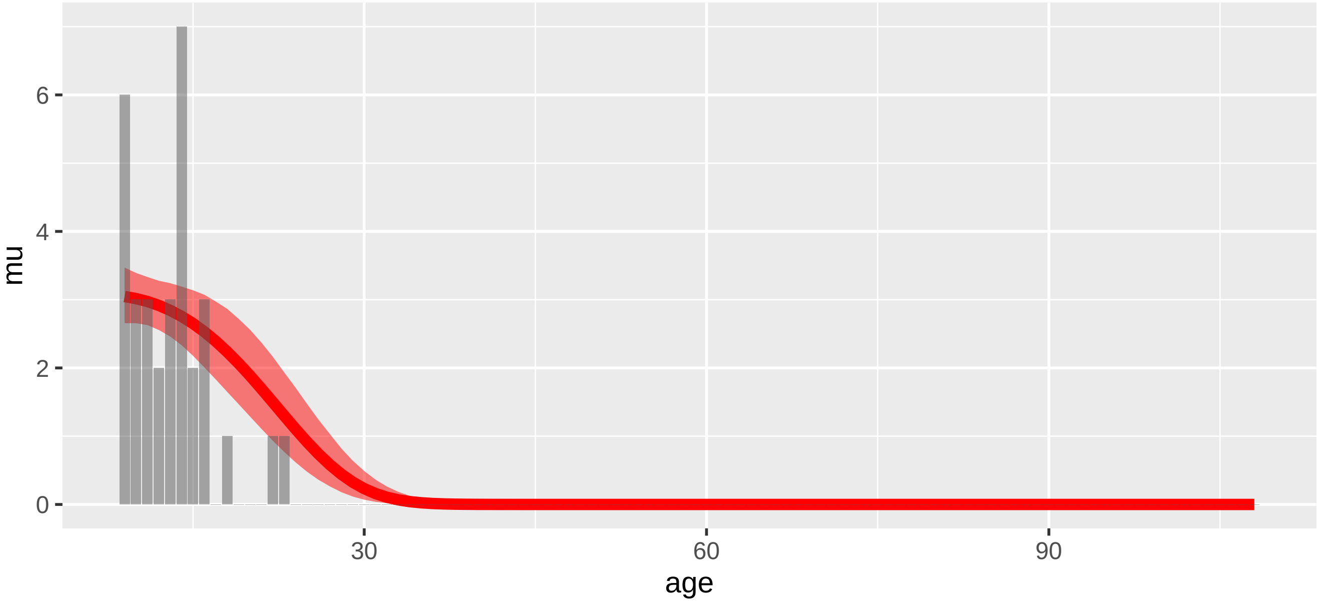


NorthernRightWhaleDolphin_F: dataset.num #3

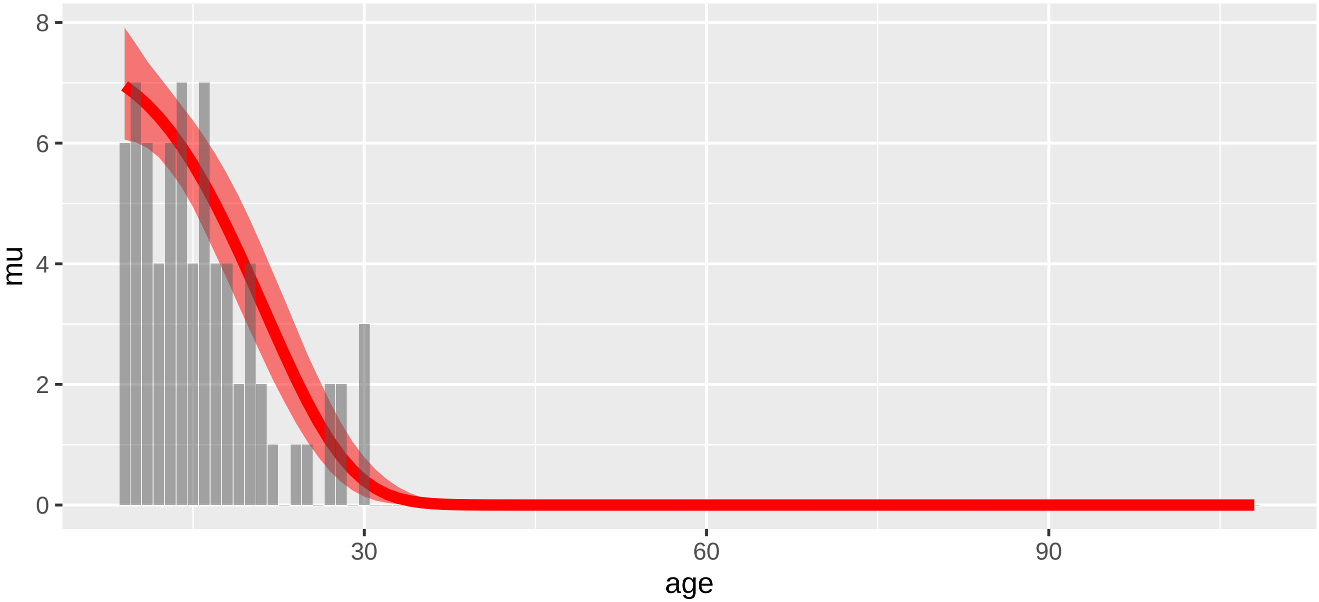

NorthernRightWhaleDolphin_M: dataset.num #2

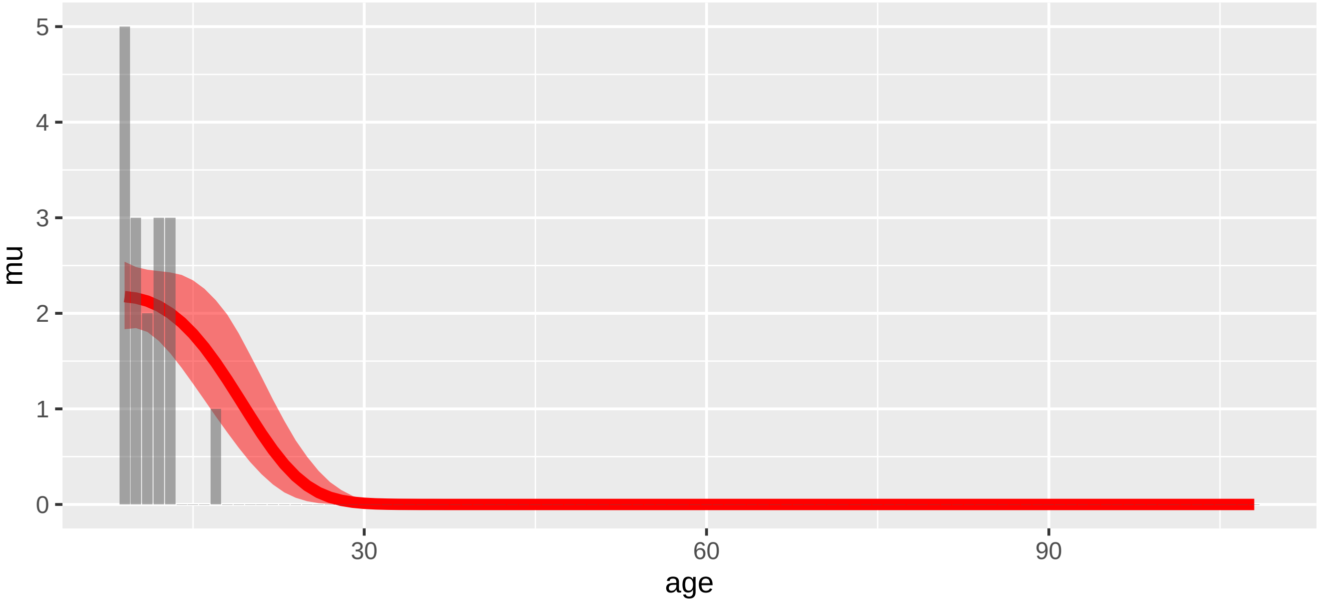


NorthernRightWhaleDolphin_M: dataset.num #4

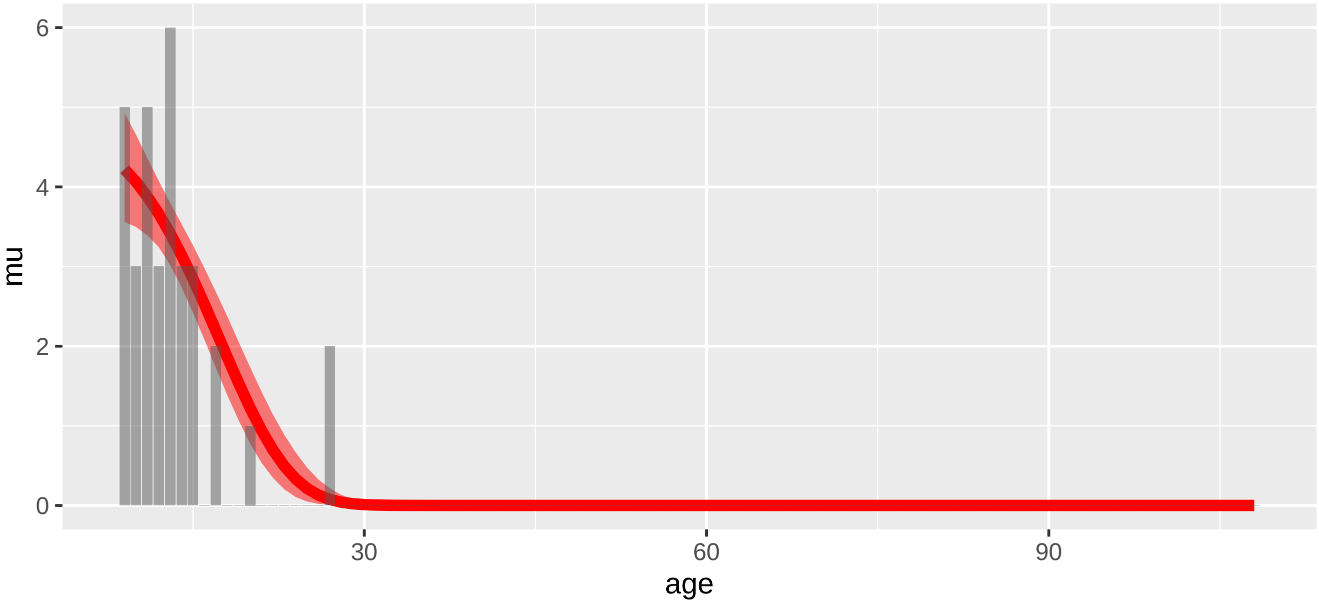

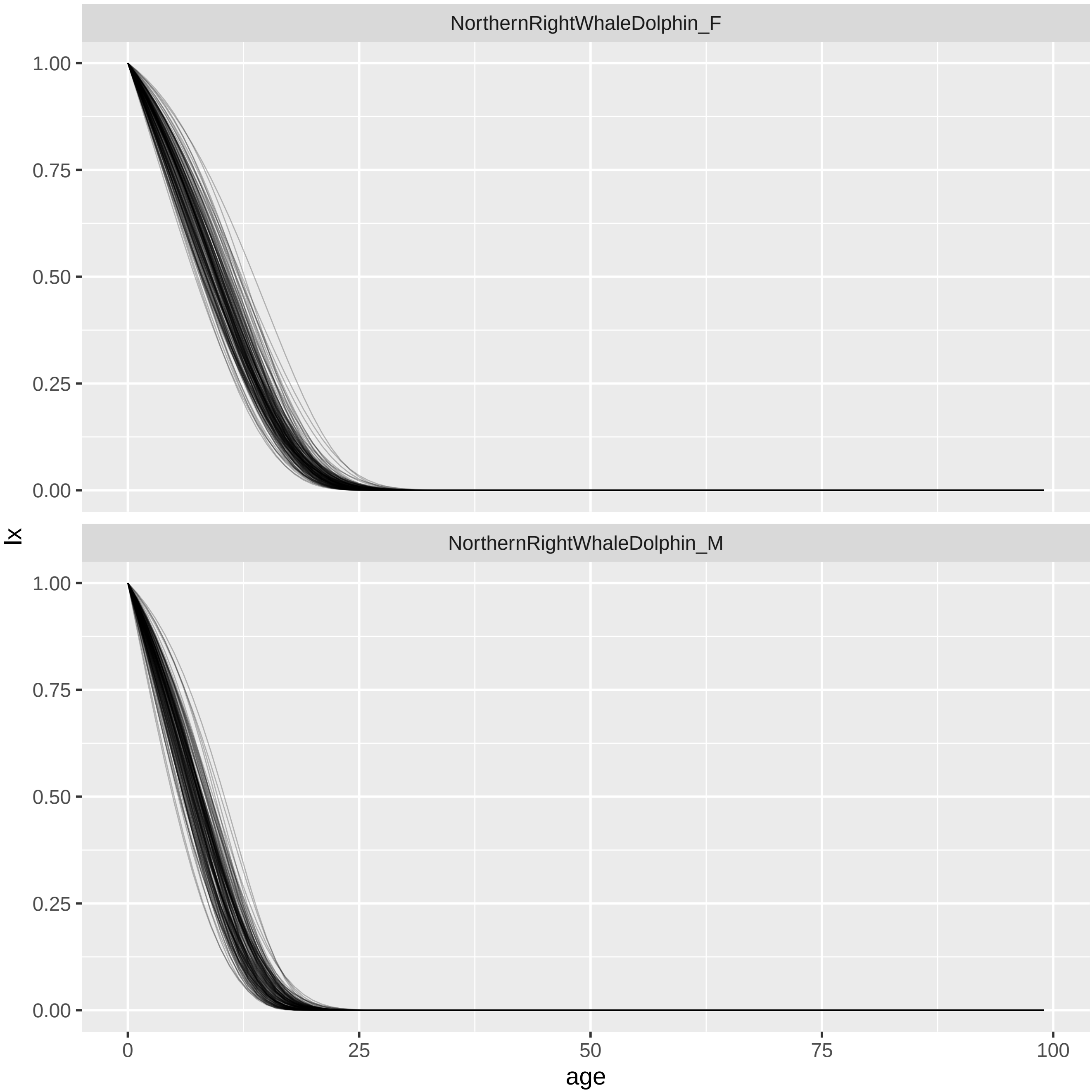
NorthernRightWhaleDolphin_F
NorthernRightWhaleDolphin_M
lx
1.00
0.75
0.50
0.25
0.00
0
25
50
75
100
age

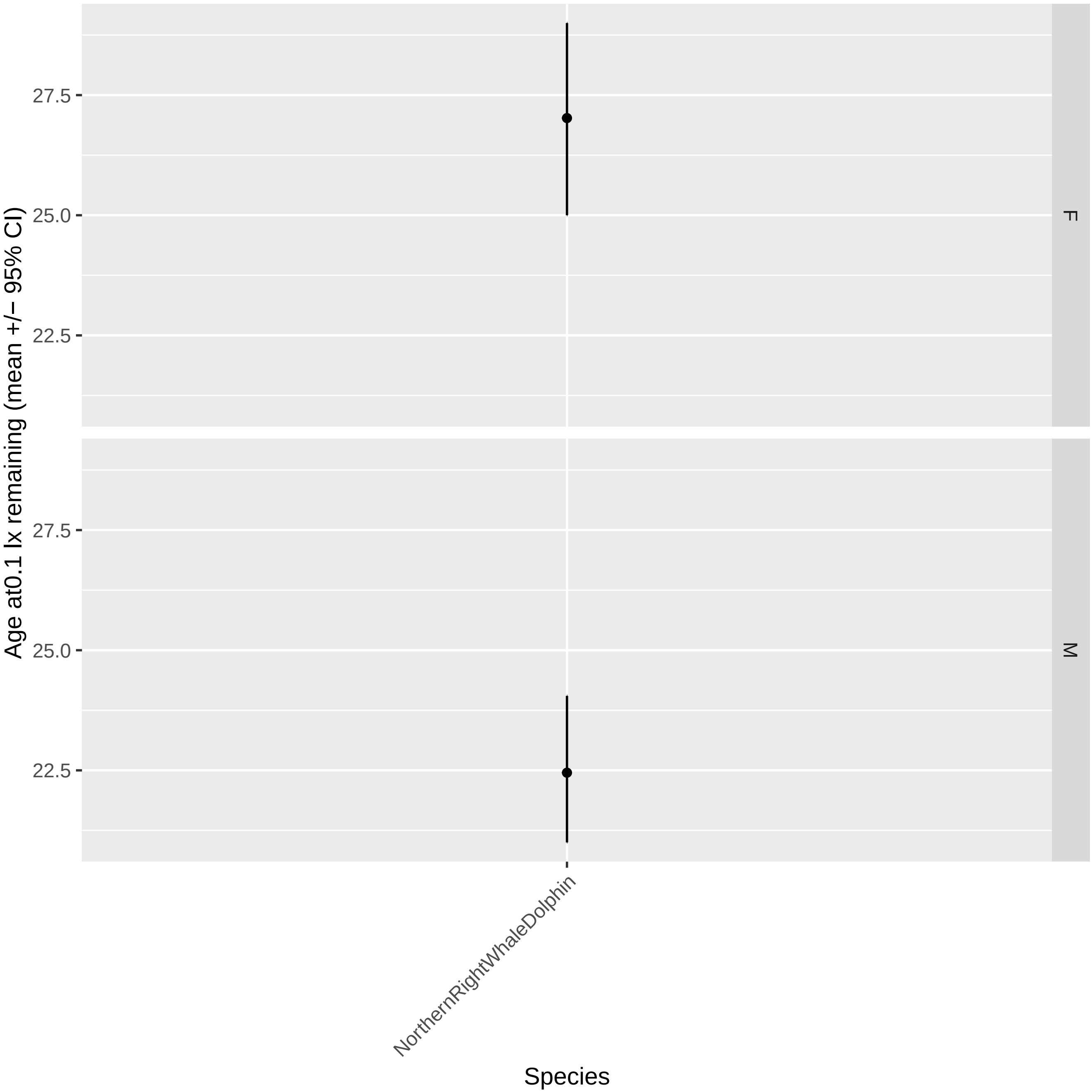

Age at0.1 lx remaining (mean +/− 95% CI)
27.5
25.0
22.5
27.5
25.0
22.5
F
M
NorthernRightWhaleDolphin
Species

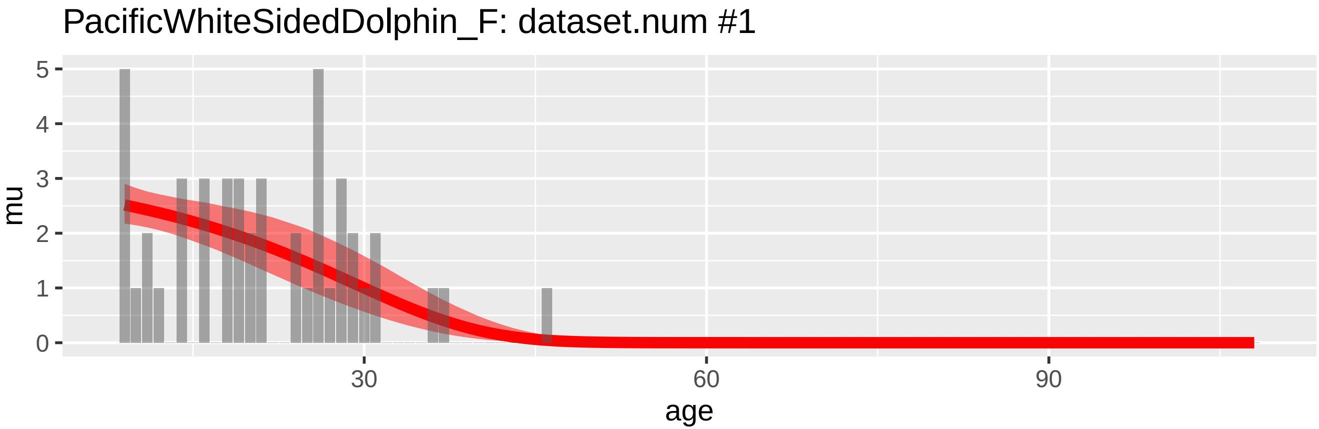
PacificWhiteSidedDolphin_F: dataset.num #1
mu
0
1
2
3
4
5
30
60
90
age

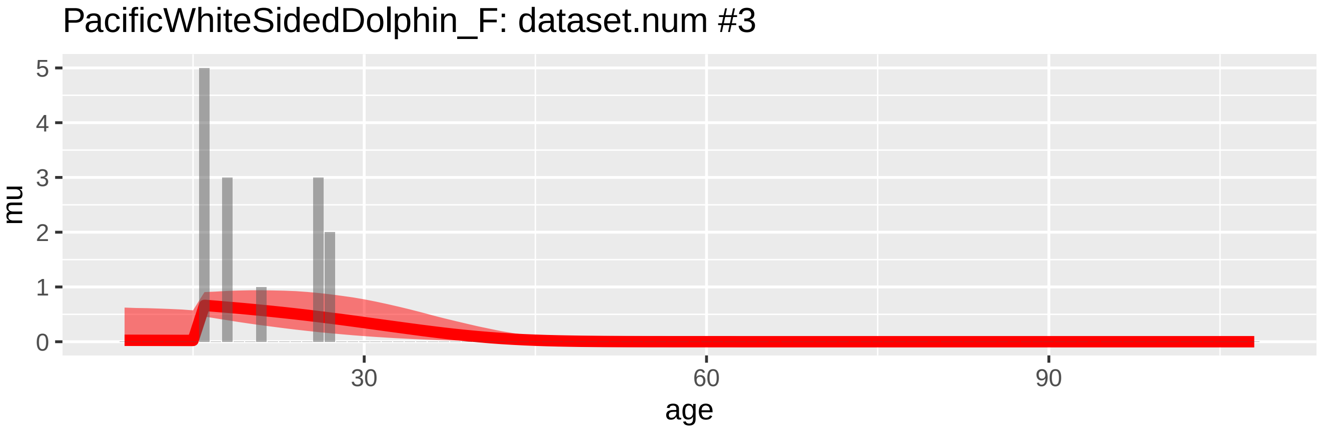
PacificWhiteSidedDolphin_F: dataset.num #3
mu
0
1
2
3
4
5
30
60
90
age

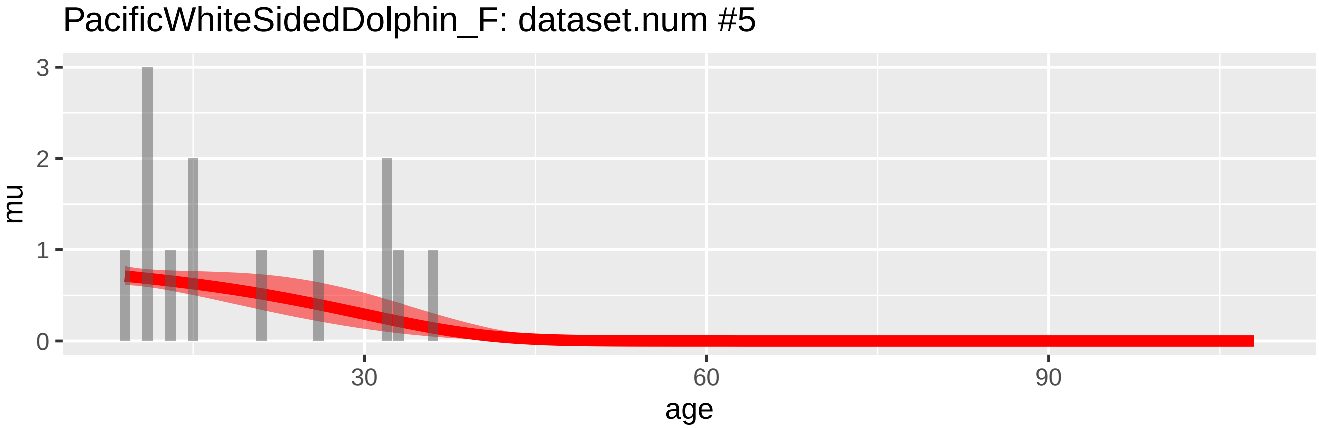
PacificWhiteSidedDolphin_F: dataset.num #5
mu
0
1
2
3
30
60
90
age

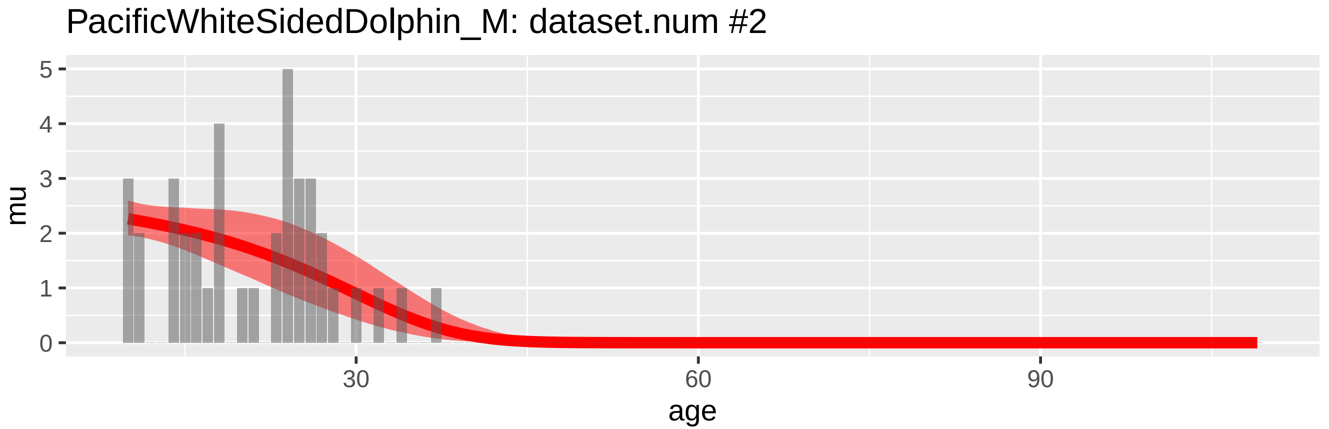
PacificWhiteSidedDolphin_M: dataset.num #2
mu
0
1
2
3
4
5
30
60
90
age

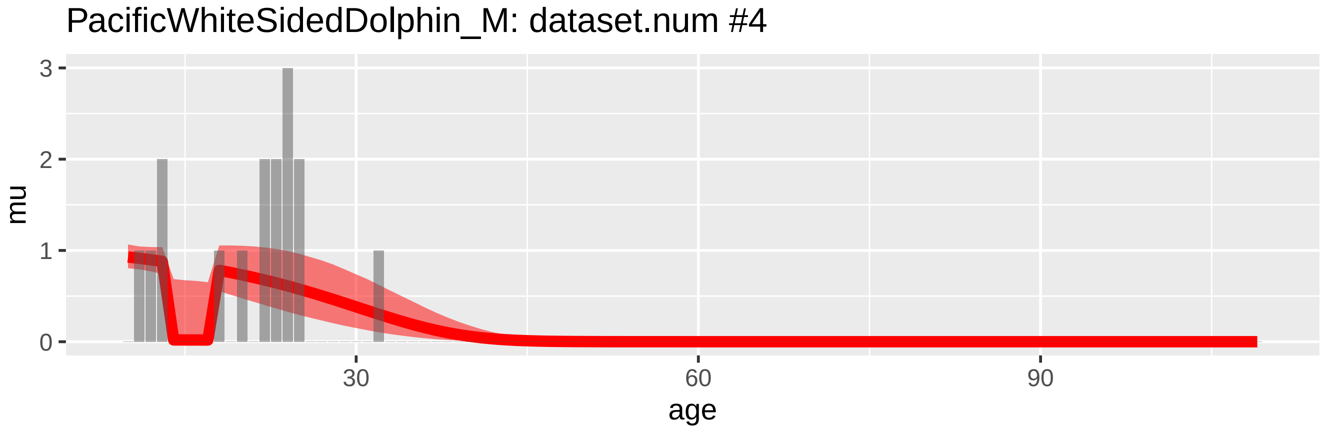
PacificWhiteSidedDolphin_M: dataset.num #4
mu
0
1
2
3
30
60
90
age

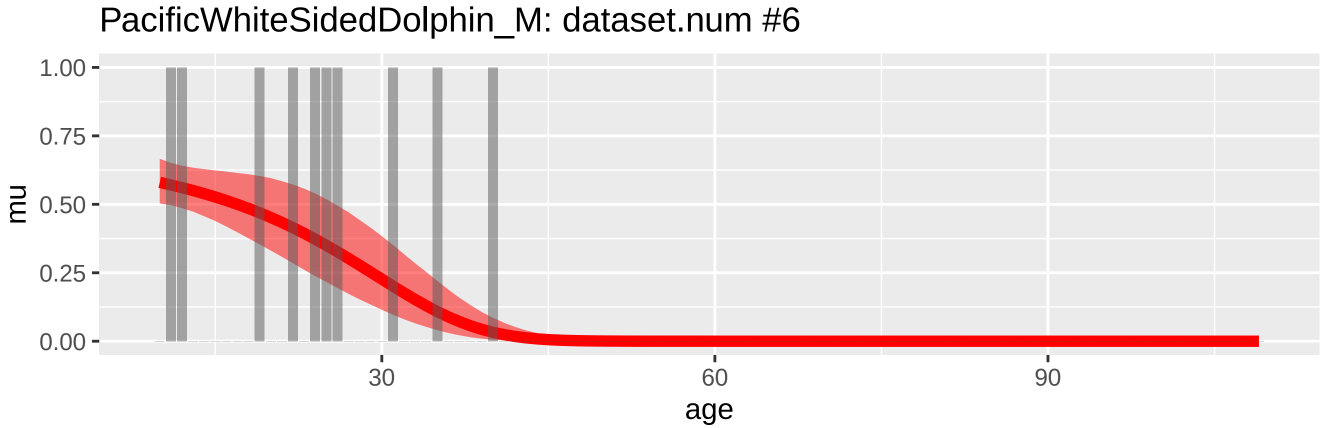
PacificWhiteSidedDolphin_M: dataset.num #6
mu
0.00
0.25
0.50
0.75
1.00
30
60
90
age

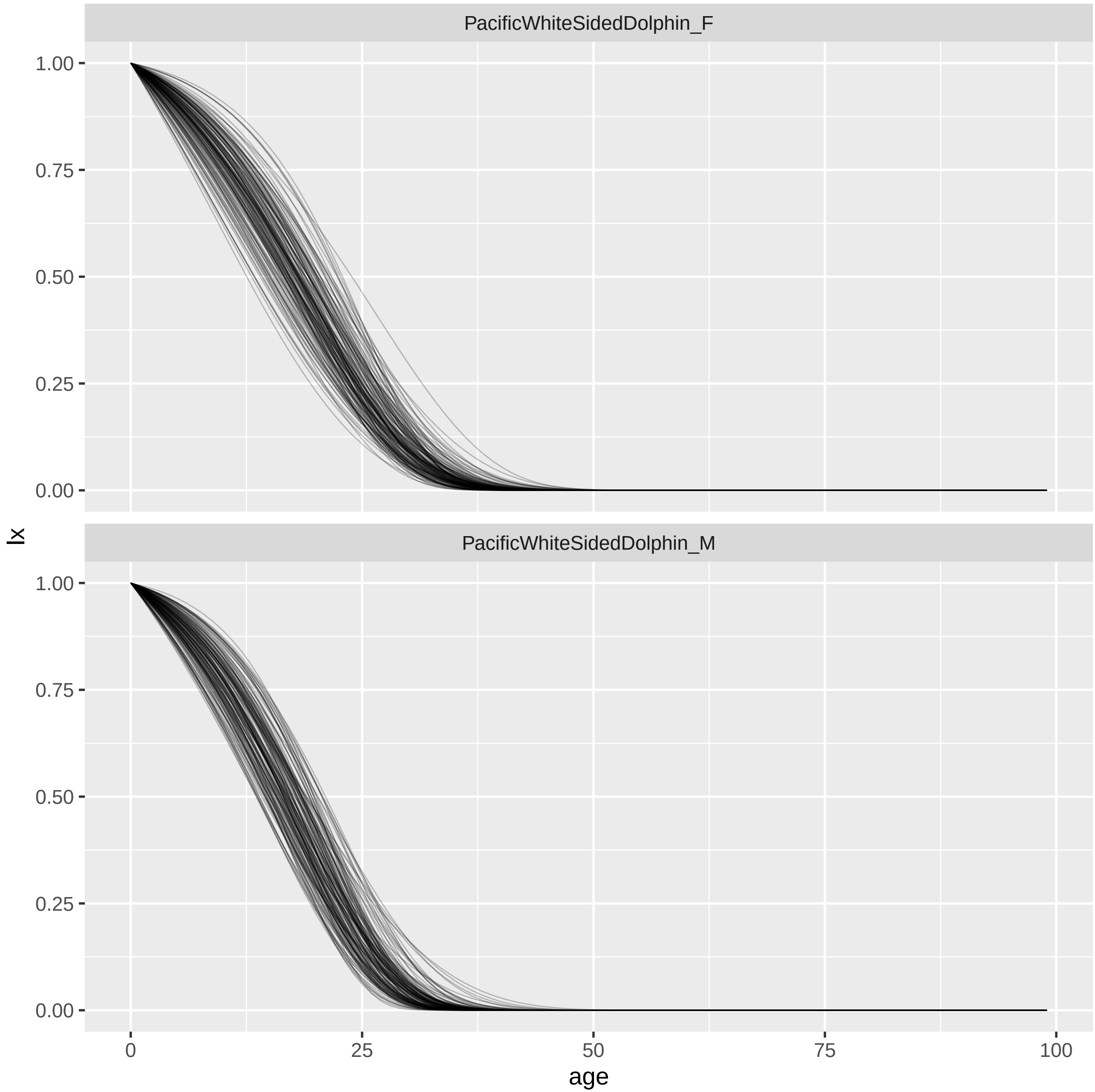
PacificWhiteSidedDolphin_F
PacificWhiteSidedDolphin_M
1.00
0.75
0.50
0.25
0.00
lx
0
25
50
75
100
age

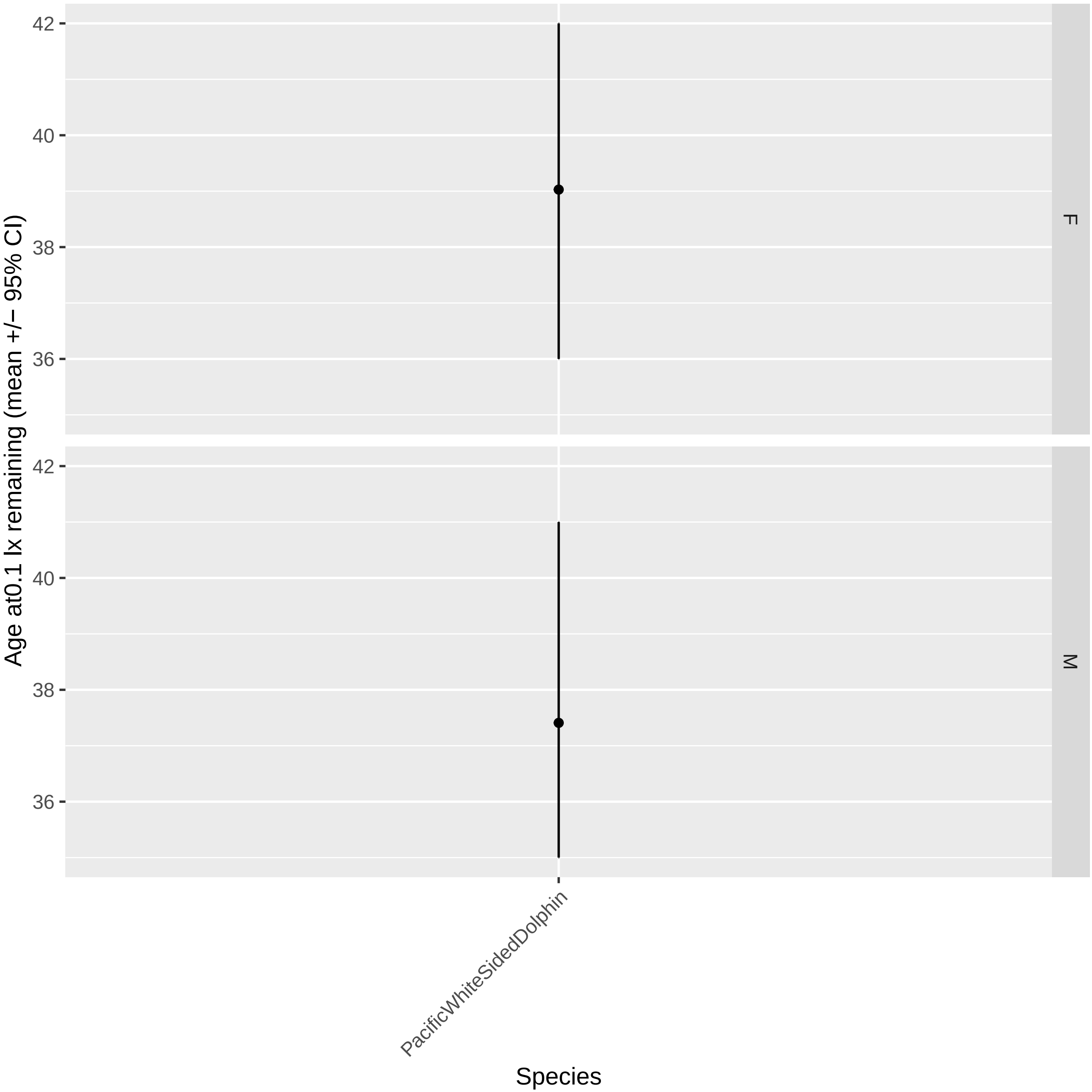

42
40
38
36
F
M
Age at0.1 lx remaining (mean +/− 95% CI)
PacificWhiteSidedDolphin
Species

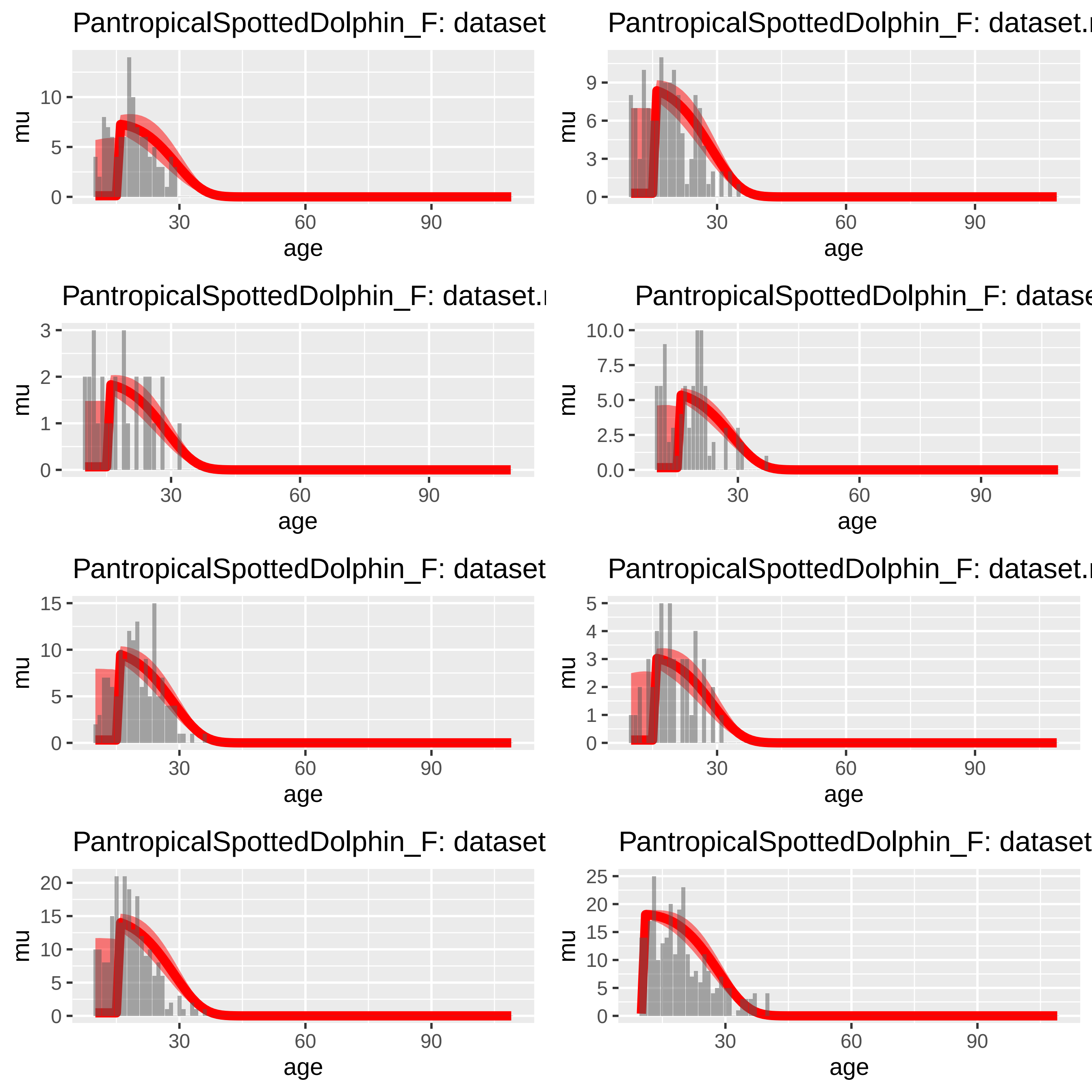

PantropicalSpottedDolphin_F: dataset
PantropicalSpottedDolphin_F: dataset.r
PantropicalSpottedDolphin_F: dataset.r
PantropicalSpottedDolphin_F: datase
PantropicalSpottedDolphin_F: dataset
PantropicalSpottedDolphin_F: dataset.r
PantropicalSpottedDolphin_F: dataset
PantropicalSpottedDolphin_F: dataset
mu
age
30
60
90

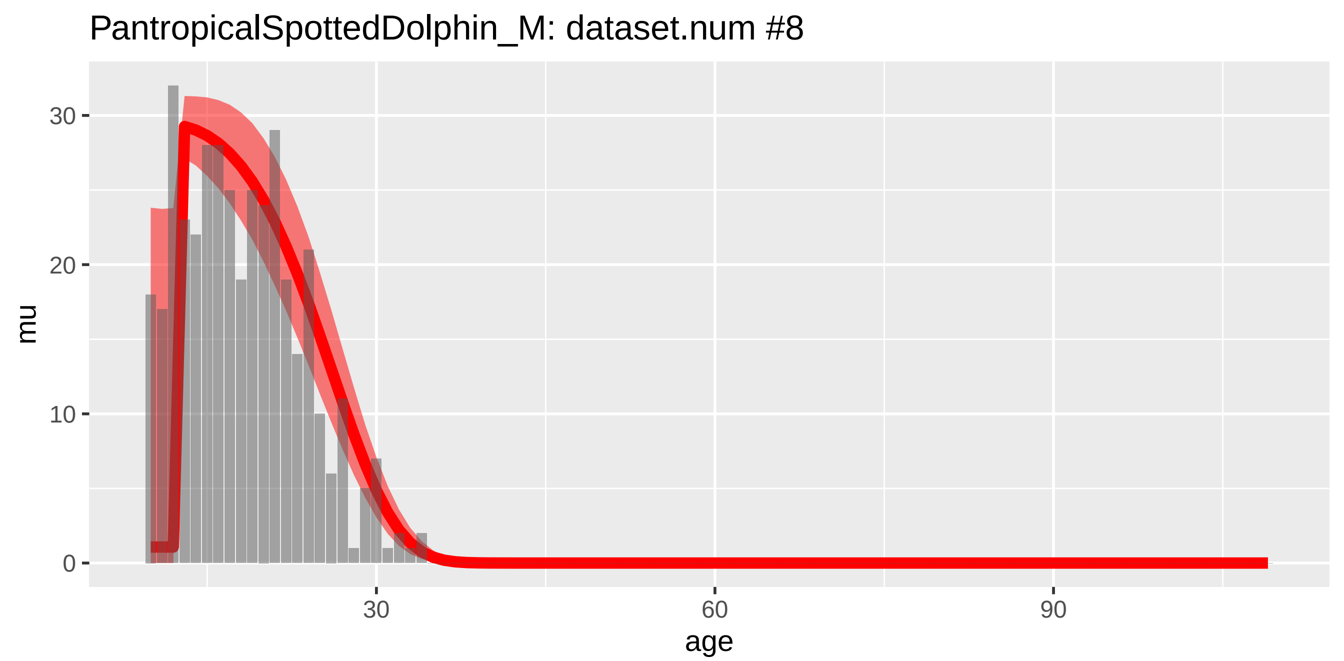
PantropicalSpottedDolphin_M: dataset.num #8
mu
age
0
10
20
30
60
90

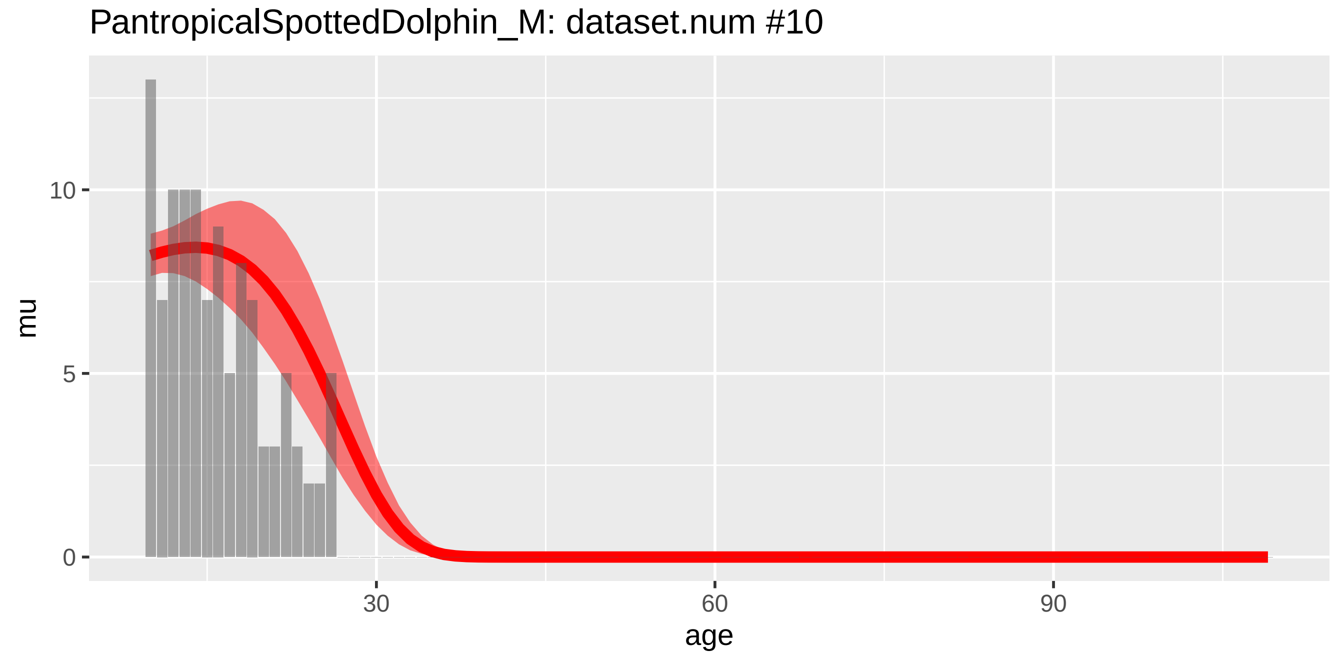
PantropicalSpottedDolphin_M: dataset.num #10
mu
age
0
5
10
30
60
90

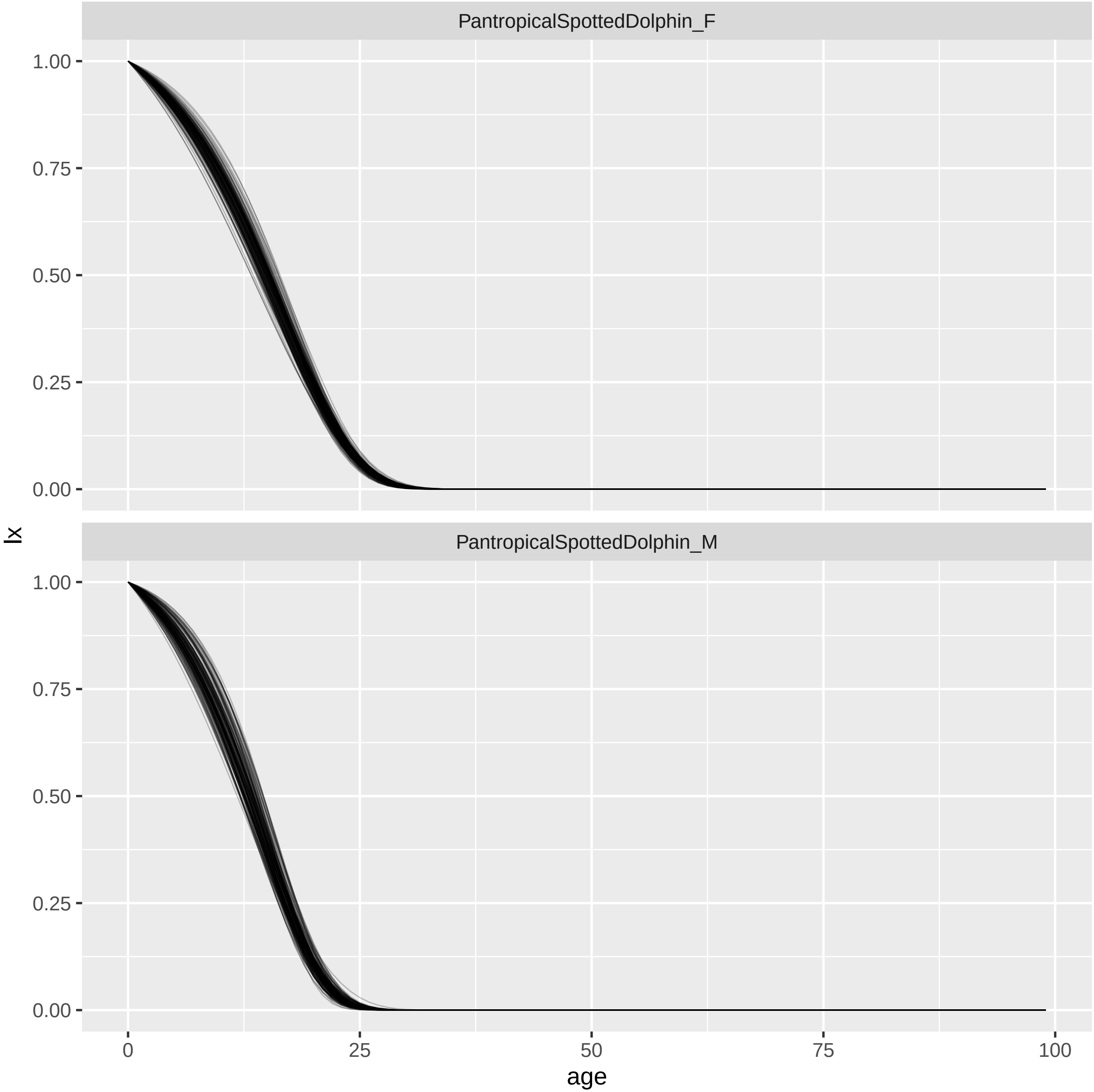
PantropicalSpottedDolphin_F
PantropicalSpottedDolphin_M
1.00
0.75
0.50
0.25
0.00
lx
0
25
50
75
100
age

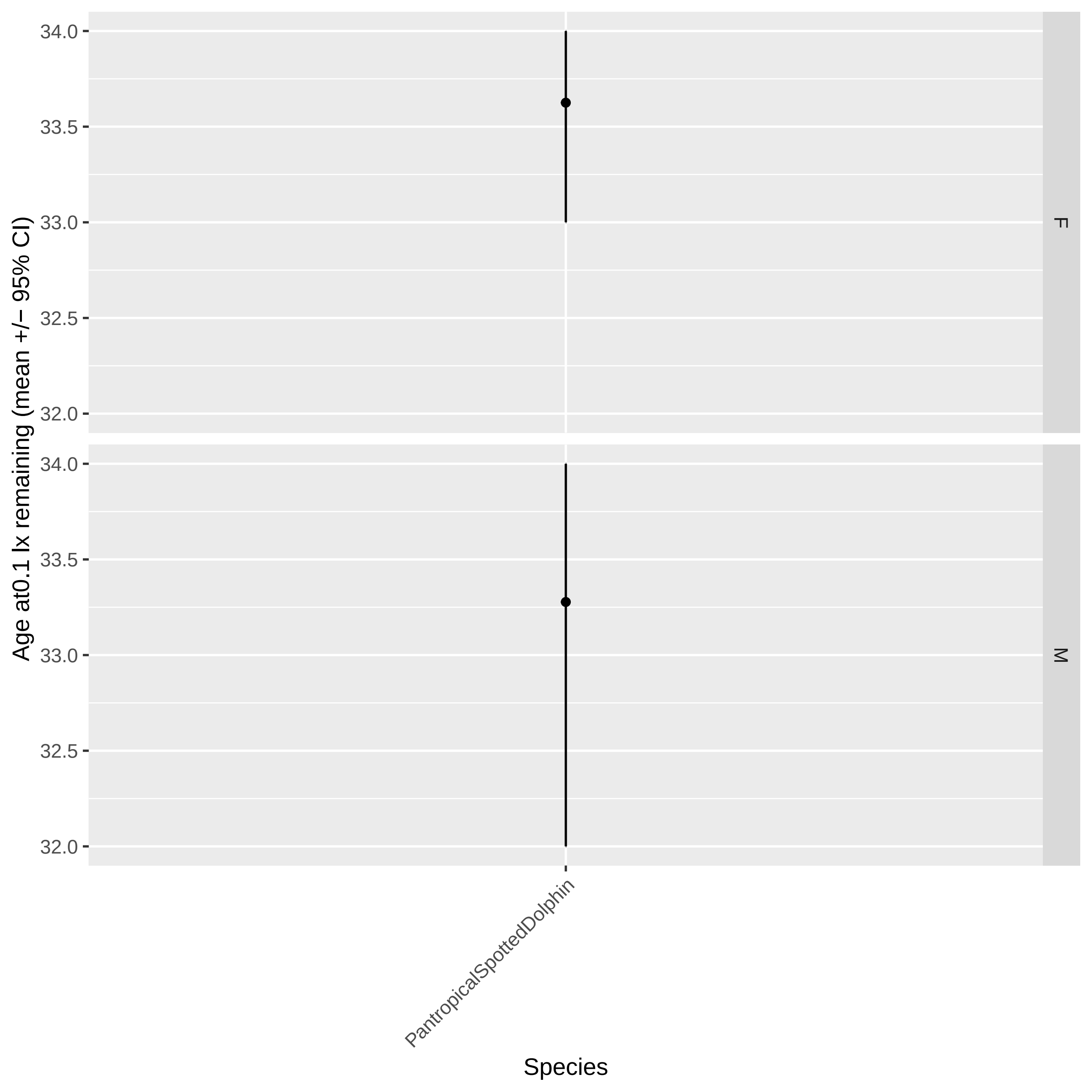
Age at0.1 lx remaining (mean +/− 95% CI)
34.0
33.5
33.0
32.5
32.0
F
M
PantropicalSpottedDolphin
Species

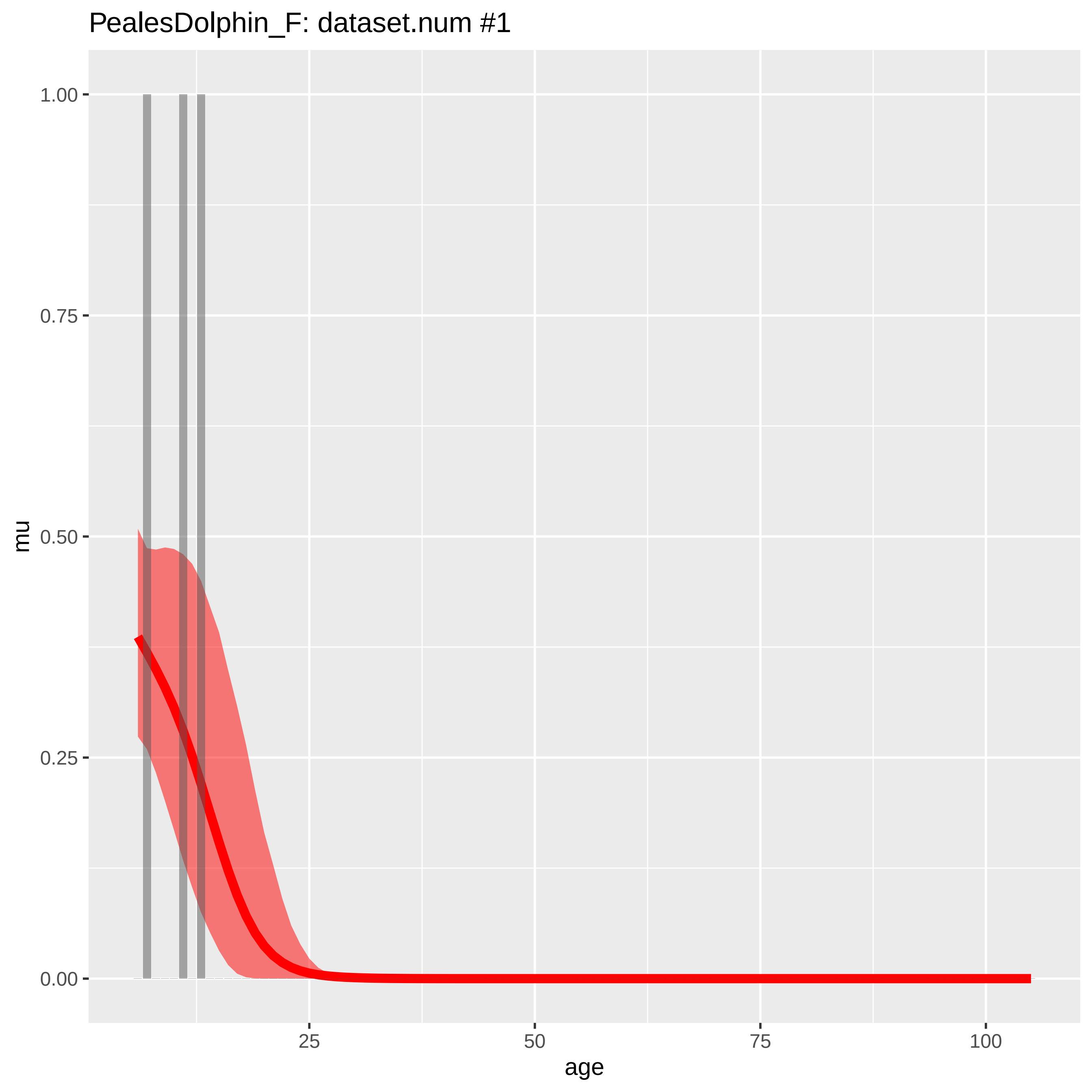
PealesDolphin_F: dataset.num #1
mu
1.00
0.75
0.50
0.25
0.00
25
50
75
100
age

PealesDolphin_M: dataset.num #2

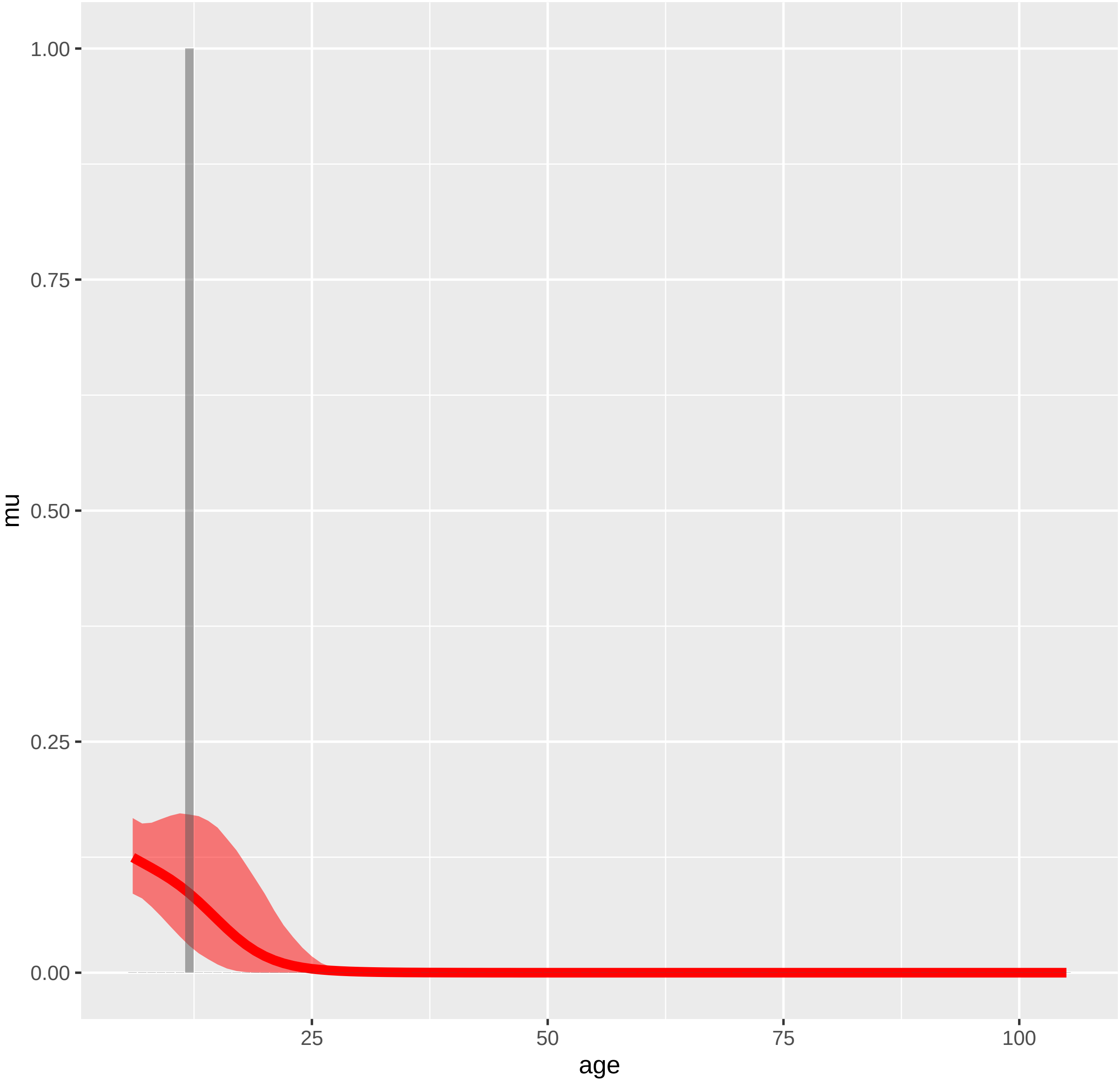

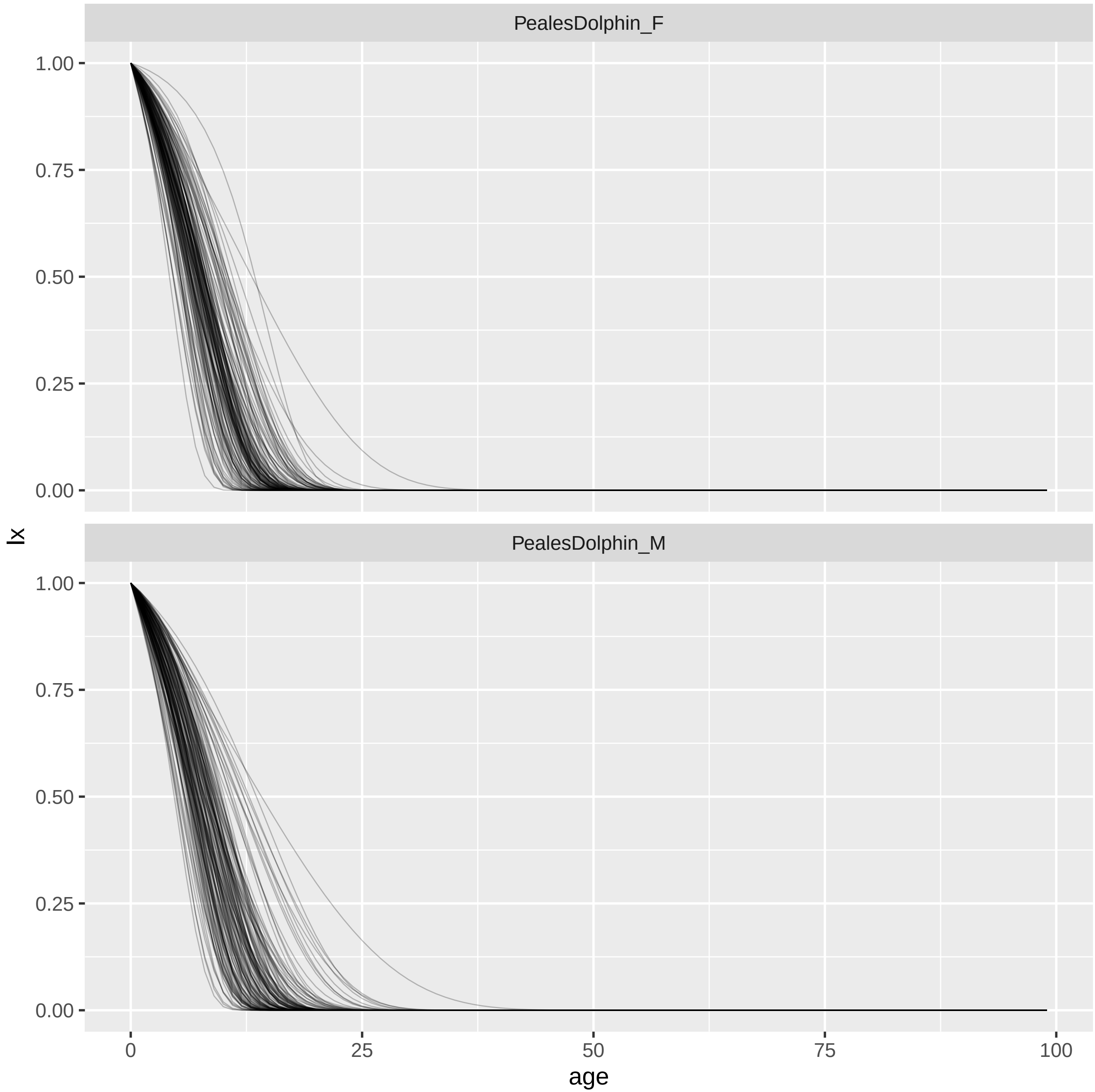
PealesDolphin_F
PealesDolphin_M
lx
age
1.00
0.75
0.50
0.25
0.00
0
25
50
75
100

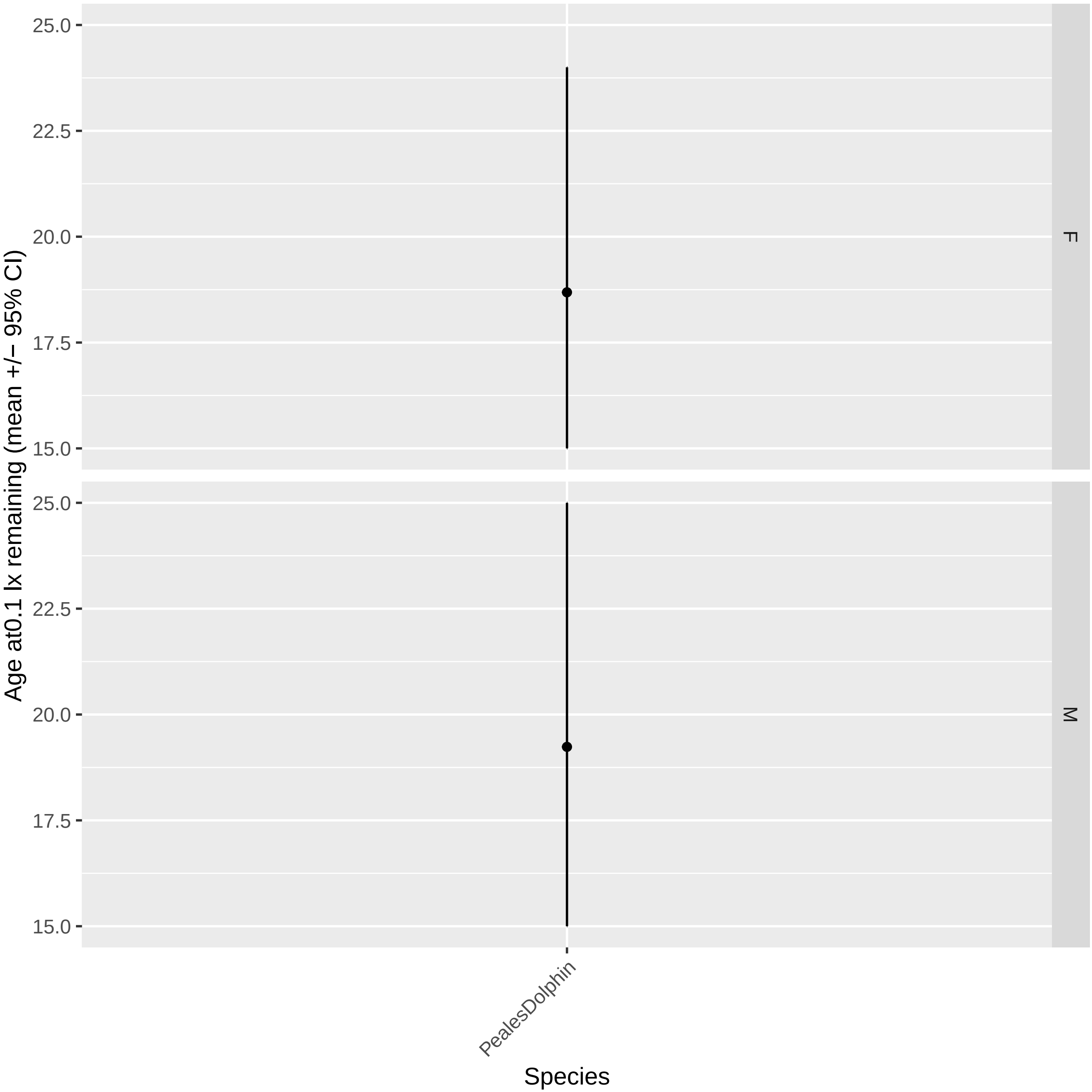

Age at0.1 lx remaining (mean +/− 95% CI)
25.0
22.5
20.0
17.5
15.0
F
M
PealesDolphin
Species

PygmySpermWhale_F: dataset.num #1

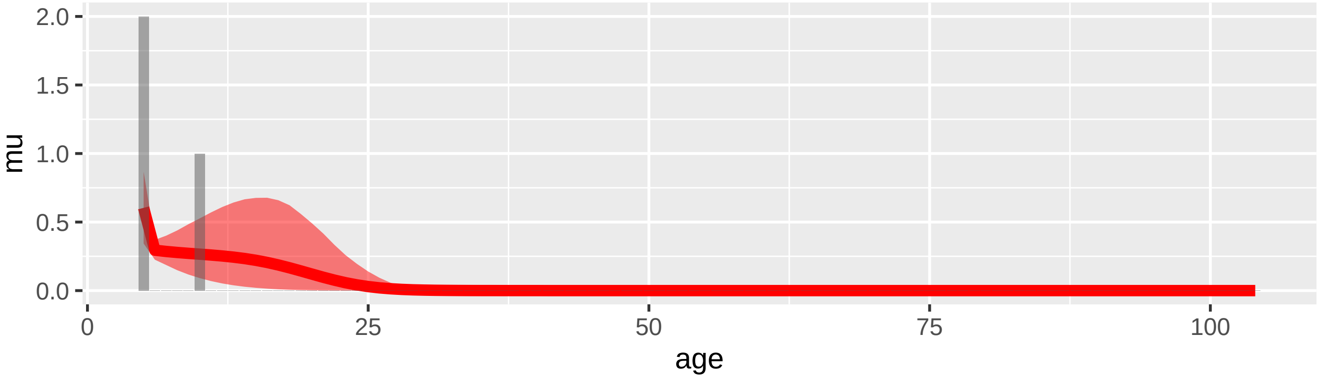


PygmySpermWhale_F: dataset.num #2

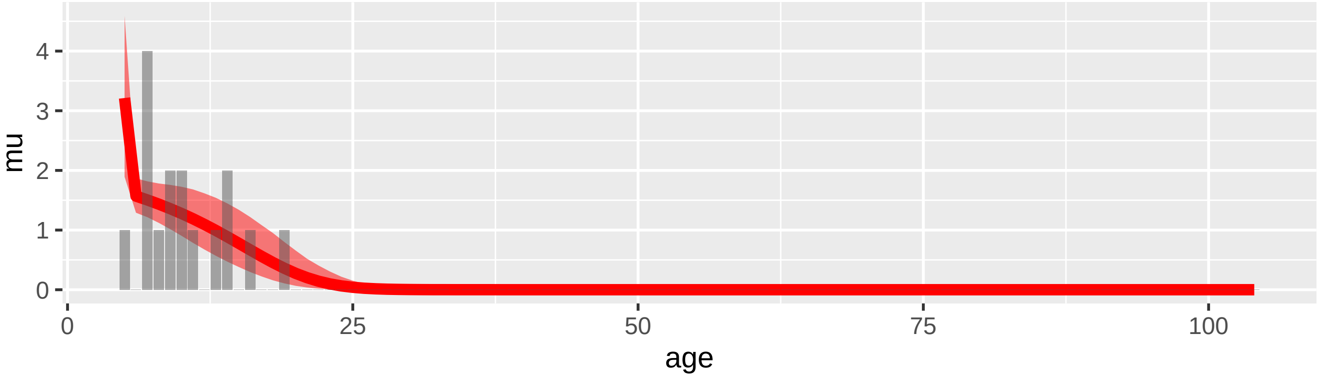


PygmySpermWhale_F: dataset.num #5

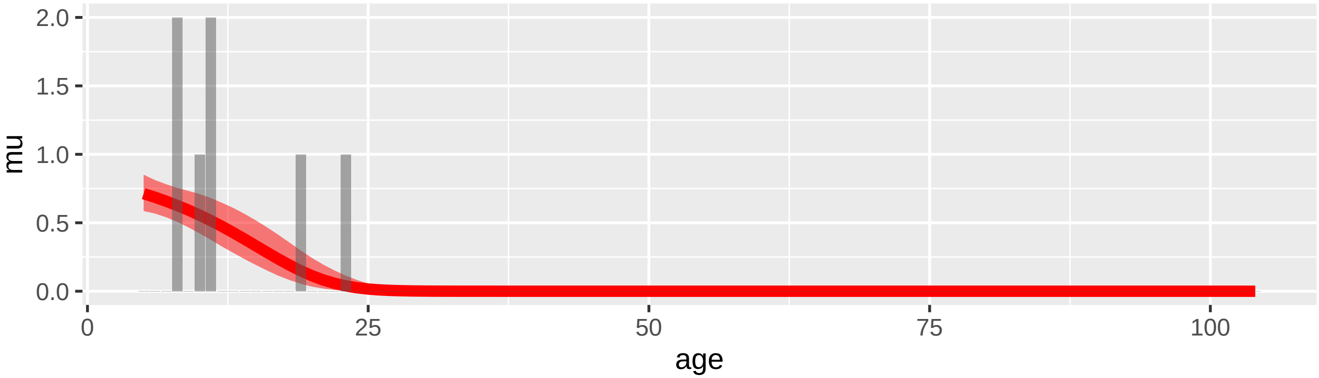

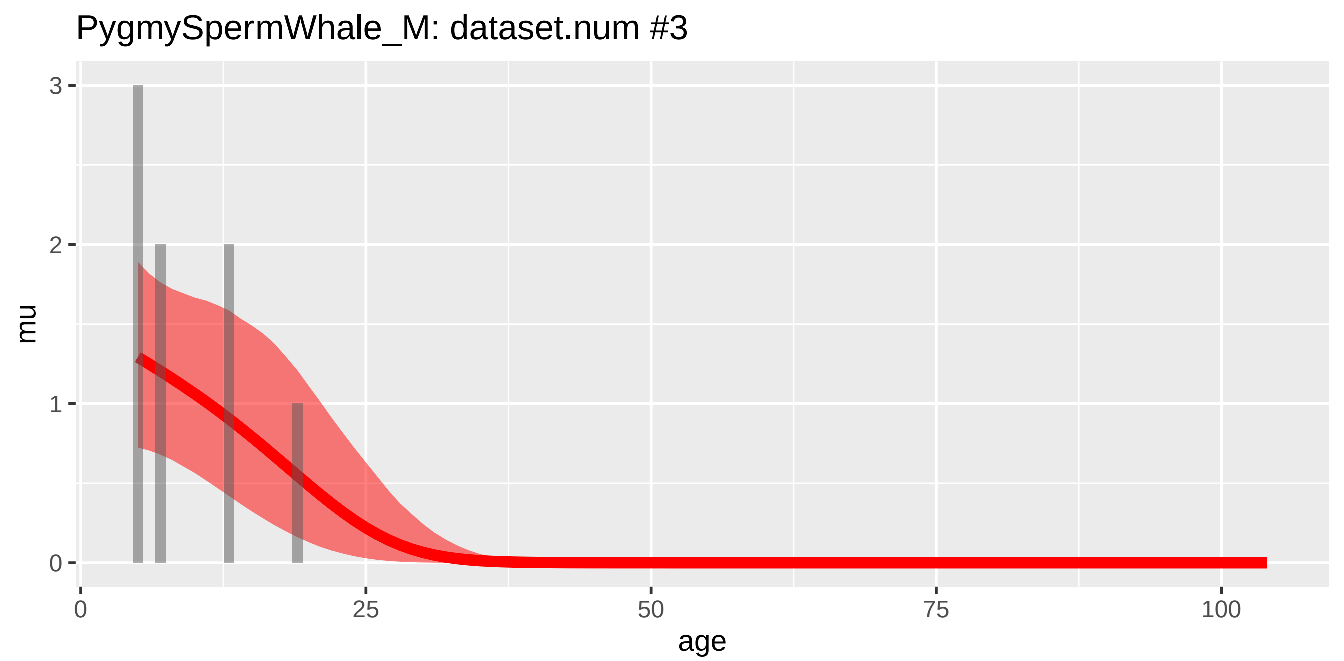
PygmySpermWhale_M: dataset.num #3
mu
age
0
1
2
3
0
25
50
75
100

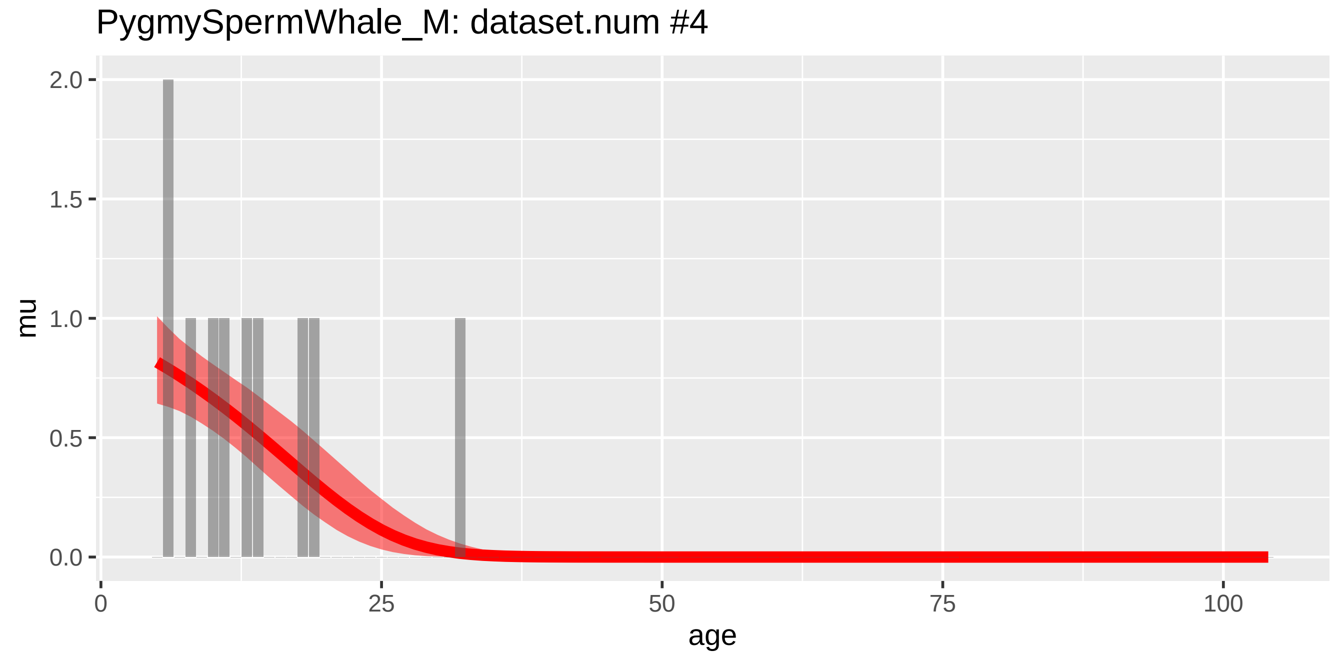
PygmySpermWhale_M: dataset.num #4
mu
age
0.0
0.5
1.0
1.5
2.0
0
25
50
75
100

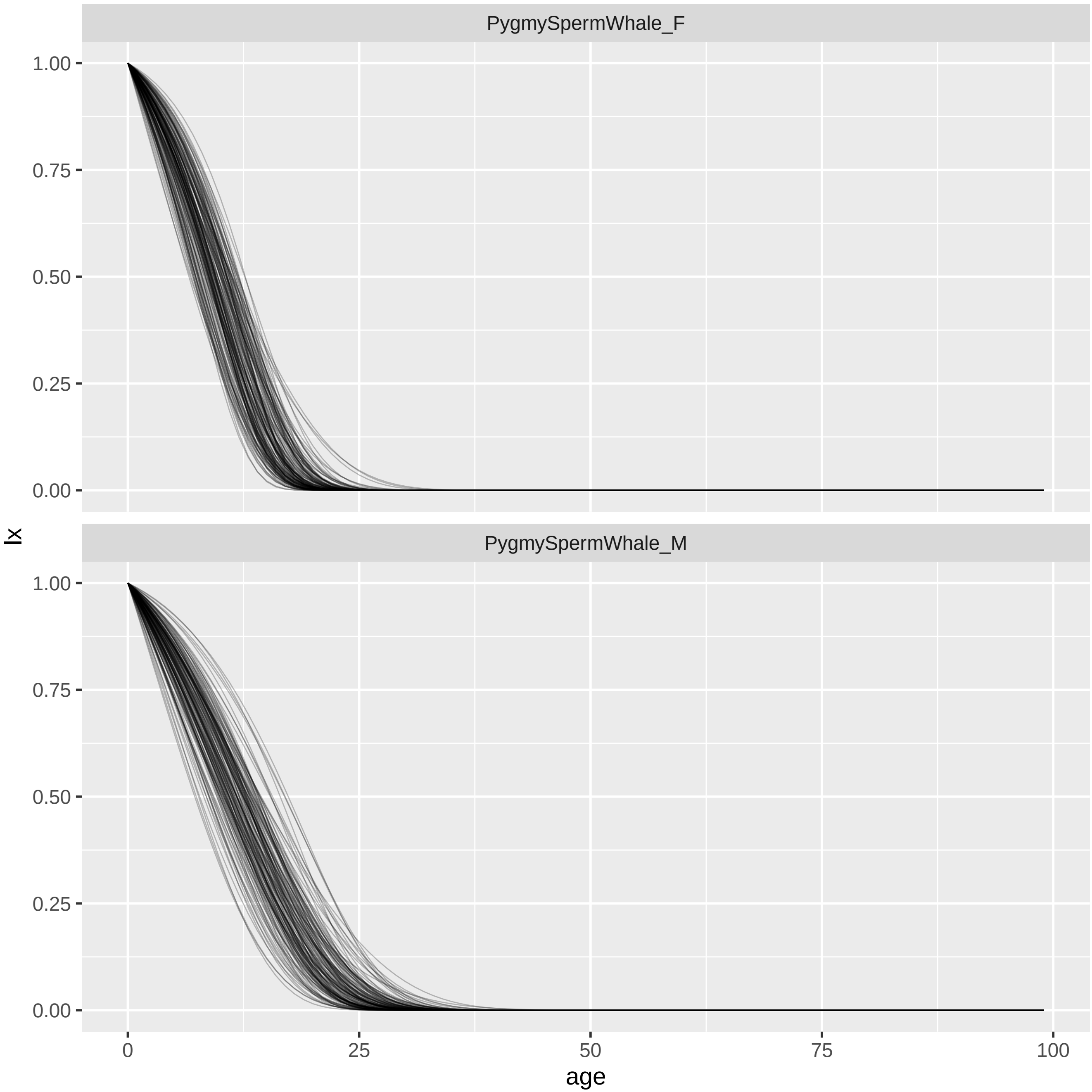
PygmySpermWhale_F
PygmySpermWhale_M
1.00
0.75
0.50
0.25
0.00
lx
0
25
50
75
100
age

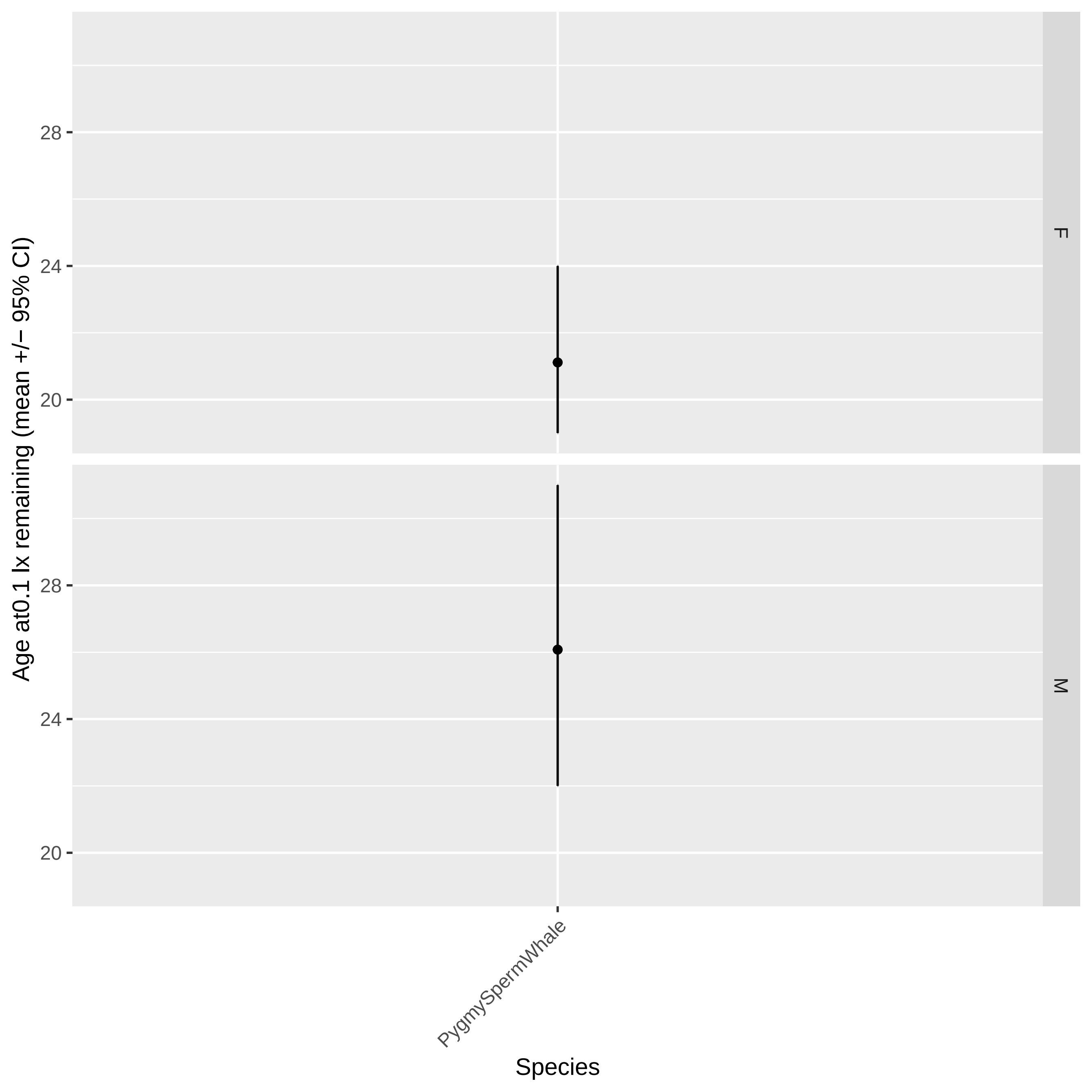

Age at0.1 lx remaining (mean +/− 95% CI)
28
24
20
28
24
20
F
M
PygmySpermWhale
Species

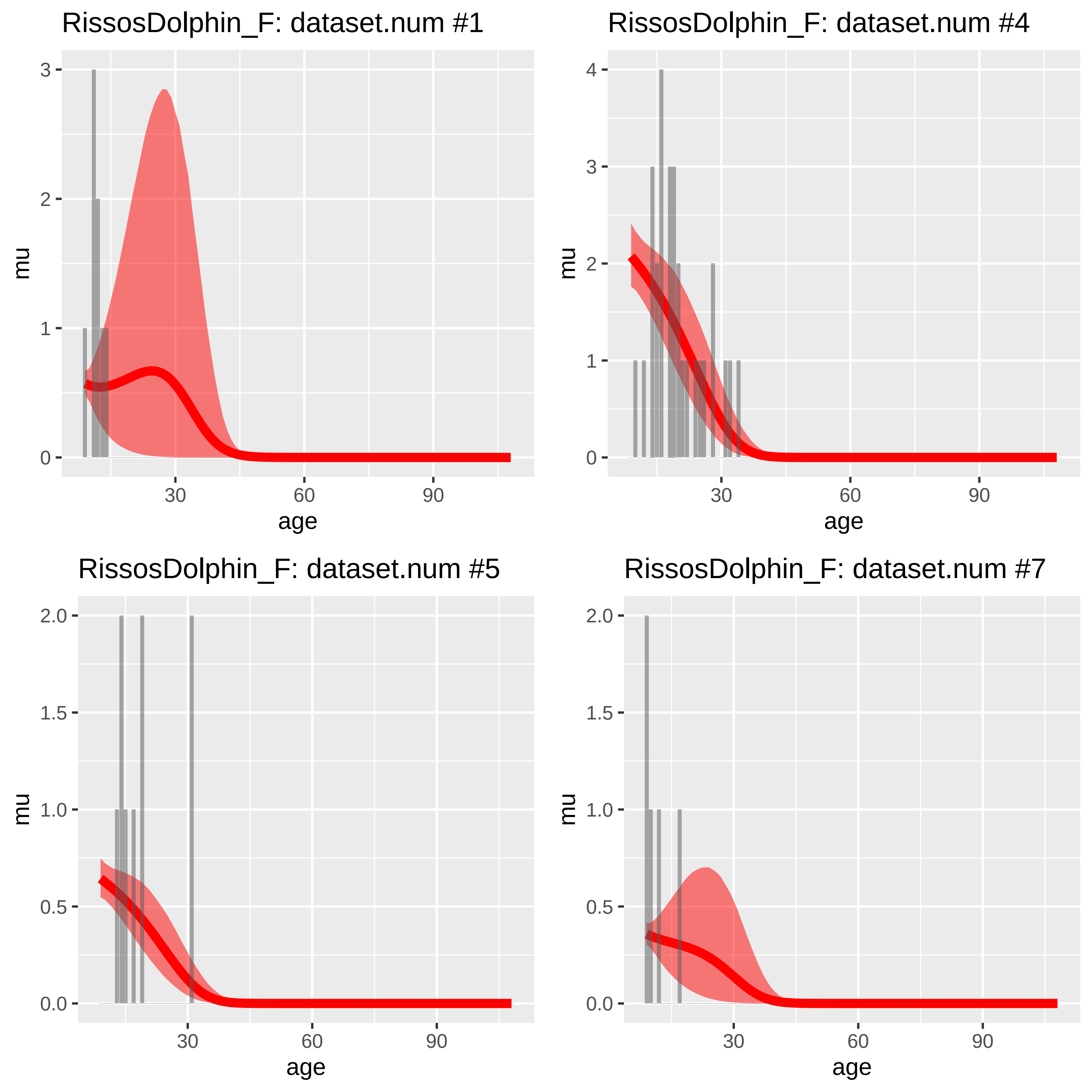
RissosDolphin_F: dataset.num #1
RissosDolphin_F: dataset.num #4
RissosDolphin_F: dataset.num #5
RissosDolphin_F: dataset.num #7
mu
age
0
1
2
3
4
0.0
0.5
1.0
1.5
2.0
30
60
90

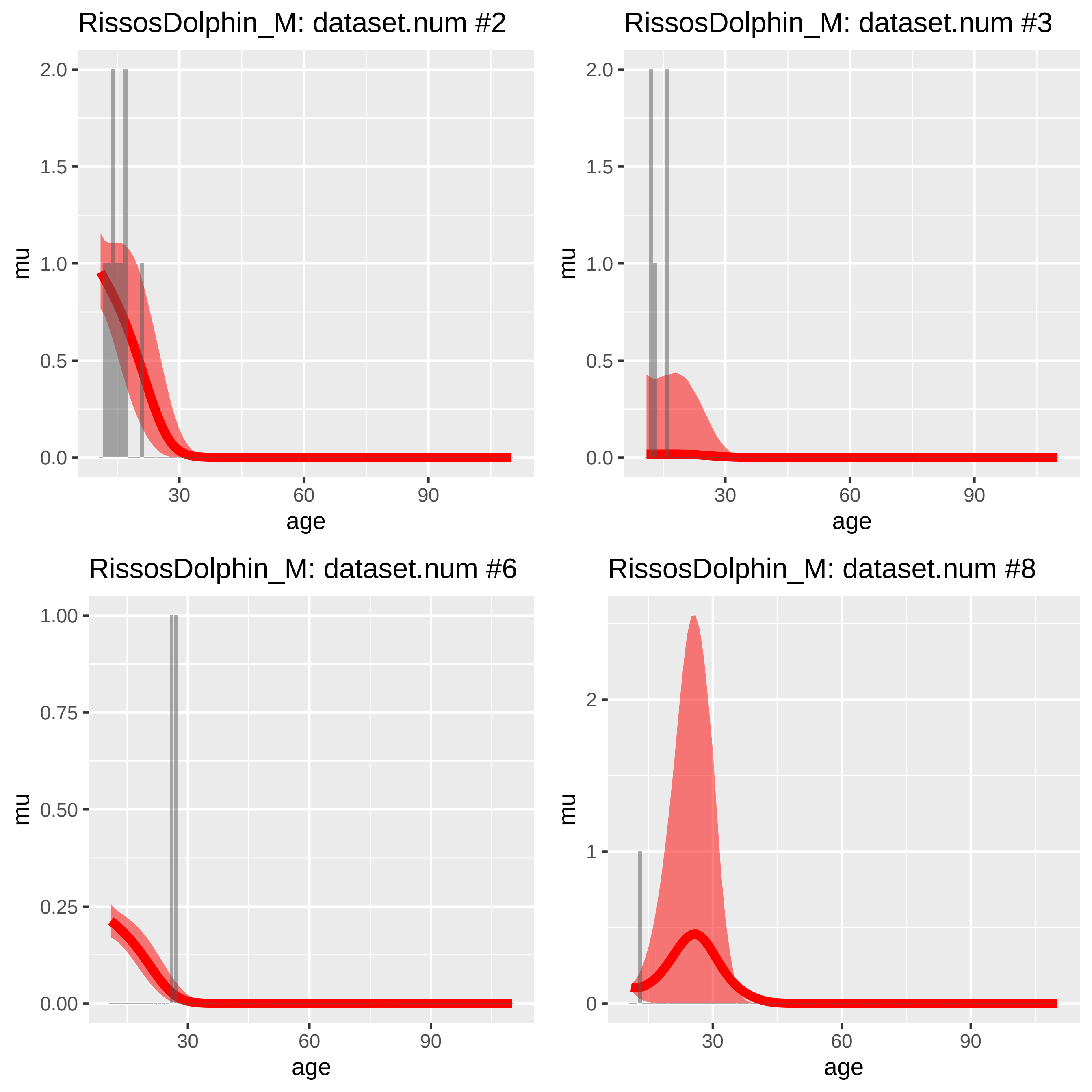
RissosDolphin_M: dataset.num #2
RissosDolphin_M: dataset.num #3
RissosDolphin_M: dataset.num #6
RissosDolphin_M: dataset.num #8
mu
age
0.0
0.5
1.0
1.5
2.0
0.00
0.25
0.50
0.75
1.00
0
1
2
30
60
90

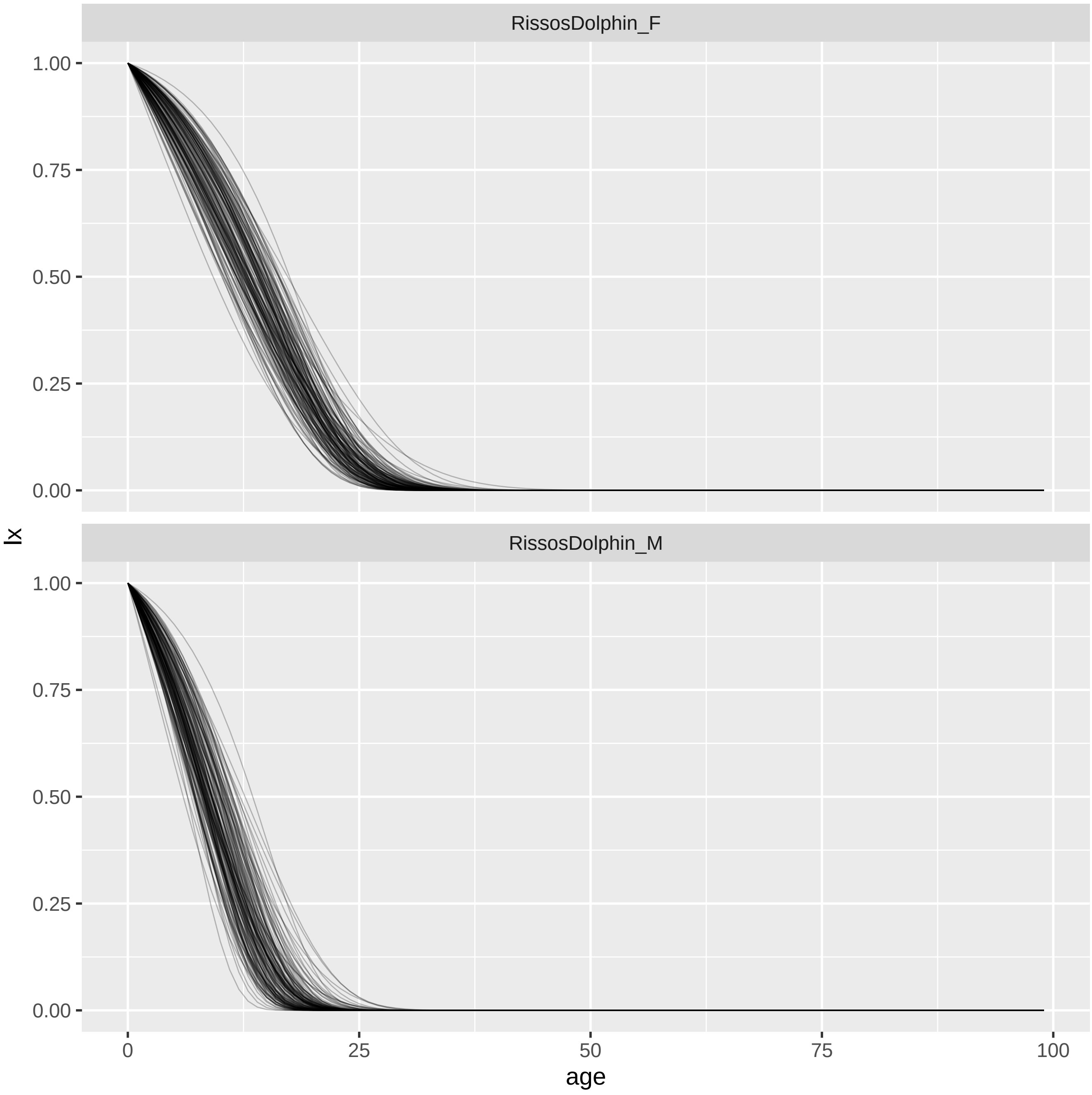
RissosDolphin_F
RissosDolphin_M
1.00
0.75
0.50
0.25
0.00
lx
0
25
50
75
100
age

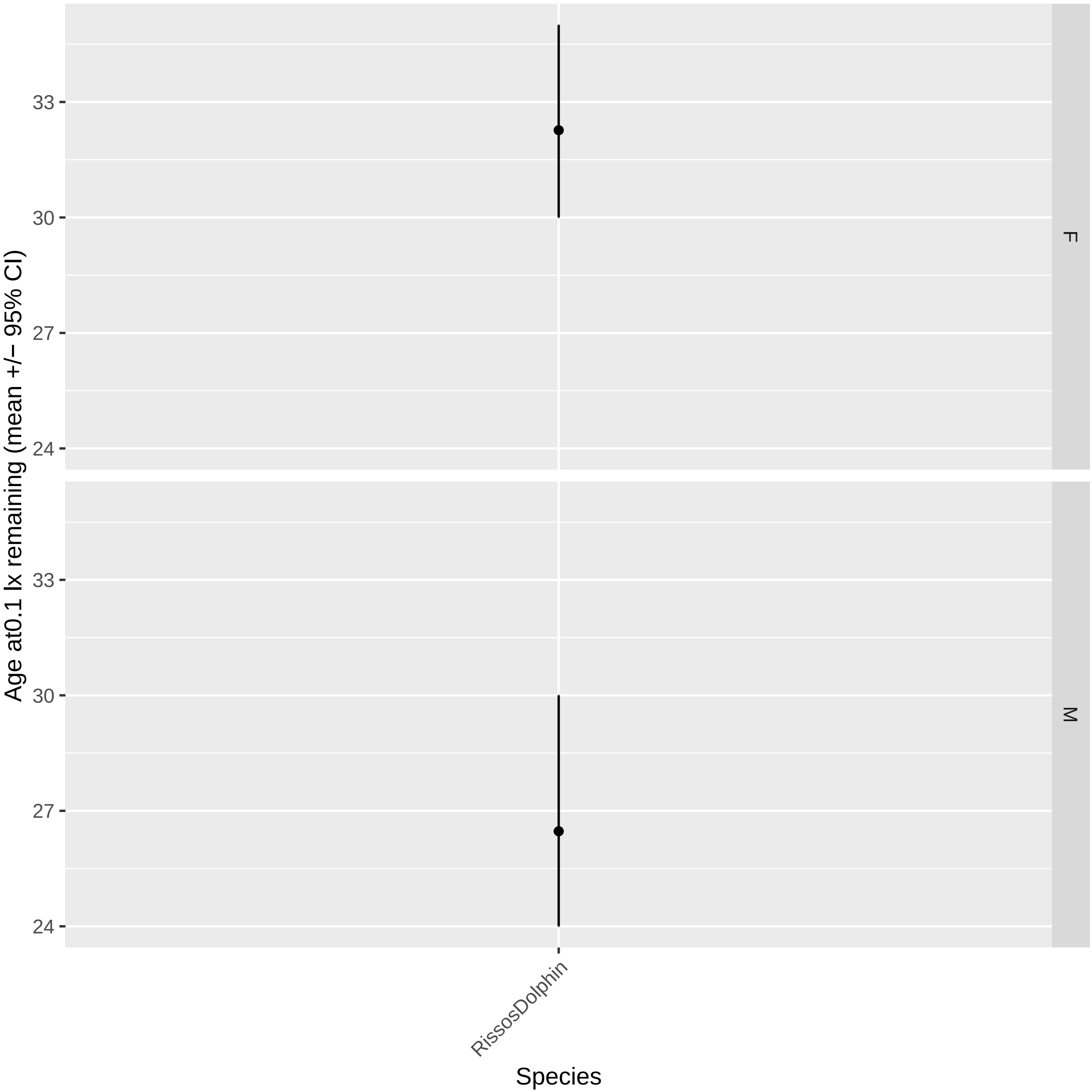

Age at0.1 lx remaining (mean +/− 95% CI)
33
30
27
24
F
M
RissosDolphin
Species

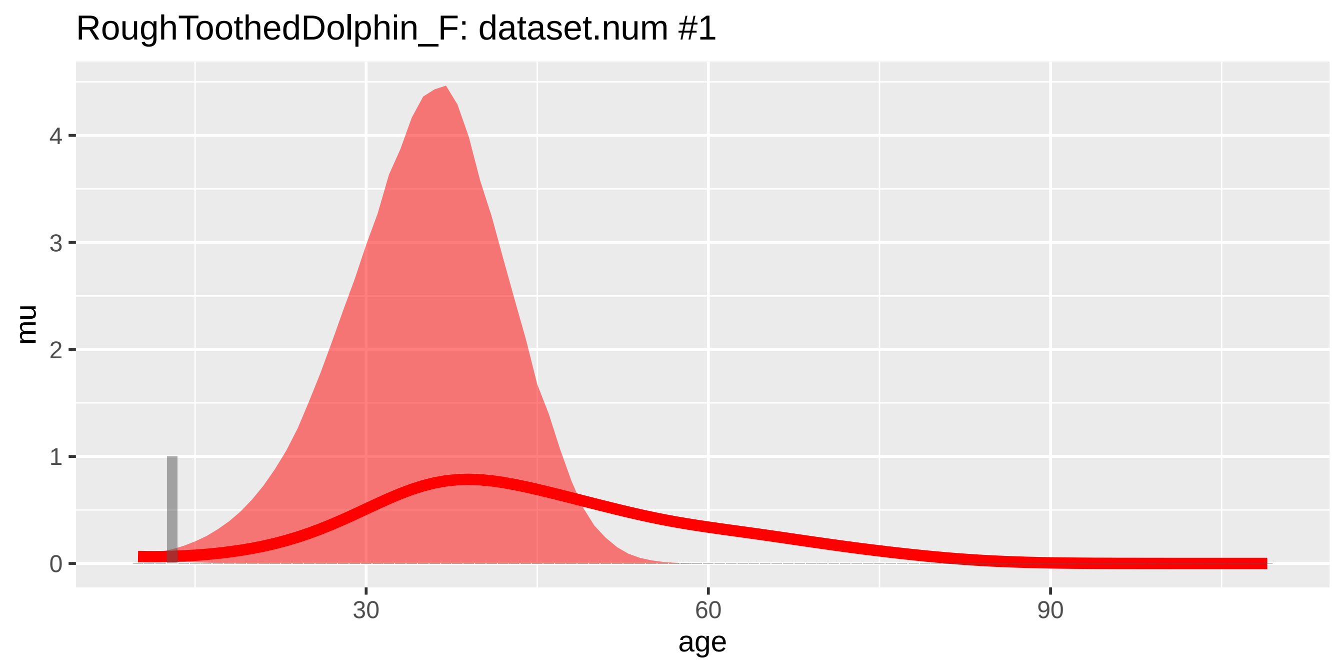
RoughToothedDolphin_F: dataset.num #1
mu
0
1
2
3
4
30
60
90
age

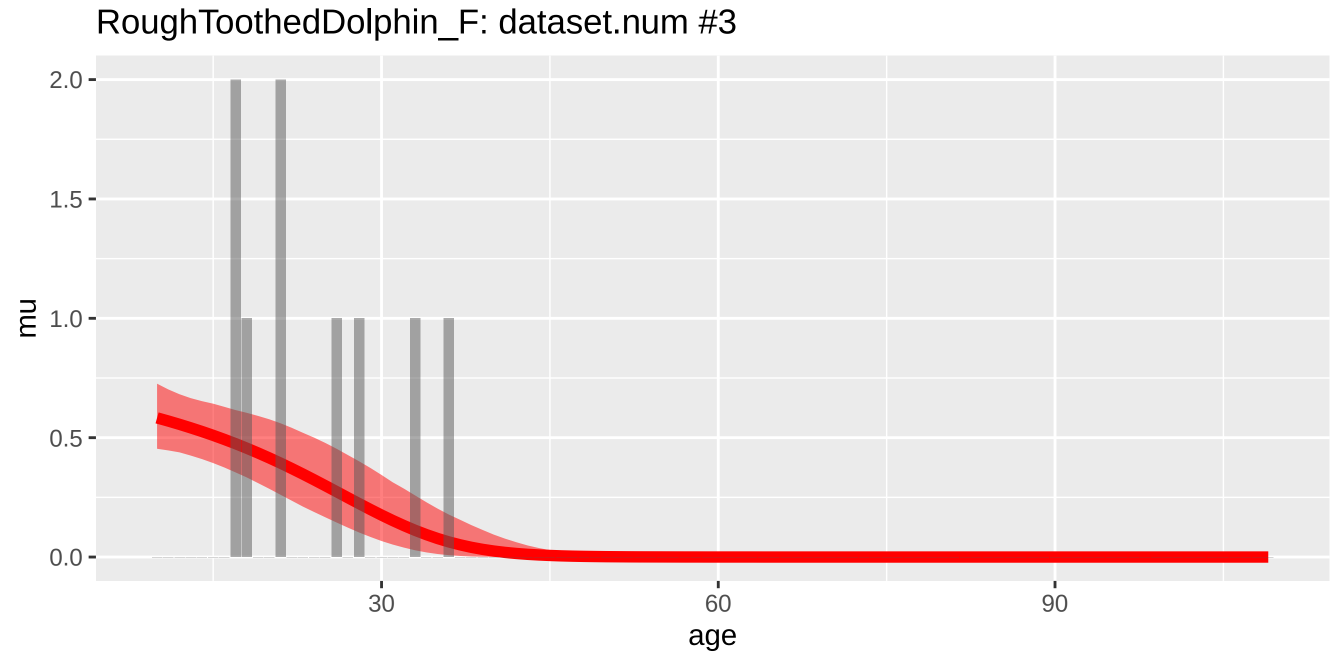
RoughToothedDolphin_F: dataset.num #3
mu
0.0
0.5
1.0
1.5
2.0
30
60
90
age

RoughToothedDolphin_M: dataset.num #2

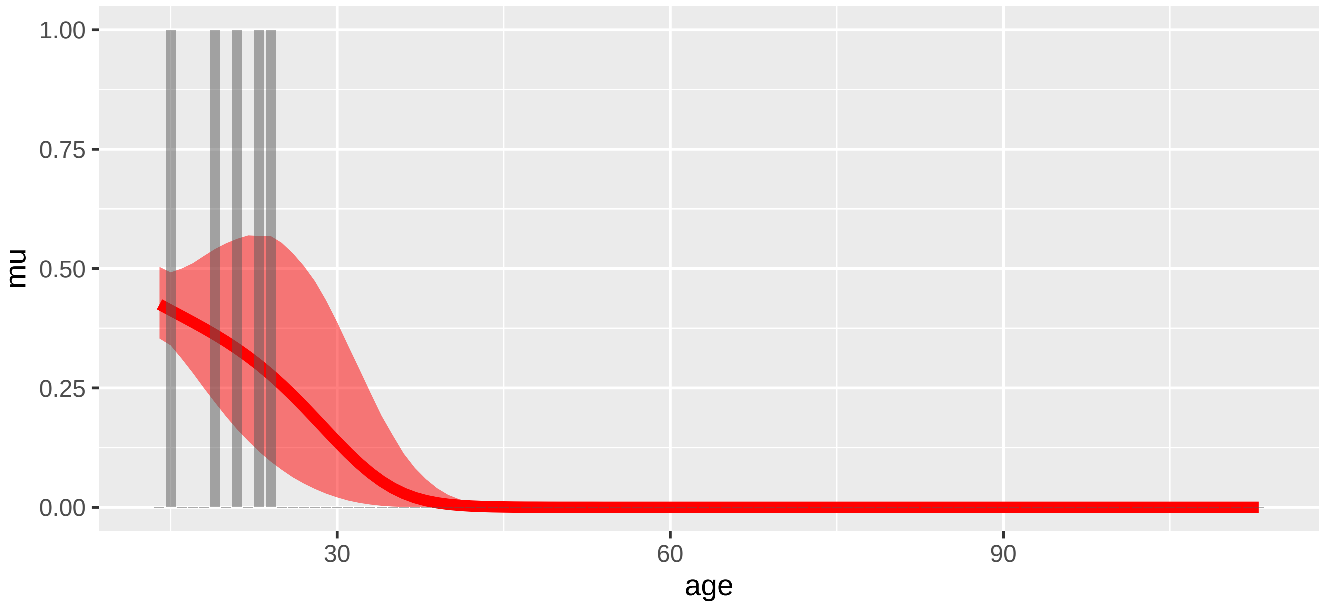


RoughToothedDolphin_M: dataset.num #4

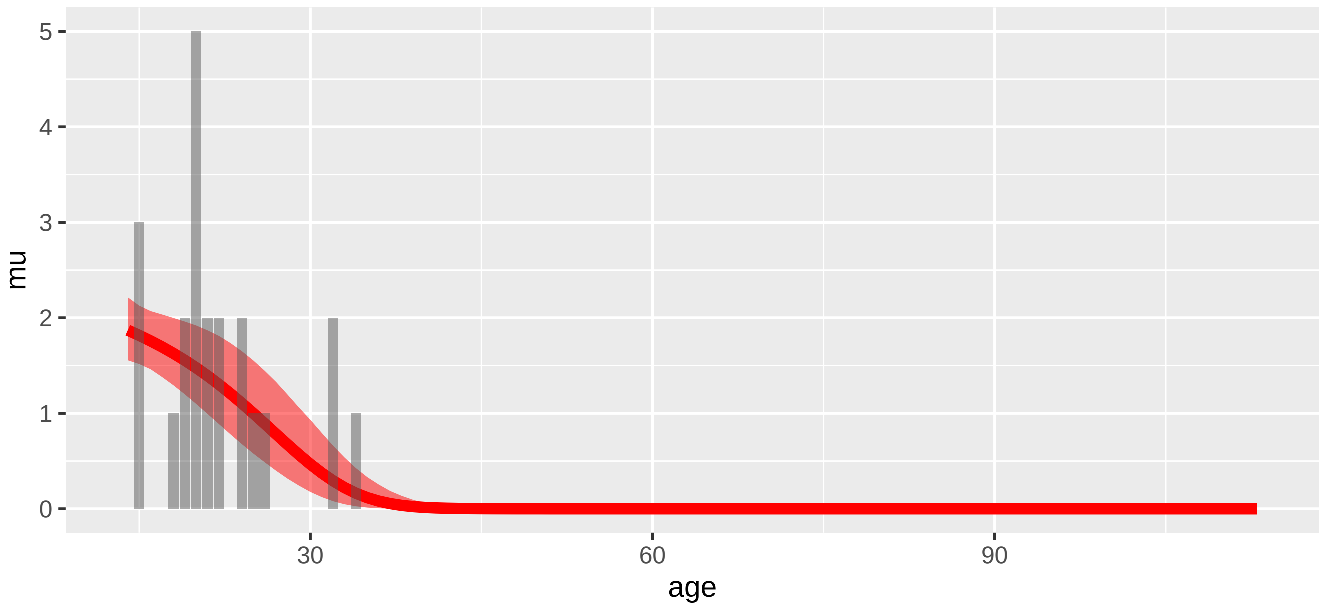

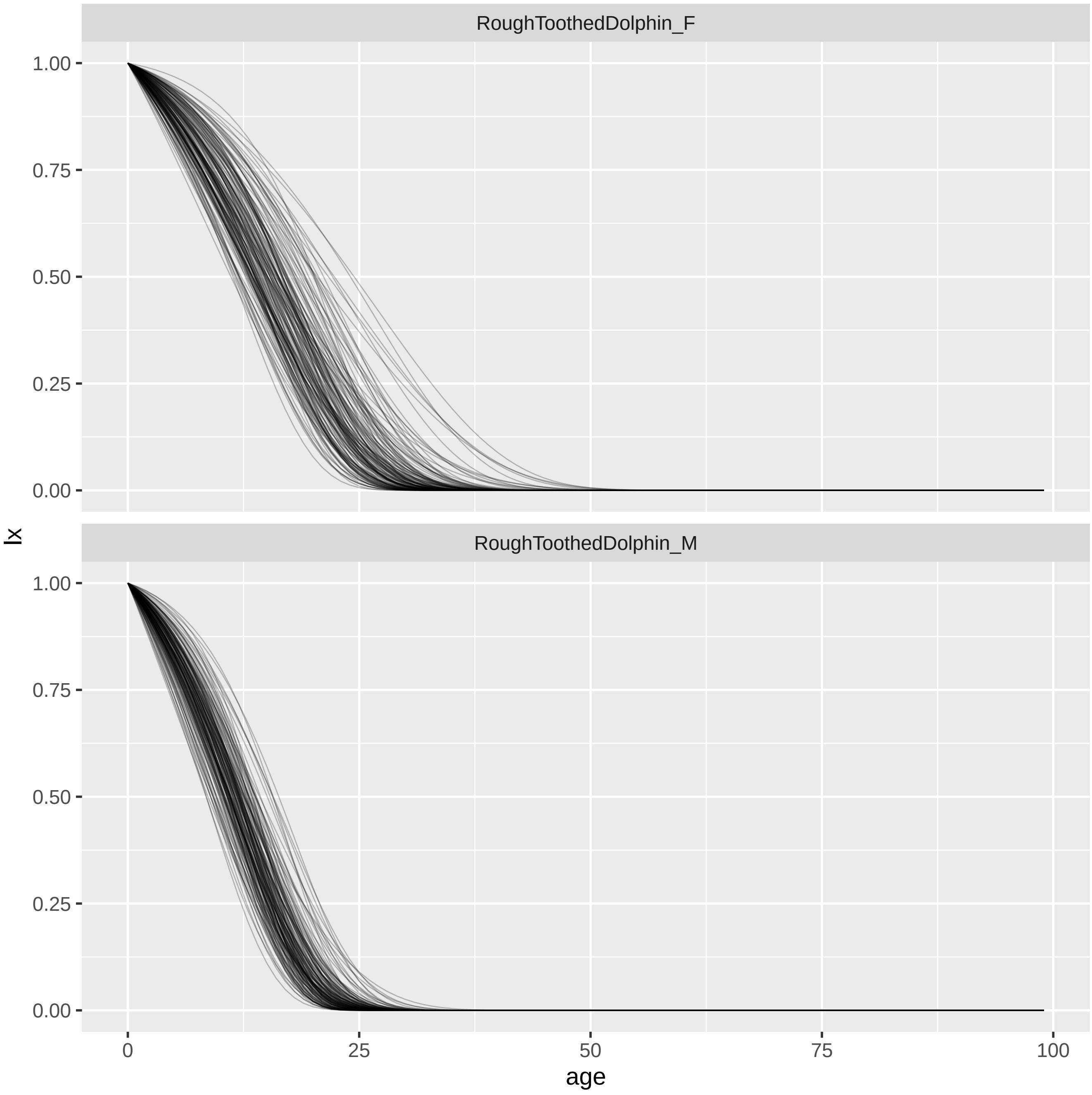
RoughToothedDolphin_F
RoughToothedDolphin_M
lx
age
1.00
0.75
0.50
0.25
0.00
0
25
50
75
100

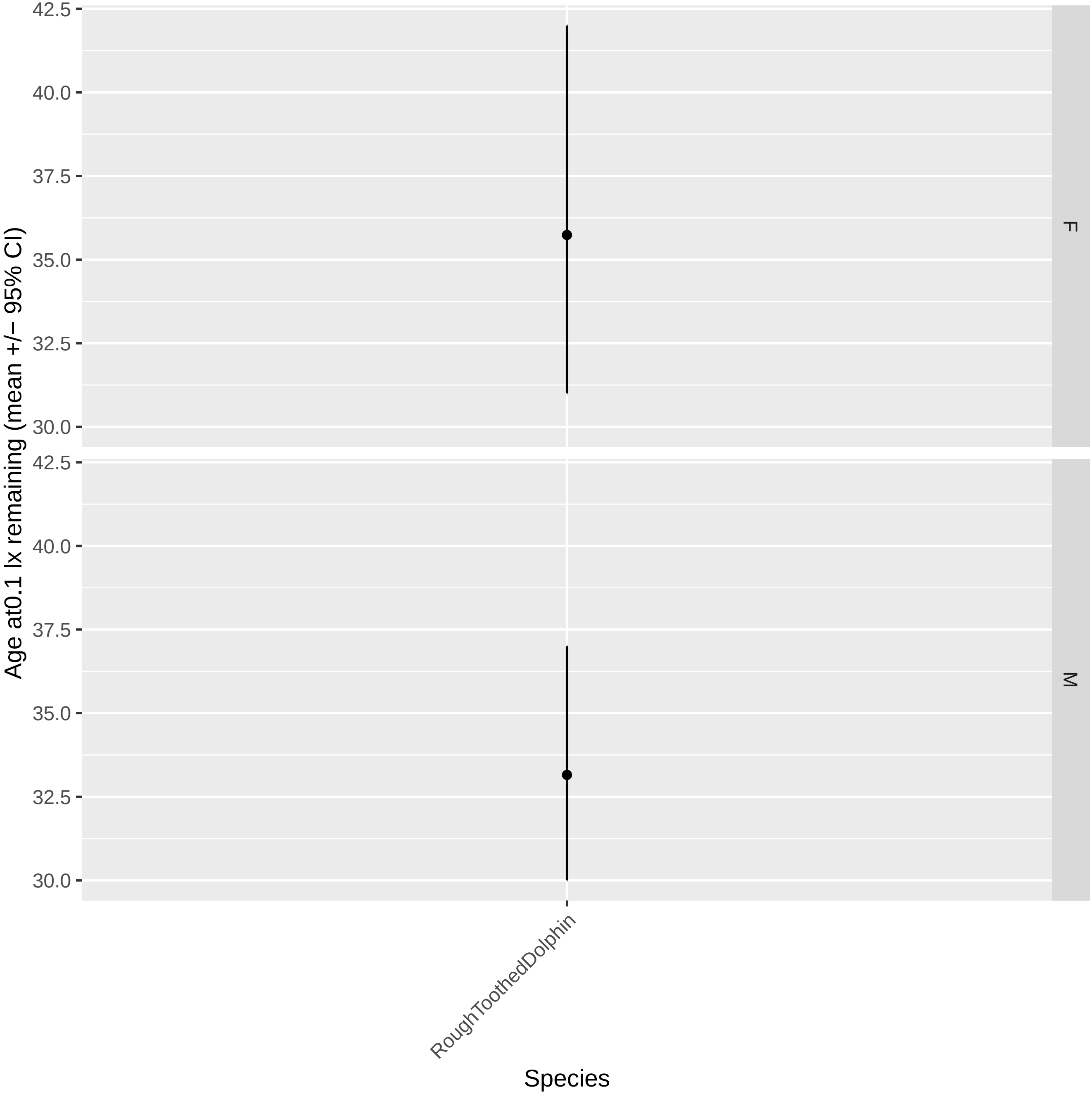

Age at0.1 lx remaining (mean +/− 95% CI)
42.5
40.0
37.5
35.0
32.5
30.0
F
M
RoughToothedDolphin
Species

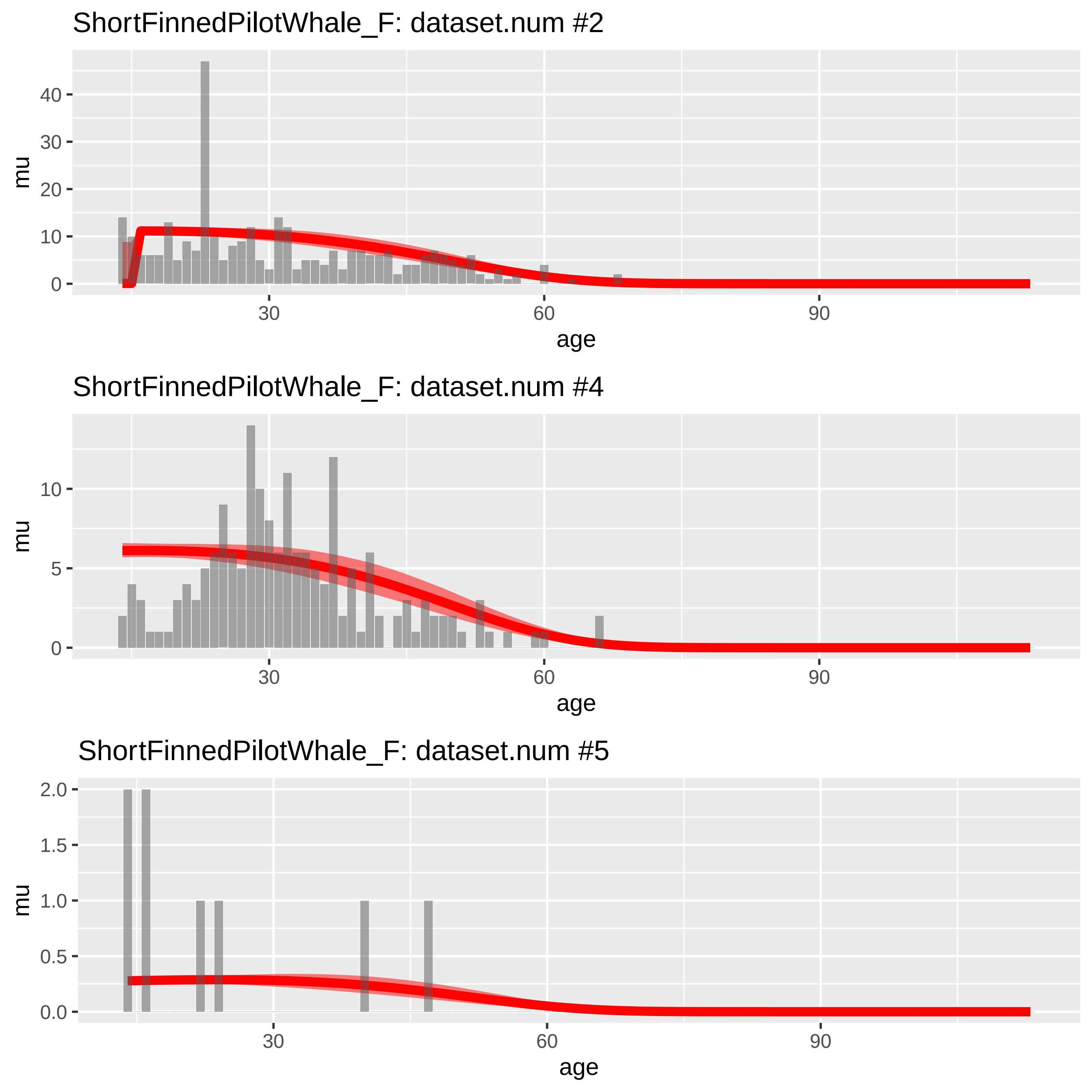
ShortFinnedPilotWhale_F: dataset.num #2
mu
40
30
20
10
0
30
60
90
age
ShortFinnedPilotWhale_F: dataset.num #4
mu
10
5
0
30
60
90
age
ShortFinnedPilotWhale_F: dataset.num #5
mu
2.0
1.5
1.0
0.5
0.0
30
60
90
age

ShortFinnedPilotWhale_M: dataset.num #1

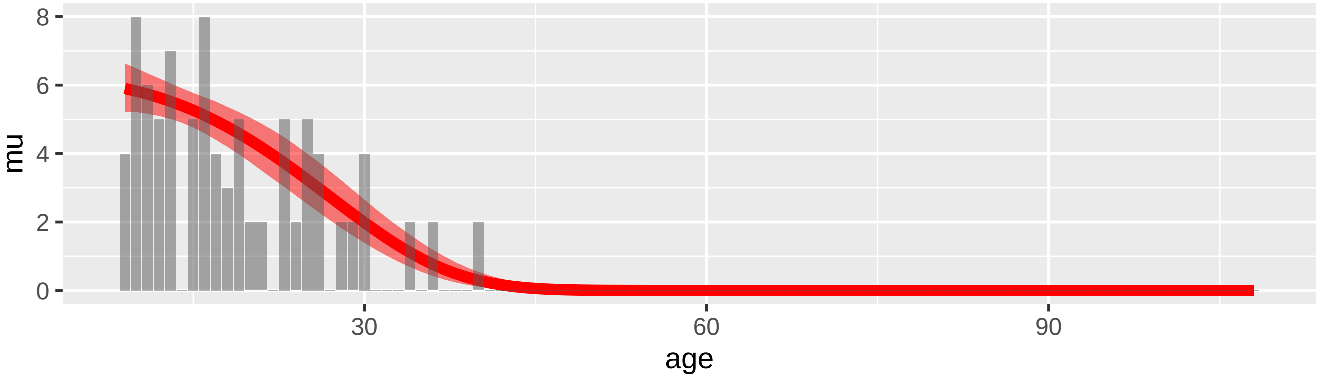


ShortFinnedPilotWhale_M: dataset.num #3

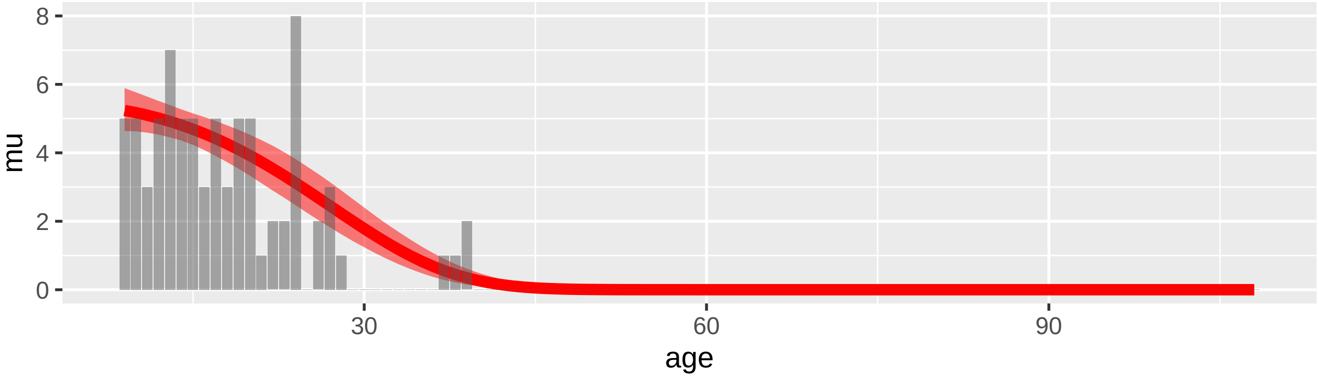


ShortFinnedPilotWhale_M: dataset.num #6

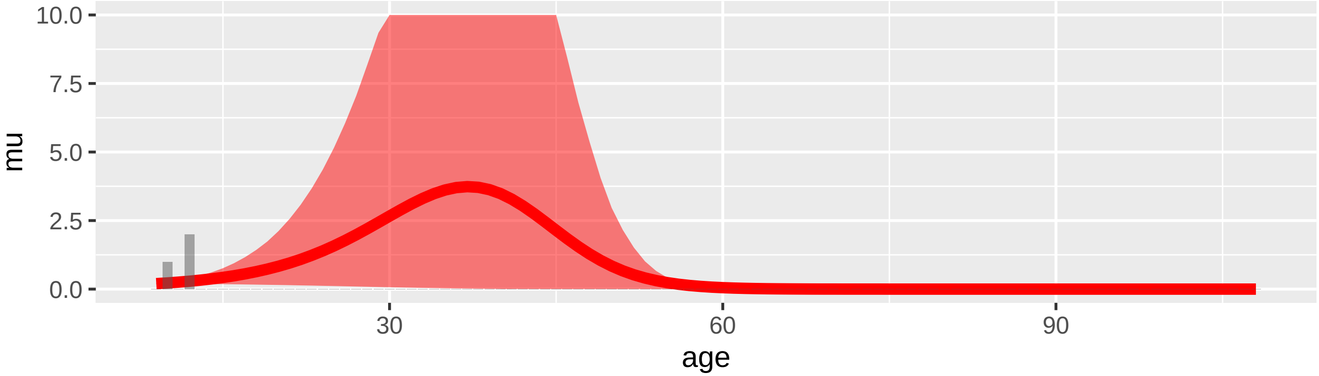

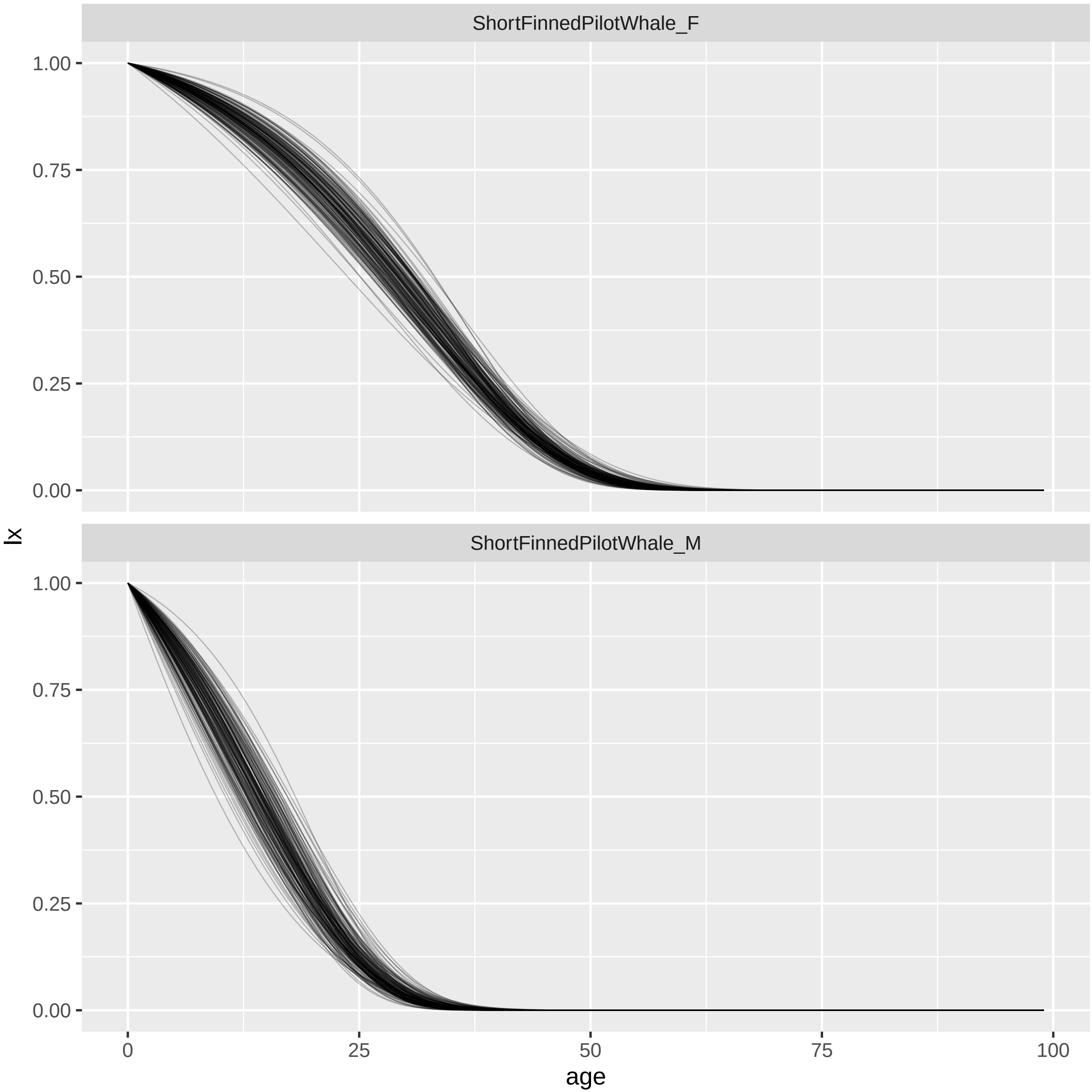
ShortFinnedPilotWhale_F
ShortFinnedPilotWhale_M
lx
1.00
0.75
0.50
0.25
0.00
0
25
50
75
100
age

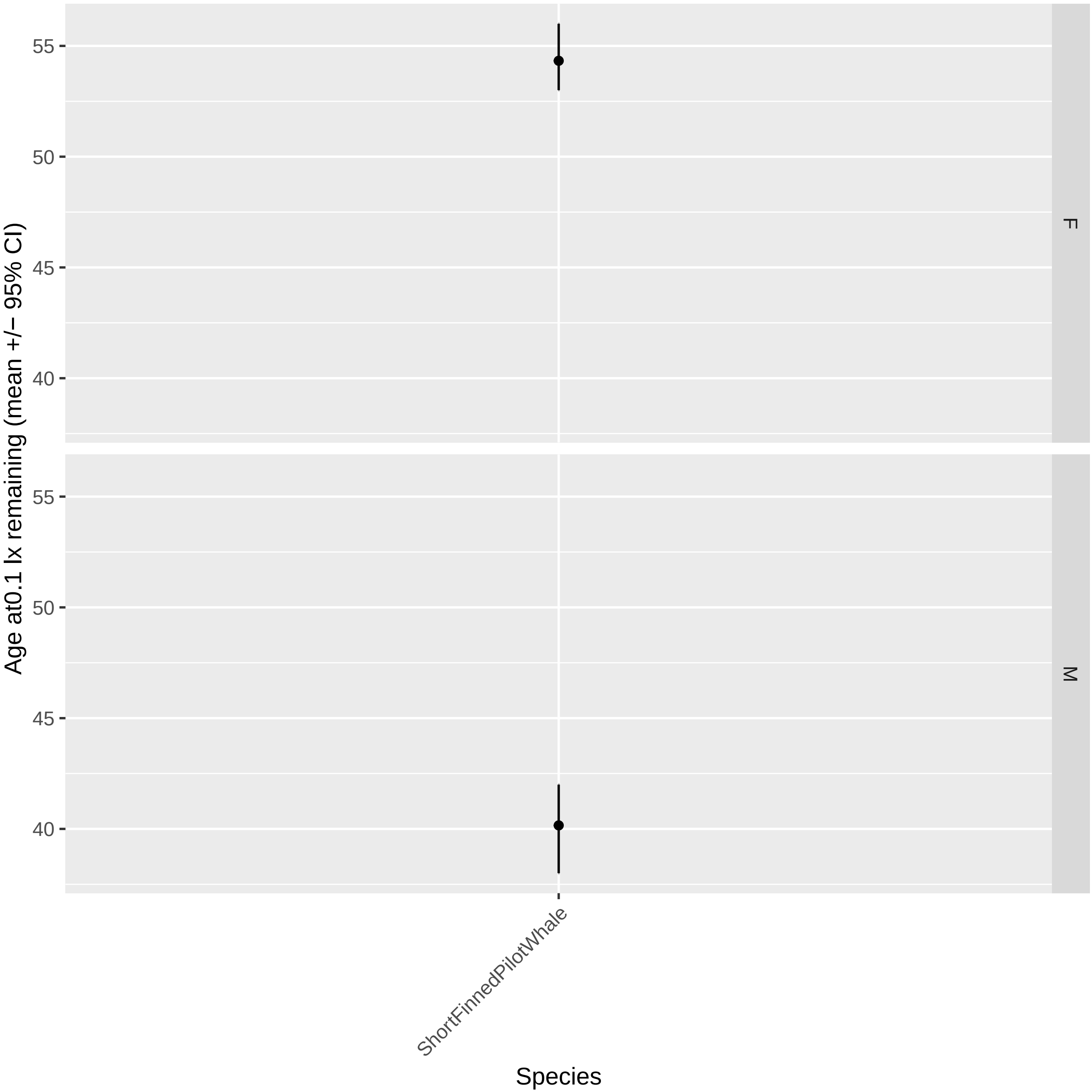

Age at0.1 lx remaining (mean +/− 95% CI)
55
50
45
40
F
M
ShortFinnedPilotWhale
Species

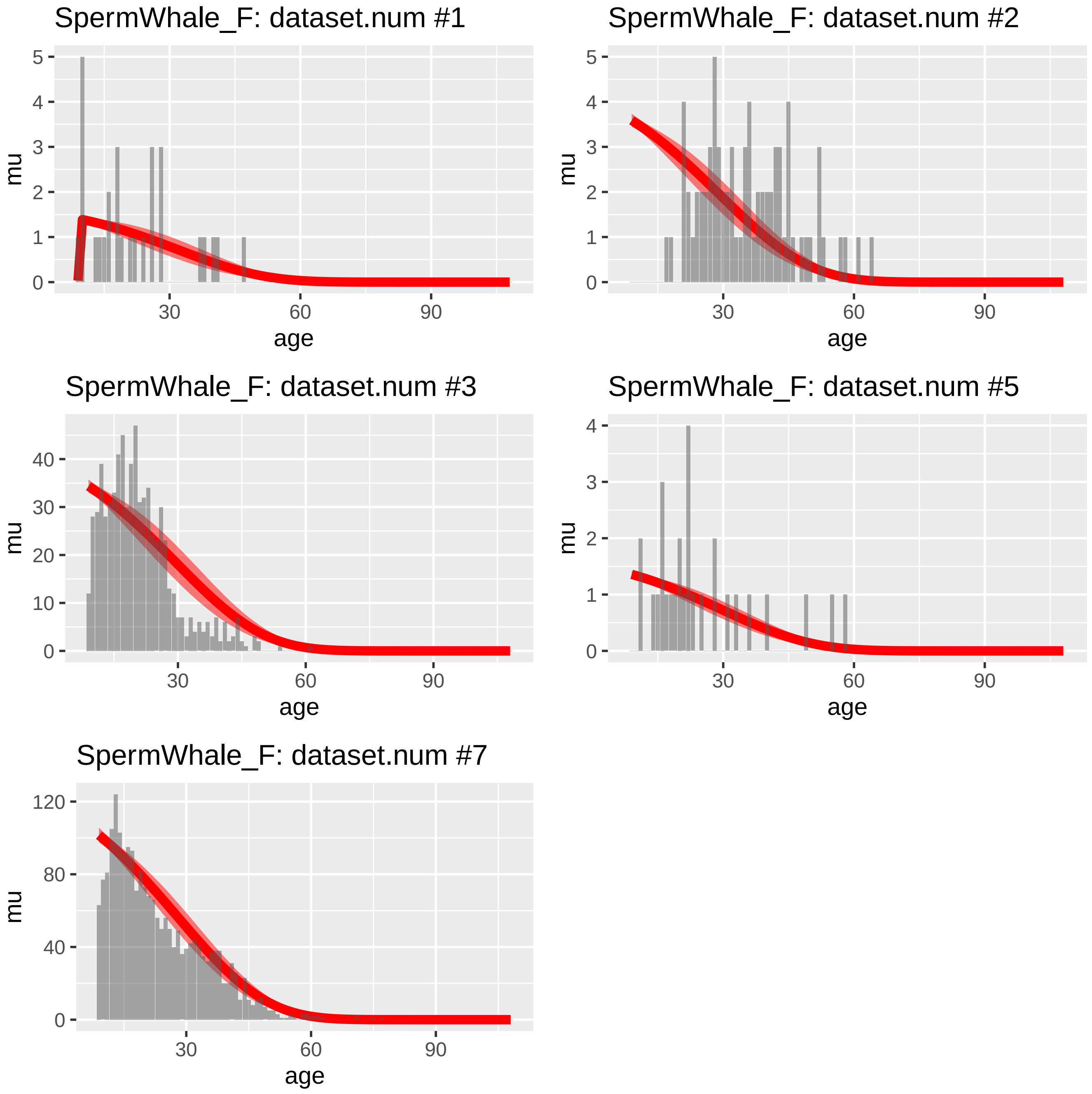
SpermWhale_F: dataset.num #1
SpermWhale_F: dataset.num #2
SpermWhale_F: dataset.num #3
SpermWhale_F: dataset.num #5
SpermWhale_F: dataset.num #7
mu
age
0
1
2
3
4
5
10
20
30
40
80
120
60
90

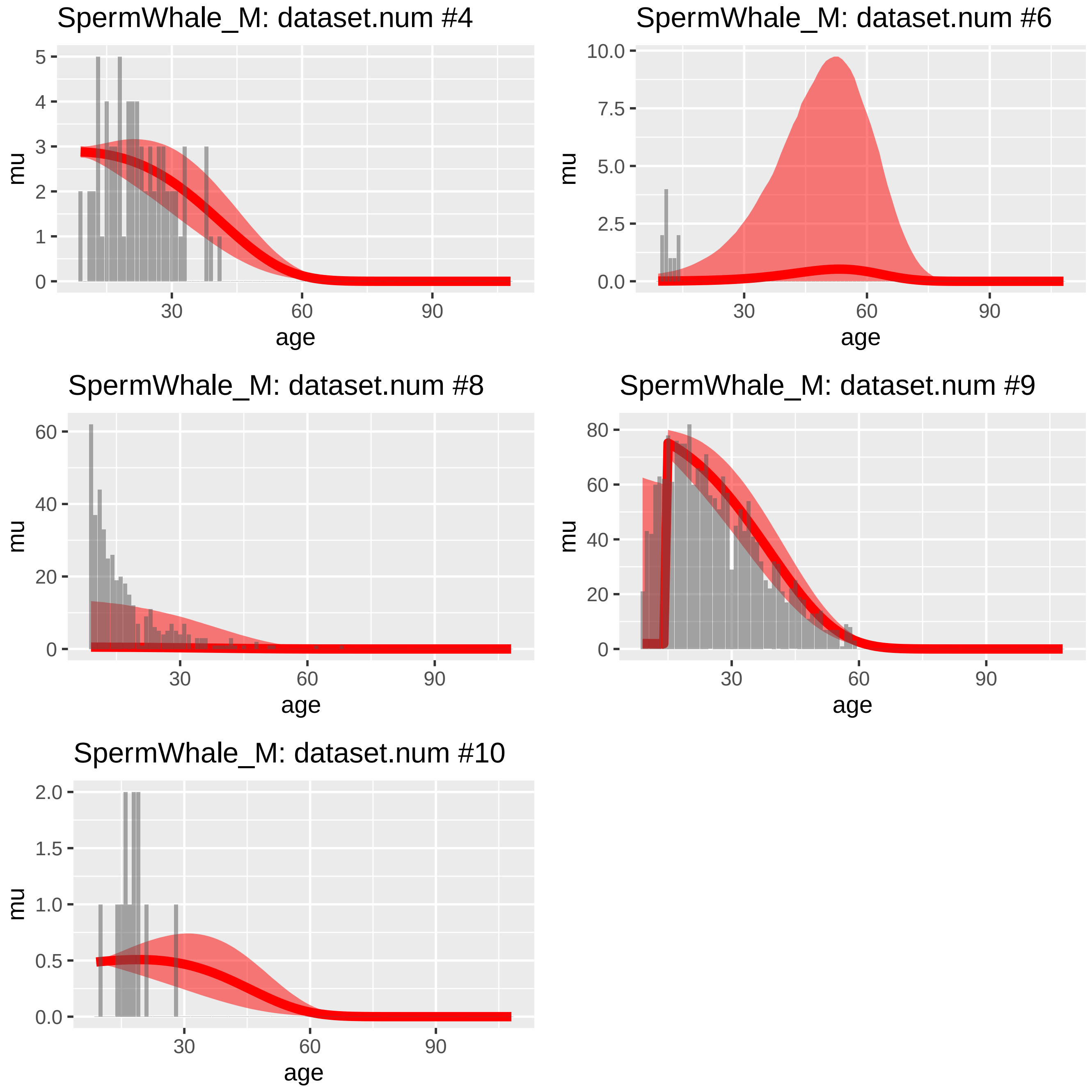
SpermWhale_M: dataset.num #4
mu
0
1
2
3
4
5
30
60
90
age
SpermWhale_M: dataset.num #6
mu
0.0
2.5
5.0
7.5
10.0
30
60
90
age
SpermWhale_M: dataset.num #8
mu
0
20
40
60
30
60
90
age
SpermWhale_M: dataset.num #9
mu
0
20
40
60
80
30
60
90
age
SpermWhale_M: dataset.num #10
mu
0.0
0.5
1.0
1.5
2.0
30
60
90
age

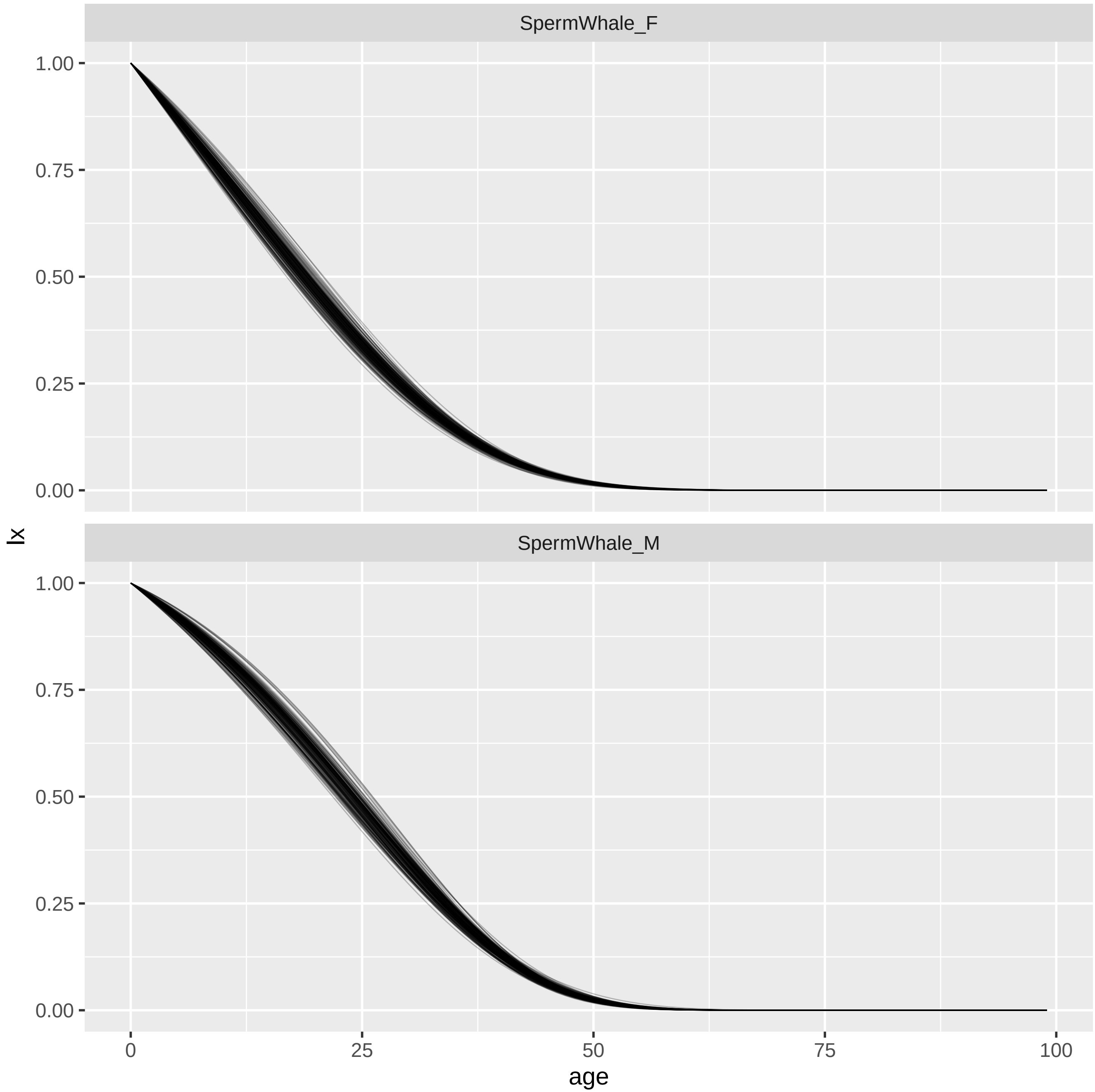

SpermWhale_F
SpermWhale_M
1.00
0.75
0.50
0.25
0.00
lx
0
25
50
75
100
age

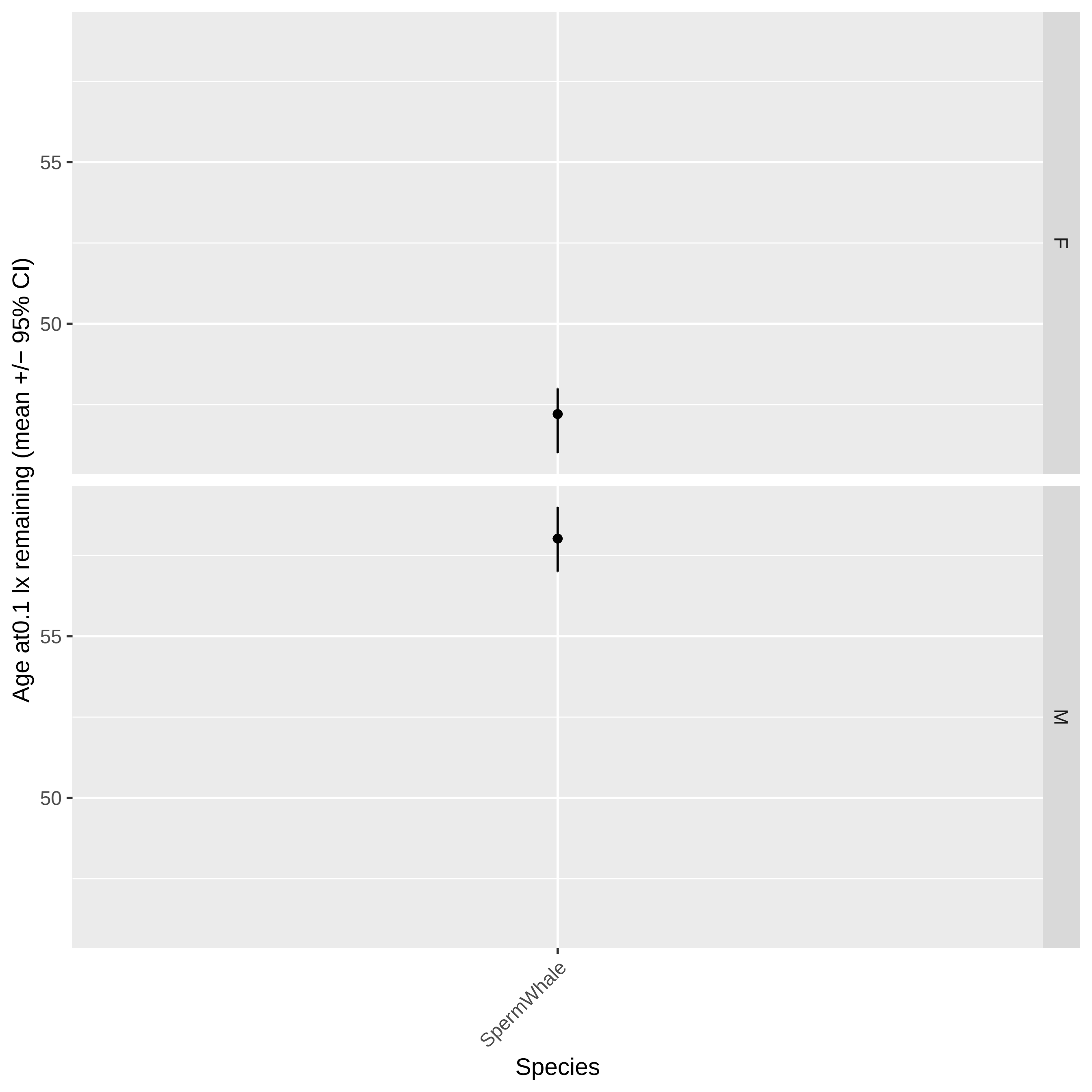
Age at0.1 lx remaining (mean +/− 95% CI)
55
50
F
55
50
M
SpermWhale
Species

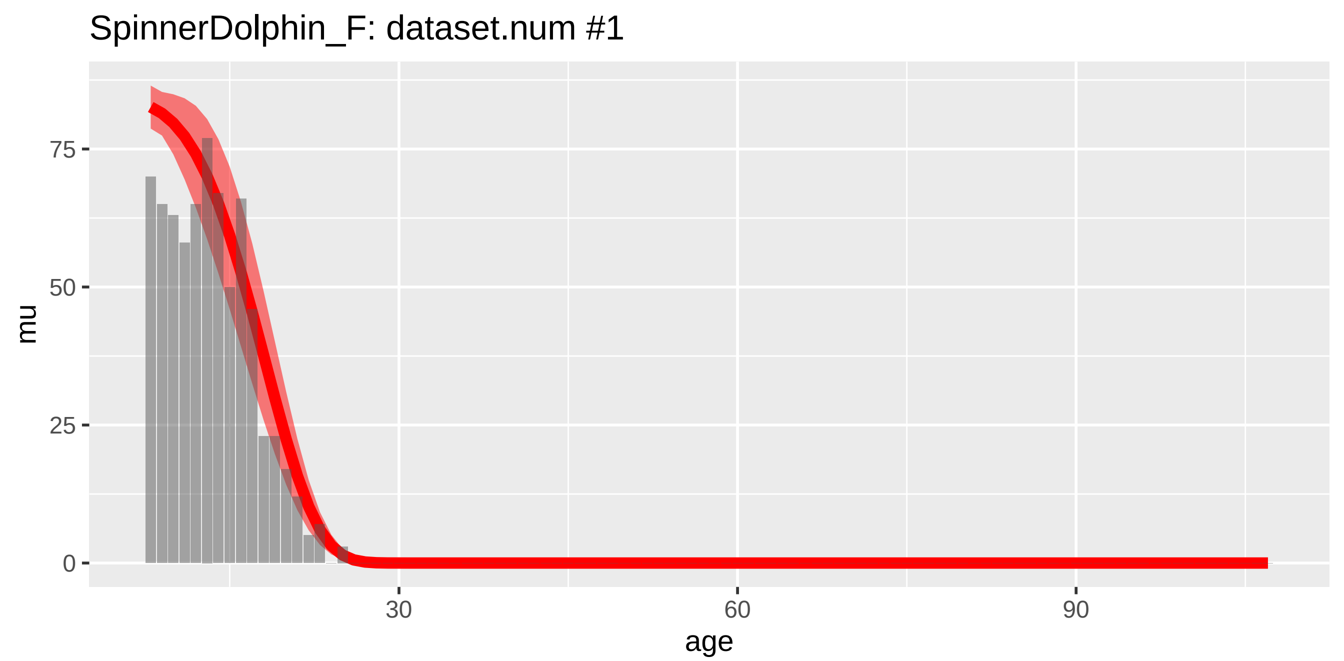
SpinnerDolphin_F: dataset.num #1
mu
75
50
25
0
30
60
90
age

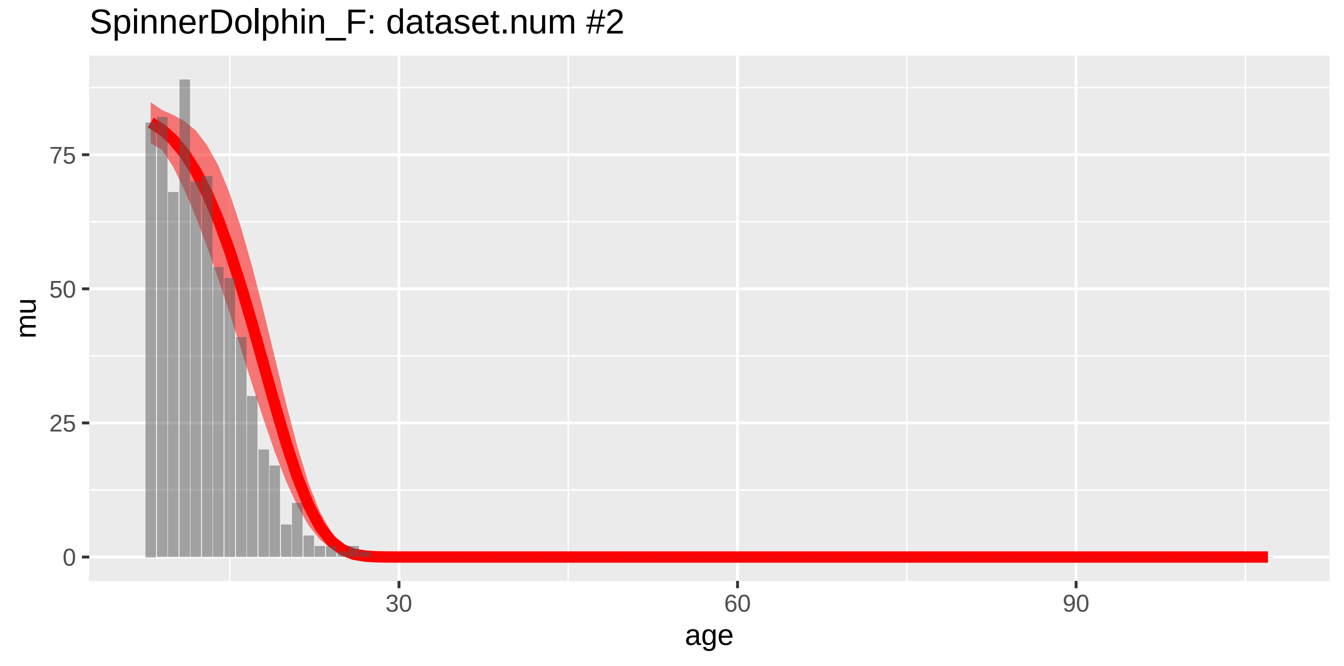
SpinnerDolphin_F: dataset.num #2
mu
75
50
25
0
30
60
90
age

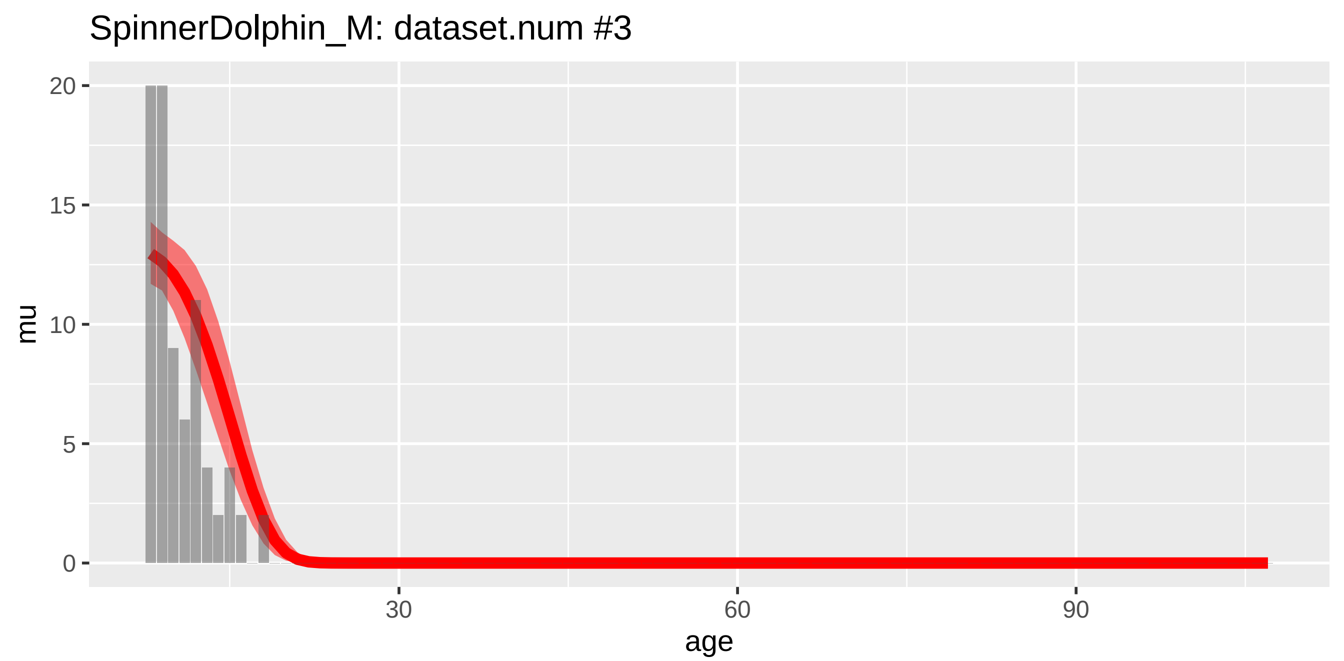
SpinnerDolphin_M: dataset.num #3
20
15
10
5
0
mu
30
60
90
age

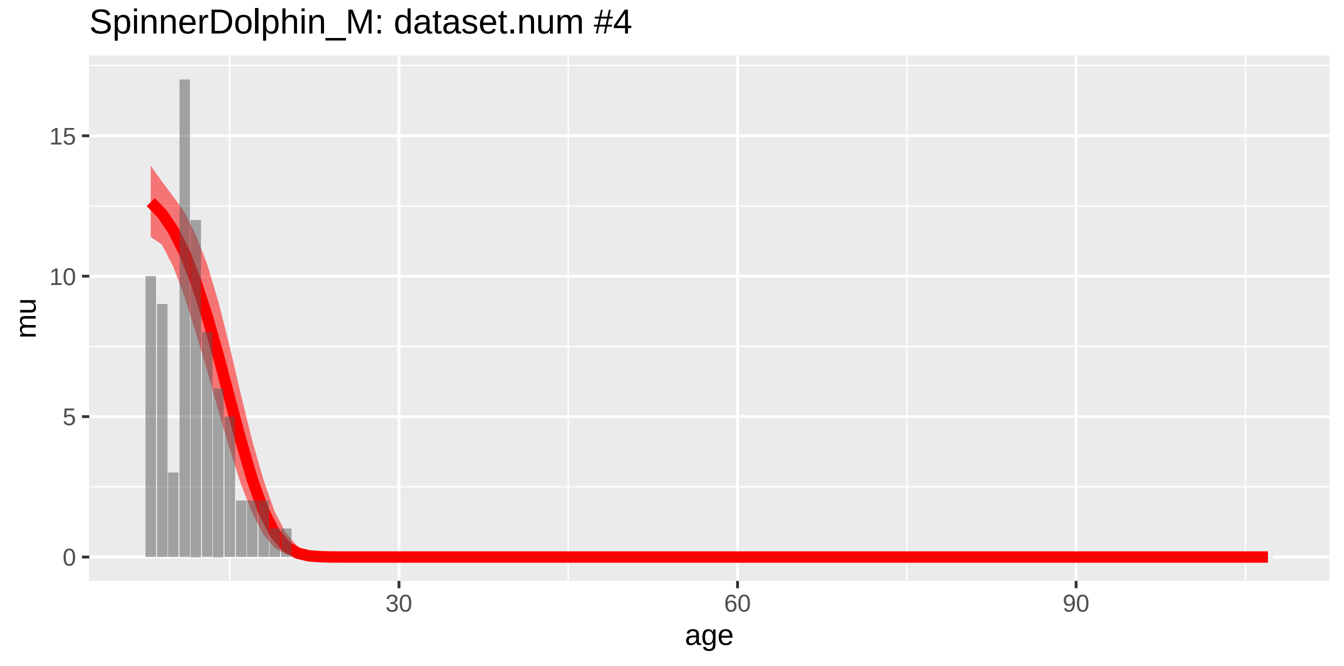
SpinnerDolphin_M: dataset.num #4
15
10
5
0
mu
30
60
90
age

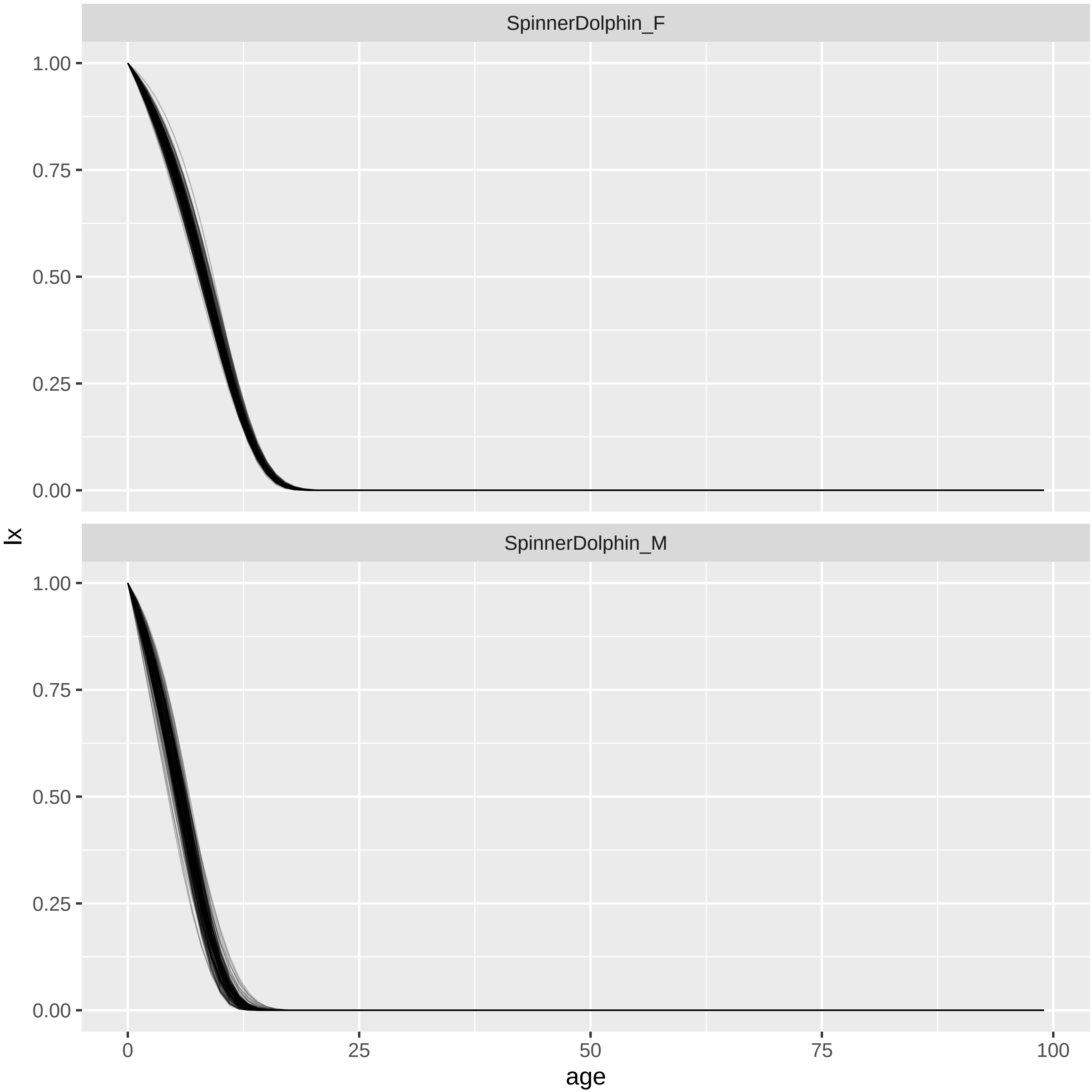
SpinnerDolphin_F
SpinnerDolphin_M
1.00
0.75
0.50
0.25
0.00
0
25
50
75
100
age
lx

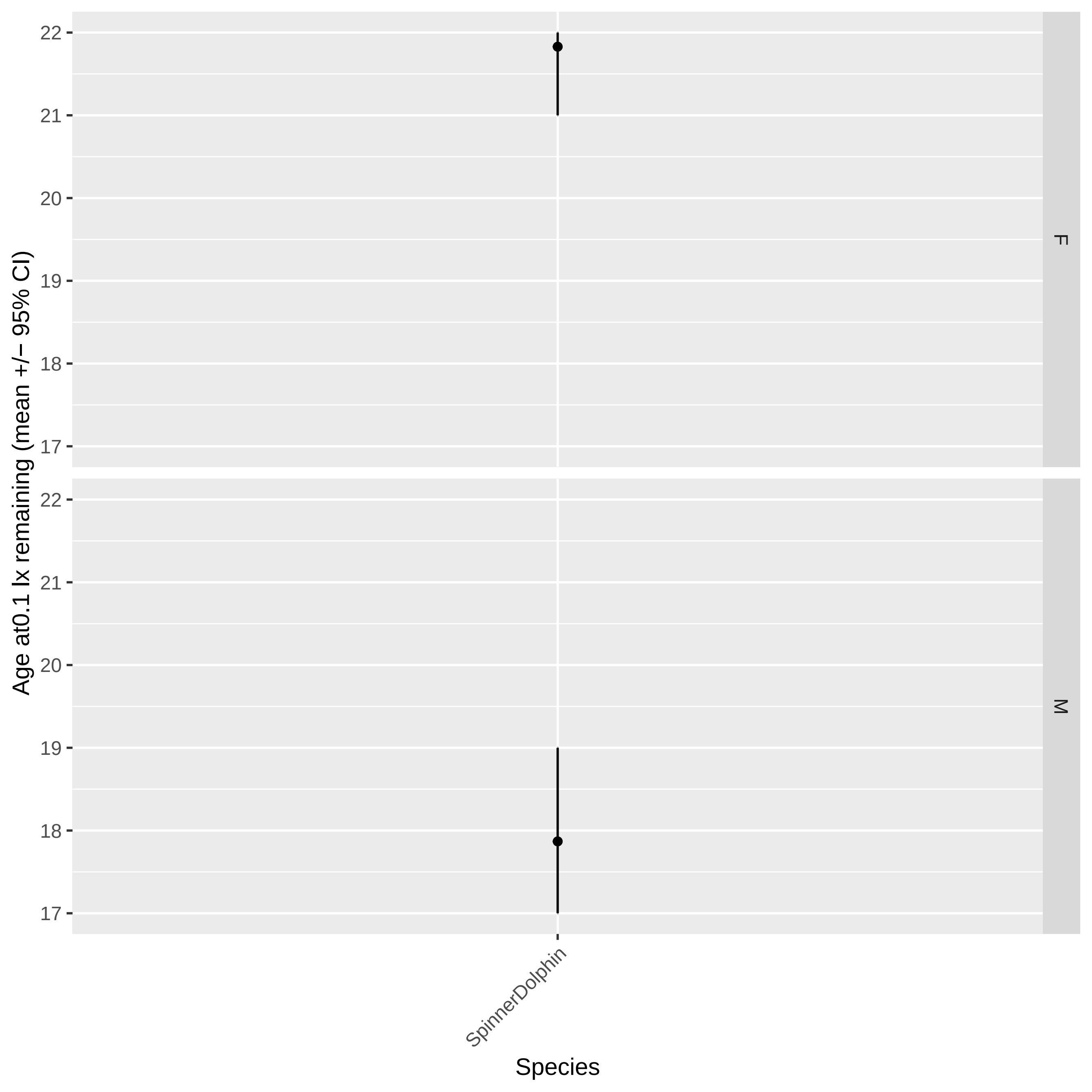

Age at0.1 lx remaining (mean +/− 95% CI)
22
21
20
19
18
17
F
M
SpinnerDolphin
Species

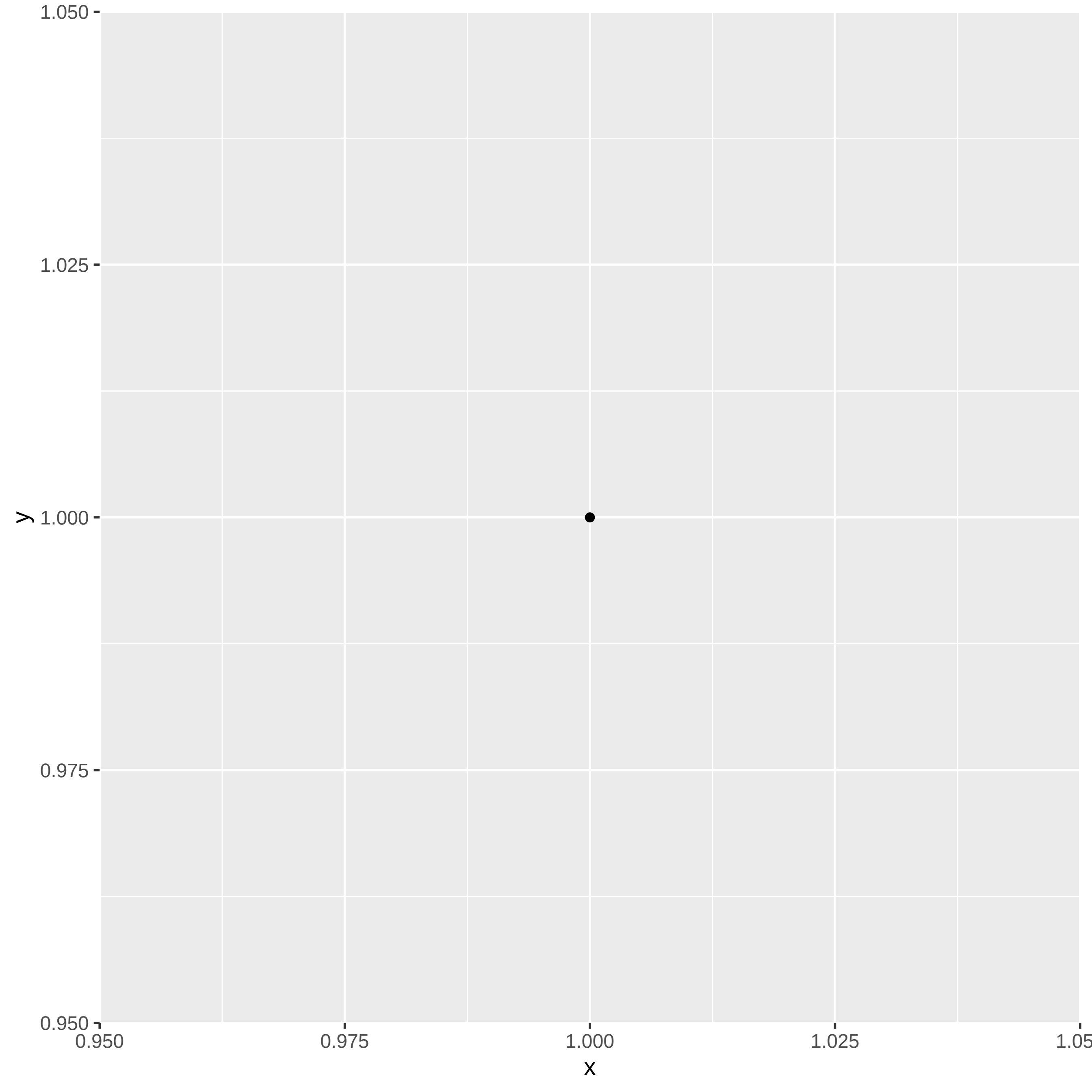
1.050
1.025
1.000
0.975
0.950
y
0.950
0.975
1.000
1.025
1.05
x

StejnegersBeakedWhale_M: dataset.num #1

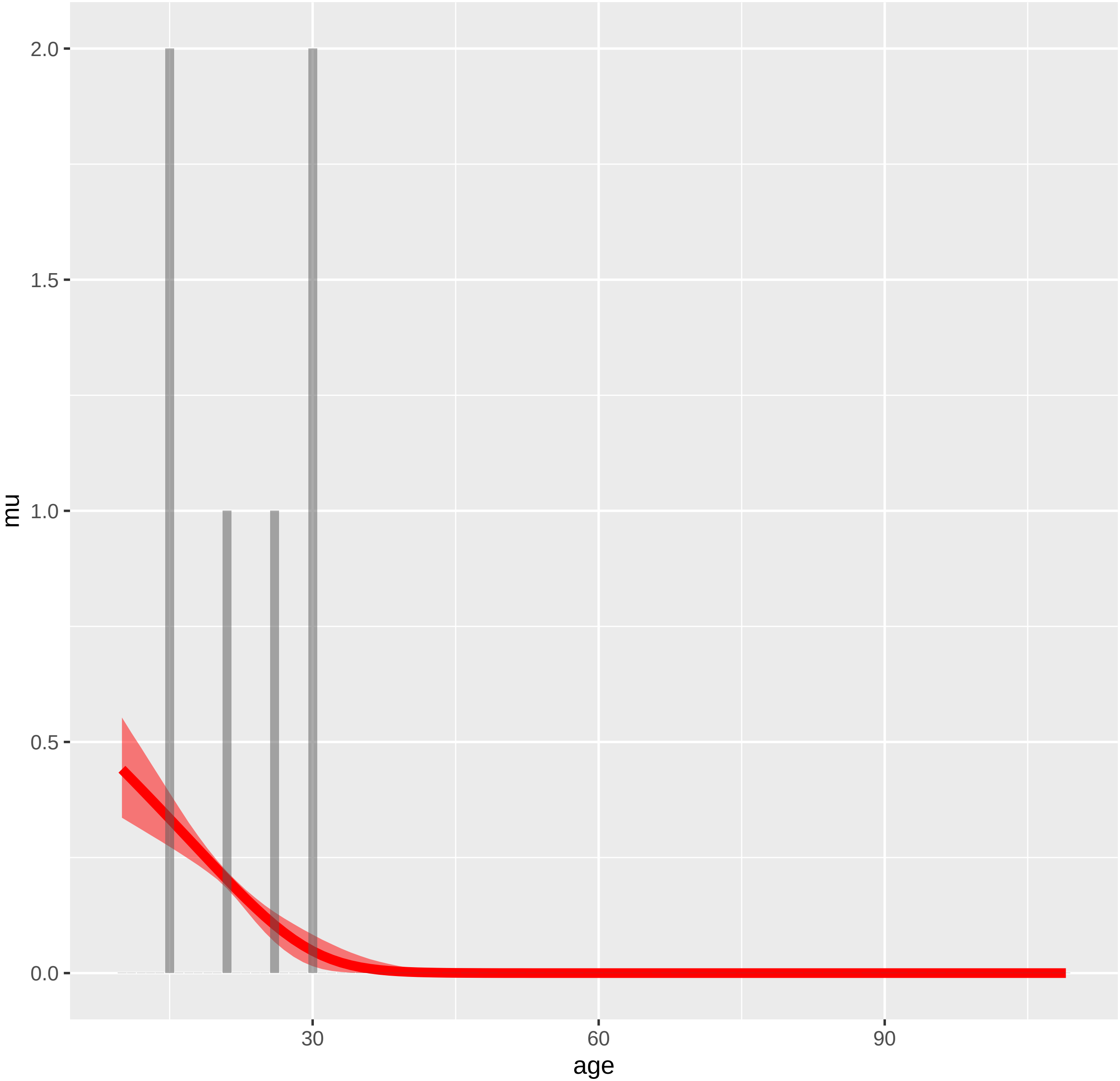

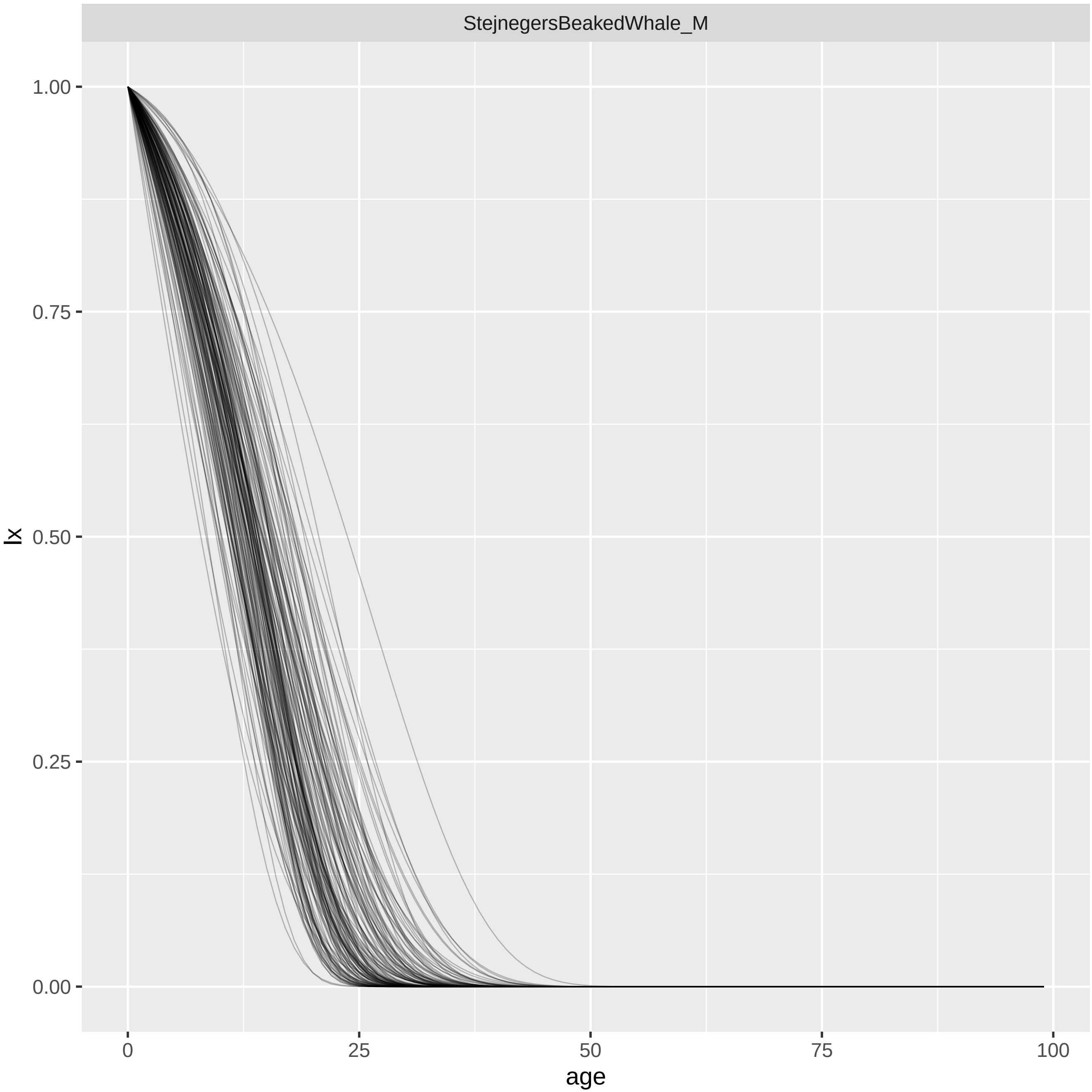
StejnegersBeakedWhale_M
lx
1.00
0.75
0.50
0.25
0.00
0
25
50
75
100
age

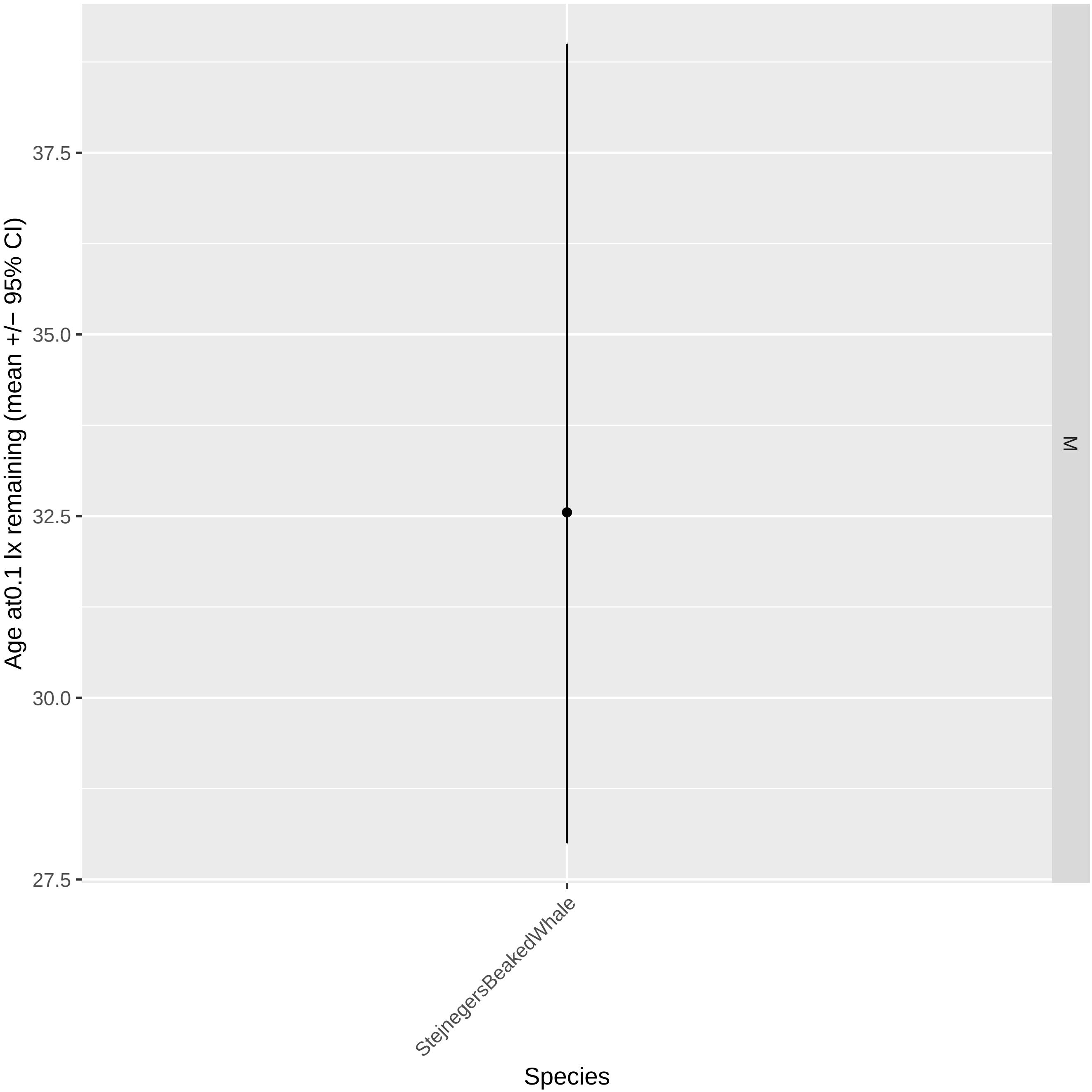
Age at0.1 lx remaining (mean +/− 95% CI)
37.5
35.0
32.5
30.0
27.5
M
StejnegersBeakedWhale
Species

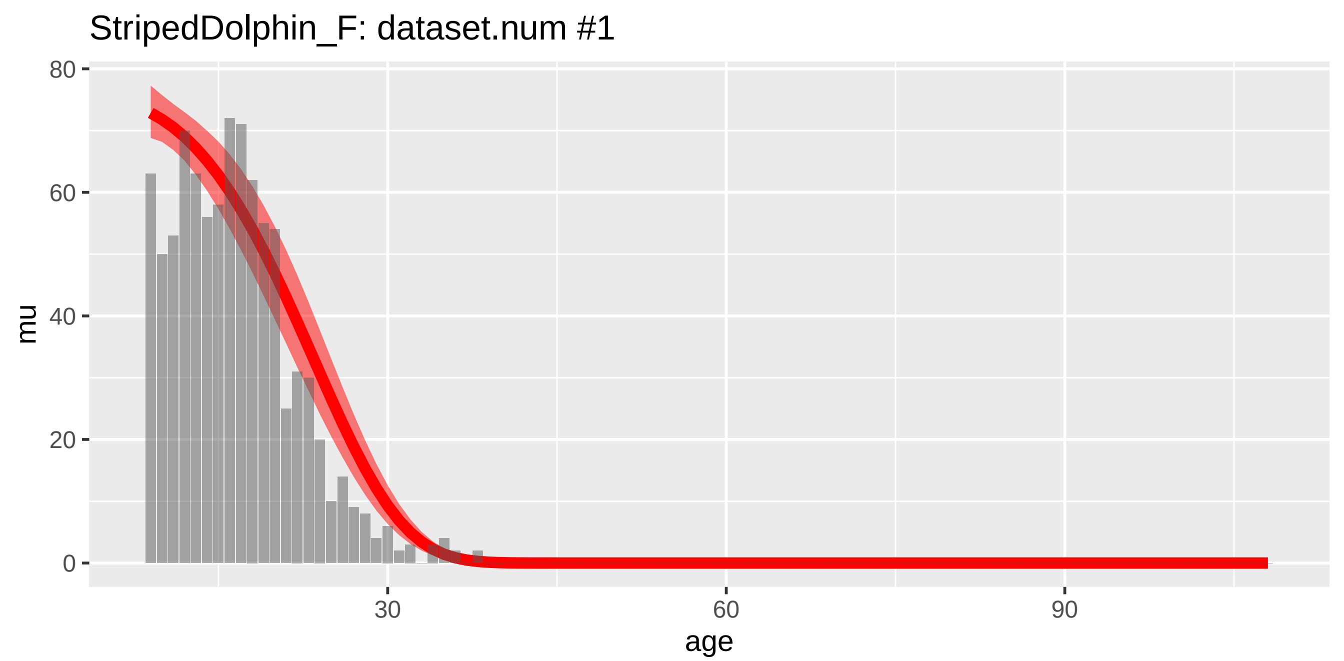
StripedDolphin_F: dataset.num #1
mu
80
60
40
20
0
30
60
90
age

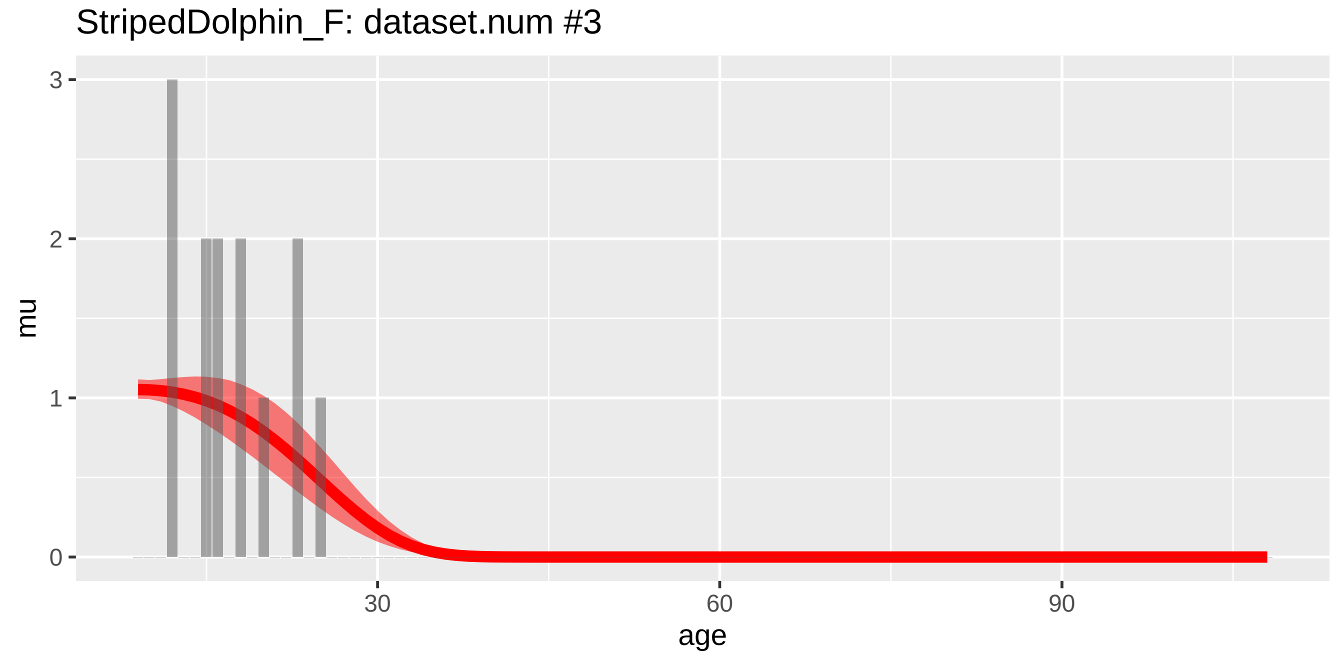
StripedDolphin_F: dataset.num #3
mu
3
2
1
0
30
60
90
age

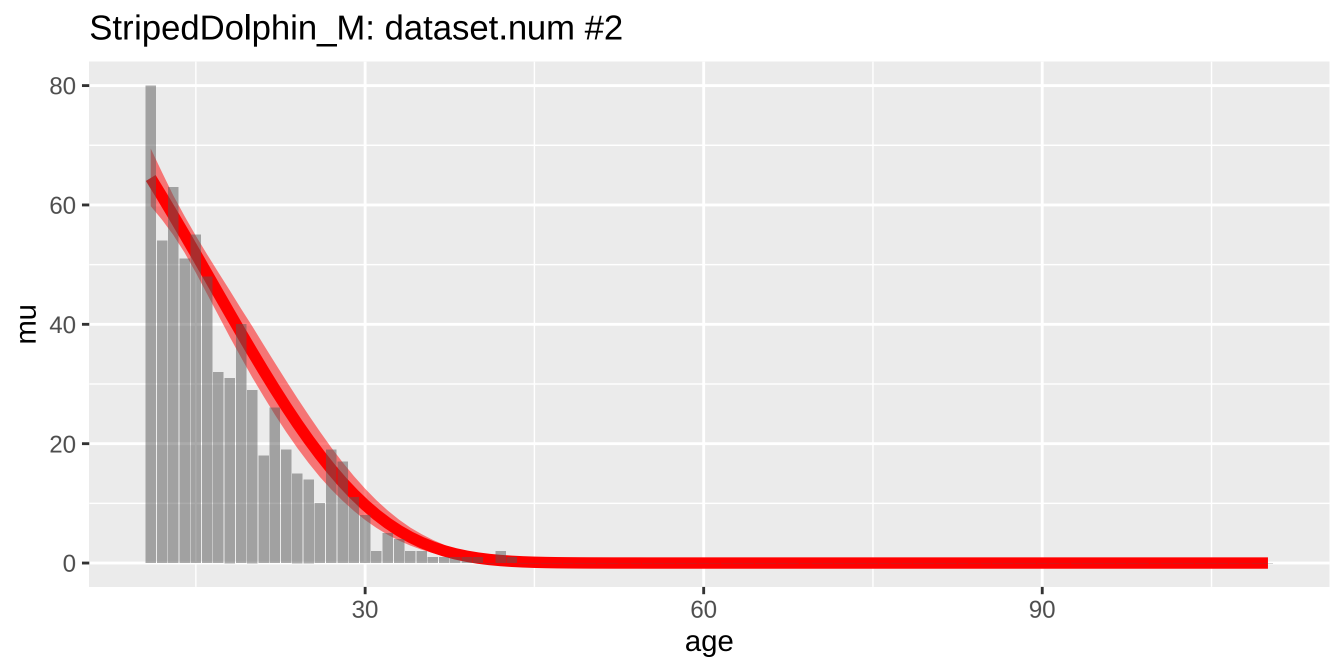

StripedDolphin_M: dataset.num #2
mu
80
60
40
20
0
30
60
90
age


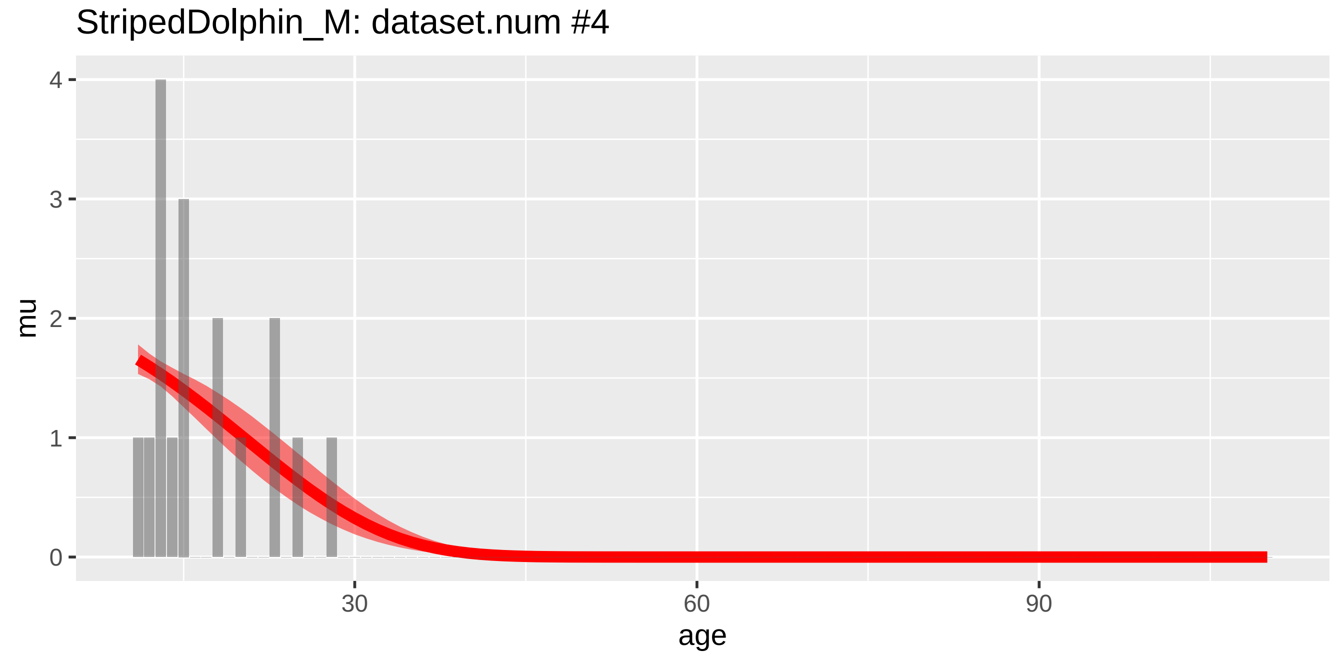

StripedDolphin_M: dataset.num #4
mu
4
3
2
1
0
30
60
90
age

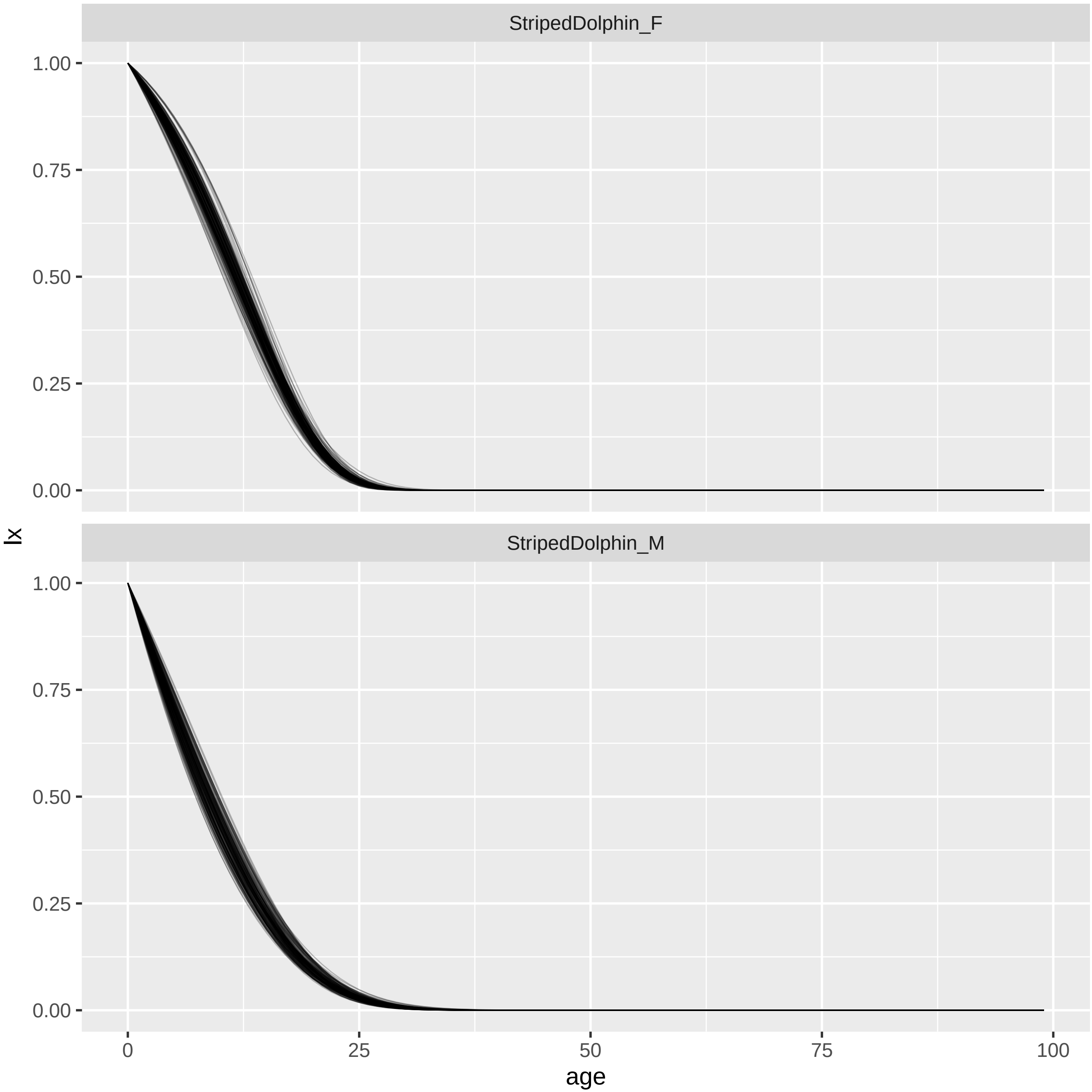

StripedDolphin_F
StripedDolphin_M
1.00
0.75
0.50
0.25
0.00
lx
0
25
50
75
100
age

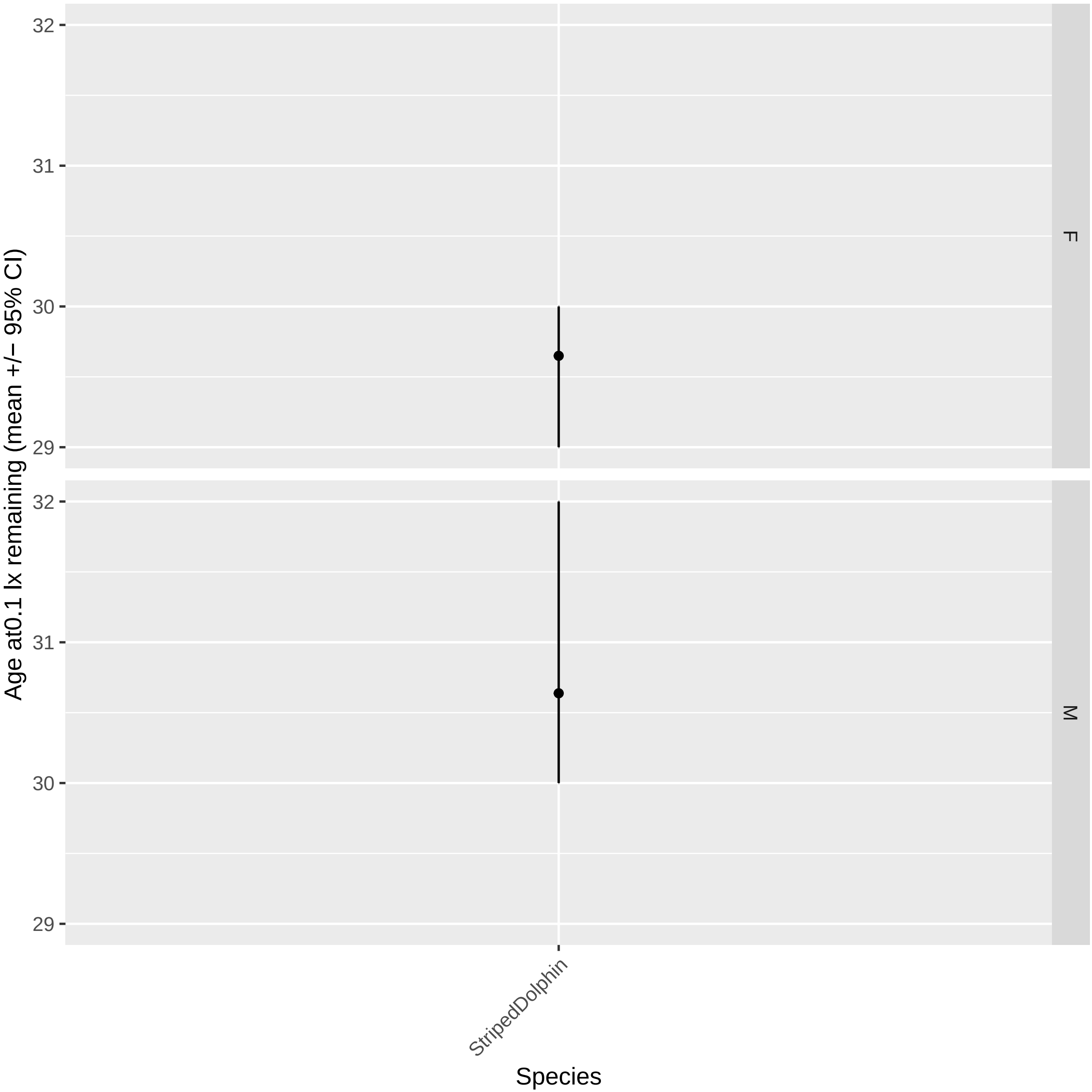

Age at0.1 lx remaining (mean +/− 95% CI)
32
31
30
29
F
M
StripedDolphin
Species

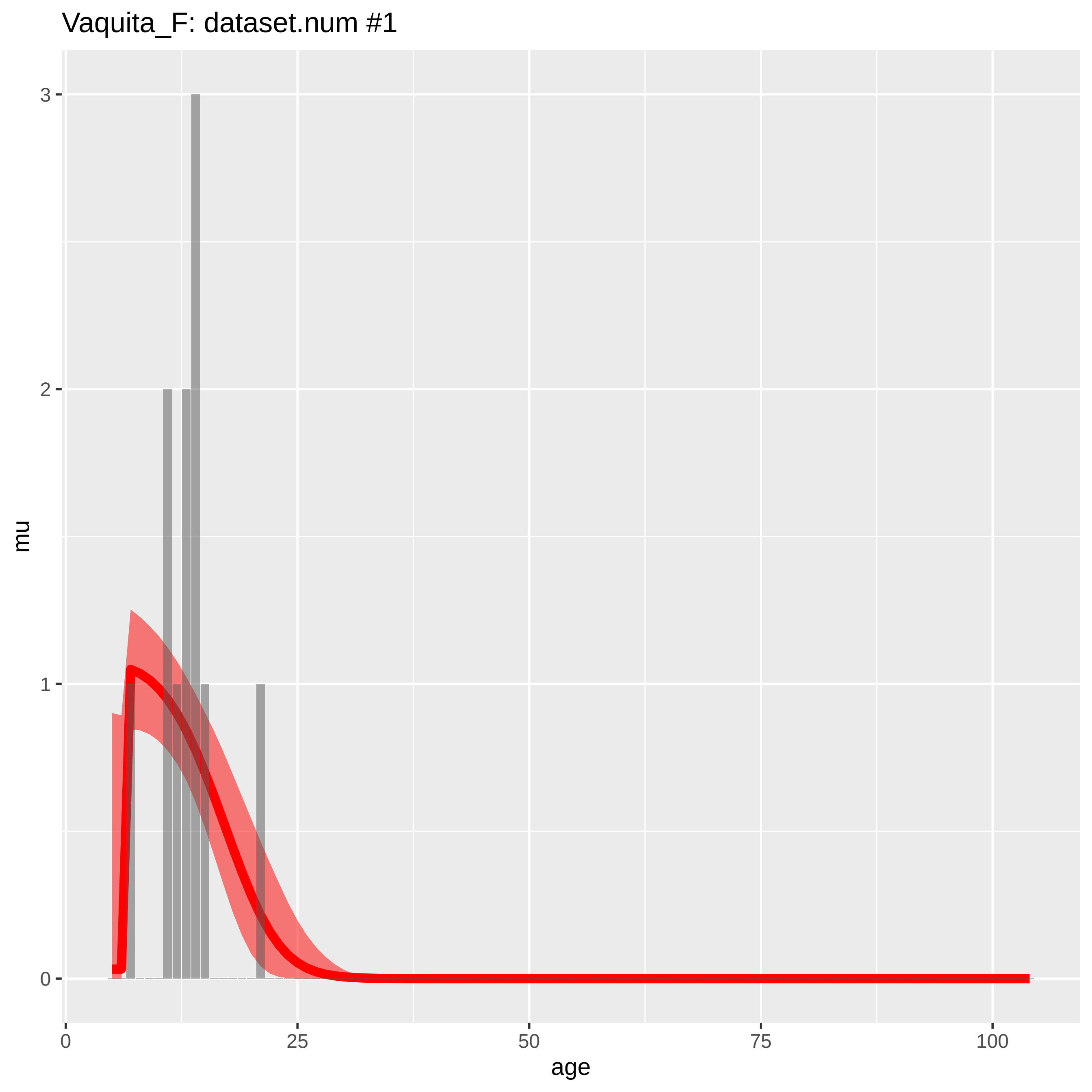

Vaquita_F: dataset.num #1
mu
3
2
1
0
age
0
25
50
75
100

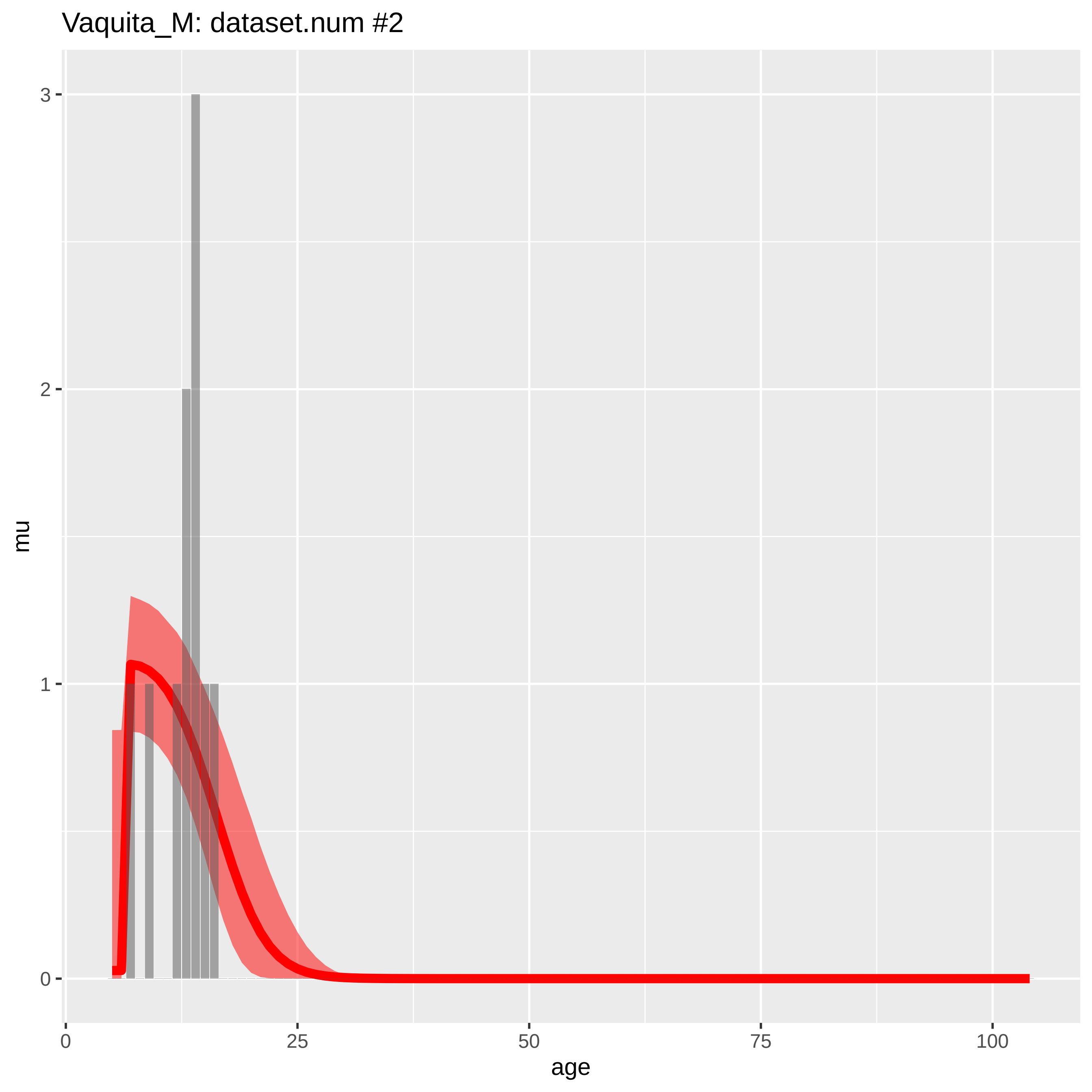
Vaquita_M: dataset.num #2
mu
3
2
1
0
age
0
25
50
75
100

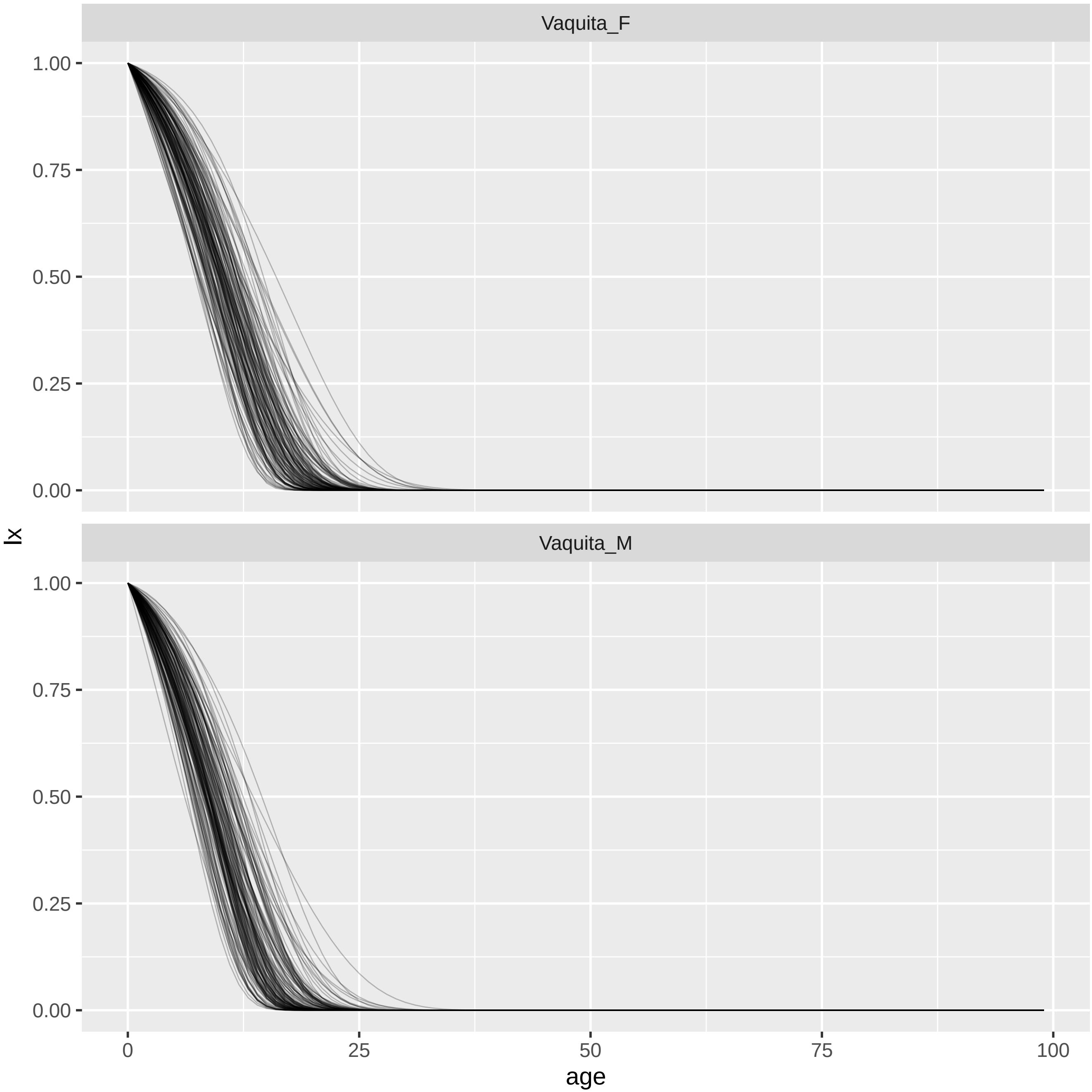

Vaquita_F
Vaquita_M
lx
1.00
0.75
0.50
0.25
0.00
age
0
25
50
75
100

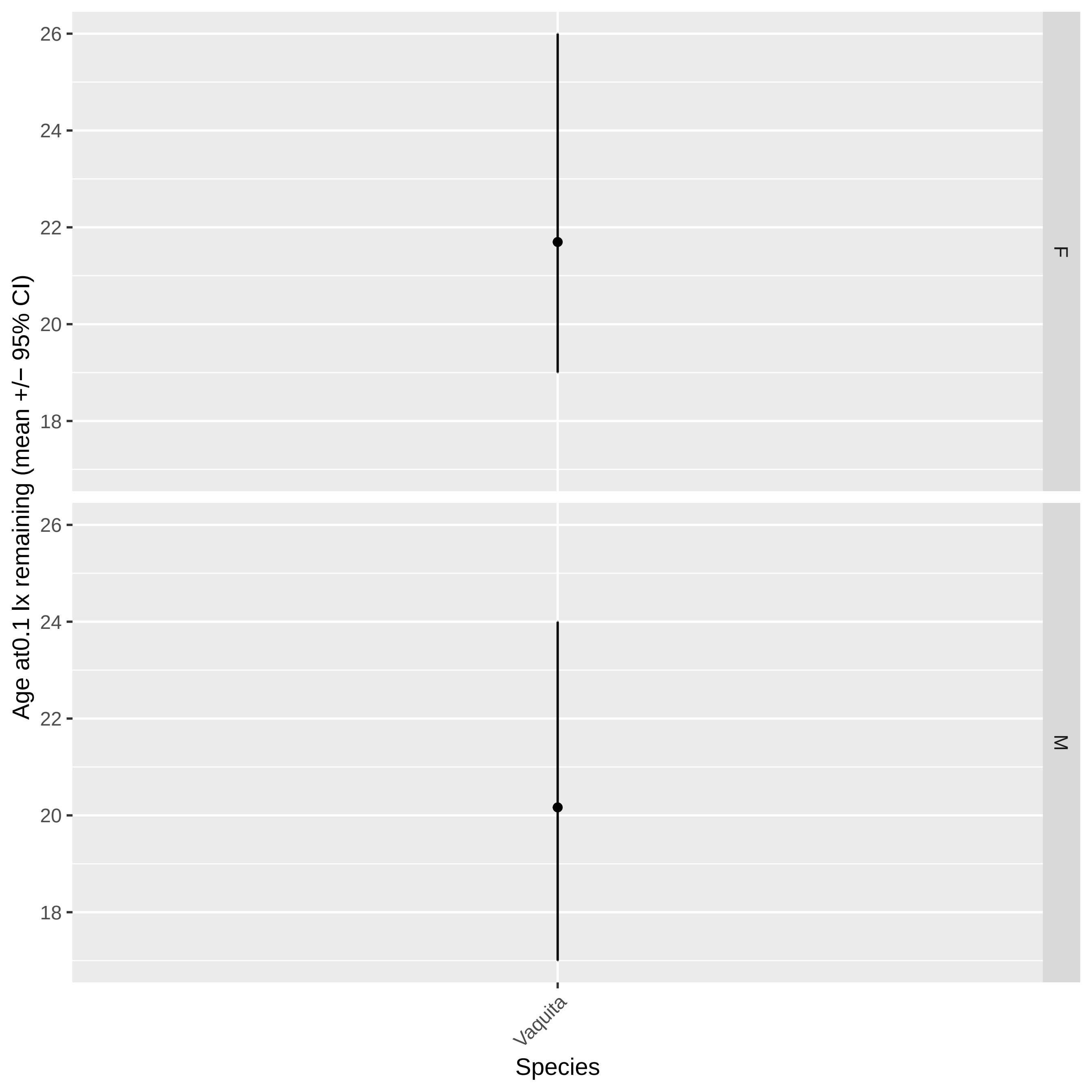

Age at0.1 lx remaining (mean +/− 95% CI)
26
24
22
20
18
F
M
Vaquita
Species

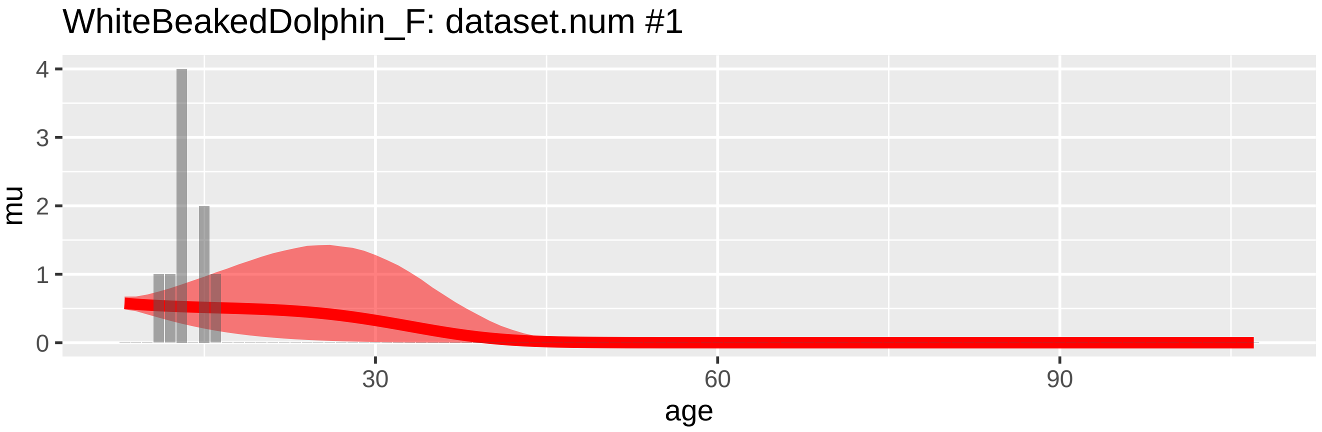
WhiteBeakedDolphin_F: dataset.num #1
mu
4
3
2
1
0
30
60
90
age

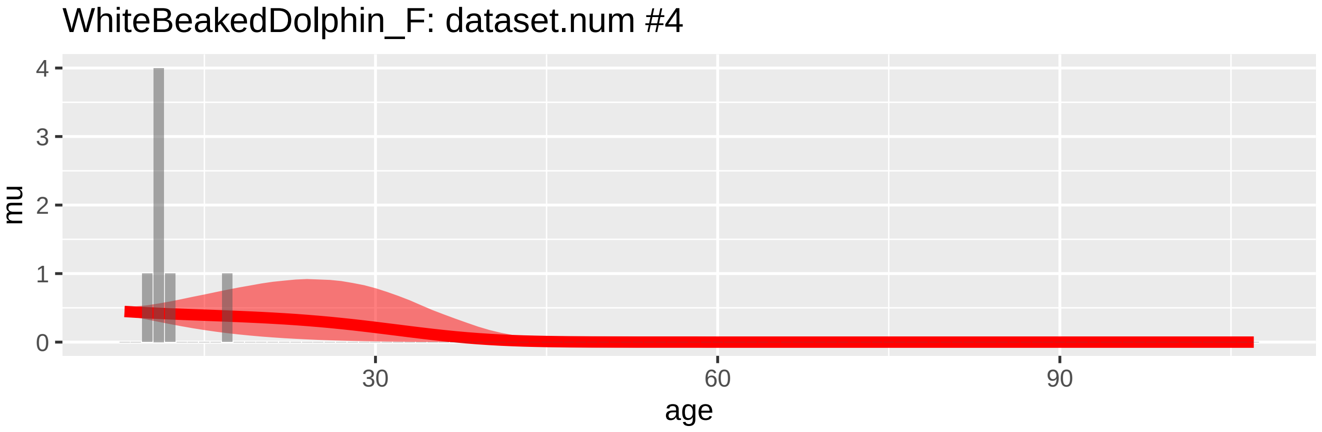
WhiteBeakedDolphin_F: dataset.num #4
mu
4
3
2
1
0
30
60
90
age

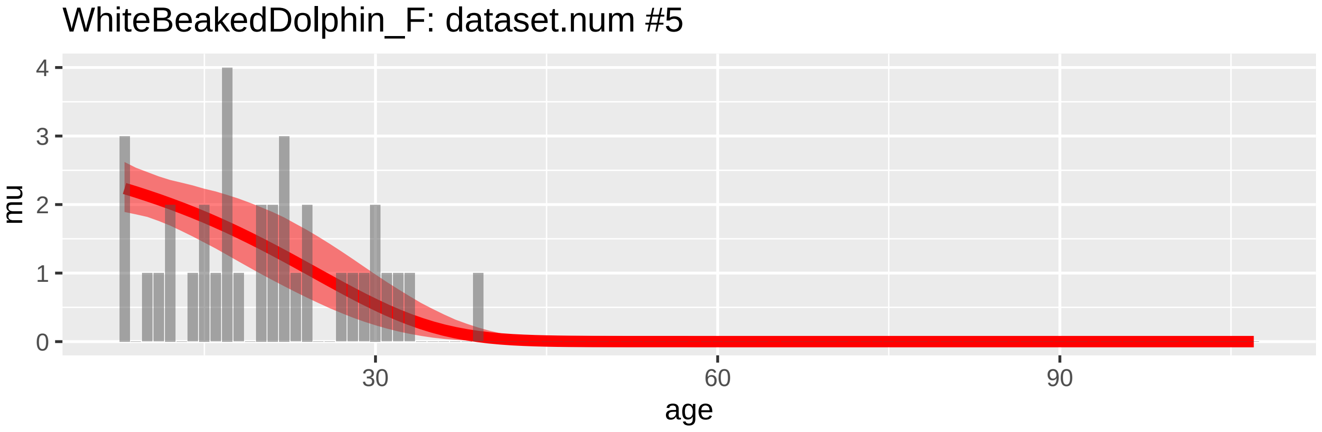
WhiteBeakedDolphin_F: dataset.num #5
mu
4
3
2
1
0
30
60
90
age

WhiteBeakedDolphin_M: dataset.num #2

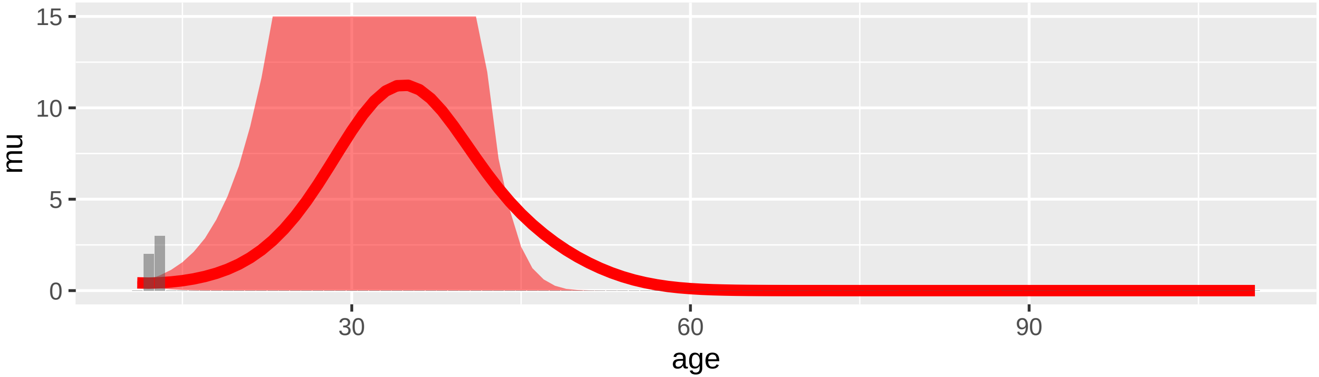


WhiteBeakedDolphin_M: dataset.num #3

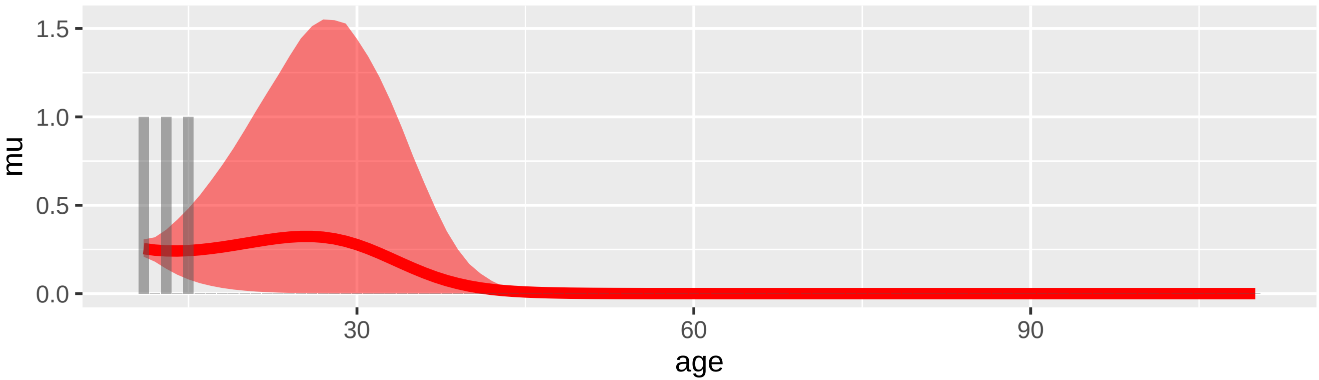


WhiteBeakedDolphin_M: dataset.num #6

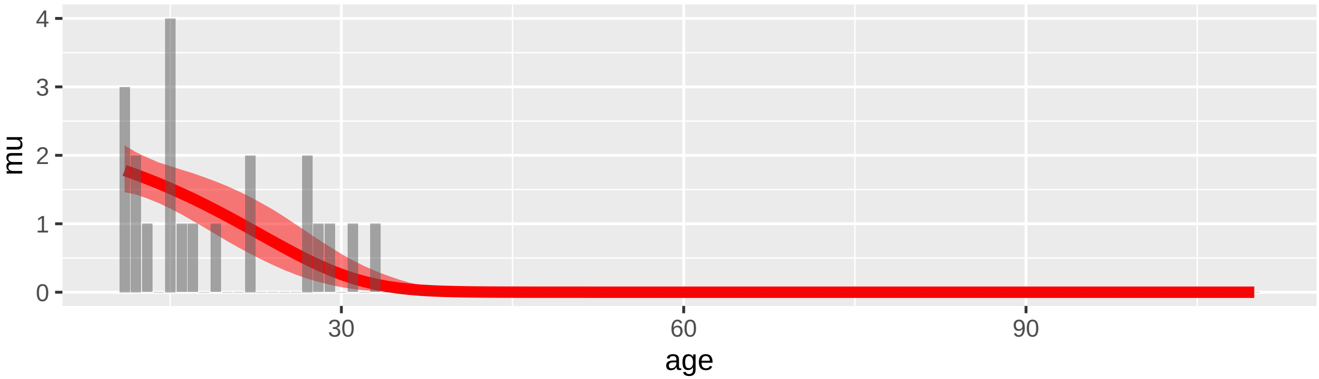

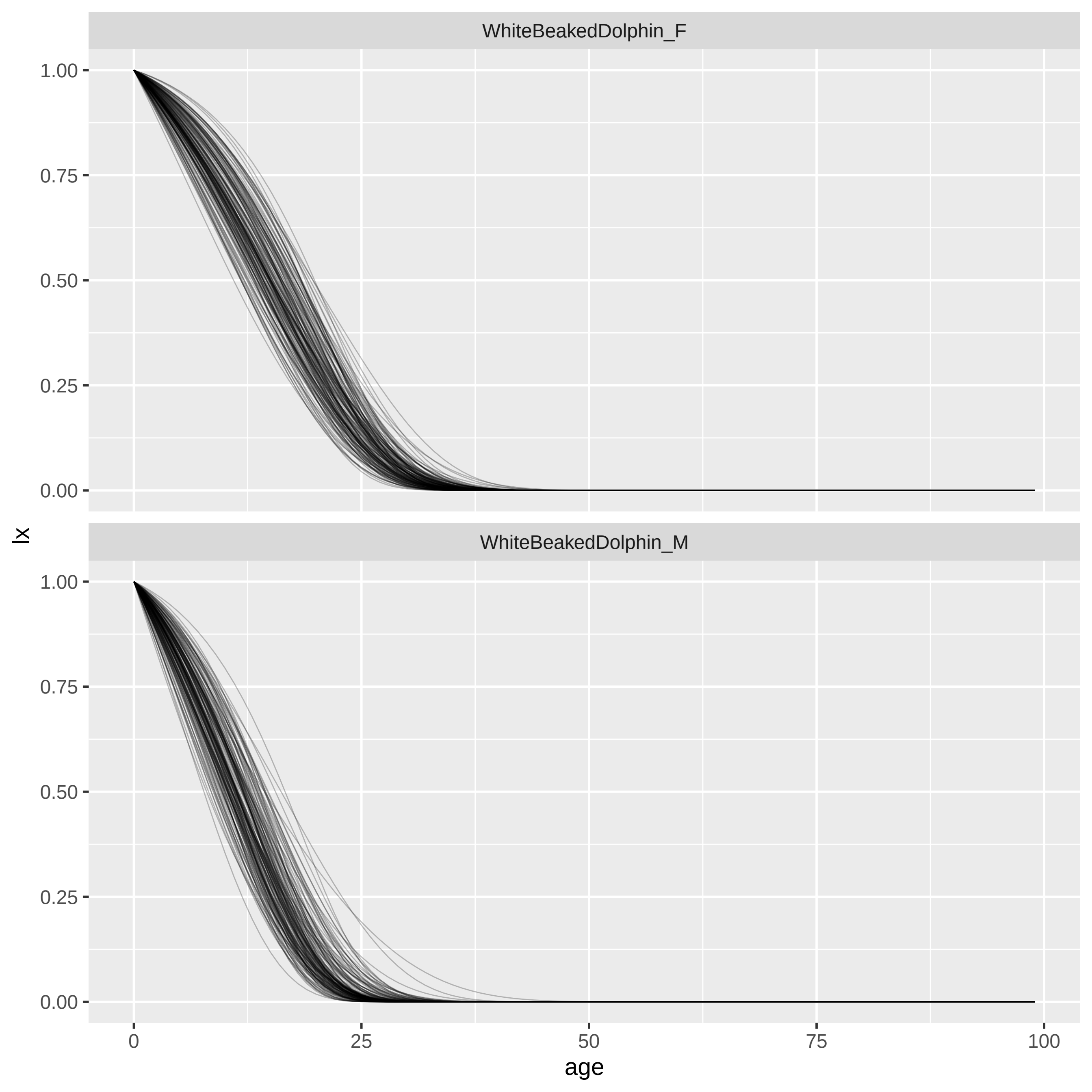

WhiteBeakedDolphin_F
WhiteBeakedDolphin_M
1.00
0.75
0.50
0.25
0.00
lx
0
25
50
75
100
age

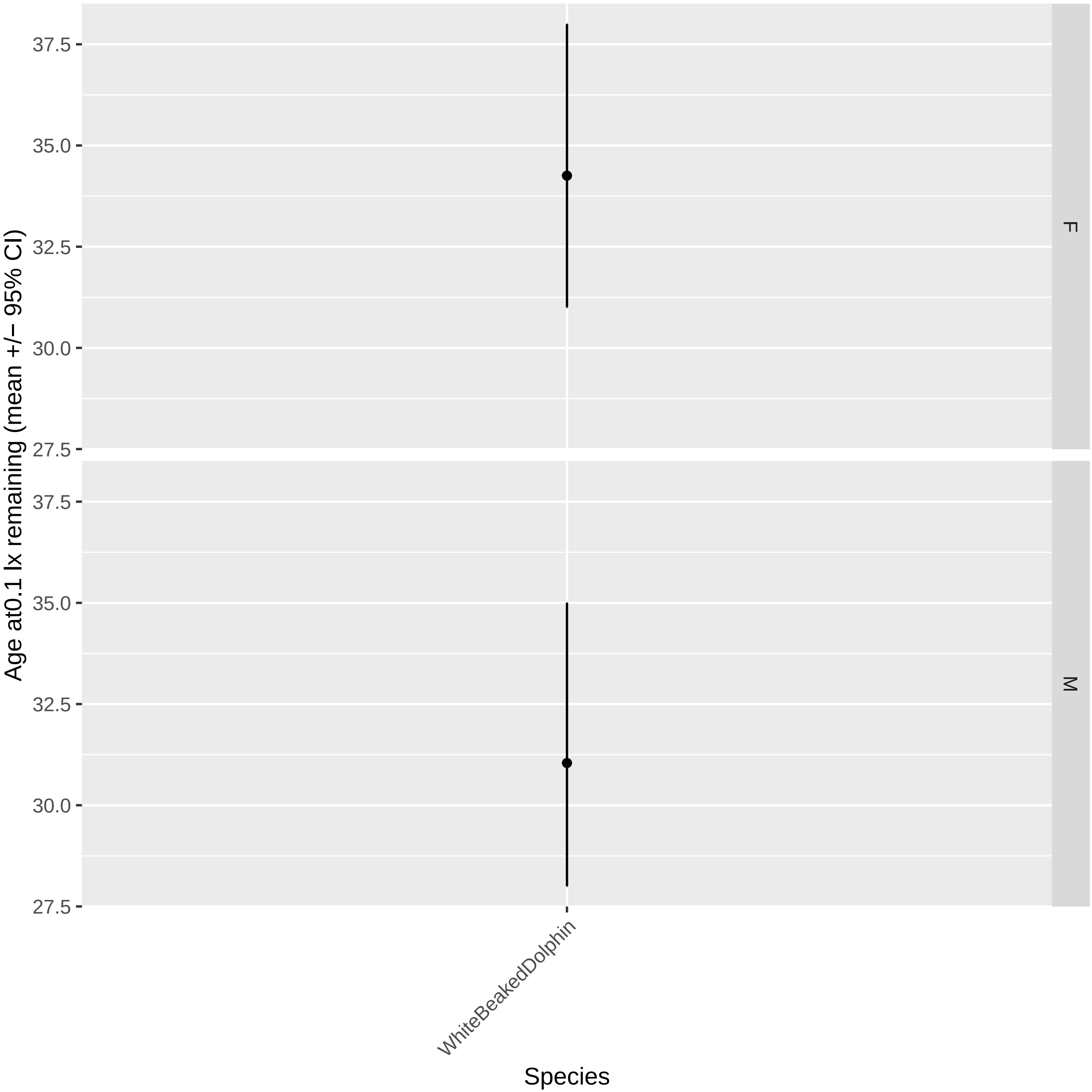

Age at0.1 lx remaining (mean +/− 95% CI)
37.5
35.0
32.5
30.0
27.5
F
M
WhiteBeakedDolphin
Species
