## Supplementary 2 for "Bayesian inference of toothed whale lifespans"

Counting tooth layer groups is subject to error. Particularly in older individuals early deposited tooth layers can become difficult to discern, or teeth can become “full” meaning newer growth layers can be harder to distinguish (Read et al. 2018; Barratclough et al. 2023). Comparing methods of ageing in Common Bottlenose Dolphins for example has highlighted that the error in age estimation from tooth growth layer groups can sometimes be very high in older individuals. Systematically higher uncertainty in older whales has the potential to affect our calculated measures of lifespan.

We explored the influence of an extreme higher age uncertainty in older whales by re-running our models with a new age structure. Our aim was to formulate a model where above a given age there is almost no information about the age of older whales, due to for example a teeth becoming “full”. To do this, for each species-sex we took the age two-thirds of the maximum observed age as our “no information” threshold. For all samples above this threshold we re-age the sample from it’s observed age to the age at a mid-point between the threshold-age, and the maximum age. For example, if the maximum observed age of a species-sex were 12, the threshold age is 8, and therefore all individuals older than 8 are re-aged to 10. We then give that age sample a very high uncertainty by adjusting the standard deviation in age-estimate to be equal to one sixth of total age range (that is: maximum age – threshold /2). The effect of this is that for the older samples the only information we assume is that there is a 50% chance that the sample is between the threshold age and the maximum observed age. In essence, we are assuming that we have very little information about the exact ages of older whales except that they are likely to be older whales. After this re-ageing we then ran the same models and framework as described in the main text. Hereafter, we refer to the models fitted to the model fitted to these restructured datasets as the ‘reaged model(s)’.

In 30 species the 95% credible of predicted ordinary maximum lifespan of the reaged model largely or completely overlaps the 95% credible interval of the original (main text reported) model for both sexes (figure S2.1). In three species – Atlantic white-sided dolphin, baji and vaquita (species names in main text) - the 95% credible interval of the reaged and original models do not overlap or only slightly overlap for both sexes (figure S2.1). And for two further species - Commerson’s dolphin and Fraser’s dolphin - the intervals do not overlap for females (figure S2.1). For all examples where intervals do not overlap the reaged models predict a longer ordinary maximum lifespan than the original models.

The results suggest that in most species our predicted ordinary maximum lifespan are robust to potential difficulties in ageing older individuals. The examples where the reaged model do differ from our original model are all species with relatively low sample sizes (figure S2.1; table 2; table S2.1)- though in several species with similarly low sample size the estimates of lifespan remains unchanged (table 2).

*S2 references*

Barratclough A, McFee WE, Stolen M, et al (2023) How to estimate age of old bottlenose dolphins (Tursiops truncatus); by tooth or pectoral flipper? Frontiers in Marine Science 10:

Read FL, Hohn AA, Lockyer CH (2018) A review of age estimation methods in marine mammals with special reference to monodontids. NAMMCO Scientific Publications 8:1–67. https://doi.org/10.7557/3.4474

Table S2.1. Sample size for each species-sex example where the reaged models differ from the original models (see table 2 for more information and other species). Empty cells are where a given species-sex pair does not differ between the two models or where that species-sex is not included in our analysis (see main text).

| Species | Female sample size | Male sample size |
|---|---|---|
| **Atlantic white-sided dolphin** | 45 | 32 |
| **Baiji** | 17 | 11 |
| **Commerson's dolphin** | 38 | |
| **Fraser's dolphin** | 39 | |
| **Vaquita** | 11 | 10 |

Figure S2.1. Comparison of ordinary maximum lifespan estimates for the original (red) and reaged (blue) models for each species-sex. Each figure shows one species, with sexes on the x axis. In each plot, the point shows the posterior mean of the distribution of ordinary maximum lifespans derived from the model, and the lines show the 95% credible interval. For clarity each figure is depicted on it's own page.

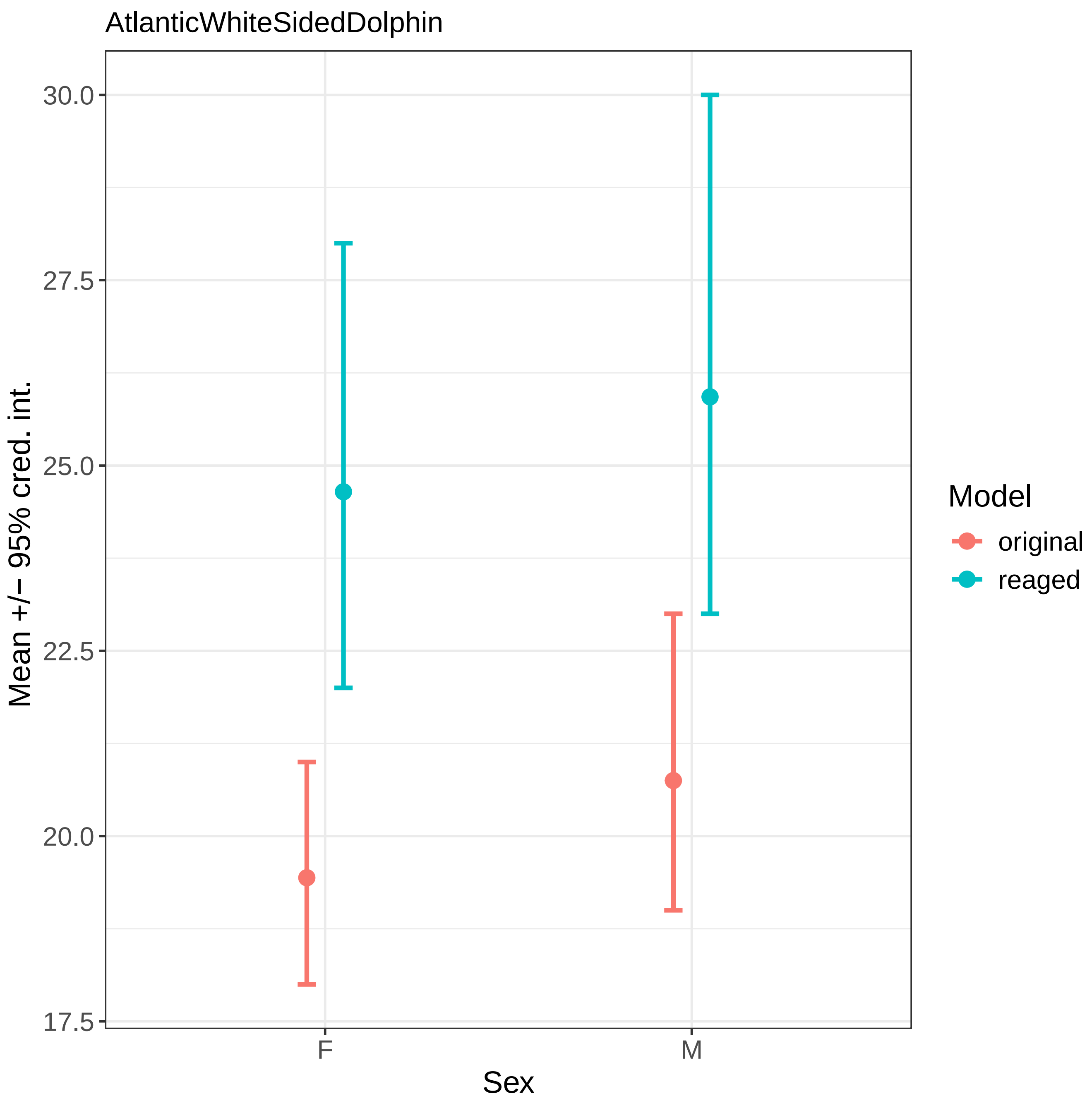
AtlanticWhiteSidedDolphin
30.0
27.5
25.0
22.5
20.0
17.5
Mean +/− 95% cred. int.
F
M
Sex
Model
original
reaged

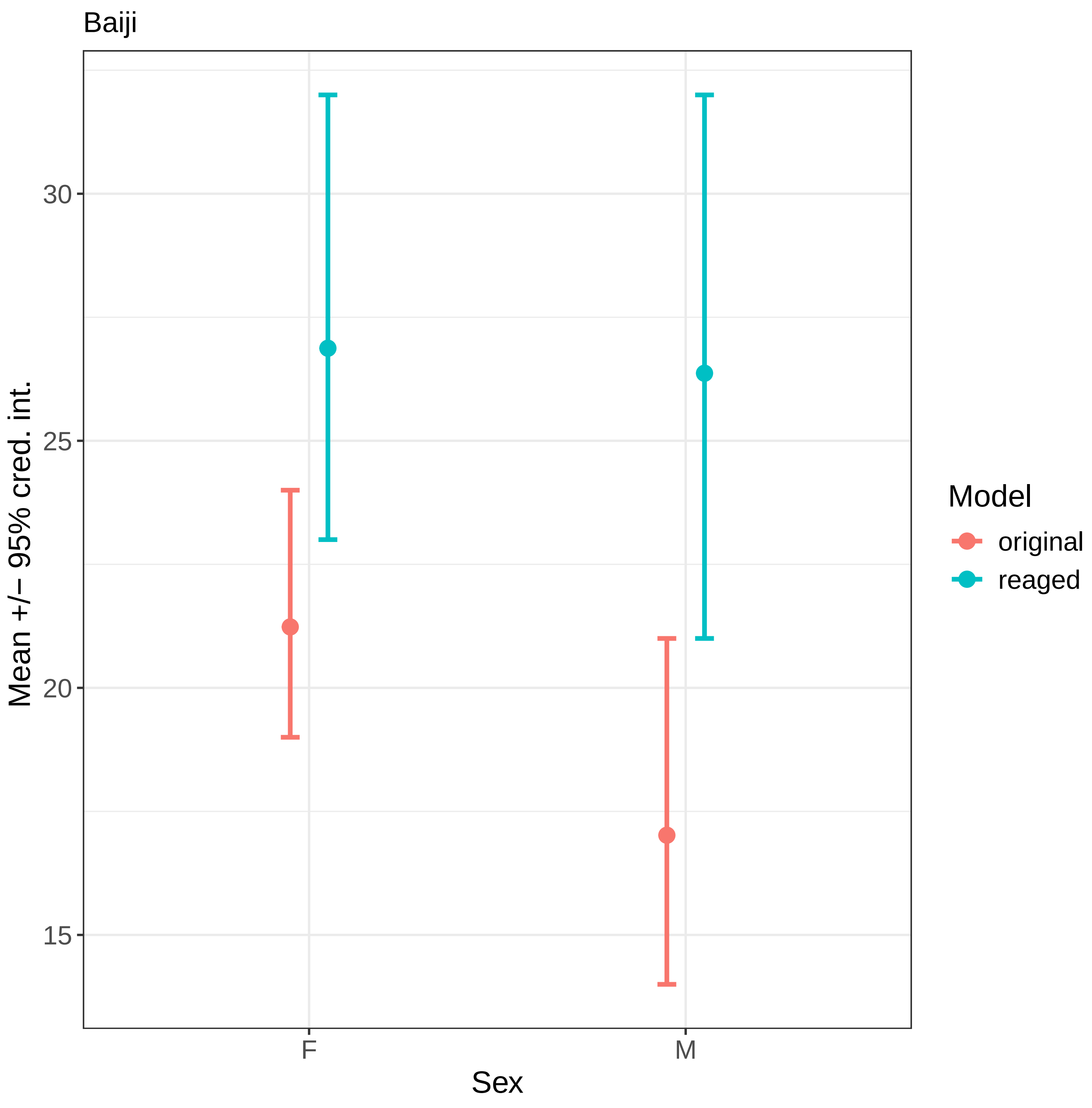
Baiji
Mean +/− 95% cred. int.
30
25
20
15
F
M
Sex
Model
original
reaged

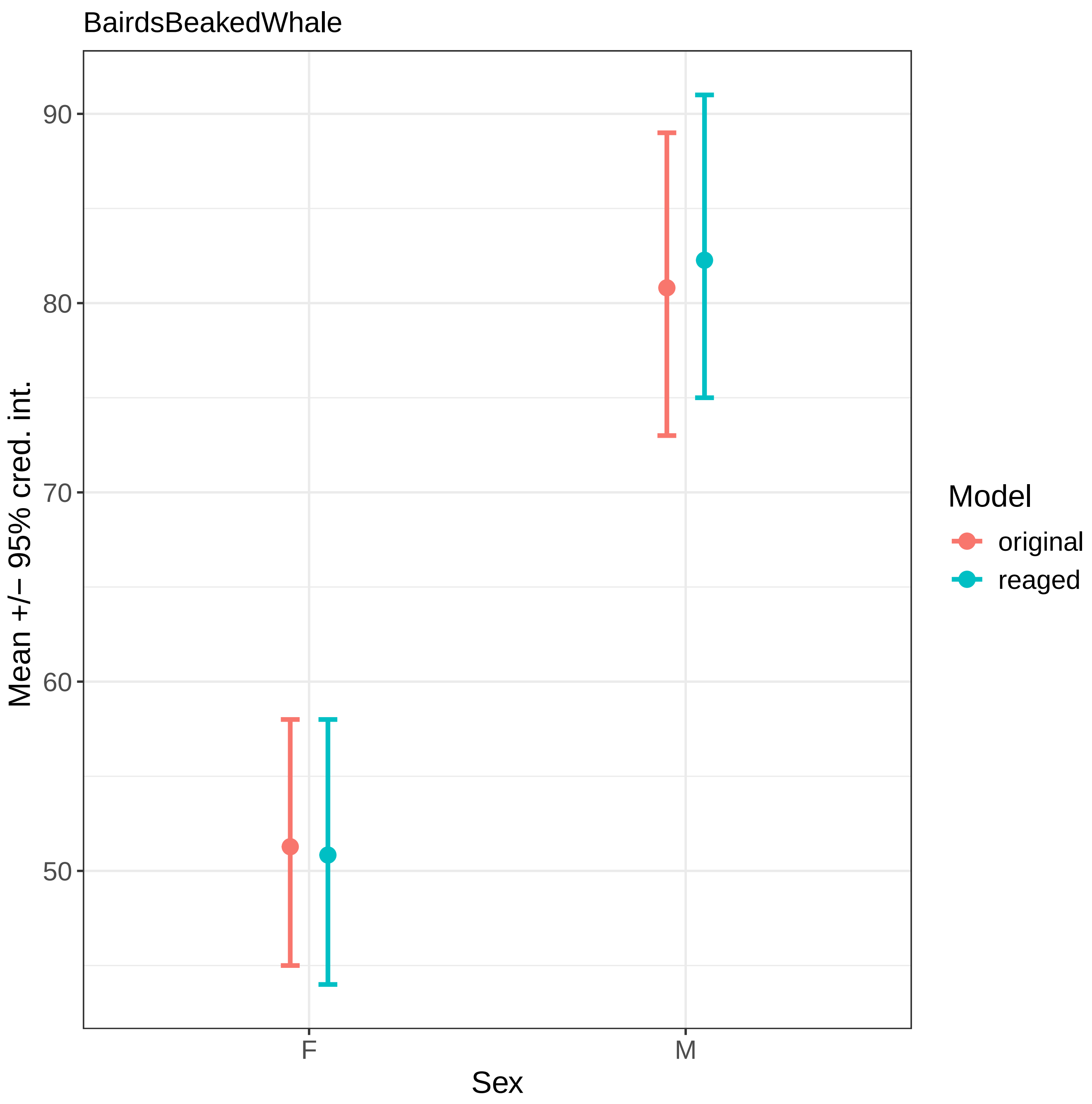
BairdsBeakedWhale
90
80
70
60
50
Mean +/− 95% cred. int.
F
M
Sex
Model
original
reaged

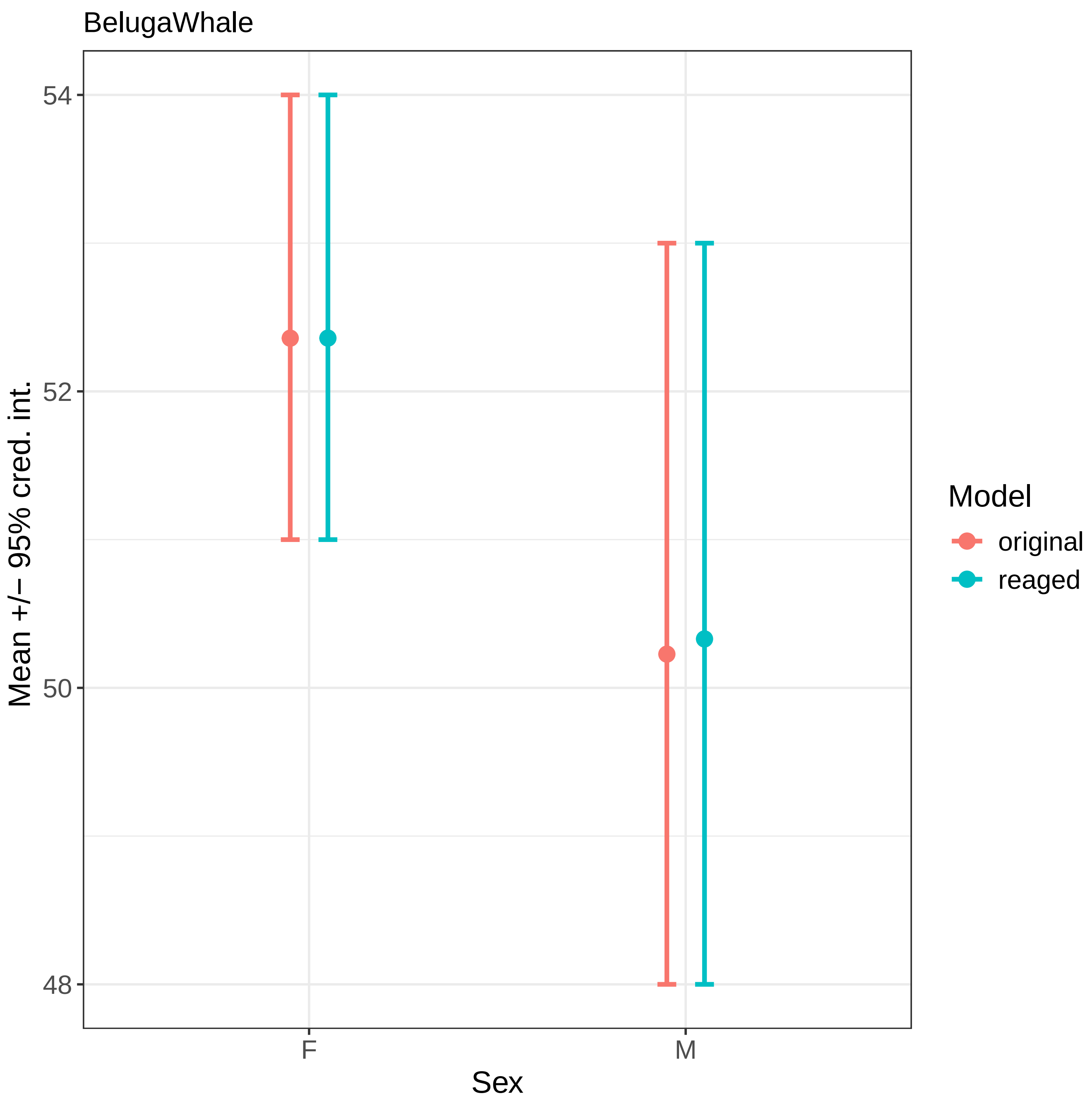
BelugaWhale
54
52
50
48
Mean +/− 95% cred. int.
F
M
Sex
Model
original
reaged

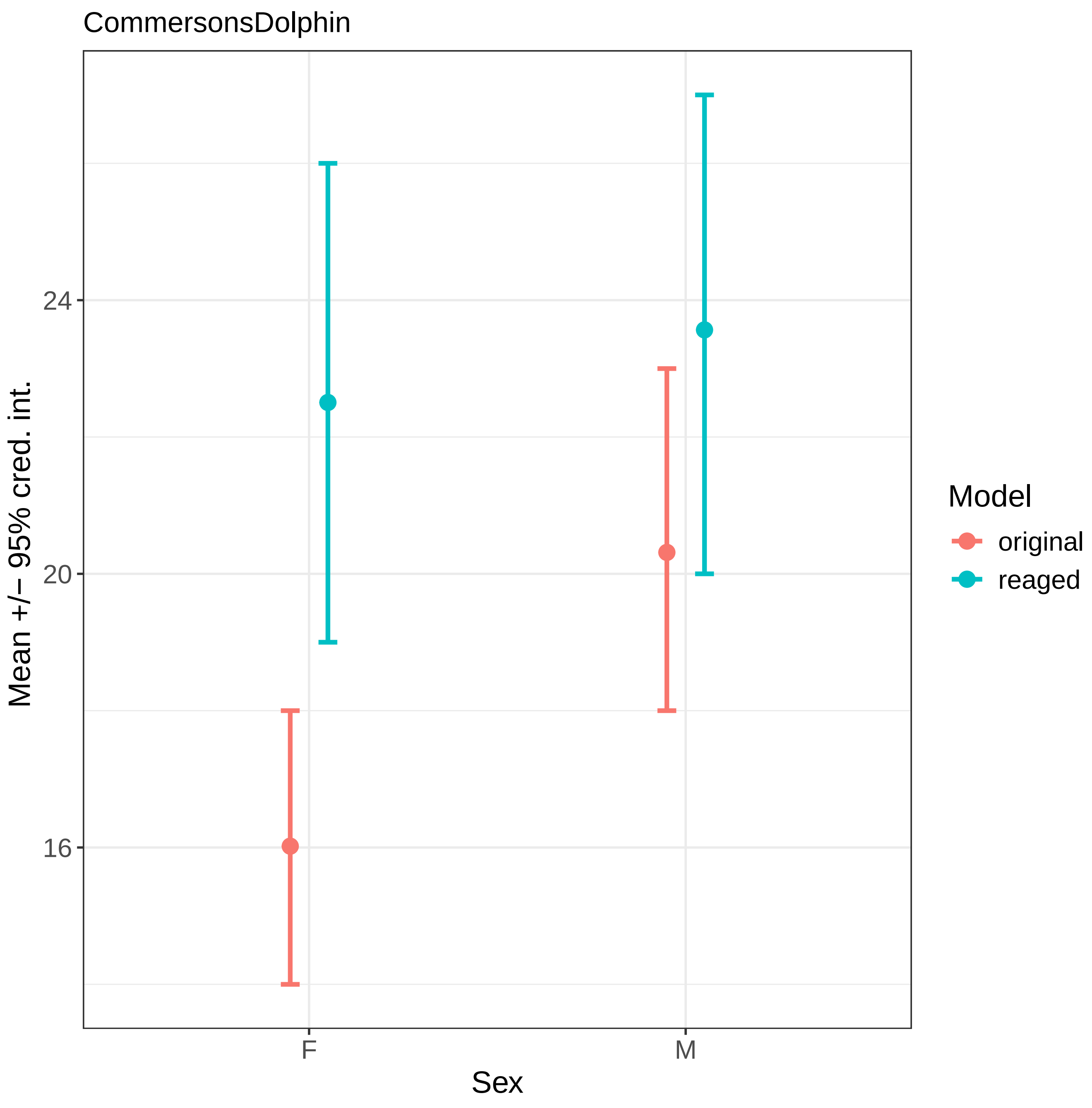
CommersonsDolphin
Mean +/− 95% cred. int.
24
20
16
F
M
Sex
Model
original
reaged

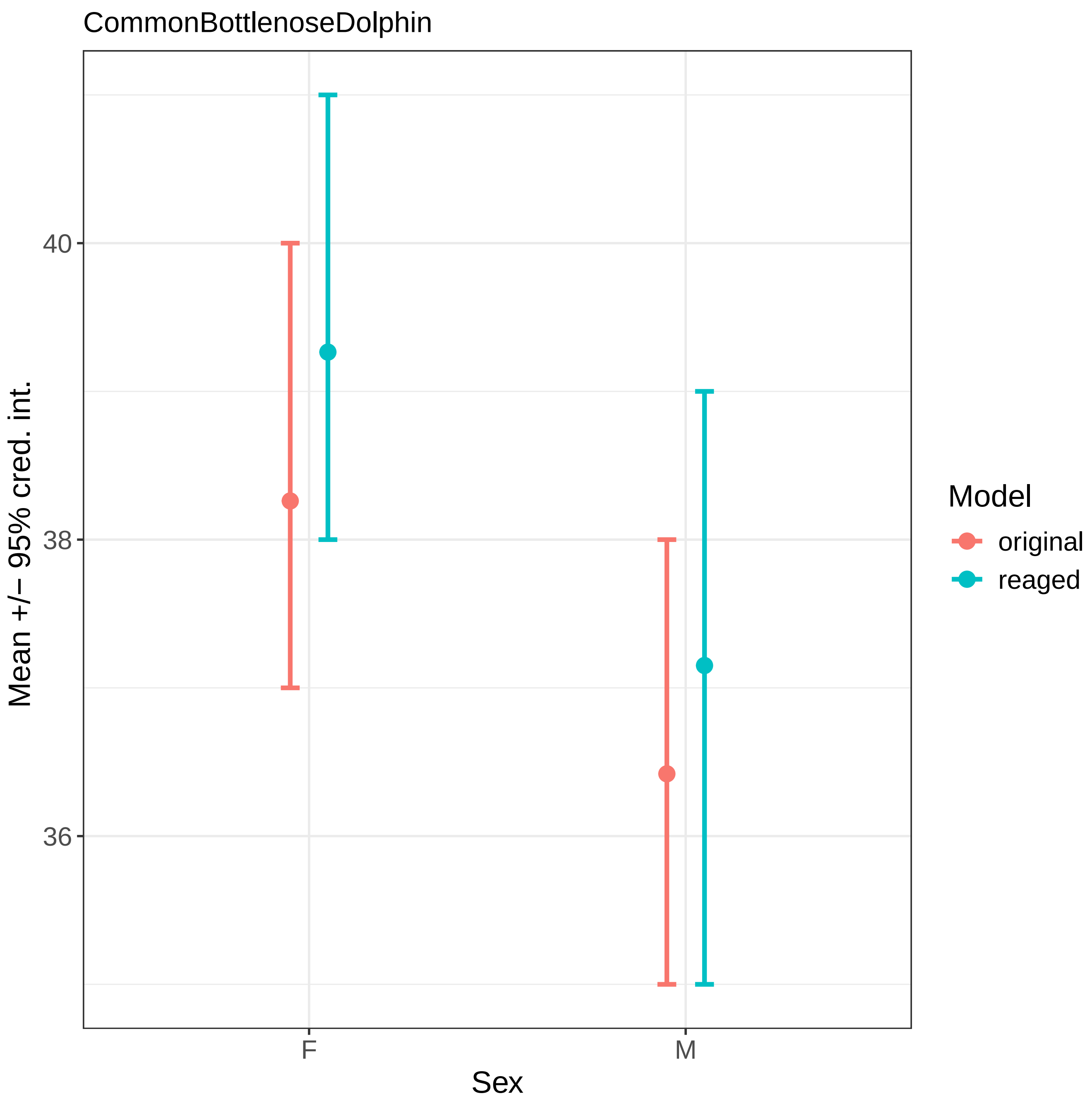
CommonBottlenoseDolphin
40
38
36
Mean +/− 95% cred. int.
F
M
Sex
Model
original
reaged

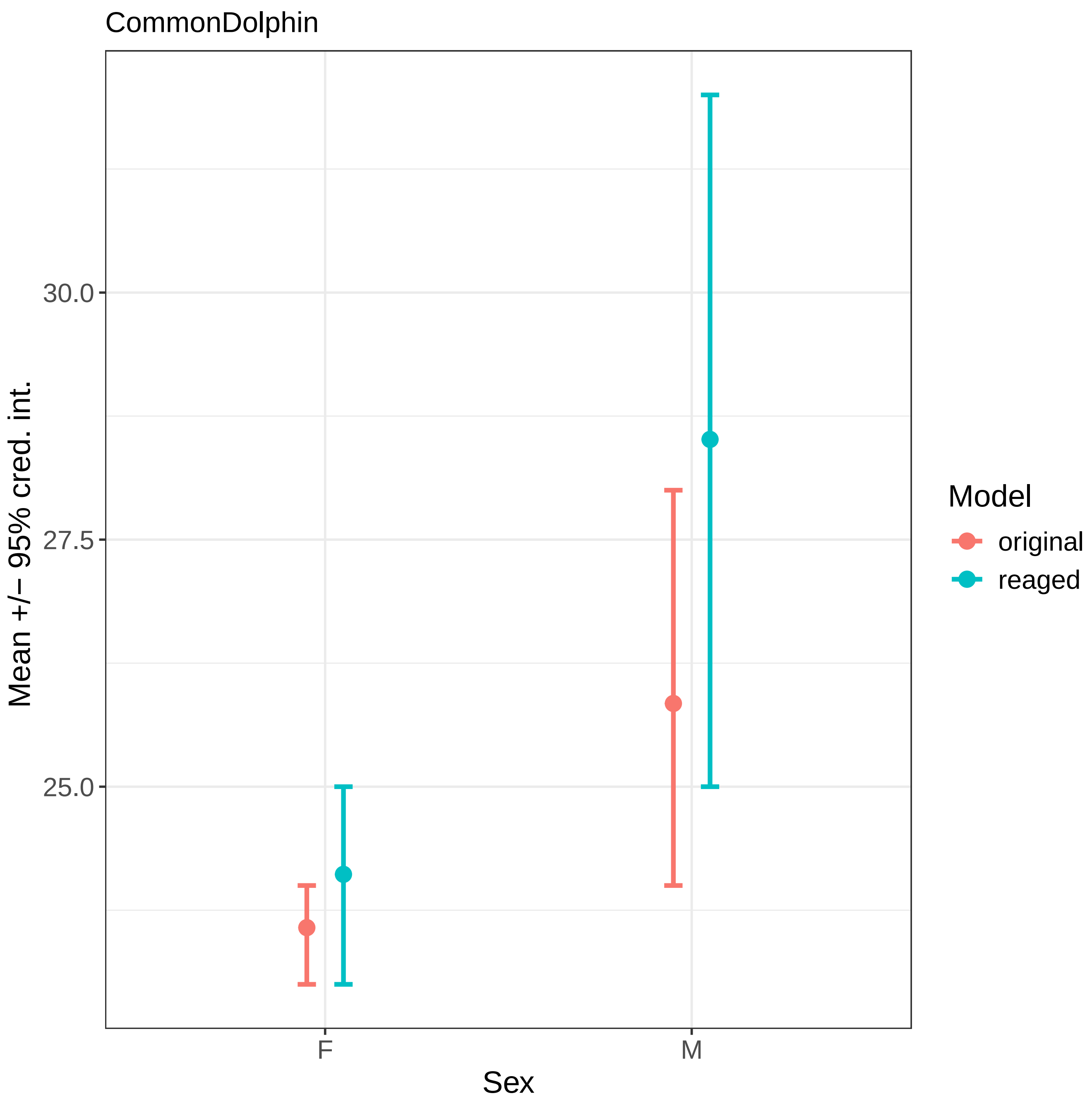
CommonDolphin
30.0
27.5
25.0
Mean +/− 95% cred. int.
F
M
Sex
Model
original
reaged

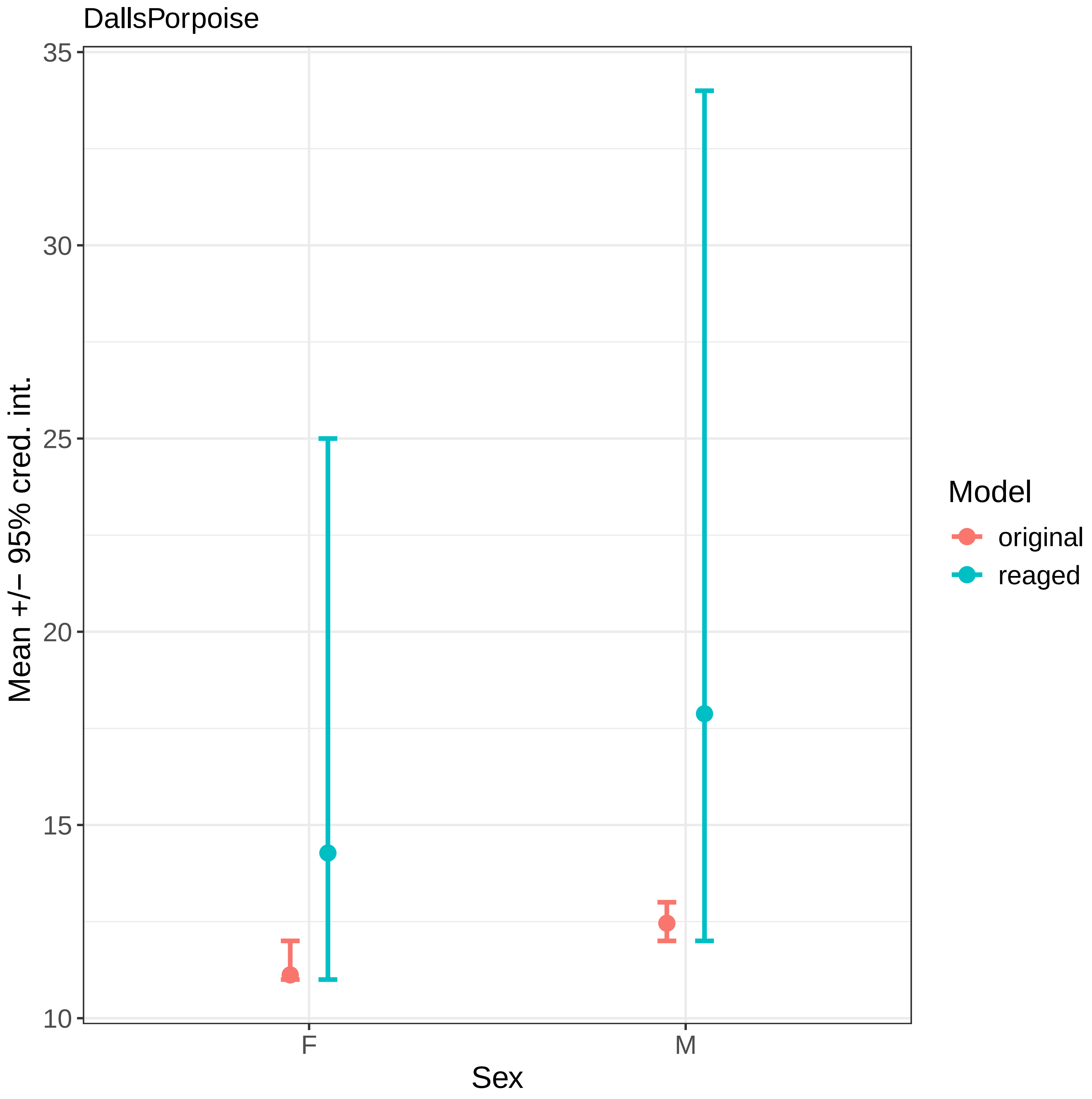

DallsPorpoise
35
30
25
20
15
10
Mean +/− 95% cred. int.
F
M
Sex
Model
original
reaged

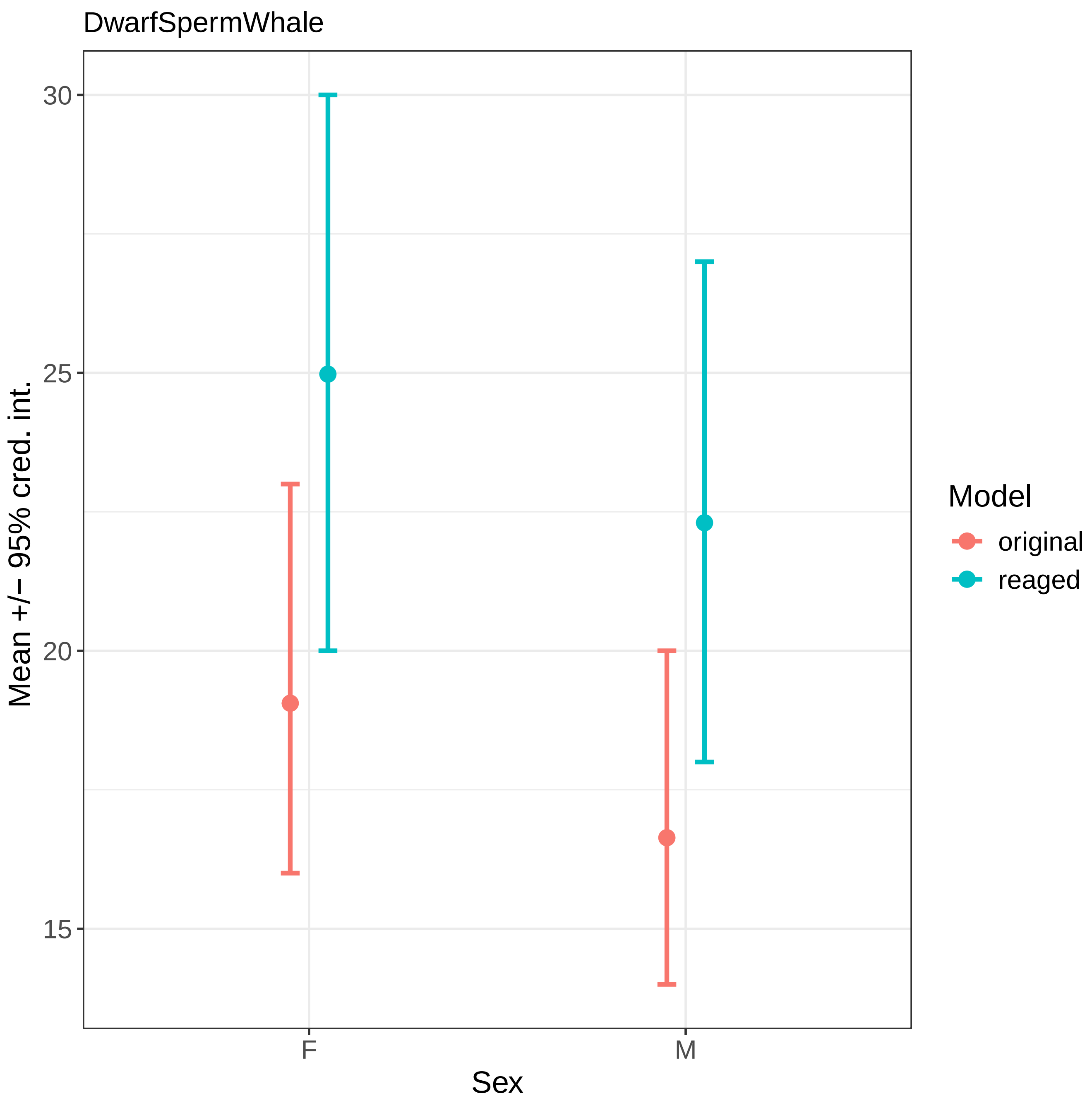

DwarfSpermWhale
Mean +/− 95% cred. int.
30
25
20
15
F
M
Sex
Model
original
reaged

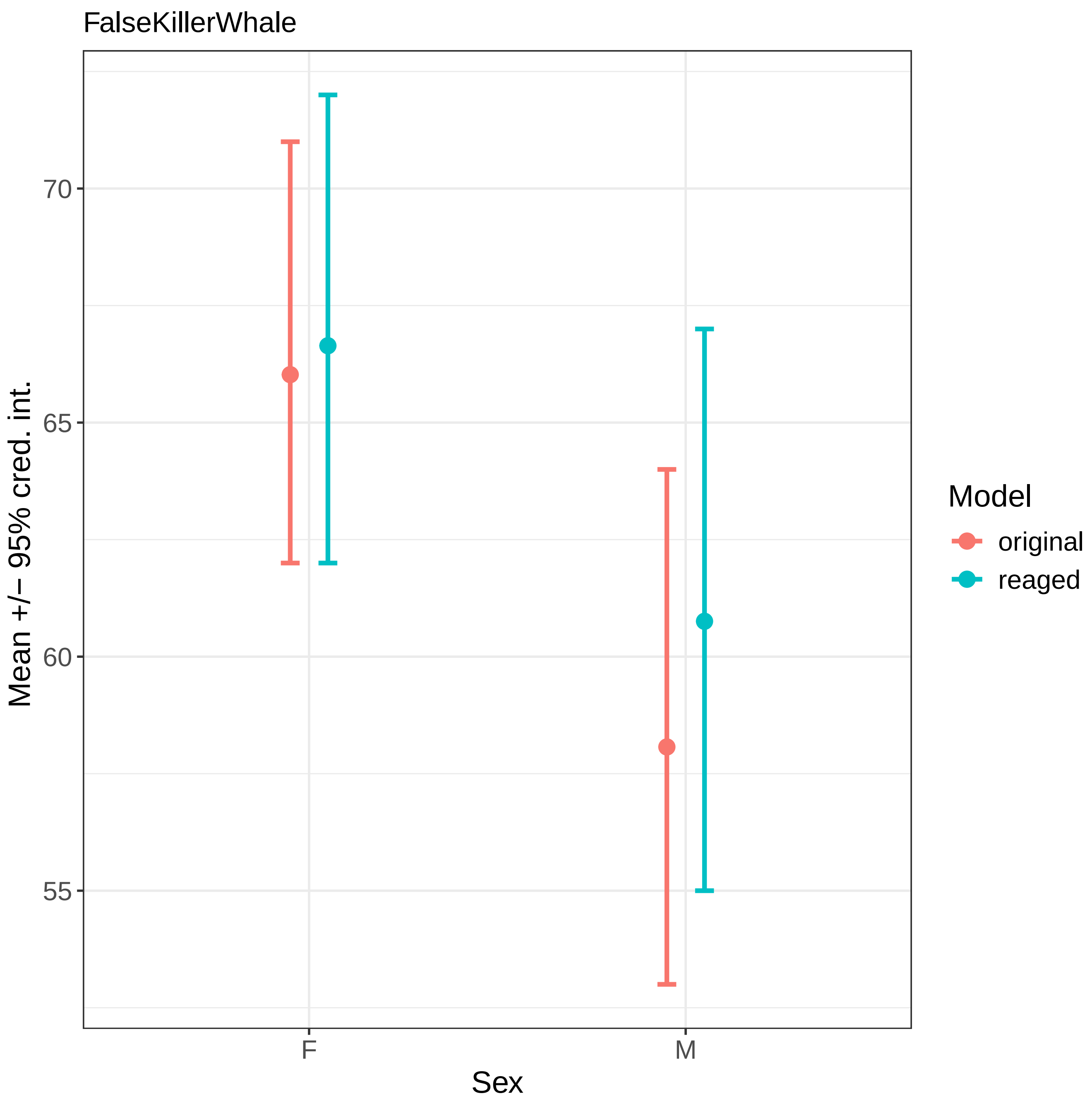
FalseKillerWhale
70
65
60
55
Mean +/− 95% cred. int.
F
M
Sex
Model
original
reaged

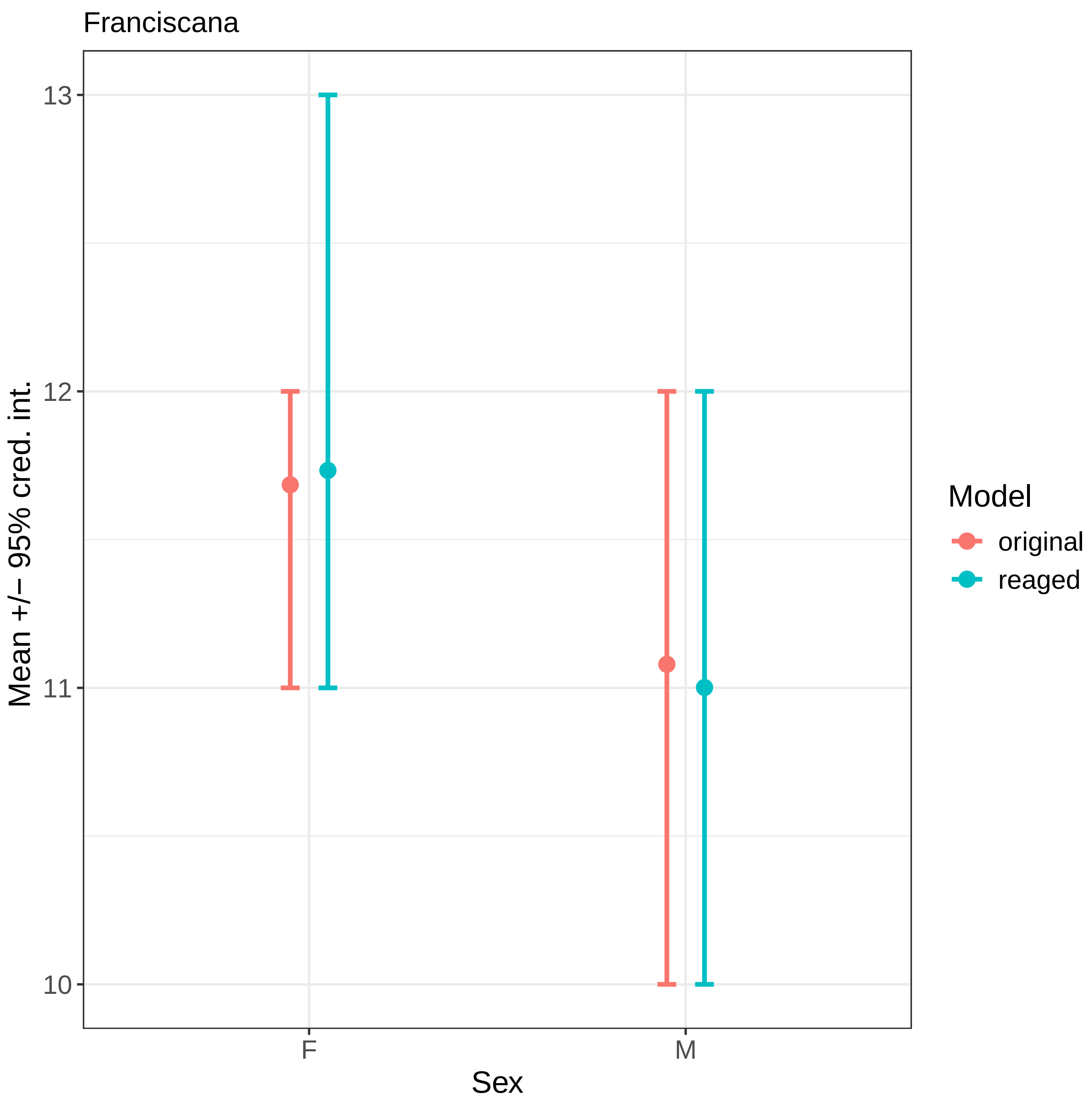

Franciscana
Mean +/− 95% cred. int.
13
12
11
10
F
M
Sex
Model
original
reaged

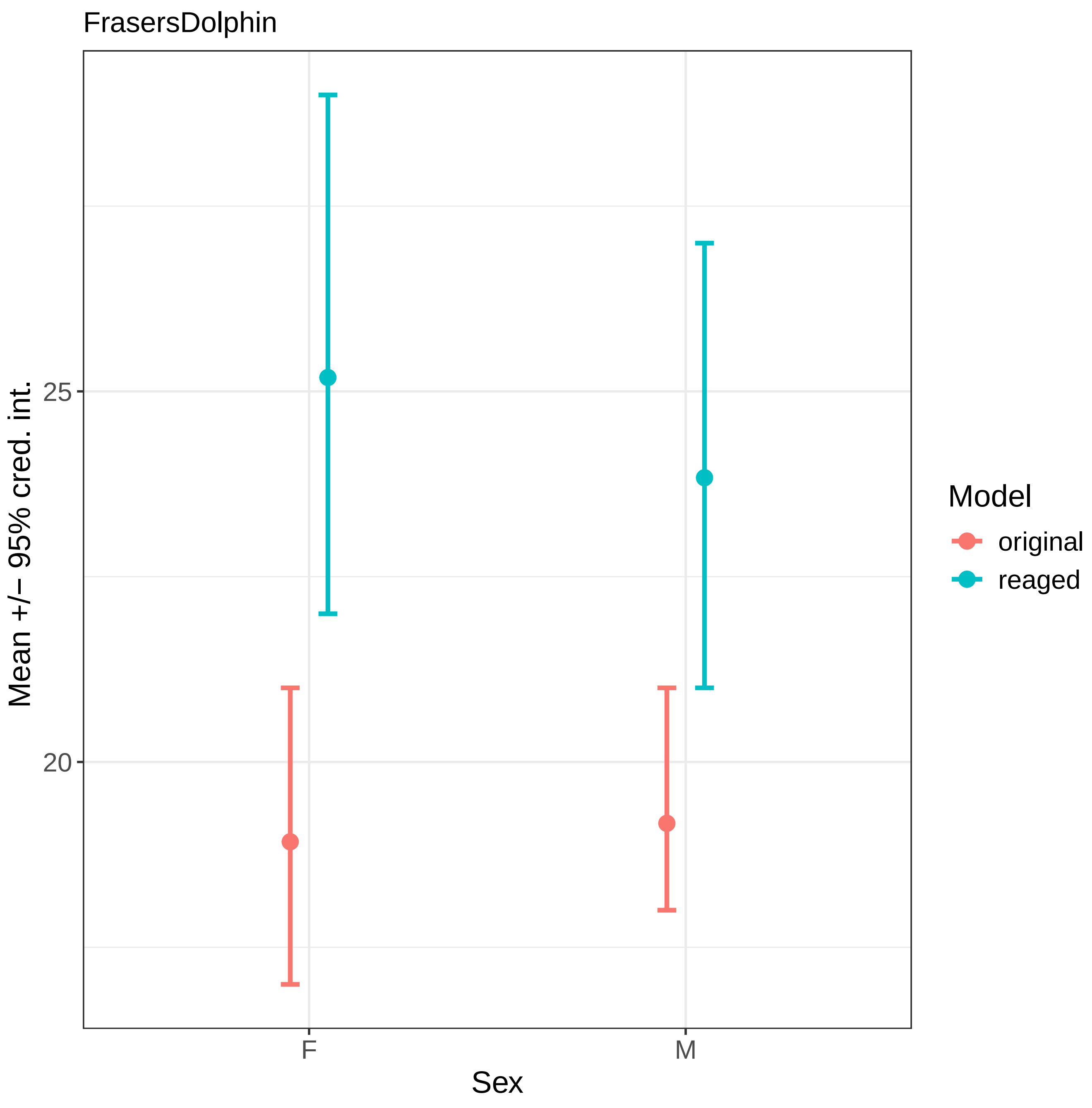

FrasersDolphin
Mean +/− 95% cred. int.
25
20
F
M
Sex
Model
original
reaged

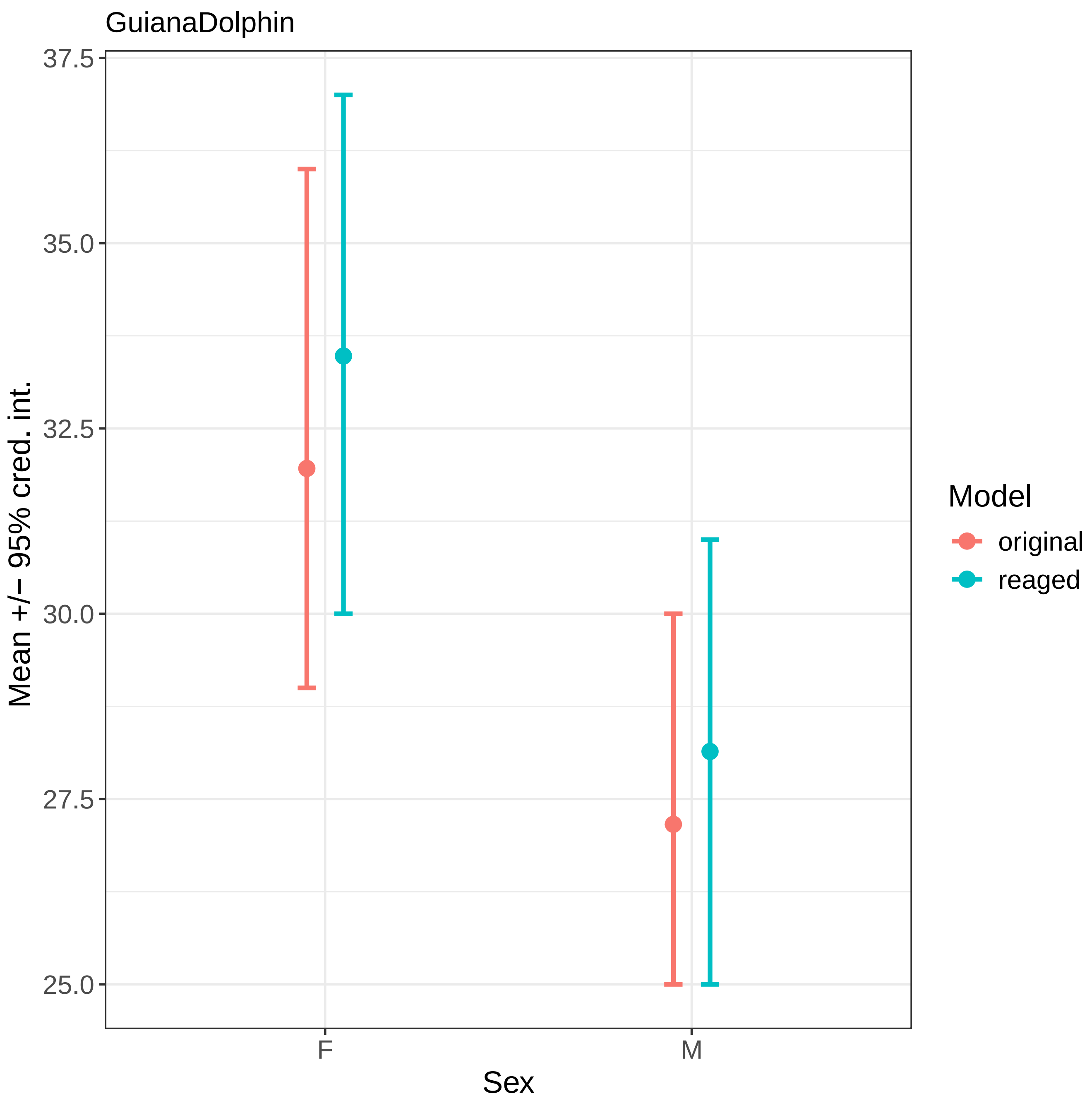

GuianaDolphin
37.5
35.0
32.5
30.0
27.5
25.0
Mean +/− 95% cred. int.
F
M
Sex
Model
original
reaged

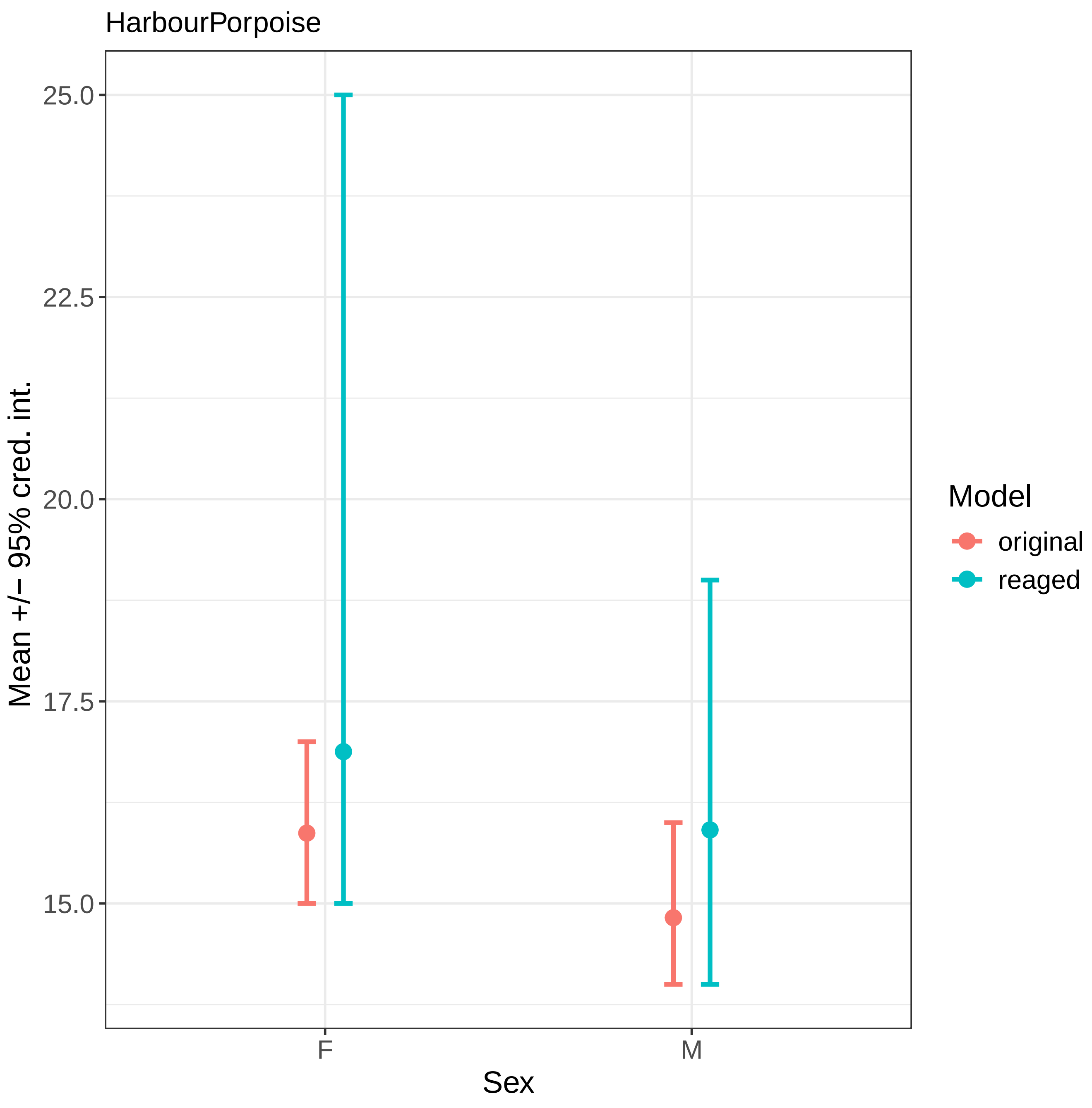
HarbourPorpoise
Mean +/− 95% cred. int.
25.0
22.5
20.0
17.5
15.0
F
M
Sex
Model
original
reaged

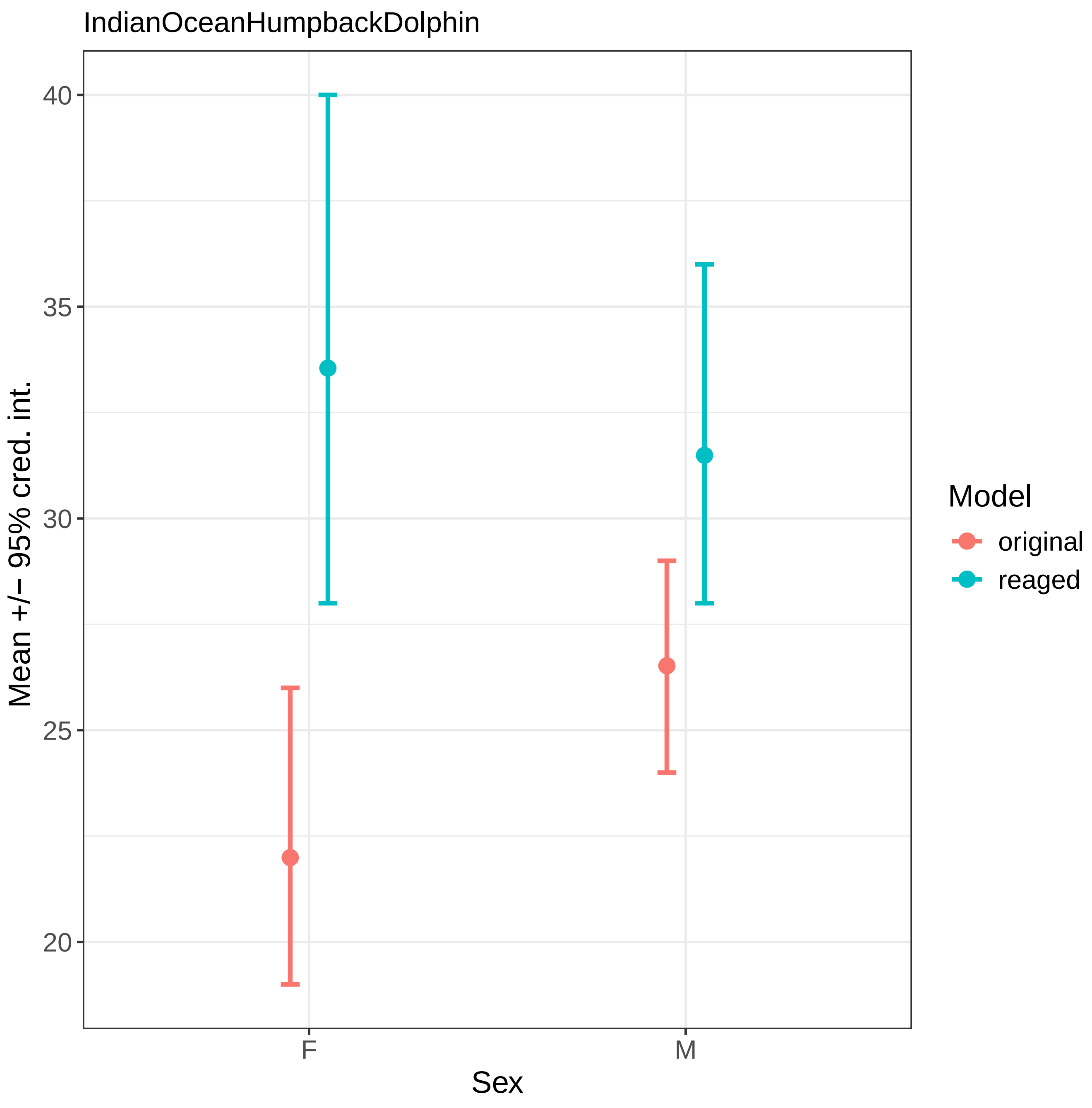

IndianOceanHumpbackDolphin
Mean +/− 95% cred. int.
40
35
30
25
20
F
M
Sex
Model
original
reaged

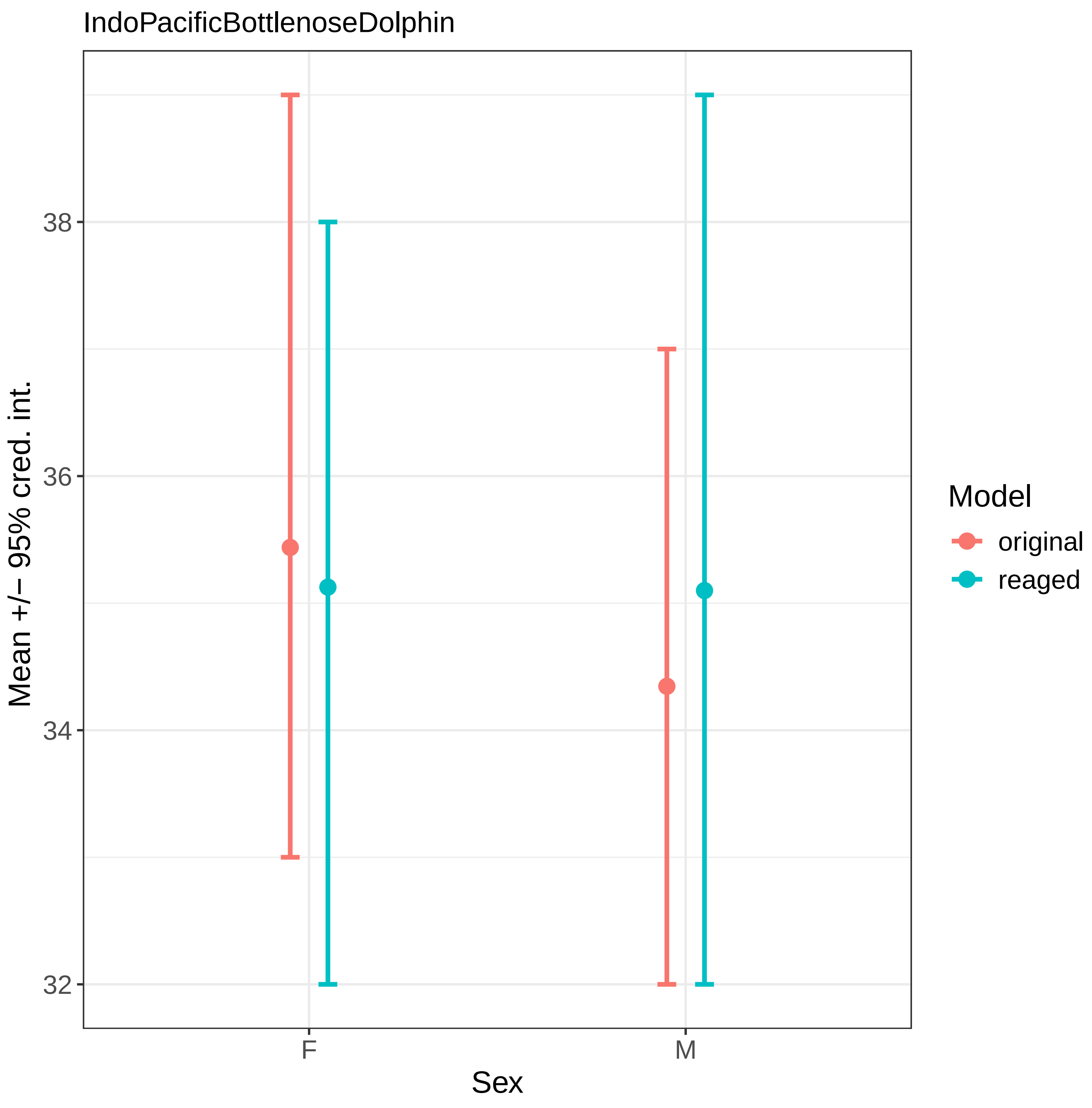
IndoPacificBottlenoseDolphin
Mean +/− 95% cred. int.
38
36
34
32
F
M
Sex
Model
original
reaged

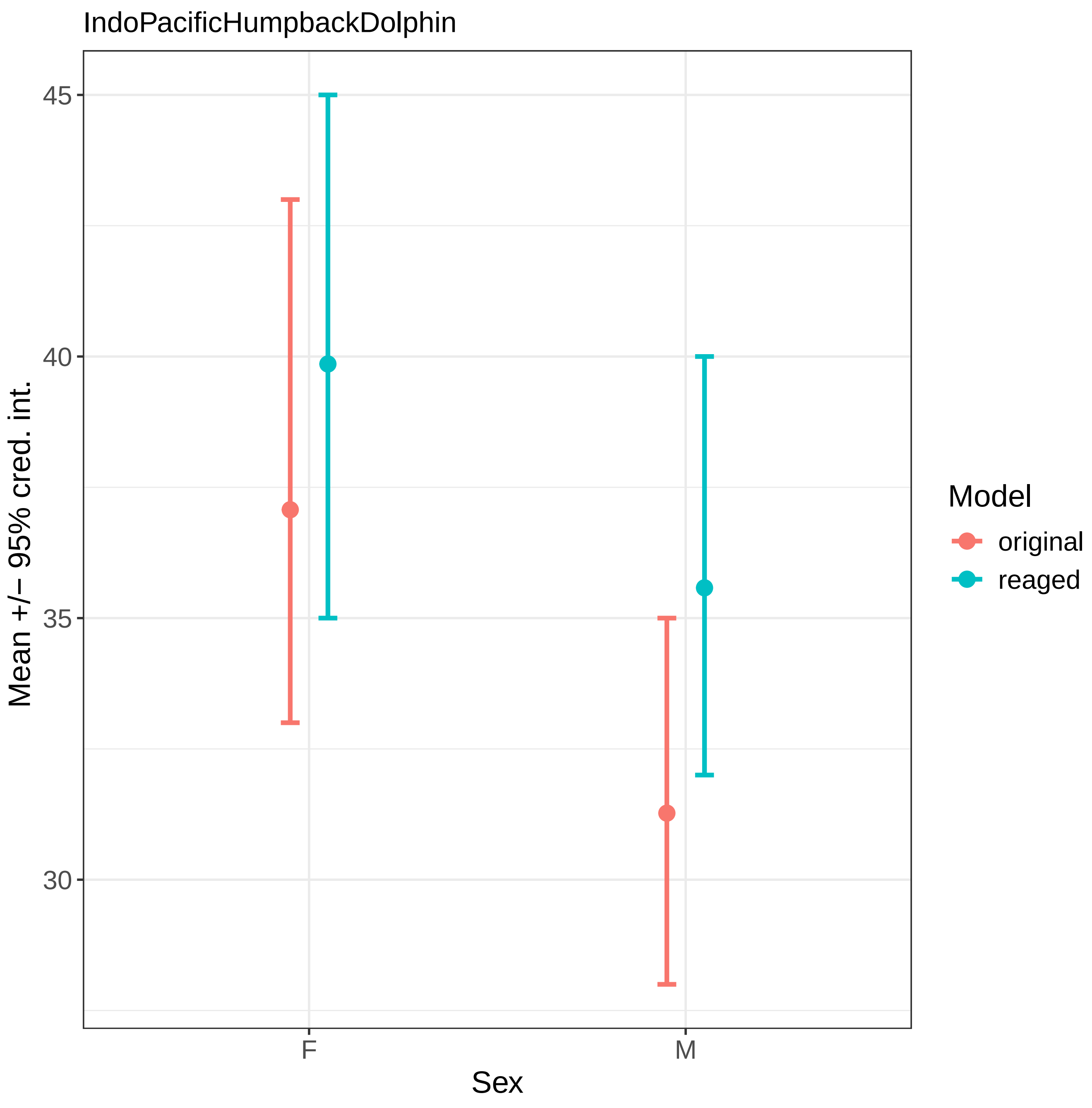
IndoPacificHumpbackDolphin
45
40
35
30
Mean +/− 95% cred. int.
F
M
Sex
Model
original
reaged

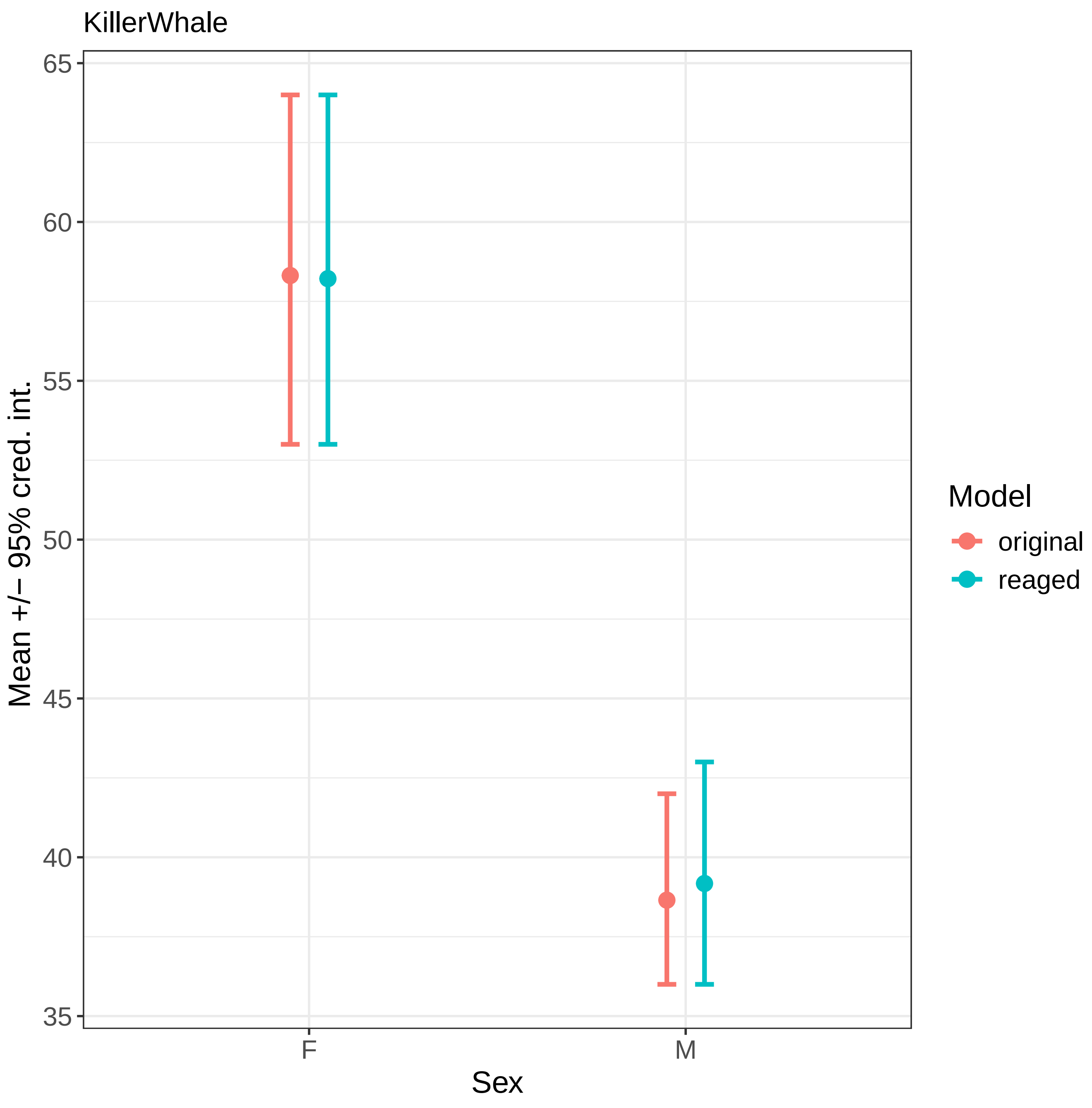

KillerWhale
Mean +/− 95% cred. int.
65
60
55
50
45
40
35
F
M
Sex
Model
original
reaged

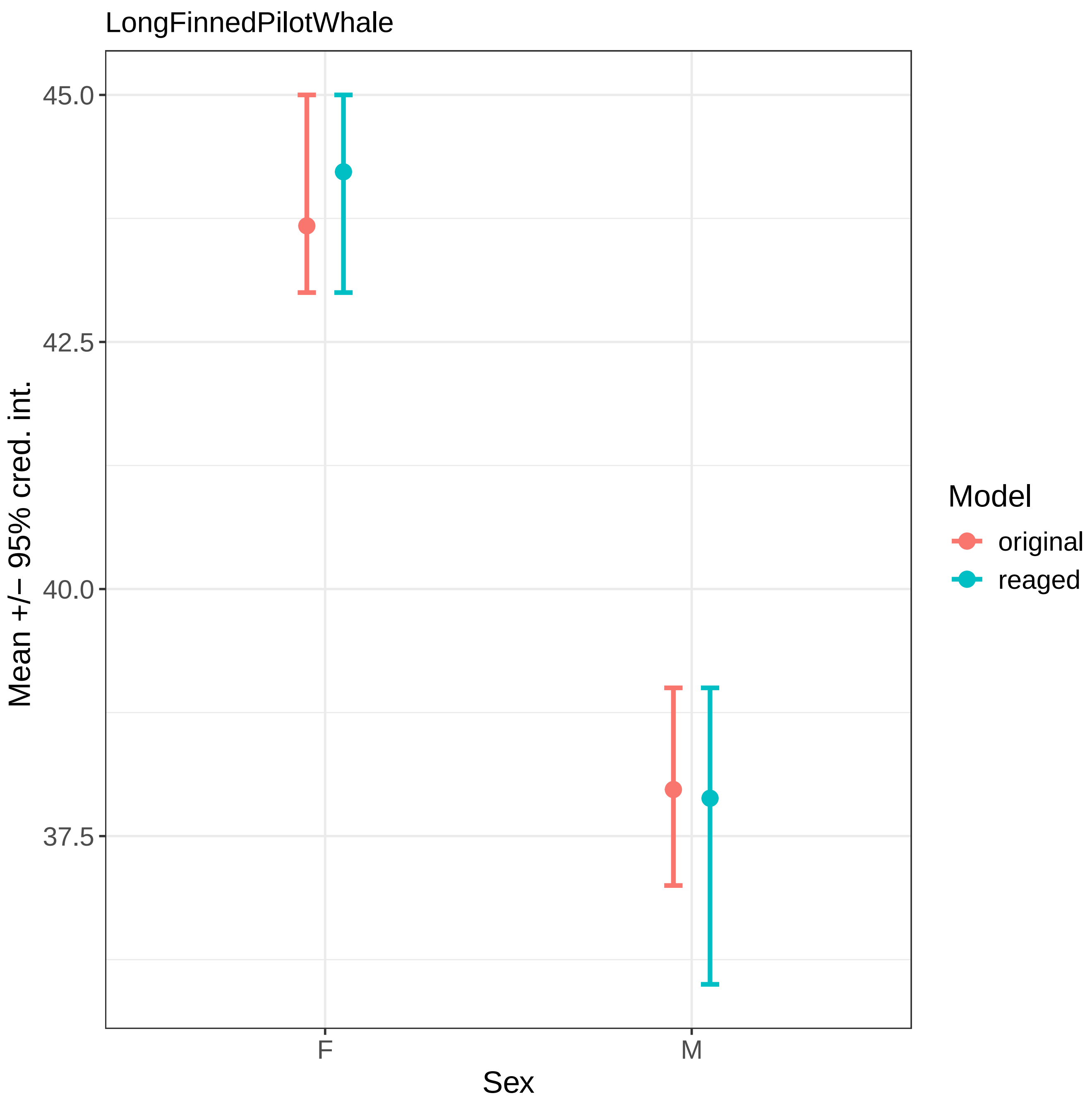
LongFinnedPilotWhale
Mean +/− 95% cred. int.
45.0
42.5
40.0
37.5
F
M
Sex
Model
original
reaged

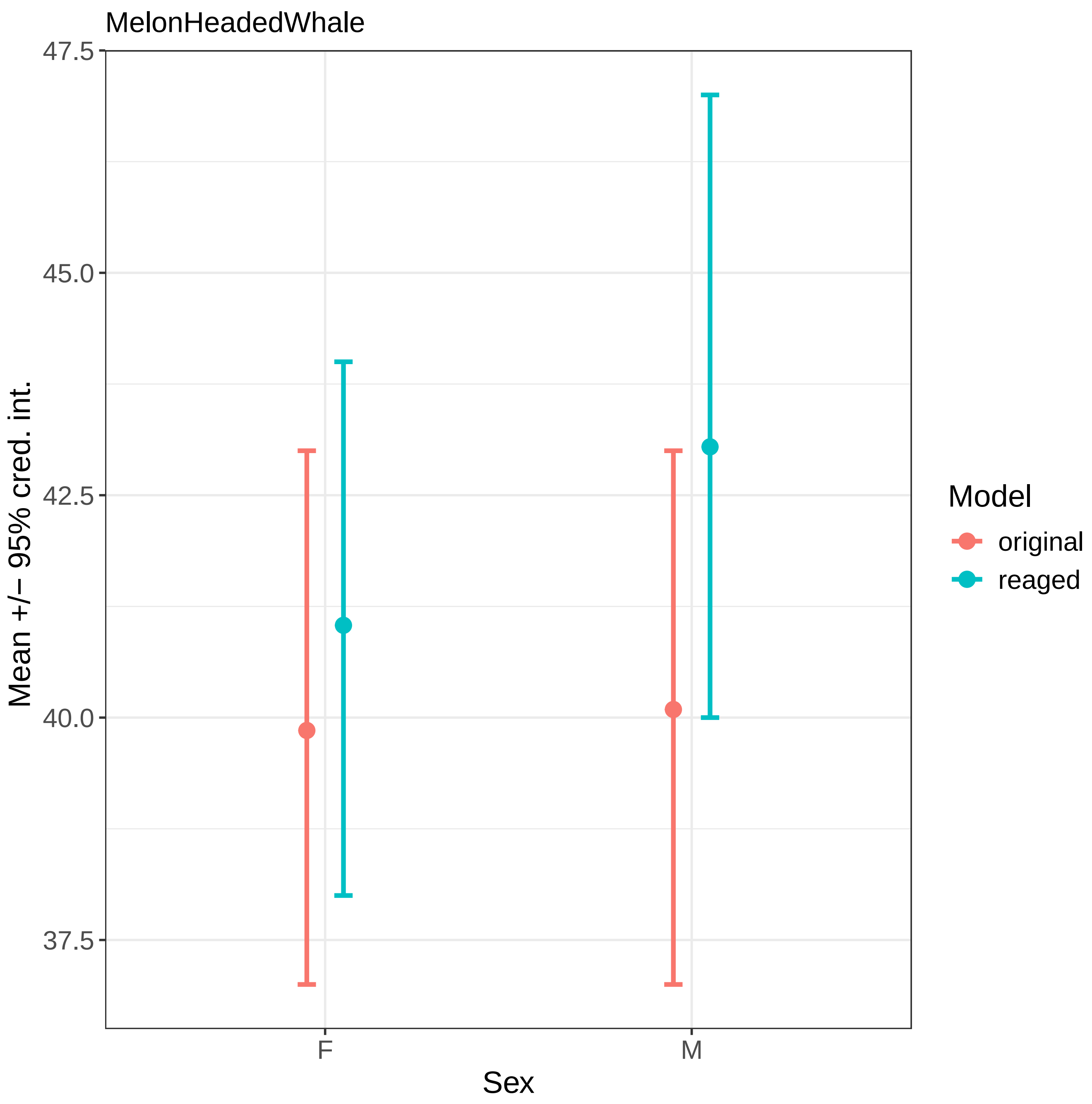

MelonHeadedWhale
47.5
45.0
42.5
40.0
37.5
Mean +/− 95% cred. int.
F
M
Sex
Model
original
reaged

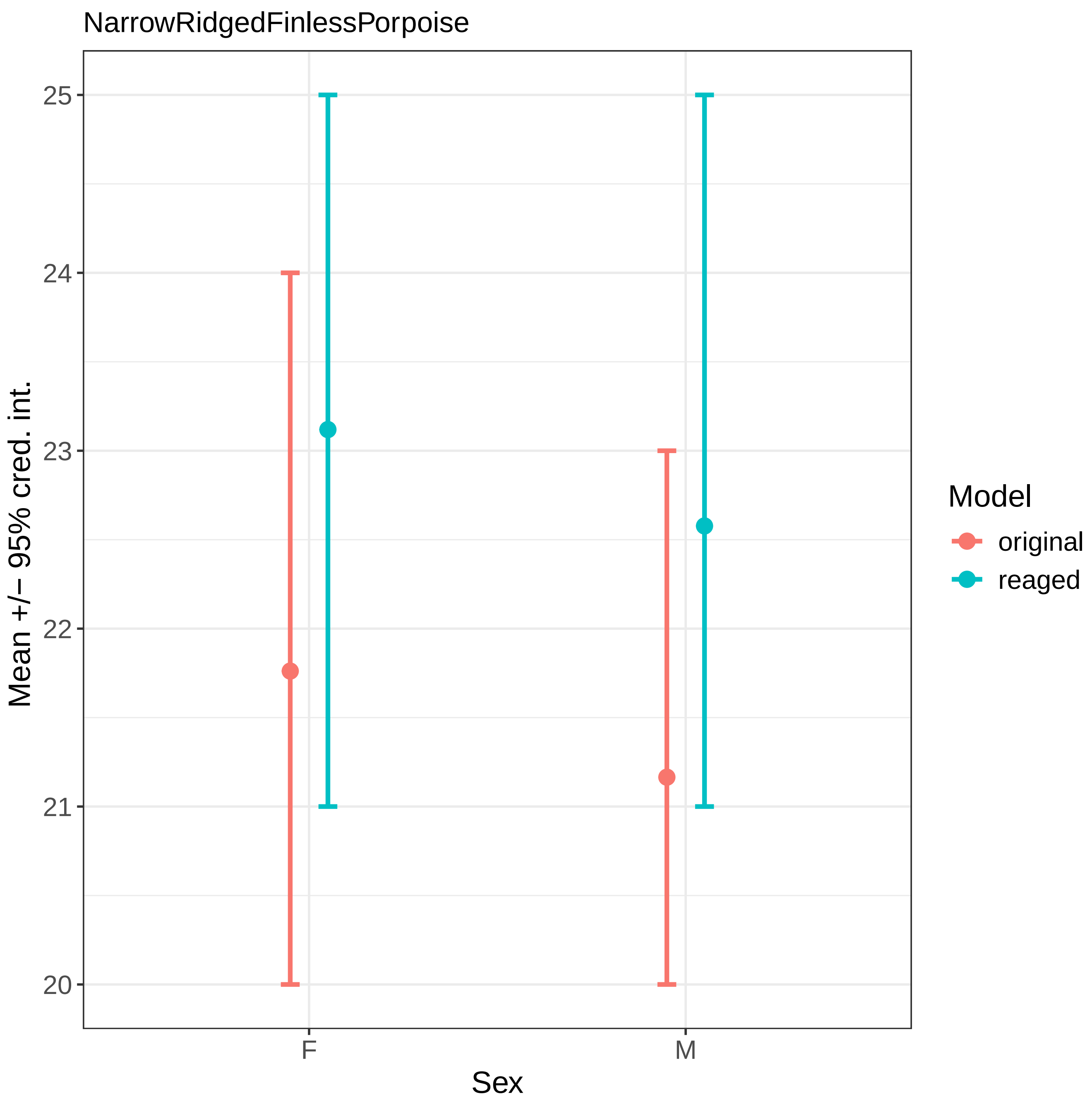
NarrowRidgedFinlessPorpoise
Mean +/− 95% cred. int.
25
24
23
22
21
20
F
M
Sex
Model
original
reaged

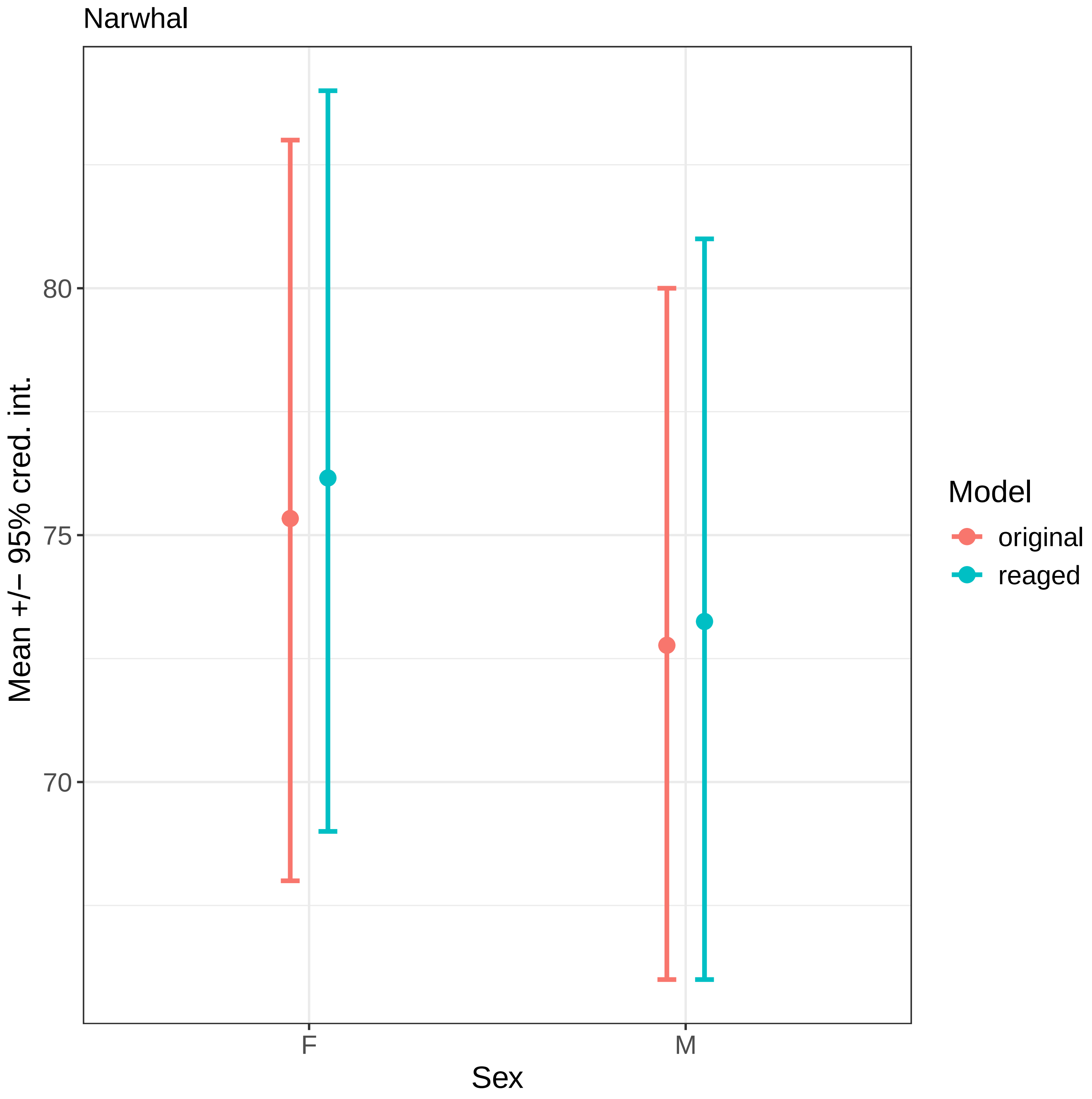
Narwhal
80
75
70
Mean +/− 95% cred. int.
F
M
Sex
Model
original
reaged

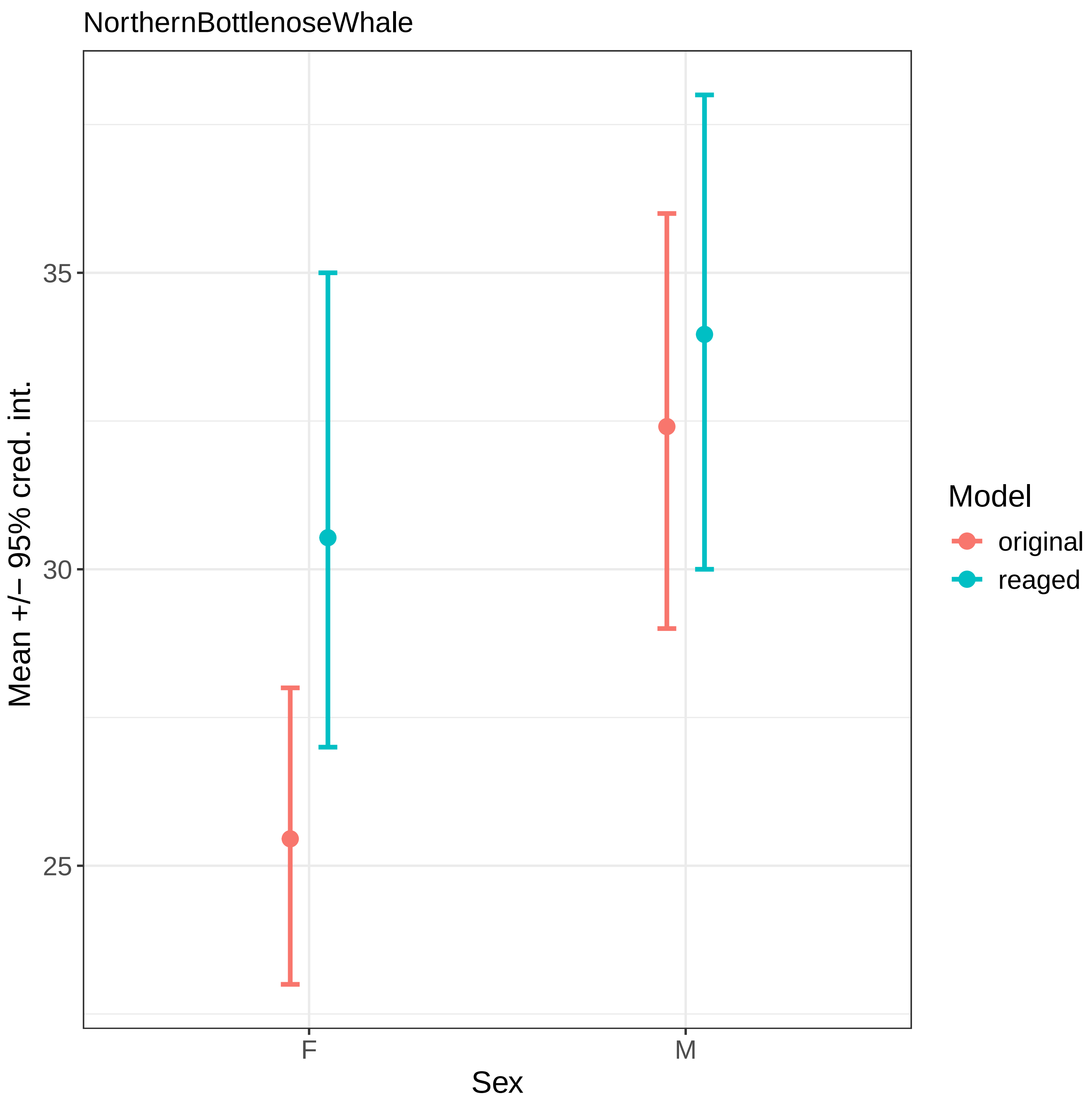
NorthernBottlenoseWhale
Mean +/− 95% cred. int.
35
30
25
F
M
Sex
Model
original
reaged

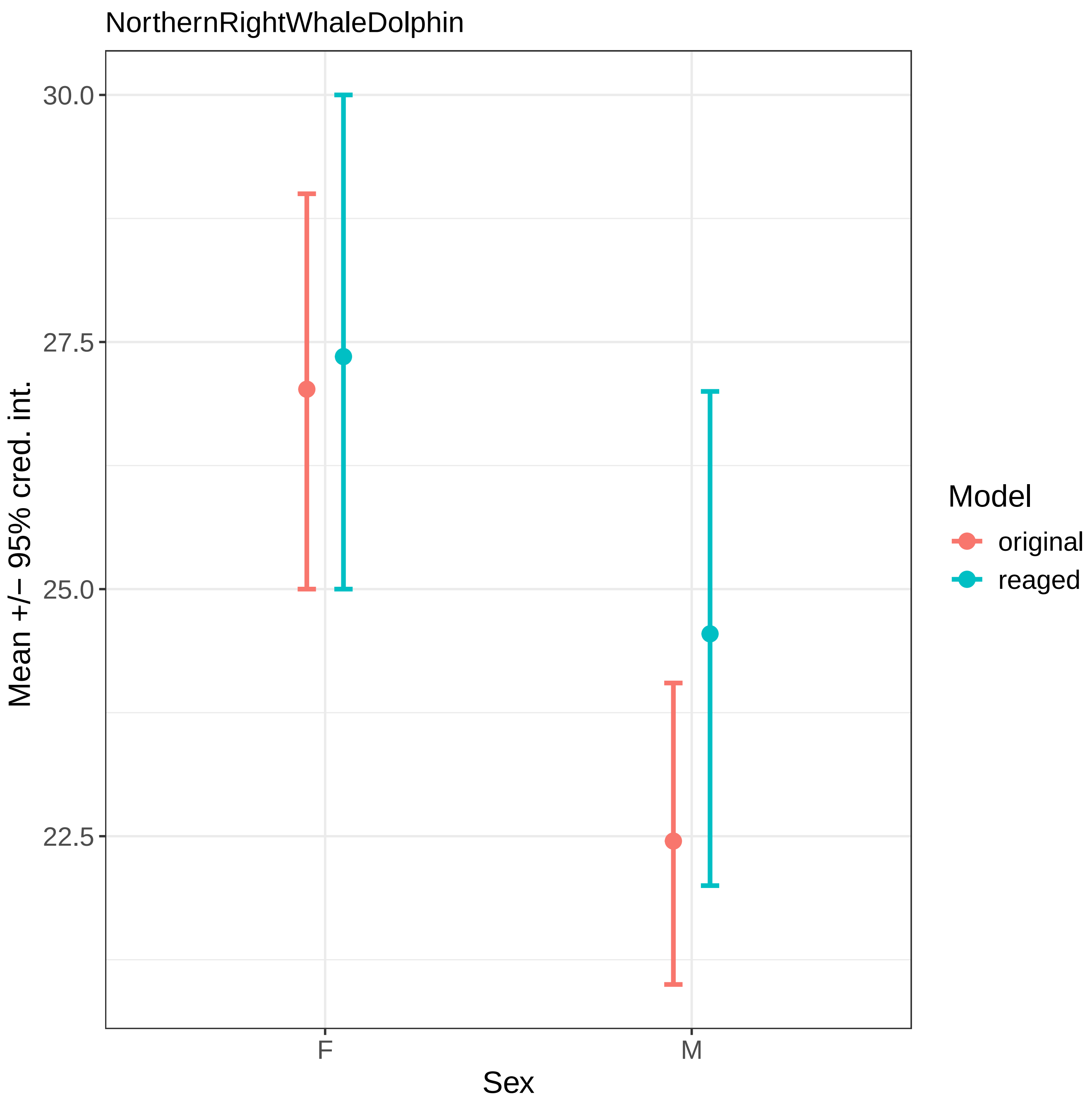

NorthernRightWhaleDolphin
30.0
27.5
25.0
22.5
Mean +/− 95% cred. int.
F
M
Sex
Model
original
reaged

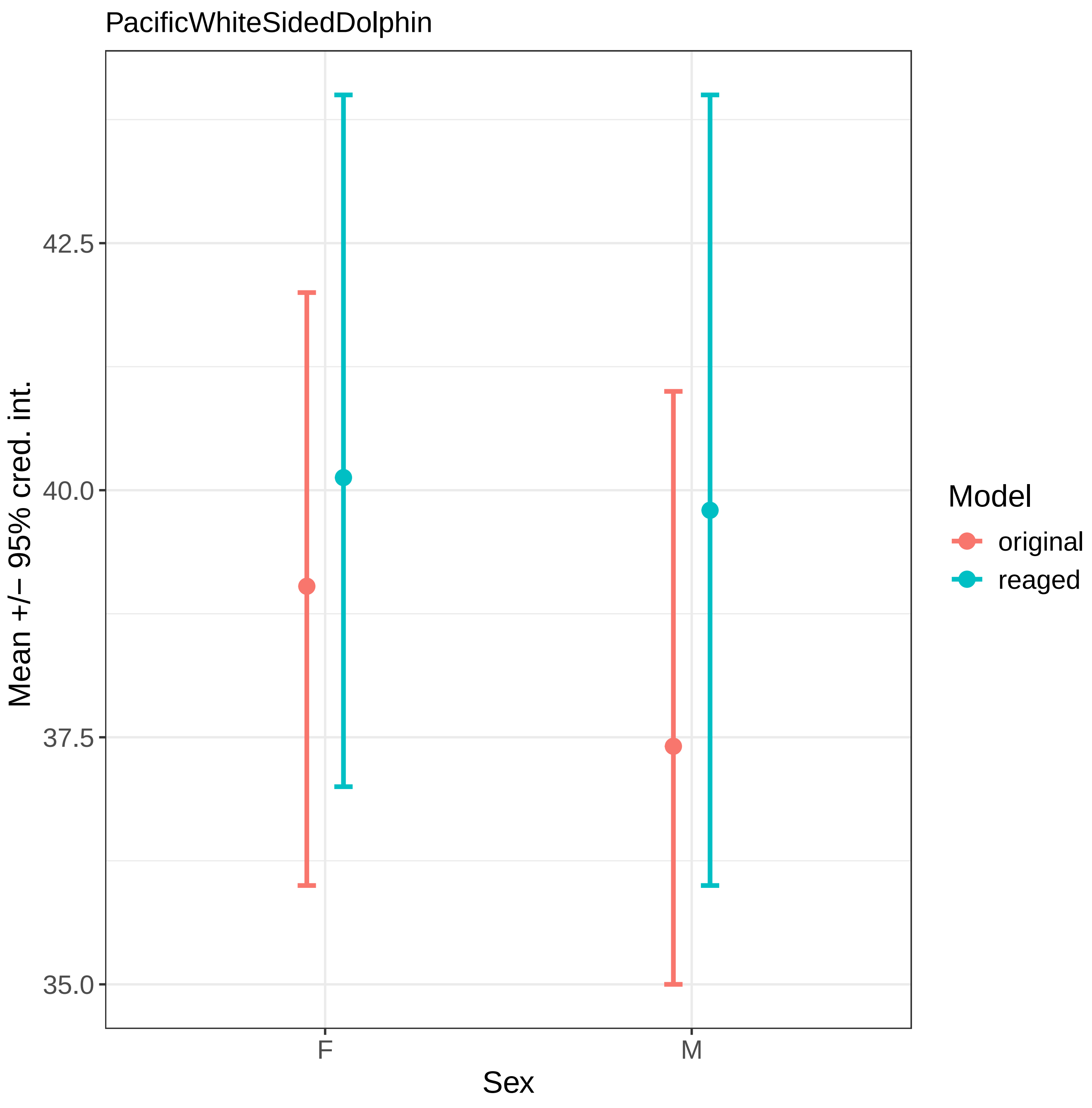
PacificWhiteSidedDolphin
42.5
40.0
37.5
35.0
Mean +/− 95% cred. int.
F
M
Sex
Model
original
reaged

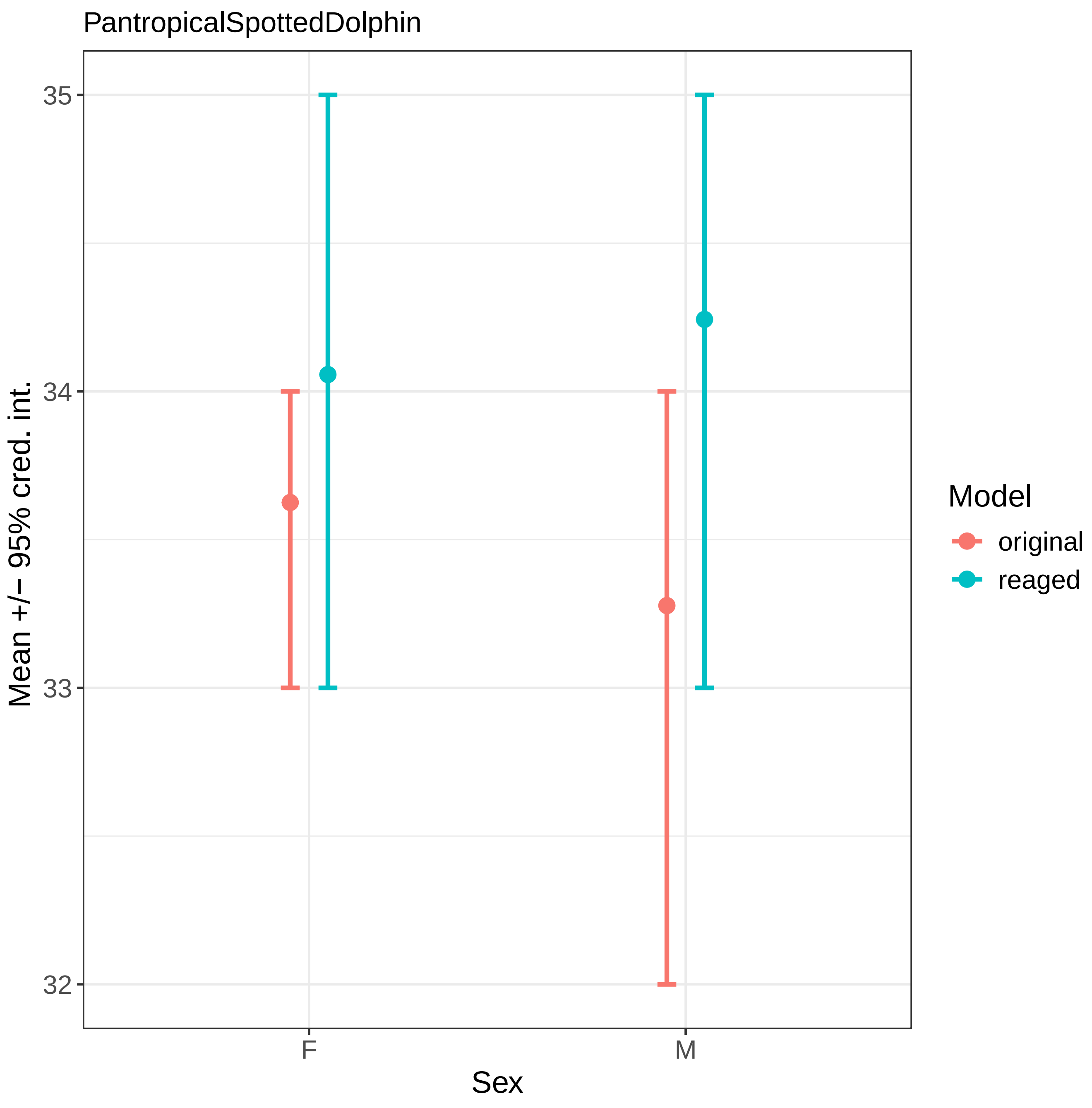
PantropicalSpottedDolphin
35
34
33
32
Mean +/− 95% cred. int.
F
M
Sex
Model
original
reaged

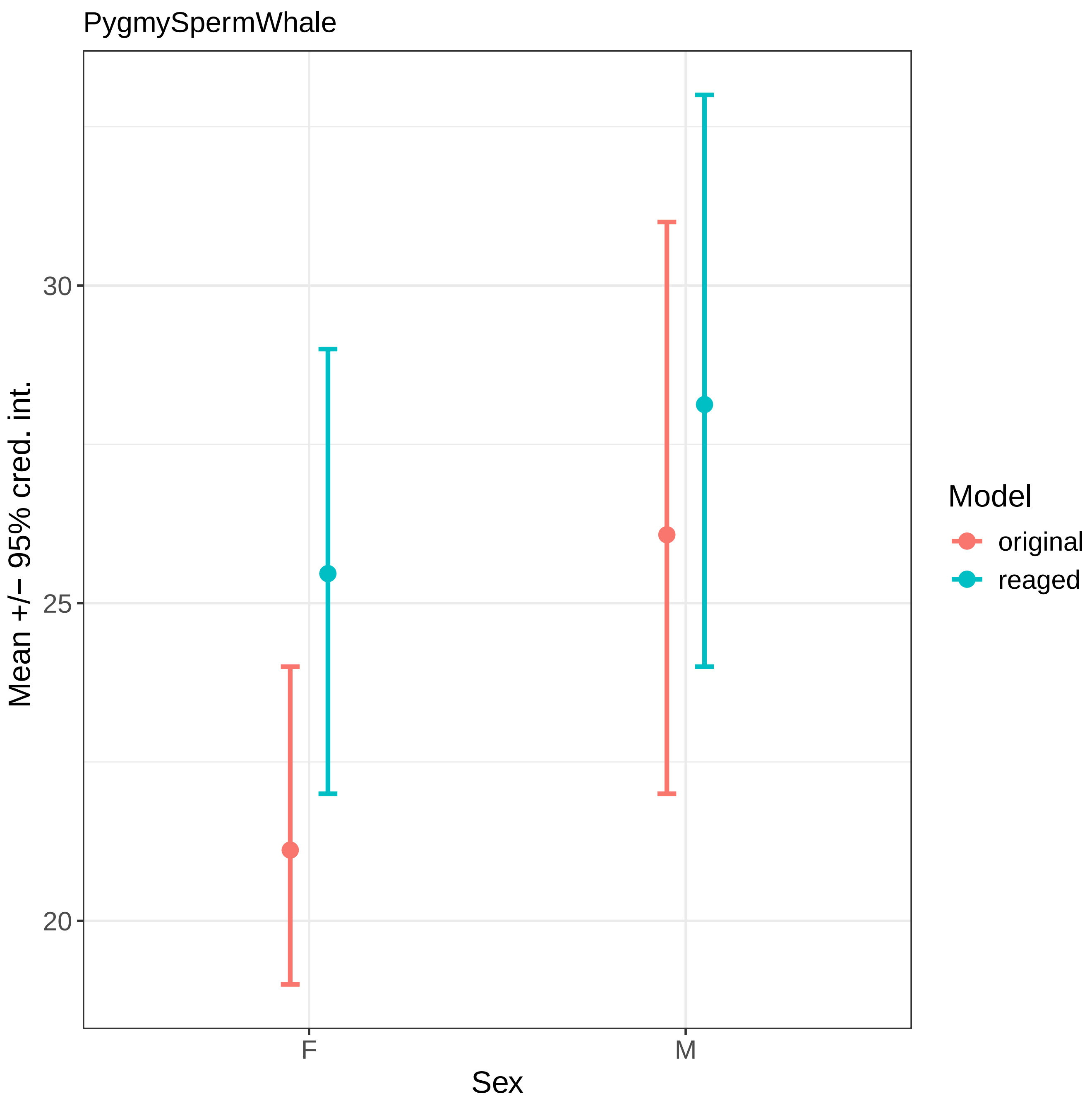
PygmySpermWhale
Mean +/− 95% cred. int.
30
25
20
F
M
Sex
Model
original
reaged

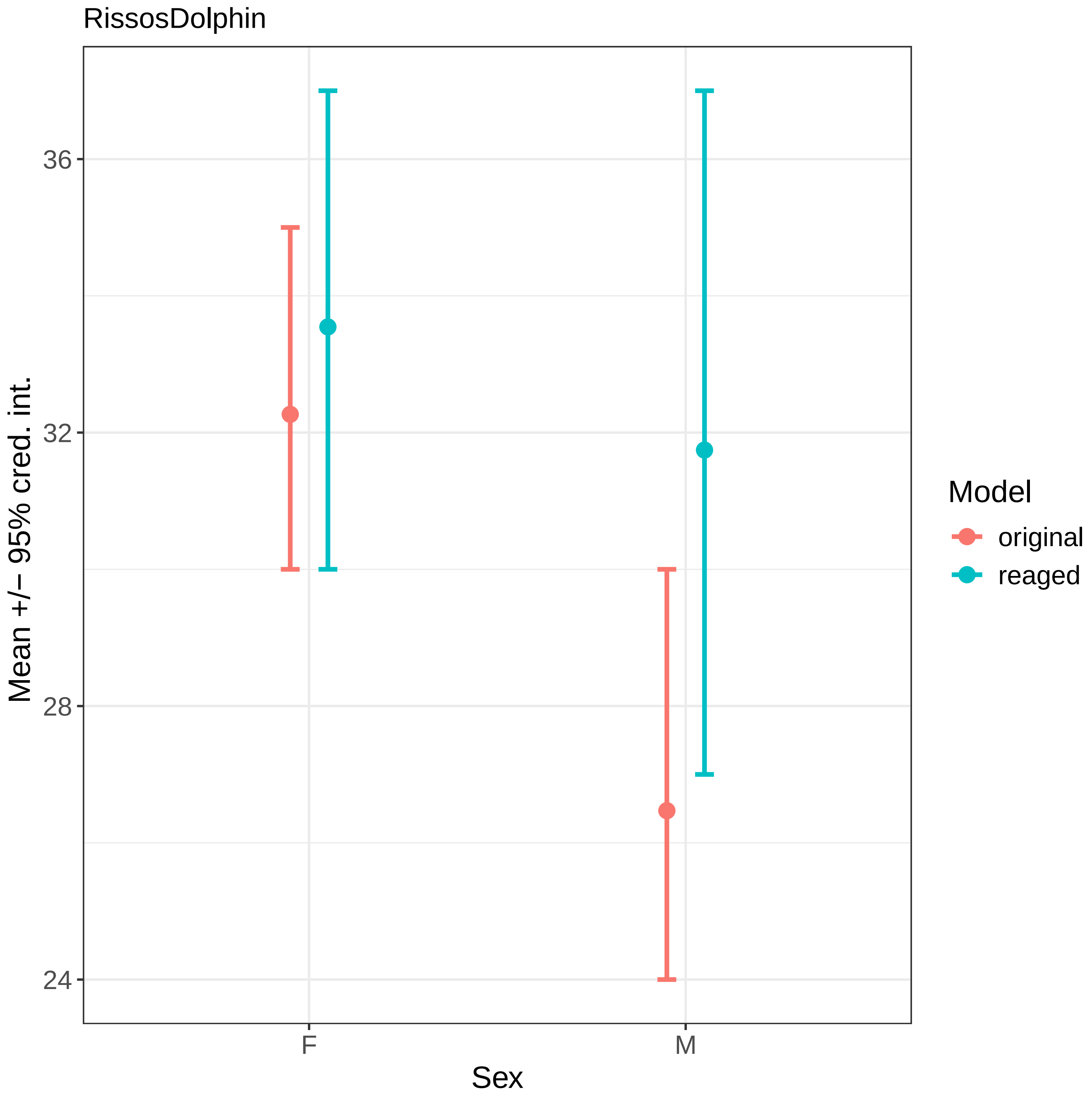
RissosDolphin
Mean +/− 95% cred. int.
36
32
28
24
F
M
Sex
Model
original
reaged

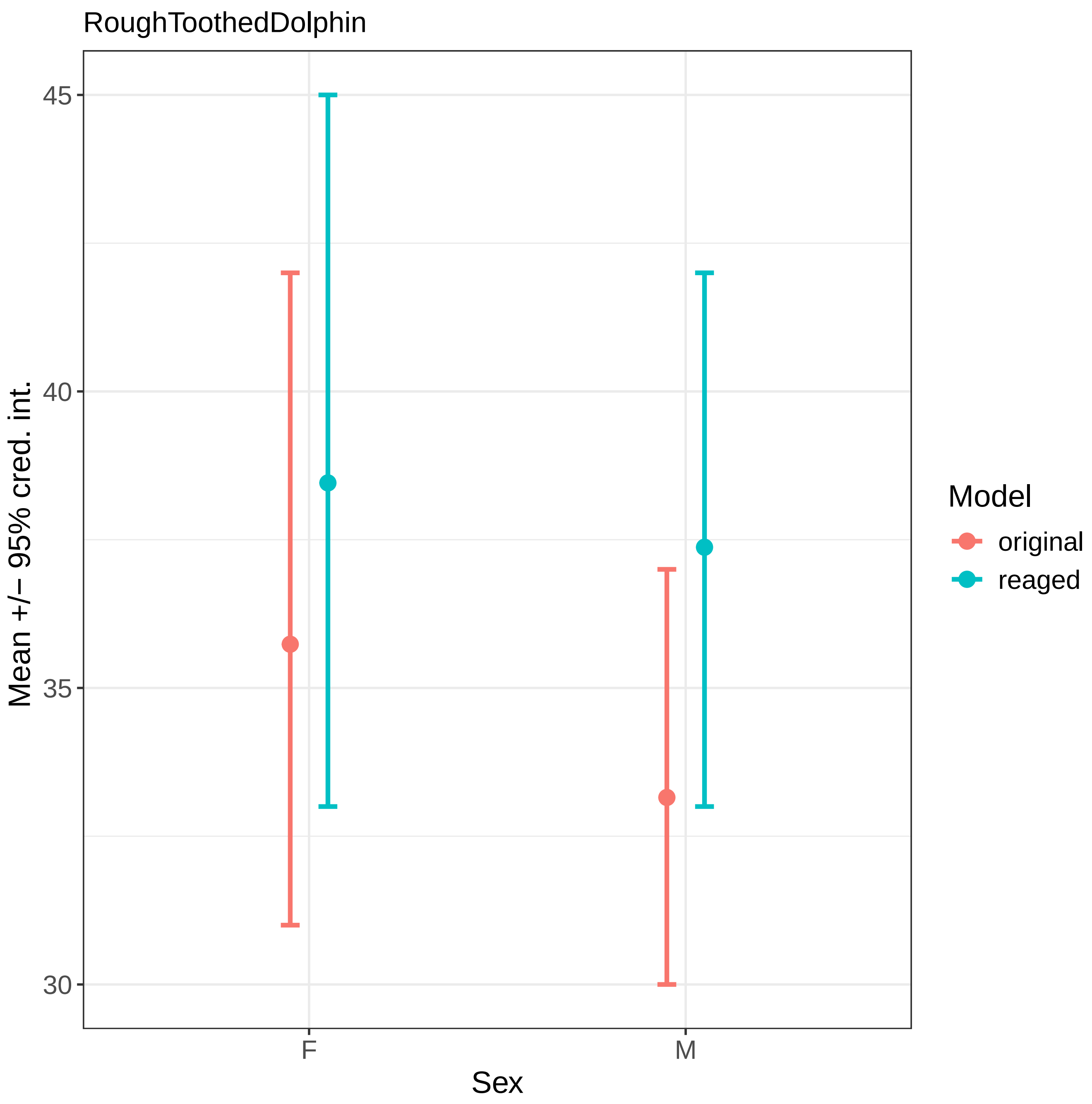
RoughToothedDolphin
Mean +/− 95% cred. int.
45
40
35
30
F
M
Sex
Model
original
reaged

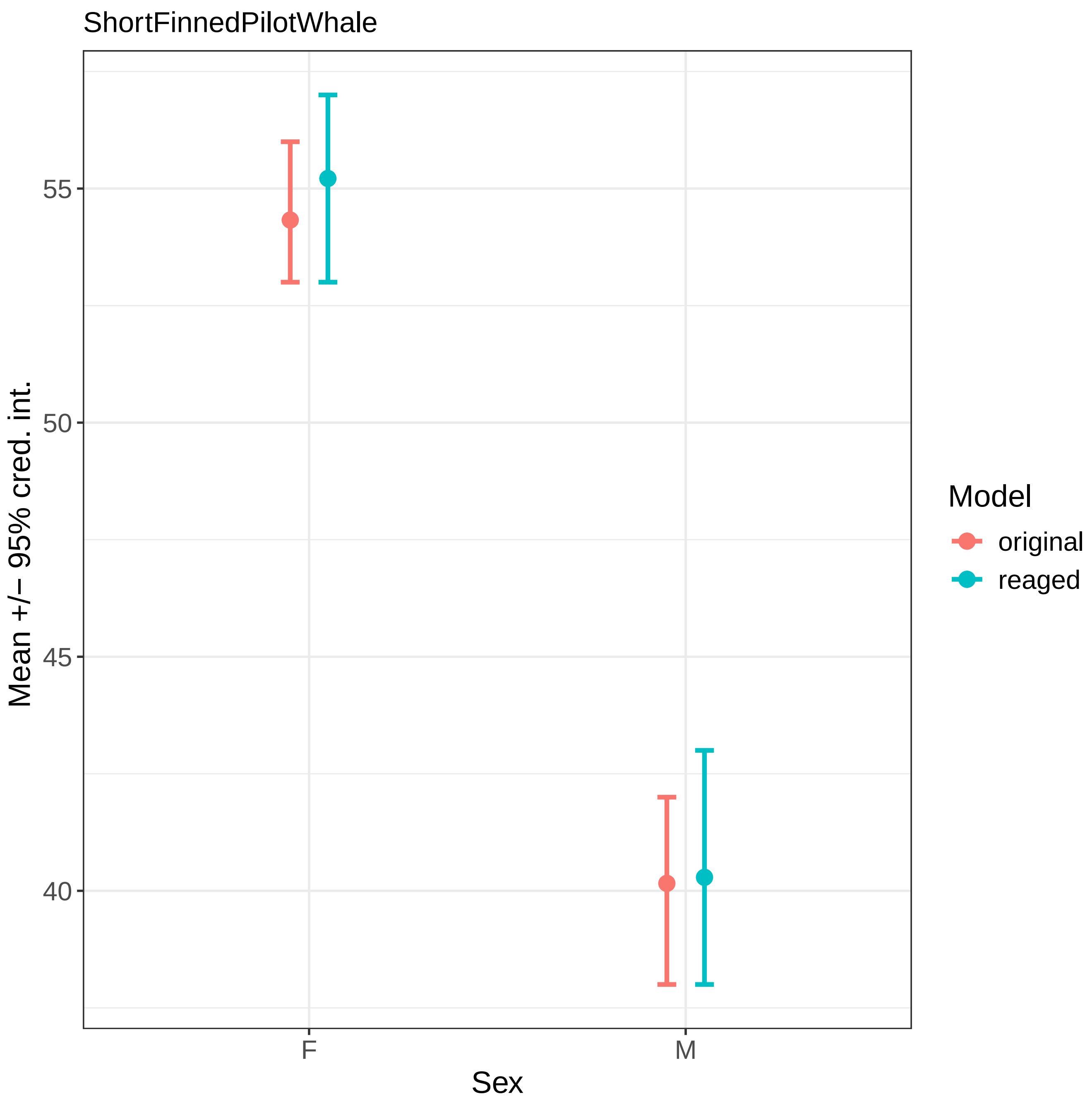
ShortFinnedPilotWhale
Mean +/− 95% cred. int.
55
50
45
40
F
M
Sex
Model
original
reaged

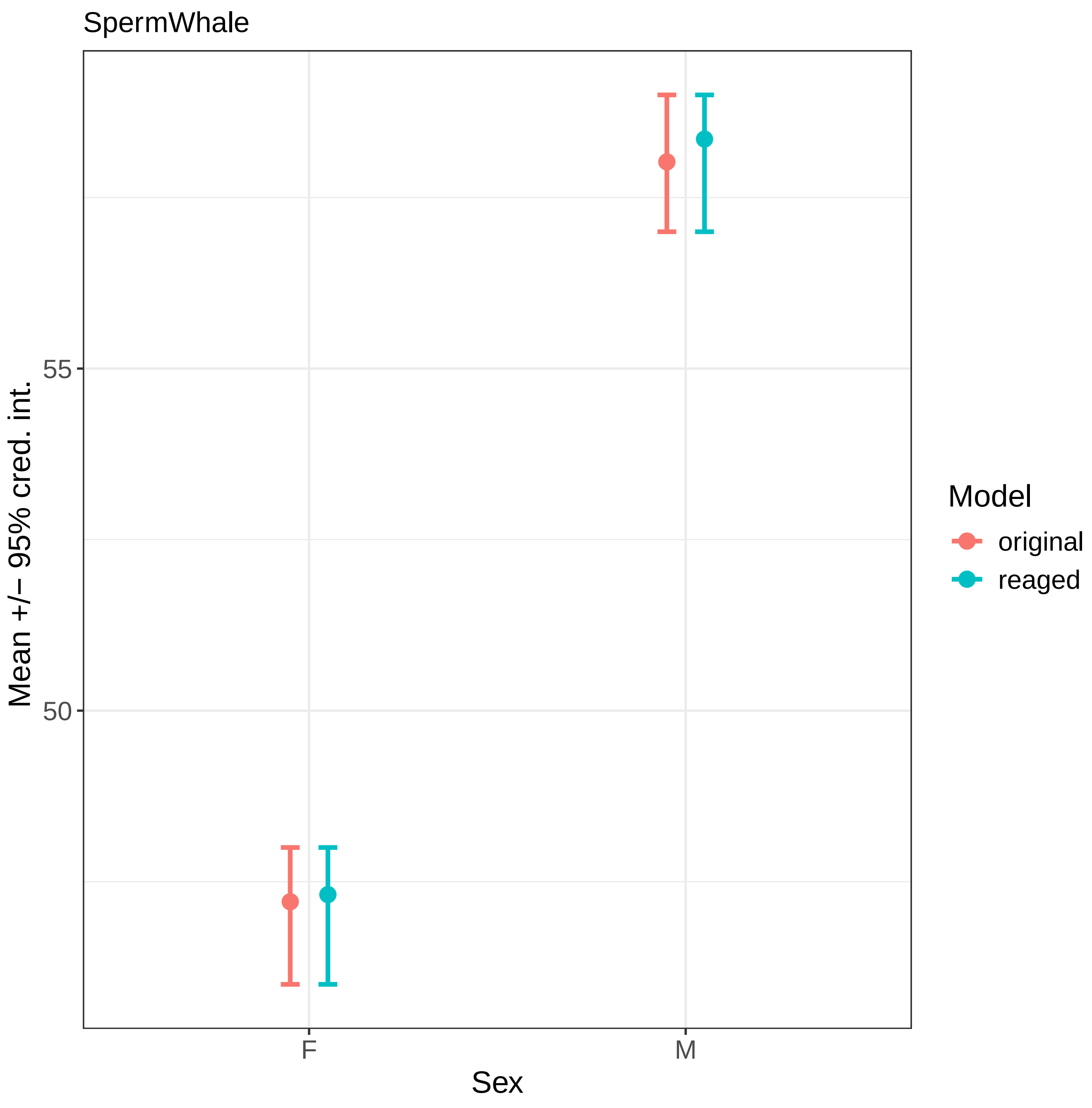
SpermWhale
Mean +/− 95% cred. int.
55
50
F
M
Sex
Model
original
reaged

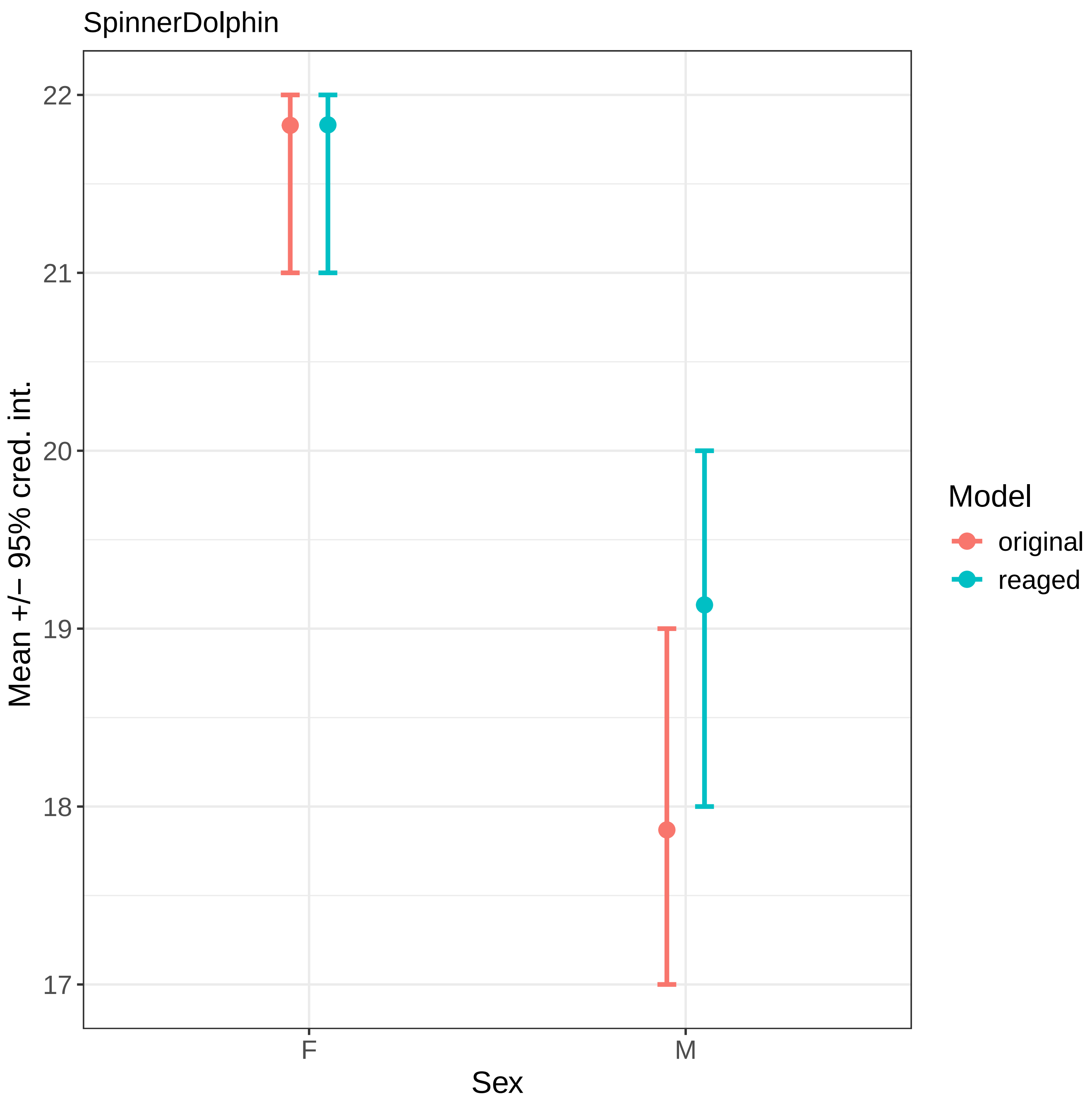

SpinnerDolphin
Mean +/− 95% cred. int.
22
21
20
19
18
17
F
M
Sex
Model
original
reaged

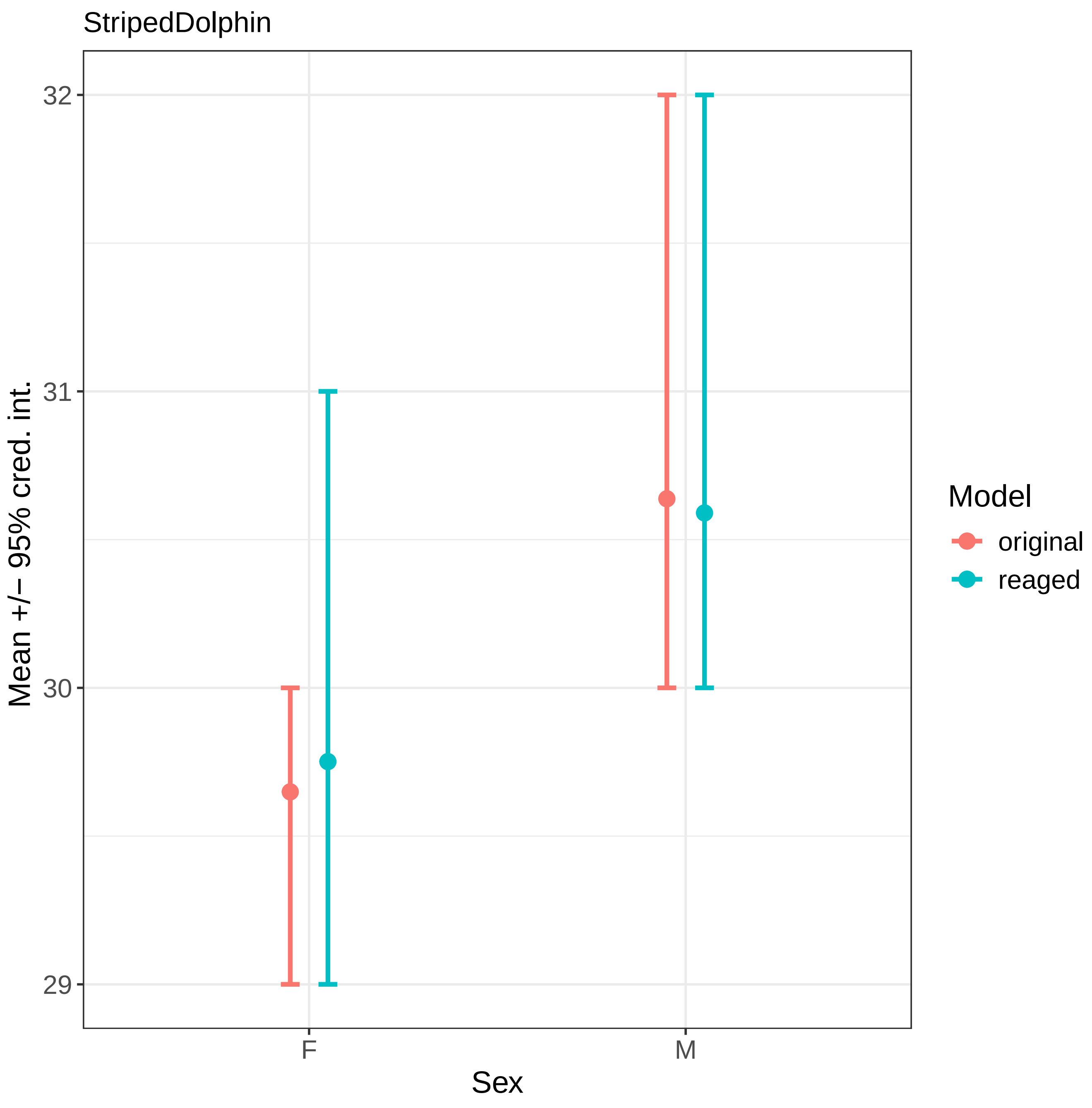
StripedDolphin
Mean +/− 95% cred. int.
32
31
30
29
F
M
Sex
Model
original
reaged

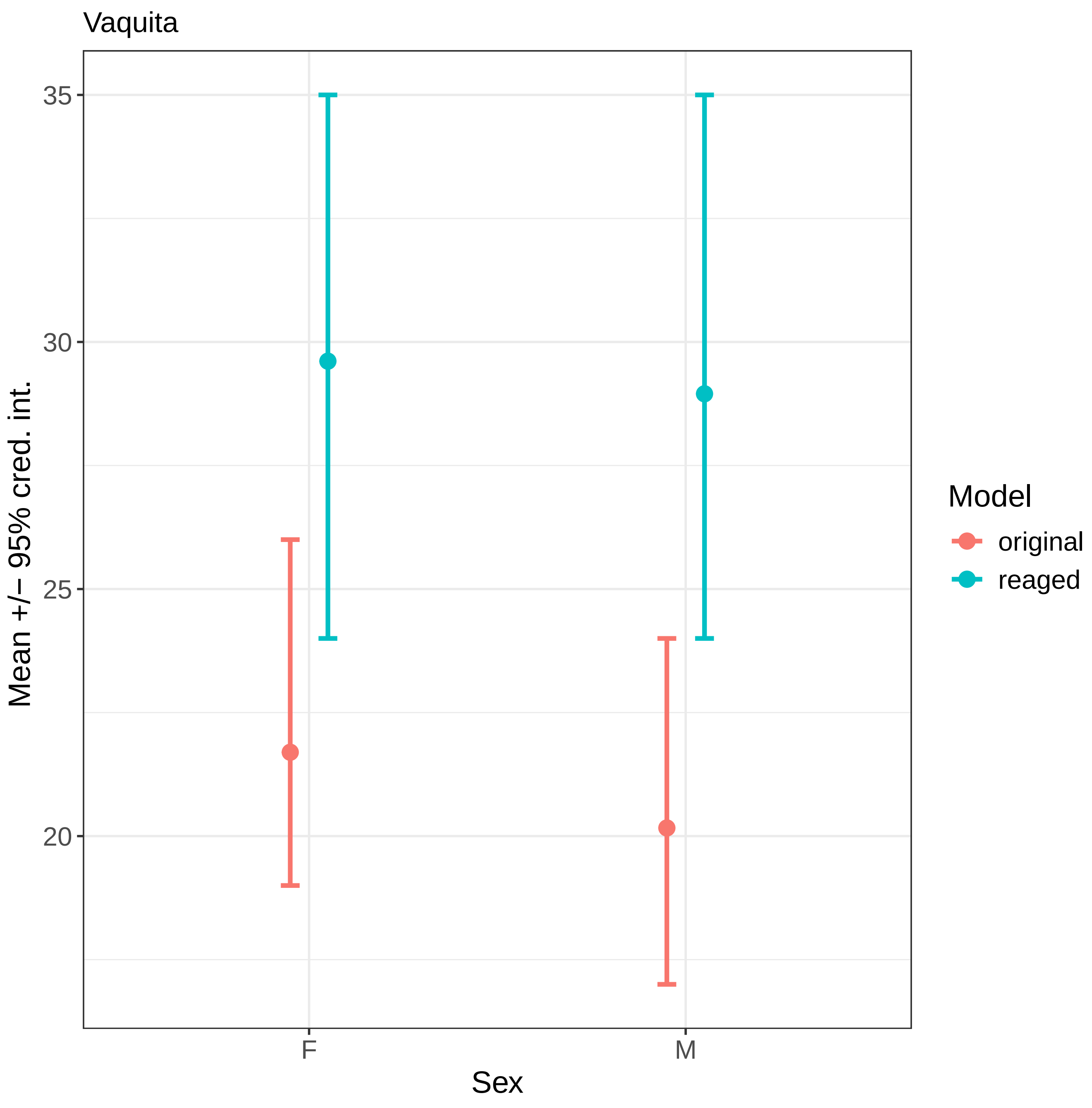
Vaquita
35
30
25
20
Mean +/− 95% cred. int.
F
M
Sex
Model
original
reaged

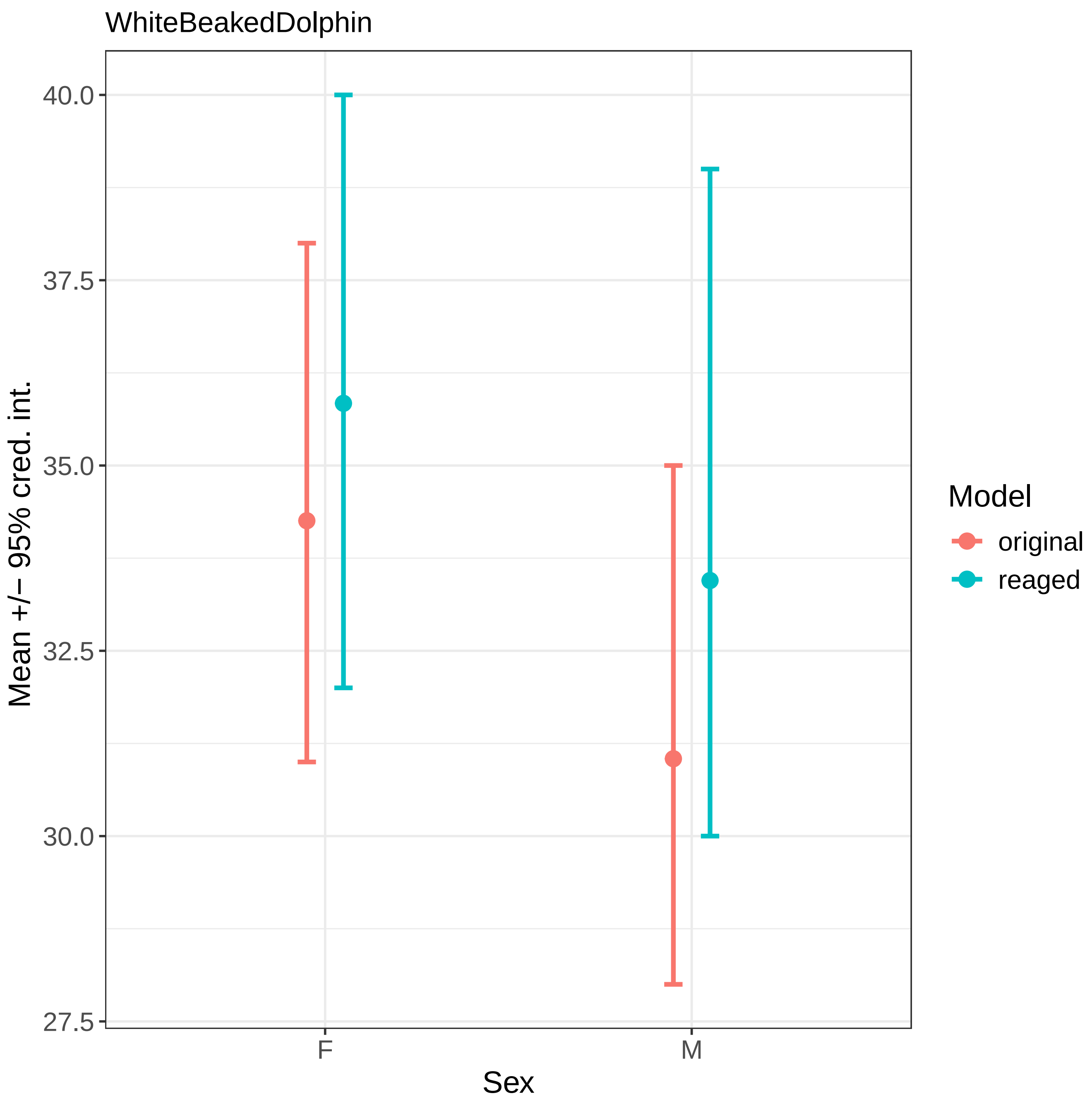
WhiteBeakedDolphin
Mean +/− 95% cred. int.
40.0
37.5
35.0
32.5
30.0
27.5
F
M
Sex
Model
original
reaged
