## Supplementary 3 for "Bayesian inference of toothed whale lifespans"

Counting tooth rings to age Odontocete samples has been an established method for several decades (Perrin & Myrick, 1980). However, the technique has evolved and changed over time (Read, Hohn, & Lockyer, 2018; Barratclough *et al.*, 2023). Here we investigate whether these changes in methodology lead to different age distributions through time.

Five species toothed whale species – beluga whale, common bottlenose dolphins, franciscana, Guiana dolphin, harbour porpoise (see text for species names)- have reasonably sized age-structured samples in four separate decades. For simplicity, where two sexes of multi-decade sampling for the same species we focus only on females, this results in all samples in this analysis being female (though qualitatively the same results occur when males are used). To compare the age distribution through time for each species we (1) collate samples from each decade of data collection (XXX0 – XXX9; $d$), (2) categorise age into five-year bracket ($a$), (3) count the number of samples from a given decade are in each age bracket ($C$), (4) and the number of samples from each decade overall ($N$). We then use these data as the input for a simple liner model of the structure:

Equation S3.1

$$C \sim Binomial(N, p)$$
$$logit(p_i) = \alpha_{a_i} * \delta_{d_i}$$

Where $\alpha$ and $\delta$ are coefficients for age bins ($a$) and decades ($d$) respectively. The model is fitted as a Bayesian in model in R, and Stan using the brms package. All coefficients have weakly informative priors.

In none of the five species is there any evidence of systematic differences in age-distributions between decades (figure S3.1). This suggests that our samples, and therefore our conclusions, are robust to changing tooth ageing methodologies.

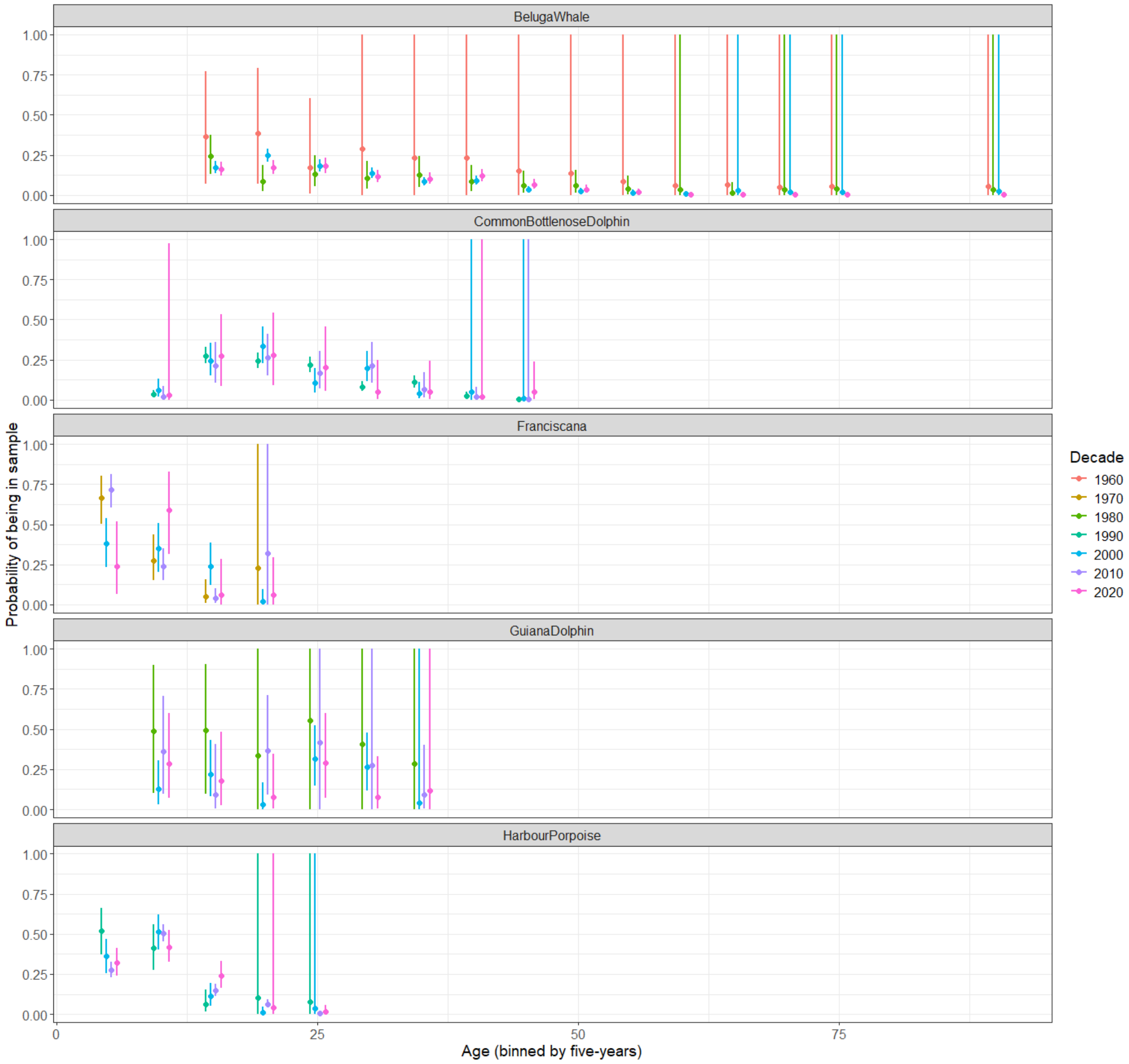


Figure S3.1. Comparison of age distributions across decades. Each panel shows differences in age distributions between decades for a given species. Points shows the mean posterior predicted probability of a given age-bin from a given decade (see equation S3.1), errors show the 95% credible interval. Point and error colour shows the decade the data are drawn from.

### S3 References

Barratclough A, McFee WE, Stolen M, Hohn AA, Lovewell GN, Gomez FM, Smith CR, Garcia-Parraga D, Wells RS, Parry C, Daniels R, Ridgway SH & Schwacke L. 2023. How to estimate age of old bottlenose dolphins (Tursiops truncatus); by tooth or pectoral flipper? *Frontiers in Marine Science* 10.

Perrin WF & Myrick AC. 1980. *Age determination of toothed whales and sirenians: report of the workshop*.

Read FL, Hohn AA & Lockyer CH. 2018. A review of age estimation methods in marine mammals with special reference to monodontids. *NAMMCO Scientific Publications* 8: 1–67.
