## Supplementary 4 for "Bayesian inference of toothed whale lifespans"

For the analysis presented in the main text we assume that the population change (*pr* parameter) can be defined by the source of the data for the population. Specifically, we assume that samples from large-scale by-catch or whaling/drive fisheries represent a declining population, and that samples from subsistence whaling represent stable populations. These assumptions are necessarily coarse, here we explore the sensitivity of our results to our assumptions be re-running all models assuming that we have no information about population change for any population (*pr = 0* for all populations). The model structure, and all other parameters are the same as presented in the main text.

The 95% credible interval predicted ordinary maximum lifespan for species largely or completely overlaps between our original model (as presented in the main text) and our model with no population change assumption (Pr0 models) for both sexes in 31 species (figure S4.1). In five species-sexes – Atlantic white-sided dolphin- both sexes, baiji-females, Commerson's dolphin- females, Fraser's dolphin-females and vaquita-males – the 95% credible intervals of original and Pr0 models do not overlap. These species are all species with smaller sample sizes which were also different between models in our reageing analysis (supplementary 2).

These results suggest that in most species our predicted ordinary maximum lifespan predictions are not sensitive to our population change assumptions, but in a few species with small sample sizes the estimates can be affected by these assumptions.

Figure S4.1. Comparison of ordinary maximum lifespan estimates for the original (red) and Pr0 (blue) models for each species-sex. Each figure shows one species, with sexes on the x axis. In each plot, the point shows the posterior mean of the distribution of ordinary maximum lifespans derived from the model, and the lines show the 95% credible interval. For clarity each figure is depicted on it's own page.

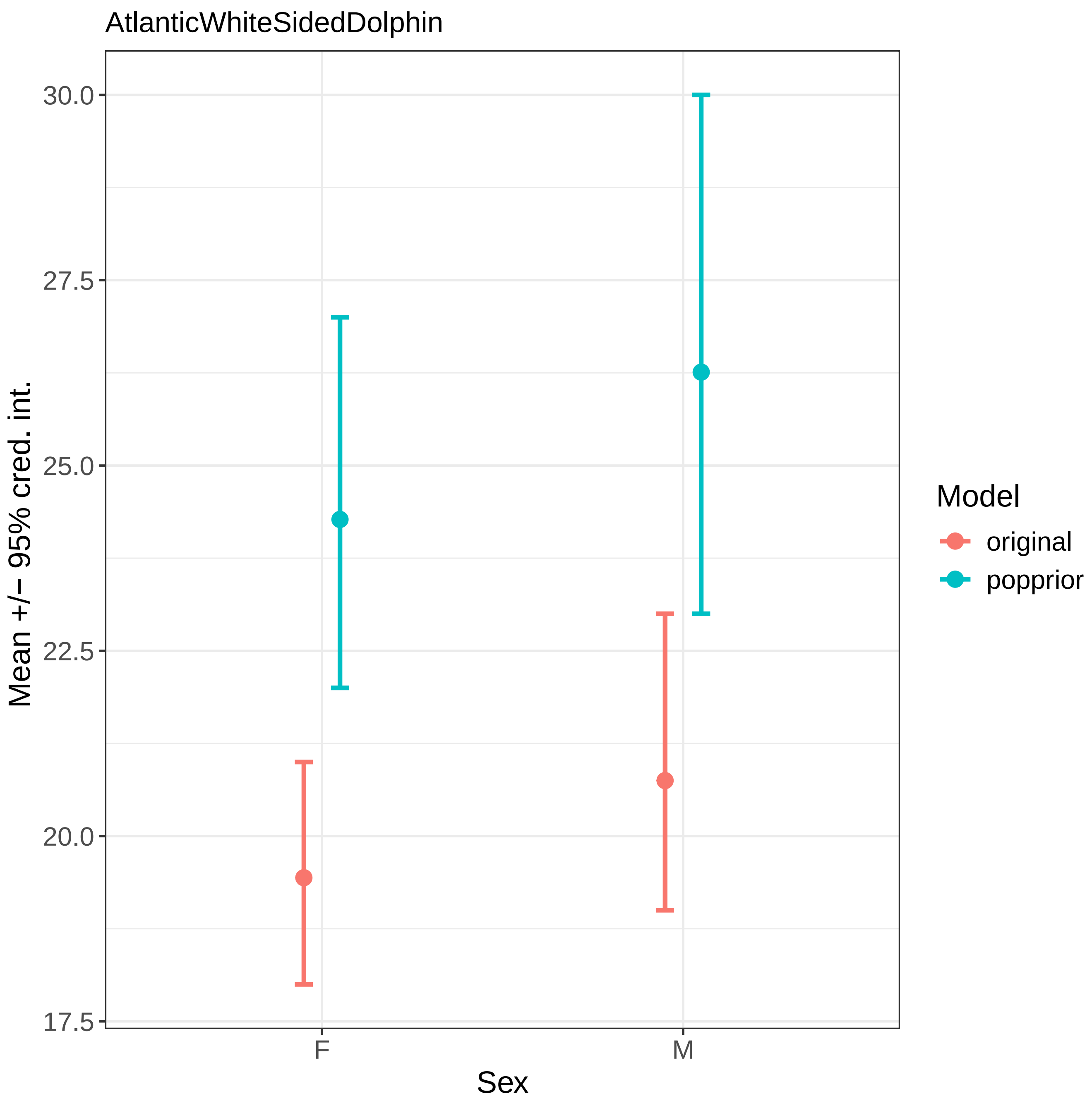
AtlanticWhiteSidedDolphin
Mean +/− 95% cred. int.
30.0
27.5
25.0
22.5
20.0
17.5
F
M
Sex
Model
original
popprior

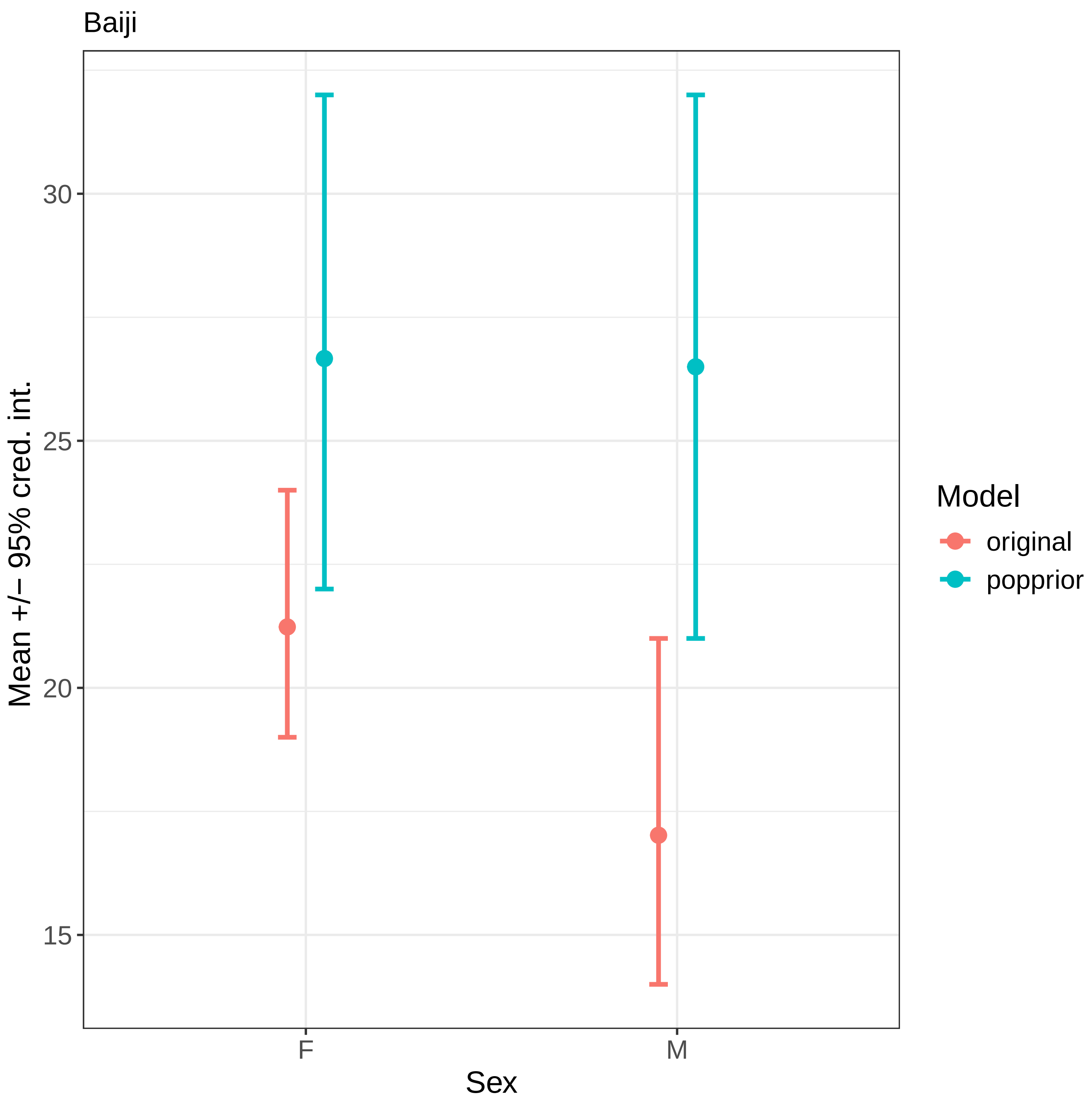
Baiji
Mean +/− 95% cred. int.
30
25
20
15
F
M
Sex
Model
original
popprior

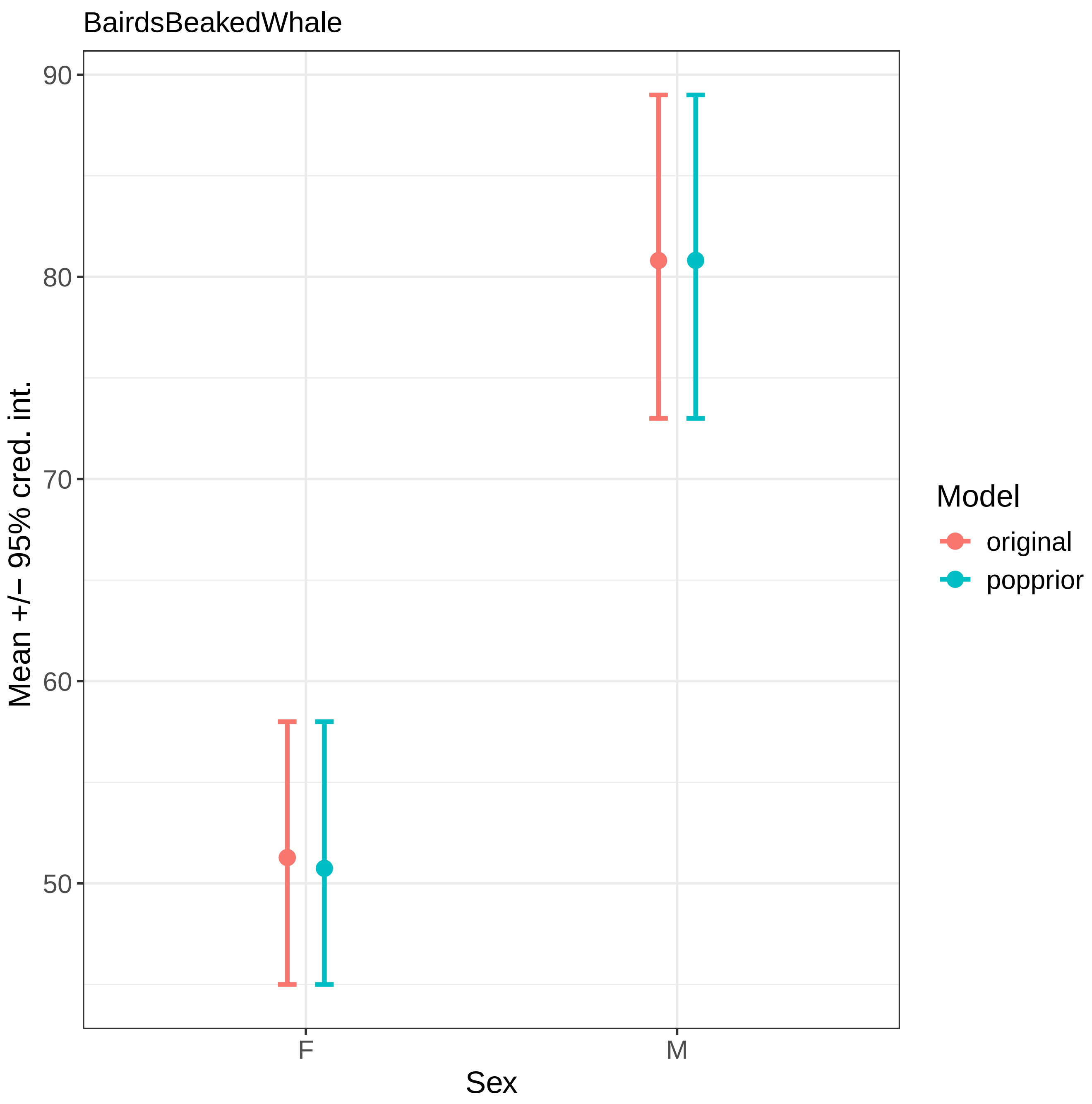

BairdsBeakedWhale
90
80
70
60
50
Mean +/− 95% cred. int.
F
M
Sex
Model
original
popprior

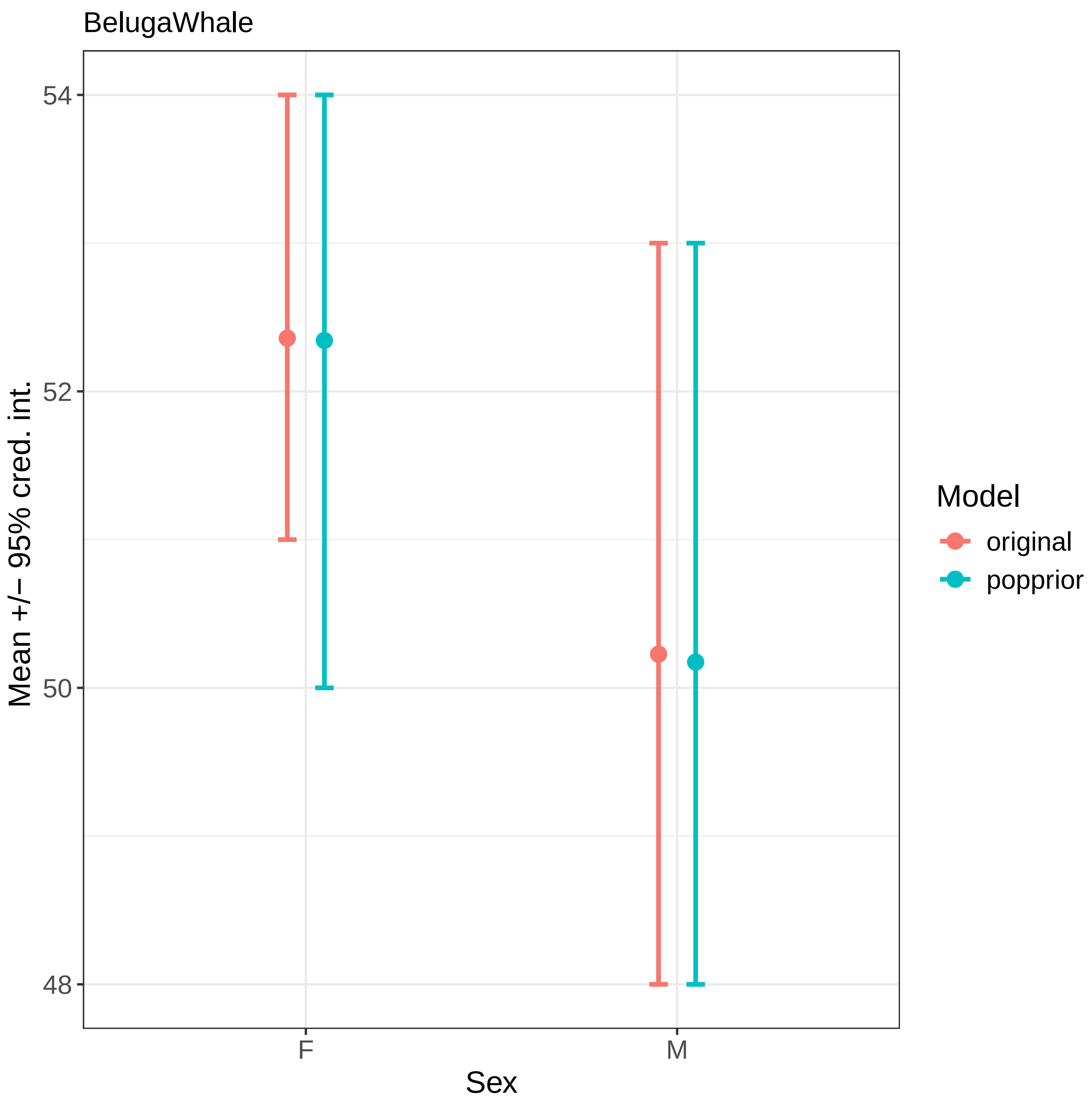

BelugaWhale
54
52
50
48
Mean +/− 95% cred. int.
F
M
Sex
Model
original
popprior

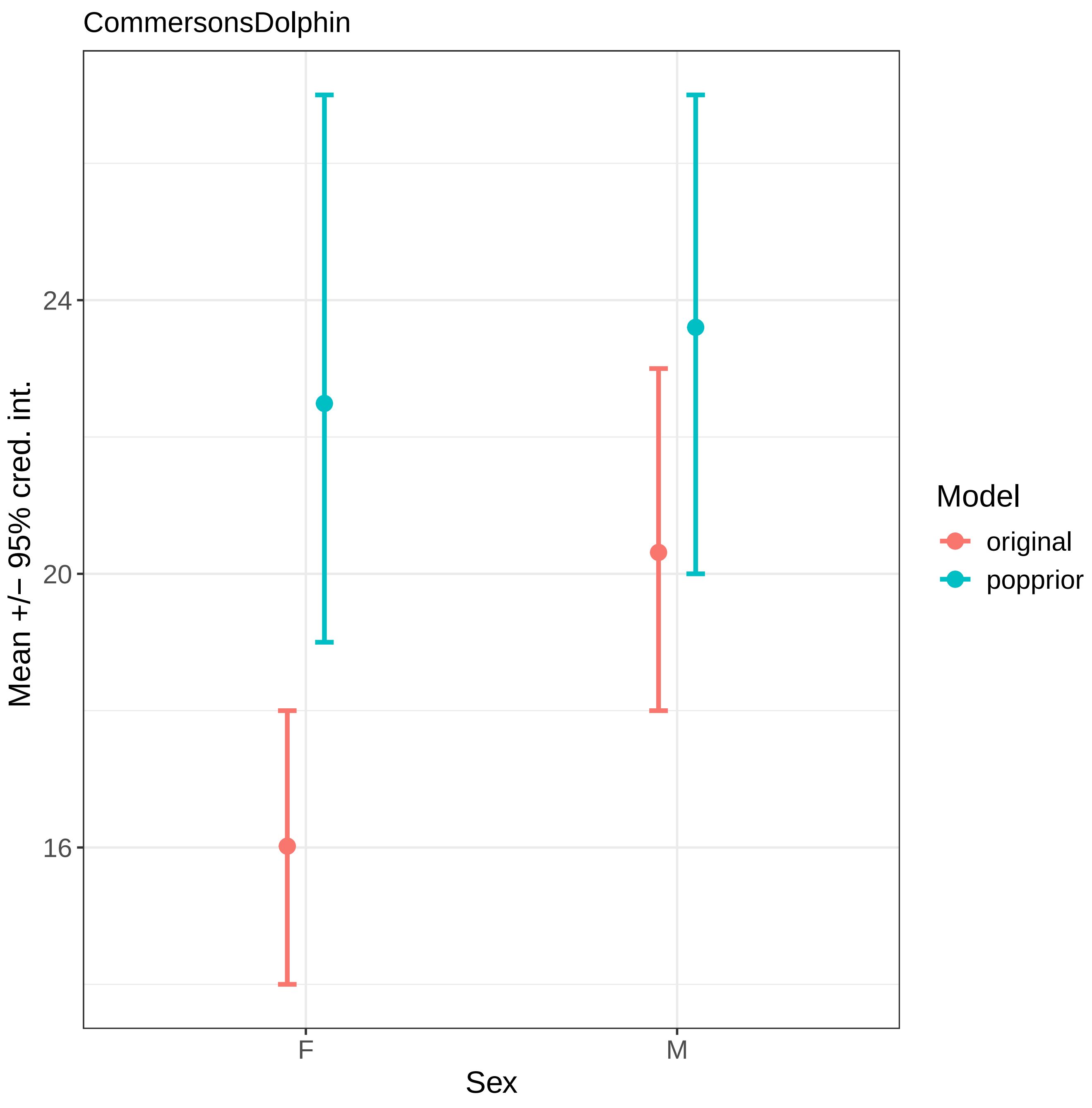
CommersonsDolphin
Mean +/− 95% cred. int.
24
20
16
F
M
Sex
Model
original
popprior

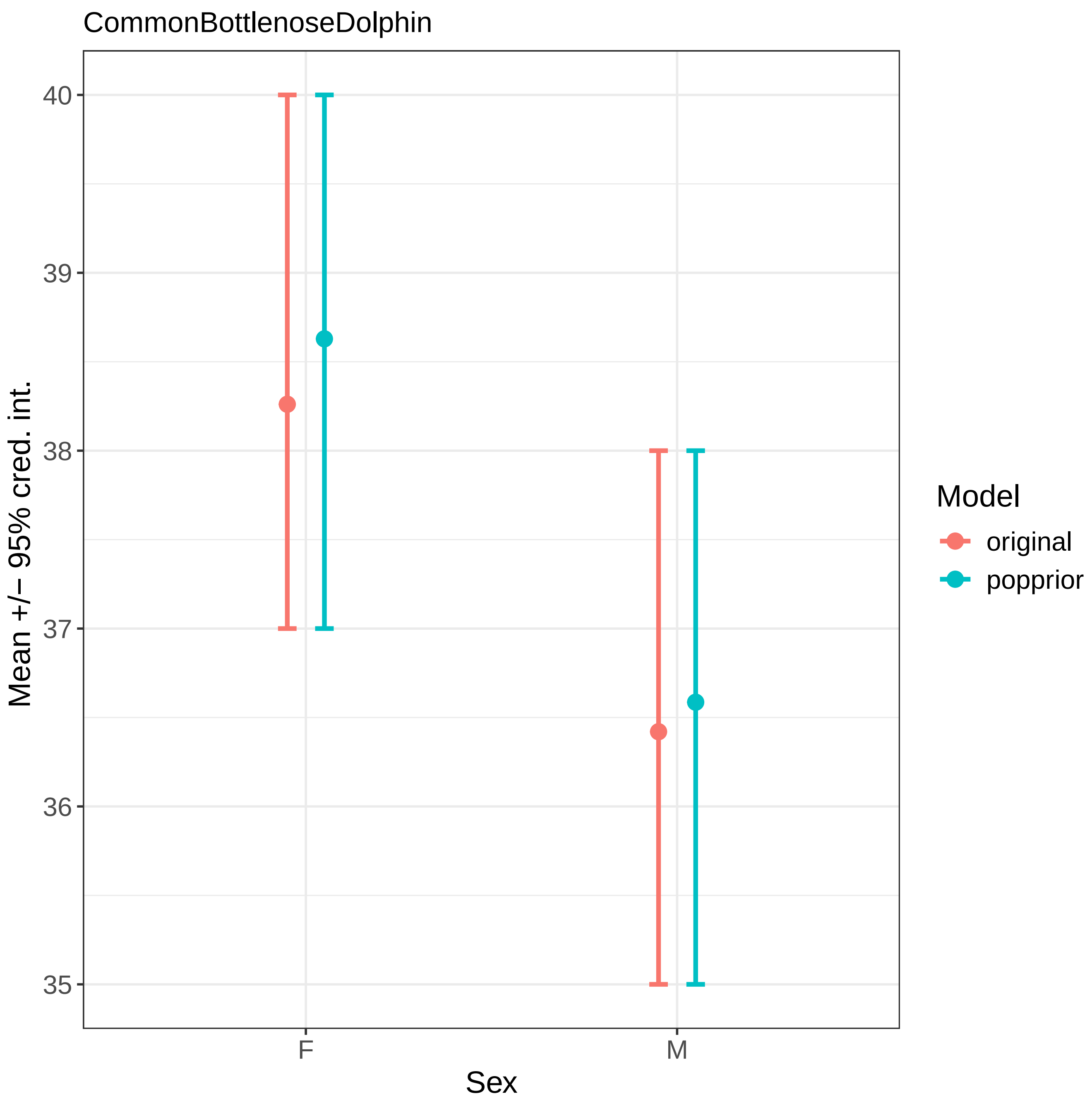

CommonBottlenoseDolphin
Mean +/− 95% cred. int.
40
39
38
37
36
35
F
M
Sex
Model
original
popprior

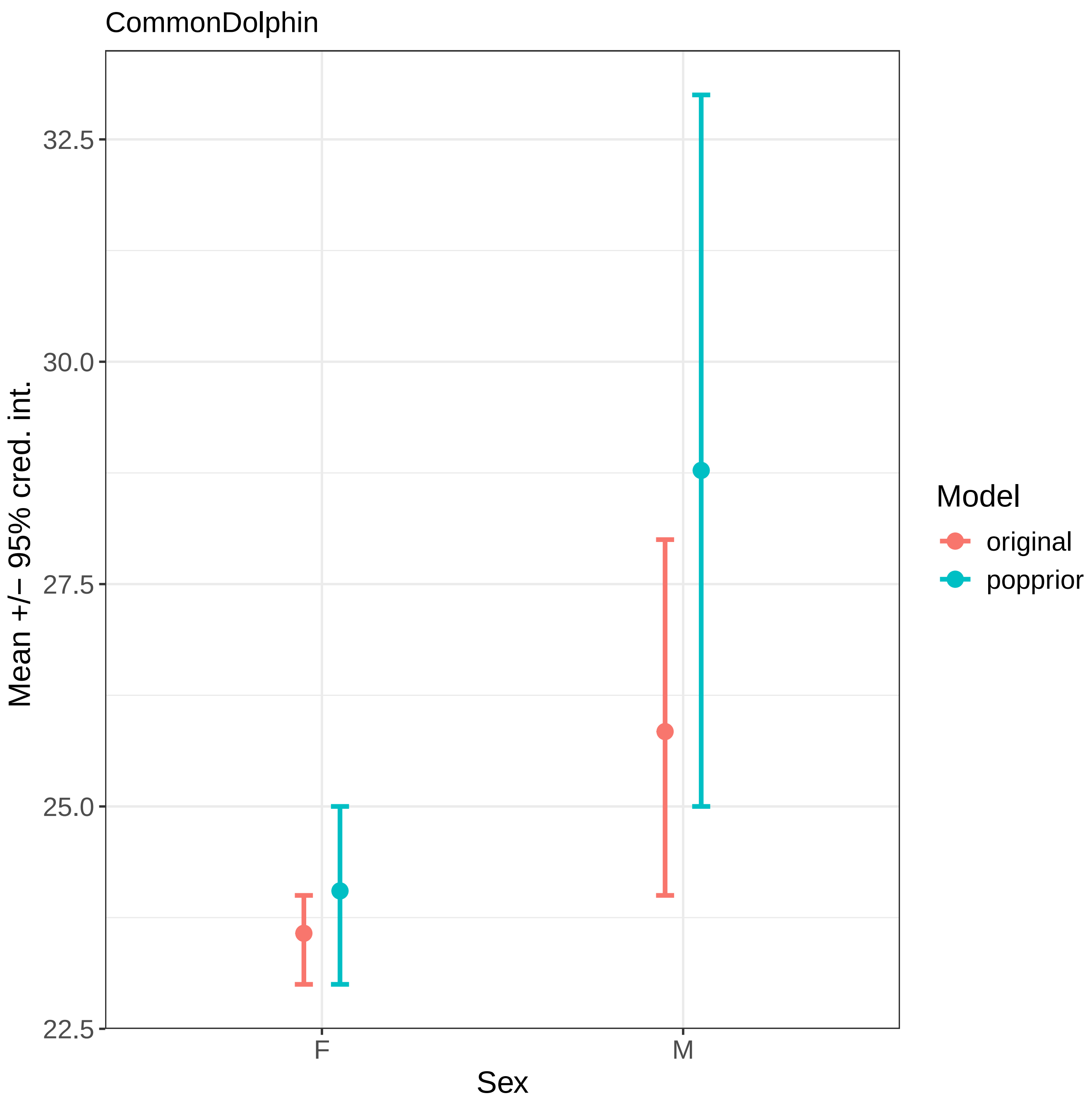

CommonDolphin
Mean +/− 95% cred. int.
32.5
30.0
27.5
25.0
22.5
F
M
Sex
Model
original
popprior

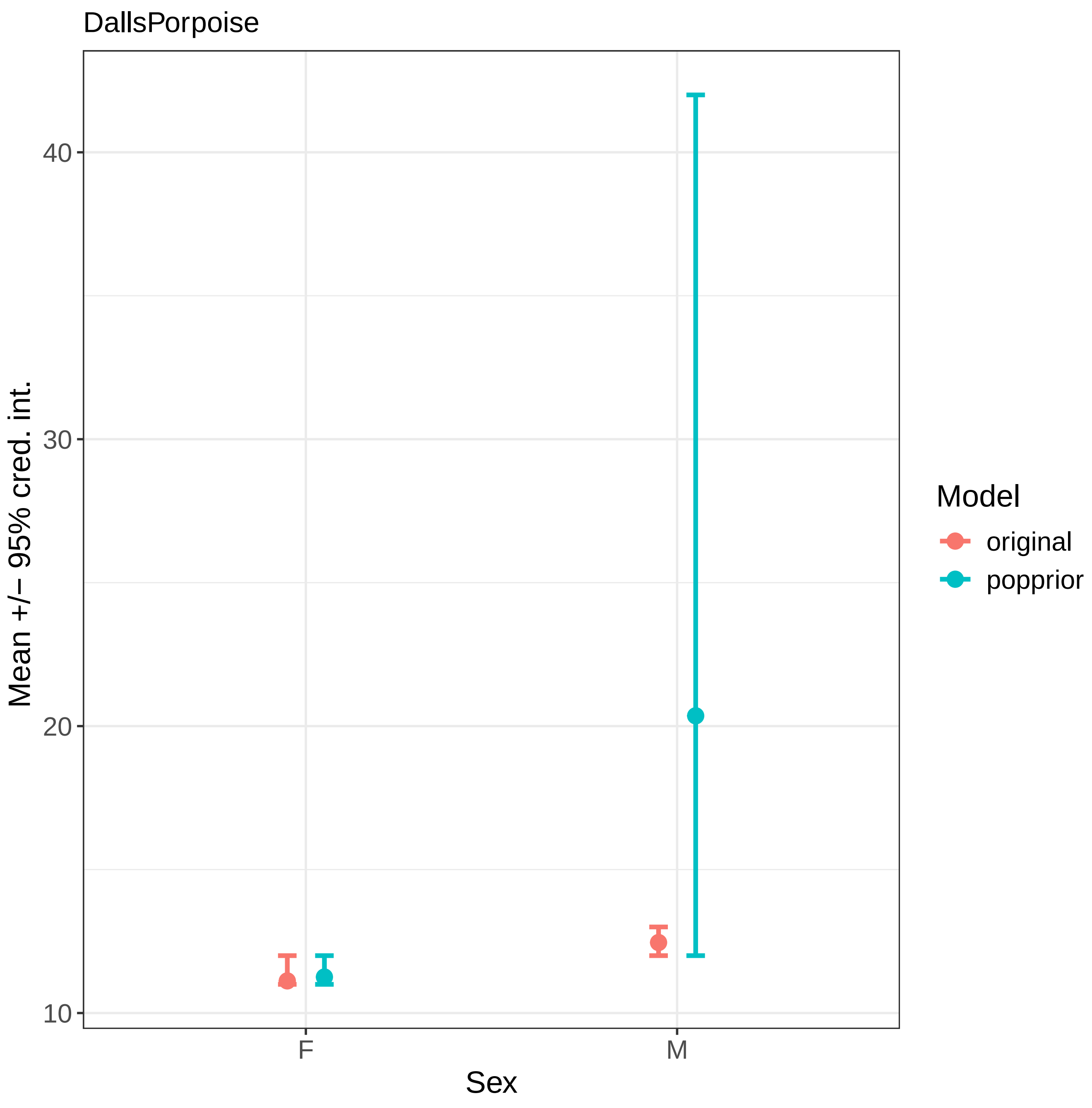
DallsPorpoise
Mean +/− 95% cred. int.
40
30
20
10
F
M
Sex
Model
original
popprior

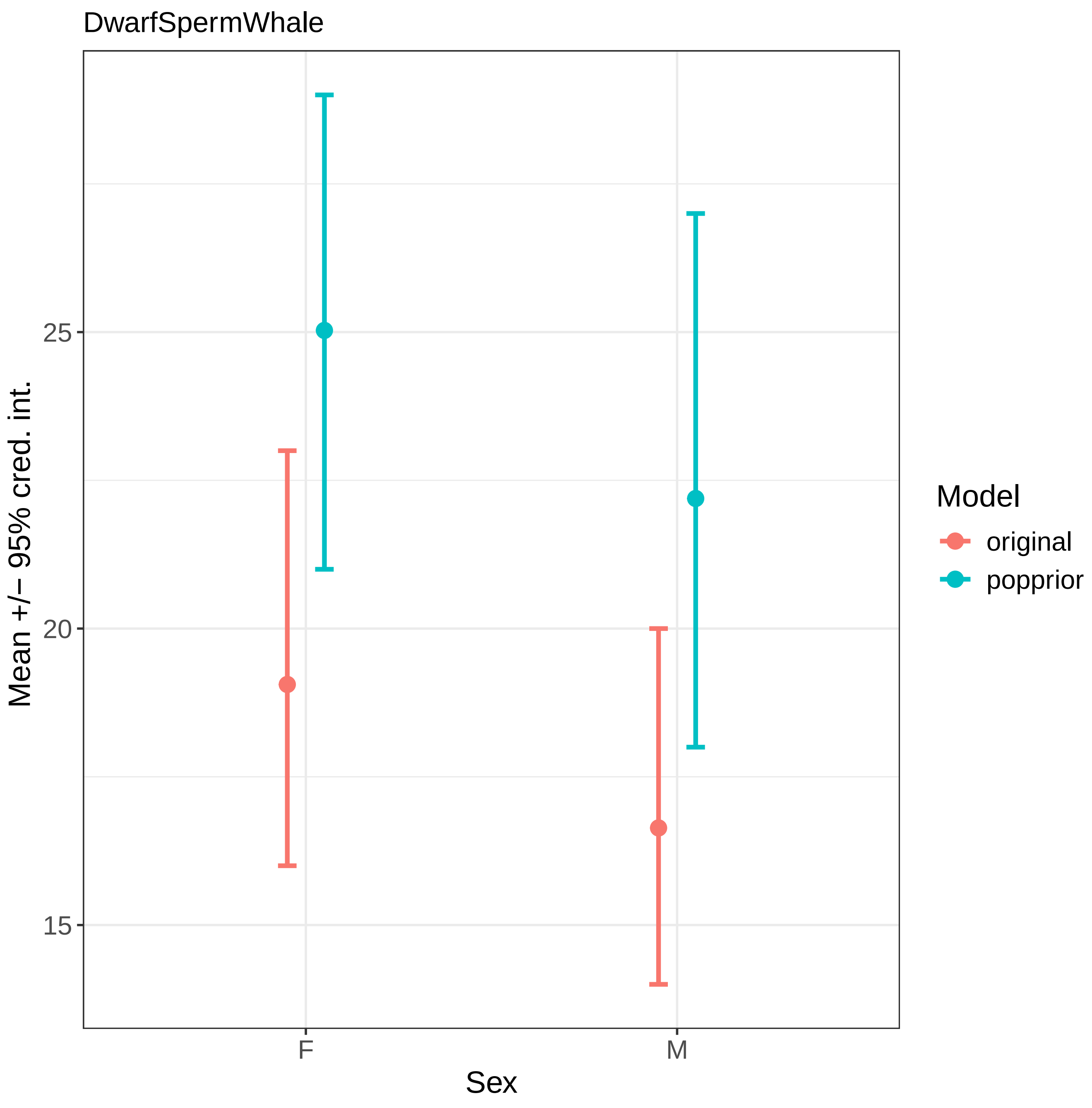
DwarfSpermWhale
Mean +/− 95% cred. int.
25
20
15
F
M
Sex
Model
original
popprior

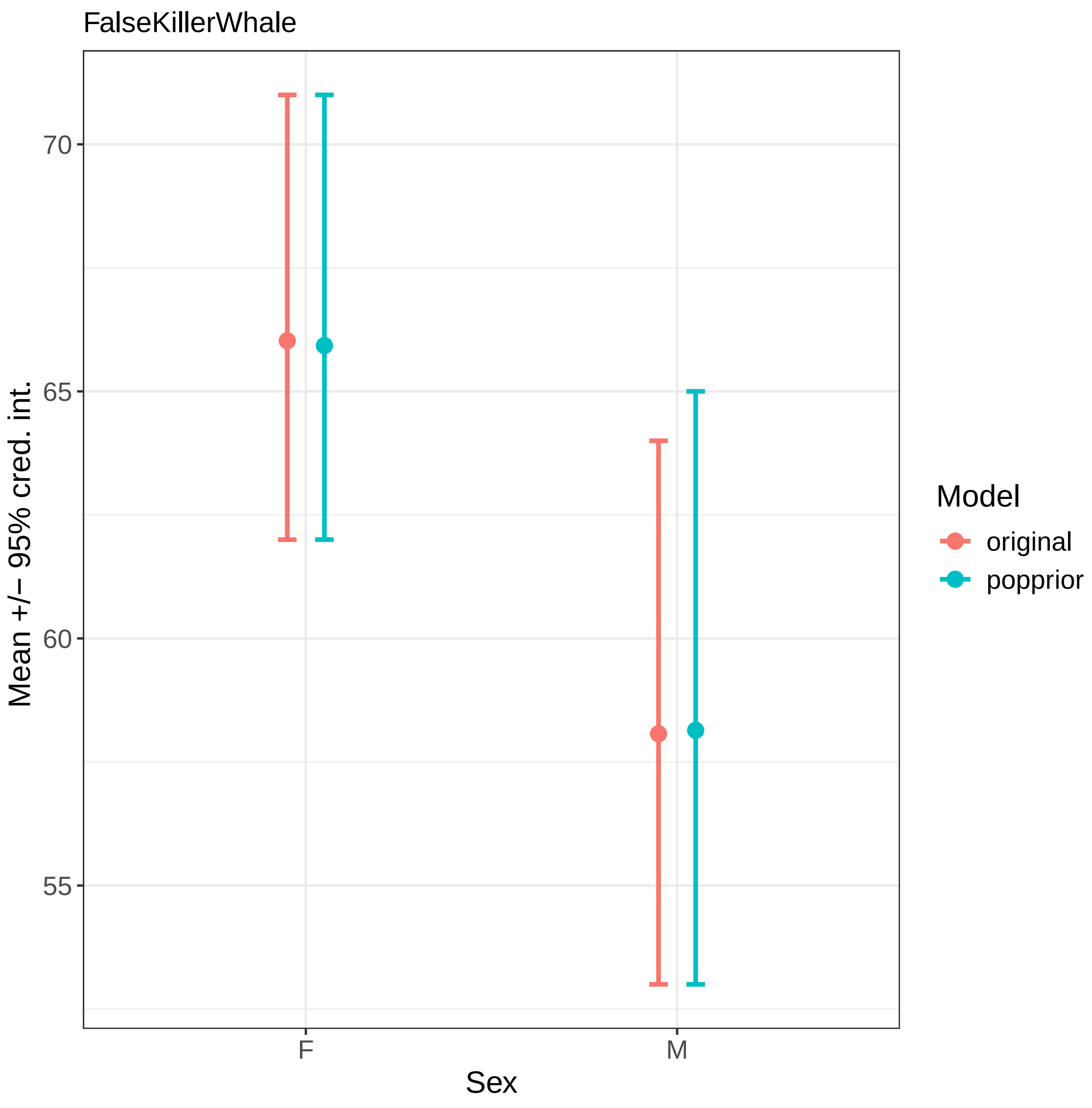

FalseKillerWhale
70
65
60
55
Mean +/− 95% cred. int.
F
M
Sex
Model
original
popprior

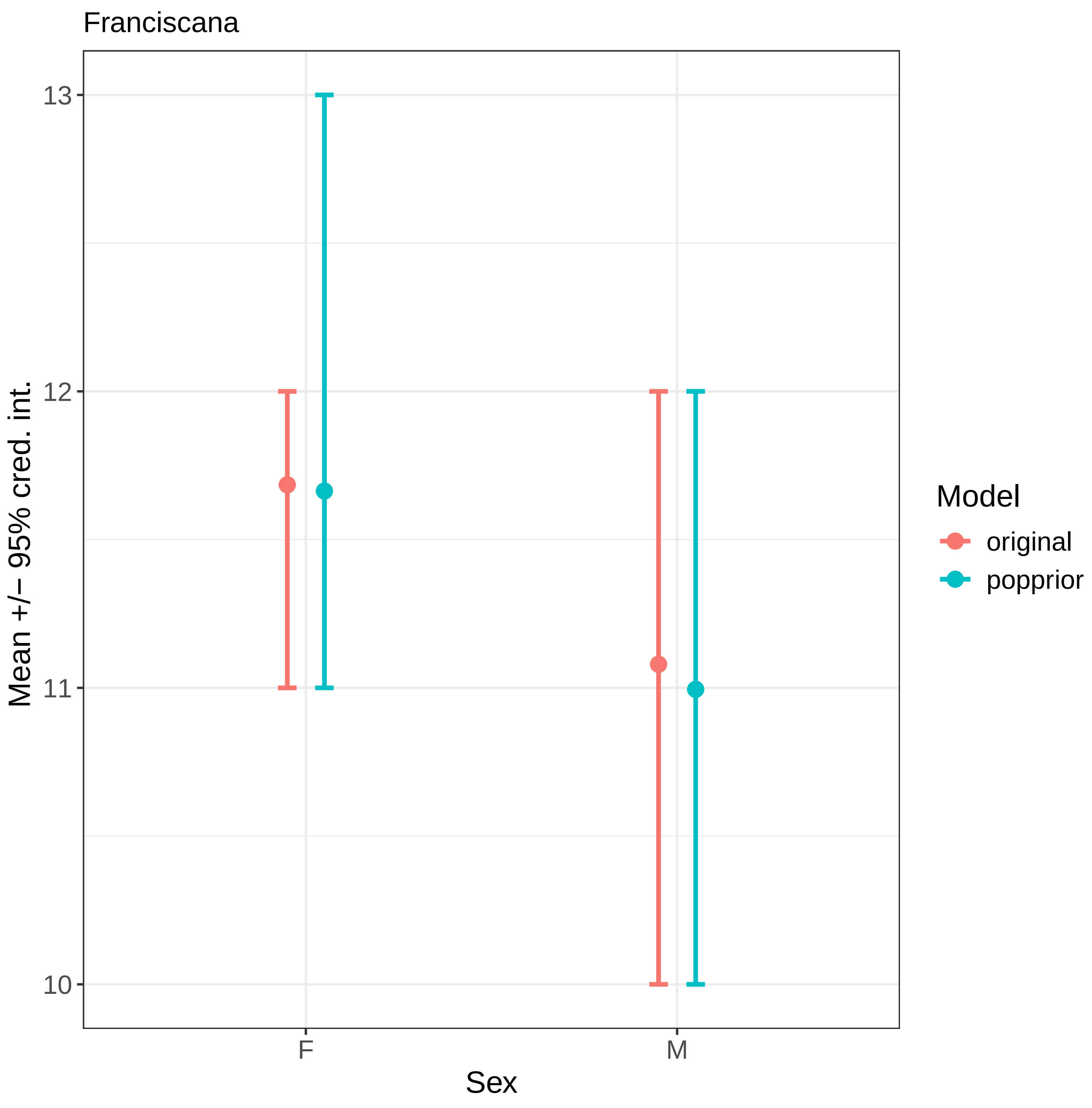

Franciscana
Mean +/− 95% cred. int.
13
12
11
10
F
M
Sex
Model
original
popprior

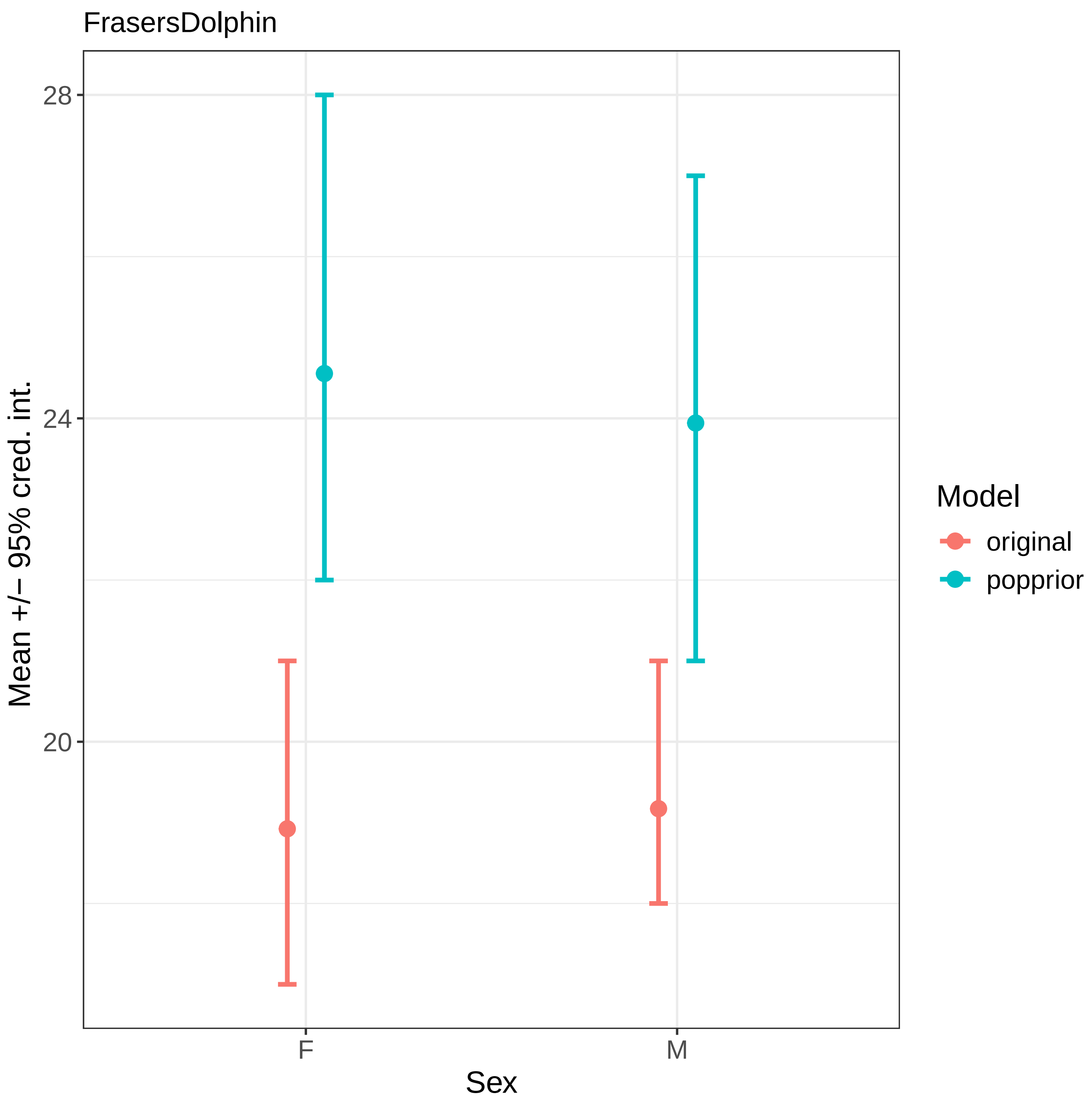
FrasersDolphin
28
24
20
Mean +/− 95% cred. int.
F
M
Sex
Model
original
popprior

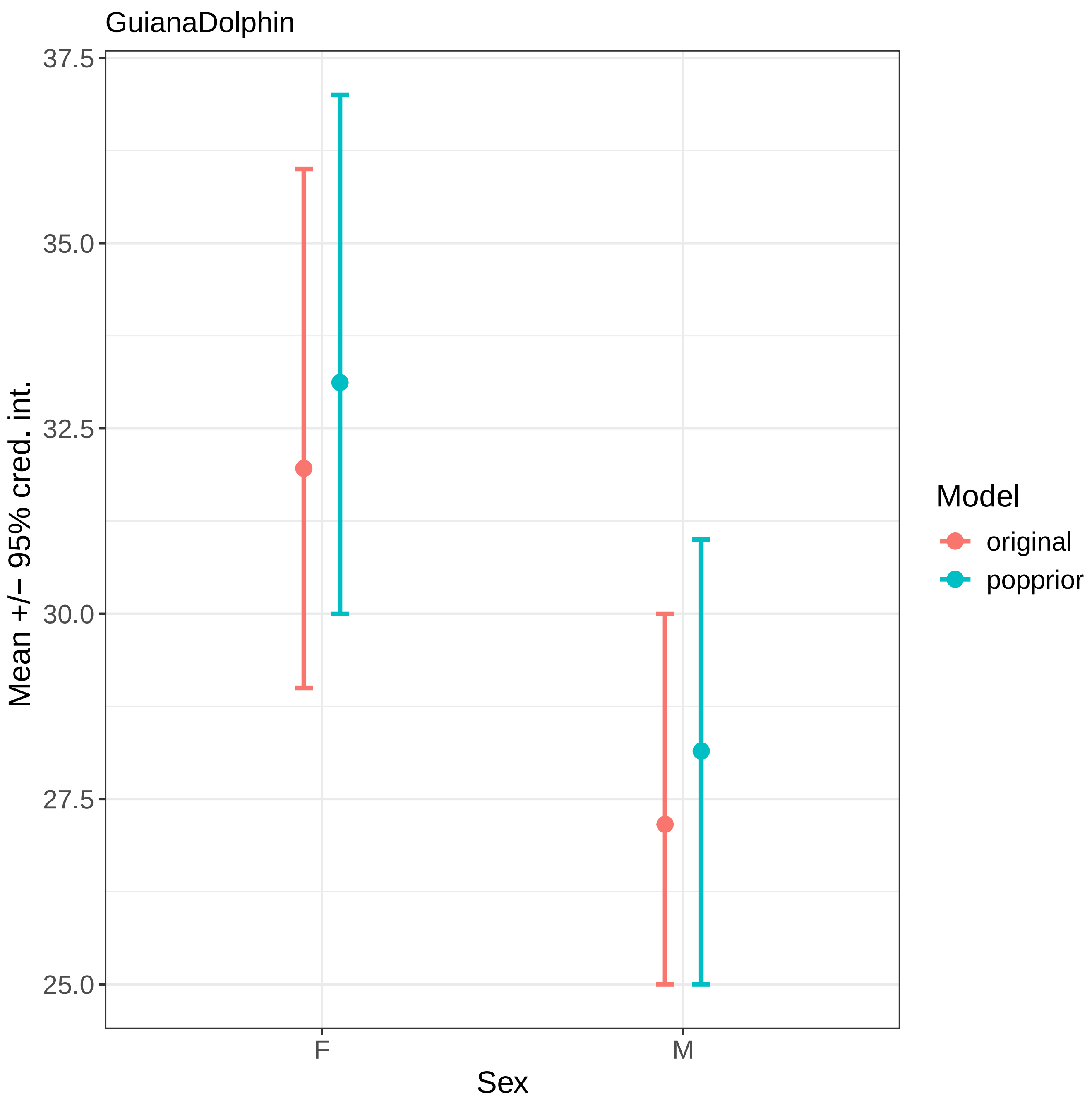
GuianaDolphin
37.5
35.0
32.5
30.0
27.5
25.0
Mean +/− 95% cred. int.
F
M
Sex
Model
original
popprior

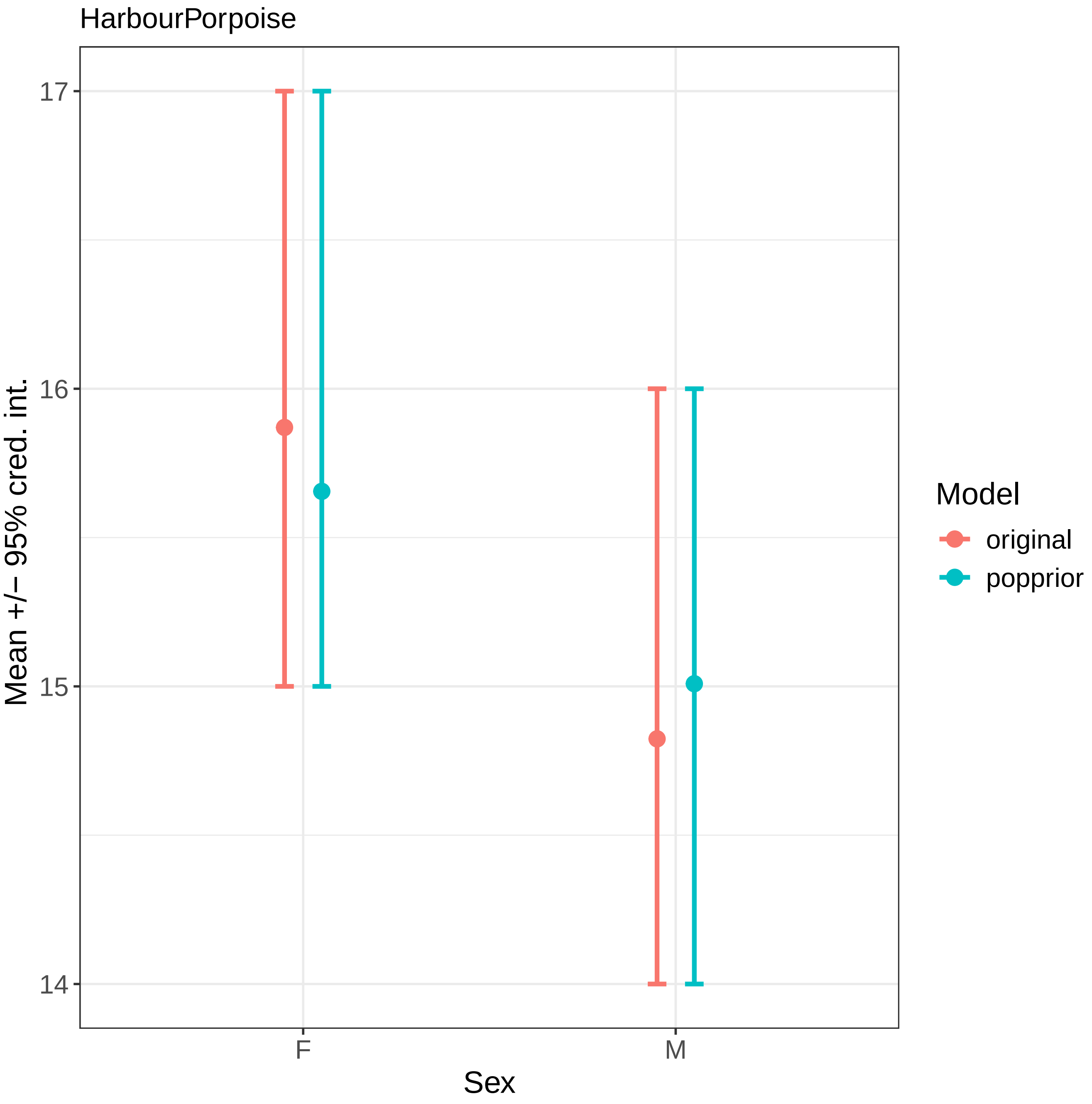

HarbourPorpoise
Mean +/− 95% cred. int.
17
16
15
14
F
M
Sex
Model
original
popprior

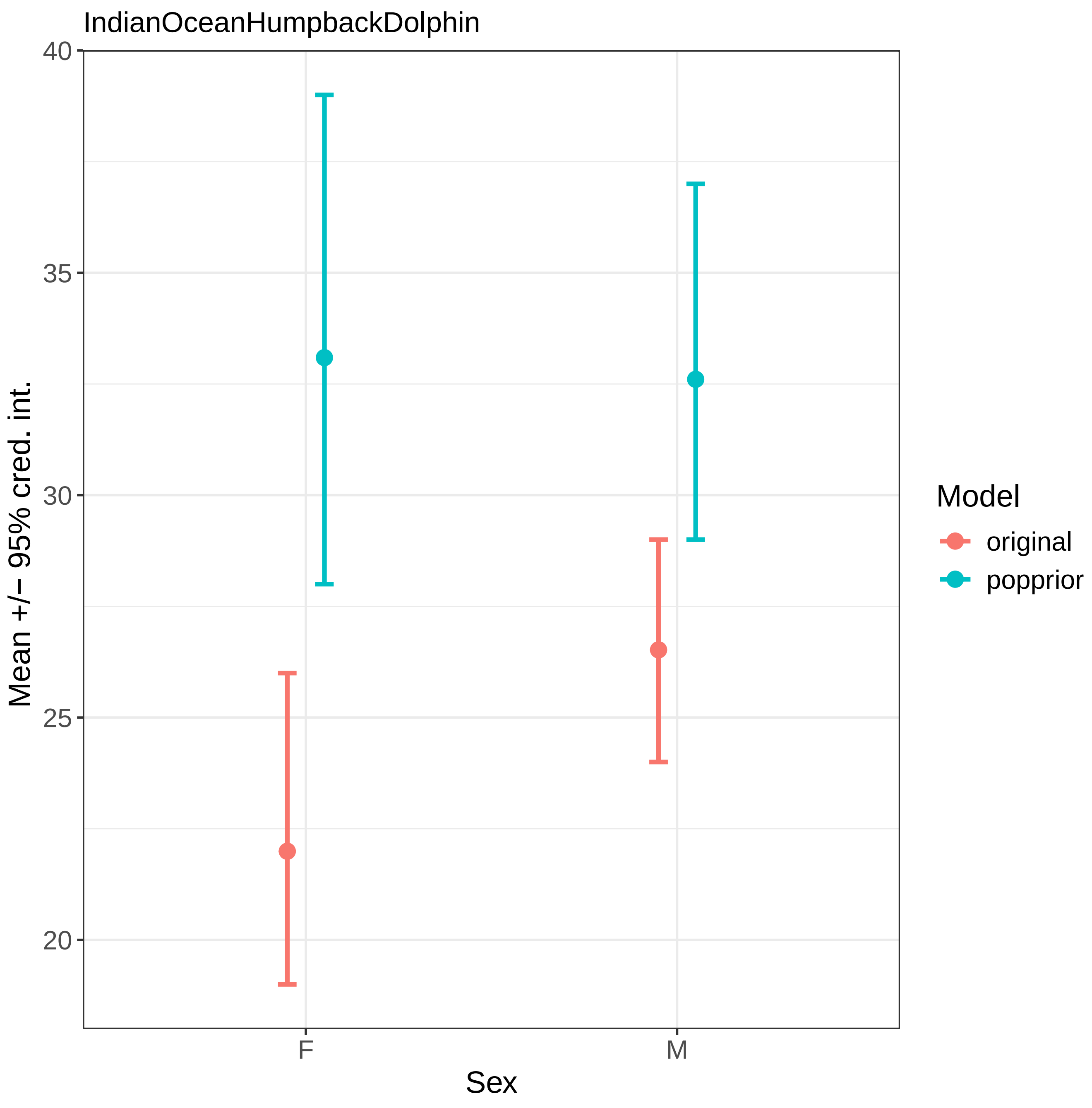

IndianOceanHumpbackDolphin
Mean +/− 95% cred. int.
40
35
30
25
20
F
M
Sex
Model
original
popprior

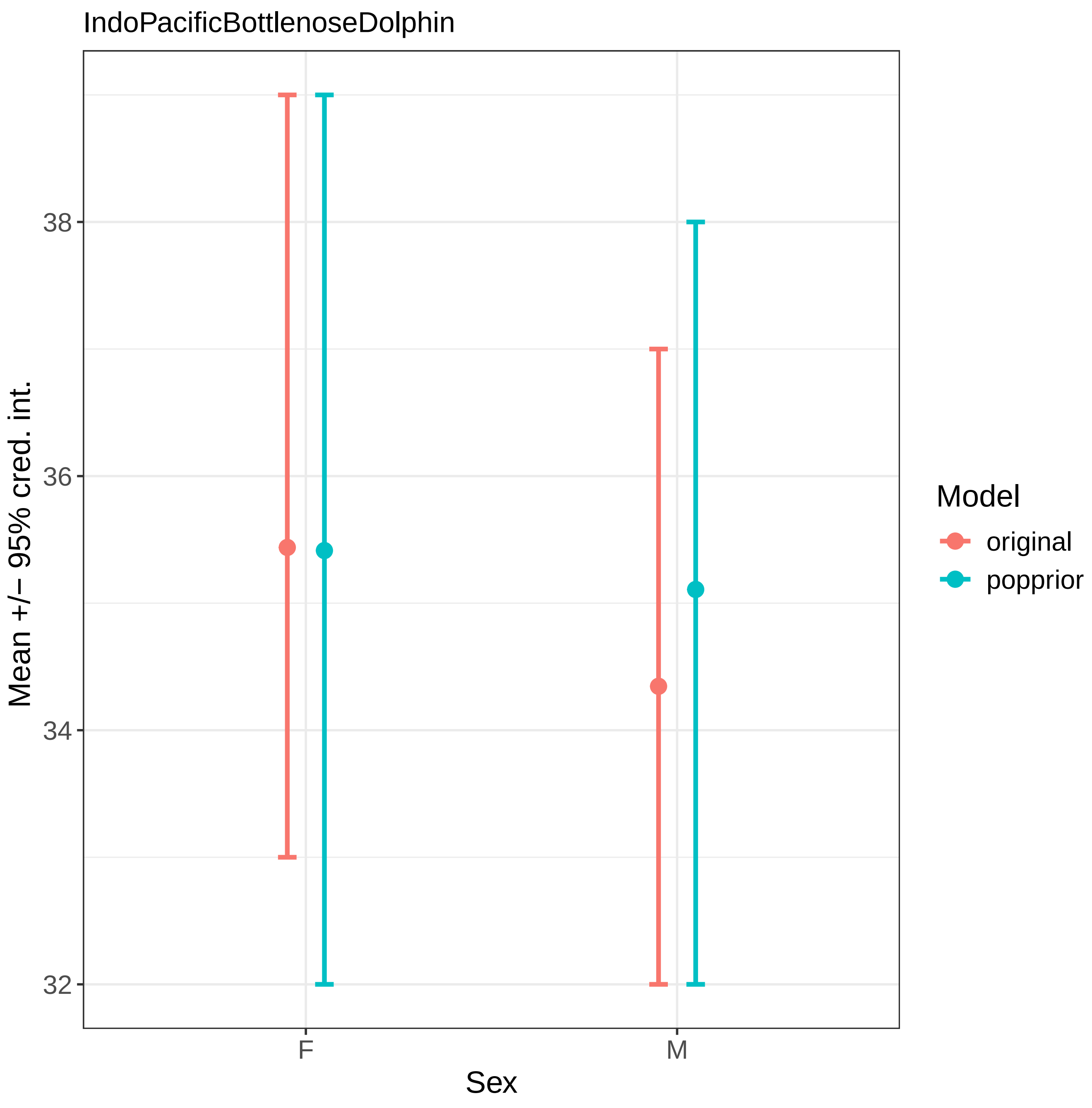

IndoPacificBottlenoseDolphin
Mean +/− 95% cred. int.
38
36
34
32
F
M
Sex
Model
original
popprior

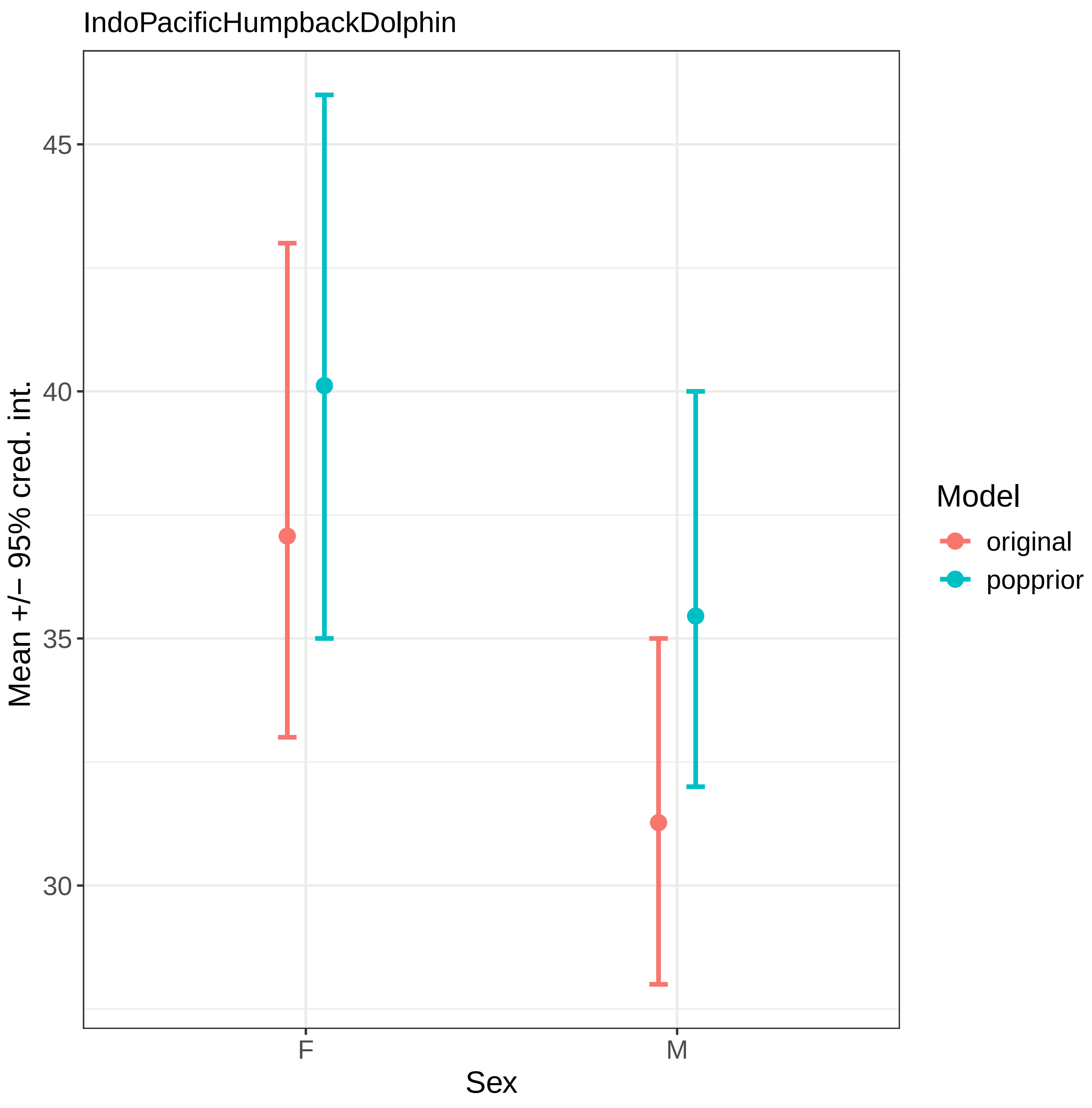

IndoPacificHumpbackDolphin
Mean +/− 95% cred. int.
45
40
35
30
F
M
Sex
Model
original
popprior

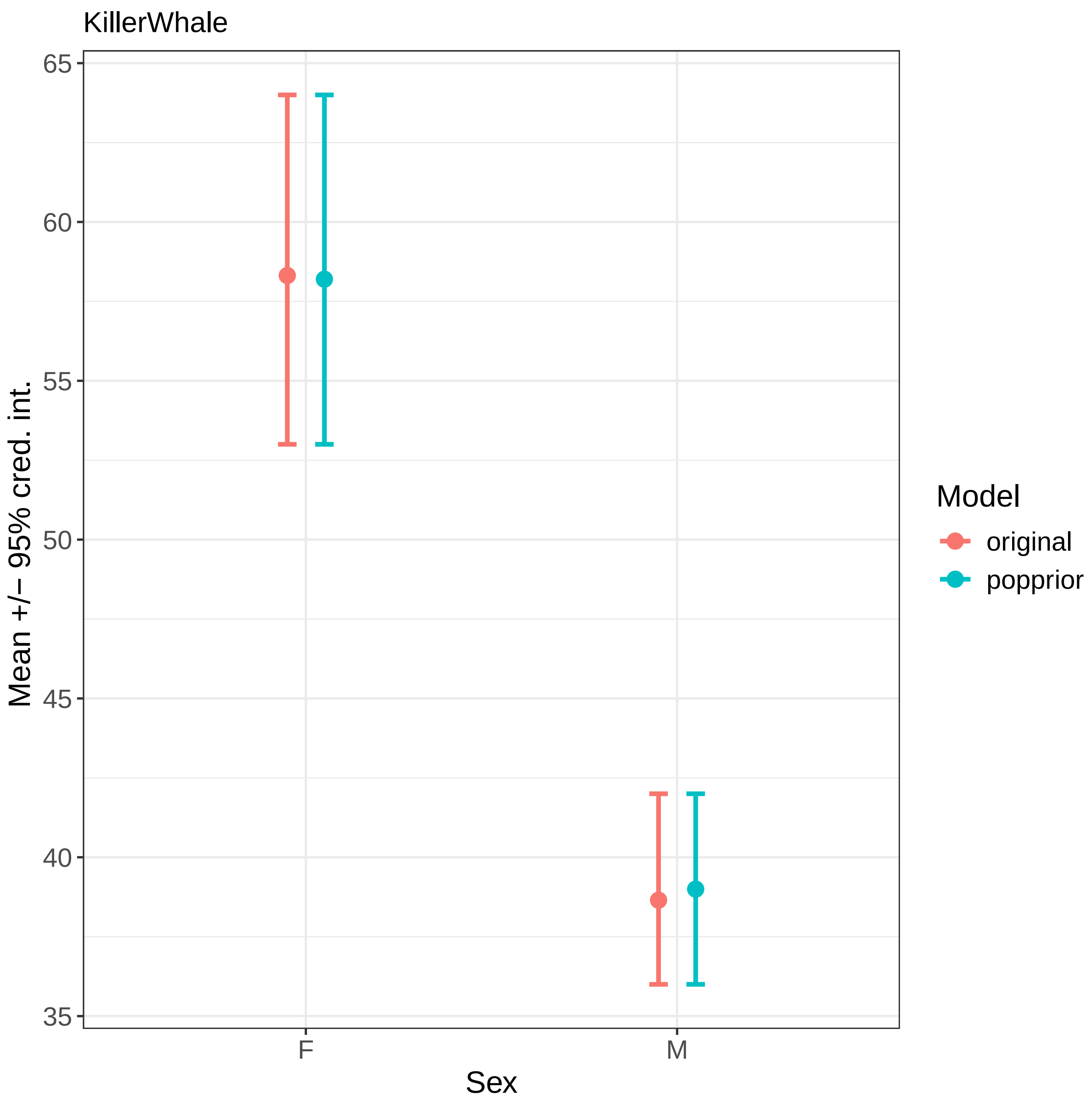
KillerWhale
65
60
55
50
45
40
35
Mean +/− 95% cred. int.
F
M
Sex
Model
original
popprior

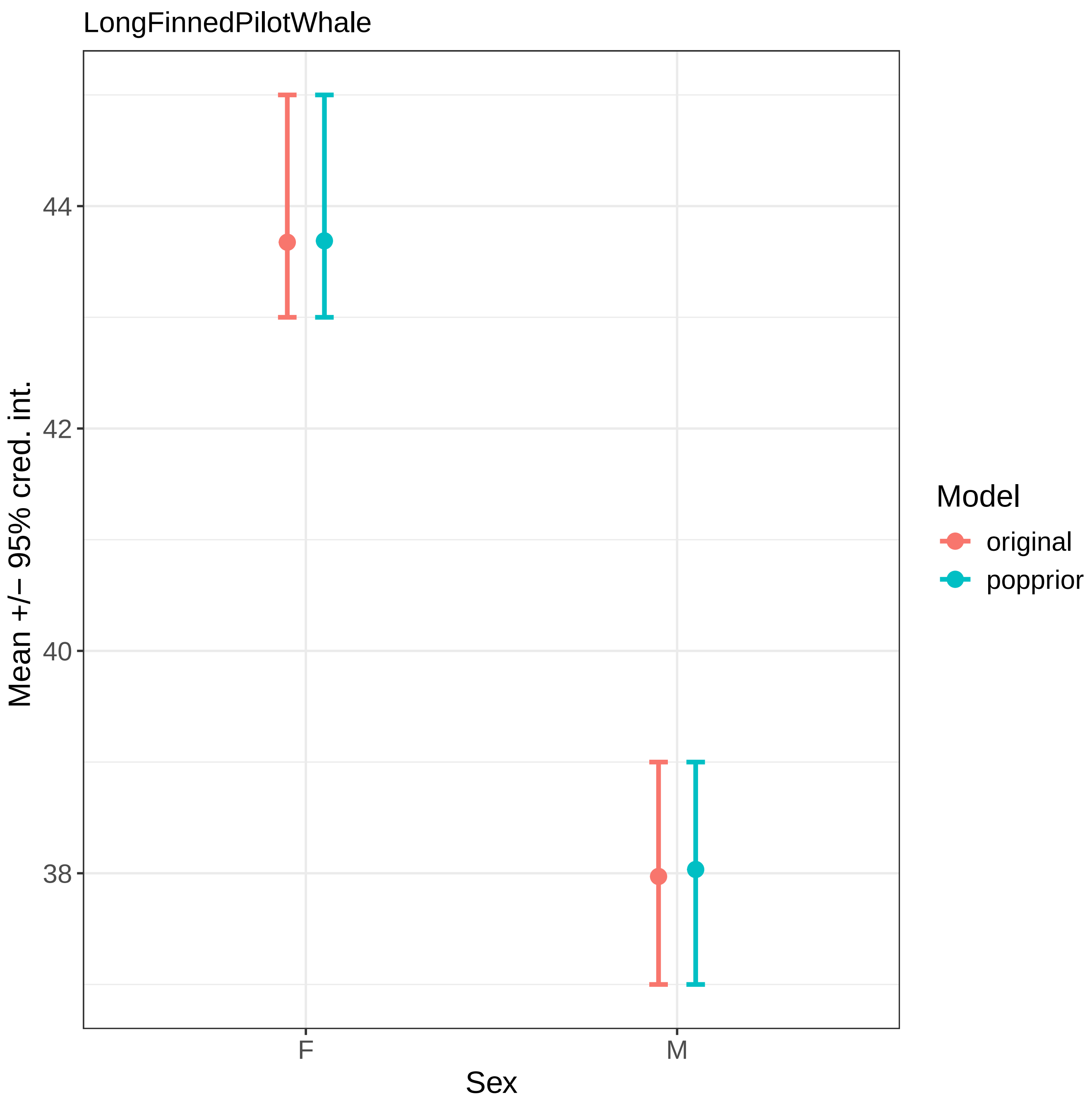

LongFinnedPilotWhale
Mean +/− 95% cred. int.
44
42
40
38
F
M
Sex
Model
original
popprior

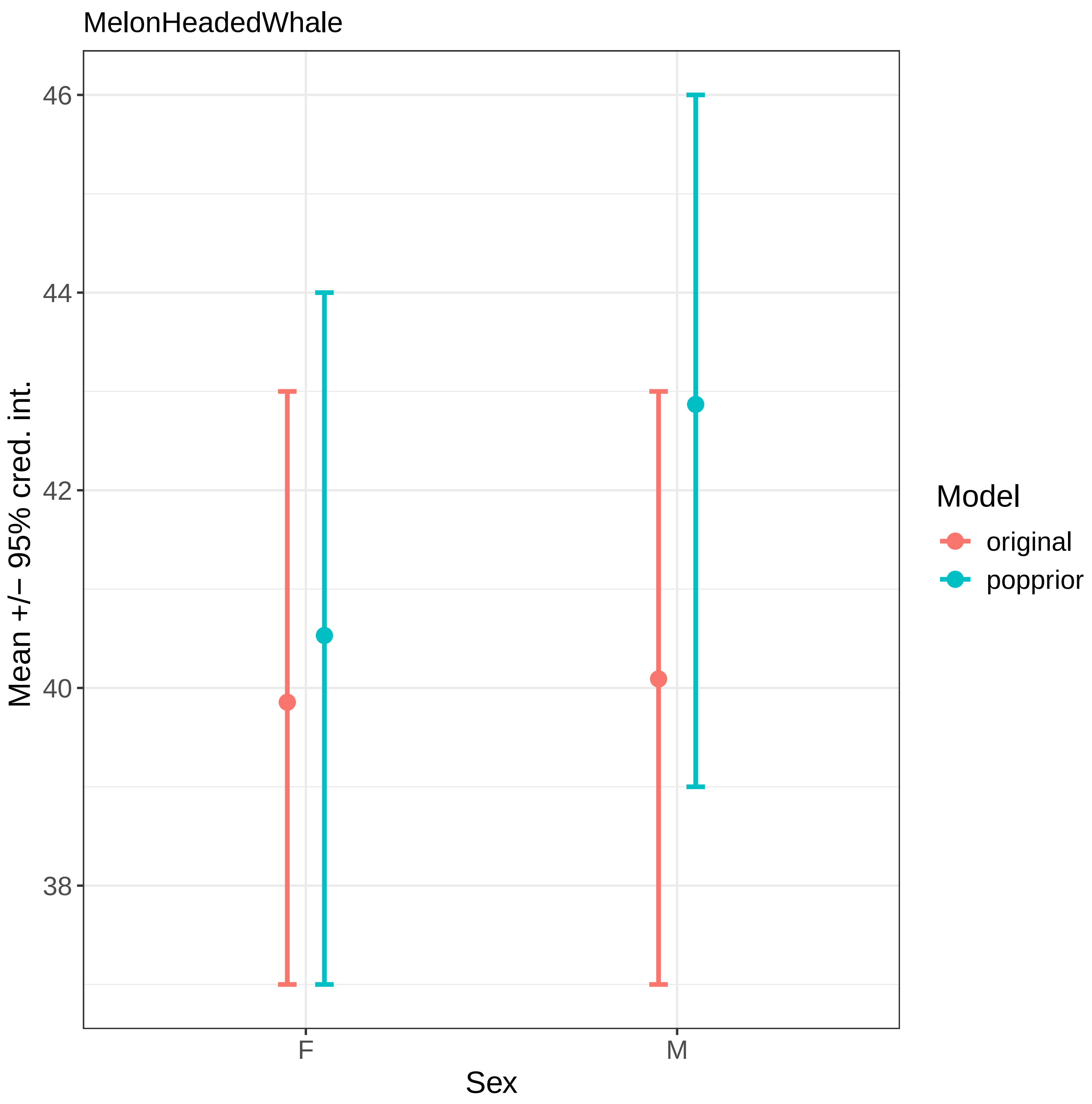
MelonHeadedWhale
46
44
42
40
38
Mean +/− 95% cred. int.
F
M
Sex
Model
original
popprior

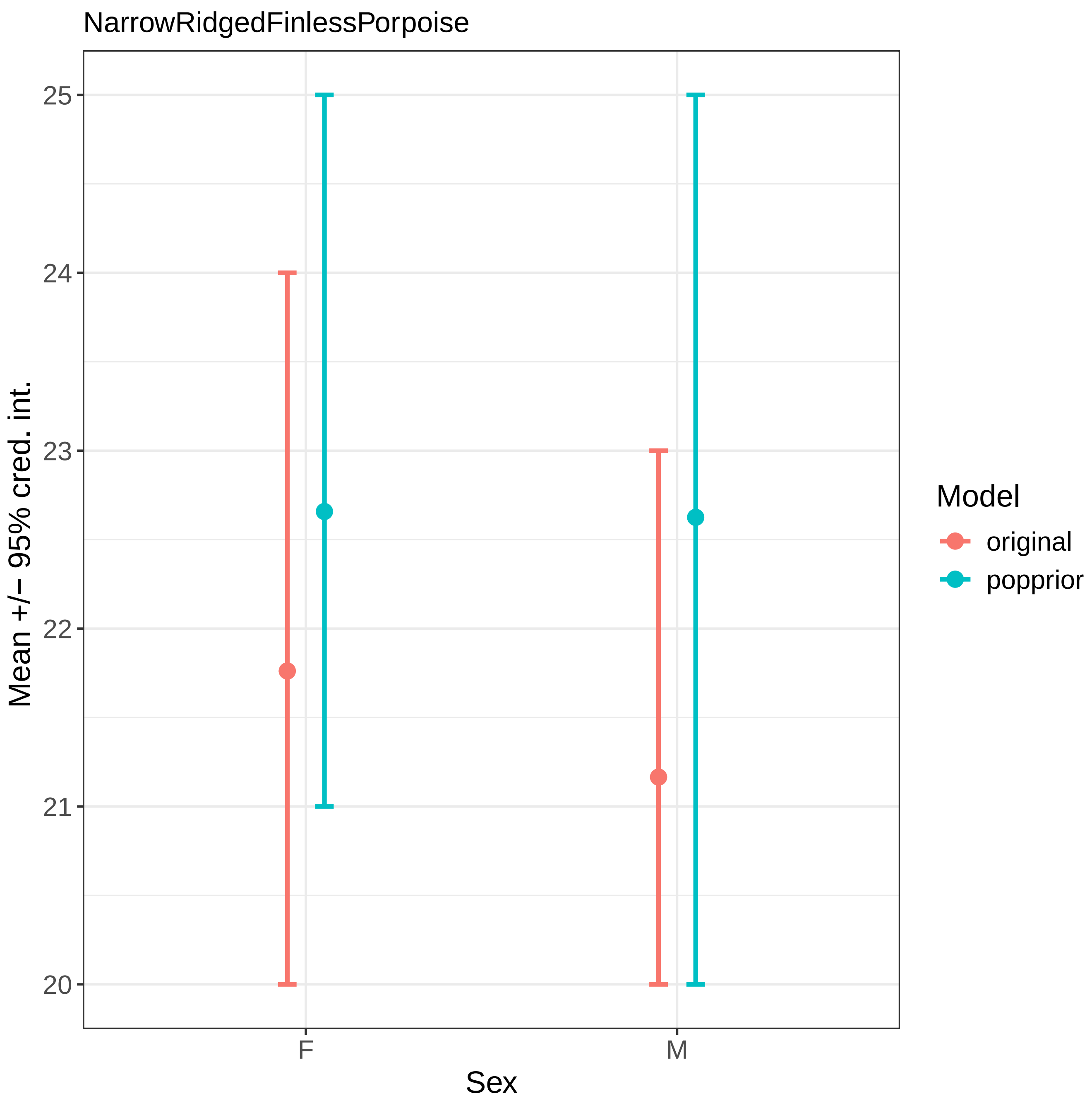
NarrowRidgedFinlessPorpoise
Mean +/− 95% cred. int.
25
24
23
22
21
20
F
M
Sex
Model
original
popprior

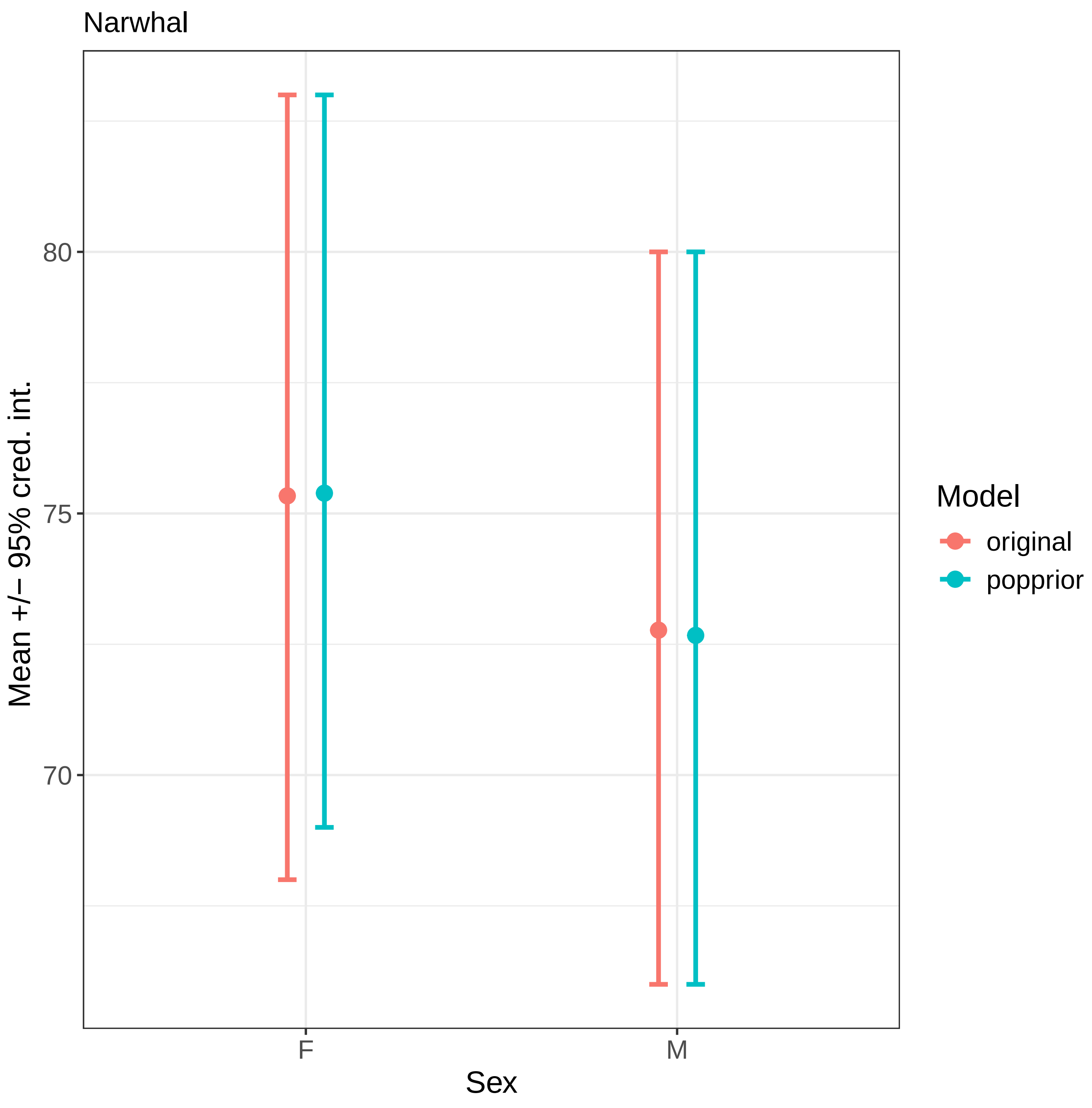
Narwhal
Mean +/− 95% cred. int.
80
75
70
F
M
Sex
Model
original
popprior

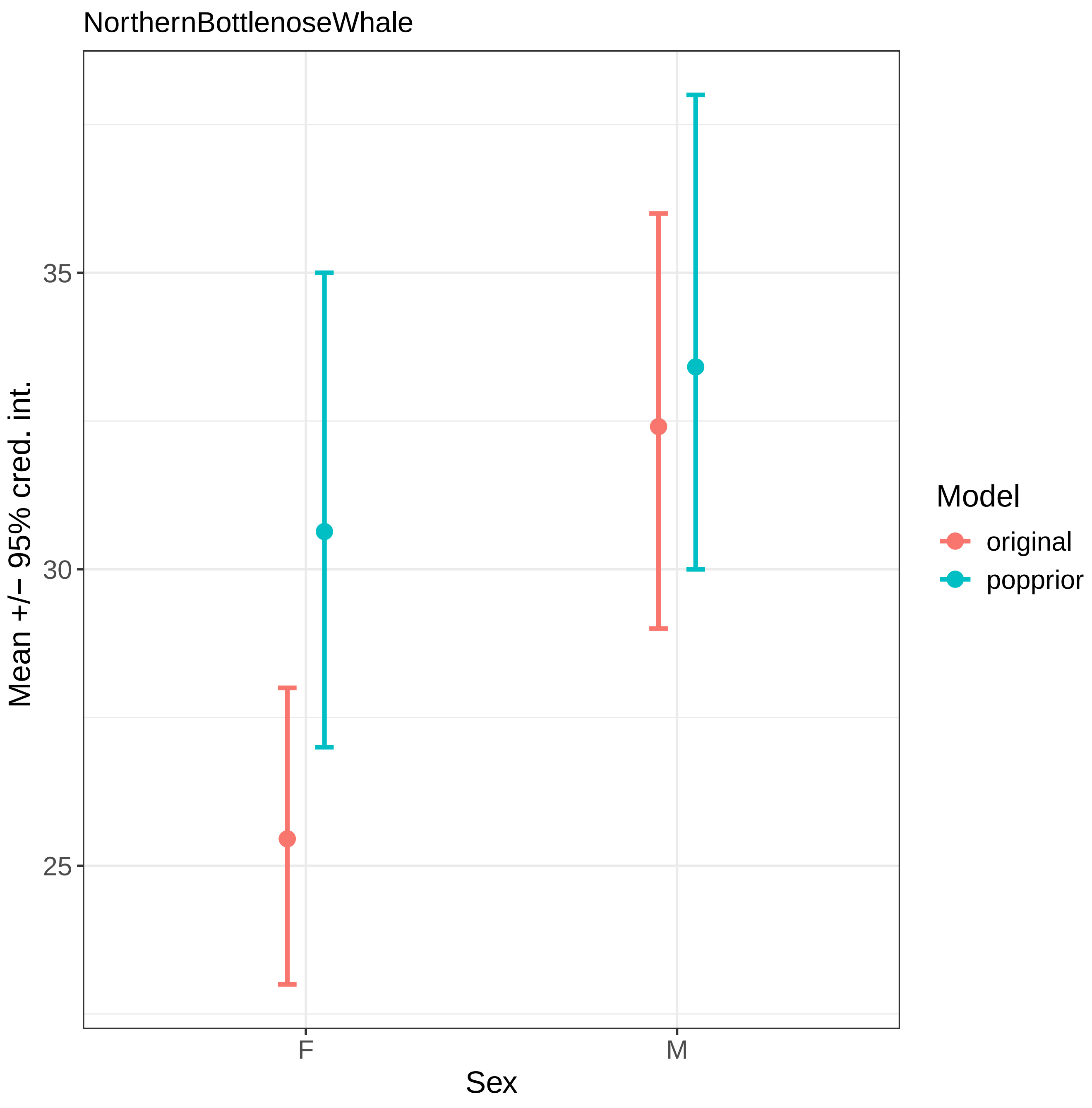

NorthernBottlenoseWhale
35
30
25
Mean +/− 95% cred. int.
F
M
Sex
Model
original
popprior

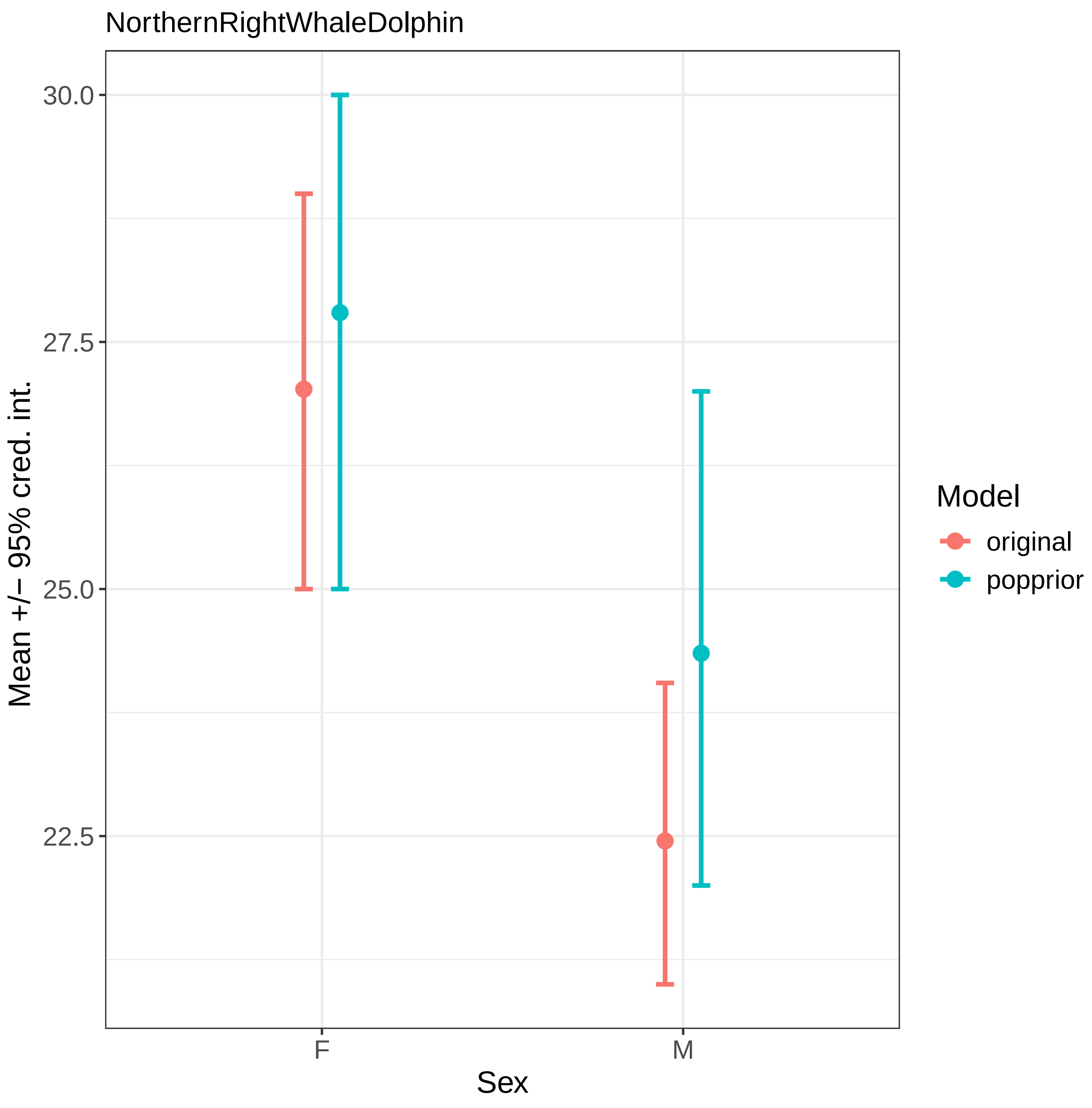

NorthernRightWhaleDolphin
30.0
27.5
25.0
22.5
Mean +/− 95% cred. int.
F
M
Sex
Model
original
popprior

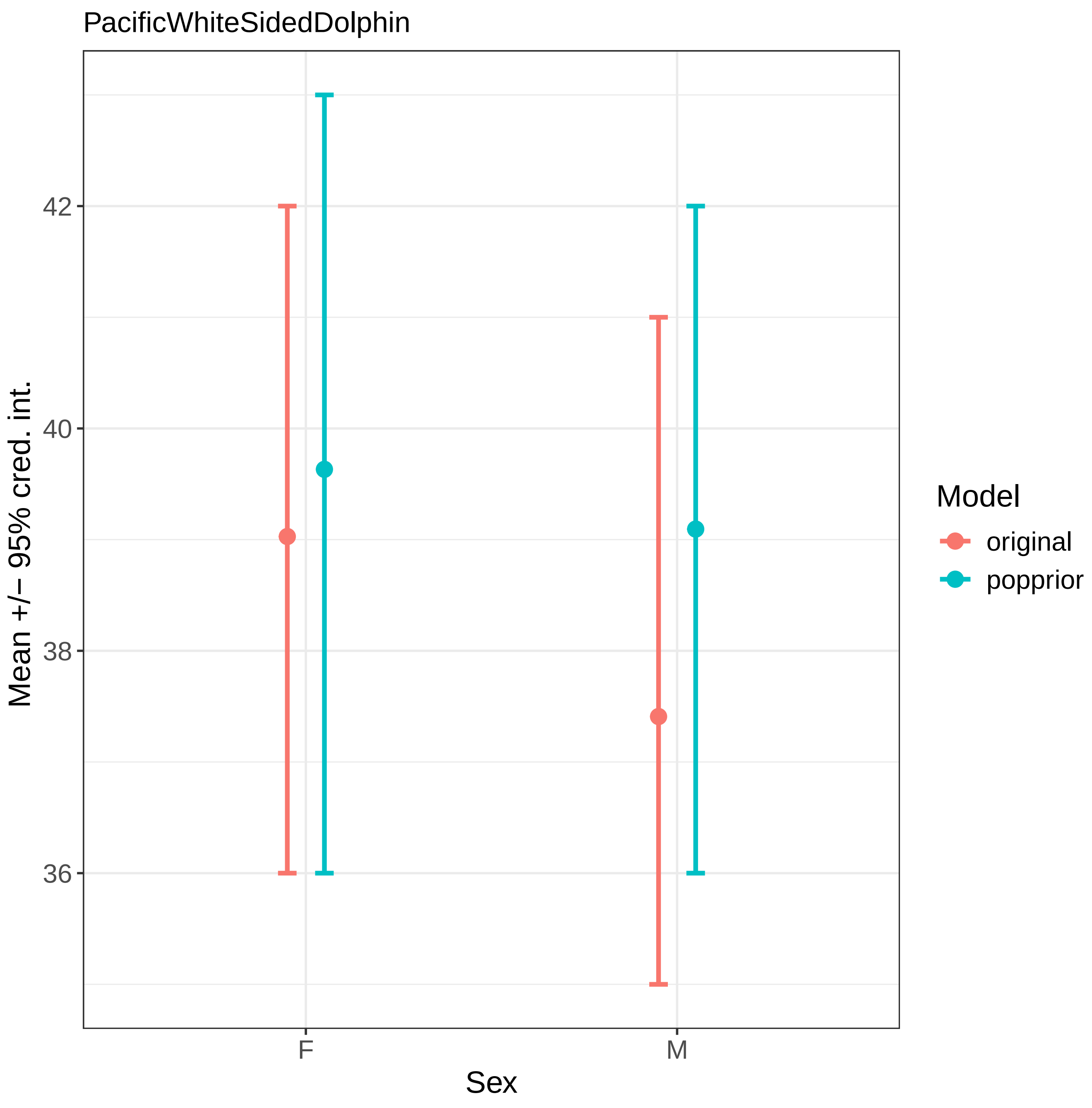

PacificWhiteSidedDolphin
Mean +/− 95% cred. int.
42
40
38
36
F
M
Sex
Model
original
popprior

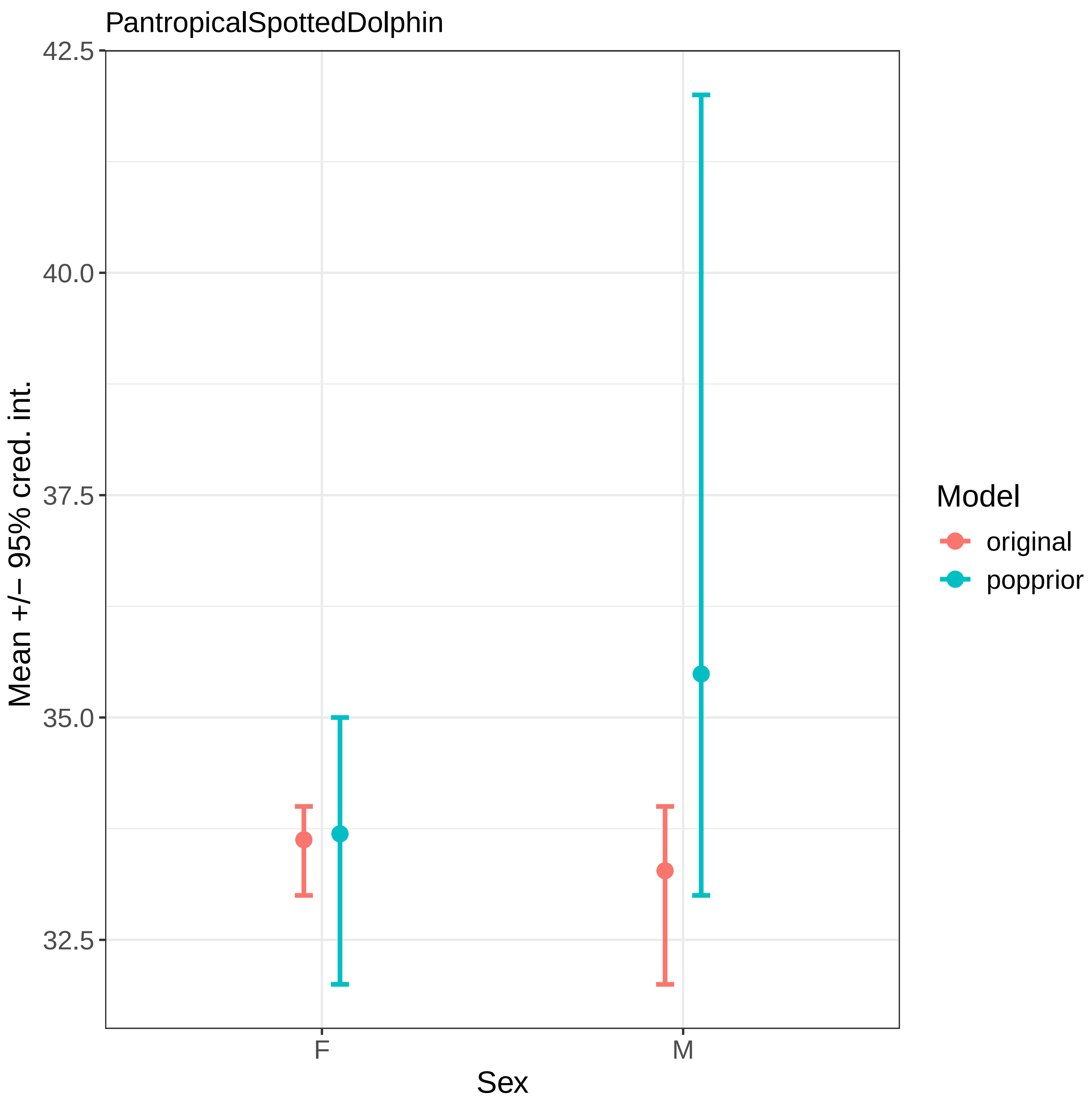
PantropicalSpottedDolphin
42.5
40.0
37.5
35.0
32.5
Mean +/− 95% cred. int.
F
M
Sex
Model
original
popprior

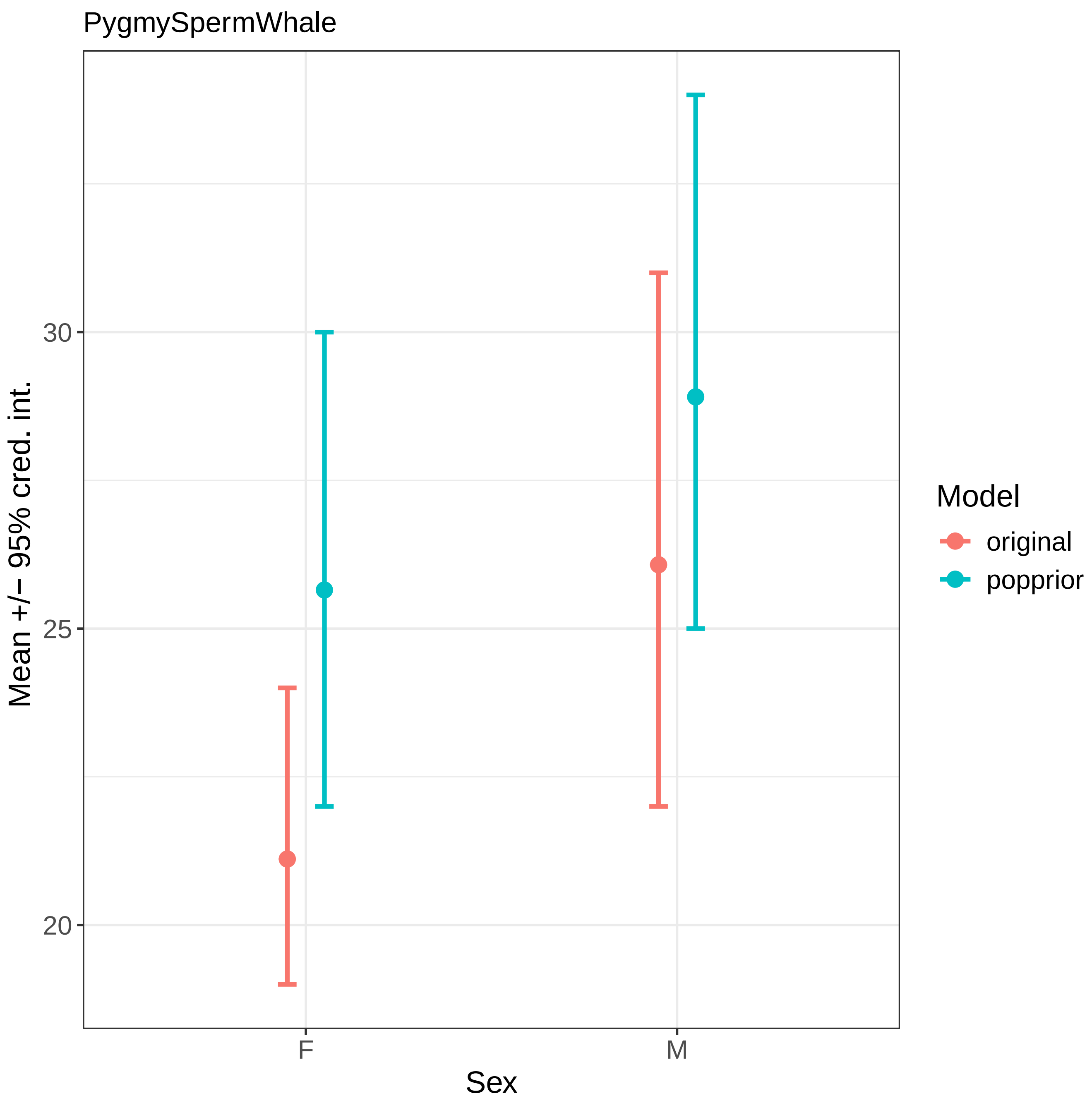
PygmySpermWhale
Mean +/− 95% cred. int.
30
25
20
F
M
Sex
Model
original
popprior

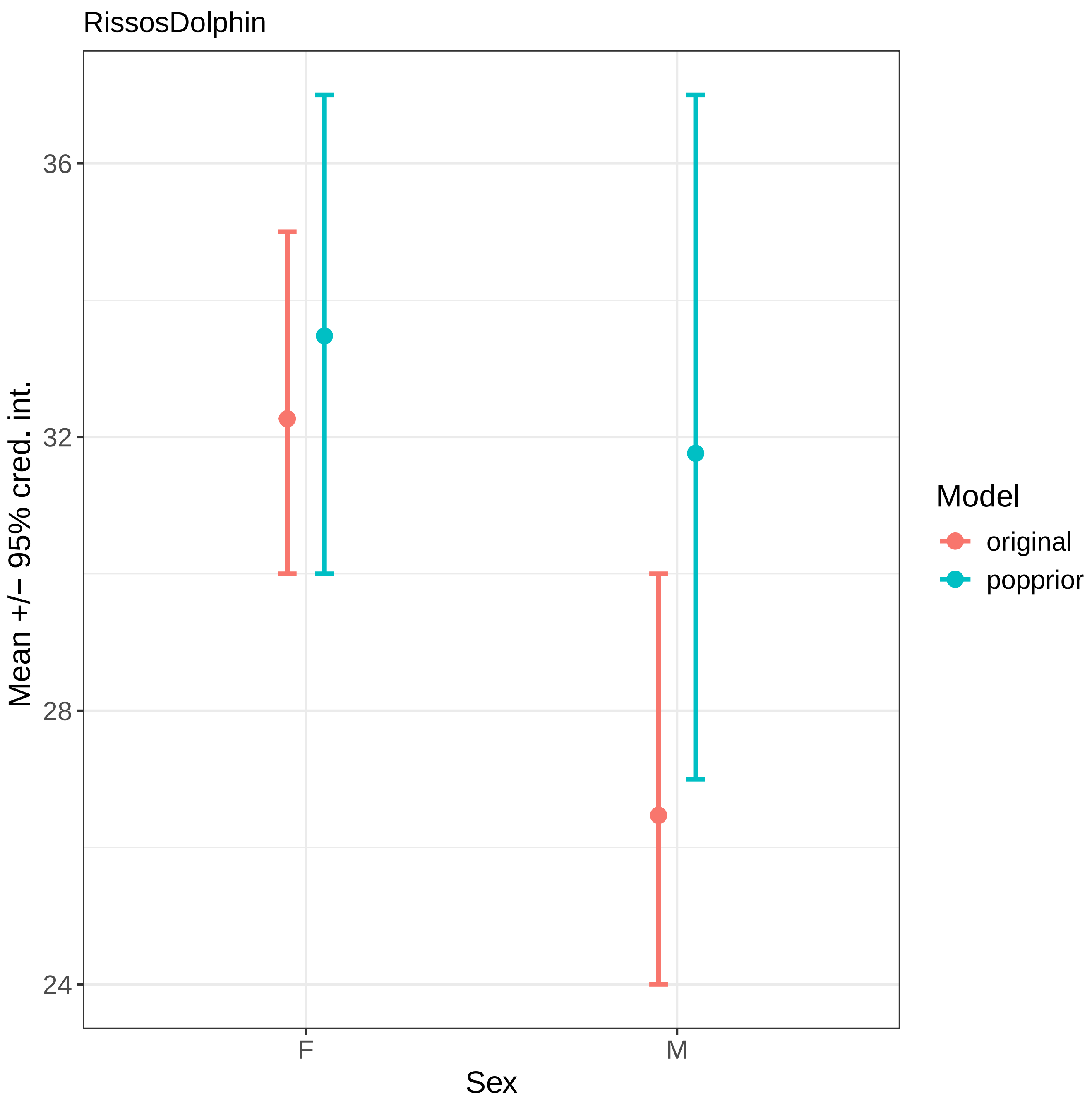

RissosDolphin
Mean +/− 95% cred. int.
36
32
28
24
F
M
Sex
Model
original
popprior

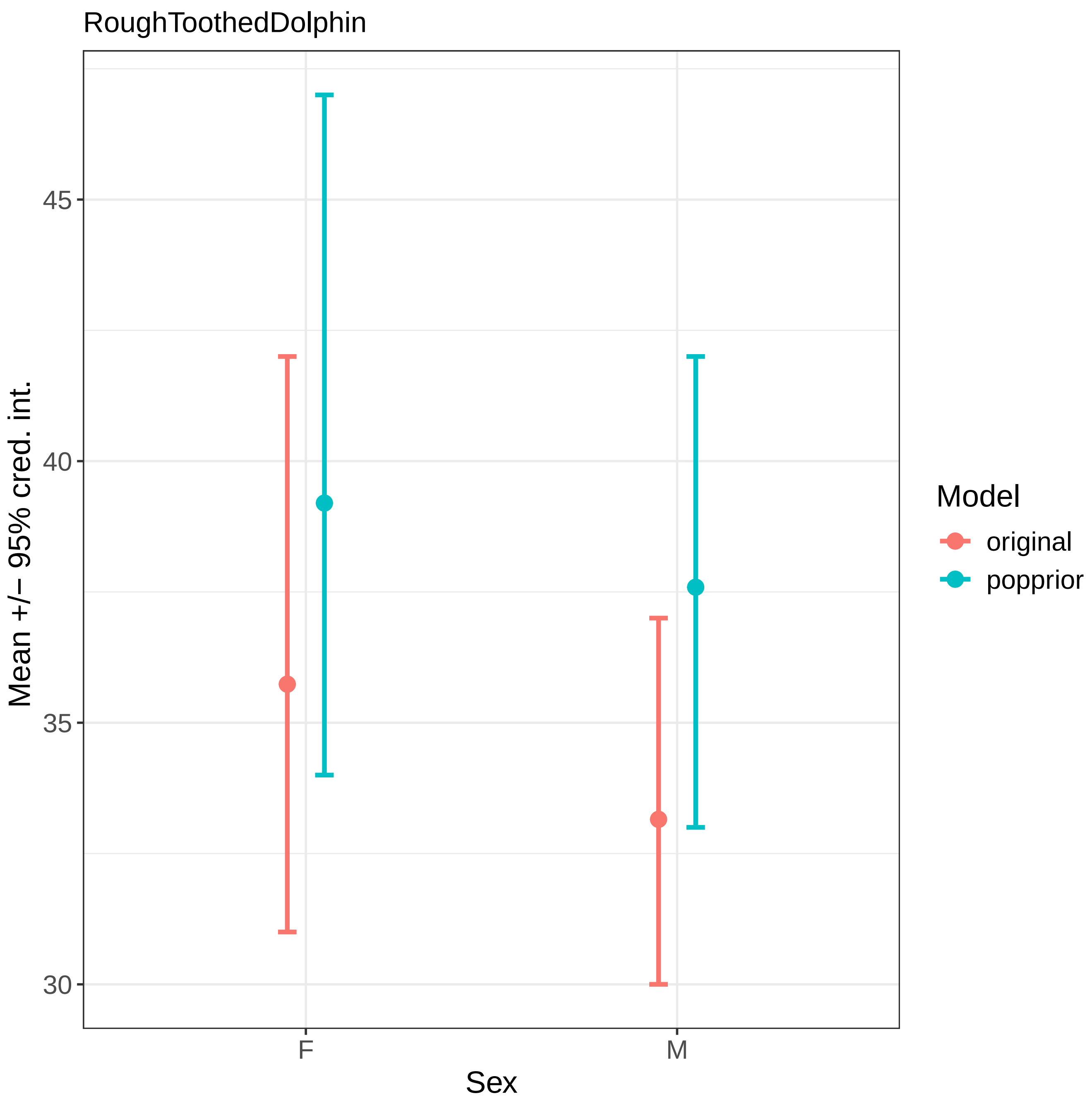
RoughToothedDolphin
Mean +/− 95% cred. int.
45
40
35
30
F
M
Sex
Model
original
popprior

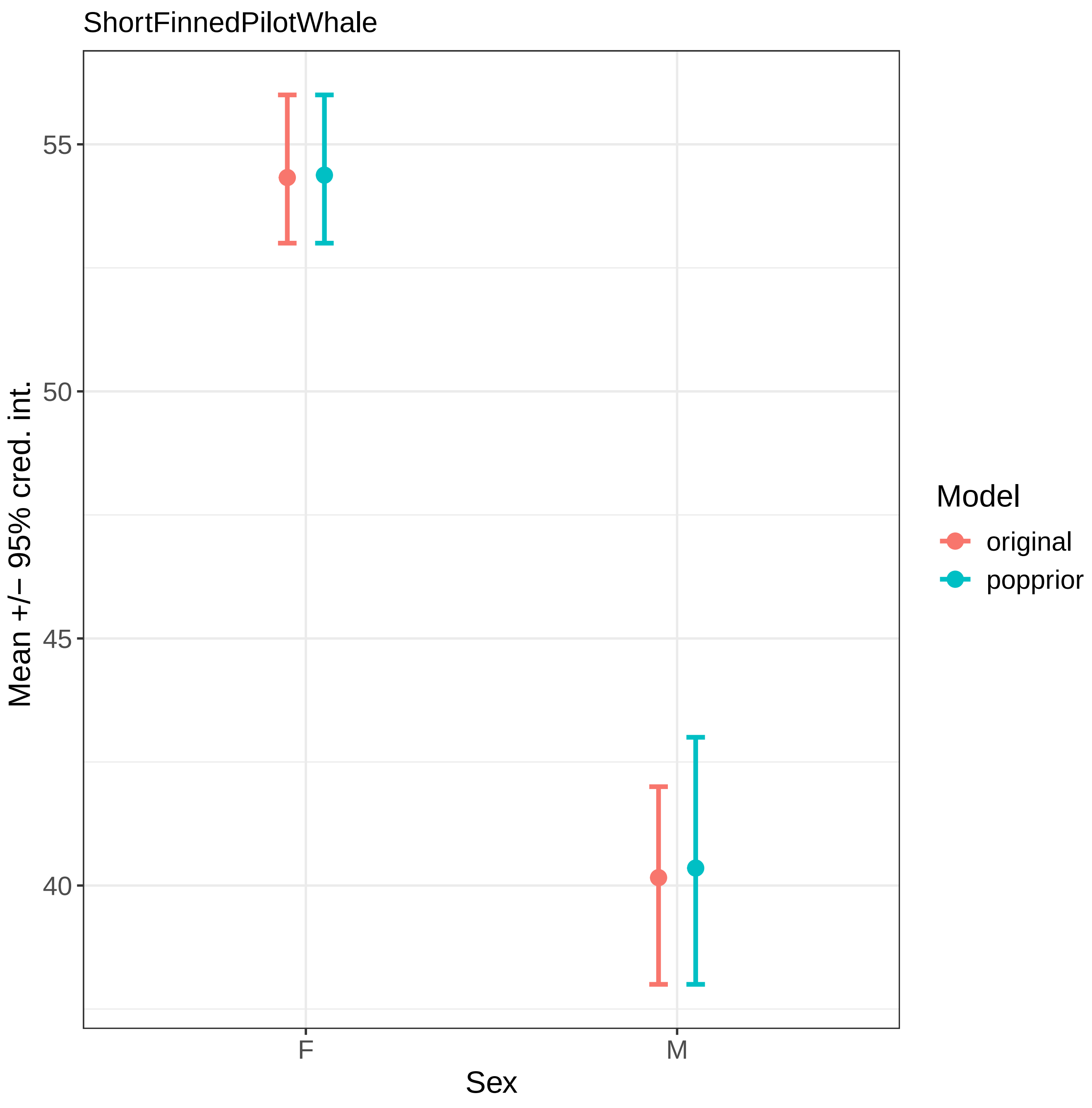
ShortFinnedPilotWhale
Mean +/− 95% cred. int.
55
50
45
40
F
M
Sex
Model
original
popprior

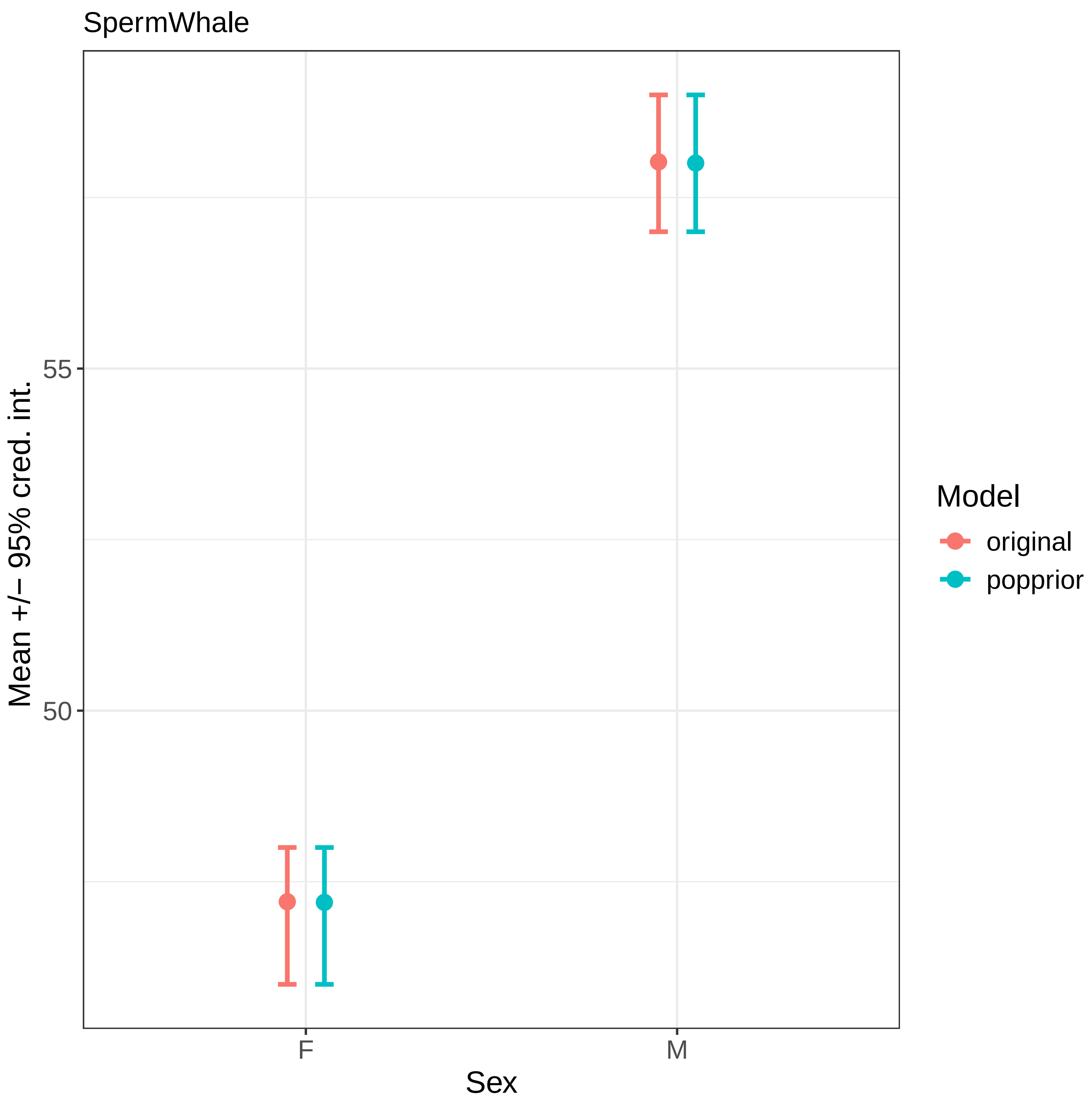
SpermWhale
Mean +/− 95% cred. int.
55
50
F
M
Sex
Model
original
popprior

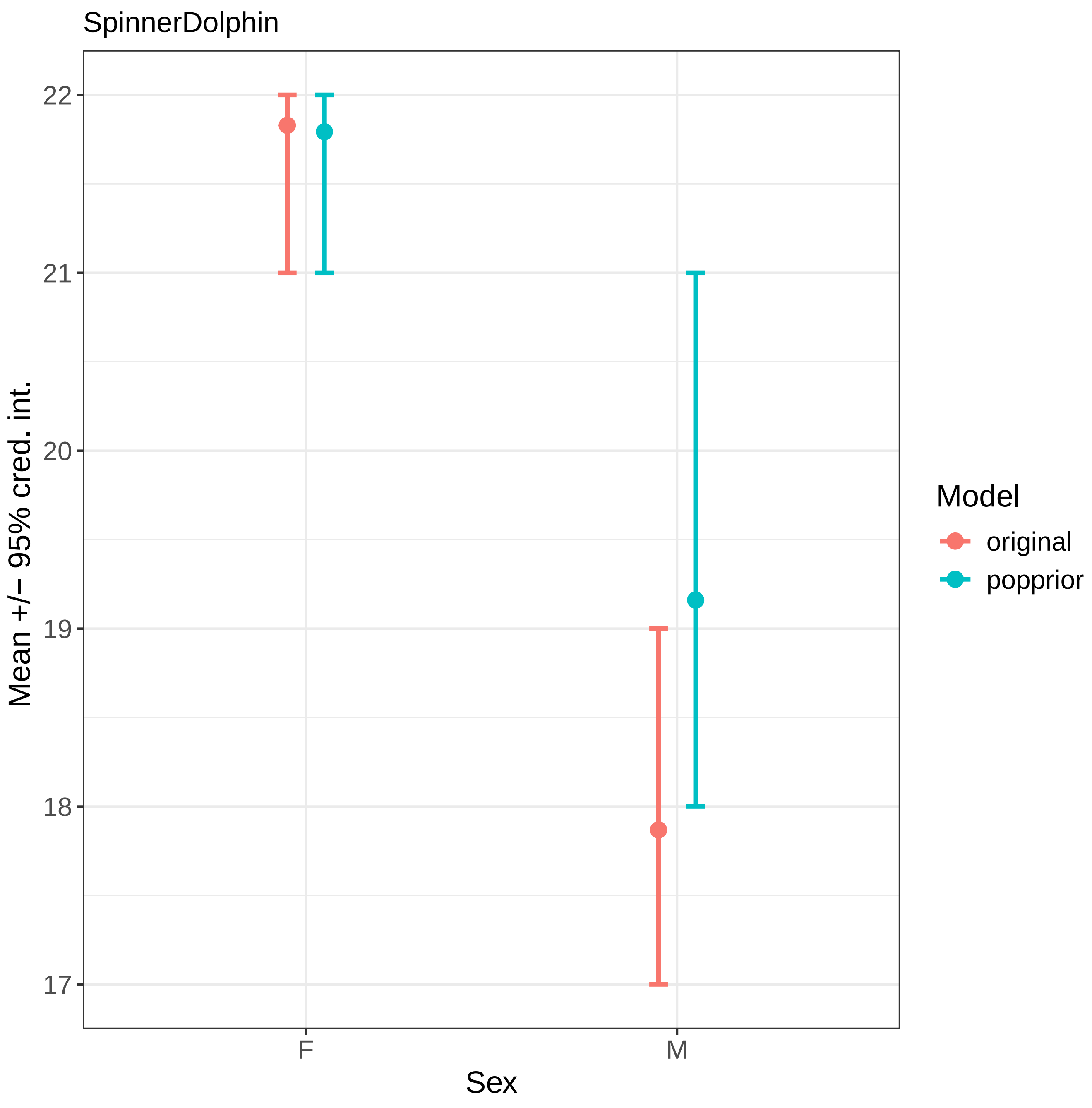

SpinnerDolphin
22
21
20
19
18
17
Mean +/− 95% cred. int.
F
M
Sex
Model
original
popprior

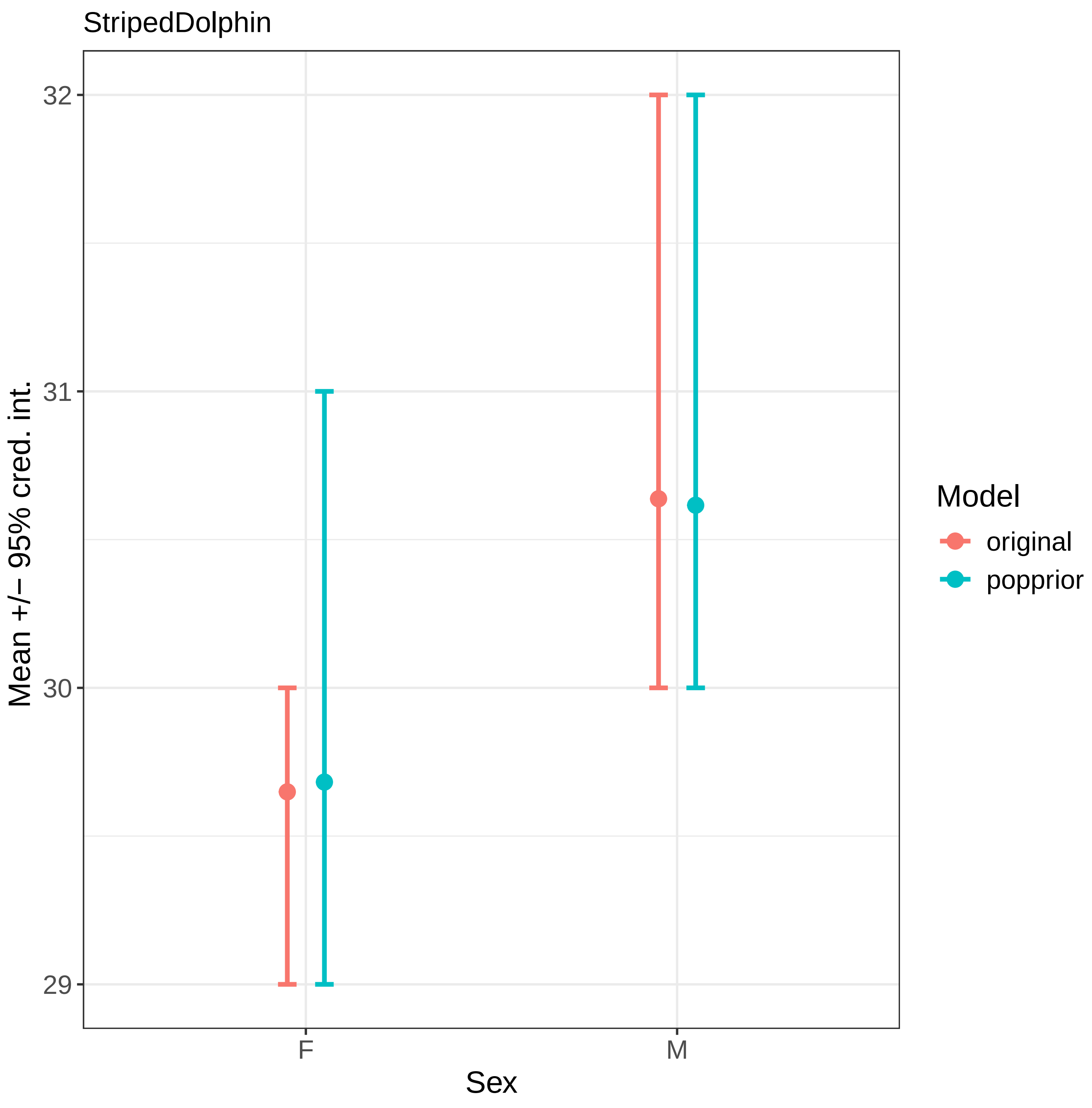
StripedDolphin
Mean +/− 95% cred. int.
32
31
30
29
F
M
Sex
Model
original
popprior

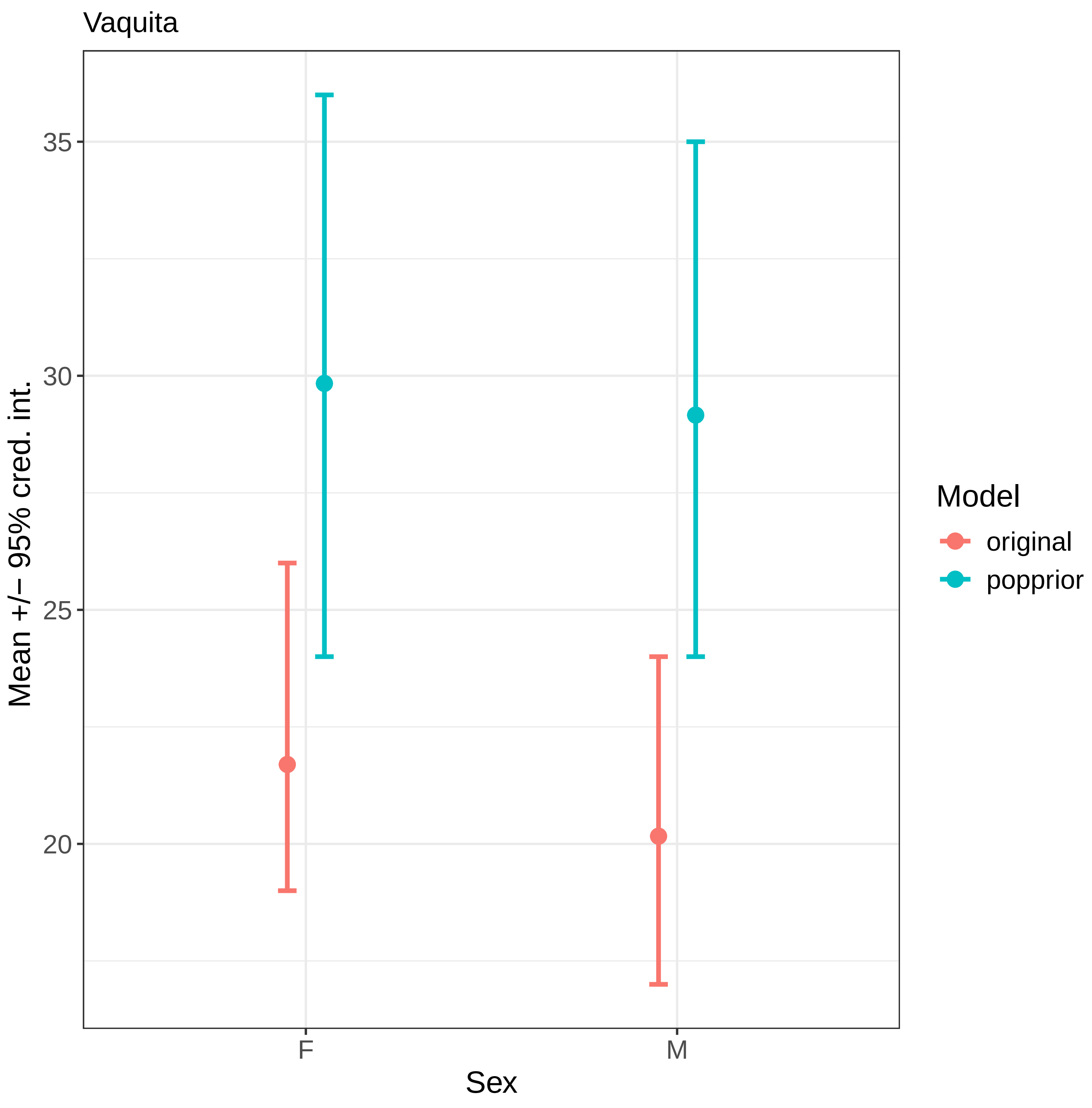
Vaquita
Mean +/− 95% cred. int.
35
30
25
20
F
M
Sex
Model
original
popprior

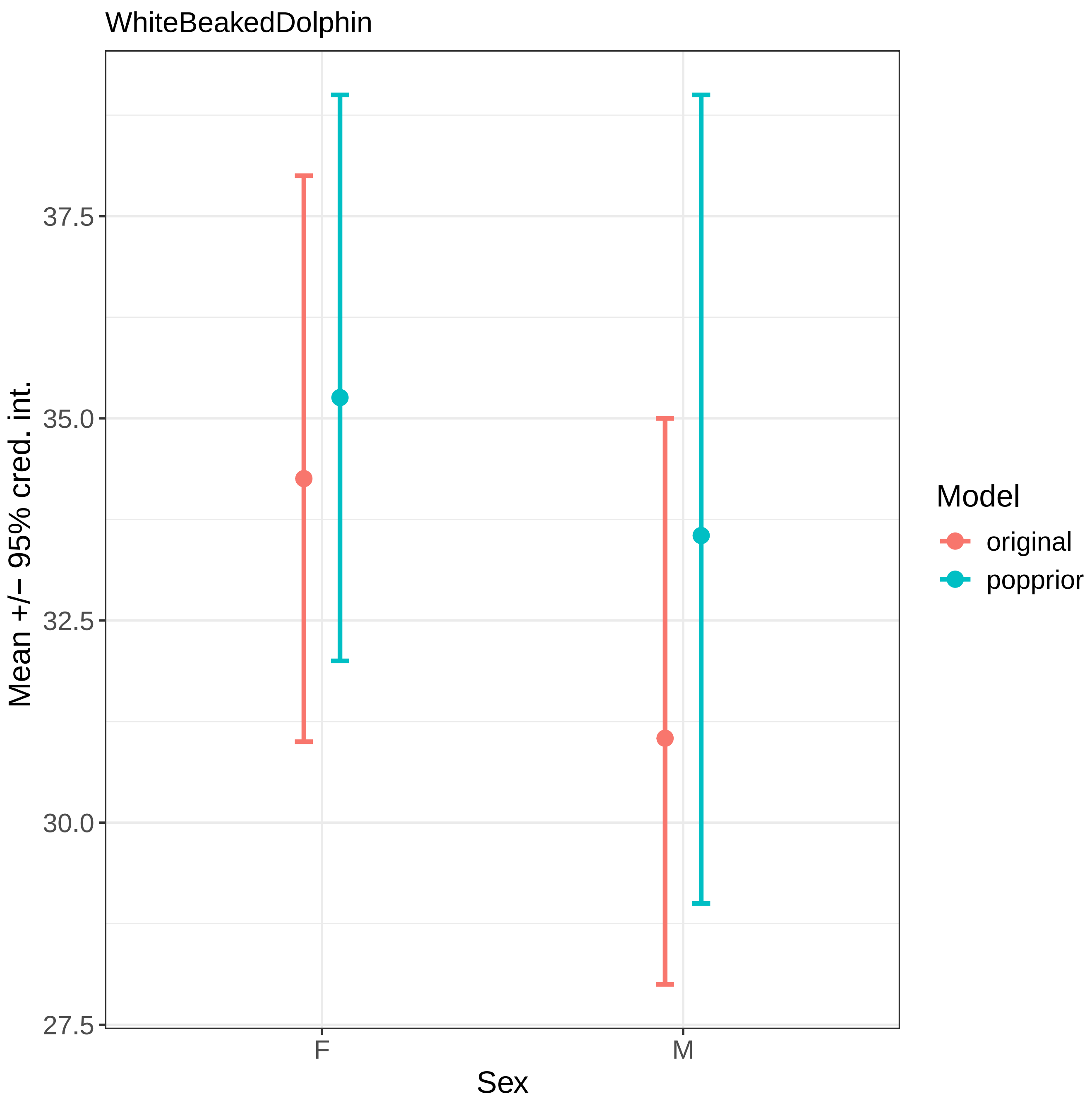
WhiteBeakedDolphin
Mean +/− 95% cred. int.
37.5
35.0
32.5
30.0
27.5
F
M
Sex
Model
original
popprior
