## Supplementary 5 for "Bayesian inference of toothed whale lifespans"

Age-at-death data is data where rather than being a cross-section of the age-cohort at a given time is instead samples of individuals who have died from "natural" causes. Age-at-death data are not common in toothed whale data because most data come from mass-strandings and by-catch- and even non-mass strandings data often consist at least in part of individuals killed and discarded as by-catch or vessel collision.

Our model can be reformulated to analyse age-at-death data. Our model is fitted to a survival function which represents the probability of surviving to or beyond a given age (equation 2). Survival functions are derived from the mortality functions which describe the probability of dying at a given age. Replacing the Gompertz survival function in our model (equation 2) with the Gompertz mortality function (equation S5.1; (Gompertz 1825); and the probability density function derived from this equation S5.2), converts our model to fit to age-at-death data. All other parameters and variables in the model remain the same (equation 11).

Equation S5.1

$$u(i) = \alpha e^{\beta \times AGE_i}$$

Equation S5.2

$$U_i = \frac{\alpha e^{\beta \times AGE_i}}{\sum_{j=0}^{n} \alpha e^{\beta \times AGE_{j-1}}}$$

We demonstrate this model using data from female Southern Resident Killer Whales. In this well studied population the ages-at-death for all whales who have died since 1976 are known or estimated (see main text). Applying the reformulated age-at-death model to these data gets a predicted ordinary maximum lifespan for female resident killer whales of 64 (43-97) . This is comparable to the ordinary maximum lifespan for female southern resident killer whales derived from the longitudinally-derived estimate 69 (62-79; main text: Results/Testing the model) and the estimate derived by applying the "survival" model formulation to the population 76 (65-88; main text: Results/Testing the model). Although the age-at-death model does have a wider credible interval.

**S5 References**

Gompertz B (1825) On the nature of the function expressive of the law of human mortality, and on a new mode of determinng the value of life contingencies. Philosophical Transactions of the Royal Society 115:513–583
