## Supplementary 6 for "Bayesian inference of toothed whale lifespans"

Our estimates of ordinary maximum lifespan allow us to, for the first time, investigate the phylogenetic signal in toothed whale life history in a broad scale analysis. Phylogenetic signal is - broadly- the amount of variation in a trait, in this case ordinary maximum lifespan, explained by the phylogenetic relationships between species (Pagel 1999). It is commonly calculated in life history studies (Garamszegi 2014).

There is a strong positive relationship between body mass and lifespan across animals, including within female toothed whales (Ellis et al. 2024). To understand the phylogenetic signal within toothed whale ordinary maximum lifespans we therefore first regress ordinary maximum lifespan with body size for both males and females while controlling for phylogeny. For each sex we fit a model of the form:

Equation S6.1

$$\begin{aligned}
&\tau Z \sim MultiNormal(\mu, K)\\
&\tau Z \sim Normal(\mu Z, \sigma Z)\\
&\tau S \sim Normal(\mu S, \sigma S)\\
&\mu_i = \alpha + \beta_{SIZE} \tau S_i\\
&K_{i,j} = R_{i.j}\sigma^2
\end{aligned}$$

Where true ordinary maximum lifespan $\tau Z$ is drawn from a multi-normal distribution the means of which are described by a linear model regressed against log true size ($\tau S$). Both true size and lifespan are considered to be drawn from a normal distribution parametrised by the means ($\mu X$) and standard deviation ($\sigma X$) derived from the data and models. The dispersion of the multi-normal distribution, $K$, is a covariance matrix derived from the phylogenetic closeness (1/distance) between species ($R$)– which effectively represents the covariance driven by phylogeny as a Brownian process (Mahler and Ingram 2014). From this model we also compute the phylogenetic signal in the relationship between ordinary maximum lifespan and body size. We use body length as our measure of size due to the difficulty in robustly measuring the mass of cetaceans (Whitehead and Mann 2000; Ellis et al. 2024). We derive our measures of body length from published expert-consensuses as a mean and standard deviation for each species-sex (described in detail Ellis et al 2024). We use a recent consensus phylogeny (McGowen et al. 2020; adapted as *per* Ellis et al 2024) as a measure of the phylogenetic distance between two species. We quantify phylogenetic signal as Pagel's λ calculated as the proportion of variance in the model (equation 14) explained by phylogenetic structure (here the phylogenetic covariance) divided by the total model variance (Bürkner 2017; but see Pearse et al. 2023)).

As expected there is a strong positive relationship between lifespan and size in both sexes (females: $\beta_{SIZE}$, post. mean = 0.75, 95% cred. int. = 0.35-1.15, males: $\beta_{SIZE}$, post. mean = 0.56, 95% cred. int. = 0.24-0.83). In neither sex is weak phylogenetic signal in the relationship between size and lifespan (females: Pagel's λ, post. mean = 0.10, 95% cred. int. = 0.01-0.32; males: Pagel's λ, post. mean = 0.13, 95% cred. int. = 0.002-0.59; figure S6.1), although for both sexes the uncertainty around the estimate is high.

Although we find some evidence for a weak phylogenetic signal in the relationship between ordinary maximum lifespan and body size in toothed whales this result should be interpreted with caution. Pagel's λ has been criticised for being hard to define (Pearse et al. 2023). This is especially in the context of hierarchical Bayesian models, and the approach we use here - although widely applied - may actually be better interpreted as heritability than phylogenetic signal itself (Pearse et al. 2023). Further, the value itself largely depends on what other factors are included in the model, and the estimate is perhaps best interpreted as "unexplained variation that correlates with phylogeny"  rather than phylogenetic signal *senus stricto* (Pearse et al. 2023). For example, various aspects of diet, social behaviour and distribution will also correlate with phylogeny and unpicking the contribution of these various factors is an important avenue for future research.

**S6 References**

Bürkner PC (2017) brms: An R package for Bayesian multilevel models using Stan. Journal of Statistical Software 80:. https://doi.org/10.18637/jss.v080.i01

Ellis S, Franks DW, Nielsen MLK, et al (2024) The evolution of menopause in toothed whales. Nature 627:579–585. https://doi.org/10.1038/s41586-024-07159-9

Garamszegi LZ (ed) (2014) Modern Phylogenetic Comparative Methods and Their Application: Concepts and Practice. Springer, London

Mahler DL, Ingram T (2014) Phylogenetic comparative methods for studying clade-wide convergnence. In: Garamszegi LZ (ed) Modern Phylogenetic Comparatve Methods and Their Application in Evolutionary Biology: Concepts and Practice. Springer, London, pp 425–450

McGowen MR, Tsagkogeorga G, Álvarez-Carretero S, et al (2020) Phylogenomic Resolution of the Cetacean Tree of Life Using Target Sequence Capture. Systematic Biology 69:479–501. https://doi.org/10.1093/sysbio/syz068

Pagel M (1999) Inferring the historical patterns of biological evolution. Nature 401:877–884. https://doi.org/10.1038/44766

Pearse WD, Davies TJ, Wolkovich EM (2023) How to define, use, and interpret Pagel's λ (lambda) in ecology and evolution. 2023.10.10.561651

Whitehead H, Mann J (2000) Female reproductive strategies of ceteceans: life histories and calf care. In: Mann J, Connor RC, Tyack PL, Whitehead H (eds) Cetacean Societies: Field Studies of Whales and Dolphins. The University of Chicago Press, Chicago and London, pp 219–246

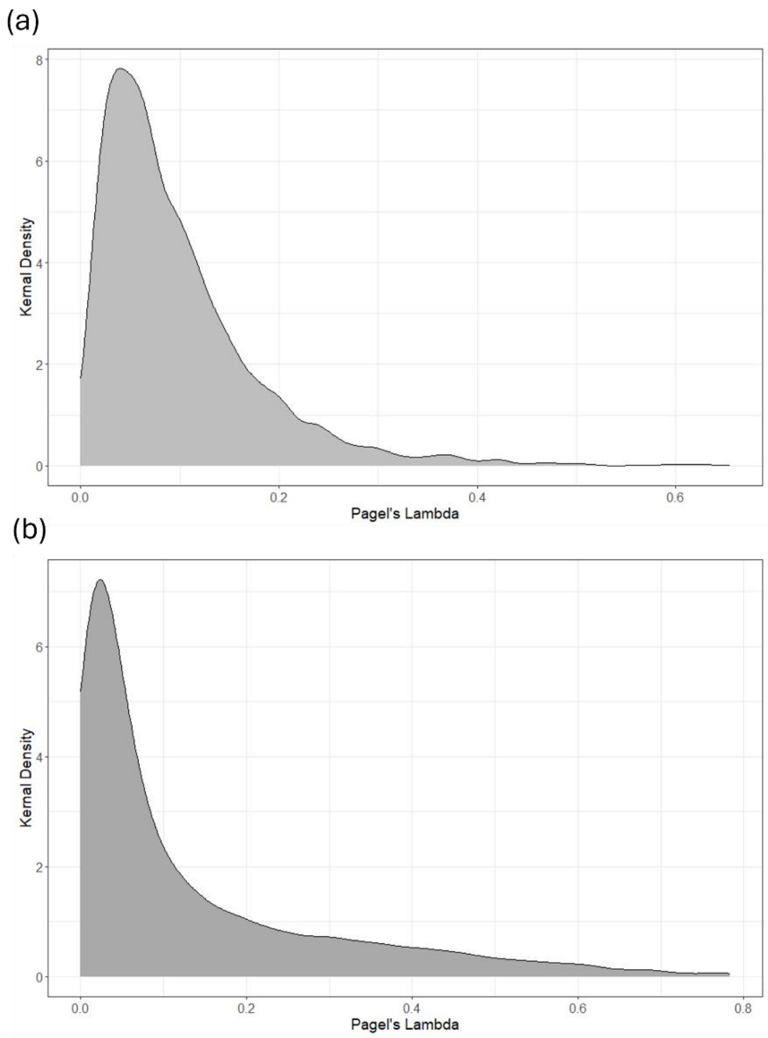


Figure S6.1. Posterior Distribution of Pagel's λ for female (a) and male(b) toothed whales.
