## Supplementary 7 for "Bayesian inference of toothed whale lifespans"

Comparison of models with age-estimation error model structure presented in the text (Model A) and an alternative structure (Model B) for five species of toothed whale.

Model A has the age-estimation error structure

$$\tau_j \sim Normal(o_j, \epsilon_j)$$

And Model B has the alternative structure

$$o_j \sim Normal(\tau_j, \epsilon_j)$$

The rest of the model structure remains the same as in Equation 11.

Both structures give very similar results (five example species presented below). But models with structure A are healthier models that fit more easily and robustly, and hence are retained in the text.

We chose the five species to give a spread of species with and without menopause, and where the sexes have both similar and different lifespans.

| SPECIES | SEX | MODEL | ORDINARY MAX. LIFESPAN POST. MEAN | ORDINARY MAX. LIFESPAN POST. 95% LCI | ORDINARY MAX. LIFESPAN POST. 95% LCI |
|---|---|---|---|---|---|
| **BAIRDS BEAKED WHALE** | F | A | 52.76 | 46 | 62 |
| | F | B | 51.8 | 46 | 59 |
| | M | A | 87.31 | 78 | 97 |
| | M | B | 82.7 | 75 | 91 |
| **BELUGA WHALE** | F | A | 53.85 | 52 | 56 |
| | F | B | 53.55 | 52 | 55 |
| | M | A | 48.72 | 46 | 52 |
| | M | B | 50.41 | 48 | 53 |
| **FALSE KILLER WHALE** | F | A | 67.51 | 62 | 74 |
| | F | B | 66.32 | 62 | 71 |
| | M | A | 55.54 | 49 | 63 |
| | M | B | 58.11 | 53 | 64 |
| **NARWHAL** | F | A | 76.81 | 69 | 86 |
| | F | B | 76.43 | 69 | 84 |
| | M | A | 66.27 | 58 | 75 |
| | M | B | 65.16 | 58 | 72 |
| **SPERM WHALE** | F | A | 48.7 | 48 | 50 |
| | F | B | 47.99 | 47 | 49 |
| | M | A | 52.5 | 51 | 54 |
| | M | B | 53.8 | 53 | 55 |
